# Revealing enzyme functional architecture via high-throughput microfluidic enzyme kinetics

**DOI:** 10.1101/2020.11.24.383182

**Authors:** C.J. Markin, D.A. Mokhtari, F. Sunden, M.J. Appel, E. Akiva, S.A. Longwell, C. Sabatti, D. Herschlag, P.M. Fordyce

**Author notes:** These authors contributed equally to this work (C.J. Markin, D.A. Mokhtari).

## Abstract

Systematic and extensive investigation of enzymes is needed to understand their extraordinary efficiency and meet current challenges in medicine and engineering. We present HT-MEK, a microfluidic platform for high-throughput expression, purification, and characterization of >1500 enzyme variants per experiment. For 1036 mutants of the alkaline phosphatase PafA, we performed >670,000 reactions to determine >5000 kinetic and physical constants for multiple substrates and inhibitors. These constants allowed us to uncover extensive kinetic partitioning to a misfolded state and isolate catalytic effects, revealing spatially contiguous “regions” of residues linked to particular aspects of function. These regions included active-site proximal residues but also extended to the enzyme surface, providing a map of underlying architecture that could not be derived from existing approaches. HT-MEK, using direct and coupled fluorescent assays, has future applications to a wide variety of problems ranging from understanding molecular mechanisms to medicine to engineering and design.

**One Sentence Summary:** HT-MEK, a microfluidic platform for high-throughput, quantitative biochemistry, reveals enzyme architectures shaping function.

## Introduction

Understanding how sequence encodes function remains a fundamental challenge in biology. Linear chains of amino acids fold into three-dimensional protein structures that carry out the physical and chemical tasks needed for life, such as highly efficient and specific catalysis. Variations in these amino acid sequences across organisms and individuals confer beneficial and deleterious effects: sequence variation throughout evolution creates proteins with improved or novel functions, but variation amongst individuals can also compromise function and cause disease (*1*–*3*). An enhanced predictive understanding of the sequence-function landscape could have profound impacts across biology, from enabling efficient protein design to improving accurate detection of rare allelic variants that drive disease (*4*–*7*).

Understanding sequence-function relationships within enzymes poses a particular challenge. Structural and biochemical studies of enzymes have revealed the sites of substrate binding and catalytic transformation, the residues directly involved in catalysis, and roles for these residues. However, the remainder of the enzyme is also essential: residues outside the binding and active sites are needed for the active site to assemble and function, and allosteric ligands and covalent modifications modulate activity through interactions with distant surface residues (*8*–*10*). Despite their importance, the roles played by residues outside the active site, which comprise the majority of amino acids in an enzyme, remain largely unexplored.

This dearth of knowledge stems from the nature of experimental approaches currently available. Site-directed mutagenesis (SDM), which entails systematically mutating specific residues, has traditionally been used to assess function via in-depth biochemical assays that yield kinetic and thermodynamic constants. However, SDM is time, resource, and labor-intensive, limiting investigation to a small number of residues. By contrast, deep mutational scanning (DMS) provides the ability to assay the effects of all 20 amino acids at every position within an enzyme (*5, 11, 12*). Nevertheless, DMS lacks the depth and dimensionality of traditional SDM studies, typically providing a scalar readout with an uncertain relationship to the multiple fundamental physical constants that are needed to describe an enzyme’s function.

Marrying the strengths of traditional SDM and emerging DMS would usher in a new era of mechanistic enzymology. Here, we present HT-MEK (High-Throughput Microfluidic Enzyme Kinetics), a platform capable of simultaneously expressing, purifying, and characterizing >1000 rationally chosen enzyme mutants in parallel with the depth and precision of traditional SDM. Each HT-MEK experiment provides 1000s of measurements and multiple kinetic and thermodynamic constants (*e*.*g*., *k*_cat_, *K*_M_, *k*_cat_/*K*_M_, *K*_i_) in days and at low cost.

To guide HT-MEK development and demonstrate its capabilities, we carried out a comprehensive mechanistic investigation of the effects of mutations to every residue within the alkaline phosphatase superfamily member PafA (Fig. 1A and fig. S1). PafA and related phosphomonoesterases are among the most prodigious catalysts known, with rate enhancements of up to ∼10^27^-fold, providing a large dynamic range to explore (*13*). We also anticipated that PafA, a secreted enzyme, would be highly stable, potentially allowing us to more deeply probe catalysis without obfuscation from global unfolding. Strikingly, we found that 702 of the 1036 mutants investigated have significant functional consequences, with none arising from equilibrium unfolding. Additional experiments revealed that many mutations promote the formation of a long-lived, catalytically-incompetent misfolded state both *in vitro* and in cells. The multidimensional measurements provided by HT-MEK allowed us to decouple misfolding from catalytic effects and to relate these catalytic effects to particular aspects of catalysis and mechanism using an approach we term Functional Component Analysis. Each functional parameter was affected by a large number of mutations, with spatially contiguous regions of effects extending from the active site to the enzyme’s surface. These regions of residues with shared functional signatures together define the enzyme’s functional architecture, and reveal different regions of the enzyme responsible for optimizing particular catalytic strategies and that can potentially be used to tune particular reaction parameters. Surface residues associated with these functional effects provide candidate allosteric handles for rational control of catalytic activity. The HT-MEK platform and the quantitative multidimensional datasets that can be obtained will have broad utility for future efforts to understand catalytic mechanisms, natural variation, and evolutionary trajectories, and to design enzymes with novel functions.

**Fig. 1.**
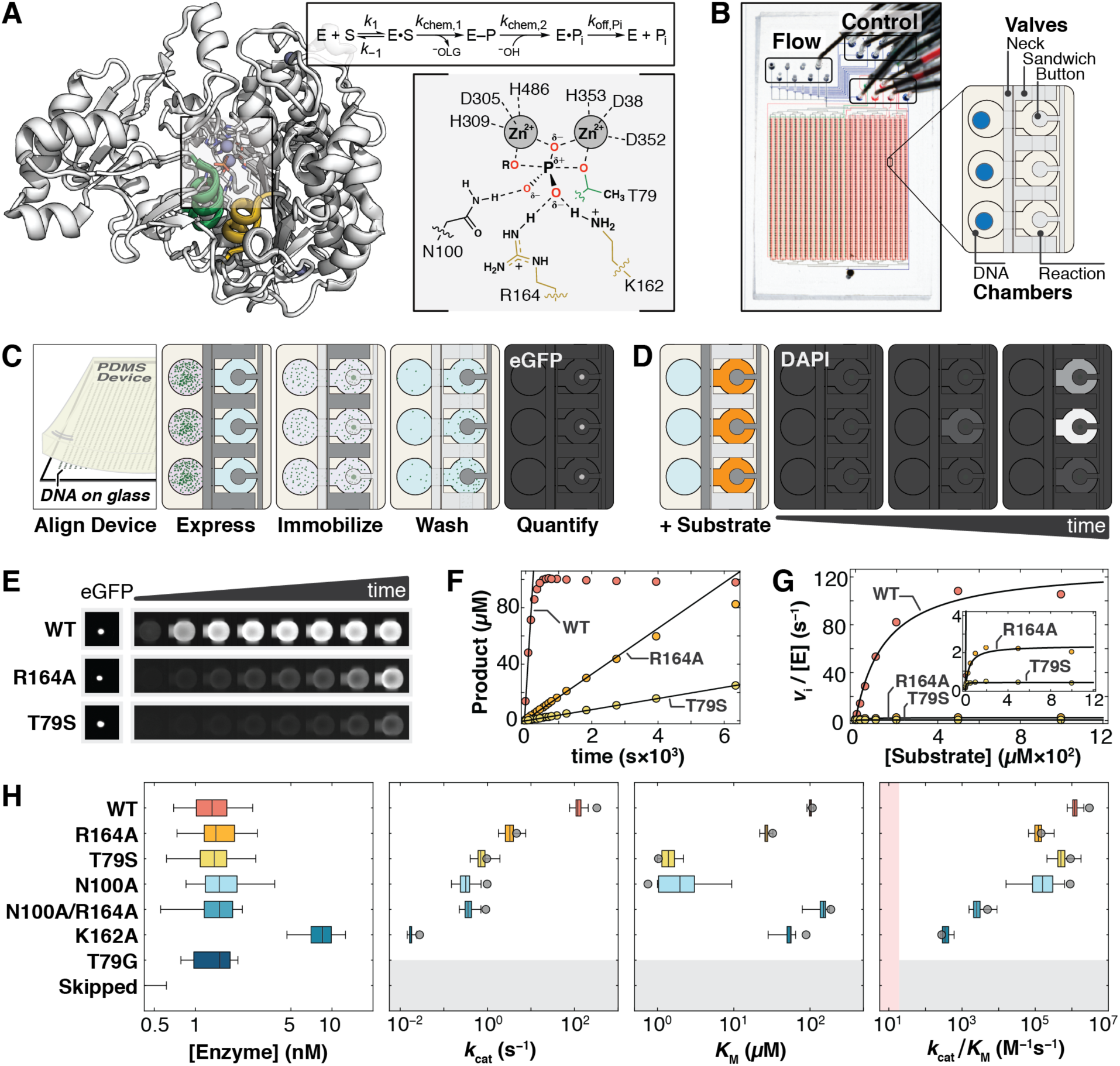
PafA and HT-MEK overview. (**A**) Crystal structure of WT PafA (PDB ID: 5TJ3, left), and representation of its active site in the transition state of phosphate monoester hydrolysis (right), showing active site residues and relevant secondary structures (“nucleophile helix,” positions 77–89, green; “monoesterase helix,” positions 161–171, yellow; this color coding is used throughout). The PafA catalytic cycle is schematized above where E–P represents the covalent phospho-threonine intermediate. HT-MEK microfluidic device image and schematic showing solution (Flow) and pneumatic manifold (Control) input ports, as well as device valves and chambers. Schematic of on-chip enzyme expression pipeline. Dark and light gray valves are pressurized (closed) and depressurized (open), respectively. (**D**) Schematic of on-chip activity assays using fluorogenic substrate in reaction chambers (orange). (**E**) Sample data images of immobilized enzyme (left) and accumulation of fluorogenic product over time for WT PafA and two active site mutants (R164A and T79S) with the substrate cMUP. (**F**) Example cMUP progress curves for chambers containing WT and two active site mutants, and initial rate fits to these data. (**G**) Michaelis-Menten fits to initial rates yield *k*_cat_, *K*_M_, and *k*_cat_/*K*_M_ for cMUP. (**H**) On-chip expressed concentrations for WT PafA and six active site mutants calculated using eGFP calibration curves (left); comparisons of on-chip (box plots) and off-chip (grey circle) values of *k*_cat_, *K*_M_, and *k*_cat_/*K*_M_ values for cMUP for seven PafA variants (WT, R164A, T79S, N100A, N100A/R164A, K162A, and T79G; table S1). K162A was expressed at higher concentration in a later experimental tier and data were merged with other active site variants from the Active Site print (table S2). “Skipped” refers to chambers without printed plasmid DNA. The boundary of the pink shaded region is 10-fold above the median apparent second order rate constant determined from controls measuring hydrolysis in T79G chambers (see Materials and Methods).

### HT-MEK device and experimental pipeline

HT-MEK is built around a two-layer polydimethylsiloxane (PDMS) microfluidic device with 1568 chambers and integrated pneumatic valves (Fig. 1B and fig. S2) (*14, 15*). Each chamber is composed of two compartments (DNA and Reaction) separated by a valve (Neck), with adjacent chambers isolated from one another by a second valve (Sandwich). A third valve (Button) reversibly excludes or exposes a circular patch of the reaction compartment surface, enabling surface patterning for on-chip protein immobilization and purification (Fig. 1C) as well as simultaneous initiation of successive on-chip reactions across the device (Fig. 1D). Each DNA compartment of each chamber is programmed with a specified enzyme variant by aligning the device to a spotted array of DNA plasmids that encode for the expression of C-terminally eGFP-tagged variants (fig. S3). After alignment, device surfaces are patterned with anti-eGFP antibodies beneath the Button valve and passivated with BSA elsewhere. All enzymes are then expressed in parallel via the introduction of an *E. coli in vitro* transcription/translation system and purified via capture by surface-patterned immobilized antibody and washing (figs. S3 and S4). Production of up to 1568 different purified enzymes takes ∼10 hours, with most steps automated. Enzymes are immobilized under Button valves that protect against flow-induced loss of enzyme during solution exchange and allow repeated synchronous initiation of reactions.

To obtain catalytic rate parameters, we quantify: (1) the concentration of immobilized enzyme in each chamber using an eGFP calibration curve (fig. S5) and (2) the amount of product formed as a function of reaction time using a chamber-specific product calibration curve (fig. S6). We then fit reaction progress curves in each chamber to obtain initial rates (*v*_i_) for each substrate concentration using a custom image processing pipeline (Fig. 1, E and F, and fig. S7). This process, repeated on a single device for multiple substrate concentrations, multiple substrates, and multiple inhibitors, provides the data necessary to obtain Michaelis-Menten parameters and other kinetic and thermodynamic constants (Fig. 1G and fig. S7).

### HT-MEK reproduces kinetic constants previously measured via traditional assays

To demonstrate the technical capabilities of HT-MEK, we applied it to study seven previously-characterized PafA variants: wild-type (WT), five active site mutants (T79S, N100A, R164A, K162A, and N100A/R164A), and one mutant lacking detectable activity (T79G, negative control) (*16*). Activities of WT PafA and the six mutants span a broad range in *k*_cat_ (>10^4^-fold), *k*_cat_/*K*_M_ (>10^4^-fold), and *K*_M_ (>10^2^-fold) for aryl phosphate monoester hydrolysis, providing a stringent initial test of HT-MEK dynamic range (table S1). Nearly all DNA-containing chambers expressed enzyme (>90%), and all mutants expressed at similar levels as determined by eGFP fluorescence (Fig. 1H and fig. S8; K162 was deliberately expressed at higher concentrations in a later experimental tier, described below).

While fluorogenic phosphate ester substrates permit kinetic assays of phosphatase activity with a high dynamic range, microfluidic assays using the commercial methylumbelliferone phosphate ester (MUP) were complicated by partitioning of the hydrophobic fluorescent product into the hydrophobic PDMS, which increases background and can distort kinetic measurements. To address this limitation, we synthesized MUP derivatives of similar reactivity (cMUP and a corresponding methyl phosphodiester, MecMUP) that bear a charged moiety on the leaving group to eliminate PDMS absorption (figs. S9 to S11 and table S1).

Accurately resolving enzymatic rates spanning four or more orders of magnitude poses technical challenges, as different acquisition times are needed at catalytic extremes and even a small concentration of contaminating fast enzyme introduced into nearby chambers during fluid exchanges can obscure the true rates for the most catalytically-compromised mutants. To address the first challenge, we expressed enzymes at two concentrations: ∼1.5 nM (for accurate measurement of fast enzymes) and ∼15 nM (to speed reactions for efficient detection of slow enzymes) (Fig. 1H and fig. S8). To identify any regions of the device with contaminating enzyme from other chambers, we interspersed chambers that were empty or contained the inactive T79G mutant and also measured their apparent activity (fig. S15 and table S2; see Materials and Methods). Per-device normalizations (0.4 and 1.4-fold) were used to account for small variations in apparent activity due to non-specific adsorption of mutant enzymes to chamber walls (see Materials and Methods). This normalization increased precision across replicates but did not affect conclusions (fig. S17). HT-MEK assays recapitulated cMUP kinetic parameters accurately and over a wide dynamic range (>10^4^-fold in *k*_cat_/*K*_M_; Fig. 1H).

### Many mutations throughout PafA affect phosphate monoester hydrolysis

To explore functional effects of mutations throughout PafA, we created mutant libraries in which we introduced two residues with widely differing side-chain properties at each position: i) glycine, to ablate site chain interactions and increase backbone flexibility, and ii) valine, to introduce a branched chain hydrophobe of average volume. Native valine and glycine residues were mutated to alanine. Nearly all of these 1052 possible mutants (1036, 98%) were successfully cloned, sequenced, expressed, and assayed via HT-MEK (fig. S18).

We first measured the catalytic effect of each substitution on the steady-state kinetic parameters for cMUP hydrolysis (*k*_cat_, *K*_M_, and *k*_cat_/*K*_M_) (Fig. 2A). To facilitate measurement of mutants at catalytic extremes, we performed experiments in three tiers based on reaction rates. In tier 1, we assayed devices containing all valine or all glycine variants, printed in duplicate and with active site mutants distributed throughout as fiducial controls. Tier 2 and 3 measurements focused successively on the slowest variants with increasing numbers of replicates to provide high-precision measurements of more deleterious mutants. Each device was used to measure 10s of cMUP progress curves and all expressed variants were stable over days (as assessed by periodic re-measurement of rate constants at a given substrate concentration), facilitating high-throughput data collection. Per-experiment data reports contain all data collected for each chamber including initial rate plots and fit Michaelis-Menten curves (example in fig. S19, full data sets in Supporting Auxiliary Files); per-mutant summaries combine data from all experiments and include estimates of statistical significance (Fig. 2, B and C, and fig. S20). In total, we acquired a median of 9 and 7 replicates for valine and glycine mutants, respectively, over 16 experiments (figs. S22–24 and table S2). The wealth and precision of these data allowed us to resolve differences across ranges of 10^4^, 10^2^, and 10^5^-fold for *k*_cat_, *K*_M_, and *k*_cat_/*K*_M_, respectively.

**Fig. 2.**
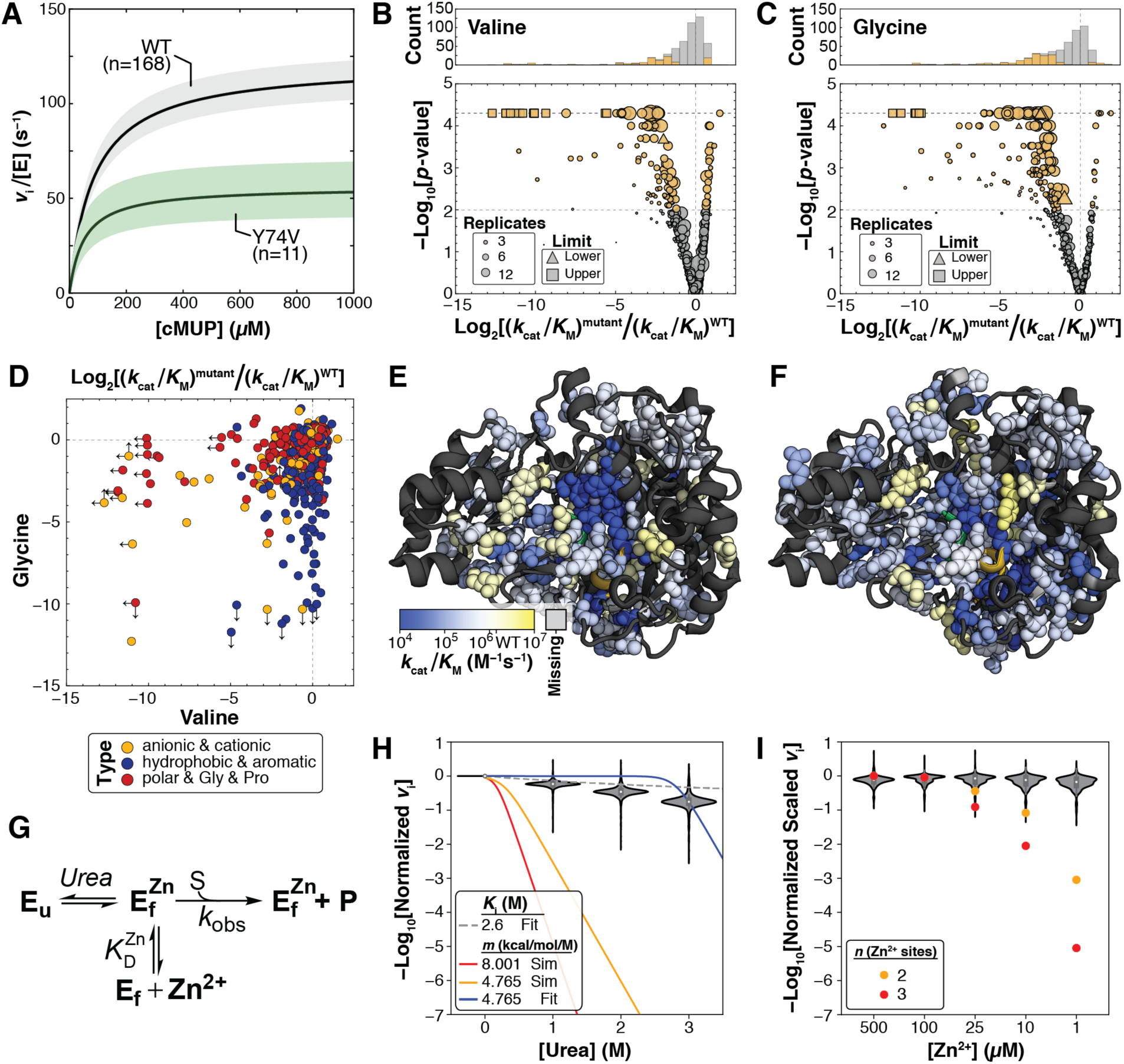
HT-MEK measurements of aryl phosphate monoester (cMUP) hydrolysis for valine and glycine scans of PafA. (**A**) Median Michaelis-Menten curves for wild-type and an example mutant with *k*_cat_ and *K*_M_ effects. The colored regions denote 99% confidence intervals (CIs) on the medians of parameters from replicate measurements. (**B**) Valine substitutions at 126 positions alter cMUP *k*_cat_/*K*_M_ at *p* < 0.01 (105 slower, 21 faster, gold markers; grey, *p*-value ≥ 0.01). (**C**) Glycine substitutions at 176 positions alter cMUP *k*_cat_/*K*_M_ at *p* < 0.01 (162 slower, 14 faster, gold; grey, *p* ≥ 0.01). (**D**) Magnitude of effects for glycine vs. valine substitutions on *k*_cat_/*K*_M_ of cMUP hydrolysis at each position, colored by identity of the native residue. Arrows denote the presence and direction of measurement limits. (**E**) Magnitude of valine substitution effects on cMUP hydrolysis shown on the PafA structure (*p* < 0.01 sites shown as spheres, *p* ≥ 0.01 and missing sites represented as ribbons). (**F**) Magnitude of glycine substitution effects on cMUP hydrolysis shown on the PafA structure (coloration as in **E**). (**G**) Model of equilibrium unfolding of PafA in the presence of varying urea and Zn^2+^. (**H**) Activity of mutants relative to WT in the presence of increasing concentrations of urea at 50 μM cMUP and compared to the urea dependencies predicted if mutants were 10% unfolded (red and yellow curves). (**I**) Activity of mutants relative to WT (normalized across experiments as described in the Materials and Methods) in the presence of decreasing concentrations of zinc. Orange and red points are expected activities given Zn^2+^-concentration dependent unfolding assuming 2 and 3 Zn^2+^ binding events, respectively, if mutants were 10% unfolded at 100 µM Zn^2+^.

As expected, mutations of active site residues and catalytic Zn^2+^ ligands were highly deleterious, and positional effects varied for valine and glycine (Fig. 2, D to F). Nevertheless, a surprisingly large number of mutations throughout the enzyme were deleterious, with decreases in *k*_cat_/*K*_M_ observed for 267 of the 1036 mutants (*p* < 0.01). We also observed several with increased activity (35; fig. S22). These measurements provide a comprehensive quantitative survey of Michaelis-Menten kinetic constants for mutations throughout a large enzyme, but do not tell us why so many mutations alter activity.

The most obvious explanation for these widespread effects would be destabilization leading to a significant fraction of unfolded enzyme. Beyond destabilization, mutations can have other repercussions, altering the catalytic effectiveness of particular active site residues, reducing Zn^2+^ affinity at the bimetallo active site, or altering enzymatic protonation states. The ability to efficiently measure catalytic activity for all mutants under different assay and expression conditions as afforded by HT-MEK allowed us to directly test each of these possibilities for PafA.

### Widespread mutational effects do not arise from equilibrium unfolding

Reflecting its role as a secreted phosphatase designed to function in harsh and variable environments, WT PafA is highly stable and remains folded, as inferred by circular dichroism (CD) spectra, even after exposure to 4 M urea for 14 days (fig. S25). This stability suggests that any individual mutation is unlikely to lead to a significant population of unfolded enzyme. To directly test this expectation, we measured cMUP activity in the presence of increasing concentrations of urea for all mutants. If a PafA variant were already partially unfolded in the absence of urea, then even low concentrations of added urea would cause substantial additional unfolding, following standard urea unfolding dependence (*17*), and proportionally lower activity (Fig. 2G and Supplementary Text S1). By contrast, we observed only minor rate effects for all mutants (6±1-fold decrease at 3 M compared to 0 M urea, ±SEM), dramatically less than the >10^9^-fold decrease predicted for an unfolding effect and consistent with inhibition by urea with *K*_i_ = 2.6 M (Fig. 2H and fig. S26).

Equilibrium unfolding would also predict a Zn^2+^ concentration dependence for the observed reaction rate (Fig. 2G). Consistent with an absence of unfolding effects, we observe no dependence on Zn^2+^ over a ∼10^3^-fold concentration range (Fig. 2I and Supplementary Text S1). This experiment also establishes that observed rate effects do not arise from loss of bound Zn^2+^ due to lowered Zn^2+^ affinity. Finally, mutants were unaltered in their pH dependencies (fig. S27 and Supplementary Text S1), ruling out altered protonation states as responsible for the observed kinetic effects.

### A general high-throughput assay for phosphate release and additional mutational effects

While fluorogenic probes provide a sensitive and convenient method for directly visualizing enzyme activity in kinetic assays, many reactions lack a direct fluorogenic readout. To allow future application of HT-MEK to a much broader range of enzymes lacking fluorogenic substrates, we developed an on-chip coupled assay in which P_i_ is detected via fluorescence emitted upon binding to a modified phosphate binding protein (PBP, Fig. 3A) (*18*). Calibration curves for P_i_ and PBP and control measurements using methyl phosphate (MeP) as a PafA substrate established that this coupled on-chip assay can detect sub-micromolar P_i_ formation and accurately reproduce off-chip kinetic constants (figs. S28–S30).

**Fig. 3.**
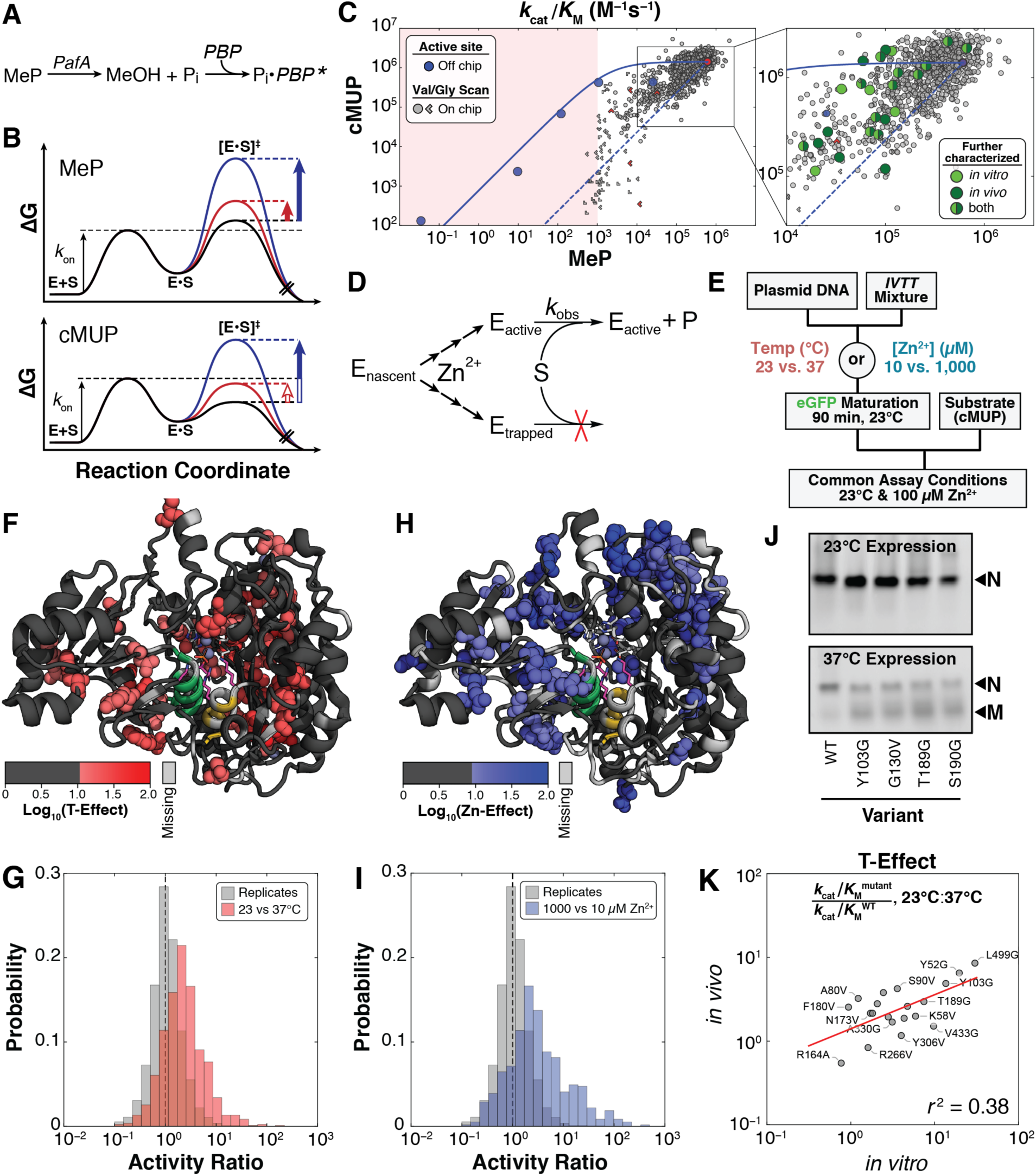
Substrates with different intrinsic reactivity reveal PafA kinetic folding traps. (**A**) Inorganic phosphate (P_i_) produced from MeP by PafA is detected in real time on chip using the Phosphate Sensor, a fluorophore-conjugated phosphate binding protein (PBP) with dramatically enhanced emission upon P_i_ binding (*18*). (**B**) Reaction coordinate diagrams for substrates with either chemistry (MeP) or binding (cMUP) as the rate-limiting step, illustrating how effects of mutations (red, blue) can be obscured for substrates with a binding-limited rate of hydrolysis (cMUP). In each case, the solid part of the arrow corresponds to the portion of the intrinsic effect on the chemical step expressed in *k*_cat_/*K*_M_. (**C**) Measured *k*_cat_/*K*_M_ for MeP vs. *k*_cat_/*K*_M_ for cMUP for active site mutants off-chip (blue points) and for valine and glycine library mutants. Limits in one or both directions are denoted by chevrons pointing in the direction or quadrant, respectively, of the limit. Green points correspond to Val and Gly library mutants subjected to further off-chip characterization *in vitro* (light green) and/or *in vivo* (dark green). The predicted functional relationship between catalytic efficiency towards cMUP and MeP is shown as a solid blue line; the relationship predicted for enzyme with WT activity but a varying fraction of inactive enzyme is shown as the dashed blue line (see Supplementary Text S2). The right boundary of the red shaded region (∼10^3^ M^−1^s^−1^) denotes an approximate on chip lower limit of detection. (**D**) Scheme showing kinetic partitioning between folded (active) and misfolded (kinetically-trapped) enzyme during or following translation. (**E**) Pulse-chase experiment to test for expression-dependent misfolding: expression under different conditions (varying temperature or [Zn^2+^]) was followed by activity measurements under a common condition. (**F**) Residues with temperature dependent misfolding shown as spheres on the PafA structure (see Materials and Methods). (**G**) Histogram of the effects of expression temperature on mutant PafA activity (red) compared to the distribution of replicate activity assays (gray; cMUP *k*_cat_/*K*_M_, measurements under standard expression conditions). (**H**) Residues with expression [Zn^2+^] dependent activity changes upon mutation shown as spheres on the PafA structure. (**I**) Histogram of the effects of expression Zn^2+^ concentration on mutant PafA activity (blue) compared to the distribution of replicate activity assays (gray; cMUP *k*_cat_/*K*_M_, measurements under standard expression conditions). (**J**) Native gels for WT PafA and a subset of mutants expressed via *in vitro* transcription/translation off-chip. After expression at 23°C, all constructs appear as a single band corresponding to the natively-folded species (“N”); after expression at 37°C, an additional band corresponding to the putatively misfolded (“M”) species appears for non-WT PafA variants. (**K**) Linear least-squares regression of expression temperature effects on-chip (*in vitro*) versus in *E. coli* (*in vivo*) (best fit obtained for log-transformed data, red line: slope = 0.41, intercept = 0.14). See table S3 for mutants and Materials and Methods for assay conditions. *In vivo* protein levels were determined from eGFP fluorescence to estimate total expressed protein concentration.

Beyond their limited availability, fluorogenic substrates are often more reactive than naturally-occurring substrates, and this can render their binding step rate-limiting and obscure mutational effects on the chemical step of catalysis; indeed, there is evidence for this behavior with PafA (*16*). Off-chip measurements of several active site mutants revealed a decrease in observed MeP activity (due to effects on the chemical step) without a concomitant change in cMUP activity, as expected for rate-limiting cMUP binding (Fig. 3B, red vertical arrows). Once transition state destabilization was sufficiently large, MeP and cMUP reactions both slowed (Fig. 3B, blue vertical arrows). The solid blue line in Fig. 3C is a fit to the rate model derived from the free energy-reaction profiles in Fig. 3B for a series of active site mutants (blue points; fig. S31 and Supplementary Text S2) (*19*), and this model and line predict the kinetic behavior expected for the PafA glycine and valine scanning library mutants.

HT-MEK kinetic measurements for Val and Gly PafA mutants revealed *k*_cat_/*K*_M_ effects for almost half the mutants (498/1035 with *p* < 0.01, fig. S32), but few exhibited the predicted behavior (Fig. 3C, solid blue line vs. grey symbols). Instead, the mutants tended to fall between the predicted line and a diagonal line representing equally deleterious effects on the reactions of both substrates (Fig. 3C, blue solid and dashed lines, respectively). Equally deleterious effects are expected for enzymes with WT activity but only a fraction of the enzyme in the active configuration, with variants having less correctly folded enzyme further down the diagonal. Thus, the observed intermediate effects could represent combinations of effects on the chemical step and on the fraction of the PafA mutant population that is active.

### Many mutations reduce catalysis by altering folding

The urea, Zn^2+^, and pH data presented above provided strong evidence that there is not a significant fraction of inactive enzyme from equilibrium unfolding for any of the PafA variants (Fig. 2H). We therefore considered and tested an alternate model in which inactive enzyme resulted from a non-equilibrium process—*i*.*e*., the formation of long-lived misfolded protein during expression (Fig. 3D) (*20*). We varied the temperature and Zn^2+^ concentration present during folding, as temperature is known to affect folding efficiency (*21, 22*), and as PafA binds multiple Zn^2+^ ions during the folding process. We then measured reaction rates under identical assay conditions, so that any observed rate changes must arise from differences during folding that persisted over time (Fig. 3E). Many mutations had differential effects on observed catalytic activity when expressed at 23 vs. 37°C (“T-Effect”) or with different concentrations of Zn^2+^ (“Zn-Effect”), with T- and Zn-Effects in different regions of the enzyme (Fig. 3, F to I, and fig. S33). These results strongly support the presence of persistent non-equilibrium folding effects (Fig. 3D) (*20, 21, 23*). A second prediction of the misfolding model was also met by our data: PafA variants with WT activity but with different misfolded fractions (*i*.*e*., variants falling along the diagonal blue dashed line in Fig. 3C) were equally impeded in *k*_cat_/*K*_M_ and *k*_cat_ but unaltered in *K*_M_ (fig. S34). PafA folding thus apparently involves one or more branchpoints that are sensitive to temperature and Zn^2+^ and lead to active PafA or a long-lived inactive state (Fig. 3D).

### Altered folding pathways promote a long-lived inactive state *in vitro* and *in vivo*

Misfolding could be an artifact of high-throughput on-chip expression, or could also arise during standard expression *in vitro* and possibly *in vivo*. To test for chip-induced misfolding effects, we selected 19 mutants for off-chip expression via *in vitro* transcription/translation and kinetic characterization (Fig. 3C, light green points, and table S3). Activities were similar off and on-chip (fig. S35), suggesting that the chip is not responsible for the observed misfolding.

Native gels and kinetic assays provided additional support for and information about the misfolded state. Mutants with reduced activity had an additional band of distinct mobility when expressed at high temperature but not when expressed at low temperature (Fig. 3J, misfolded state “M”). Transient treatment with thermolysin, a protease that cleaves within exposed hydrophobic regions that occur in unfolded or misfolded proteins, resulted in loss of M but not the native state (native state “N”, fig. S36A) (*24*). Nevertheless, despite degradation of the majority of the protein (present as M), enzyme activity was unaffected (fig. S36B), indicating that M lacked significant activity and that N and M did not equilibrate over the hours taken to express and carry out these experiments.

Our observation of a long-lived inactive state for PafA raised the question of whether analogous misfolding occurs in cells, where cellular machinery can assist protein folding. To test folding in cells, we recombinantly expressed 20 variants in *E. coli* that did and did not undergo temperature-dependent misfolding *in vitro* (Fig. 3C, dark green points, and table S4). Expression *in vivo* was also temperature dependent, with changes in apparent *k*_cat_/*K*_M_ values for the *in-vivo* expressed PafA mutants that correlated with the change observed *in vitro* (*r*^2^ = 0.38, Fig. 3K). Circular dichroism (CD) spectra of a purified misfolding mutant provided independent evidence for a structural alteration that accompanied misfolding *in vivo*. WT PafA exhibits identical CD spectra when expressed at 37 or 23°C; in contrast, the CD spectrum for Y103G matches WT when the mutant is expressed at low temperature (23°C) but not at higher temperature (37°C; fig. S37). The observed difference at 37°C suggests a loss of about one third of PafA’s α-helical character in M (table S5). Together, these results suggest that cellular folding conditions and chaperones are insufficient to prevent mutations from causing PafA to misfold and form a long-lived inactive state. A tendency to form kinetically-stable misfolded states may therefore exert a selective pressure and influence the fitness landscape of proteins in cells (*25*–*29*).

### Dissecting the origins of observed catalytic effects

HT-MEK assays allow us to quantify and dissect the degree to which observed changes in activity arise from changes in the amount of expressed protein, the amount that is correctly folded, and the catalytic efficiency of the correctly folded enzyme. Below, we isolate the catalytic effects for our PafA variants. We then take advantage of HT-MEK’s ability to provide quantitative kinetic and thermodynamic constants for multiple substrates and inhibitors and use these data to probe PafA’s functional architecture and catalytic mechanisms at a global level.

To remove expression and folding effects, we quantified the amount of PafA expressed in each reaction chamber, as described above (see “HT-MEK device and experimental pipeline”). Then, after combining measurements across chambers and experiments for each variant, we calculated the fraction active by using the *k*_cat_/*K*_M_ values measured for substrates with different rate-limiting steps (cMUP and MeP; Fig. 3C; Supplementary Text S3 and S4). Mathematically, we represented the datapoint for each PafA variant in Fig. 3C as a superposition of a catalytic component (Fig. 3C, blue solid line) and a misfolded component (Fig. 3C, diagonal blue dashed line) and solved for both (Fig. 4A and fig. S38). The solid line in Fig. 3C defines the relationship between cMUP and MeP hydrolysis by *active* PafA and the dashed line describes the equal loss in observed activity for the two substrates due to *inactive* PafA (fig. S38). With this procedure, we were able to quantify catalytic effects 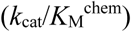 for 946 of the 1036 PafA variants and obtain upper limits of 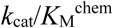 for an additional 65 variants (fig. S39). Deleterious catalytic effects were found for mutations at 161 of PafA’s 526 positions (Fig. 4B, figs. S40 and S41, and table S6). Mutations at an even larger number of positions, 232, gave folding effects.

**Fig. 4.**
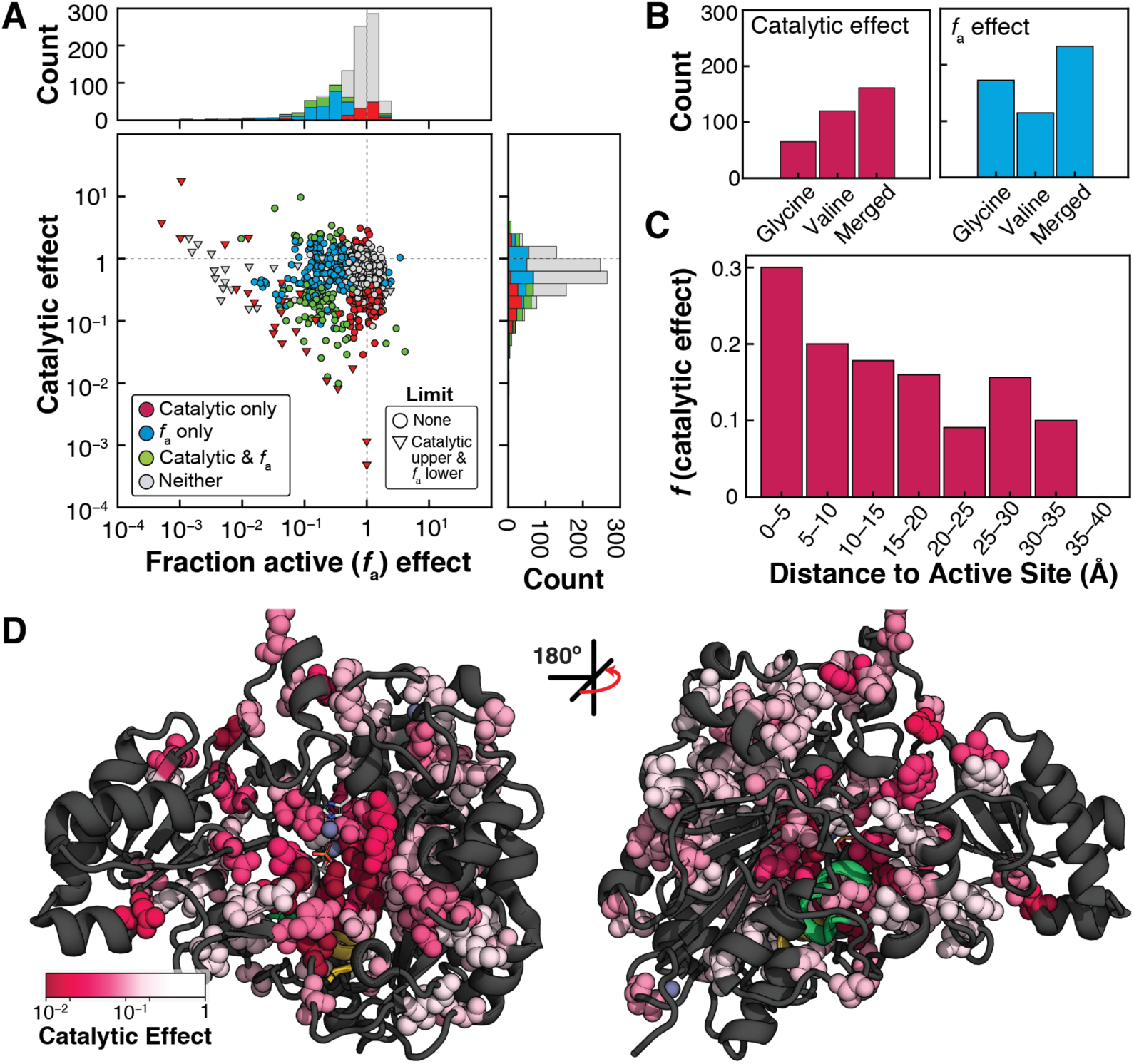
Separating mutational effects on catalysis from changes in the enzyme fraction active (*f*_a_). (**A**) Catalytic effects and *f*_a_ effects (relative to wild-type PafA) for all variants, colored by effect (see Materials and Methods), with the distributions of effects projected along the axes (stacked). (**B**) Total counts of mutants with catalytic (*p* < 0.05) and *f*_a_ effects (*p* < 0.01) for the valine and glycine libraries, and the total number of positions (of the 525 total measured residues) with effects (“Merged”; see fig. S41 for details). (**C**) Fraction of non-active-site mutants with deleterious catalytic effects (*p* < 0.05) as a function of distance from the active site (5 Å bins, see table S6). Representation of the 161 positions with deleterious catalytic effects on the PafA structure, corresponding to the “Merged” set in panel **B**. Residues with deleterious effects (>5-fold down from WT) are colored and shown as spheres; positions at which both mutations have catalytic effects are colored by the more deleterious effect.

The largest catalytic effects cluster in and around the active site, with the fraction of residues with mutational effects diminishing with distance from the active site (Fig. 4C and table S6). While positions giving catalytic effects tend to cluster, the pattern of effects is asymmetric and complex (Fig. 4D and see below). To better understand these patterns and to relate overall effects to the specific mechanisms that PafA uses in catalysis, we developed an approach we refer to as Functional Component Analysis (FCA). In FCA, we first define Functional Components (FCs), which describe different energetic and functional relationships in the catalytic cycle based on prior mechanistic insights for PafA and other AP superfamily members (*16, 30, 31*). We then attempt to attribute observed mutational effects to alterations in specific FCs using the extensive quantitative measurements enabled by HT-MEK, thereby defining the functional architecture of PafA.

### Functional Component 1: Effects through the O2 phosphoryl oxygen atom

Our first Functional Component is derived from the observation that removal of two active site side chains, K162 and R164, renders PafA an equally potent phosphate mono- and diesterase (*16*) (fig. S42); these residues interact with the phosphoryl oxygen that is anionic in monoesters but esterified in diesters (O2, Fig. 5A) (*30*). Thus, interactions that support catalysis by K162 and R164 are expected to disrupt monoester but not diester hydrolysis, and we define FC1 to reveal these effects (FC1 = Δ^diester^/ Δ^monoester^, where Δ = (***k***_**cat**_/***K***_**M**_)^mutant^/(***k***_**cat**_/***K***_**M**_)^WT^). While the simplest expectation is that mutations to residues neighboring K162 and R164 will have FC1 effects, we cannot predict how large and how varied these effects are, how far they extend from the active site, whether there are remote regions that have large effects, or whether effects extend to the enzyme surface. In addition, we cannot predict whether residues affecting FC1 also contribute to other catalytic mechanisms, represented as other Functional Components below. A comprehensive accounting of each residue’s effects on each identified FC is a necessary step to understand an enzyme’s functional architecture—how the residues and structures that surround and extend from the active site potentiate the functions carried out within the active site.

**Fig. 5.**
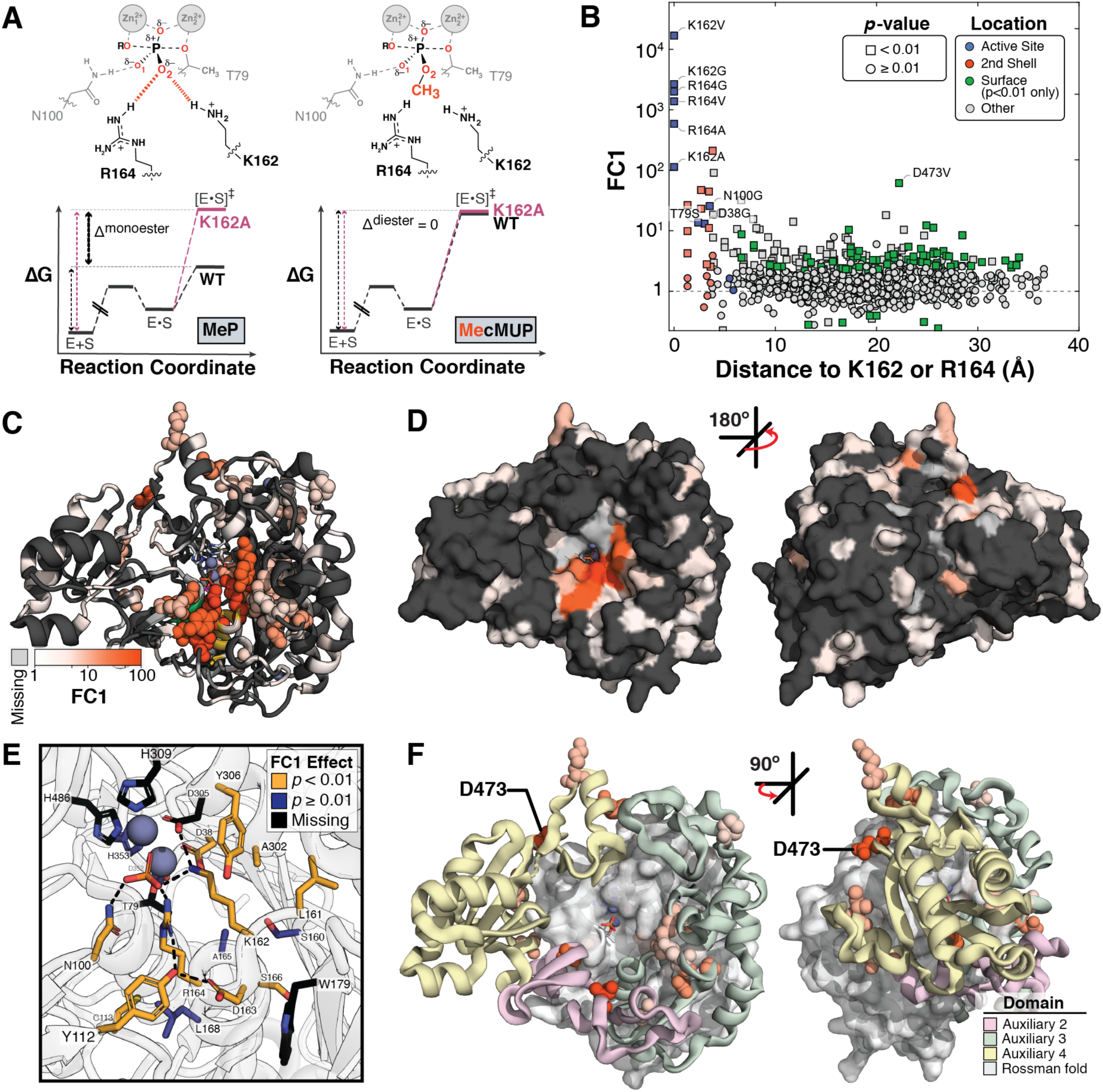
Functional Component 1: catalytic effects through the O2 phosphoryl oxygen atom. (**A**) Schematic of the transition states for reaction of MeP (left panel) and MecMUP (right panel), highlighting the position of the methyl group on the O2 phosphoryl atom of MecMUP (orange) and its interactions with K162 and R164. Thin black and pink arrows are proportional to *k*_cat_/*K*_M_ and the thick black arrow denotes their difference (Δ ^monoester^ and Δ ^diester^). The ratio of *k*_cat_/*K*_M_ effects (or the difference between Δ ^monoester^ and Δ ^diester^ in Δ G space) gives FC1. (**B**) FC1 values for variants as a function of the minimum distance of each to K162 or R164, with active site, 2^nd^ shell, and significantly affected surface residues colored. (**C**) PafA positions with FC1 effects (*p* < 0.01) when mutated to valine or glycine, colored by FC1 magnitude: >5-fold effects are shown as spheres, with ribbon coloring for positions with ≤5-fold FC1 effects. (**D**) PafA surface representation with FC1 effects colored as in **C**. (**E**) FC1 effects of active site and 2^nd^ shell residues. Residues with significant effects are colored yellow; those without FC1 effects and unmeasured residues are blue and black, respectively (see table S7). (**F**) Distal (≥3^rd^ shell) positions with >5-fold FC1 effects shown as colored spheres. The PafA Rossmann core is shown as a gray surface and auxiliary domains 2–4 as colored ribbons.

To measure diester activity on-chip, we synthesized a fluorogenic PafA phosphate diester substrate suitable for HT-MEK (MecMUP, see Materials and Methods), then measured *k*_cat_/*K*_M_ for the PafA mutant libraries (high *K*_M_ values for the noncognate diesterase activities preclude estimation of *k*_cat_ and *K*_M_ separately) (fig. S43) (*16*). We obtained *k*_cat_/*K*_M_ values for 857 of the 1036 mutants, and upper limits for an additional 178 (fig. S44, A and B). As for the monoesterase measurements described above, on-chip diester rate constants matched off-chip measurements (fig. S44C). To enhance detection of mutations with preferentially-reduced monoesterase activity (FC1), we compared measured rates directly, without correcting for folding effects (fig. S45A), as folding affects both substrates equally and the direct comparison yields stronger statistical inference (fig. S45B).

Many mutants have FC1 effects: 88 Val and 93 Gly mutations (fig. S45C), corresponding to 156 of the 494 measurable positions throughout PafA (of a total of 526 positions; Fig. 5, B to D). Seven of the ten measurable non-active site residues directly contacting K162 or R164 (2^nd^ shell residues) exhibited FC1 effects (Fig. 5E, fig. S46, A and B, and table S7A), consistent with frequent 2^nd^ shell effects observed in directed evolution experiments in multiple systems (*32*– *34*). Of the three active site Zn^2+^ ligands with measurable effects upon mutation, we observe an FC1 effect for D38G (which accepts a hydrogen bond from K162) but no effect for D352G and H353V (which do not interact with K162 or R164) (Fig. 5E and table S7B). The active site variants T79S and N100G also had FC1 effects (Fig. 5E and tables S7A and S9), consistent with coupling between active site residues due to shared contacts with K162 and R164 (Fig. 5E).

Although the largest effects were observed for active site residues and next-largest for the 2^nd^ shell residues, there was no additional drop in effect size after the 3^rd^ shell; indeed, the majority of residues with >10-fold FC1 effects were found in the 3^rd^ shell and beyond (15 of 23; fig. S46 and table S9). Several of these residues (4) lie at the enzyme surface (Fig. 5D and fig. S46), consistent with the hypothesis that enzymes possess a reservoir of allosteric potential and suggesting that HT-MEK can be used to identify regions that are potential sites for allosteric inhibitors and drugs (*35*–*39*).

PafA has three of four possible non-terminal auxiliary domains found within the AP superfamily (ADs 2–4, see Supplementary Text S5) that sit around the surface of the universally conserved Rossmann fold (Fig. 5F and fig. S47). ADs 2 and 4 are present in both AP superfamily phosphate mono- and diesterases, whereas AD 3 contains K162 and R164 and is considerably more extensive in the monoesterases than in the diesterases (figs. S47 and S48, and table S10). Despite these apparent functional and evolutionary differences, FC1 effects are found extensively in all three ADs (fig. S49A and B, table S11) and the effects within each AD span similar magnitudes (fig. S49C). The largest FC1 effect outside of the active site or 2^nd^ shell comes from a solvent-exposed surface residue, D473, within AD 4 (Fig. 5F, yellow); a dramatically larger effect for the valine substitution (>60-fold vs. <2-fold for D473G) suggests that a change in local folding may allosterically disrupt the O2 site 20 Å away. Intriguingly, the truncation of AD 3 in AP superfamily phosphate diesterases eliminates the direct interaction between ADs 3 and 4 seen in PafA and other monoesterases (figs. S1 and S47). We speculate that the residues in AD 3 that exhibit FC1 effects functionally link PafA’s surface (D473) to its active site (K162 and/or R164) (Fig. 5F). Future multi-mutant cycle experiments (*40*–*42*) via HT-MEK may allow dissection of these functional pathways and their underlying structural properties.

### Functional Components 2 and 3: Effects on phosphate affinity

To provide catalysis, enzymes must bind their transition states more strongly than they bind their substrates, as otherwise the energetic barrier for the reaction and reaction rate would remain the same as in solution (*43, 44*). Enzymes must also limit the binding of substrates and products to allow sufficient turnover in the presence of higher substrate and product concentrations (*45, 46*). For these reasons, ground state destabilization has been considered a possible mechanism for enhancing enzyme function, and there is evidence for electrostatic ground state destabilization by PafA and other AP superfamily members via electrostatic repulsion between the anionic nucleophile, T79 in PafA, and the negatively-charged phosphoryl oxygens (Fig. 6A) (*16, 31, 47*). For PafA, mutating T79 to serine increases the affinity for inorganic phosphate (P_i_), the reaction’s product and a ground-state analog, by 100-fold, while in *E. coli* AP the nucleophile S102G mutation increases affinity >1000-fold (*16, 31*); the S102G mutation ablates the negatively-charged group adjacent to the phosphoryl anion and the chemically conservative T79S substitution presumably allows greater mobility and an ability to reduce electrostatic repulsion (*16*). We therefore define the second Functional Component, FC2, by *strengthened* P_i_ binding 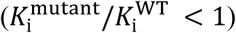 that suggests a reduction in ground state destabilization upon mutation.

**Fig. 6.**
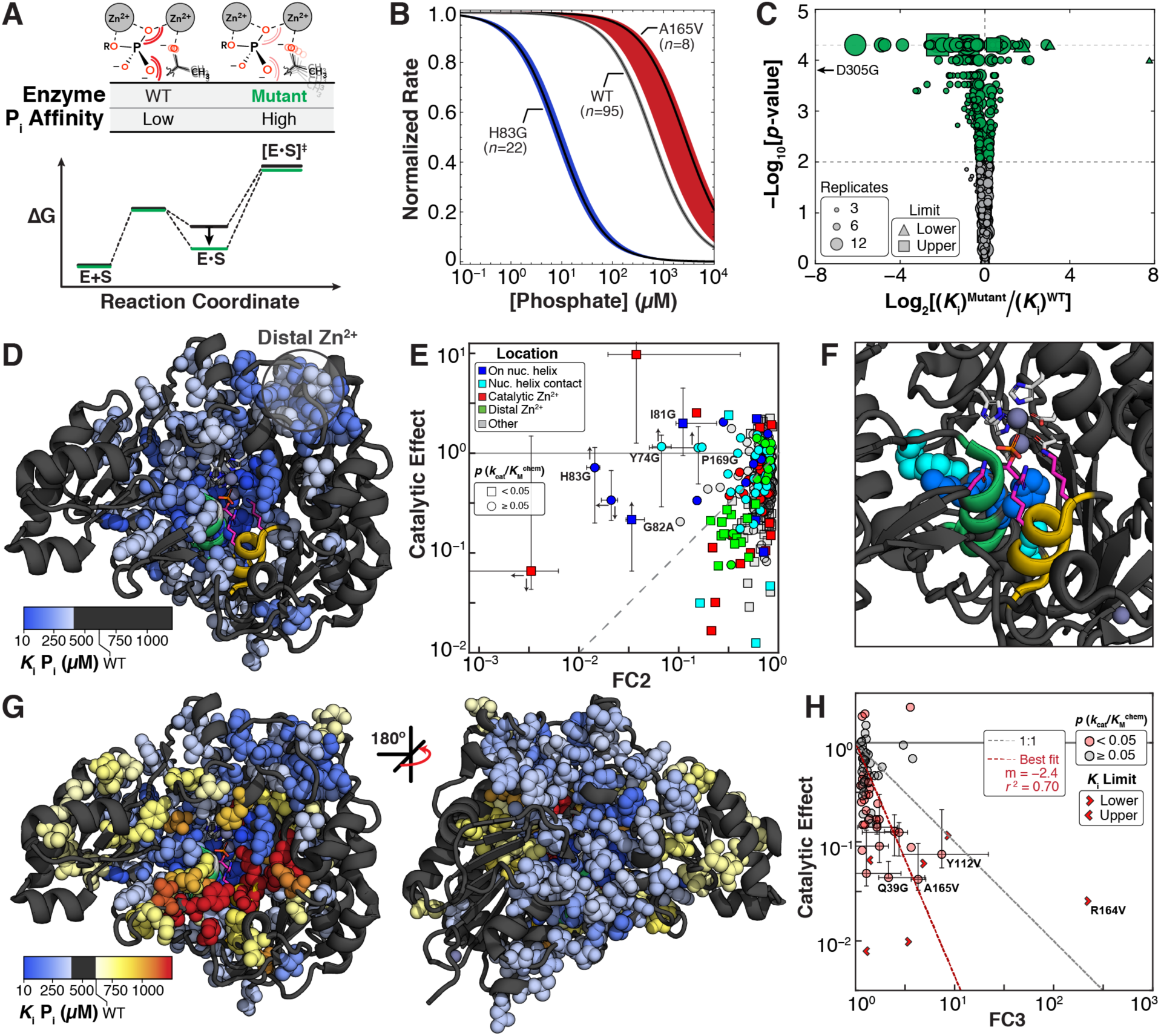
Functional Components 2 and 3: mutational effects on affinity for inorganic phosphate. (**A**) Reaction coordinate diagram for hypothetical mutant (green) in which ground state destabilization is diminished relative to the WT (black). The schematic illustrates the model of a mutation increasing flexibility of the T79 nucleophile yielding an FC2 effect. (**B**) Median P_i_ inhibition curves for wild-type and two representative mutants with FC2 (H83V) and FC3 (A165V) effects. The colored regions denote 99% confidence intervals (CIs) on the medians of replicate *K*_i_ measurements. Inhibition curves were measured using between 6 and 12 P_i_ concentrations, with an average of seven replicates per mutant. (**C**) Volcano plot of *K*_i_ P_i_ effects for glycine and valine scan mutants (*p* < 0.01, green markers; *p* ≥ 0.01, gray markers). Plotted values with log_2_[(*K*_i_)^mutant^/[(*K*_i_)^WT^] < 0 are equivalent to log_2_(FC2) and those with log_2_[(*K*_i_)^mutant^/[(*K*_i_)^WT^] ≥ 0 are equivalent to log_2_(FC3). (**D**) PafA structure with positions for which *K*_i_ P_i_ is >1.5-fold tighter than the WT (at *p* < 0.01) in either or both scans shown as spheres, coloring by the larger effect when both had FC2 effects. The nucleophile helix (green) is largely obscured by spheres representing FC2 effects on and abutting it. (**E**) Scatter plot of catalytic and FC2 effects (*p* < 0.01), colored by location in PafA. Error bars correspond to 95% confidence intervals (CIs) determined from bootstrapping, and the mutants were labeled if the upper bound of the 95% CI was >5-fold below the WT *K*_i_ P_i_. Up and down arrows denote upper and lower 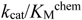>limits (see Materials and Methods), and left and right arrows denote upper and lower *K*_i_ P_i_ limits. (**F**) PafA active site with the five mutants possessing large FC2 effects without 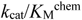 effects shown as spheres, corresponding to the labeled mutants in **E**. Active site residues are colored pink. (**G**) Front and back views of FC2 and FC3 effects (spheres); FC2 effects are those shown in **D** and FC3 effects are shown for *p* < 0.01. (**H**) Scatter plot of catalytic and FC3 effects for mutants with FC3 effects (*p* < 0.01). Error bars correspond to 95% CIs, as in **E**. The red dashed line is the best fit line to mutants with significant FC3 and catalytic 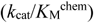 effects, excluding the active site mutants.

Conversely, active site residues typically make both ground state and transition state interactions, so their removal weakens binding and diminishes catalysis, in some instances to a similar extent (so-called “uniform binding”) and in other cases preferentially destabilizing the transition state (*48*–*51*). As expected, mutations to the PafA active site residues that interact with the phosphoryl O1 and O2 oxygen atoms, N100, K162, and R164, weaken P_i_ binding, and N100 and K162 mutations have even larger effects on catalysis, indicating preferential transition state stabilization (fig. S51) (*16*). We therefore define the third Functional Component (FC3) by effects that weaken P_i_ binding 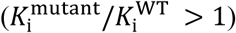, and we assess FC2 and FC3 below.

To measure inhibition constants for the valine and glycine scanning libraries, we quantified rates of cMUP hydrolysis as a function of P_i_ concentration and fit the observed initial rates to a standard competitive inhibition model (Fig. 6B). HT-MEK-determined inhibition constants agreed with those measured previously in off-chip assays for active site mutants (fig. S52) and are of higher precision than kinetic constants (Fig. 6C), as uncertainty in total enzyme concentration and the fraction of active enzyme does not affect measured inhibition constants (see Materials and Methods; Supporting Auxiliary Files contain per-experiment reports of all inhibition measurements).

We uncovered 331 mutants that increased P_i_ affinity (FC2) and 73 that decreased P_i_ affinity (FC3) (Fig. 6C and fig. S53). Thus, about one-third of all mutants measurably altered phosphate affinity, and, remarkably, four times as many mutations enhanced binding as weakened it. As it is highly unusual to *enhance* function by random variation, this observation suggests that residues at many positions are evolutionarily selected to prevent tight P_i_ binding. Mutations with ground state destabilization effects (FC2) were located in an extended yet spatially-contiguous region that included the helix containing the T79 oxyanion (“nucleophile helix”), the catalytic Zn^2+^ ions, and the distal Zn^2+^ site (Fig. 6D and tables S12 and S13).

The catalytic effects 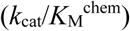 for mutants with FC2 effects ranged from insignificant to 100-fold reductions (Fig. 6E and fig. S54). For most mutants, the magnitude of the catalytic effects was greater than that of the FC2 effects, consistent with multiple inactive (or less active) mispositioned conformations of the T79 oxyanion (or other catalytic groups), only some of which ameliorate electrostatic repulsion. Nevertheless, five mutants with the largest FC2 effects had little or no catalytic effect. These residues formed a spatially contiguous subregion on and adjacent to the nucleophile helix (Fig. 6E and F, and fig. S54). These results suggest that the resting mutant enzyme remains aligned for catalysis but that the T79 oxyanion can rearrange to reduce electrostatic repulsion in the presence of bound Pi.

Mutants that weakened P_i_ binding (FC3 effects) were also located in a region that was contiguous (Fig. 6G). The largest FC3 effects were near active site residues K162 and R164 and yielded qualitatively similar 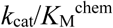 but quantitatively smaller effects (Fig. 6H, red; fig. S55). These results, like those for FC2, are consistent with a preponderance of mutations having larger effects on catalysis than binding, presumably reflecting the greater constraints present in the transition state (*52*).

Residues with FC2 and FC3 effects formed an interface (Fig. 6G), with a small number of residues at this interface yielding either an FC2 or FC3 effect depending on the identity of the mutation (fig. S56). FC2 effects extended significantly further from the active site than FC3 effects (Fig. 6G and figs. S57 and S58), perhaps reflecting a need for extensive interactions to mutually constrain the relative positions of the T79 anionic nucleophile and the substrate in the presence of strong electrostatic repulsion.

FC2 and FC3 are mutually exclusive by definition, as mutations that strengthen P_i_ binding cannot simultaneously weaken P_i_ binding. However, a systematic comparison of mutational effects across Functional Components 1–3 reveals that many mutations outside the active site preferentially affect either FC1 or FC2 (fig. S59). Further, several mutations preferentially alter FC2 without dramatically altering FC1, reducing the fraction of active enzyme, or reducing overall catalysis 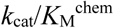; fig. S19). The ability to selectively alter particular enzyme properties via specific mutations provides a potential starting point for attempts to engineer enzymes with desired kinetic and thermodynamic constants and behaviors.

### Functional Component 4: Rates of phospho-enzyme hydrolysis

Linking observed mutational rate effects to their physical and chemical origins requires knowledge of the step that is being observed—*i*.*e*., the rate-limiting step. While pre-steady state approaches (*e*.*g*., stopped flow, rapid quench) are the gold standard for determining rates of individual reaction steps, these approaches do not readily scale to large libraries. Here, we leverage prior mechanistic knowledge and our ability to measure multiple reactions for multiple substrates to distinguish individual PafA reaction steps and determine mutational effects on these for 992 PafA variants.

For PafA, each steady state kinetic constant can have a different rate-limiting step (Fig. 7A). The steady state kinetic constant *k*_cat_/*K*_M_ can be limited by substrate binding or chemical cleavage of the substrate to form the covalent enzyme-phosphate (E–P_i_) species (Fig. 7A, *k*_1_ and *k*_chem,1_ steps). As described above, comparing *k*_cat_/*K*_M_ values for reactions of cMUP and MeP allowed us to determine mutational effects on the chemical step and to discover that many PafA mutants form a long-lived misfolded state (Figs. 3 and 4). The steady state rate constant *k*_cat_ can be limited by hydrolysis of E–P (*k*_chem,2_) or by dissociation of P_i_ subsequent to hydrolysis (*k*_off,Pi_; Fig. 7A). Here, we determined mutational effects on *k*_chem,2_ using measurements of *k*_cat_ for cMUP and *K*_i_ for P_i_, and *k*_chem,2_ effects define FC4 (fig. S60, and see Supplementary Text S6). Seven mutants changed the rate-limiting step from E–P_i_ hydrolysis to P_i_ release (*k*_off,Pi_ < *k*_chem,2_; fig. S61). The ease of this transition is consistent with the observation of naturally-occurring alkaline phosphatases of the AP Superfamily with either of these steps rate limiting (*53*).

**Fig. 7.**
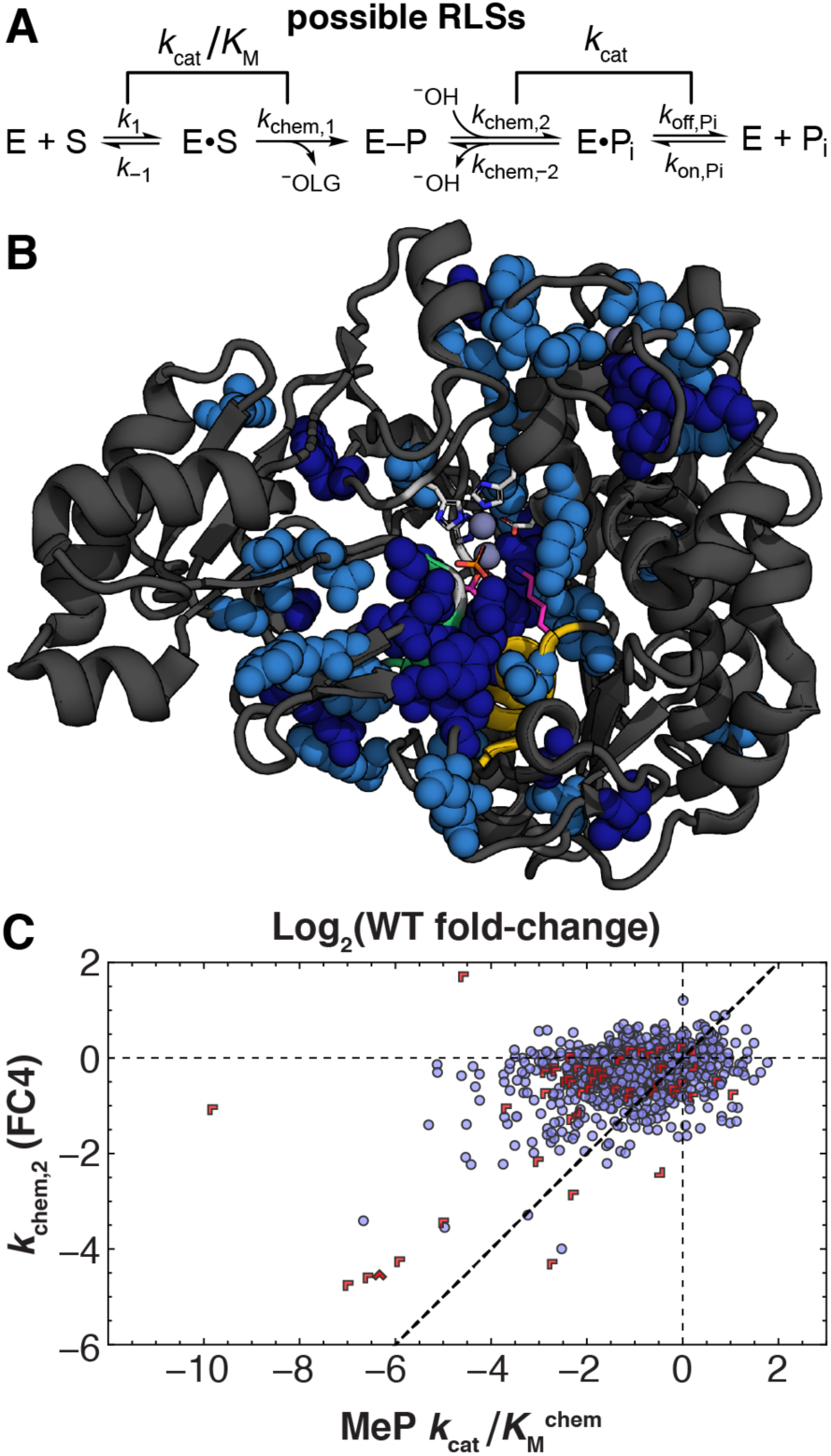
Functional Component 4: mutational effects on phospho-enzyme intermediate hydrolysis. (**A**) Schematic of the PafA catalytic cycle with possible rate-limiting steps under saturating (*k*_cat_) and sub-saturating (*k*_cat_/*K*_M_) conditions. (**B**) Structure showing positions with decreases in *k*_chem,2_ (*p* < 0.05 and *p* < 0.1 in dark and light blue, respectively) upon Val and/or Gly substitution (spheres). Positions lacking an estimate of *k*_chem,2_ for both substitutions are light gray ribbons. (**C**) Scatter plot comparing the magnitude of mutational effects on the rate of phospho-enzyme hydrolysis (*k*_chem,2_, or FC4) and effects on 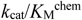 for MeP. Limits are denoted by red chevrons pointing in limit directions (see Supplementary Text S6).

Overall, we find 18 Val and 36 Gly mutants that reduce turnover via reductions in *k*_chem,2_ (fig. S60), and these spatially cluster near the three active site phosphoryl oxygen binding residues (N100, R164, and K162) and extend to the distal Zn^2+^ site (Fig. 7B). The FC4 residues overlap substantially with those affecting 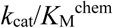 and Functional Components 1–3 (table S15). In the simplest scenario, the mutations reducing *k*_chem,2_ would be a subset of those reducing 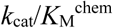, as 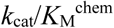 includes a phosphoryl transfer chemical step but also involves an additional step (ligand binding). Our data support this expectation, as mutations affect either 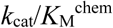 alone or 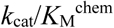 and *k*_chem,2_ to a similar or smaller extent (Fig. 7C).

### Evolutionary constraints revealed by comparisons between phylogeny and functional parameters

Natural evolution provides a mutagenesis and selection experiment at a massive scale. Highly conserved residues typically perform critical functions (*e*.*g*. active site residues) and coevolution between residues can reveal molecular contacts required for folding and stability (*54*–*56*). Nevertheless, conservation cannot reveal all aspects of function, most basically because multiple functional properties combine to give the observed conservation parameter. HT-MEK and FCA have allowed us to dissect and quantify effects of mutations at every residue on misfolding, catalysis, and four Functional Components. Here, we assess the correlation between conservation and these functional effects using a metagenomic alignment of 14,505 AP superfamily sequences with PafA-like active site residues (T79, K162, and R164; fig. S62).

Fig. 8A shows conservation at each position throughout the PafA structure and provides the starting point for these comparisons. Residues with high conservation scores often have no apparent functional effect, and some with lower scores have effects (figs. S63 and S64). Overall, observed *k*_cat_/*K*_M_ values (MeP hydrolysis) correlated with the information content at a given position (Spearman rho = −0.4; Fig. 8B and fig. S63A), with this correlation increasing slightly for more perturbative substitutions (fig. S63B). Systematic comparisons of information content and the mutational effects on misfolding, catalysis 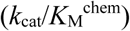 and each of the four Functional Components gave Spearman’s correlation coefficients with magnitudes ranging from 0.1 to 0.25 (Fig. 8B). The observation of multiple correlations with these functional parameters underscores the complexity of evolutionary pressure and responses and the need for additional information to understand and interpret the sequence record.

**Fig. 8.**
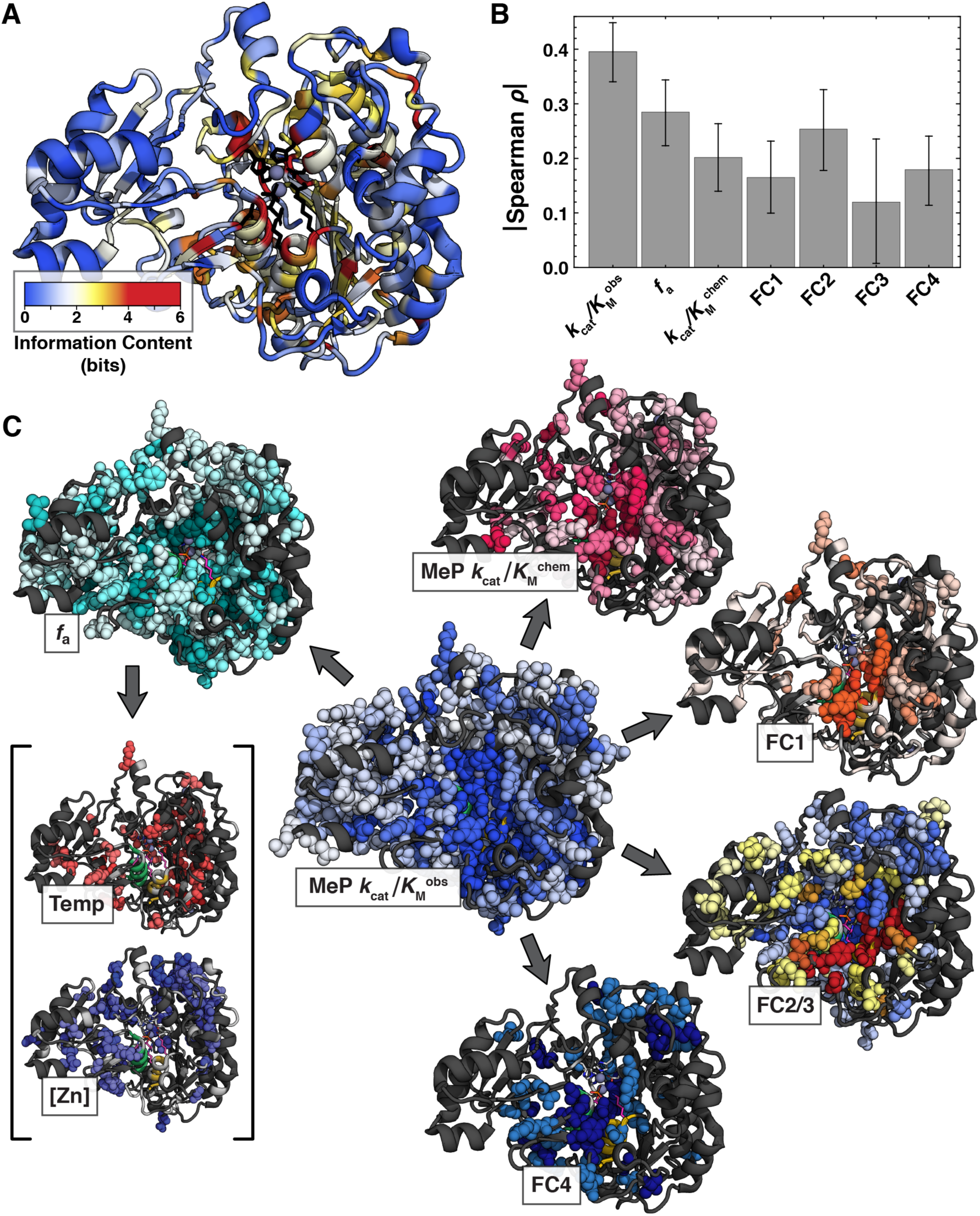
Anatomy of an enzyme. (**A**) Conservation of residues across a broad metagenomic alignment of PafA-like AP superfamily members. (**B**) Spearman’s rank correlation coefficients for comparisons of functional parameters against residue conservation (information content). Error bars denote bootstrap 95% CIs. (**C**) Decomposition of PafA function in terms of Functional Component Analysis and misfolding effects (*f*_a_), including the folding effects sensitive to temperature or Zn^2+^. Structures show the composite of Val and Gly substitution effects for each parameter. Temperature and Zn effect, MeP 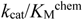, and FC1–4 structures are reproduced from Figs. 3 to 7; the *f*_a_ and MeP 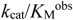 structures show the union of Val and Gly mutant effects (spheres; *p* < 0.01), coloring by the larger effect when both were measured. The FC2/3 structure shows the union of FC2 (blue) and FC3 (yellow/red) effects.

## Discussion

HT-MEK uses automatic valved microfluidics to carry out high-throughput expression, purification, and comprehensive biochemical characterization of enzymes at an unprecedented scale. Using HT-MEK, we quantified the effects of 1036 single amino acid substitutions within PafA phosphatase, measuring >650,000 reaction time courses to obtain over 5000 kinetic and thermodynamic constants for multiple substrates and inhibitors across our mutant library. HT-MEK can be applied to the vast number of enzymes whose activity can be monitored via fluorescence either directly or with a coupled assay. Here, we used both fluorogenic substrates and a sensitive coupled assay with a previously established fluorescent P_i_ sensor (*18*). This coupled assay renders HT-MEK immediately applicable to ATPases, GTPases, helicases, protein chaperones, polymerases (with pyrophosphatase present), and many additional enzymes.

Remarkably, initial assays quantifying the effects of glycine and valine substitutions at every PafA position revealed deleterious *k*_cat_/*K*_M_ effects for nearly half of the mutants. Many large effects clustered around the active site, but effects also extended throughout the enzyme and to the surface, and many of the largest effects were distant from the active site (Fig. 8C, fig. S32). These findings mirror results obtained from DMS studies of other enzymes that often reveal distal effects on product formation or organismal fitness (*12, 57*–*59*). However, HT-MEK allowed us to determine how mutations affect steady state rate parameters and individual reaction steps and to interrogate their mechanistic origins.

HT-MEK-based measurements of *k*_cat_/*K*_M_ values for phosphate monoester substrates with different rate limiting steps suggested the presence of a long-lived inactive population for most PafA variants (Fig. 3C); additional experiments indicated that none of the effects originated from equilibrium unfolding and instead resulted from mutations promoting the formation of a misfolded state (Fig. 3D to I). The analogous behavior observed for these mutants in *E. coli* and correlation of misfolding effects with phylogenetic conservation is consistent with a selective pressure to avoid misfolding *in vivo* and with growing evidence that kinetic factors affect stable protein expression in cells (Fig. 3K) (*20, 26, 60, 61*). The strong dependence of activity on temperature and Zn^2+^ concentration present during expression suggests that mutations promoting formation of the inactive state may disrupt co-translational folding (*20, 62*). We speculate that highly stable proteins like PafA and other secreted enzymes may be more prone to forming long-lived kinetically-trapped states. Moreover, these results highlight the need to explicitly decouple mutational effects on folding and catalysis, whether in cell-based or *in vitro* high-throughput assays, as a prerequisite for interpreting effects and understanding the functional roles of residues throughout an enzyme.

By removing effects from misfolding and ensuring that chemistry was rate limiting, we were able to quantify mutational effects on intrinsic catalytic activity and on a panoply of additional kinetic parameters. To gain insights from this multidimensional dataset, we expressed these effects in terms of previously identified PafA catalytic mechanisms using a framework we term Functional Component Analysis (FCA; table S16). FCA exploits the rich data provided by HT-MEK to generate an atlas of PafA functional “anatomy” with unprecedented detail, revealing that PafA is comprised of large, well-defined, and spatially contiguous regions of residues that share catalytic signatures upon mutation (Fig. 8C). While catalytic effects are largest for mutations to residues in and contacting the active site, as expected, effect sizes do not decay continuously with distance from the active site. Instead, catalytic (and Functional Component) effects extend from the active site to the distal Zn^2+^ site, previously assumed to play a purely structural role, and to other surface regions. These observations affirm that the enzyme beyond the active site is not a passive, monolithic scaffold, but rather aids in positioning active site residues and/or modulating dynamics that contribute to function.

Comparing patterns of mutational effects between FCs reveals the idiosyncratic nature of atomic environments and reveals “architectural” solutions for different catalytic interactions and strategies. From prior mechanistic studies, we expected residues with FC1 effects (monoesterase:diesterase specificity) to contact either the K162 and R164 residues that directly interact with the O2 phosphoryl oxygen atom or to lie on or contact the “monoesterase helix” present only in AP superfamily monoesterases (Fig. 5A and figs. S1 and S47). For FC2 (ground state electrostatic destabilization), we anticipated effects around the T79 anionic nucleophile and associated “nucleophile helix” (Fig. 6, D to F, and fig. S1). While residues with each effect generally appeared in the expected vicinities, mutations on and around the monoesterase helix largely did not give FC1 effects (Fig. 5C to F) while those on and around the nucleophile helix gave the largest FC2 effects (Fig. 6F and fig. S54). These differences may reflect a need for more interactions to secure the nucleophile helix against ground state electrostatic repulsive forces and prevent its rotation and translation. Consistent with this model, many more glycine than valine mutations led to increased P_i_ binding (222 Gly vs. 109 Val, fig. S53), potentially because the additional space afforded by side chain ablation allows structural rearrangements to reduce electrostatic repulsion without disrupting favorable binding interactions.

While the functional regions we observe have superficial similarities to sectors and other measures of evolutionary co-variation and co-conservation (*63*–*67*), our data report directly on sequence-function relationships in ways that sequence analyses alone cannot. Positions at which mutations promote misfolding tend to be more conserved, consistent with the fact that all AP superfamily members must fold to function. However, decoupling mutational effects on folding from those on catalysis reveals that 702 of our 1036 PafA mutants influence at least one functional parameter, with different mutations affecting different aspects of function (Fig. 8C).

For sequence analysis, even if a particular algorithm could identify these positions, it could not link the sequence conservation or variation to the particular aspect(s) of function under adaptive selection. Selective pressures likely vary, and vary in unknown ways, through evolution. For example, the selective pressures on PafA, and thus its adaptive responses, will differ with the available P_i_ at the organism’s physical location, whether available P_i_ varies temporally, and whether there is competition for P_i_ from other organisms present in the same ecological niche. In sequence comparisons, a mutation with a critical role in responding to these variable adaptive pressures would be poorly conserved, changing frequently despite being tightly linked to survival, and thus appear to be unimportant. HT-MEK opens new opportunities to link molecular sequence and function to evolutionary outcomes in the future.

Most distal mutations with catalytic effects preferentially affect the transition state over binding, underscoring the complexity of proteins as catalysts (Fig. 6H). The need for interactions between residues throughout the enzyme to ensure optimal catalysis highlights the tremendous challenges inherent in *de novo* design of efficient enzymes, as it suggests computational efforts must consider interactions between prohibitively large numbers of residues. The detailed anatomical maps provided by HT-MEK and FCA—and the ability to distinguish catalytic from folding effects—can guide computational and experimental mutagenesis to improve enzymes by identifying residues and regions that affect particular aspects of catalysis. In addition, the ability to rapidly iterate through design-build-test cycles will generate information not only about which designs were successful, but the mechanisms by which other designs failed, information that should prove valuable in developing next-generation *de novo* enzyme design.

Enzymes are the targets of many therapeutics, are altered in genetic diseases, serve as workhorse tools for modern molecular biology, and play critical roles in industrial processes. The rapidity and low per-assay cost, combined with the wide availability of fluorescence-based enzymatic activity assays, make HT-MEK a powerful tool for future applications across all these areas. In basic research, the large and highly quantitative datasets provided by HT-MEK can greatly extend and even supplant mechanistic studies that now use traditional site-directed mutagenesis approaches in many cases. When combined with recent advances in gene synthesis, HT-MEK can facilitate rapid functional characterization of metagenomic variants across superfamilies, providing a critically-needed additional dimension to phylogenetic analyses. In medicine, HT-MEK can rapidly determine functional effects of human allelic variants of unknown significance and systematically identify candidate allosteric surfaces within currently “undruggable” therapeutic target enzymes. We anticipate HT-MEK contributing to these and still more areas of basic and applied biology, medicine, and engineering.

## Acknowledgements

We thank members of the Herschlag and Fordyce laboratories for discussions and review of the manuscript, Michael Madsen for technical assistance, Kara Brower for microfluidic mold fabrication and optics assistance, and Shawn Costello and Susan Marqusee for helpful discussions.

## Author contributions

C.J.M., D.A.M., and F.S. designed experiments and collected and analyzed data, M.J.A. collected and analyzed data. C.S. provided assistance with statistical analyses. D.A.M. and E.A. generated the sequence alignments and performed phylogenetic analyses. S.A.L. contributed to hardware automation and software development. D.H. and P.M.F. conceived and supervised the project and acquired funding. C.J.M., D.A.M., D.H., and P.M.F. wrote and revised the manuscript with input from all authors.

## Data and materials availability

Summary tables of all kinetic and thermodynamic parameters measured for each mutant are included in the Supplementary Materials. All data acquired in this study are available in an Open Science Foundation Repository (https://osf.io/ez9xv/); code used to obtain and process images and fit kinetic and thermodynamic parameters is available at the Fordyce lab Github repository (https://github.com/FordyceLab).

