## Supplementary Materials for "Revealing enzyme functional architecture via high-throughput microfluidic enzyme kinetics"

**Figures S1 to S64** **35**

**Tables S1 to S16** **99**

**Captions for Auxiliary Supplementary Materials, Data S1 and S2** **115**

**Captions for OSF Repository Data S1 to S14** **116**

**References** **117**

### Materials and Methods

#### HT-MEK device fabrication

##### Photolithographic mold production

Molding masters for flow and control layers of microfluidic devices were designed in AutoCAD (Autodesk, Inc.) and fabricated on 4″ single-sided silicon test-grade wafers (University Wafer) using transparency masks printed at 30000 dpi (Fineline Imaging) via standard photolithography procedures (*68*, *69*). Flow molds were composed of the following layers: (1) a ~5 µm uniform layer of SU-8 2005 (Microchem) covering the entire wafer surface to improve feature adhesion, (2) a ~15 µm layer of AZ50 XT (Capitol Scientific) subjected to a slow final hard bake to generate rounded valve features, and (3) a ~15 µm layer of SU-8 2015 (Microchem) to generate square flow channels. Control molds were composed of a single ~25 µm layer of SU-8 2025 (Microchem). All mask designs are available on our lab website (http://www.fordycelab.com).

##### PDMS device fabrication

Two layer "push-down" valved MITOMI polydimethylsiloxane (PDMS) devices (**Fig. S2**) were fabricated from molding masters largely as described previously (*68*). Prior to fabrication, silicon wafer molding masters were silanized via vapor deposition of trichloro(1H,1H,2H,2H-perfluorooctyl)silane (Sigma) for 10 min to prevent PDMS adhesion. To fabricate thick control layers, a 1:5 ratio of RTV 615 (R.S. Hughes) crosslinker:base was mixed using a THINKY centrifugal mixer for 3 min, degassed for 3 min at 4000 rpm, and then poured onto control molds positioned within an aluminum foil holder. After pouring, thick PDMS layers were degassed for ~1 h in a vacuum chamber under house vacuum. To fabricate thin flow layers, a 1:20 ratio of RTV 615 (R.S. Hughes) crosslinker:base was mixed in a THINKY centrifugal mixer for 3 min and then spin coated onto the flow mold (10 s ramp at 500 rpm, 133x acceleration; 75 s spin coating at 1725 rpm, 266x acceleration). Flow and control molds were subsequently baked at 80°C for 40 min and 60 min, respectively. After baking, the thick control layer was peeled off of the silicon wafer, individual devices were cut out of the layer using a scalpel, and control inlet holes were punched using a 20 gauge needle and a small drill press (Technical Innovations). Control layers were then aligned by eye to flow layers remaining on their molding master using a stereoscope and this entire assembly was baked for an additional 50 min. Fully baked devices were cut off of the flow wafer using a scalpel and stored for subsequent alignment to printed plasmid arrays, as described below. All fabrication was performed in the Stanford Microfluidics Foundry class 10,000 clean room.

#### Construct and library generation

##### WT and active site PafA-eGFP fusion construct generation

Wild-type PafA was C-terminally tagged with enhanced green fluorescent protein (eGFP) via a flexible glycine/serine linker (GGGSGGGGSG) and subcloned into the optimized PURExpress expression plasmid (referred to as the "DHFR control template" in the manufacturer protocol) (**Fig. S4** and **OSF Repository Data S1**).

##### Generation of scanning mutant libraries

Mutant libraries were generated using QuikChange site-directed mutagenesis (Agilent Technologies, Inc.), in a 96-well plate format, detailed below and diagrammed in **Fig. S18**).

**Mutagenic primer design** Mutagenic primers optimized for QuikChange site-directed mutagenesis were designed in batch using a custom script (https://github.com/FordyceLab/designQuikChangePrimers). Given a desired single-residue mutation, the script chooses an optimal mutant codon based on the minimum number of nucleotide changes required and generates a series of potential primers 30–45 nt in length with the mutation site <2 nt from the center of the primer. The algorithm then scores each primer based on its annealing temperature ($T_\mathrm{m}$, calculated using Primer3) (*70*), propensity to form hairpins, and the presence or absence of G or C nucleotides at either end. To compute a final primer score, each of these parameters was weighted by its relative importance for successful primer design, with the $T_\mathrm{m}$ weighted most heavily. $T_\mathrm{m}$ scores were assigned a maximum value for $T_\mathrm{m}$>78°C, a half-maximal score for 77°C<$T_\mathrm{m}$<78°C, and a score that linearly decreased to zero for $T_\mathrm{m}$ <77°C. The full code, weights used, and associated comments are available in the GitHub repository.

Optimized primers were ordered in 96-well plates at 10 nmol synthesis scale and purified using standard desalting (Integrated DNA Technologies, Inc.). Forward and reverse primers were premixed and normalized to 6 nmol per well. Primer pairs were re-suspended in 120 µL Milli-Q water and then diluted to working stock solutions at 1.25 µM for use in downstream PCR reactions.

**PCR mutagenesis and digestion of template DNA** Mutagenic PCRs were performed in 96-well PCR plates using a modified version of the QuikChange site-directed mutagenesis protocol (Agilent Technologies, Inc.). First, we added 6 µL of a 1.25 µM working primer solution to each well. Next, we added all additional components of the reaction (including PfuTurbo AD polymerase; Agilent Technologies Inc., catalog no. 600259) from a master mix prepared on ice and added immediately before thermocycling. Final reactions contained 2.5 µL 10X Cloned Pfu DNA polymerase reaction buffer AD, 300 nM forward and reverse primer, 200 µM dNTPs, 5% (v/v) DMSO, 2.5–5.0 ng template plasmid, and 1.25 U PfuTurbo AD polymerase in a total volume of 25 µL. Reactions were heated to 94°C for 60 s and then cycled 14 times using the following parameters: a 60 s denaturation step at 95°C, a 60 s annealing step at 55°C, and a 12 min extension step at 68°C. After cycling, a final elongation step was performed for 60 s at 68°C and reactions were cooled to 4°C. For reactions that did not yield the desired mutation (assessed by Sanger sequencing, described below), repeated reactions were performed with an increased annealing temperature of 60°C. To digest remaining template DNA after PCR, we added 20 U of DpnI (New England Biolabs Inc., catalog no. R0176S) and 2 µL CutSmart buffer (New England Biolabs Inc., catalog no. B7204S) to 10 µL of PCR reaction and incubated at 37°C overnight.

**Transformation and plasmid purification** For bacterial transformation of mutant plasmids, we added 1 µL of DpnI-digested plasmid to 5 µL of ice-cold NEB 5-alpha competent *E. coli* (high efficiency; New England Biolabs Inc., catalog no. C2987). Cells were incubated with DNA on ice for 30 min, heat-shocked for 30 s at 42°C in a thermocycler, and then recovered on ice for 2 min. Outgrowth at 37°C was carried out for >1 h in 300 µL SOC medium (New England Biolabs Inc., catalog no. B9020S) while shaking. We then plated the entire volume of outgrowth solution for each mutagenesis reaction on an LB agar plate containing either 100 µg/mL ampicillin or 50 µg/mL carbenicillin and incubated overnight at 37°C. A well-isolated single colony was picked from each plate and used to inoculate 5–10 mL of LB broth containing the same concentration of antibiotic. Plates with more than one colony were stored at 4°C in case picking additional colonies was necessary, as discussed below. Following overnight growth at 37°C, cells were pelleted at 1800×g for 10 min in an Eppendorf 5810R centrifuge. Plasmid DNA from cell pellets was either purified immediately or the pellet was frozen and stored at −20°C for later purification. To minimize the risk of contamination between mutants, all plasmid purifications were carried out individually using commercially available spin miniprep kits (either a QIAprep spin miniprep kit, Qiagen, catalog no. 27104, or GeneJet plasmid miniprep kit, ThermoFisher Scientific, catalog no. K0502).

**Confirmation of desired mutation by Sanger sequencing** Prior to mutagenesis, we sequenced the full-length (~2 kb) wild-type PafA-eGFP template construct for all stock solutions to ensure that no unwanted mutations were propagated to the entire library due to errors in the template DNA. After mutagenesis, each mutant was confirmed by Sanger sequencing using a sequencing primer designed to cover the region containing the putative mutation. While the majority of reactions (~75%) were successful on the first attempt, we encountered 3 types of unsuccessful outcomes: (1) lack of transformants after PCR and digestion, (2) transformants that harbored either the wild-type plasmid or an unanticipated mutation in the primer region (presumably due to a DNA synthesis error), or (3) a mixture of wild-type and mutant plasmids within a single colony, evident as mixed peaks in the Sanger sequencing chromatogram. In cases without transformants, we redesigned primers and/or attempted new PCR reactions using a different annealing temperature. For cases with transformants but not the correct mutation, we picked and sequenced additional colonies two or three additional times; if still unsuccessful, we redesigned mutagenic primers and began the process again.

**Probability of incorporating an unwanted mutation outside of the sequencing read** As PfuTurbo is a high-fidelity polymerase with an error rate of $1.3\times10^{-6}$ mutations/bp (*71*), the probability of the polymerase introducing an error in the PafA-eGFP construct outside of the primer region is approximately 0.003 per mutant (0.3%). As a result, in the course of making the 1036 mutants in this work, we expect approximately 3 errors in the entire library. Given that many nucleotide mutations would be synonymous, this provides an upper bound on the expected number of unwanted amino acid substitutions within the library.

#### Optics, pneumatics, and custom software

**Imaging setup** We imaged devices using a Nikon Ti-S Microscope equipped with a motorized XY stage (Applied Scientific Instrumentation, MS-2000 XYZ stage), cMOS camera (Oxford Instruments, Andor Zyla 4.2), solid-state light source (Lumencor, SOLA SE Light Engine), and automated filter turret equipped with an eGFP filter set (Chroma Technology Corp., part no. 49002), DAPI filter set (Semrock Inc., catalog no. DAPI-1160B-NTE), and a custom "PBP" filter set (Semrock Inc., 427/10 bandpass excitation filter, catalog no. FF01-427/10-25; 470/22 bandpass emission filter, catalog no. FF01-470/22-25; 442 nm dichroic beamsplitter, catalog no. DI03-R442-T1- 25X36; mounted in a TE2000 filter cube, catalog no. NTE). All imaging was performed using a 4X objective (CFI Plan Apochromat $\lambda$ 4X NA 0.20, Nikon) at 2x2 binning (1024x1024 pixels) with exposure times as follows: eGFP, 500 ms; DAPI, 50 ms or 200 ms; PBP, 100 ms. Apparatus temperatures were measured throughout assays using a thermistor sensor (Thorlabs Inc., catalog no. TSP01, data logger; catalog no. TSP-TH NTC, external thermistor).

**Pneumatic control** Microfluidic devices were run using a custom-built pneumatic manifold (*72*) with pneumatic valve state controlled by custom open-source software (https://github.com/FordyceLab/RunPack).

**Hardware automation and image processing software** To automate device valve actuation, reagent flow, and image acquisition during HT-MEK experiments, we developed a custom, publicly-available Python package (https://github.com/FordyceLab/RunPack). This package provides an extensible unified hardware interface for imaging and stage control via the µManager (*73*) *MMCore* Python API; valve control using AcqPack, an in-house package for actuation of the HT-MEK WAGO-controlled pneumatic apparatus (https://pypi.org/project/acqpack) (*74*); and temperature measurement using PyVISA (https://pypi.org/project/PyVISA/). Images were processed using a publicly-available in-house image processing package written in Python (https://github.com/FordyceLab/ProcessingPack).

We analyzed images to yield progress curves and kinetic and thermodynamic binding constants using a custom suite of Mathematica notebooks (Wolfram Research, Inc.), available at https://github.com/FordyceLab/HT-MEK_KineticThermodynamicFitting. Data from each device was initially processed individually and then replicates across multiple devices were combined and normalized using an additional notebook (*cf. Multi-Tier measurement strategy*, below). Summary CSV and PDF files containing the data and summary plots from each experiment, and for each mutant across experiments, are available in the OSF Repository Data (https://osf.io/ez9xv/).

#### Expressing and purifying enzymes on HT-MEK

##### Printing plasmid library and aligning the device

Sequence-validated plasmid solutions were transferred directly into a series of 96-well plates for storage and array printing. For each plate, we generated a series of "daughter" plates by diluting 20 µL of each DNA solution from the "parent" 96-well plate to a final volume of 50 µL with Milli-Q water. For printing, DNA solutions (10 µL from each well of the daughter plates) were transferred from 96-well to 384-well plates using a Biomek FX Automated Workstation (Beckman Coulter, model A31843). After transfer, all 384-well plate solutions were evaporated to dryness and the DNA was re-suspended in 15 µL of 1% (w/v) BSA (UltraPure, ThermoFisher Scientific, catalog no. AM2616), 12 mg/mL trehalose dihydrate (Sigma Life Science, catalog no. T9531), and 200 mM NaCl (Sigma Life Science, catalog no. 71376). Final plasmid DNA concentrations in the 384-well plate ranged from ~40–100 ng/µL.

Plasmid DNA arrays were printed in tandem (two per slide) on custom 2″ x 3″ epoxysilane-coated slides (ThermoFisher Scientific SuperChip) using a custom microarrayer fitted with 75 µm silicon tips (Parallel Synthesis Technologies, catalog no. SMT-S75). To prevent carry over between spots on the array, we washed silicon tips two times after depositing each mutant for 10 s in hot water (>70°C) followed by 8 s of drying under vacuum. After printing, PDMS devices were manually aligned to DNA arrays under a stereo microscope and then bonded to the slides for 12 h at 95°C on an aluminum hot plate (Torrey Pines Scientific). Device alignment was performed in the Stanford Microfluidics Foundry class 10,000 clean room to minimize the risk of particulates interfering with device function.

##### Surface patterning for enzyme immobilization

Surface functionalization was carried out largely as described previously (*14*), with several critical modifications (**Fig. S3**). Initial flow and control line pressures were set to 32 and 3–4 psi, respectively. To prevent premature solubilization of the DNA spots from osmotic transfer of water from the control layer, we filled control lines with a 0.55 M NaCl solution. To begin surface functionalization, the Button and Neck valves were pressurized (closed) to prevent solubilization of the plasmid DNA spots and protect Buttons and areas beneath them from fluid flow, respectively. To passivate PDMS walls and the epoxysilane surface outside of the Button valve, we introduced a 5 mg/mL BSA solution (ThermoFisher Scientific, catalog no. AM2616) throughout all channels. Solutions introduced into the device were initially primed through the inlet tree and directed to waste for ~5 min to purge any air from the sample lines and inlet. After this initial purge step, solutions were flushed through the chip for 30 min with the Neck and Button valves pressurized (closed); between injections, we flowed 1X phosphate buffered saline (PBS, ThermoFisher Scientific, catalog no. 10010023) for 10–15 min to fully wash device channels. To selectively functionalize slide surfaces beneath the Button valve for recruitment and surface-immobilization of eGFP-tagged enzymes, we introduced a 2 mg/mL solution of biotinylated BSA (bBSA; ThermoFisher Pierce, catalog no. 29130) for 5 min with the Button valves pressurized (to ensure channels were filled with bBSA), then opened Button valves and continued to flow bBSA for an additional 30 min to allow binding. After bBSA introduction, we washed the entire device with PBS for 10 min. Next, we introduced 1 mg/mL neutravidin (NA) (Thermo Scientific, 31000) for 30 min to bind to the bBSA previously deposited on epoxysilane slide surfaces beneath Button valves and the Button itself, followed by another 10 min PBS wash. To saturate any NA bound outside of the Buttons, we again pressurized (closed) the Button valves (to protect NA-functionalized surfaces) and introduced bBSA for an additional 30 min, followed by another 10 min PBS wash. Finally, we depressurized (opened) the Button valves and introduced either a 100 µg/mL solution of biotinylated anti-GFP antibody (b-anti-eGFP; Abcam, catalog no. ab6658) or a mixture of biotinylated anti-GFP antibody and bBSA (final concentrations of 100 µg/mL and 50 µg/mL, respectively) for 15 min, thereby tuning the density of potential attachment sites and ultimate effective enzyme concentration (as described below). This surface patterning procedure ultimately yielded a bBSA/NA/b-anti-eGFP stack non-specifically adsorbed to epoxysilane slide surfaces (**Fig. S4**) capable of recruiting expressed enzymes to surfaces. We then again pressurized (closed) Buttons, washed with PBS for 10 min, closed the outlet valve (to prevent evaporation) and maintained the device under positive pressure to await the manual expression step. The device was stable in this state for at least 18 h.

##### On-chip expression of PafA-eGFP constructs

Printed PafA-eGFP mutant DNA spots were solubilized and protein was expressed using the PURExpress expression system (New England Biolabs Inc., catalog no. E6800L). To prepare the expression mixture, we mixed 10 µL of PURExpress Solution A with 7.5 µL of Solution B and allowed this mixture to incubate on ice for 30–45 min, as this incubation period increased yields of PafA-eGFP >2-fold. Immediately before flowing the reaction mixture into the chip, we added 2.5 µL of a 1 mM $ZnCl_2$ solution, 0.5 µL of 40 U/L RNasin ribonuclease inhibitor (Promega Corporation, catalog no. N2515), and 4.5 µL of nuclease-free water, and mixed gently. With the Neck valves closed, we flowed this expression mixture over the device for 12–15 min to ensure that PURExpress solution fully filled all channels. To introduce the PURExpress mixture into all expression chambers and solubilize the DNA, we closed the chip outlet valve, opened the Neck valve, and increased the flow pressure from 3.5 to 4–4.5 psi, thereby dead-end filling DNA chambers under pressure; the filling process was followed visually using a stereo microscope. When chambers were 90–95% full, we closed the Button, Neck, and Sandwich valves, isolating adjacent protein chambers from one another and closing the Button valves to prevent premature binding of contaminating enzyme in the case of any leakage between expression chambers within the flow path. We then placed the slide bearing the device onto a pre-warmed aluminum hot plate (Torrey Pines Scientific) and allowed enzyme constructs to express for 45 min at 37°C. During expression, we opened the Sandwich valves and flowed 1X PafA reaction buffer (100 mM MOPS, 500 mM NaCl, 100 µM $ZnCl_2$, pH 8.0) continuously through the device to counteract evaporation and wash out any enzyme leaking from expression chambers that would otherwise contaminate other chambers of the chip. Following expression, we removed devices from heat and incubated them for 90 min at room temperature in the dark to allow for maturation of the eGFP fluorophore.

##### Surface immobilization of expressed enzyme and subsequent buffer exchange

After enzyme expression, we removed device-slide assemblies from hot plates and mounted them on a fully automated Nikon Ti-S microscope (components described above). To bind expressed enzymes to antibody-functionalized regions of the glass protected by the Button and to the Button itself, we closed Sandwich valves (to isolate adjacent reaction chambers from one another), opened Button valves (to expose binding surfaces), and then opened the Neck valve, allowing diffusion of expressed enzyme into the reaction chamber. Binding was allowed to proceed for 15–60 min. During binding, we imaged chambers across the device periodically in the eGFP channel, and then closed Button valves to stop binding when the desired enzyme concentration ([E]) was reached.

As wild-type PafA is inhibited by inorganic phosphate ($P_i$), accurate measurement of kinetic rate constants requires that any $P_i$ produced during the on-chip transcription and translation reactions be washed out prior to kinetic assays. To fully remove any remaining $P_i$ from reaction chambers, we flushed the entire device with 1X PafA reaction buffer with the Neck valves open for 60 min immediately after completion of binding.

##### Removing cross-chamber enzyme contamination with surfactant washing

Accurate measurement of kinetic and thermodynamic parameters for mutants with compromised catalysis requires ensuring that chambers are free from even very low levels of contaminating WT-like enzyme molecules from other chambers non-specifically adsorbed to chamber walls or slide surfaces. Initial kinetic measurements for "empty" chambers (lacking a plasmid) were consistent with approximately 0–0.3 nM levels of contamination with WT-like enzyme molecules. To eliminate this cross-contamination, we washed each device (with Button valves closed) with four 5-min treatments of 1% (w/v) sodium dodecyl sulfate (SDS) in MES buffer at pH 6.0. This treatment lowered background enzymatic activity of non-specifically adsorbed enzyme by 50-fold (on average) while preserving the fluorescence of the enzyme-GFP fusion constructs under the Button and the activity of specifically bound enzyme. While we also tested use of chaotropes to denature non-specifically adsorbed enzyme, these agents resulted in death of Button-protected surface-immobilized enzymes. We suspect that this enzyme death under the Button occurs because the small hydrodynamic radius of these chaotropes allows them to quickly diffuse beneath the Button valve, unlike monomeric and micellular SDS.

#### Imaging and quantification of expressed enzymes

##### Imaging of eGFP signals

To detect and quantify expressed enzymes, we imaged devices in the "eGFP channel" using an eGFP filter set (Chroma Technology Corp., part no. 49002) with acquisition times of 500 ms per image. Using 2x2 binning, imaging of the full device typically required 49 tiled images with 10% overlap (~32 chambers per tile).

##### Automated detection of eGFP fluorescence spots

Measuring expression for a given mutant requires mapping each chamber to a unique positional index within the array (each of which contains a known spotted plasmid) and quantifying the eGFP signal beneath the Button valve in that chamber.

**Initial image processing** To correct for position-dependent differences in excitation intensities, we first applied a flat-field correction to each tile (*75*). We then generated complete images of each device by stitching the overlapped flat-field corrected tiles. After stitching, we corrected for PDMS autofluorescence by background-subtracting an image of the device acquired before construct expression. Finally, we divided each stitched image into 28x56 sub-images (stamps), each centered on and containing a single chamber, using the 4 corner chambers as reference positions. The ID of each mutant was then assigned to its corresponding stamp.

**Quantification of eGFP fluorescence within each chamber** Prior to the start of each kinetic assay, we quantified eGFP intensities beneath Button valves. To find the centroid positions of Button valve regions containing bound enzyme within each stamp, we used a combination of a sparse grid search (to identify the circular region of maximal summed fluorescence intensity) and a dense local grid search (to refine this position). We then summed intensities from all pixels within a fixed radius of 15 pixels, larger than the physical radius of the Button. To account for spatially-dependent differences in background intensities, we calculated a local background intensity considering pixels in an annulus concentric with, abutting, and larger than the Button bounding circle. Local background summed intensity was normalized to the Button area (#pixelsButton/#pixelsannulus) and subtracted from the summed Button region pixel intensity to yield a background-subtracted summed intensity used to quantify [E] in all downstream analyses.

##### eGFP intensity calibration curves

Quantitative measurement of the kinetic constants $k_{\mathrm{cat}}$ and $k_{\mathrm{cat}}/K_{\mathrm{M}}$ hinges on accurate measurement of enzyme concentration ([E]). To convert measured eGFP intensities to effective [E], we patterned a device with anti-GFP antibody as described above, iteratively introduced 3 nM solutions of eGFP (BioVision, catalog no. 4999) with 2% (w/v) BSA (UltraPure, ThermoFisher Scientific, catalog no. AM2616), incubated for 1200–7200 s, and then quantified bound fluorescence at saturation (**Fig. S5**). Prior to titrations, chamber walls were pre-equilibrated with eGFP by flowing a 3 nM or 5 nM solution of eGFP with 2% (w/v) BSA (UltraPure, ThermoFisher Scientific, catalog no. AM2616) in 1X PafA reaction buffer for 10 min, washing for another 10 min with 1X PafA reaction buffer alone, and repeating this procedure three times. Control experiments in which Button valves were opened to expose anti-GFP-antibody-coated surfaces in the absence of introduced eGFP solution established that non-specifically-bound eGFP does not significantly migrate to Button surfaces and increase observed intensity. Bound fluorescence intensities underneath the Button were quantified as described above. Prior to saturation, measured summed intensities increased linearly with each additional injection of eGFP (as expected, given that the $K_D{}^{\alpha\text{-eGFP}} \ll 3$ nM). A linear fit to these first (typically 4–9) titration points (~0–20 nM) yielded calibration parameters sufficient to convert measured intensity to [E] with an error of generally <±5% (95% CI) in the linear fit region. We performed this procedure for every microscope and light source setup (and again upon any adjustments to the instruments), as differences in illumination intensity and optics can alter fit parameters.

#### Synthesis of fluorogenic cMUP and MecMUP substrates

Syntheses of cMUP and MecMUP were adapted from previously published work. For both substrates, methyl 7-hydroxycoumarin-4-acetate (Sigma-Aldrich, IDF00040) was used as the coumarinyl (phenolic) starting material, and intermediates were not purified.

To synthesize the phosphate monoester 2-(2-oxo-7-(phosphonooxy)-2H-chromen-4-yl)acetic acid (cMUP; **Fig. S9** compound **3**), we first generated the diethyl phosphate triester (compound **1**) analogously to previous reports by reacting the coumarin phenol (1.0 eq) with triethylamine (2.0 eq) in dry dichloromethane, adding dimethyl chlorophosphate (2.0 eq) dropwise while stirring under argon overnight (*76*). Crude products were subjected to an aqueous workup consisting of aqueous HCl, saturated $NaHCO_3$, and saturated NaCl, and organics were dried over $MgSO_4$ and concentrated to dryness under reduced pressure to afford the crude diethyl triester (compound **1**) as a golden oil. Adapted from a previously reported procedure (*77*), the crude phosphotriester coumarinyl methyl ester was subjected to alkaline saponification with stoichiometric LiOH monohydrate (2.0 eq) in 1:1 THF:$H_2O$. Products were acidified to pH 2–4 with HCl and extracted into dichloromethane. Organics were washed with saturated NaCl, dried over $MgSO_4$, and concentrated to dryness under reduced pressure to afford the crude deprotected diethyl triester **2** as an oily residue. Finally, deprotection of the phosphate ester was achieved with trimethylsilyl iodide (TMSI), also analogously to a previous reported synthesis (*77*). An oven-dried flask under an atmosphere of argon was charged with a solution of crude compound **2** in dry dichloromethane via syringe. To the stirring solution, TMSI (4.0 eq) was added via syringe, drop-wise, and the reaction was allowed to stir for 2 h in the dark. Then, the reaction was quenched by drop-wise addition into excess $H_2O$ and adjusted to pH ~8 with 0.5 M NaOH. The biphasic mixture was allowed to stir, and the mixture was readjusted to pH 8 repeatedly with 0.5 M NaOH until the pH stopped dropping. The aqueous layer was separated and then washed with two portions of dichloromethane. The aqueous layer was then dried under reduced pressure in a heated bath to give a white-yellow solid of crude phosphate monoester **3** (subsequently purified and characterized, below).

The phosphate diester substrate 2-(7-((hydroxy(methoxy)phosphoryl)oxy)-2-oxo-2H-chromen-4-yl)acetic acid (MecMUP; **Fig. S9** compound **6**), was also prepared via a phosphate triester. First, the phosphate methyl triester was prepared as described for ethyl triester (**1**), to afford crude compound **4** as an oil. Then, crude reaction products were singly deprotected with LiCl (2 eq) in refluxing acetone for 2 h, as previously reported, with altered workup as follows (*78*). Upon evaporation to remove solvent, products were resuspended in water and acidified to pH 2–3 with 0.5 M HCl, washed with two portions of dichloromethane, and the aqueous phase was again concentrated, yielding an oil of impure compound **5**. Finally, to generate the desired phosphodiester carboxylic acid, the carbonyl ester was deprotected by basifying an aqueous solution of crude products in water to ca. pH 8 with 0.5 M NaOH, and reacting with LiOH (2 eq) in THF:$H_2O$ (1:1) for ~7 h at room temperature under air, analogously to deprotection of the phosphate monoester. The reaction was halted by acidifying to pH 7 with 0.5 M HCl, and solvent was removed under reduced pressure to afford crude phosphate diester **6** (subsequently purified and characterized, below).

Crude phosphate monoester and phosphate diester products were purified via semi-preparative HPLC using an Agilent 1260 HPLC equipped with a C18 reverse-phase column and eluting with a gradient of 0.1% TFA in $H_2O$ into MeCN. Phosphate monoester substrate eluted as the largest peak (detecting absorption at 360 nm); phosphate diester substrate eluted as a the second principal peak (again detecting absorption at 360 nm). Product-containing fractions were subsequently pooled and lyophilized, yielding off-white powders.

HPLC-purified material was characterized via $^1H$ and $^{31}P$ NMR (externally-referenced to 85% $H_3PO_4$) with a Varian Mercury 400 MHz magnet or Varian Inova 500 MHz magnet and processed with MestReNova 11 (Mestrelab Research) (**Figs. S10** and **S11**). Compound **3** (monoester): $^1H$ NMR (400 MHz, $D_2O$) $\delta$ 7.65 (d, $J$ = 8.5 Hz, 1H), 7.16–7.19 (m, 2H), 6.37 (s, 1H), 3.94 (s, 2H); $^{31}P$ NMR (202 MHz, $D_2O$, externally referenced to 85% $H_3PO_4$ at 0.00ppm) $\delta$ -4.00. Compound **6** (diester): $^1H$ NMR (400 MHz, $D_2O$) $\delta$ 7.66 (d, $J$ = 8.7 Hz, 1H), 7.15–7.19 (m, 2H), 6.38 (s, 1H), 3.93 (app. d, 2H), 3.65 (d, $^3J_{31P,1H}$ = 11.1 Hz, 3H); $^{31}P$ NMR (202 MHz, $D_2O$, externally referenced to 85% $H_3PO_4$ at 0.00ppm) $\delta$ -2.72. Stock solution concentration determination was performed spectrophotometrically by complete hydrolysis using an *E. coli* alkaline phosphatase mutant not inhibited by $P_i$ (R166S) and quantification of generated free leaving group (fluorogenic hydrolysis product) using a standard series of 7-hydroxycoumarinyl-4-acetic acid (cMU; Sigma-Aldrich, IDF00040).

#### Multi-Tier measurement strategy

Measuring kinetic constants for all mutants within the 1036 member valine and glycine scanning libraries requires the ability to accurately quantify initial rates spanning >5 orders of magnitude (Main Text **Fig. 1**). Measurements at the limits of this range pose distinct technical challenges: fast mutants require low concentrations of expressed enzyme to slow reactions to allow multiple measurements before substrate is exhausted, while slow mutants require high concentrations of expressed enzyme and have activities easily obscured by even small amounts of cross-contamination from neighboring chambers. To maximize measurements efficiency and enhance the accuracy and precision of returned catalytic and thermodynamic parameters, we therefore measured rates for all experiments using a three-Tier experimental pipeline (**Table S2**), in which mutants with similar rates were measured in separate experiments ("Tiers"). During tier 1, we assayed all valine and glycine mutants. During tier 2, we reprinted and assayed all mutants >10-fold down in cMUP or MeP hydrolysis, on the "Slow cMUP" or "Slow MeP" devices, respectively. In tier 3, we reprinted and assayed all mutants that remained un-resolvable during tier 2.

#### *In vivo* expression, purification, and assay of control WT PafA and active site mutants

To compare the Michaelis-Menten parameters determined from on-chip measurements with those from traditional off-chip assays, we recombinantly expressed wild-type PafA and 5 active site mutants, and measured their activities using the same substrates as used in on-chip assays. C-terminally Strep-tagged PafA WT and mutants were over-expressed and purified as previously described (*16*). Briefly, 20 mL of starter culture was prepared from plasmid-transformed SM547(DE3) *E. coli* cells in Luria Broth (LB) with 50 µg/mL carbenicillin. The starter culture was added to 2 L of Luria broth and grown at 37°C to an OD600 of 0.6. IPTG was then added to a final concentration of 300 µM to induce protein expression, and cells were incubated on a rotor overnight at room temperature. Enzymes were affinity purified from filtered cell lysate over Streptactin Sepharose resin (Cytiva, catalog no. 17511301) and purity was assessed via SDS-PAGE gels to be >95% for all preps. Enzyme kinetic measurements were collected on Synnergy H4 and Tecan Infinite M200 plate readers, as described previously (*16*), except the build-up of fluorescent cMU from cMUP and MecMUP reactions was followed using excitation and emission wavelengths of 365 nm and 455 nm, respectively.

#### Measuring cMUP hydrolysis on-chip

##### Imaging of fluorescent leaving group (cMU) fluorescence

To detect and quantify the fluorescent product cMU (7-hydroxycoumarinyl-4-acetic acid) of cMUP and MecMUP hydrolysis, we imaged devices in the "DAPI channel" using a DAPI filter set (Semrock Inc., catalog no. DAPI-1160B-NTE) with acquisition times ranging from 50–200 ms and using 2x2 binning.

**Measuring cMU fluorescence standard curves**

To convert product intensities measured during activity assays to product concentrations and correct for position-dependent intensity differences, we measured product (cMU) standard curves for each chamber of each device by sequentially introducing increasing concentrations of cMU product in 1X PafA reaction buffer (0, 2, 5, 10, 25, 50, and 100 µM) and imaging the device after enzyme expression and purification but before kinetic assays. We generated a separate standard curve for each chamber of the device in order to correct for position-dependent excitation intensity differences in the DAPI channel. For each concentration, we (1) closed the Button valves in all chambers across the device to protect surface-immobilized enzyme. (2) We then flowed the solution over the device for 6–8 min to fully equilibrate all chambers. (3) Sandwich valves were then closed to isolate adjacent reaction chambers from one another, before opening the Button valve and iteratively imaging all chambers across the device over time.

**Image processing for cMU fluorescence standard curves** First, we mapped chambers and generated image stamps as described previously (*cf.*, *Imaging and quantification of expressed enzymes*). We then automatically detected chamber center positions from stamps acquired at the highest cMU concentration using a Hough circle transform via OpenCV (*79*). If no circles were initially found, we increased permissiveness of the Hough circle transform using a grid-search of Hough transform "accumulator threshold" and "gradient" parameters. If multiple circles were found, in either case, we selected the circle with highest median pixel intensity. For chambers in which a "best" circle was found following this procedure, we circumscribed the chamber using the center of this circle and a fixed chamber radius, and used the median pixel brightness within this region for all downstream kinetic analysis. These chamber boundaries were subsequently used for all other cMU concentrations and downstream analysis of kinetic assays.

**Standard curve fitting** To generate product standard curves, we identified chamber boundaries as described above, calculated the median intensity for each chamber for each product concentration, and collated these data by chamber. For each chamber, we performed a linear least-squares fit to measured intensities as a function of concentration and used these fit parameters to convert measured intensities to product concentrations in downstream assays.

**Performing kinetic assays**

On-chip cMUP hydrolysis assays were performed in 100 mM MOPS, 500 mM NaCl, 100 µM $ZnCl_2$, pH 8.0 (1X PafA reaction buffer). Substrate concentrations used were generally 10, 25, 50, 100, 200, 500, and 1000 µM, with the addition of 2 µM in a subset of experiments. Before starting reactions, substrate solution was flowed for 6–8 min to fully exchange the solution in the device, with the Button valves closed. To start reactions, the Sandwich valves were closed (to separate chambers from one another and prevent leakage of product) and the Buttons were opened to expose the immobilized enzyme to substrate and start the reaction.

Assays were acquired in ascending order of [cMUP], and, when possible, we performed replicate 10 or 50 µM [cMUP] assays after the 1000 µM assay to check for enzyme death over the course of the full experiment.

**Kinetic assay image acquisition** Enzymatic turnover was quantified by acquiring a time series of images in the DAPI channel (for detection of cMU). Images were iteratively acquired for all chambers across the device over time, saving images with a timestamp that indicated the precise time of acquisition. Exposure times were either 50 or 200 ms. As for eGFP imaging, cMU imaging required 49 tiles (10% overlap, 2x2 binning), so that each full chip image required ~75 s. Most assays were acquired for 65 min, but ranged from 24 min to ~6 h, depending on which experimental tier was being measured. To measure initial rates for both fast and slow enzyme mutants, we initially acquired images as quickly as possible (10 full images spaced 75 s apart) and then acquired images at pseudo-logarithmically increasing time intervals. For our standard 65 min assay, acquisition times were 75

(×9), 150, 300, 600, 900, and 1200 s. Actual acquisition times for each chamber are reported in the per-experiment CSV files.

#### Analysis and quality control of cMUP hydrolysis measurements

All code for analyzing experimental data exist in Mathematica notebooks available on our lab GitHub page (https://github.com/FordyceLab/HT-MEK_KineticThermodynamicFitting). To identify and flag chamber measurements likely contaminated by enzymes from neighboring chambers, we developed and applied a series of quality control procedures as described below (**Figs. S13–S16**). All data and quality control flags are included in the per-experiment CSV and PDF files in the **OSF Repository Data** (https://osf.io/ez9xv).

##### Obtaining initial rate measurements

To process images and obtain kinetic measurements for each chamber, we first calculated the median intensity at each time point using the chamber boundaries defined from the cMU standard curve (see above).

To determine initial rates, we then fit median chamber intensities as a function of time to a linear model and converted fitted rates from RFU to [cMU] using the per-chamber standard calibration curves described above. To ensure that fits consider only time-points in the linear regime, we included only timepoints at which cMU concentrations were <30% of the initial cMUP concentration, $[S_0]$ ([P] $<0.3[S_0]$).

##### Fitting Michaelis-Menten parameters

The parameters $k_{cat}$, $K_M$, and $k_{cat}/K_M$ were determined from a non-linear least-squares fit of the initial rates at each substrate concentration to the Michaelis-Menten equation (**Eq. 1**):

$$v_i = \frac{k_{cat}[E][S]}{K_M + [S]} \tag{1}$$

In all cases, Michaelis-Menten parameters were determined using estimated enzyme concentrations from images acquired immediately prior to starting the series of kinetic assays, based on control experiments in which we measured the activity loss over the full assays series by comparing the rates of the first 10 µM [cMUP] assay to those of the replicate acquired at the end of the full series. Scaling by the enzyme concentrations determined from eGFP intensity measurements obtained immediately before each assay resulted in the rates of the replicate assay being significantly higher than the original measurement, while the unscaled rates are in good agreement (**Fig. S12**). This indicates that the eGFP fluorescence loss is decoupled from any PafA activity loss, which itself is not significant over a single cMUP titration series. This is likely due to enzyme lost from the Button non-specifically binding the chamber walls (rather than remaining in solution and being flushed out). We therefore used the [E] measured before the first assay of the series to determine kinetic parameters.

##### Flagging $K_M$ values outside the range of [cMUP] used

Accurate determination of $K_M$ requires the ability to measure initial rates at concentrations significantly above and below this value. For measurements with fit $K_M$ values >2-fold lower than the lowest [substrate] ($K_M$ < $[cMUP]_{lowest}/2$) for more than 50% of the total replicates, we set $K_M = [cMUP]_{lowest}/2$ and flagged these values as upper limits (8 mutants, including the active site mutants T79S and N100A). $K_M$ upper limits result in $k_{cat}/K_M$ being lower limits, while $k_{cat}$ values can still be determined.

We applied an analogous criterion to flag as lower limits mutants with $K_M$ values higher than the measurable range (where $K_M > 2[cMUP]_{highest}$, for more than 50% of the total replicates); however, no mutants met this criterion.

##### Lower limit of resolution due to non-PafA-driven hydrolysis

Hydrolysis by water and other buffer components imposes an absolute lower limit on the measurable initial rate of *enzymatic* substrate hydrolysis. To quantify rates of non-PafA-mediated hydrolysis on the device, we performed the standard on-chip surface patterning procedure, introduced a catalytically incompetent PafA mutant, T79G, expressed off chip via *in vitro* transcription/translation into one half of the HT-MEK device, and then measured rates of hydrolysis of cMUP across all chambers. To determine apparent first order rate constants, we fit the last 7 points of this time course, avoiding early timepoints in which measured intensities are significantly affected by photobleaching and diffusion between solutions within the flow and control layers of the device, resulting in negative rates. We measured first order rate constants of ~$5.0\times10^{-5}$ $h^{-1}$ and $1.1\times10^{-5}$ $h^{-1}$ at 500 µM cMUP for T79G-containing and empty chambers, respectively (median, $n = 671$ and 895, respectively). These observed rate constants are approximately 5-fold higher and the same as the rate expected for cMUP hydrolysis by water at pH 8.0 based on published values for para-nitrophenylphosphate (pNPP) ($9.7\times10^{-6}$ $h^{-1}$) (*13*), suggesting that the practical limit of resolution for on-chip reactions is largely defined by the uncatalyzed rate of hydrolysis.

##### Flagging based on detected PafA-eGFP expression

To flag chambers with levels of expressed PafA below the limit of accurate on-chip quantification or containing high-intensity spot artifacts resulting from dust, we automatically eliminated any chambers containing a calculated [E] <0.3 nM (**Fig. S13**). Chambers were also manually inspected and flagged based on Button morphologies and the presence of high-intensity pixel artifacts using a scripted GUI.

##### Flagging 2-point initial rate fits to minimize error in measurements of fast enzymes

Imaging all chambers within an HT-MEK device requires ~75 s. This minimum time delay between successive images of a given chamber sets an upper limit on rates that can accurately be measured, as WT PafA and the fastest mutants may hydrolyze a significant fraction (>30 %) of substrate within the first 2 acquisition time-points, leading to underestimation of initial rates (**Fig. S14A**).

To determine the conditions under which systematic underestimation of initial rates leads to systematic errors on fit Michaelis-Menten parameters, we: (1) simulated progress curves for WT-like enzymes hydrolyzing cMUP for enzyme concentrations between 0.1 and 7.5 nM (**Fig. S14B**), (2) simulated sampling at different acquisition times ($t = 2$ s, 37.5 s, and 75 s, corresponding to chambers at the top, middle, and bottom of the device, respectively) (**Fig. S14B**), (3) estimated rates from either a linear fit (for simulated curves in which <30 % of substrate has been hydrolyzed by the second timepoint) or from a 2 point fit (for simulated curves in which <30 % of substrate has been hydrolyzed by the second timepoint), (4) compared these estimated rates to the "true" rates (**Fig. S14B**), (5) repeated this process for every cMUP concentration used in the experiment, and then (6) compared fitted Michaelis-Menten parameters to the "true" parameters used in the simulation (**Fig. S14C**). While fitted $k_{\mathrm{cat}}$ values were not significantly affected, chambers at the bottom of device had fitted $K_{\mathrm{M}}$ values that were inflated by ~20% for chambers with >5 two-point fits (of 7 total) and became progressively more inflated as the number of two-point fits increased (**Fig. S14D**).

To test if we observed these predicted trends in the HT-MEK data, we plotted the fitted $K_{\mathrm{M}}$ for all WT PafA constructs within tier 1 (fast enzymes, low [E]) and tier 2 (slow enzymes, high [E]) experiments. As predicted, fitted $K_{\mathrm{M}}$ values for chambers with >5 two-point fits showed systematic increases. To ensure accurate $K_{\mathrm{M}}$ estimates, we therefore flagged all chambers in which >5 of the 7 initial rates (corresponding to each [cMUP]) were determined from two-point fits and eliminated them from downstream analysis.

##### Assessing measurement quality using "fiducial activity measurements" and local Lower Limit of Detection (LLoD)

As the kinetic parameters of enzymes within the mutant library span a wide dynamic range, even very small amounts of contaminating "fast" enzymes within chambers containing "slow" enzymes could yield artificially high observed rates. To detect any such contamination, each printed array included "fiducial activity" spots interspersed throughout containing either no plasmid ("Skipped") or a catalytically-incompetent or compromised mutant (T79G and/or K162A). Initial experiments probing PafA active site mutants included "Skipped" and T79G chambers every other chamber (1248 chambers total). tier 1 and 2 scanning library experiments included either "Skipped" chambers or chambers containing T79G or K162A (mutants with a $> 10^4$-fold decrease in catalysis compared to WT) constructs at every seventh chamber within a channel; this number of fiducial chambers increased in tiers 2 and 3 (as enumerated in **Table S2**). For tier 3, the K162A mutant became measurable; thus, we estimated LLoDs via linear interpolations of the rates for the T79G and "Skipped" chambers as a function of row number for each column on the device at each [cMUP].

For each kinetic assay, we calculated observed rates of hydrolysis for these fiducial chambers and then estimated an observed local background hydrolysis rate (Lower Limit of Detection, LLoD) for all chambers using a linear interpolation of these observed rates along a given channel (column) and the previously measured background rate of non-PafA-mediated hydrolysis (**Fig. S15A**):

$$\text{LLoD} = \frac{v_{\text{i}}^{\text{obs}} - v_{\text{bg}}^{\text{true}}}{v_{\text{bg}}^{\text{obs}} - v_{\text{bg}}^{\text{true}}} \tag{2}$$

where $v_{\text{i}}^{\text{obs}}$ denotes the observed initial rate, $v_{\text{bg}}^{\text{obs}}$ denotes the interpolated background rate from catalytically-inactive or compromised chambers, and $v_{\text{bg}}^{\text{true}}$ denotes the on-chip measured background rate ($-4 \times 10^{-5}$ $\mu$Ms$^{-1}$). In contrast to prior efforts to measure a true background rate (*cf. Lower limit of resolution due to non-PafA-driven hydrolysis*), where we considered only later timepoints to minimize contributions from photobleaching and diffusion across PDMS layers, to determine this number we considered only the early timepoints to accurately mimic observed effects during kinetic assays, yielding the slightly negative observed rate.

To determine the minimum activity threshold above the LLoD that would minimize false positives without unnecessarily culling data, we simulated initial rates as a function of [cMUP] for mutant $k_{\text{cat}}$ values from 0.2–500 s$^{-1}$ and $K_{\text{M}}$ values from 0.05–500 µM in the presence of a variable amount of a "contaminant" enzyme with WT parameters ($1 \times 10^{-5}$ to 0.2 nM, with 0.2 nM representing the total amount of enzyme immobilized under the Button valve). For each simulation, we: (1) divided the simulated rate (for the mutant enzyme plus WT-like contaminant) by the contribution from "background" contaminant only (**Eq. 2**, assuming $v_{\text{bg}}^{\text{true}}$ for the purposes of simulation) to calculate a "fold-change over the limit of detection", (2) fit observed rates to the Michaelis-Menten equation to yield simulated observed $k_{\text{cat}}$ and $K_{\text{M}}$ values, and (3) plot the ratio of the observed $k_{\text{cat}}$ and $K_{\text{M}}$ values to the true value as a function of the fold-change over the limit of detection. This analysis suggests that a fold-change over the limit of detection of $\geq 5$ allows accurate measurement of $k_{\text{cat}}$ and $K_{\text{M}}$ values spanning the range from wild-type PafA to all known active site mutants within 2-fold (**Fig. S15B**). We therefore flagged all chambers in which observed rates were <5-fold above the per-chamber estimated LLoD and removed them from downstream analysis.

#### Aggregating cMUP hydrolysis data across experiments

To aggregate data across experiments and across tiers, identify likely outlier measurements, estimate error on returned parameters, and test for statistically significant differences from WT values, we carried out multiple replicate experiments within each tier to obtain multiple replicates of each mutant. At this stage, we also applied additional quality control filters and normalizations as described below (and see **Figs. S16–S17**). We provide final aggregated estimates for each mutant in the **OSF Repository Data** (https://osf.io/ez9xv/).

##### Aggregating measurements for mutants measured across multiple tiers

Many mutants have rates that cannot be resolved during tier 1 (low [E], fast mutant experiments). For these mutants, rare replicates that pass quality control filters in tier 1 most likely result from cross-chamber contamination, and inclusion of these replicates could skew measured rates.

**Estimating false positive probabilities for mutants with many replicates** To estimate the probability of a slow mutant replicate erroneously passing the LLoD filter, we re-examined the initial active site mutant experiments, in which true rates for all mutants are known and every second chamber is a "fiducial rate" chamber. To quantify the number of slow mutants that might erroneously pass a LLoD filter with fewer "fiducial rate" spots, we: (1) sub-sampled the "fiducial rate" chambers to mimic having such chambers at only every $7^{\text{th}}$ position (as in the tier 1 experiments), (2) calculated an apparent interpolated local LLoD for each channel, and (3) quantified the number of slow mutants that would erroneously pass this local LLoD due to local contamination. These simulations returned a median false positive rate of 11±3% (SD).

Next, we tested whether this predicted false positive rate was consistent with experimental observations in tier 1. In the glycine and valine scanning library measurements, we expect that: (1) most mutants will behave similarly to the WT enzyme, (2) chambers with artificially high rates from cross-chamber contamination will, therefore, most likely be contaminated with WT-like enzyme, and (3) that cross-chamber contamination will disproportionately affect mutants with the slowest kinetic parameters resolved within a given experimental tier. The T79S, N100A, and R164A mutants have significantly tighter $K_{\mathrm{M}}$ values compared to WT, are near the lower dynamic range limit for tier 1 experiments ($k_{\mathrm{cat}}$ values of 1 $\mathrm{s}^{-1}$ for T79S and N100A and 5 $\mathrm{s}^{-1}$ for R164A), and each device contained between 4 and 20 replicates of each mutant (with [E]>0.3 nM). We therefore compared the fitted $K_{\mathrm{M}}$ values for these mutants with their known values to test if the measured rates in these chambers arise from WT-like contaminant or from the mutants themselves. For each mutant, we calculated: (1) the number of chambers passing the tier 1 local LLoD filter, and (2) the Michaelis-Menten parameters for chambers that did and did not pass the tier 1 local LLoD filter. While the R164A mutant $K_{\mathrm{M}}$ values recapitulated the off-chip value for this mutant (establishing that R164A can be accurately measured in tier 1), the majority of T79S and N100A chambers had significant deviations from their expected $K_{\mathrm{M}}$ values (**Fig. S16A**). The majority of these T79S and N100A chambers were successfully culled (**Fig. S16B**); however, ~10% and ~4% of chambers passed quality control and had WT-like $K_{\mathrm{M}}$ values (**Fig. S16A**, red points). This indicates that these are likely false positives and provides a second conservative estimate of the false positive rate of ~10%.

**Estimating false positive measurements in tier 1 as a function of number of replicates** Previous off-chip measurements and high replicate numbers allowed us to estimate the false positive probability for T79S and N100A (~100 replicates, $f_{\text{passed}} = 0.1$). However, most mutants have a smaller number of total replicates (~5) (**Fig. S22C**), so the observed false positive probabilities will likely be higher. To estimate false positive probabilities for smaller replicate numbers, we simulated the number of replicates likely to erroneously pass the LLoD filter as a function of number of replicates for a mutant too slow to be measured in tier 1 using the 10% false positive rate (**Fig. S16C**). Based on these simulations for 5 replicates, we selected a conservative cutoff of $f_{\text{passed}} = 0.4$ for tiers 1 and 2. For mutants not meeting this criterion, we incorporated only measurements from tiers 2 and 3. For tier 3, we did not implement this cutoff as all mutants were separated by a "fiducial" chamber in these experiments.

##### Normalizing $k_{\mathrm{cat}}$ and $k_{\mathrm{cat}}/K_{\mathrm{M}}$ measurements across experiments and tiers

**Rationale for cross-experiment normalization** Within and across experimental tiers, we observed systematic variations in measured $k_{\mathrm{cat}}$ values for WT PafA and other active site mutants (**Fig. S17A**). One model for variability in measured $k_{\mathrm{cat}}$ values between experimental replicates is that small and variable amounts of enzyme can remain non-specifically adsorbed to chamber walls even after stringent washing and contribute to hydrolysis. To test for this, we performed the standard expression procedure (including on-chip expression, a 1 h incubation post-expression to allow expressed constructs to bind, and subsequent SDS washing) for a tier 1 library on a device in which we patterned BSA across all device surfaces in the first step (to eliminate neutravidin binding beneath the Button) and additionally flowed bBSA across the device with Button valves open prior to introducing anti-eGFP. This protocol should eliminate any anti-eGFP binding sites underneath the Button, thereby preventing specific recruitment of enzymes to the surface. We then measured cMUP standard curves and initial rates of turnover across all chambers to obtain a full Michaelis-Menten curve and fit observed rates to the following equation:

$$V_{\max}^{\text{obs}} = k_{\text{cat}}^{\text{intrinsic}}([\text{E}]_{\text{self}}^{\text{Button}} + [\text{E}]_{\text{self}}^{\text{wall}}) \tag{3}$$

Returned apparent concentrations of enzymes bound non-specifically to chambers were distributed around a median of 0.38 nM with a long tail extending past 0.1 nM (**Fig. S17B**). However, the amount of non-specifically adsorbed mutant enzyme likely varies within and between devices based on actual flow rates within device channels and the duration of incubation times between expression and washout. We therefore normalized data between experiments prior to aggregation using a linear normalization strategy, as described below.

**Linear normalization procedure** To normalize data across experiments, we rely on measured values for WT PafA and the R164A mutants as follows:

(1) We normalize tier 1 experiments to one another by: (1) calculating the median $k_{\text{cat}}$ for all WT replicates within *each* tier 1 experiment, (2) calculating the median $k_{\text{cat}}$ for *all* WT replicates across all tier 1 experiments, and then (3) scaling each all $k_{\text{cat}}$ measurements within each experiment by the ratio of the WT grand median to the per-experiment median.

(2) tier 2 and 3 experiments typically did not contain WT replicates, and, at the high [E] used, these are extremely prone to two-point initial rate fits (see above). We therefore perform the same normalization for the tier 2 and 3 experiments using the R164A measurements instead of WT.

(3) As R164A mutants are present across all tiers, the tier 1 and 2/3 experiments are normalized using the R164A grand medians in tier 1 and tiers 2 and 3.

Mathematically, this process is expressed as:

$$k_{\text{cat}}^{\text{Norm.}} = k_{\text{cat}}^{\text{obs.}} \left( \frac{\widetilde{k}_{\text{cat}}^{\text{obs.,WT,Tier 1}}}{\widetilde{k}_{\text{cat}}^{\text{WT,expt.i}}} \right) \left( \frac{\widetilde{k}_{\text{cat}}^{\text{R164A,All tiers}}}{\widetilde{k}_{\text{cat}}^{\text{R164A,Tier 1}}} \right) \tag{4}$$

where $k_{\text{cat}}^{\text{obs.}}$ is the fitted $k_{\text{cat}}$ for each individual mutant replicate, $\widetilde{k}_{\text{cat}}^{\text{WT,expt.i}}$ is the median WT $k_{\text{cat}}$ within the same experiment as the mutant, $\widetilde{k}_{\text{cat}}^{\text{WT,Tier 1}}$ is the median WT $k_{\text{cat}}$ across all tier 1 experiments (grand median), $\widetilde{k}_{\text{cat}}^{\text{R164A,All tiers}}$ is the R164A grand median used to normalize across tiers 1 and 2/3, and $k_{\text{cat}}^{\text{Norm.}}$ is the normalized $k_{\text{cat}}$.

For mutant replicates in tier 2 and 3 experiments:

$$k_{\text{cat}}^{\text{Norm.}} = k_{\text{cat}}^{\text{obs.}} \left( \frac{\widetilde{k}_{\text{cat}}^{\text{R164A,Tier 2/3}}}{\widetilde{k}_{\text{cat}}^{\text{R164A.i}}} \right) \left( \frac{\widetilde{k}_{\text{cat}}^{\text{R164A,All tiers}}}{\widetilde{k}_{\text{cat}}^{\text{R164A,Tier 2/3}}} \right) \tag{5}$$

where $\widetilde{k}_{\text{cat}}^{\text{R164A,expt.i}}$ is the median R164A $k_{\text{cat}}$ within the same experiment as the mutant, $\widetilde{k}_{\text{cat}}^{\text{R164A,Tier 2/3}}$ is the median R164A $k_{\text{cat}}$ across all tier 2 and 3 experiments (grand R164A median), and $k_{\text{cat}}^{\text{Norm.}}$ is the normalized $k_{\text{cat}}$. The grand median used to normalize tiers 1 and 2/3 ($\widetilde{k}_{\text{cat}}^{\text{R164A,All tiers}}$) was calculated by taking the median of the R164A tier 1 and tier 2/3 median values.

Normalized $k_{\text{cat}}/K_{\text{M}}$ values were determined using the same normalization factors (*i.e.* replacing $k_{\text{cat}}^{\text{obs}}$ with $k_{\text{cat}}/K_{\text{M}}{}^{\text{obs}}$ in **Eqs. 4** and **5**). Final normalization factors were all within ~2-fold, and this procedure significantly reduced cross-experiment variance compared with unnormalized rates (**Fig. S17**, **C** and **D**).

**Specifying lower limits for very slow mutants**

A total of 24 of the 1036 mutants expressed but were too slow to measure even in tier 3 experiments. For these mutants, we defined an estimated upper limit $k_{\text{cat}}/K_{\text{M}}$ by: (1) assuming that the lowest [cMUP] (typically 10 µM) is below the $K_{\text{M}}$ of the mutant, which is reasonable based on the distribution of $K_{\text{M}}$ values for all mutants (**Fig. S24**), (2) converting the initial rate measured at the lowest [cMUP] into an observed second-order rate constant ($k_{\text{obs}}$), and (3) denoting this as an upper limit.

#### Testing statistical significance of measured effects

The large number of replicate measurements obtained for each mutant allow us to leverage statistical techniques to estimate effect magnitudes, probe the quality of the evidence behind each estimate, and estimate the probability that a mutant's measured kinetic parameters differ from those of the WT enzyme.

After aggregating all replicates for each mutant, we calculated $p$-values under the null hypothesis that the observed median estimate of a mutant parameter did not differ from that of the WT enzyme. For each kinetic parameter, we compared the distribution of measurements for a given mutant with the distribution of measurements for the WT enzyme using a bootstrap hypothesis test (*80*). For each individual mutant, we: (1) pooled the $m$ observations of the mutant distribution with the $n$ WT observations, (2) randomly selected bootstrap samples by sampling from this pool $m$ and $n$ times, with replacement, corresponding to the number of mutant and WT observations, respectively (denoted by WT$^*$ and mutant$^*$), and then (3) calculated the difference between median values for these bootstrap samples ($t^*$, where $t^* = |\widetilde{\mathrm{WT}}^* - \widetilde{\mathrm{mutant}}^*|$). After iterating this procedure multiple times, we computed $p$-values by counting the fraction of times the observed difference between the medians of the two samples was smaller than the difference recorded in the bootstrap procedure:

$$p = \frac{\#(t_{\mathrm{obs}} < t^*)}{N} \tag{6}$$

where $t_{\mathrm{obs}}$ represents the experimentally observed difference between the mutant and WT medians and $N$ represents the number of bootstrap iterations. To enhance computational efficiency, we initially calculated $p$-values using 100 iterations and increased the number of iterations to 1000 and $1 \times 10^5$ only when necessary to resolve small $p$-values for mutants whose parameter estimates were very different from wild-type ($p \leq 0.1$ and $p \leq 0.01$, respectively). For mutants with cMUP $k_{\mathrm{cat}}/K_{\mathrm{M}}$ values resolvable only as upper limits, we calculated $p$-values in a similar manner using upper limit $k_{\mathrm{obs}}$ values; in this case, resulting $p$-values are also upper limits.

We defined $p < 0.01$ as a conservative threshold for declaring a mutational effect (rejecting the null hypothesis) for a given parameter. As we assayed 1036 individual mutants, we expect some number of false positives at this $p$-value threshold. Thus, we implemented a Benjamini-Hochberg false discovery rate (FDR) controlling procedure at a type I error rate of $\alpha = 0.05$ (*81*). This procedure suggested that, generally, mutants with $p > 0.01$ are at a higher than desired probability of being false discoveries (**fig. S21**), which supported our choice to use $p < 0.01$ as a metric for strong evidence of a mutational effect.

#### Probing for equilibrium unfolding, changes in $Zn^{2+}$ affinity, and changes in enzymatic p$K_a$

To test three models that could account for changes in the fraction of active enzyme on the device, we obtained independent on-chip measurements of cMUP as a function of urea concentration, $Zn^{2+}$ concentration, and pH, as described below (see **Supplementary Text S1**).

##### Urea titration

To measure activities in the presence of 0–2 M urea, we expressed enzymes on-chip using the standard protocol described above and then measured enzyme progress curves at 50 µM [cMUP] in the standard reaction buffer (100 mM MOPS, 500 mM NaCl, 100 µM $Zn^{2+}$, pH 8.0) with added [urea] at 0, 1, and 2 M. Measurements were acquired sequentially in order of increasing [urea] and per-experiment quality control was applied as described above.

To determine the urea dependence of each mutant, we scaled the rates of each individual replicate by the enzyme concentration and then normalized to the [E]-scaled rate at 0 M. For mutants with >1 replicate, we used the mean of all normalized rates in subsequent analyses. We then plotted normalized rates as a function of [urea] and compared them to the theoretical curves expected for $[\mathrm{urea}]_{1/2}$ values of 0 and 1 M (Main Text **Fig. 2H**).

The urea dependency of enzyme activity was measured off-chip by following turn over of the chromogenic phosphate diester methyl-paranitrophenylphosphate (MepNPP) (0.5 mM) by WT PafA (0.1 µM) with the following concentrations of urea added immediately before measuring turnover: 0, 0.1, 0.2, 0.5, 12, and 4M urea. WT PafA was expressed and turnover was measured using a Tecan plate reader as described above. Initial rates of turnover were fit to a competitive model of inhibition, as described below (**Eq. 18**), yielding an apparent $K_i$.

##### Equilibrium $Zn^{2+}$ titration

To measure the [$Zn^{2+}$] dependence of activity, we expressed enzymes on-chip using the standard protocol and then measured rates of hydrolysis for 50 µM [cMUP] under standard buffer conditions in the presence of 100 µM $Zn^{2+}$. We then measured rates of hydrolysis in the presence of 500, 100, 25, 10, and 1 µM [$Zn^{2+}$], in order of descending [$Zn^{2+}$]. To ensure that the [$Zn^{2+}$] was completely exchanged and that we have reached equilibrium at each [$Zn^{2+}$], we carried out 6 sequential assays at each [$Zn^{2+}$] (~6.5 hours total at each [$Zn^{2+}$]). We then calculated mean rates for each mutant at each [$Zn^{2+}$], normalized them to the median rate for each mutant at 100 µM [$Zn^{2+}$], and plotted normalized rates as a function of [$Zn^{2+}$] (Main Text **Fig. 2I**).

##### Measuring enzymatic p$K_a$s

Mutations might shift the enzymatic p$K_a$ from the WT-value of ~8.0 (*16*) to more acidic values, diminishing apparent activity under assays at pH 8. To test for this, we acquired Michaelis-Menten curves at pH 7, 7.5, and 8.5 in addition to the standard pH of 8. To minimize chip-to-chip variability, we acquired all pH data on the same device by acquiring Michaelis-Menten curves at five concentrations (10, 250, 100, 200, and 1000 µM) rather than the standard 7, and also acquired per-chamber cMU standard curves at only three concentrations (2, 10, and 50 µM). Each turnover assay was 2925 s in duration and we acquired Michaelis-Menten curves in the following pH order: 8, 7.5, 7.0, and 8.5. We determined $k_{cat}/K_M$ values for each mutant at each pH and compared these to theoretical curves assuming a 2-state pH-dependent inactivation (competitive inhibition) model with p$K_a$s between 5 and 8 (see **Supplementary Text S1** and **Fig. S27**).

#### Measuring methyl phosphate hydrolysis using HT-MEK

To measure hydrolysis of methyl phosphate (MeP), we developed an on-chip coupled assay in which we detect production of inorganic phosphate ($P_i$) using a commercially available fluorophore-derivatized phosphate binding protein (PBP; Phosphate Sensor, ThermoFisher, catalog no. PV4406) (*18*). Binding of ($P_i$) by PBP leads to an increase in fluorescence that we detected via excitation at 427/10 and collection of emitted light at 470 nm using a custom "PBP" filter set (Semrock Inc., 427/10 bandpass excitation filter, catalog no. FF01-427/10-25; 470/22 bandpass emission filter, catalog no. FF01-470/22-25; 442 nm dichroic beamsplitter, catalog no. DI03-R442-T1-25X36; mounted in a TE2000 filter cube, catalog no. NTE).

#### Measuring enzyme concentration for MeP assays

Small amounts of PBP can non-specifically bind to the chamber surface (including the Button valve) and interfere with eGFP quantification. To avoid this, we measured enzyme concentrations by measuring eGFP intensities before introducing PBP into the device and acquiring the standard curve.

##### On-chip [$P_i$] standard curves

To obtain per-chamber standard curves relating fluorescence of PBP to [$P_i$], we prepared standards containing between 0 and 150 µM $P_i$ (typically 0, 1, 3, 6, 15, 30, 75, and 150 µM) and 30 µM PBP in 10 mM MOPS, 50 mM NaCl, 100 µM $ZnCl_2$, at pH 8.0 and 5 mg/mL BSA (UltraPure, ThermoFisher Scientific, catalog no. AM2616); the BSA functions as a sacrificial protein to minimize non-specific adsorption of PBP to device walls. As binding to PBP weakens with increasing ionic strength, we carried out MeP kinetic assays at a reduced ionic strength of ~60 mM as compared to the ~600 mM used for cMUP kinetic assays. All BSA was exchanged into ultrapure water using an Amicon microcentrifuge concentrator (3K or 10K MWCO, Millipore, catalog no. UFC5003 and UFC5010, respectively) over multiple spin cycles to remove residual $P_i$.

Unlike the linear cMU standard curves, binding of $P_i$ to PBP is defined by a single-site binding isotherm relating median fluorescence within each chamber to $[P_i]$:

$$I([\mathrm{P_i}]) = 0.5A(K_D + [\mathrm{P_i}] + [\mathrm{PS}] - \sqrt{(K_D + [\mathrm{PS}] + [\mathrm{P_i}])^2 - 4[\mathrm{PS}][\mathrm{P_i}]}) + I(0\mu\mathrm{M}) \quad (7)$$

where $I([\mathrm{P_i}])$ is the median fluorescence at a given $[\mathrm{P_i}]$, [PS] is the concentration of PBP, $K_\mathrm{D}$ is the dissociation constant, $A$ is the scaling factor necessary to relate fraction PBP bound to observed fluorescence, and $I(0\mu\mathrm{M}[\mathrm{P_i}])$ is the median fluorescence intensity of PBP in the absence of added $P_i$. To determine chamber-specific fitted parameters required to calculate $[P_i]$ from observed fluorescence, we then performed a nonlinear least-squares fit of observed intensities as a function of $A$, $I(0\mu\mathrm{M}[\mathrm{P_i}])$, [PS], and $K_\mathrm{D}$. Fitting all four parameters effectively provides an interpolation of the standard curve that makes it possible to determine $[P_i]$ from observed fluorescence within the pseudo-linear range of the sensor's response (rather than providing an accurate estimate of $K_\mathrm{D}$).

##### On-chip MeP hydrolysis assays

We performed MeP hydrolysis reactions in the same reaction buffer as the standard curve (10 mM MOPS pH 8.0, 50 mM NaCl, 100 mM $ZnCl_2$, 5 mg/mL BSA, and 30 µM PBP). To equilibrate the device, we flowed reaction solution through all chambers for 6–8 min with the Buttons closed. To start reactions, we opened the Button valves and imaged as described previously (*cf. Measuring cMUP hydrolysis on-chip*) to quantify fluorescence over time. Images were processed as for the cMUP assays with the exception that individual chambers were found using images from the highest $[P_i]$ in the standard curve.

To determine product concentration ($[P_i]$) from the measured fluorescence within each chamber, we substituted the fitted parameters from **Eq. 7** into the inverse of the isotherm function:

$$I^{-1}(I[P_i](t)) = [P_i](t) \quad (8)$$

where $I[\mathrm{P_i}](t)$ and $[\mathrm{P_i}](t)$ are the fluorescence intensity and $[P_i]$ at time $t$, respectively.

##### Measuring initial rates of MeP hydrolysis

To determine initial rates at each [MeP], we used a similar algorithm to the determination of initial rates for cMUP-based assays. However, we limit the fitted range to include only calculated $P_i$ concentrations below two-thirds of the total available [PBP], as the accuracy of inferred $[P_i]$ from **Eq. 8** decreases sharply as PBP approaches saturation.

Rates of MeP hydrolysis were measured for substrate concentrations 0–50 µM. Having substrate concentrations both below and above the [PBP] results in three different regimes under which we need to determine initial rates:

1. [MeP] <[PBP]
2. 0.3 [MeP] >(2/3)[PBP]
3. 0.3 [MeP] <(2/3)[PBP]

In regime 1, where [MeP] is less than [PBP] (*i.e.* where [MeP] < 30 µM) and for which we can follow the complete progress curve without saturating the PBP, we fit either $\leq$ 30% of the total turnover or the first two points in cases where $\geq$ 30% is turned over before the second point is acquired, as described for cMUP assays. In regime 2, we only fit points corresponding to [PBP] <20 µM (the linear regime of the calibration curve). In regime 3, we fit all points in which $[P_i]$ < 0.3 [MeP].

Initial rates were fit to the Michaelis-Menten equation (**Eq. 1**) as for cMUP hydrolysis.

##### Quality control and aggregation of MeP $k_{cat}/K_M$ values across experiments

Quality control of MeP hydrolysis assays was performed identically to that for cMUP assays (*cf. Analysis and quality control of cMUP hydrolysis measurements*). To obtain final estimates of $k_{cat}/K_M$ for MeP hydrolysis, we aggregated data from tier 1 (low [E]) and tiers 2 and 3 (high [E]) experiments (**Table S2**) as for the cMUP assays, with the following changes to the normalization procedure: (1) We calculated normalization factors using $k_{cat}/K_M$ values rather than $k_{cat}$, as the low [MeP] used and high MeP $K_M$ values for most mutants preclude accurate estimation of $k_{cat}$ and $K_M$; (2) We normalized tier 1 experiments to one another based on the WT value in each experiment; (3) We normalized tier 2/3 experiments to one another based on the T79S value; and (4) We did not perform additional cross-Tier normalization as there were no mutants that could be resolved in both tier 1 and tier 2/3 experiments.

##### Determining significance of MeP $k_{cat}/K_M$ effects

To determine the significance of observed changes in MeP $k_{cat}/K_M$ relative to WT, we used the same procedures for bootstrapping probability estimates and defining upper limits as described above for cMUP measurements.

##### Off-chip MeP hydrolysis assays

As for cMUP, we compared rates obtained on-chip for active site mutants hydrolyzing Me-P with rates obtained for the same recombinant enzymes expressed off-chip. Off-chip experiments used the same substrate, $P_i$, and PBP concentrations as on-chip measurements and detected build-up of $P_i$ via PBP fluorescence using a Tecan Infinite M200 plate reader at excitation and emission wavelengths of 415 nm and 470 nm, respectively.

#### Quantifying enzyme activity as a function of temperature and [$Zn^{2+}$] during expression

To measure the dependence of catalytic activity on expression temperature, we carried out on-chip PURExpress reactions at either 23°C or 37°C for 45 min. During the last 10 min of this reaction, we flowed reaction buffer containing 500 µM chloramphenicol. After the 45 min incubation, we closed the Sandwich valves and opened the Necks, and allowed the enzyme to fold for 90 min at 23°C in the presence of the chloramphenicol, thereby halting translation to prevent formation of a mixture of enzymes produced at different temperatures. After the 90 min incubation, we opened the Buttons to bind enzyme to antibody-coated surfaces for 15–60 min and then closed the Buttons and performed the standard flush using normal PafA reaction buffer.

To measure the dependence of catalytic activity on added $ZnCl_2$ during expression, we substituted either 10 or 1000 µM $ZnCl_2$ for the 100 µM $ZnCl_2$ typically added to PURExpress reactions and flowed reaction buffer with the same concentration of $ZnCl_2$ during expression. For the subsequent enzymatic activity assays, we switched to standard reaction buffer containing 100 µM $ZnCl_2$ and carried out all assays as for standard cMUP hydrolysis experiments.

In both cases, we again measured Michaelis-Menten parameters for cMUP hydrolysis under standard on-chip reaction conditions, as described above. For all mutants, we aggregated data across experiments and tiers largely as described previously to yield median estimates of $k_{cat}$, $K_M$, and $k_{cat}/K_M$ at each expression temperature and [$Zn^{2+}$]. However, data aggregation procedures did not cull for mutants based on the percentage of observations exceeding LLoD thresholds due to a more limited number of replicates. In addition, this limited number of replicates precluded determination of significance using bootstrap hypothesis testing. Instead, we estimated significance by comparing the overall distribution of the ratios of the median $k_{cat}/K_M$ values at the two temperatures and $Zn^{2+}$ concentrations, normalized to the median WT PafA $k_{cat}/K_M$ for each experiment, to the distribution of the ratios between experimental replicates of the same mutant under identical conditions (Main

Text **Fig. 3, G and I**). These temperature effect (T-Effect) and zinc-effect (Zn-Effect) ratios are defined below (**Eqs. 9** and **10**, respectively).

$$\text{T-Effect} = \frac{\left[\frac{(k_{cat}/K_M)^{\text{Mutant}}}{(k_{cat}/K_M)^{\text{WT}}}\right]^{23°\text{C}}}{\left[\frac{(k_{cat}/K_M)^{\text{Mutant}}}{(k_{cat}/K_M)^{\text{WT}}}\right]^{37°\text{C}}} \tag{9}$$

$$\text{Zn-Effect} = \frac{\left[\frac{(k_{cat}/K_M)^{\text{mutant}}}{(k_{cat}/K_M)^{\text{WT}}}\right]^{1000\ \mu\text{M Zn}^{2+}}}{\left[\frac{(k_{cat}/K_M)^{\text{mutant}}}{(k_{cat}/K_M)^{\text{WT}}}\right]^{10\ \mu\text{M Zn}^{2+}}} \tag{10}$$

We plotted the top 5th percentile of T-Effects as spheres on the structure of PafA (Fig. 3F and G). The number of mutants for which we measured the Zn-Effect was lower than that in the T-Effect experiments; we therefore used a less stringent threshold (top 20th percentile of Zn-Effects) when plotting effects on the structure (Fig. 3, H and I).

#### Characterization of the misfolded state

##### Further characterization of mutants off-chip *in vitro*

To test if on-chip expression alters enzyme folding, we expressed and assayed a subset of library variants (9 Valine, 8 Glycine, 1 Active Site, and the WT) off-chip *in vitro* (**Table S3**). These mutants spanned more than an order of magnitude in apparent $k_{cat}/K_M$ MeP and included mutants with and without likely fraction active effects (Main Text **Fig. 3C**, closer to the solid blue line and dashed blue line, respectively). In parallel, we expressed and assayed the catalytically inactive Gly library mutant T79G as a control for any background activity stemming from contaminating phosphatase activity in PURExpress.

**Off-chip expression via *in vitro* transcription/translation** We expressed variants in **Table S3** *in vitro* with PURExpress off chip. First, we gently mixed four parts of PURExpress component A with 3 parts of PURExpress component B (a $\frac{10}{7}$X stock) and incubated on ice for 45 min. We then added 40 U/L recombinant ribonuclease inhibitor (0.8 U/µL final), 1 mM $ZnCl_2$ (100 µM final), and nuclease-free water to bring the A and B two-component mixture to 1X. Then, we partitioned 11-µL aliquots of this master mix into PCR tubes and added 0.5 µL of variant plasmid DNA to each. We expressed enzyme in a thermocycler at 37°C for 45 min, followed by a 90 min incubation at room temperature in the dark. To quantify protein yields, we diluted the full reaction volumes with PafA reaction buffer (100 mM MOPS, pH 8, 500 mM NaCl, 100 uM $ZnCl_2$) to 200 µL total, and measured the GFP fluorescence intensity using a Denovix fluorometer/spectrophotometer (model DS-11 FX+). Enzyme concentration was calculated using an eGFP standard curve measured on the same instrument, prepared using commercial eGFP (Biovision, catalog no. 4999).

**pNPP hydrolysis assays** We measured activities of enzymes expressed *in vitro* via turnover assays with the chromogenic substrate pNPP. First, we diluted the IVTT expression mixture for each enzyme an additional 10-fold in water and prepared kinetic assays as follows: we mixed 100 µL of 2X PafA reaction buffer with 50 µL of 4X pNPP substrate at concentrations ranging from 31.2–2000 µM in a 96-well optically-clear plate and then added 50 µL of the diluted IVTT reactions (200-µL reactions). We then measured absorbance at 400 nm over time using a Tecan Infinite M200 plate reader, fit initial rates of turnover to linear models, and fit the Michaelis-Menten equation to linear rates as a function of substrate concentration as described above (**Eq. 1;** using KaleidaGraph software, version 4.5, Synergy Software). Finally, we calculated $k_{cat}/K_M$ for each mutant using the fit enzyme-concentration-independent Michaelis Menten parameters and enzyme concentrations inferred from the eGFP quantifications described above.

**Native polyacrylamide gel electrophoresis of temperature-sensitive mutants** To assess the temperature-dependence of misfolding off-chip, we expressed WT PafA and four temperature-sensitive mutants (Y103G, G130V, T189G, and S190G) with PURExpress for 45 min at either 37°C or 22°C, respectively. Expression was halted by doubling the reaction volumes with 0.5 mM chloramphenicol, and reactions were then incubated at room temperature for 90 min in the dark. This procedure mimicked the protocol employed on-chip for expression at low and high temperature.

Aliquots (two parts) of each sample were mixed with one part 3X native gel loading buffer (62 mM Tris, 0.01% (w/v) bromophenol blue, and 40% (v/v) glycerol at pH 6.8) and loaded onto a polyacrylamide gel (BioRad, Mini-PROTEAN 4–20% TGX gel, catalog no. 4561096) and electrophoresed at 120 V, 4°C, for 3 h in a running buffer containing 25 mM Tris and 190 mM Glycine at pH 8.3. Gels were characterized via in-gel fluorescence imaging (measuring eGFP fluorescence) with a Typhoon FLA 9500 imaging system (excitation at 473 nm with a long pass blue, LPB 510LP, emission filter).

**Thermolysin sensitivity assays of temperature-sensitive mutants** To test if mutants exhibiting reduced activity when expressed at higher temperatures were more susceptible to thermolysin proteolysis, we diluted expression mixtures from WT and five temperature-sensitive mutants (Y52G, Y103G, G130V, T189G, and S190G) into PafA reaction buffer. We then performed native gel electrophoresis with samples of WT and mutant PafA expressed at low and high temperature both with and without treatment with thermolysin protease. Protease-treated samples were prepared as follows: To each sample of IVTT expression product diluted into PafA reaction buffer, we added a 100X stock solution of 10 mg/mL thermolysin (Sigma-Aldrich, catalog no. T7902) containing 10 mM $CaCl_2$ and 2.5 M NaCl to give a final reaction containing 0.05 mg/mL thermolysin (200-fold dilution). Reactions were incubated at room temperature for 10 min and then quenched by adding an equivalent volume of 50 mM sodium EDTA (to inactivate thermolysin via chelation of calcium). Untreated (no thermolysin cleavage) samples consisted of dilutions of the original IVTT mixture in PafA reaction buffer. Then, we loaded both thermolysin treated and untreated samples and electrophoresed in a polyacrylamide gel under non-denaturing conditions, as described above. In-gel fluorescence of eGFP-bearing constructs was quantified with a Typhoon imaging system, also as described previously.

Separately, we performed substrate turnover assays with WT and mutant PafAs both in the presence and absence of thermolysin to correlate observed loss in native-gel band intensity with loss of activity. Expressed enzymes diluted in PafA reaction buffer were further diluted 10-fold in water, and 200-µL reactions were prepared as follows: to 100 µL of 2X PafA reaction buffer, 50 µL of 800 µM pNPP, either 2 µL of water (no thermolysin) or 2 µL of 100X thermolysin stock solution (with thermolysin, final concentration of 0.1 mg/mL thermolysin), and 50 µL of diluted enzyme were added and the reactions were mixed. Reaction progress was monitored via time-dependent changes in absorbance at 400 nm, and no significant curvature was observed over at least 15 minutes (a time frame longer than the thermolysin cleavage reactions complementarily characterized via native gel electrophoresis). Initial rates of hydrolysis were fit as described for other off-chip assays.

##### Further characterization of possible misfolding mutants in *E. coli*

To test if misfolding-induced activity changes also arise *in vivo*, we recombinantly expressed 22 variants in *E. coli*: a subset of the 14 variants characterized *in vitro* off-chip (5 Val, 7 Gly, 1 Active Site, and the WT), an additional 7 variants (4 Val and 3 Gly), and T79G as a control for background activity from contaminating phosphatases *E. coli* cell lysate. After purification, we assayed each for hydrolytic activity towards pNPP (**Table S4**).

**Recombinant *E. coli* expression of enzymes** We expressed PafA-eGFP mutant variants **Table S4** in *E. coli* using leaky expression of the PURExpress vector. Competent BL21 *E. coli* cells (made in house) were transformed with mutant or WT PafA-eGFP constructs (the same plamid stocks use for on- and off-chip expression *in vitro*), plated onto LB agar with 50 mg/mL carbenicillin, and grown overnight at 37°C. For each mutant, 20 mL LB media with carbenicillin was innoculated with a single colony, and cultures were grown overnight at 37°C (high temperature expressions). To express enzyme at room temperature, we used 100 µL of each high-temperature expression overnight culture to inoculate 10 mL LB with carbenicillin, and those cultures were grown overnight at room temperature (room temperature expressions).

The high temperature expression cultures were partitioned into two 10-mL aliquots and pelleted via swinging bucket centrifugation. Plasmid was isolated for one aliquot (each mutant) using a Qiagen miniprep kit and sequence verified with Sanger sequencing; the other aliquot of each high-temperature expression culture was frozen at –20°C for subsequent kinetic assays, described below. The 10-mL room temperature expression cultures were similarly pelleted and frozen for kinetic assays.

Frozen pellets were defrosted and re-pelleted in a swinging bucket centrifuge at 4°C, supernatant was decanted, and the pellets were re-suspended in 1 mL of 10 mM MOPS, 50 mM NaCl, 100 M $ZnCl_2$ at pH 8 (PafA storage buffer). The suspension was then re-pelleted in a bench-top centrifuge and again suspended in 1 mL PafA storage buffer. Cells were lysed via sonication (using a Fisherbrand Model 705 Sonic Dismembrator), on ice, as follows: 5 repetitions of 5 s on, at 25% power, and 30 s rest. The crude lysate was pelleted and the supernatant was isolated and clarified by centrifugation in a bench-top centrifuge for 10 min at 13,000 rpm and 4°C. Enzyme concentration in the clarified supernatant was quantified using eGFP fluorescence as measured using a DeNovix spectrophotometer/fluorometer and using the eGFP standard described previously. The component of total eGFP signal from full-length enzyme-eGFP construct was assessed via in-gel fluorescence (denaturing with sodium dodecyl sulfate, BioRad, Mini-PROTEAN 4–20% TGX polyacrylamide gel) using a Typhoon imaging system, as described previously, and found to be approximately comparable to or greater than the signal arising from free eGFP and truncated enzyme-eGFP. This suggested that eGFP fluorescence-inferred enzyme concentration was comparable with the actual concentration of full-length enzyme construct.

**pNPP hydrolysis assays** The clarified supernatant of each mutant expressed *in vivo* (cell lysate) was further diluted 5-fold into PafA storage buffer, then diluted 10-fold into water. Kinetic assays were then performed as follows: 75 µL of 2X PafA reaction buffer was mixed with 37.5 µM pNPP substrate at concentrations ranging from 31.2–2000 µM and 37.5 µL of diluted cell lysate (150-µL reactions) in a 96-well optically-clear plate. Product turnover measurement, initial rate fitting, and Michaelis-Menten parameter calculation were performed as described for assays of *in vitro* expressed enzymes.

##### Circular dichroism measurements

We over-expressed WT PafA and Y103G constructs with a Strep-tag in *E. coli* at 37°C and room temperature and purified constructs via QuikChange mutagenesis, as previously described in (*16*). Aliquots of enzyme stocks in PafA storage buffer were then buffer-exchanged into both a buffer of 10 mM Tris at pH 8 and a buffer of 10 mM $KP_i$ at pH 8, respectively. Circular dichroism (CD) spectra were obtained for concentrated enzyme stocks in PafA storage buffer, Tris, and potassium phosphate buffers via a J-815 Jasco Spectrophotometer from 210–250 nm (1 nm bandwidth, 50 nm/min scanning speed) at 20°C in a 0.1 cm cuvette (Hellma Analytics). Additionally, WT PafA expressed at room temperature was subjected to incubation with urea for 2 weeks, then characterized by CD to assess loss of secondary structure. This was performed by incubating concentrated protein in reaction buffer with urea on the benchtop, and CD was performed with urea present in $\frac{1}{2}$X reaction buffer, as described (**Fig. S25**).

#### Deconvolving mutational effects on folding and catalysis

Theory and derivation of the expected relationship between the observed MeP and cMUP $k_{\rm cat}/K_{\rm M}$ for catalytic and misfolding effects is detailed in **Supplementary Text S2** and **S3**.

##### Determining statistical significance of fraction active and catalytic effects within the active fraction

We determined the significance of $f_{\mathrm{a}}$ and intrinsic $k_{\mathrm{cat}}/K_{\mathrm{M}}$ effects using a modification of the bootstrap hypothesis testing procedure described above. As determination of the active fraction ($f_{\mathrm{a}}$) depends on measured $k_{\mathrm{cat}}/K_{\mathrm{M}}$ values for both cMUP and MeP hydrolysis, we: (1) pooled the mutant and wild-type MeP $k_{\mathrm{cat}}/K_{\mathrm{M}}$ observations, (2) pooled the mutant and wild-type cMUP $k_{\mathrm{cat}}/K_{\mathrm{M}}$ observations separately, and then (3) determined bootstrap samples of MeP and cMUP measurements by resampling with replacement, and then calculated $f_{\mathrm{a}}$ and intrinsic MeP and cMUP $k_{\mathrm{cat}}/K_{\mathrm{M}}$ values using **Eqs. 31–35** and the number of MeP ($m$) and cMUP ($n$) replicates for the mutant and WT, respectively. We then calculated the difference between the medians of $f_{\mathrm{a}}$ and $k_{\mathrm{cat}}/K_{\mathrm{M}}^{\mathrm{chem}}$ for the mutant and WT bootstrap samples ($t^* = |m - n|$), iterated this procedure 100–1000 times, and computed a $p$-value using **Eq. 6**.

As $k_{\mathrm{cat}}/K_{\mathrm{M}}^{\mathrm{chem}}$ depend on both cMUP and MeP $k_{\mathrm{cat}}/K_{\mathrm{M}}$, and its dependence on these two observed rate constants is non-linear, their errors are typically larger than those on the observed cMUP and MeP $k_{\mathrm{cat}}/K_{\mathrm{M}}$ measurements. We therefore use $p < 0.05$ to determine $k_{\mathrm{cat}}/K_{\mathrm{M}}^{\mathrm{chem}}$ effects, with the caveat that we expect more false positives. The relationship between $f_{\mathrm{a}}$ and cMUP and MeP $k_{\mathrm{cat}}/K_{\mathrm{M}}$ is more linear, so we use our standard $p < 0.01$ to determine $f_{\mathrm{a}}$ effects.

#### Measuring hydrolysis of the non-cognate substrate MecMUP

We assayed rates of MecMUP hydrolysis using a procedure similar to that used for cMUP hydrolysis, again acquiring cMU standard curves at the start of each experiment. However, we modified the pipeline for obtaining initial rates of MecMUP hydrolysis in two ways. First, the non-cognate MecMUP hydrolysis reaction catalyzed by WT PafA is $10^5$-fold slower than cMUP hydrolysis. Therefore, to maximize substrate turnover within a reasonable acquisition time, we measured all rates at high [E] (10–30 nM) and high [S] (0.5–2 mM) over 1–3 hr/assay. Second, trace amounts of the monoester (cMUP) within MecMUP stocks (~0.1% mol/mol) led to an initial burst phase (corresponding to fast hydrolysis of trace cMUP monoester) followed by a longer linear phase (corresponding to slower hydrolysis of MecMUP diester). We therefore fit fluorescent product buildup to a function including a linear term for MecMUP hydrolysis and the integrated form of the Michaelis-Menten equation (*82*) for trace cMUP hydrolysis:

$$[\mathrm{cMU}](t) = v_{\mathrm{i}}^{\mathrm{MecMUP}} t + [\mathrm{cMUP}]_0 - K_{\mathrm{M}} W\left(\frac{[\mathrm{cMUP}]_0}{K_{\mathrm{M}}} \mathrm{e}^{\frac{[\mathrm{cMUP}]_0 - k_{\mathrm{cat}}[E]t}{K_{\mathrm{M}}}}\right) \tag{11}$$

Here, $[\mathrm{cMU}](t)$ is the concentration of fluorescent product at time $t$, $v_{\mathrm{i}}^{\mathrm{MecMUP}}$ is the initial rate of MecMUP hydrolysis, $k_{\mathrm{cat}}^{\mathrm{cMUP}}$ and $K_{\mathrm{M}}^{\mathrm{cMUP}}$ are the Michaelis-Menten parameters of cMUP hydrolysis, [E] is the enzyme concentration, $[\mathrm{cMUP}]_0$ is the contaminating concentration of cMUP in the assay present during the first fluorescence measurement, and $W$ is the Lambert Omega function (*82*). To reduce the number of free parameters, we substituted measured $k_{\mathrm{cat}}^{\mathrm{cMUP}}$ and $K_{\mathrm{M}}^{\mathrm{cMUP}}$ for cMUP hydrolysis for each mutant and restrained $[\mathrm{cMUP}]_0$ to be $\leq 0.1\%$ of [MecMUP]. We then determined $v_{\mathrm{i}}^{\mathrm{MecMUP}}$ for each mutant by non-linear least squares fitting of the entire progress curve.

To measure $k_{\mathrm{cat}}/K_{\mathrm{M}}$ for MecMUP hydrolysis for nearly all mutants, we measured initial rates at 3–4 MecMUP concentrations from 500–2000 µM and fit these to the Michaelis-Menten equation. For the tier 3 library, we measured rates by quantifying MecMUP hydrolysis at a single high concentration (2000 µM) over 22 hours. For this experiment, we determined $k_{\mathrm{cat}}/K_{\mathrm{M}}$ by simply dividing the fitted rate by the [E] and [MecMUP] concentrations as [MecMUP] was not appreciably depleted over the total measurement time (with the assumption of being subsaturating).

##### Quality control of MecMUP hydrolysis measurements

Quality control of MecMUP hydrolysis assays was carried out in a similar manner to cMUP assays with two differences: (1) the local LLoD interpolations considered only T79G and Skipped chambers and not K162A (as K162A has a WT-like rate of diester hydrolysis), and (2) we culled any chambers with measured MecMUP rates for a given mutant within 5-fold of the cMUP rate during the last 5 timepoints of the reaction, reasoning that these rates likely represent hydrolysis of trace monoester contaminant.

##### Obtaining final estimates of MecMUP hydrolysis and determining statistical significance

After quality control, MecMUP assays were aggregated and normalized in an analogous manner to the cMUP and MeP measurements, but with the following differences: (1) K162A was used as the normalizing mutant instead of WT, as its MecMUP $k_{\mathrm{cat}}/K_{\mathrm{M}}$ is WT-like, and every experiment contained multiple replicates. (2) Instead of a two-stage normalization procedure, K162A was used to normalize all experiments across all tiers. As MecMUP hydrolysis is intrinsically orders of magnitude slower than cMUP and MeP hydrolysis, there were no 2-pt fits.

To determine the statistical significance of MecMUP $k_{\mathrm{cat}}/K_{\mathrm{M}}$ effects, we again used bootstrap hypothesis testing to compare mutant $k_{\mathrm{cat}}/K_{\mathrm{M}}$ distributions relative to that of wild-type PafA, as described for cMUP and MeP assays above.

#### Functional Component 1: Effects through the O2 phosphoryl oxygen atom

We quantified Functional component 1 (FC1) for each mutant by comparing observed fold-changes in $k_{\mathrm{cat}}/K_{\mathrm{M}}$ from the WT value for MecMUP versus MeP hydrolysis:

$$\mathrm{FC1} = \Delta\mathrm{MecMUP}/\Delta\mathrm{MeP} \tag{12}$$

where

$$\Delta = \frac{k_{\mathrm{cat}}/K_{\mathrm{M}}^{\mathrm{mutant}}}{k_{\mathrm{cat}}/K_{\mathrm{M}}^{\mathrm{WT}}} \tag{13}$$

for MeP and MecMUP hydrolysis. As noted in the main text, as misfolding is expected to affect both MeP and MecMUP equally, we compared the observed $k_{\mathrm{cat}}/K_{\mathrm{M}}$ values rather than the values corrected for $f_{\mathrm{a}}$ effects.

##### FC1 values for mutants below the dynamic range of on-chip assays

There are 3 instances in which upper limits in MeP and MecMUP result in limits on calculated FC1 values:

(1) MecMUP $k_{\mathrm{cat}}/K_{\mathrm{M}}$ upper limit but MeP $k_{\mathrm{cat}}/K_{\mathrm{M}}$ measured. In this case, rates of MecMUP hydrolysis are much slower than cMUP and MeP, limiting assay dynamic range. The calculated value therefore represents an upper limit; however, as these mutants could have FC1 values anywhere between 0 and this upper limit, we do not consider them significant in downstream analyses.

(2) MecMUP $k_{\mathrm{cat}}/K_{\mathrm{M}}$ measured, but MeP $k_{\mathrm{cat}}/K_{\mathrm{M}}$ upper limit, resulting in lower FC1 limits. We conservatively estimate the significance of these mutants using this lower limit value, as true FC1 values must actually be larger and more significant.

(3) Both MecMUP and MeP $k_{\mathrm{cat}}/K_{\mathrm{M}}$ upper limits, resulting in indeterminate FC1 values and significance.

##### Determining statistical significance of FC1 effects

We determined the statistical significance of FC1 effects using bootstrap hypothesis testing, as follows. For each mutant, (1) we merged the observations of mutant MeP and MecMUP $k_{\mathrm{cat}}/K_{\mathrm{M}}$ with the WT observations, (2) randomly sampled from these observations to generate bootstrapped observations with the same number of mutant observations in both assays ($\mathrm{MecMUP}^{*}_{\mathrm{mutant}}$ and $\mathrm{MeP}^{*}_{\mathrm{mutant}}$), (3) similarly generated bootstrapped observations with the same number of WT replicates in both assays ($\mathrm{MecMUP}^{*}_{\mathrm{WT}}$ and $\mathrm{MeP}^{*}_{\mathrm{WT}}$), (4) calculated the medians of these bootstrap observations, and (5) calculated the differences ($t^{*}$) between the MecMUP and MeP $k_{\mathrm{cat}}/K_{\mathrm{M}}$ medians for the mutant and WT from the bootstrap observations:

$$\delta\text{mutant} = \log_2 \widetilde{\text{MecMUP}}^*_{\text{mutant}} - \log_2 \widetilde{\text{MeP}}^*_{\text{mutant}} \tag{14}$$

$$\delta\text{WT} = \log_2 \widetilde{\text{MecMUP}}^*_{\text{WT}} - \log_2 \widetilde{\text{MeP}}^*_{\text{WT}} \tag{15}$$

$$t^* = \mid \delta\text{mutant} - \delta\text{WT} \mid \tag{16}$$

We then repeated this process up to 500 times and calculated the $p$-value as the number of times the bootstrapped difference was greater than the difference between the medians of the experimentally-observed distributions ($t^{\text{obs}}$).

$$p = \frac{\#(t_{\text{obs}} < t^*)}{N} \tag{17}$$

#### Determination of residues within each K162/R164 interaction shell

To define residues within K162 and R164 interaction shells, we used GetContacts (https://getcontacts.github.io/) as follows. First, we ran GetContacts using the WT PafA crystal structure (PDBID: 5TJ3) as input to identify all the potential contacts between residues in PafA. We defined the active site to consist of all residues making direct contacts to the substrate (T79, N100, K162 and R164) as well as the six catalytic $Zn^{2+}$-liganding residues. We then defined the second shell as the set of residues having at least one contact to either K162 or R164 (but excluding the active site residues themselves).

To define the third shell, we then determined the set of residues making at least one contact to any of the second shell residues, including contacts between residues via bridging waters. As some of the second shell residues interact with one another, we needed to remove these from the third shell set. We therefore determined the subset of the third shell residues not present in either the second shell or the active site set. Higher order shells were also determined by enumerating the full set of contacts and then finding the unique subset of residues not present in any lower interaction shells.

In addition to the K162 and R164 interaction shells, we also defined the sets of residues within each interaction shell of any of the active site residues (T79, N100, K162 and R164 and the catalytic $Zn^{2+}$-liganding residues) in an analogous manner.

#### Determination of surface-exposed residues

Surface-exposed residues were determined by calculating relative accessible surface area (ASA) using Biopython (Bio.PDB.DSSP) (*83*) and the normalization values of Tien *et al.* (*84*). Residues having a relative ASA >0.16 were defined as surface-exposed (*85*). The N- and C-terminal residues unmodeled in the PafA crystal structure were also assumed to be solvent exposed.

#### Measuring phosphate inhibition constants using HT-MEK

We determined inhibition constants $K_{\text{i}}$ for inorganic phosphate ($P_{\text{i}}$) via competitive inhibition assays in which we varied inhibitor concentrations at a constant [cMUP]. We performed tier 1 and tier 2 assays for the full valine and glycine-scanning libraries at either 10 or 50 µM cMUP in 100 mM MOPS (pH 8.0) with 500 mM NaCl and 100 µM $ZnCl_2$ at 10–12 inhibitor concentrations ranging from 0–8000 µM. For tier 3 assays, we reduced the number of inhibitor concentrations to 6 due to the longer acquisition times required for each measurement. When possible, we included replicate assays (typically at 0 µM $P_{\text{i}}$) to control for activity loss over time. On-chip assays were otherwise acquired as previously described for cMUP.

For each chamber, we quantified initial rates of product formation at each inhibitor concentration as described previously. We then determined inhibition constants for each chamber by fitting the observed rates to a model of competitive inhibition:

$$v_i([I]) = \frac{v_i^{\text{uninhibited}}}{1 + [I]/K_i} \tag{18}$$

where $v_i$ is the initial rate at a given inhibitor concentration, $K_i$ is a fitted inhibition constant, and $v_i^{\text{uninhibited}}$ is a fitted initial rate in the absence of inhibitor.

As for the other measurements, all data, fits and quality control flags for each chamber within each experiment are in the per-experiment CSV and PDF files in the **OSF Repository Data**.

**Obtaining final estimates of phosphate inhibition constants and determining significance**

To obtain final estimates of phosphate inhibition constants for each mutant, we performed quality control and aggregated data across experiments in a similar manner as for cMUP, MeP, and MecMUP experiments. To pass all quality control filters, we required: (1) that chambers contain [E] >0.3 nM, (2) that eGFP spots be free of apparent defects or artifacts upon visual inspection, (3) that initial rates were >5-fold above the LLoD (calculated using the 0 µM [$P_i$] assay), and (4) that inhibition curves included <6 2-pt fits. In addition, we only considered tier 1 and 2 measurements for mutants in which >40% of replicates passed the LLoD filter during that tier.

We corrected for any systematic errors in either the stock phosphate concentrations or buffer pH between experiments by normalizing fitted $K_i$ values to the median $K_i$ of the 3–17 WT replicates within each experiment that passed quality control. As the R164A $K_i$ is a lower limit, we could not apply a two-stage normalization analogous to that for cMUP; therefore, the $K_i$ values from tier 2 and 3 experiments lacking WT replicates were not normalized. The per-experiment median WT $K_i$s (and thus normalization factors) were all within ~2-fold of the cross-experiment median.

Apparent competitive inhibition constants (${K_i}^{\text{obs}}$) are sensitive to substrate $K_M$, with the magnitude of the delta between ${K_i}^{\text{obs}}(P_i)$ and ${K_i}^{\text{intrinsic}}(P_i)$ related to $K_M$ of cMUP for our mutants. In traditional competitive inhibition assays, the substrate concentration is chosen to be $[S] < K_M$, such that ${K_i}^{\text{obs}} \approx {K_i}^{\text{intrinsic}}$. Here, we used 10 and 50 µM cMUP concentrations to meet this criterion for the majority of mutants. In addition, to correct for systematic overestimation of measured (observed) $K_i(P_i)$ values arising for mutants with low cMUP $K_M$ values, we used the Cheng-Prusoff relationship, which relates the intrinsic competitive $K_i$ to the observed competitive $K_i$ and substrate $K_M$:

$$K_i = \frac{K_i^{\text{obs}}}{1 + [S]/K_M} \tag{19}$$

We corrected each individual measurement using the median cMUP $K_M$ for that mutant, the experimental [cMUP] concentration used, and **Eq. 19**. After correction, we combined all $K_i$ measurements and calculated median estimates for each mutant. Raw fitted $K_i$ values are reported in the per-experiment CSV and PDF summaries, and Cheng-Prusoff corrected values are given in the aggregated estimate summary CSV and PDF files.

**Identifying lower and upper $K_i$ limits** We identified 3 mutants for which $K_i$ values were lower limits and 6 for which $K_i$ values were upper limits. Lower $K_i$ limits arise when $K_i \gg$ the highest [$P_i$] assayed. During the aggregation process, we flagged individual measurements as lower limits if the observed fit $K_i$ was >2-fold above the highest [$P_i$] used in the experiment and set a conservative lower limit of the $K_i$ to be 2-fold above the highest [$P_i$]. In addition, we defined the median $K_i$ estimate for a given mutant to an upper limit if >50% of the underlying measurements for that mutant were $K_i$ lower limits.

Upper limits in $K_i$ can arise when: (1) the fit $K_i$ is below the lowest non-zero [$P_i$] concentration used (1 µM in on-chip assays), or (2) when measured cMUP $K_M$ values are upper limits, propagating a limit through (preventing accurate correction via) the Cheng-Prusoff relationship described above. In the first case, as for the lower limits, we defined the median $K_i$ value as an upper limit if >50% of the individual replicates were $K_i$ upper limits and set the $K_i$ of these replicates to be 2-fold less than the lowest [$P_i$]. In the second case, if the median cMUP $K_M$ was a lower limit, we automatically denoted the median $K_i$ value as an upper limit, as this will introduce systematic error on all $K_i$ replicates.

**Determining significance of $K_i$ measurements** We again determined the significance of mutational effect estimates using bootstrap hypothesis testing comparing the distribution of corrected $K_i$ values to the WT corrected $K_i$ distribution (as described above for cMUP measurements). We conservatively estimated the statistical significance of upper and lower $K_i$ limits in the same way, as limit values will only underestimate the true significance. For mutants with catalytic rates below the lower limit of detection, we cannot determine a significant change in $K_i$ of phosphate binding and, therefore, do not report either $K_i$ or significance.

#### PafA phylogenetic analysis

##### Generating a multiple sequence alignment of public database and metagenomic PafA-like AP superfamily member sequences

**Obtaining, aligning, and filtering publicly-available sequences** To assess conservation within metagenomic sequences, we first generated a heavily curated alignment of PafA-like sequences originating in public databases.

To identify candidate sequences, we first assembled a comprehensive collection of 156,929 alkaline phosphatase (AP) superfamily members from the UniProtKB database (*86*). Next, we added the following additional AP superfamily sequences from publicly available databases: (1) sequences that possess the appropriate AP fold, as defined by the CATH (*87*) database (the 3.40.720.10 domain), (2) sequences with sequence patterns manually selected from the InterPro (*88*) and Pfam (*89*) databases that are associated with AP functions (IPR000917, IPR001952, IPR002591, IPR004245, IPR006124, IPR007312, IPR010869, IPR013973, IPR017849, IPR017850, IPR026263, IPR029879, IPR029881, IPR029885, IPR029889, IPR029890, IPR029895, IPR029896, IPR029897, IPR032506, PF00245, PF00884, PF01663, PF01676, PF02995, PF04185, PF07394, PF08665, PF14707, PF16347), and (3) sequences from the UniProtKB/SwissProt, BRENDA (www.brenda-enzymes.org) (*90*) and Sabio-RK (*91*) databases with any of the following EC numbers (2.7.8.20, 2.7.8.21, 2.7.8.43, 2.7.8.44, 3.1.3.1, 3.1.3.2, 3.1.3.27, 3.1.3.39, 3.1.3.54, 3.1.4.1, 3.1.4.3, 3.1.4.12, 3.1.4.38, 3.1.4.39, 3.1.6.1, 3.1.6.12, 3.1.6.13, 3.1.6.14, 3.1.6.2, 3.1.6.4, 3.1.6.4, 3.1.6.6, 3.1.6.8, 3.1.7.6, 3.10.1.1, 3.11.1.2, 3.6.1.29, 3.6.1.9, 5.4.2.1, 5.4.2.12, 5.4.2.7).

Next, we used the Structure Function Linkage Database (SFLD) (*92*) to generate a sequence similarity network (SSN) of the entire superfamily, and used a previously described procedure (*93–95*) to delineate the relevant SSN cluster that includes PafA. Briefly, by mapping experimentally validated reactions to the SSN, we evaluated different edge inclusion cutoffs to identify a threshold that: (1) maintains PafA homologs in one cluster, and (2) separates this PafA cluster from other clusters that include sequences that are clearly not PafA-like (*e.g.* annotated in SwissProt to catalyze a different reaction). This process yielded a list of 1,384 PafA homologs used for subsequent analysis.

Next, we aligned a subset of these initial 1384 PafA-like sequences. First, we clustered sequences by identity using CD-HIT (*96*) and then extracted unique representatives from 75% identity clusters for subsequent analysis. Next, we aligned these representatives, including WT PafA, with PASTA (https://github.com/smirarab/pasta) (*97*) using the following non-default parameters: "--num-cpus=6, -datatype=Protein".

The resulting alignment was restricted to sequences bearing a subset of possible residue identities at positions corresponding to the nucleophile and $Zn^{2+}$ ligands in the PafA reference sequence (Uniprot Accession Q9KJX5). This gappy restricted alignment was then trimmed by removing alignment positions at which gaps existed at $\geq$ 95% of sequences, and both PafA or the other structurally-characterized relative SPAP (Uniprot Accession A1YYW7) were gapped. The resulting alignment was then de-gapped and re-aligned using PASTA (using the same non-default parameters) to yield a final curated "public database alignment" of 478 sequences.

**Obtaining, aligning, and filtering metagenomic sequences** To identify metagenomic PafA-like sequences, we queried the IMG/M database (August 2018) (*98*) with a profile Hidden Markov Model (HMM) generated from the multiple sequence alignment of 478 highly-curated PafA-like sequences collected from public databases ("public database alignment") with hmmbuild (of HMMR, www.hmmer.org) using default arguments. The query was performed with National Energy Research Scientific Computing Center (NERSC) (*99*) resources and enabled by the JGI-EMSL Facilities Integrating Collaborations for User Science (FICUS) program (https://jgi.doe.gov/user-programs/program-info/ficus-overview/). This query returned 2,374,441 sequences that were subsequently restricted to 34,654 sequences meeting the following criteria: 250–1200 residues in length, an HMM bitscore of at least 120, and bearing a Met, Val, or Leu N-terminal residue.

Filtered sequences were concatenated with the public database sequences originally collected (giving rise to the "public database alignment") and aligned with UPP (*100*). Creating this "large alignment" first required a smaller "backbone alignment," which we generated from representative metagenomic sequences identified via CD-HIT aligned initially using PASTA, then heuristically subdivided into sub-alignments and re-aligned and merged using MAFFT (*101*). Finally, we removed sequences in the crude backbone alignment contributing to excessive gappiness via a heuristic-based procedure to generate the final backbone alignment composed of 1402 sequences. Then, we generated a Maximum Likelihood (ML) tree using this "backbone alignment" using FastTree (*102*) with default parameters. The "large alignment" was generated using UPP with the following non-default parameters: "-t $BBTree -d $NumCores -m amino", where $BBTree refers to the tree inferred from the "backbone alignment," and $NumCores was the number of logical cores employed in the calculation. This yielded an alignment of 36,036 sequences with many gapped positions.

The "large alignment" was subsequently restricted to non-gapped positions in the reference PafA sequence (Uniprot Accession Q9KJX5) to yield a "de-gapped large alignment". This masked alignment was further filtered on the presence of the canonical PafA active site residues (T79, N100, K162, and R164) and removing sequences for which PafA non-gapped positions consisted of $\geq$20% gapped identities. This procedure yielded a sieved subset of 14,505 aligned metagenomic and public database sequences (the "culled trimmed metagenomic alignment") used for downstream analyses.

We generated a ML tree of culled PafA-like metagenomic and public database ("culled trimmed metagenomic alignment") sequences using FastTree as described above. The resulting un-rooted tree was plotted using the ggtree package in R (**Fig. S62**) (*103*).

##### Calculating phylogenetic conservation

To test if on-chip measured functional parameters were correlated with conservation, we used the "culled trimmed metagenomic alignment" described above to quantify information content within the alignment at each position within the PafA ORF. We calculated information content using the information_content function in the Bio.Align.AlignInfo module of Biopython (*83*) using a custom amino-acid frequency table derived from the amino acid composition of UniProtKBSwiss-Prot (https://web.expasy.org/protscale/pscale/A.A.Swiss-Prot.html) and ignoring the following characters: "-", "X", "B", "Z", "J", "U", "O".

### Supplementary Text

#### S1. Equilibrium measurements to test if activity depends on [urea], [$Zn^{2+}$], and pH

Altered catalytic efficiencies for a given mutant could simply reflect changes in the equilibrium population of active enzyme under our assay conditions. We identified three molecular mechanisms that could alter the fraction of active enzyme at equilibrium: (1) thermodynamic destabilization that shifts the equilibrium distribution of folded and unfolded PafA, (2) diminished affinity for $Zn^{2+}$ at the bimetallo site, and (3) alteration in the observed inhibitory enzymatic p$K_a$. We systematically tested each of these three possibilities on-chip for our valine and glycine scanning mutant libraries as described below.

##### Equilibrium unfolding of PafA upon mutation in the presence of urea

For a two-state equilibrium between folded and unfolded states given by

$$\mathrm{E_f} \rightleftharpoons \mathrm{E_u} \tag{20}$$

the dependence of the Gibbs free energy of unfolding on denaturant concentration is given by

$$\Delta G = \Delta G^{\mathrm{H_2O}} - m[\mathrm{urea}] \tag{21}$$

where $m$ describes the dependence of ΔG on the concentration of denaturant. Using the lower bound, and therefore most conservative, $m$-value for urea denaturation for wild-type PafA predicted using the Protein $m$-value Calculation Webserver (*104*), ($m_{\mathrm{pred}}$ = 4.8 kcal/mol/M), we can predict the unfolding of PafA as a function of [urea] by recasting **Eq. 21** as follows and solving for the folded fraction, $f_{\mathrm{f}}$:

$$-RT\ln\frac{1-f_{\mathrm{f}}([\mathrm{urea}])}{f_{\mathrm{f}}(\mathrm{urea})} = -RT\ln\frac{1-f_{\mathrm{f}}(0\mathrm{M})}{f_{\mathrm{f}}(0\mathrm{M})} - m[\mathrm{urea}] \tag{22}$$

$$f_{\mathrm{f}}([\mathrm{urea}]) = \frac{1}{1+e^{\frac{m[\mathrm{urea}]}{RT}}(-1+\frac{1}{f_{\mathrm{f}}(0\mathrm{M})})} \tag{23}$$

where $T$ is the temperature and $R$ is the ideal gas constant. In order for an observed significant decrease in activity (*i.e.*, a >2-fold decrease in $k_{\mathrm{cat}}/K_{\mathrm{M}}$) upon mutation to result purely from unfolding, the unfolded fraction (denoted here as $f_{\mathrm{u}} = 1 - f_{\mathrm{f}}$) would have to be >0.5. Under these conditions, **Eq. 23** predicts that the relative populations within the folded and unfolded fractions (and thus the observed activity) would be highly sensitive to the addition of denaturant (Main Text **Fig. 2H**). For example, if 50% of enzyme were unfolded due to thermodynamic destabilization under our standard assay conditions, **Eq. 23** predicts that observed rates at 1 and 2 M would drop by $>10^3$ and $>10^7$-fold, respectively. Although these predicted changes are based on a computed $m$-value, even if this were underestimated by 2-fold, we would still expect a >10 and $>10^3$-fold change at 1 and 2 M urea, respectively. In contrast to these predictions, we observe effects of <4-fold and <10-fold for 99% of mutants, and <2-fold and <3-fold for 95% of mutants, at 1 and 2 M urea, respectively. These small effects are consistent with an apparent competitive inhibition of folded active PafA by urea (**Fig. S26**). Overall, this discrepancy between predicted and observed behaviors establishes that mutational effects on $k_{\mathrm{cat}}/K_{\mathrm{M}}$ cannot be due to equilibrium destabilization.

##### Altered affinity of $Zn^{2+}$ for the bimetallo site upon mutation

PafA binds two essential catalytic $Zn^{2+}$ ions in the folded state (*16*) and upon unfolding, these $Zn^{2+}$ ions should be released to solution. Therefore, a necessary corollary to the model of equilibrium unfolding is that excess [$Zn^{2+}$] will push the following equilibrium towards the folded state:

$$\mathrm{E_f} \rightleftharpoons \mathrm{E_u} + 2\mathrm{Zn}^{2+} \tag{24}$$

where the equilibrium constant, $K_\mathrm{u}$, is defined by

$$K_\mathrm{u} = \frac{[\mathrm{E_u}][\mathrm{Zn}^{2+}]^2}{[\mathrm{E_f}]} \tag{25}$$

or equivalently, in terms of the folded fraction of enzyme, $f_\mathrm{f}$,

$$K_\mathrm{u} = \frac{(1 - f_\mathrm{f})[\mathrm{Zn}^{2+}]^2}{f_\mathrm{f}} \tag{26}$$

If observed mutational effects on activity were due to equilibrium unfolding, decreasing the solution $[\mathrm{Zn}^{2+}]$ would significantly reduce $f_\mathrm{f}$. Rearranging **Eq. 26** yields the relationship between $f_\mathrm{f}$ and $[\mathrm{Zn}^{2+}]$:

$$f_\mathrm{f} = \frac{[\mathrm{Zn}^{2+}]^2}{K_\mathrm{u} + [\mathrm{Zn}^{2+}]^2} \tag{27}$$

To test this model, we measured rates of hydrolysis for PafA valine scan mutants over a range of $[\mathrm{Zn}^{2+}]$ and compared these measured rates to those what would be predicted by **Eq. 27**. Consistent with the absence of significant equilibrium unfolding, we observed no significant decreases in activity over a 100-fold range of $[\mathrm{Zn}^{2+}]$. As fractional occupancies have not been measured for the two distal $\mathrm{Zn}^{2+}$ ions seen in the WT PafA crystal structure, we do not include them in this calculation. However, their inclusion would predict even more dramatic effects.

##### Altered enzymatic p$K_\mathrm{a}$ upon mutation

PafA possesses an inhibitory enzymatic p$K_\mathrm{a}$ of ~8.0 (*16*, *105*). This may be due to deprotonation of a bimetallo zinc-coordinated water, generating an inhibitory hydroxide ion bound at the active site (*19*, *31*). We therefore tested the possibility that mutants with observed decreases in $k_\mathrm{cat}/K_\mathrm{M}$ relative to WT possess shifted enzymatic p$K_\mathrm{a}$s (apparently competitive; **Eq. 28**).

$$\mathrm{E_{active}} \xrightleftharpoons{\mathrm{p}K_\mathrm{a}^\mathrm{inhib} \approx 8.0} \mathrm{E_{inhibited}} \tag{28}$$

This two-state model predicts that a change in assay pH (a change from pH A to pH B) will change the observed $k_\mathrm{cat}/K_\mathrm{M}$ (**Eq. 29**).

$$\frac{(k_\mathrm{cat}/K_\mathrm{M})^\mathrm{pHA}}{(k_\mathrm{cat}/K_\mathrm{M})^\mathrm{pHB}} = \frac{1 + 10^{\mathrm{pHB}-\mathrm{p}K_\mathrm{a}}}{1 + 10^{\mathrm{pHA}-\mathrm{p}K_\mathrm{a}}} \tag{29}$$

Comparisons between experimentally measured $k_\mathrm{cat}/K_\mathrm{M}$ values and those predicted by the previously observed p$K_\mathrm{a}$ (**Eq. 29**) for WT and active site mutant PafA constructs at pH 8.0 (standard conditions), 7.5, and 7.0 revealed that, in contrast with the expected 1.5-fold and 1.8-fold increases in $k_\mathrm{cat}/K_\mathrm{M}$ upon reducing pH to 7.5 and 7.0, the WT enzyme either lost or only slightly gained in activity. These observations are consistent with time-dependent enzyme death that may be aggravated upon acidification. Overall, 97% of glycine and valine library mutants had pH-dependent changes in activity within the range observed for WT PafA enzymes across four pH values. These results strongly suggest that enzymatic p$K_\mathrm{a}$ values for mutants do not significantly differ from the WT values and cannot explain observed mutational effects.

#### S2. Deriving the predicted relationship between $k_{cat}/K_M$ for cMUP and MeP for PafA

At subsaturating concentrations, WT PafA hydrolysis of cMUP is limited by the rate of binding to the enzyme ($k_1$; Main Text **Fig. 7A**) (*13*, *19*). By contrast, hydrolysis of MeP at subsaturating concentrations is limited by the chemical step ($k_{\mathrm{chem},1}$; Main Text **Fig. 7A**). The relative intrinsic reactivities of these substrates is proportional to the ratio of their hydrolysis rates on the enzyme (the rate of the chemical step, enzyme-substrate complex hydrolysis, for each) (*13*):

$$f_{\mathrm{r}} = k_{\mathrm{chem},1}^{\mathrm{cMUP}} / k_{\mathrm{chem},1}^{\mathrm{MeP}} \tag{30}$$

Using $f_{\mathrm{r}}$ (**Eq. 30**) and the substrate-independent binding rate $k_1$, we can then derive a mathematical relationship between $k_{\mathrm{cat}}/K_{\mathrm{M}}$ values for cMUP and MeP (**Eq. 31**, below). We can use **Eq. 31** and previously-determined cMUP and MeP $k_{\mathrm{cat}}/K_{\mathrm{M}}$ values for PafA active site mutants to solve for $f_{\mathrm{r}}$. This value ($f_{\mathrm{r}}$), the experimentally-estimated value of $k_1 = 1.42 \times 10^6$ $\mathrm{M^{-1}s^{-1}}$ (see below), and **Eq. 31** predict the kinetic behavior of our PafA Val and Gly mutants in the absence of any misfolding effects.

$$(k_{\mathrm{cat}}/K_{\mathrm{M}})^{\mathrm{cMUP}} = \frac{f_{\mathrm{r}} k_1 (k_{\mathrm{cat}}/K_{\mathrm{M}})^{\mathrm{MeP}}}{k_1 + (f_{\mathrm{r}} - 1)(k_{\mathrm{cat}}/K_{\mathrm{M}})^{\mathrm{MeP}}} \tag{31}$$

Specifically, the value of $f_{\mathrm{r}}$ was fit using the NMinimize function of Mathematica with $\log_{10}$-scaled on-chip cMUP and MeP hydrolysis rates for wild-type and the five PafA active site mutants. MeP measurements were made with the Phosphate Sensor (PBP) under reaction conditions used on chip. While MeP $k_{\mathrm{cat}}/K_{\mathrm{M}}$ for K162A was too slow to accurately measure off chip using the PBP, a point estimate was made by scaling the previously published pNPP K162A rate constant by 6-fold to account for the previously observed ionic strength dependency of $k_{\mathrm{cat}}/K_{\mathrm{M}}$ (*16*, *106*). To generate a fit $f_{\mathrm{r}}$ consistent with on-chip data, the cMUP and MeP estimates were scaled by the ratio of $(k_{\mathrm{cat}}/K_{\mathrm{M}}^{\text{on-chip}})/(k_{\mathrm{cat}}/K_{\mathrm{M}}^{\text{off-chip}})$ for WT PafA (0.47). For fitting, $k_1$ was restrained at $1.42 \times 10^6$ $\mathrm{M^{-1}s^{-1}}$, the median on-chip WT $k_{\mathrm{cat}}/K_{\mathrm{M}}$ for cMUP. The best-fit value of $f_{\mathrm{r}}$ was 776±126 (Median±SEM), with standard error of the mean calculated from a Monte Carlo simulated distribution of $f_{\mathrm{r}}$ assuming normally-distributed error on the underlying measurements.

#### S3. A two-state model of unfolding estimates mutational misfolding effects on ($f_a$)

Assays measuring effects of altering expression temperature and zinc concentration strongly suggest that many mutations promote the formation of non-exchanging misfolded and inactive enzymes. To deconvolve and quantify mutational effects on misfolding and catalysis, we used a two-state model to quantitatively estimate the fraction of mutant enzymes in these misfolded, inactive states. As activities for active site mutants are well-fit by expected relative reactivities, we assume that each active site single mutant is fully folded and that observed losses in activity reflect purely catalytic effects ($f_{\mathrm{a}} = 1$) (**Figs. S38 and S39**).

This assumption, together with the fit relative reactivity of cMUP and MeP substrates ($f_{\mathrm{r}}$, see **Supplementary Text S3**), makes it possible to solve for the active enzyme fraction as a function of observed cMUP and MeP $k_{\mathrm{cat}}/K_{\mathrm{M}}$ values (**Eq. 32**):

$$f_{\mathrm{a}} = \frac{(f_{\mathrm{r}} - 1)(k_{\mathrm{cat}}/K_{\mathrm{M}})^{\mathrm{MeP,obs}}(k_{\mathrm{cat}}/K_{\mathrm{M}})^{\mathrm{cMUP,obs}}}{k_1 (f_{\mathrm{r}} (k_{\mathrm{cat}}/K_{\mathrm{M}})^{\mathrm{MeP,obs}} - (k_{\mathrm{cat}}/K_{\mathrm{M}})^{\mathrm{cMUP,obs}})} \tag{32}$$

We used this relationship to estimate $f_{\mathrm{a}}$, $k_{\mathrm{cat}}/K_{\mathrm{M,chem}}$, and associated confidence intervals for mutants in our valine and glycine libraries; we also estimated bootstrapped statistics and $p$-values as described in **Materials and Methods**. In two cases, we could not calculate $f_{\mathrm{a}}$: (1) mutants with $k_{\mathrm{cat}}/K_{\mathrm{M}}^{\mathrm{MeP,obs}} > k_{\mathrm{cat}}/K_{\mathrm{M}}^{\mathrm{cMUP,obs}}$, and (2) mutants with limits for both $k_{\mathrm{cat}}/K_{\mathrm{M}}^{\mathrm{MeP,obs}}$ and $k_{\mathrm{cat}}/K_{\mathrm{M}}^{\mathrm{cMUP,obs}}$ in the same direction. These mutants were therefore excluded from downstream analysis to estimate mutational effects on intrinsic catalytic parameters.

#### S4. Estimating mutational effects on intrinsic catalytic parameters

To estimate the effects of mutations on catalysis alone, we estimated $f_{\mathrm{a}}$ as described above and used this value to compute intrinsic catalytic parameters as follows:

$$(k_{\mathrm{cat}})^{\mathrm{cMUP,chem}} = (k_{\mathrm{cat}})^{\mathrm{cMUP,obs}}/f_{\mathrm{a}} \tag{33}$$

$$(k_{\mathrm{cat}}/K_{\mathrm{M}})^{\mathrm{cMUP,chem}} = (k_{\mathrm{cat}}/K_{\mathrm{M}})^{\mathrm{cMUP,obs}}/f_{\mathrm{a}} \tag{34}$$

$$(k_{\mathrm{cat}}/K_{\mathrm{M}})^{\mathrm{MeP,chem}} = (k_{\mathrm{cat}}/K_{\mathrm{M}})^{\mathrm{MeP,obs}}/f_{\mathrm{a}} \tag{35}$$

In each case, we again estimated bootstrapped statistics and $p$-values as described in **Materials and Methods**. In addition, we define the intrinsic "Catalytic effect" of mutation, using **Eq. 35**, as follows:

$$\text{Catalytic effect} = \frac{(k_{\mathrm{cat}}/K_{\mathrm{M}})^{\mathrm{MeP,chem}}_{\mathrm{mutant}}}{(k_{\mathrm{cat}}/K_{\mathrm{M}})^{\mathrm{MeP,chem}}_{\mathrm{WT}}} \tag{36}$$

#### S5. Identifying and classifying non-conserved auxiliary domains

Alkaline phosphatase (AP) superfamily members contain a universally-conserved Rossmann fold domain, the hydrophobic core of the enzyme, which bears the conserved bimetallo $Zn^{2+}$ ligands and a nucleophilic serine or threonine. AP superfamily members also contain up to 6 "auxiliary" domains (ADs) of varying length, secondary structure, and tertiary structure that are inserted at spatially-conserved junctures in the core of the Rossmann fold (**Figs. S47** and **S48**, and **Table S11**). Although the presence or absence and approximate length of particular ADs is conserved among superfamily members sharing cognate reactivities, AP phosphomonoesterases still display variability in the structure and precise length of these insertions. However, the six bimetallo $Zn^{2+}$-site ligands are conserved across all known mono- and diesterases.

To define and enumerate ADs for all AP superfamily phosphomonoesterases and phosphodiesterases for which crystal structures are available (**Table S11** and **Fig. S47**), we first aligned the six bimetallo $Zn^{2+}$-site ligands within each structure (selecting one chain to perform alignment) to those of PafA using the "align" command of PyMOL 2. Next, we identified the conserved Rossmann core within each member, defined as the regions present with shared secondary structure and well-aligned backbones (as assessed qualitatively from the alignment) in all enzymes within the structural alignment. Finally, we defined ADs to comprise any stretches of $1^{\underline{\mathrm{o}}}$ sequence present in individual superfamily members but not structurally shared among all of them. This procedure identified 3–6 ADs for each enzyme, indexed based on their point of insertion in the Rossmann core (using PafA numbering). ADs 2–4 are present are present in both phosphomonoesterases and phosphodiesterases.

AD3 is composed of two smaller domains (ADs 3.1 and 3.2) separated by only 7 or 8 residues in the Rossmann core. In monoesterases, AD 3.1 contains the K162 and R164 active site residues essential for phosphomonoesterase specificity and AD 3.2 is significantly larger, suggesting an evolutionary association between ADs 3.1 and 3.2.

#### S6. Estimating mutational effects on phospho-enzyme intermediate hydrolysis

For the three-metal AP superfamily phosphomonoesterase EcAP, phosphate release ($k_{\mathrm{off,Pi}}$) is the $k_{\mathrm{cat}}$ rate-limiting step at pH 8.0; at lower pH, hydrolysis of the covalent phospho-enzyme intermediate ($k_{\mathrm{chem,2}}$) becomes rate limiting (*53*). Independent evidence suggests that EcAP mutations alter $k_{\mathrm{off,Pi}}$ and $k_{\mathrm{chem,2}}$ (*107*). As the WT PafA catalytic cycle is highly similar to that of EcAP, PafA $k_{\mathrm{cat}}$ is likely rate-limited by one or both of these catalytic steps, and mutations may alter these rates. To quantify $k_{\mathrm{chem,2}}$ for WT PafA and all mutants, we first estimated the rate of phosphate release, $k_{\mathrm{off,Pi}}$ (**Eq. 37**):

$$k_{\text{off,Pi}} = K_{\text{i}}(\text{P}_{\text{i}}) \times k_{\text{on,Pi}} \tag{37}$$

Here, we use the measured $K_{\text{i}}$ for $\text{P}_{\text{i}}$ from on-chip HT-MEK experiments and assume that $k_{\text{on,Pi}}$ is approximately the apparent rate constant for substrate binding to the enzyme. As the $k_{\text{cat}}/K_{\text{M}}$ for activated aryl substrates (*e.g.* cMUP) is binding-limited (*19*) and we expect that $k_{\text{on,Pi}}$ will not vary substantially upon mutation (*107*), we can estimate $k_{\text{on,Pi}}$ using the WT PafA $k_{\text{cat}}/K_{\text{M}}$ for cMUP ($k_{\text{on,Pi}} \approx k_{\text{cat}}/K_{\text{M}}^{\text{cMUP,WT}} = 1.42 \times 10^6\ \text{M}^{-1}\text{s}^{-1}$).

Next, we estimated the rate of covalent phospho-enzyme intermediate hydrolysis ($k_{\text{chem,2}}$) for all mutants with an estimated $k_{\text{off,Pi}}$ and measured $k_{\text{cat}}^{\text{cMUP,chem}}$ (the $f_{\text{a}}$-corrected $k_{\text{cat}}$, **Eq. 33**) for cMUP (992/1036). To do this, we first derived an expression for the PafA $k_{\text{cat}}$ using the method of net rate constants (*108*), assuming that $k_{\text{chem,2}} \gg k_{\text{chem,-2}}$ (**Eq. 38**). This assumption is supported by the observations that $k_{\text{chem,2}}/k_{\text{chem,-2}} = 100$ and 40 for WT EcAP and R166S EcAP, respectively (*107*).

$$k_{\text{cat}}^{\text{cMUP,chem}} = \frac{1}{\frac{1}{k_{\text{off,Pi}}} + \frac{1}{k_{\text{chem,2}}} + \frac{1}{k_{\text{chem,1}}} + \frac{1}{k_1 \frac{k_{\text{chem,1}}}{k_{\text{-1}} + k_{\text{chem,1}}}}} \approx \frac{1}{\frac{1}{k_{\text{off,Pi}}} + \frac{1}{k_{\text{chem,2}}}} = \frac{k_{\text{off,Pi}} k_{\text{chem,2}}}{k_{\text{off,Pi}} + k_{\text{chem,2}}} \tag{38}$$

Next, we substituted the expression for $k_{\text{off,Pi}}$ from **Eq. 37**:

$$k_{\text{cat}}^{\text{cMUP,chem}} = \frac{k_{\text{on,Pi}} K_{\text{i,Pi}} k_{\text{chem,2}}}{k_{\text{on,Pi}} K_{\text{i,Pi}} + k_{\text{chem,2}}} \tag{39}$$

Finally, solving for $k_{\text{chem,2}}$ and substituting with **Eq. 33**

$$k_{\text{chem,2}} = \frac{k_{\text{on,Pi}} K_{\text{i,Pi}} k_{\text{cat}}^{\text{cMUP,chem}}}{k_{\text{on,Pi}} K_{\text{i,Pi}} - k_{\text{cat}}^{\text{cMUP,chem}}} = \frac{k_{\text{on,Pi}} K_{\text{i,Pi}} k_{\text{cat}}^{\text{cMUP,obs}}}{f_{\text{a}} k_{\text{on,Pi}} K_{\text{i,Pi}} - k_{\text{cat}}^{\text{cMUP,obs}}} \tag{40}$$

Using this expression, we estimated $k_{\text{chem,2}}$ for all mutants with measurements for $K_{\text{i}}(\text{P}_{\text{i}})$ and $k_{\text{cat}}^{\text{cMUP,chem}}$. Limits in $K_{\text{i}}(\text{P}_{\text{i}})$ and/or $f_{\text{a}}$ yield limits in $k_{\text{chem,2}}$, and these limit cases were i) $k_{\text{chem,2}}$ upper limit, arising from a lower $f_{\text{a}}$ limit and a lower, or no, limit in $K_{\text{i}}(\text{P}_{\text{i}})$; ii) $k_{\text{chem,2}}$ lower limit, arising from an upper limit in $K_{\text{i}}(\text{P}_{\text{i}})$ and an upper, or no limit, in $f_{\text{a}}$; iii) undefined $k_{\text{chem,2}}$ limit, arising from a lower limit in $K_{\text{i}}(\text{P}_{\text{i}})$ and an upper limit in $f_{\text{a}}$, or an upper limit in $K_{\text{i}}(\text{P}_{\text{i}})$ and a lower limit in $f_{\text{a}}$ (these estimates were omitted from plots and downstream analyses). Finally, a lower limit in $K_{\text{i}}(\text{P}_{\text{i}})$ yields a range of potential values of $k_{\text{chem,2}}$ (case iv.), but we treated this limit case as a point estimate (*i.e.*, no limit) for plots and analyses as the bounded range is small for most mutants.

Estimates of mutational effects on $k_{\text{chem,2}}$ rely on measurements of the $K_{\text{i}}$ for $\text{P}_{\text{i}}$ and $k_{\text{cat}}^{\text{cMUP,chem}}$, and the latter relies on accurate estimation of mutational effects on the active enzyme fraction ($f_{\text{a}}$). Measurement error in any of these quantities renders estimates of mutational effects on $k_{\text{chem,2}}$ less precise. To account for this uncertainty, we calculated $p$-values for mutant $k_{\text{chem,2}}$ estimates via bootstrap hypothesis testing in an analogous manner as for $k_{\text{cat}}/K_{\text{M}}^{\text{chem}}$ and $f_{\text{a}}$ effects (see **Materials and Methods**, *Determining statistical significance of fraction active and catalytic effects within the active fraction*).

### Figures S1 to S64

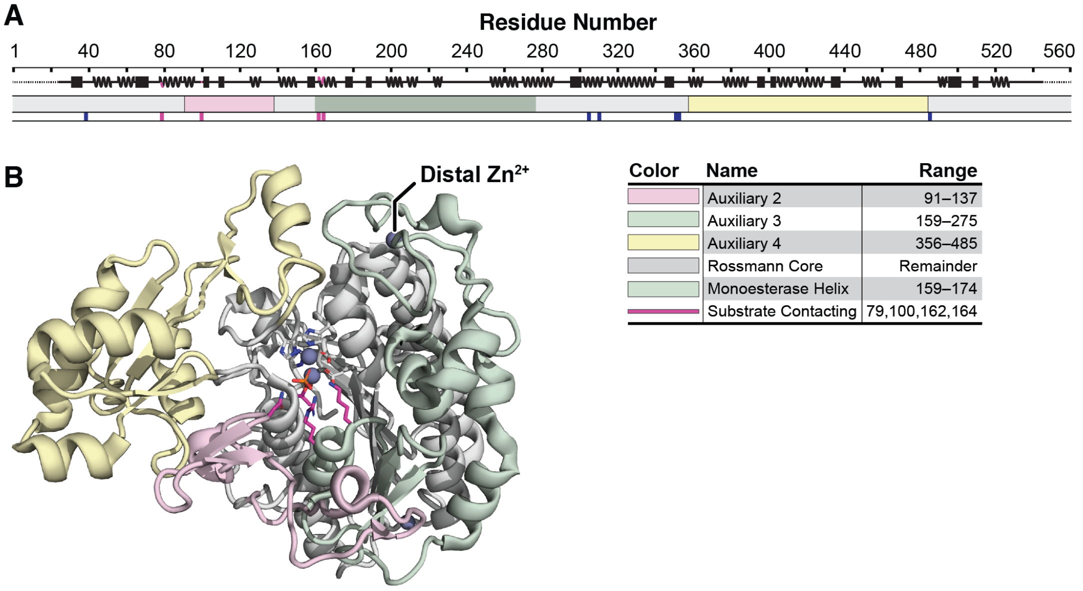

**Fig. S1.** Structural overview of PafA phosphatase. (A) Linear topology of PafA showing secondary structure, named domains and structural elements (colored as in the legend below and to the right), and key catalytic residues, including active site residues (magenta) and $Zn^{2+}$ ligands (dark blue). (B) Crystal structure of WT PafA (PDB ID: 5TJ3) with conserved superfamily core fold (Rossmann fold) in white and insertion ("Auxiliary") domains colored as in panel A. Besides the two active site $Zn^{2+}$ ions, one of two additional $Zn^{2+}$ ions is annotated ("Distal $Zn^{2+}$").

A

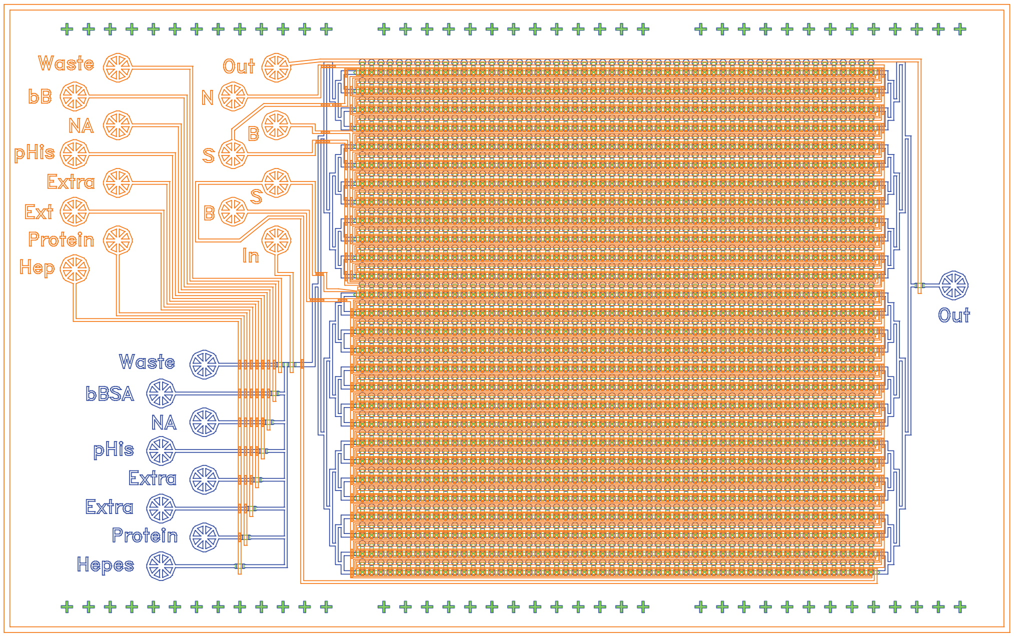

B

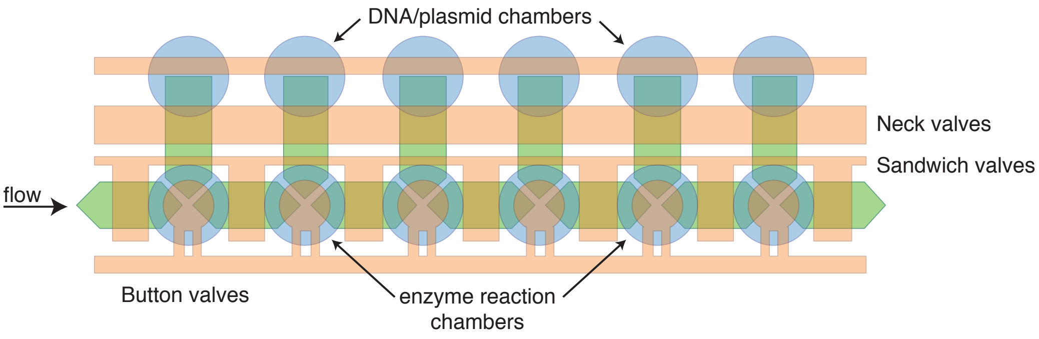

**Fig. S2.** Detailed schematic of the microfluidic device used for HT-MEK measurements. (A) PDMS device "control" (orange) and "flow" (blue) layers, with input ports labeled (*68*, *69*). Each device contains 1,568 chambers (56 chambers arrayed along each of 28 fluidically-connected channels). (B) Detailed schematic for six example reaction chambers along a single channel. Each chamber contains two compartments, "DNA/plasmid" (DNA) and "enzyme reaction" (Reaction), with fluid flow within and between chambers controlled by three valves: a Sandwich valve (to control fluid flow between chambers), a Neck valve (to control fluid flow between compartments), and a Button valve that restricts fluidic access to a particular patch of the device surface for surface patterning, enzyme immobilization, and fluid exchange.

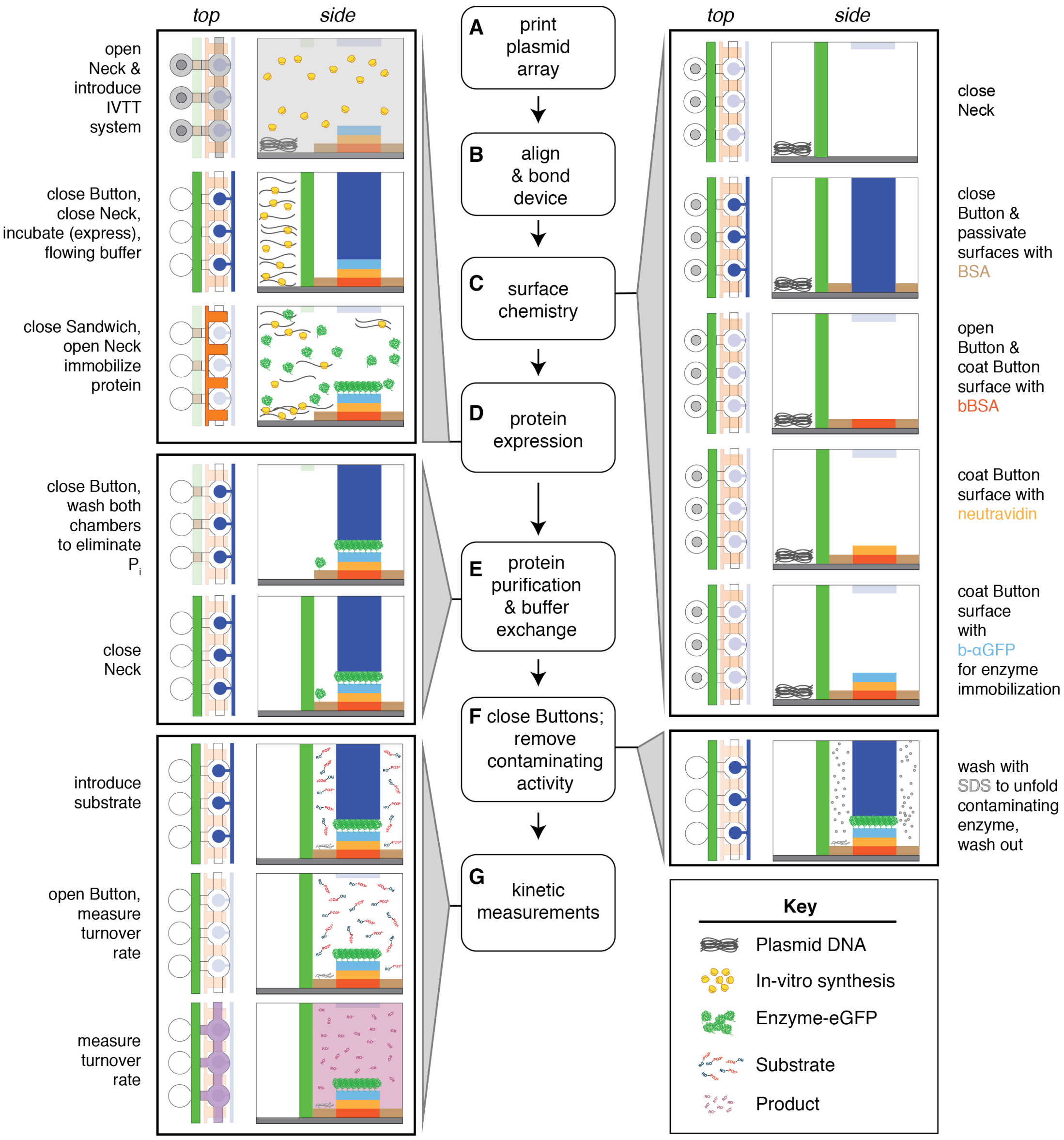

**Fig. S3.** Schematic showing workflow for on-chip surface patterning, enzyme expression and purification, and subsequent kinetic assays via HT-MEK.

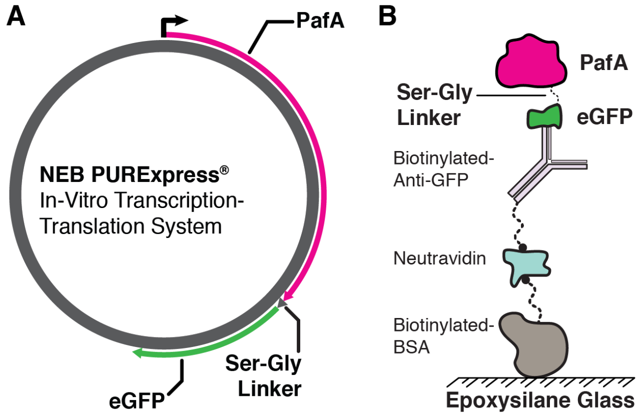

**Fig. S4.** Protein expression constructs and linkages for surface immobilization. (A) Vector map of the PafA-eGFP PURExpress construct with flexible Ser-Gly linker. (B) Molecular linkages tethering expressed PafA-eGFP constructs to epoxysilane glass surfaces within the HT-MEK device.

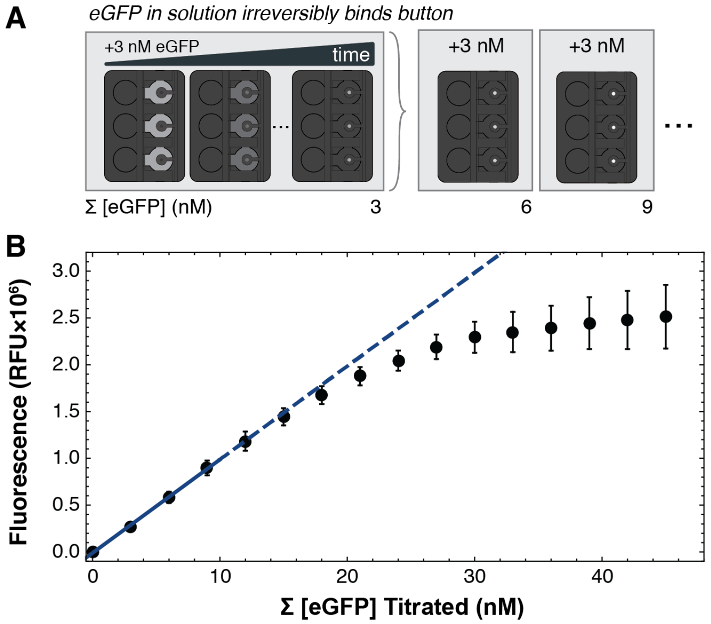

**Fig. S5.** Calibration curves relating surface-immobilized eGFP intensities to effective enzyme concentrations in each chamber. (A) Schematic of calibration curve acquisition showing iterative introduction of 3 nM eGFP into each reaction chamber, opening of the Button valves to expose anti-eGFP patterned device surfaces, and binding of eGFP until saturation. Each titration repeated this process at least 12 times (to a total summed concentration of at least 36 nM). (B) Measured eGFP intensities beneath the Button valve after 15 iterative introductions of 3 nM eGFP. Points denote the median fluorescence in all chambers across the device; error bars indicate the standard error on the mean (SEM). A linear fit (blue) to measured fluorescence intensity as a function of [eGFP] for the first 4–9 points yielded fit parameters that were subsequently used to convert measured intensities within each chamber to product concentrations during kinetic assays (slope = ~100,000–200,000 RFU/nM).

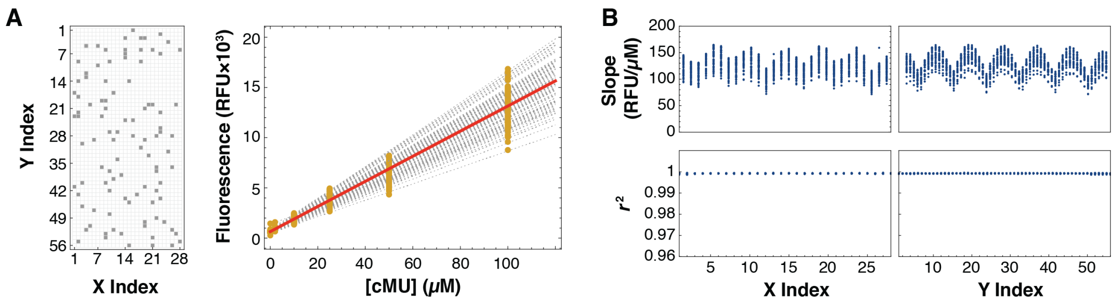

**Fig. S6.** Per-chamber on-chip product calibration curves for an example fluorescent product (cMU, see **Materials and Methods**). (A) Measured fluorescence intensities as a function of introduced cMU concentration (right) for 100 chambers at various positions within the device (left). Median intensities for each chamber are shown in gold, calibration curves for each chamber are shown as black dashed lines, and a linear fit to the median of all chamber intensities at a given concentration is shown in red. (B) Observed variation in slope as a function of chamber position within the device (top) and returned coefficients of determination for linear fits (bottom). The observed regular variation in slope, as a function of device position, is expected as a result of image vignetting.

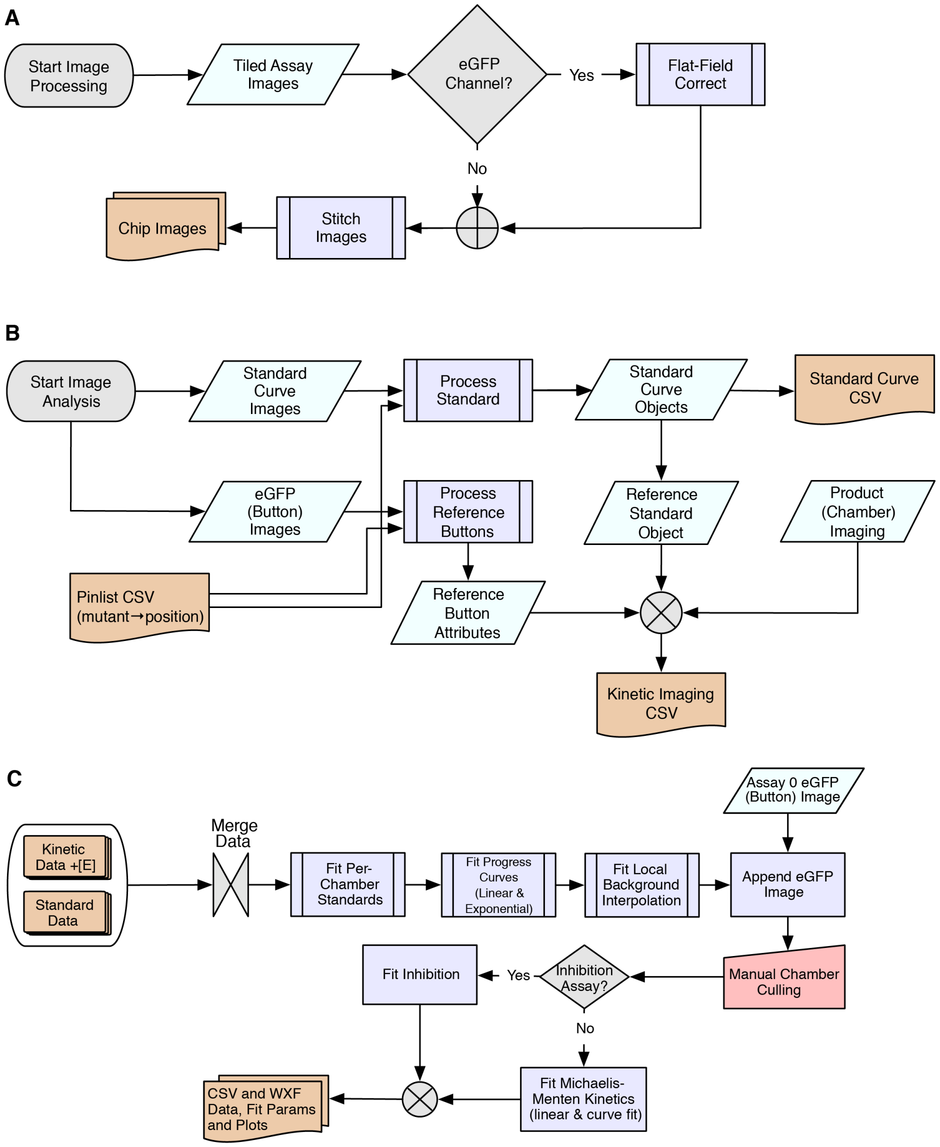

**Fig. S7.** Image processing and Michealis-Menten fitting pipelines (code online at https://github.com/FordyceLab/; CSVs and data summaries available in the OSF Repository Data, https://osf.io/ez9xv/). (A) Image processing workflow. Overlapping tiled chip images (1024 × 1024, 16-bit depth) are flat-field corrected if collected in the eGFP channel, then stitched to generate single images of the entire device at each assay time point. (B) Image post-processing pipeline. Product standard curve images, eGFP Button images, and enzymatic turnover chamber images are combined, associated with chamber variant identities (Pinlist), and processed to yield a data table describing kinetic progress curves for each chamber. (C) Pipeline for processing kinetic data and applying quality control criteria to yield Michaelis-Menten parameters and inhibition constants.

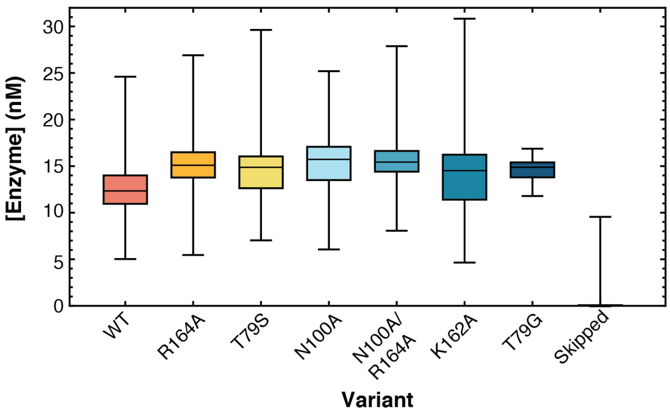

**Fig. S8.** On-chip expression of PafA active site mutants. Calibrated per-chamber enzyme concentrations for seven PafA-eGFP variants and chambers in which no plasmid was printed (Skipped). Apparent expression in several Skipped chambers is inflated by enzyme leakage from adjacent DNA-containing chambers; in kinetic experiments, these chambers would be identified and eliminated from further analysis (*cf.* **Materials and Methods**, *Analysis and quality control of cMUP hydrolysis measurements*).

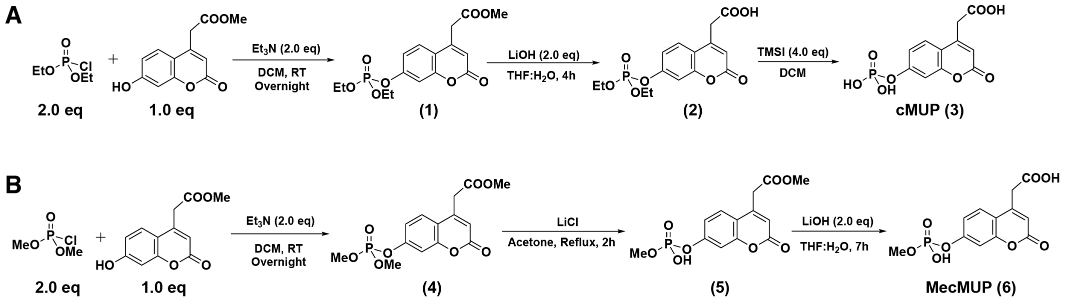

**Fig. S9.** Synthesis of carboxy-coumarinyl phosphomonoester and phosphodiester substrates. (A) Synthesis of cMUP. (B) Synthesis of MecMUP.

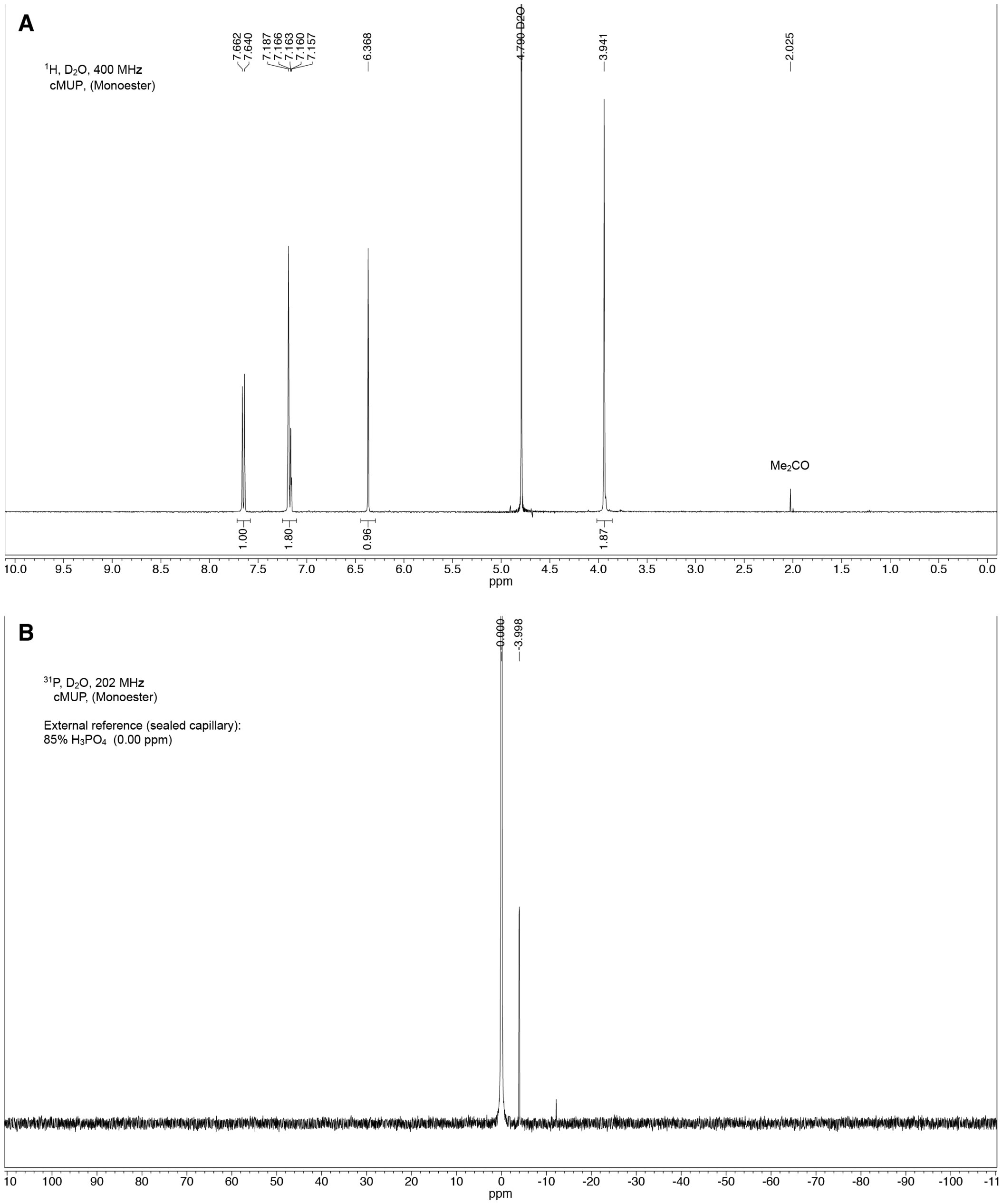

**Fig. S10.** $^{1}$H (A) and $^{31}$P (B) NMR spectra of 2-(2-oxo-7-(phosphonooxy)-2H-chromen-4-yl)acetic acid (cMUP; **Fig. S9A** compound **3**).

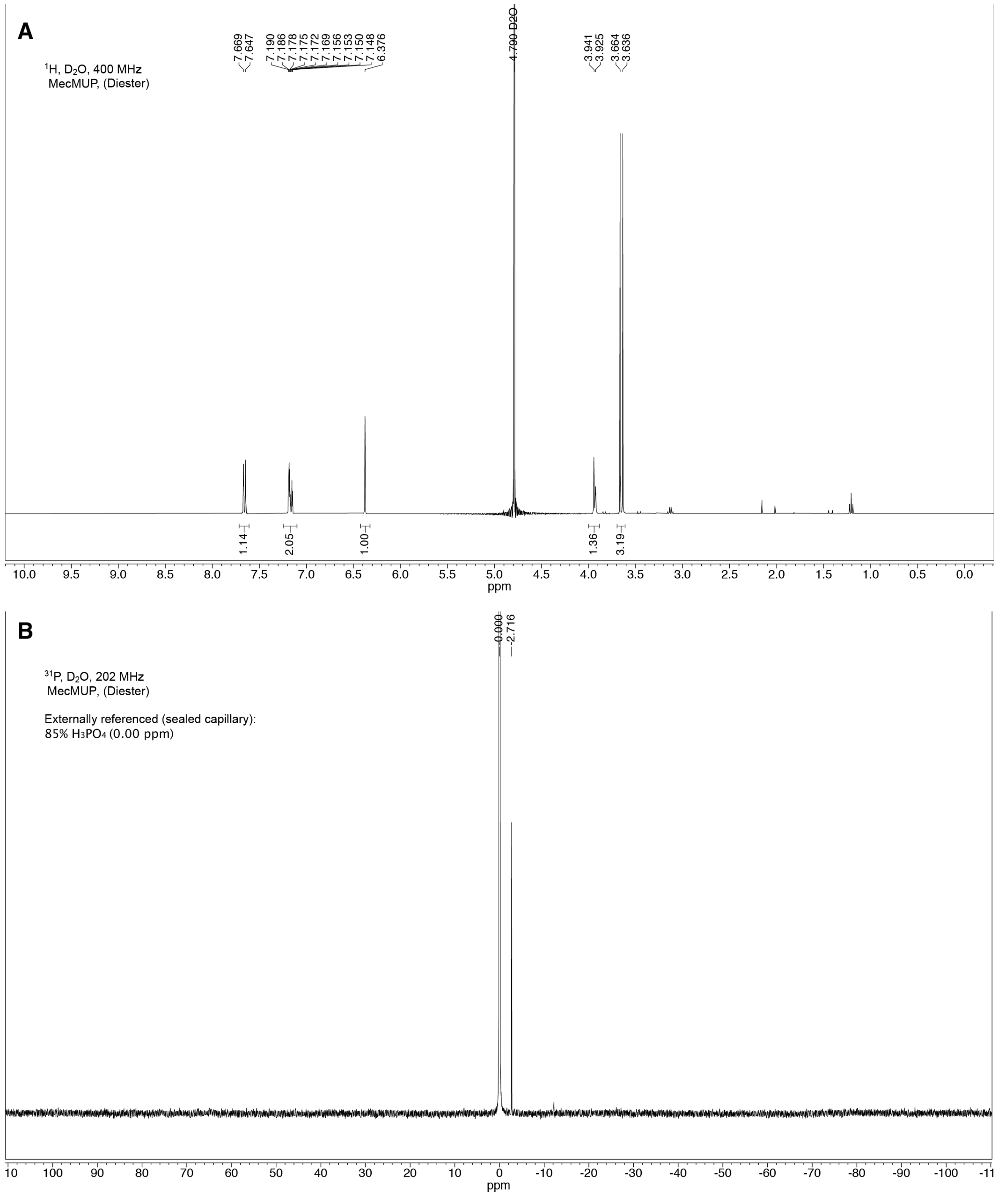

**Fig. S11.** $^{1}$H (A) and $^{31}$P (B) NMR spectra of 2-(7-((hydroxy(methoxy)phosphoryl)oxy)-2-oxo-2H-chromen-4-yl)acetic acid (MecMUP; **Fig. S9B** compound **6**).

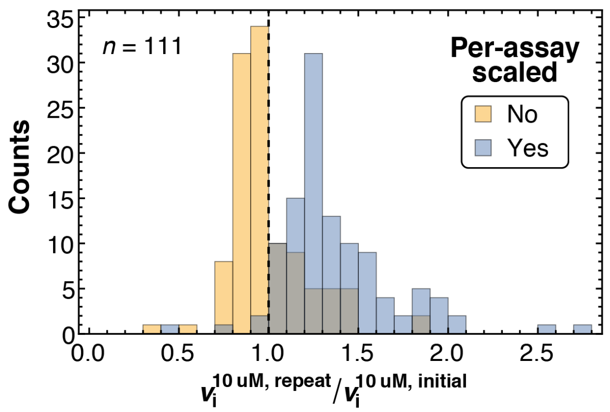

**Fig. S12.** Scaling each rate by the enzyme concentration ([E]) determined before each assay results in systematic overestimation of later rates. Comparison of the ratios of initial rates in each chamber obtained for an initial 10 µM cMUP assay and a replicate assay acquired ∼6 hours later, scaling rates measured in both assays by the first eGFP measurement only (yellow), and those obtained after scaling by the [E] determined before each assay (blue).

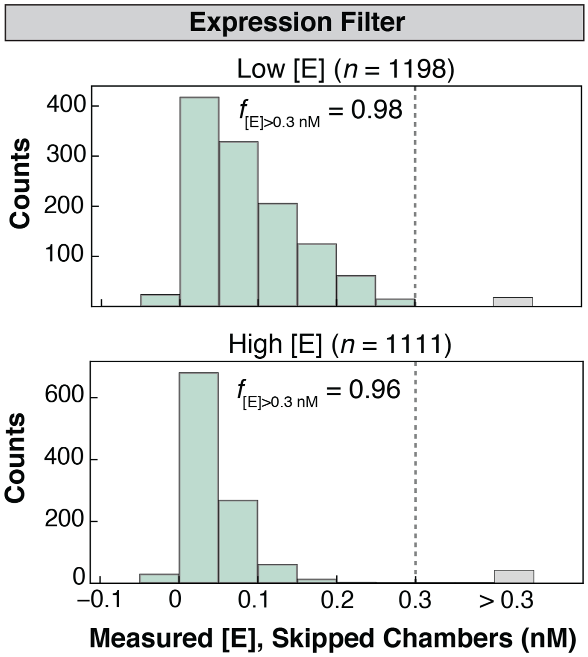

**Fig. S13.** Eliminating chambers with poor enzyme expression from further analysis. Distribution of measured eGFP intensities in "Skipped" chambers (in which no plasmid was printed) for low [E] (top) and high [E] (bottom) devices. For both sets of experiments, >95% of "Skipped" chambers had measured [E] $\leq$ 0.3 nM (grey dashed line). We therefore flagged any chambers with $\leq$ 0.3 nM immobilized enzyme as lacking successful expression.

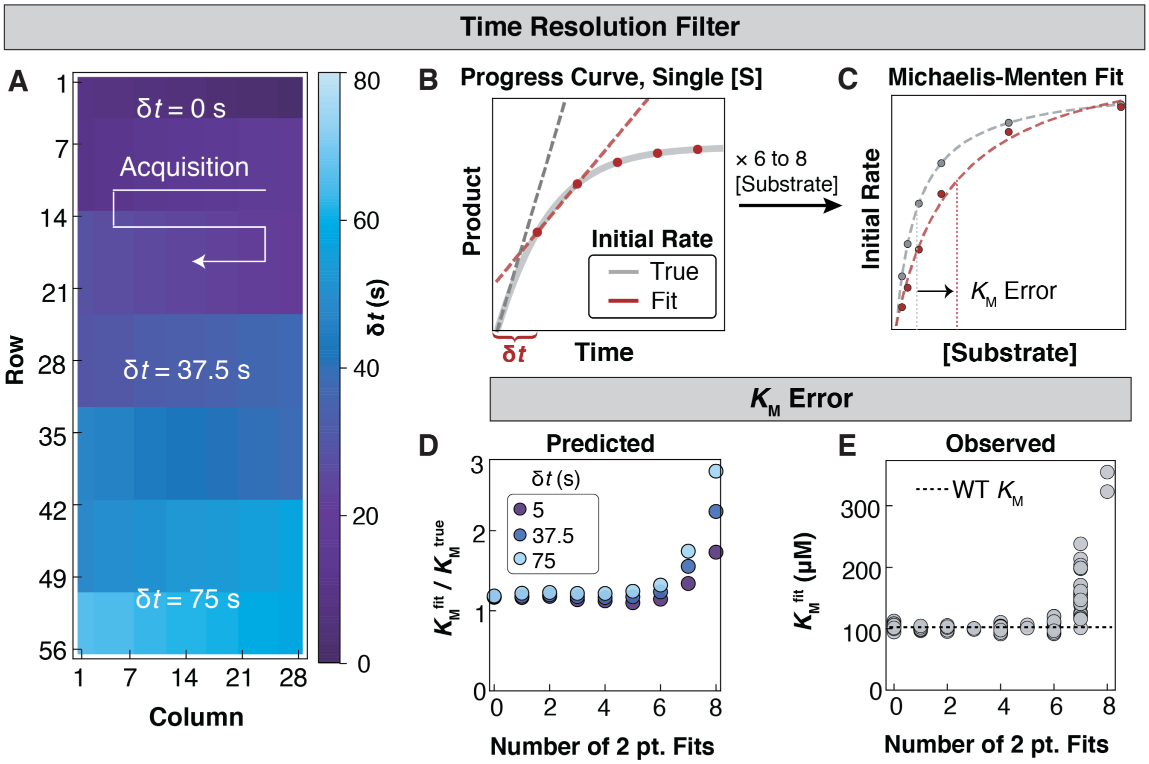

**Fig. S14.** Eliminating chambers with rates too fast to be measured accurately. (A) Heat map showing initial image acquisition delay time as a function of device position. (B) Simulated progress curves for a WT enzyme illustrating how a linear fit (red dashed line) to measurements sampled with a long initial delay (red circles) can underestimate the true rate (gray dashed line). (C) Simulated Michaelis-Menten curves illustrating how Michaelis-Menten fits to initial rates estimated from measurements with a long initial delay (red points) can lead to systematic overestimates of $K_M$ (red curve; "true" values and associated fit shown in gray). (D) $K_M$ values obtained from Michaelis-Menten fits of simulated progress curves assuming WT parameters ($k_{cat}$ = 200 $s^{-1}$, $K_M$ = 100 μM and [E] = 10 nM), for three different values of $\delta t$. Underestimation ($K_M^{fit}/K_M^{true}$) increases with increasing $\delta t$ or the number of initial rates with two-point fits used to generate the Michaelis-Menten curve. (E) Fit $K_M$ values for WT chambers as a function of the number of two-point initial rate fits in the Michaelis-Menten curve. Observed systematic errors match those predicted from the simulations in panel D.

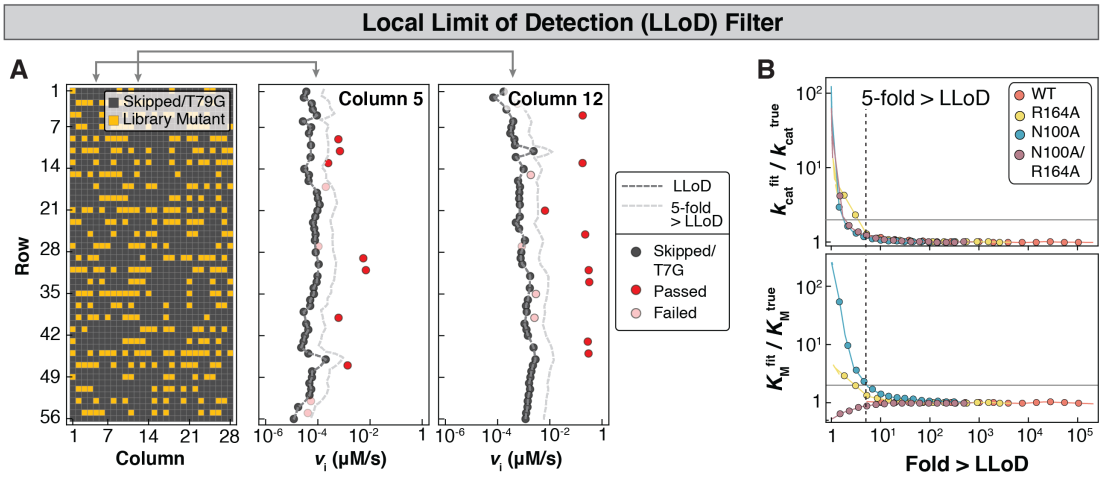

**Fig. S15.** Quality control to eliminate chambers with artificially high observed rates resulting from WT-like contamination. (A) Heat map (left) of a representative tier I device showing the location of library mutants (gold), and Skipped and T79G chambers (gray); observed rates as a function of row position for two example columns (channels) for Skipped/T79G chambers (gray) and mutant chambers that either passed (red) or failed (pink) the quality control criteria (observed rate >5-fold above the interpolated local limit of detection (LLoD, light gray dashed line). (B) Simulations used to determine a threshold of 5×LLoD. Fractional errors in $k_{\mathrm{cat}}$ (top) and $K_{\mathrm{M}}$ (bottom) as a function of the fold change above LLoD (itself a function of the amount of contaminant included in the simulations).

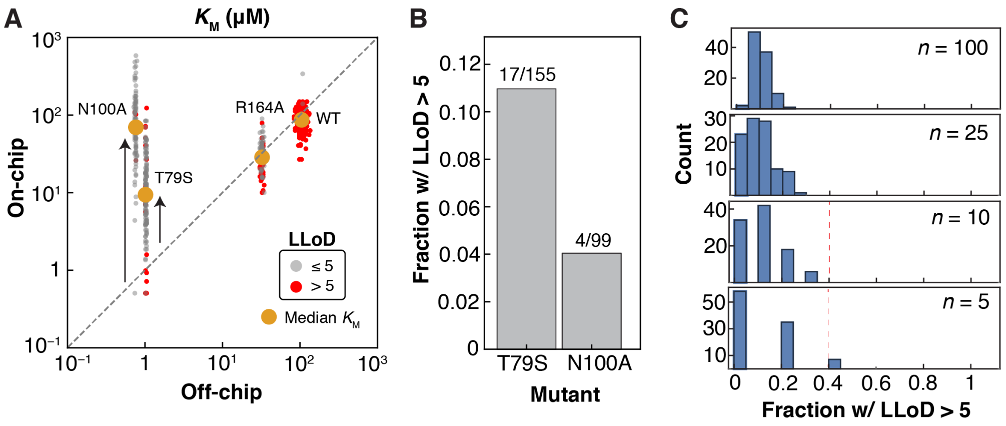

**Fig. S16.** Additional culling of LLoD false positives (>5-fold above the LLoD). (A) Determination of the LLoD false positive rate on tier I devices using a comparison of all on-chip measurements against the off-chip $K_M$ values of PafA active site mutants. The tier I values of $K_M$ indicate that T79S and N100A cannot be measured accurately (black arrows) due to contamination from WT-like enzyme. (B) The large number of T79S and N100A replicates (155 and 99, respectively) allows estimation of the false positive rate. The fraction of total T79S and N100A chambers that passed LLoD culling (*cf.* red and gray points in panel A) estimate a false positive rate of 10%. (C) Effect of a 10% false positive rate on the observed fraction of replicates >5-fold above the LLoD, as a function of the number of mutant replicates obtained ($n$, where $n$ = 0, 10, 25, and 100). One hundred simulations were performed at each number of replicates. As we typically obtained 5–10 replicates of each mutant, the rates for any mutant for which <40% of total replicates passed the LLoD culling (red dashed line) on tier I devices was considered to be due to contamination. These tier I replicates were therefore not included in subsequent analyses and measurements were taken from the tier 2 and 3 devices only.

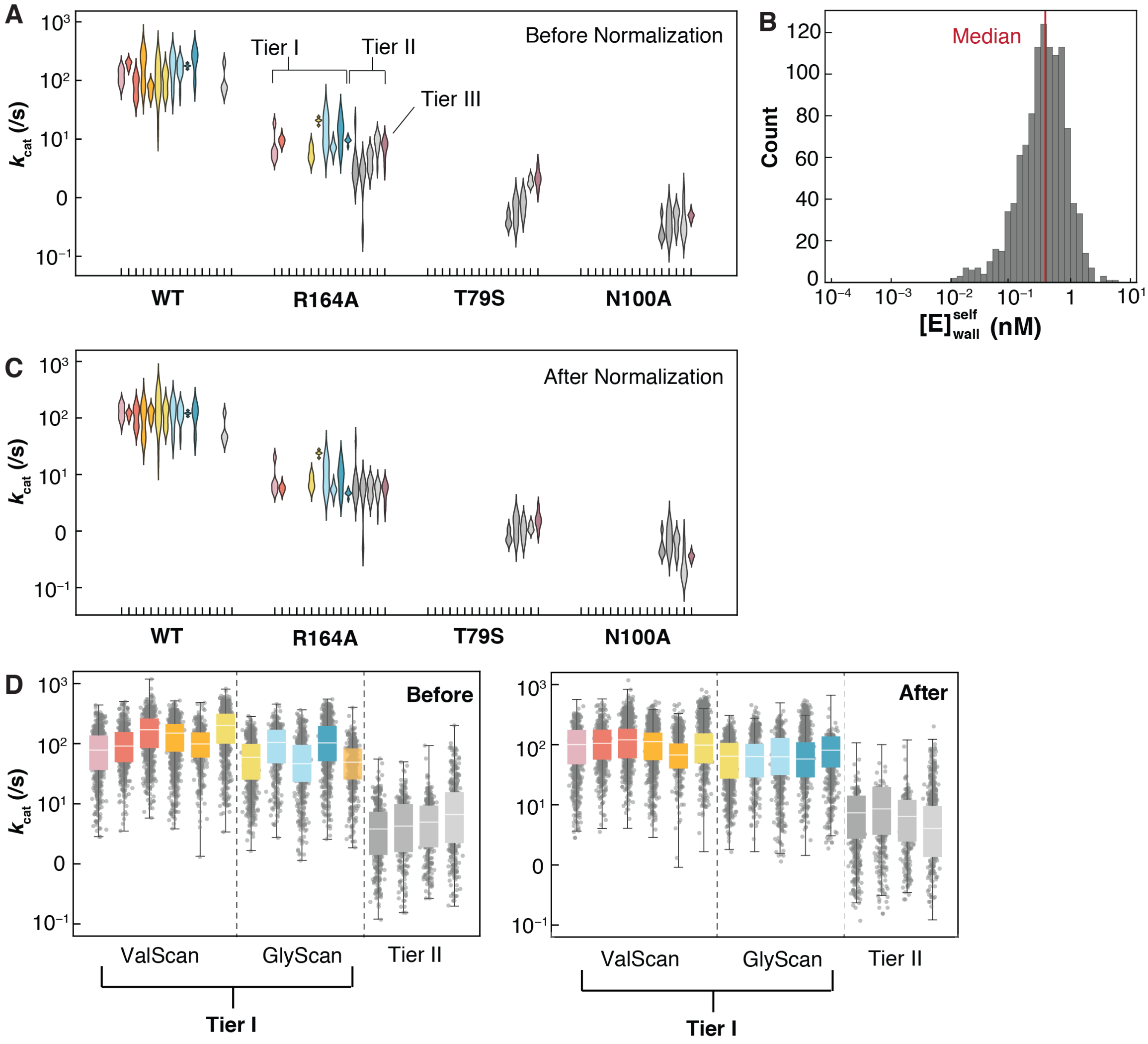

**Fig. S17.** Normalizing data across experiments and tiers. (A) $k_{cat}$ distributions within each experiment for WT and active site single mutants before normalization. (B) Distribution of contributions from non-specifically bound enzyme ($E_{wall}^{self}$) to the total [E] within each chamber. (C) $k_{cat}$ distributions within each experiment for WT and active site single mutants after normalization. (D) $k_{cat}$ distributions within each experiment for all chambers passing quality control filters, before and after cross-experiment normalization for tiers 1 and 2. White lines denote the median and boxes span 25% and 75% quantiles.

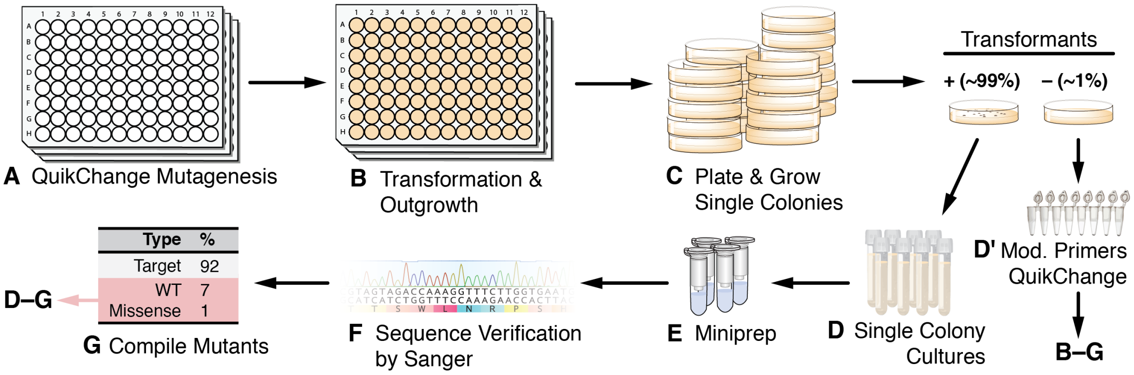

**Fig. S18.** High-throughput mutagenesis protocol to generate comprehensive single-site mutant libraries. (A) Parallelized QuikChange site-directed mutagenesis performed in 96-well plates. (B) Transformation of ultra-competent DH5alpha *E. coli* with QuikChange products and subsequent culture in deep well plates in SOC media for 1 h. (C) Plating of reactions on LB agar with ampicillin prior to growth overnight. (D) Picking single colonies for growth in 10 mL LB with ampicillin along with additional rounds of primer design, QuikChange, and transformation for reactions that did not yield colonies. (E) Miniprep plasmid isolation from 10 mL liquid cultures. (F) Sanger sequencing to verify construct sequence. (G) Picking, culturing, and sequencing additional colonies in cases where first-round sequencing identified WT constructs, undesired missense mutations, or insertions/deletions.

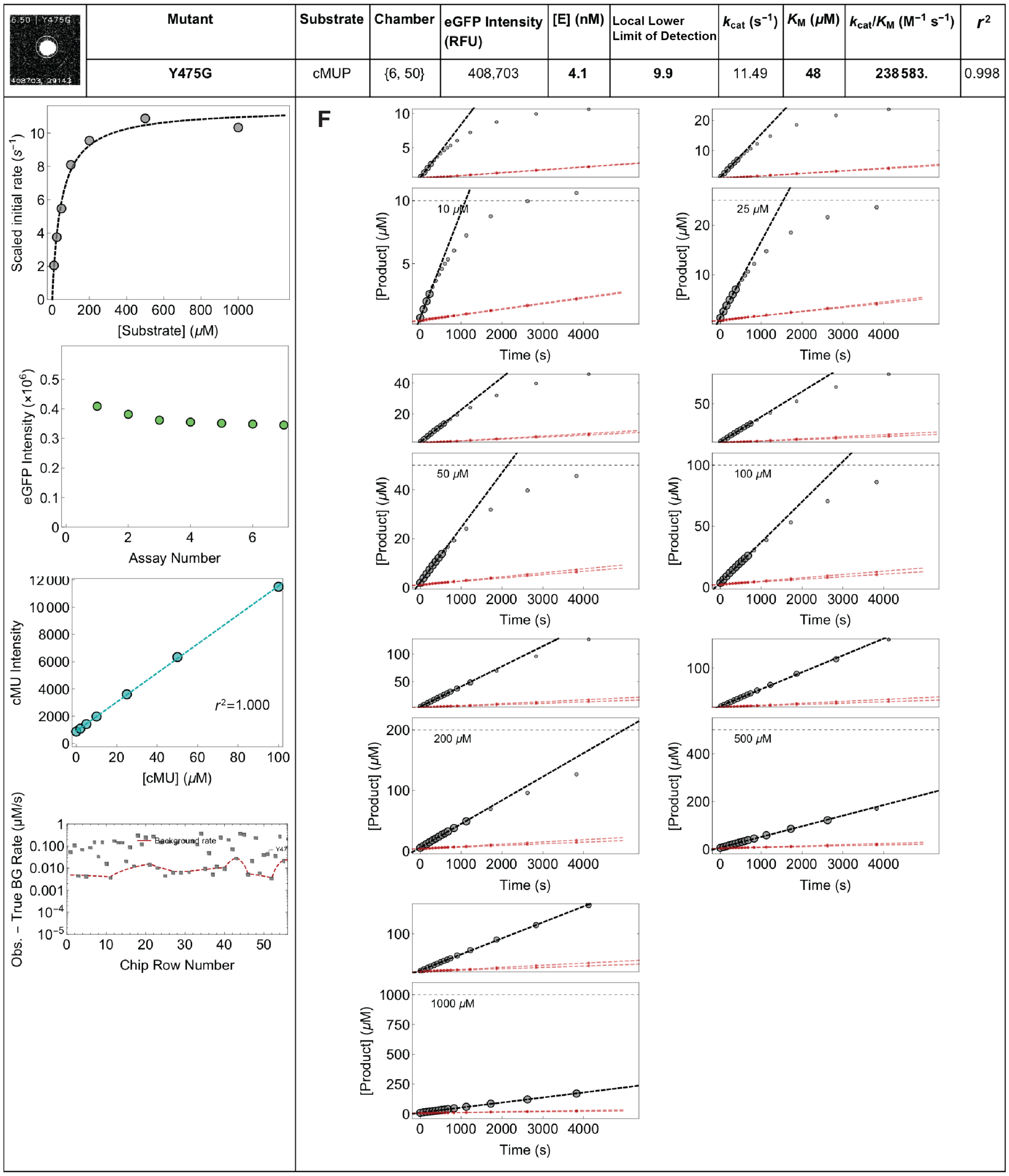

| Mutant | Substrate | Chamber | eGFP Intensity (RFU) | [E] (nM) | Local Lower Limit of Detection | $k_{cat}$ ($s^{-1}$) | $K_M$ ($\mu$M) | $k_{cat}/K_M$ ($M^{-1}$ $s^{-1}$) | $r^2$ |
|---|---|---|---|---|---|---|---|---|---|
| Y475G | cMUP | {6, 50} | 408,703 | 4.1 | 9.9 | 11.49 | 48 | 238583. | 0.998 |

**Fig. S19.** Example data report for a single chamber following an HT-MEK cMUP kinetic experiment (7 individual cMUP assays). Each experiment returns 1,568 unique per-chamber reports containing: (A) Mutant identity, image of immobilized enzyme (eGFP channel), experimental parameters (substrate, chamber column and row, eGFP intensity, enzyme concentration, the rate compared to the local background (Local Lower Limit of Detection, or LLoD, see (E) and **Materials and Methods**), and Michaelis-Menten fit parameters ($k_{\mathrm{cat}}$, $K_{\mathrm{M}}$, $k_{\mathrm{cat}}/K_{\mathrm{M}}$, and associated $r^2$ values); (B) Michaelis-Menten fits to enzyme-concentration-scaled initial rates (gray); (C) Measured eGFP intensity prior to each assay; (D) Per-chamber product fluorescence standard curve and fit $r^2$; (E) Comparison of fitted rates (gray points) with the interpolated Local Lower Limit of Detection (LLoD, red dashed line) for all chambers in the same column; (F) Full reaction progress curves (lower panels) and zoomed in plots (upper panels) for each substrate concentration with fitted initial rates (dashed gray lines). Large gray points were used to determine the linear fit; small gray points were measured but outside the linear region and not used to fit initial rates. Red points and lines denote the progress curves and linear fits from the nearest upstream and downstream background chambers used to determine the local background.

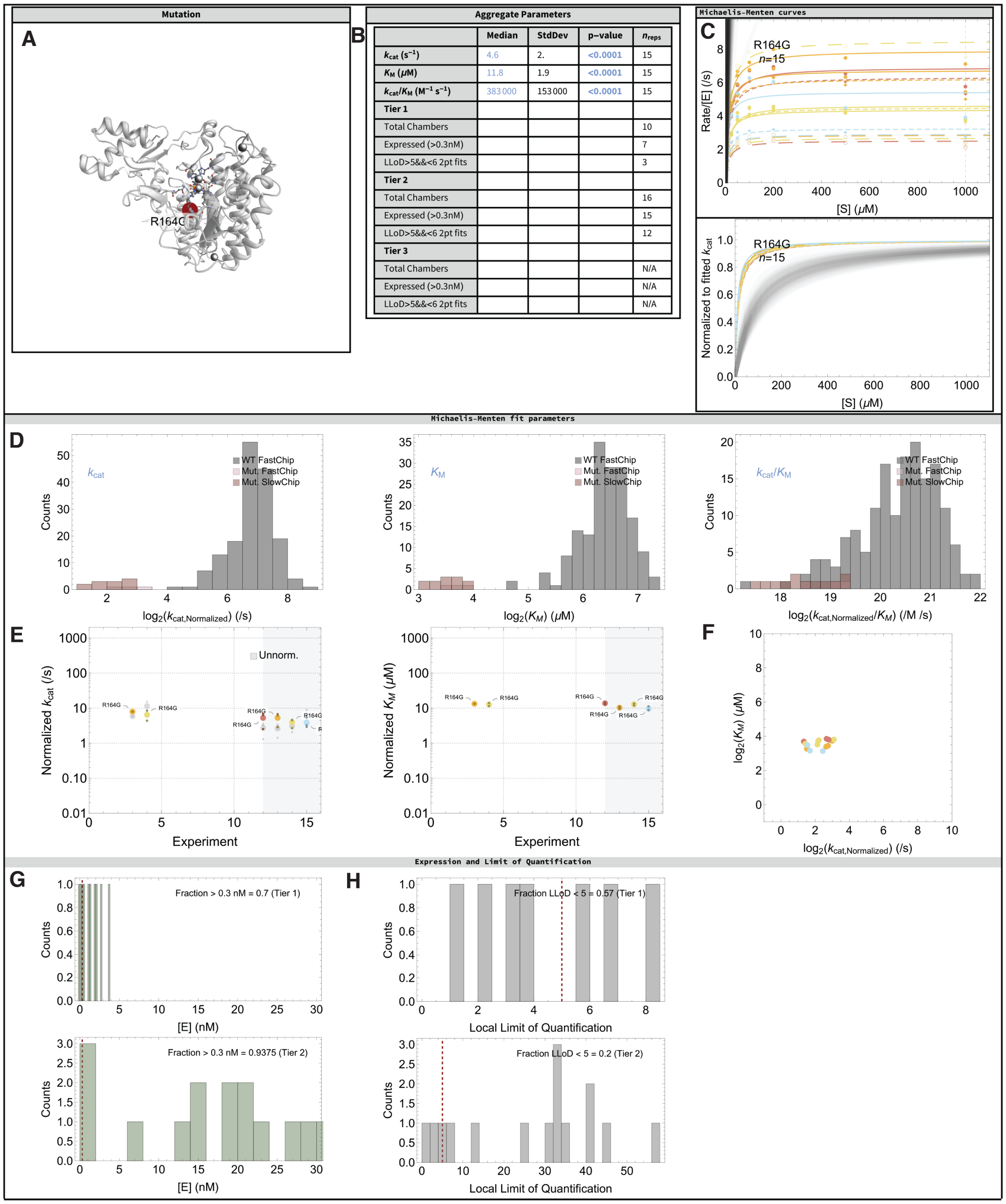

| | Median | StdDev | p-value | $n_{reps}$ |
|---|---|---|---|---|
| $k_{cat}$ ($s^{-1}$) | 4.6 | 2. | <0.0001 | 15 |
| $K_M$ (μM) | 11.8 | 1.9 | <0.0001 | 15 |
| $k_{cat}/K_M$ ($M^{-1} s^{-1}$) | 383 000 | 153 000 | <0.0001 | 15 |
| Tier 1 | | | | |
| Total Chambers | | | | 10 |
| Expressed (>0.3nM) | | | | 7 |
| LLoD>5&&<6 2pt fits | | | | 3 |
| Tier 2 | | | | |
| Total Chambers | | | | 16 |
| Expressed (>0.3nM) | | | | 15 |
| LLoD>5&&<6 2pt fits | | | | 12 |
| Tier 3 | | | | |
| Total Chambers | | | | N/A |
| Expressed (>0.3nM) | | | | N/A |
| LLoD>5&&<6 2pt fits | | | | N/A |

**Fig. S20.** Example aggregate cMUP summary for a representative mutant. (A) Name and location of mutation in PafA crystal structure (red sphere). (B) Aggregate parameter table. Number of replicates expressed and number that passed the LLoD filter in each experimental tier. (C) Fitted Michaelis-Menten curves for this mutant (colored lines) and for WT PafA (gray lines). (D) Distributions of mutant $k_{\mathrm{cat}}$, $K_{\mathrm{M}}$, and $k_{\mathrm{cat}}/K_{\mathrm{M}}$ (red) relative to WT distributions (gray). (E) $k_{\mathrm{cat}}$ and $K_{\mathrm{M}}$ for replicates within each experiment. The gray region denotes tier 2 and 3 experiments. (F) Scatter plot of $k_{\mathrm{cat}}$ vs. $K_{\mathrm{M}}$. (G) Histograms of [E] for each replicate. Red dashed line denotes expression filter cutoff of 0.3 nM. (H) Histograms of $v_{\mathrm{i}}$/LLoD, for all replicates. Red dashed line denotes the LLoD cutoff (5-fold above the LLoD). Aggregate pages also contain a table of $k_{\mathrm{cat}}$, $K_{\mathrm{M}}$, and $k_{\mathrm{cat}}/K_{\mathrm{M}}$ values for each replicate (not shown).

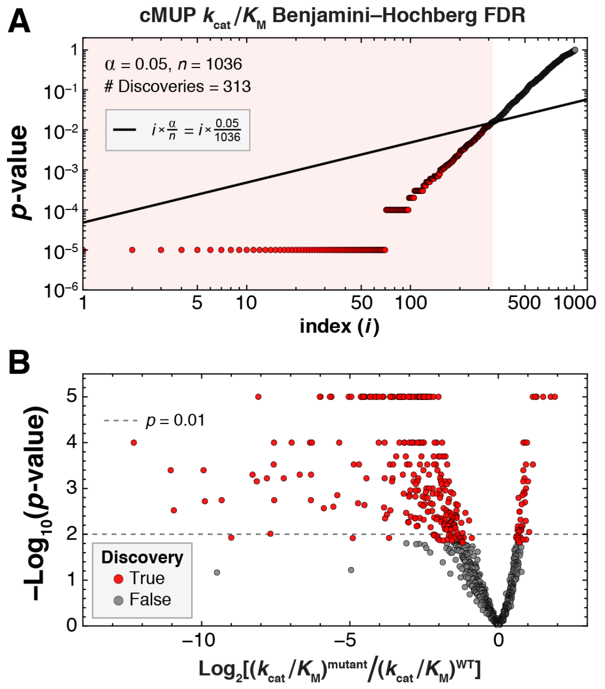

**Fig. S21.** Benjamini-Hochberg false discovery rate (FDR) controlling procedure. (A) $p$-values for median cMUP $k_{\text{cat}}/K_{\text{M}}$ values for 1036 Val and Gly mutants (markers), sorted by $p$-value. Black line indicates predicted Benjamini-Hochberg FDR at $\alpha = 0.05$, calculated using the equation shown in the inset. The pink box denotes the range of observations considered discoveries under our FDR controlling procedure. (B) Volcano plot showing true discoveries declared under this FDR controlling procedure (red) co-plotted with all other observations (gray). The horizontal dashed line denotes $p = 0.01$, the threshold used to define strong evidence of mutational effects.

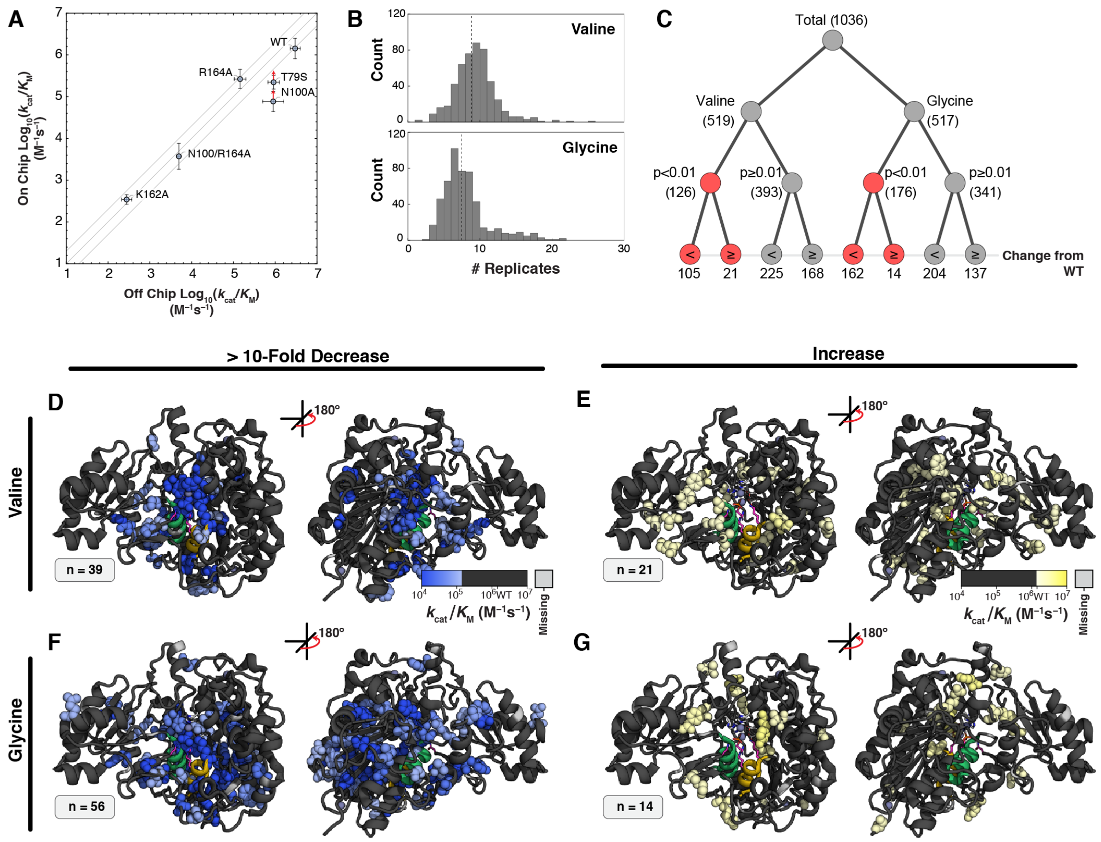

**Fig. S22.** Summary of Val and Gly library mutant cMUP $k_{\mathrm{cat}}/K_{\mathrm{M}}$ values measured on chip. (A) Comparison of on- and off-chip cMUP $k_{\mathrm{cat}}/K_{\mathrm{M}}$ measurements for WT PafA and 5 active site mutants. Red arrows denote limits. (B) Histograms showing number of experimental replicates for all valine (top) and glycine (bottom) mutants. (C) Binary tree summarizing fraction of mutations with changes in activity from WT ($p$<0.01) for valine and glycine mutant libraries. (D) Measured cMUP $k_{\mathrm{cat}}/K_{\mathrm{M}}$ values for valine substitutions with decreases of at least ten-fold in activity relative to WT projected on the PafA structure (front and rear views). (E) Measured cMUP $k_{\mathrm{cat}}/K_{\mathrm{M}}$ values for valine substitutions with increases in activity relative to WT ($p$<0.01) projected on the PafA structure (front and rear views). (F) Measured cMUP $k_{\mathrm{cat}}/K_{\mathrm{M}}$ values for glycine substitutions with decreases of at least ten-fold in activity relative to WT projected on the PafA structure (front and rear views). (G) Measured cMUP $k_{\mathrm{cat}}/K_{\mathrm{M}}$ values for glycine substitutions with increases in activity relative to WT ($p$<0.01) projected on the PafA structure (front and rear views).

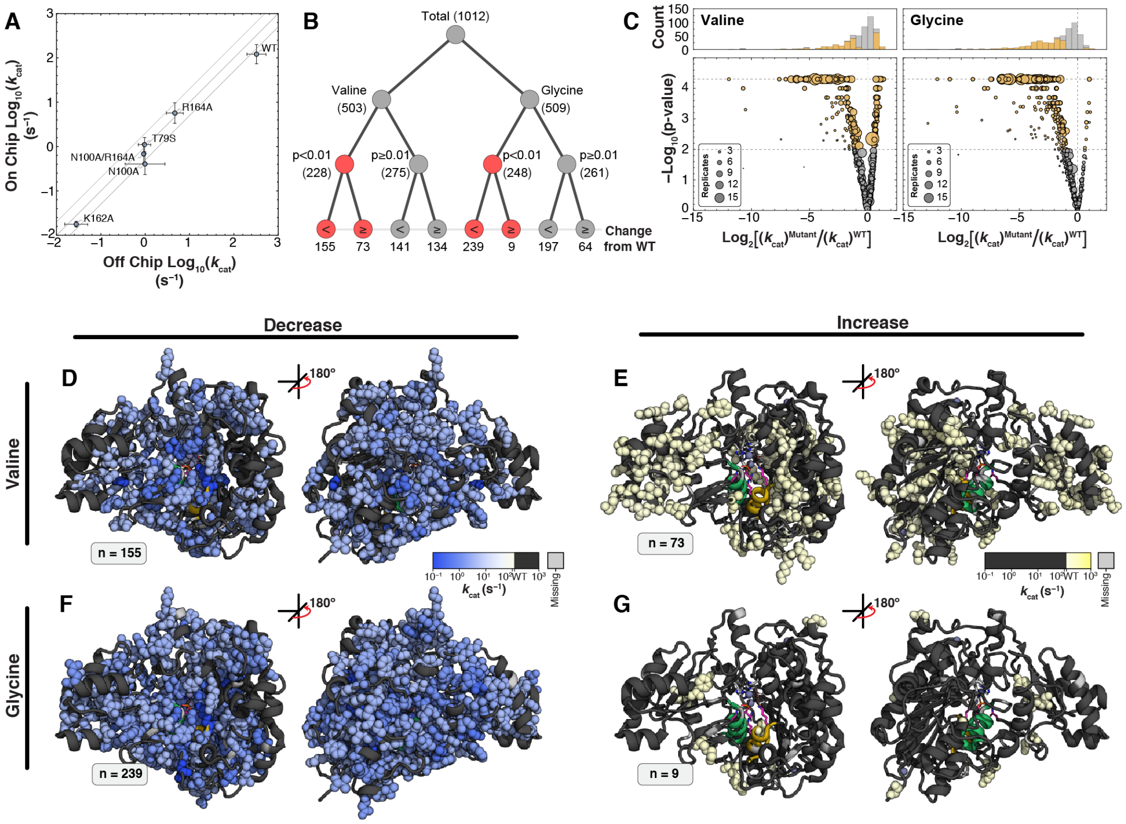

**Fig. S23.** Summary of on-chip measured cMUP $k_{\mathrm{cat}}$ values for Val and Gly library mutants. (A) Comparison of on- and off-chip cMUP $k_{\mathrm{cat}}$ measurements for WT PafA and 5 active site mutants. (B) Binary tree diagram summarizing fraction of mutations with changes in activity from WT ($p$<0.01) for valine and glycine mutant libraries. (C) Volcano plots showing $p$-value (as calculated from bootstrap analysis) as a function of measured effect size for valine (left) and glycine (right) mutants; mutants with differences in cMUP $k_{\mathrm{cat}}$ are shown in gold ($p$<0.01). (D) Measured cMUP $k_{\mathrm{cat}}$ values for valine substitutions with decreases in activity relative to WT ($p$<0.01) projected on the PafA structure (front and rear views). (E) Measured cMUP $k_{\mathrm{cat}}$ values for valine substitutions with increases in activity relative to WT ($p$<0.01) projected on the PafA structure (front and rear views). (F) Measured cMUP $k_{\mathrm{cat}}$ values for glycine substitutions with decreases in activity relative to WT ($p$<0.01) projected on the PafA structure (front and rear views). (G) Measured cMUP $k_{\mathrm{cat}}$ values for glycine substitutions with increases in activity relative to WT ($p$<0.01) projected on the PafA structure (front and rear views).

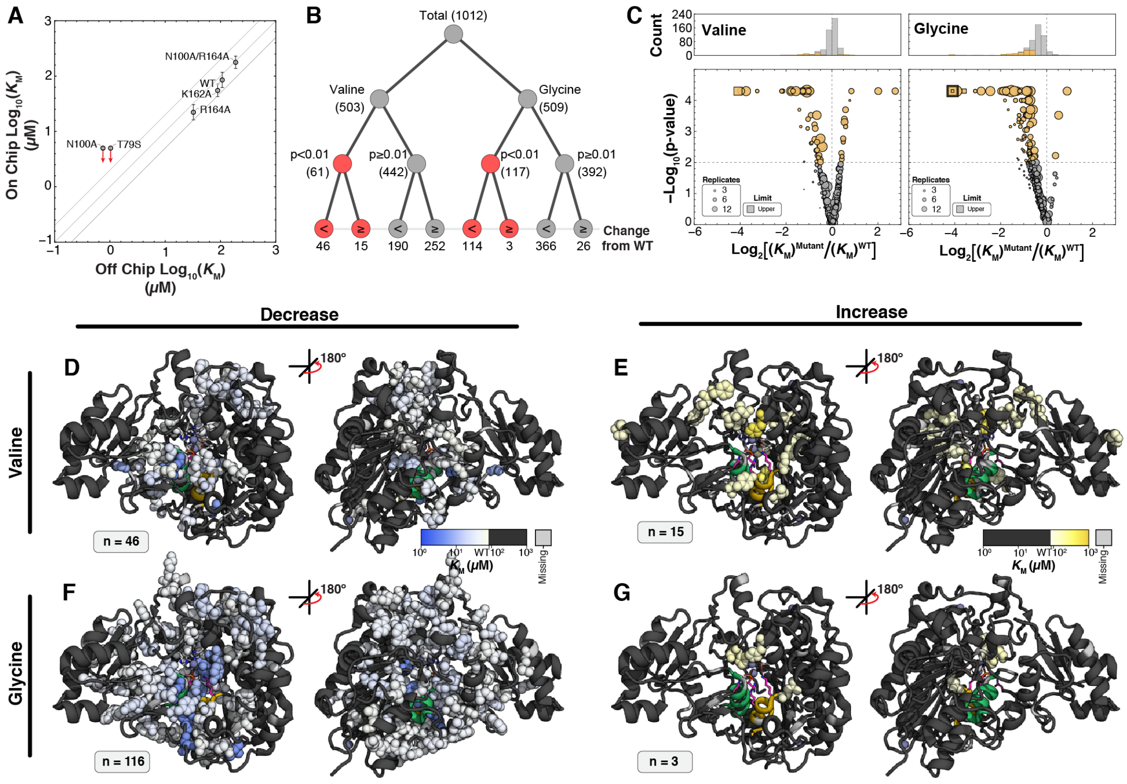

**Fig. S24.** Summary of on-chip measured cMUP $K_{\mathrm{M}}$ values for Val and Gly library mutants. (A) Comparison of on- and off-chip cMUP $K_{\mathrm{M}}$ measurements for WT PafA and 5 active site mutants. Red arrows denote limits. (B) Binary tree diagram summarizing fraction of mutations with changes in activity from WT ($p<0.01$) for valine and glycine mutant libraries. (C) Volcano plots showing $p$-value (as calculated from bootstrap analysis) as a function of measured effect size for valine (left) and glycine (right) mutants; mutants with differences in cMUP $K_{\mathrm{M}}$ are shown in gold ($p<0.01$). (D) Measured cMUP $K_{\mathrm{M}}$ values for valine substitutions with decreases in activity relative to WT ($p<0.01$) projected on the PafA structure (front and rear views). (E) Measured cMUP $K_{\mathrm{M}}$ values for valine substitutions with increases in activity relative to WT ($p<0.01$) projected on the PafA structure (front and rear views). (F) Measured cMUP $K_{\mathrm{M}}$ values for glycine substitutions with decreases in activity ($p<0.01$) relative to WT projected on the PafA structure (front and rear views). (G) Measured cMUP $K_{\mathrm{M}}$ values for glycine substitutions with increases in activity relative to WT ($p<0.01$) projected on the PafA structure (front and rear views).

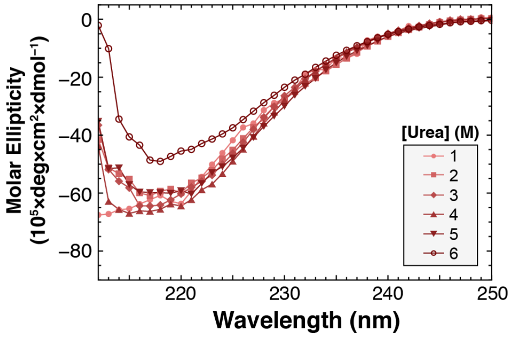

**Fig. S25.** Circular dichroism (CD) spectra of WT PafA over-expressed in *E. coli*, purified, and subjected to 2 weeks of incubation with urea.

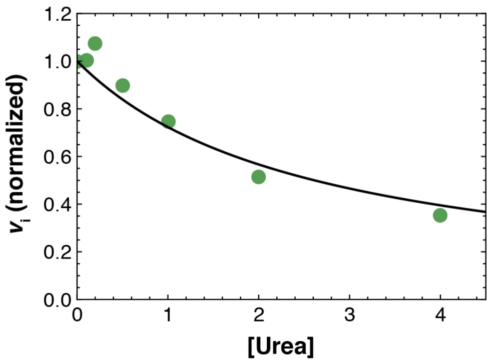

**Fig. S26.** Apparent competitive inhibition of PafA by urea. Initial rates of WT PafA hydrolysis (500 µM MepNPP) as a function of added urea concentration (green), normalized to that at 0 M urea, are well fit by a model of competitive inhibition (**Eq. 18**) with a $K_{\mathrm{i}}$ of 2.6 M ($r^2 = 0.99$).

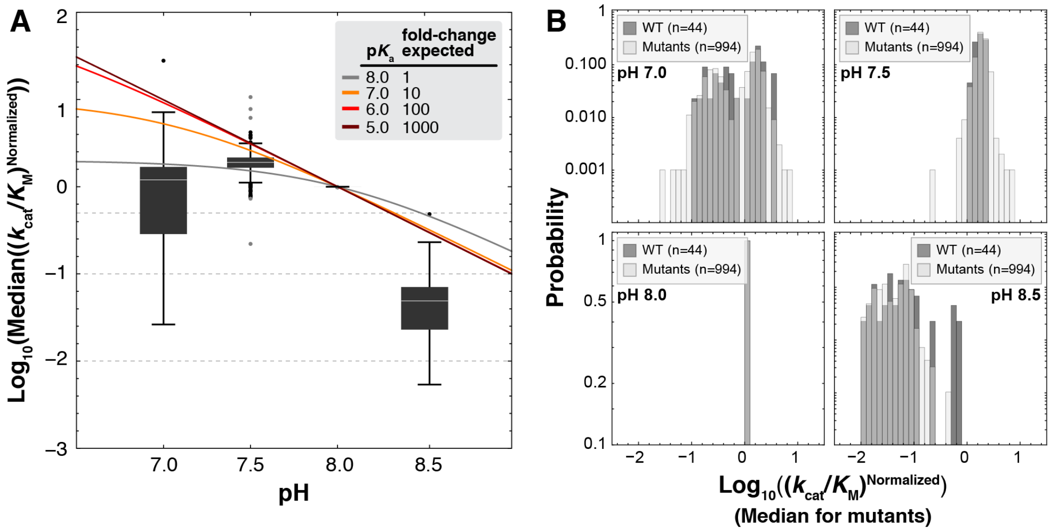

**Fig. S27.** pH-dependence of observed cMUP $k_{\mathrm{cat}}/K_{\mathrm{M}}$ for valine and glycine scan mutants. (A) Box and whisker plots of cMUP $k_{\mathrm{cat}}/K_{\mathrm{M}}$ values measured on-chip for all mutants (normalized to $k_{\mathrm{cat}}/K_{\mathrm{M}}$ at pH 8.0) at pH 7, 7.5, 8, and 8.5. Curves denote predicted $k_{\mathrm{cat}}/K_{\mathrm{M}}$ values (also normalized to the pH 8.0 values) for enzymes with p$K_{\mathrm{a}}$ values of 5.0, 6.0, 7.0, and 8.0 (the value previously reported for WT PafA) (*16*); all curves assume a single inhibitory p$K_{\mathrm{a}}$ and that the enzyme is full inactivated at high pH. The observed behavior is most consistent with mutants having an unchanged p$K_{\mathrm{a}}$ of 8.0 (B) Distributions of observed normalized $k_{\mathrm{cat}}/K_{\mathrm{M}}$ values for mutants and WT PafA at each pH.

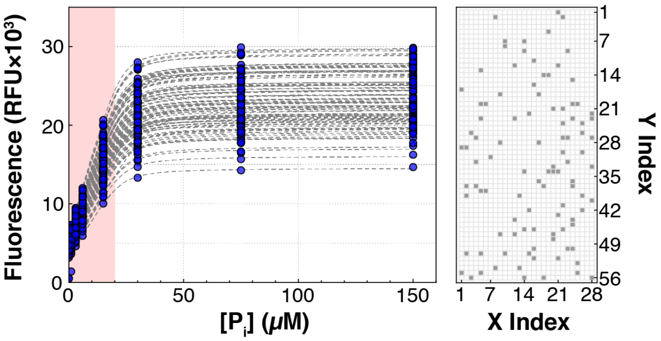

**Fig. S28.** Fluorescence standard curves relating measured Phosphate Sensor (*18*) fluorescence intensities to inorganic phosphate ($P_i$) concentrations on chip. Measured fluorescence intensities for chambers throughout the device containing 30 µM PBP in the presence of increasing amounts of $P_i$ (left, blue markers). Individual chamber intensities were fit to a single-site binding isotherm model (gray dashed lines; see **Materials and Methods**) to allow subsequent per-chamber conversion of measured intensities to $P_i$ concentrations. Initial rate fits were limited to points in the region indicated by the pink window (up to 20 µM $P_i$). Locations of randomly selected plotted chambers within the device (right).

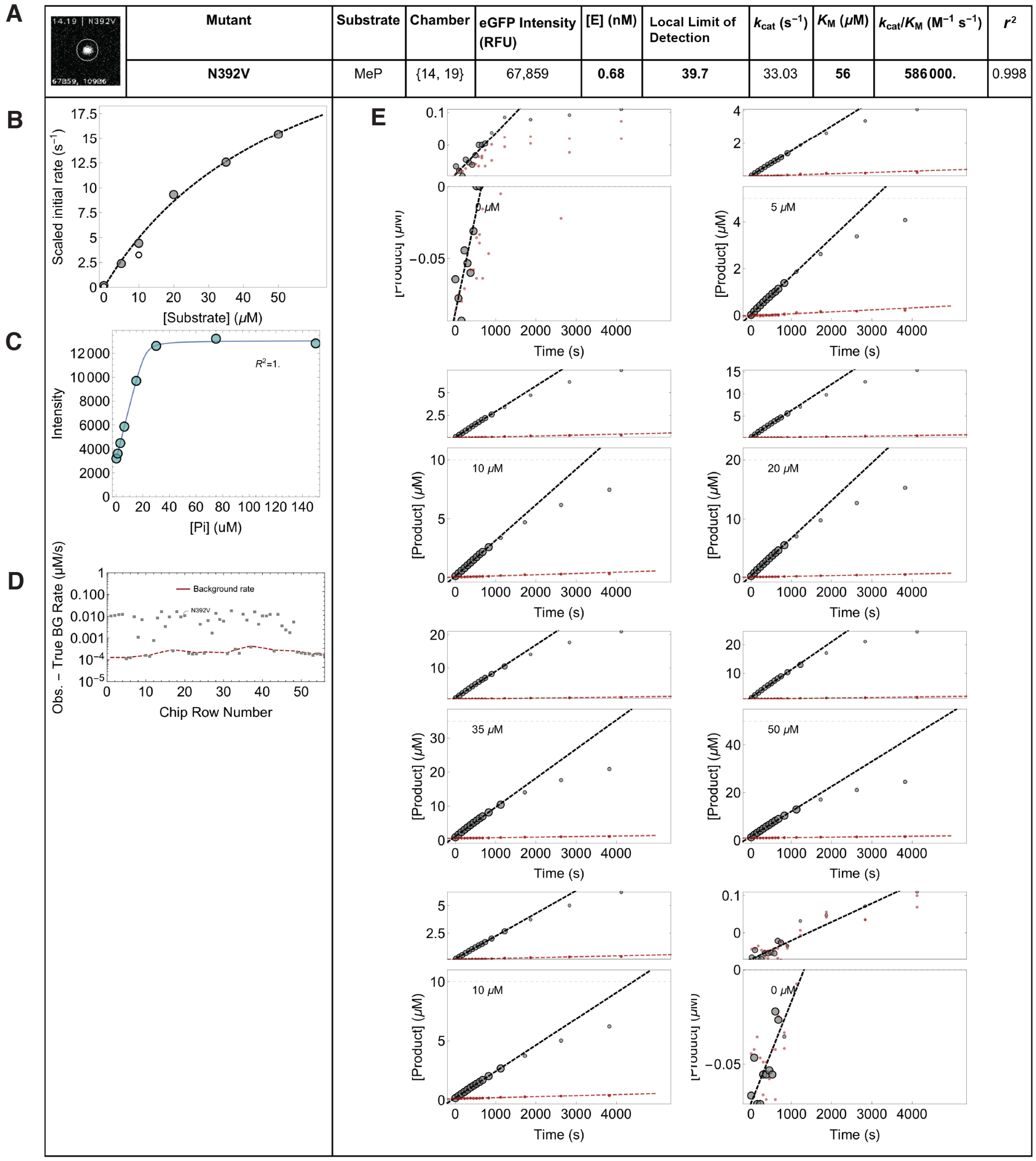

| | Mutant | Substrate | Chamber | eGFP Intensity (RFU) | [E] (nM) | Local Limit of Detection | $k_{cat}$ ($s^{-1}$) | $K_M$ (μM) | $k_{cat}/K_M$ ($M^{-1}$ $s^{-1}$) | $r^2$ |
|---|---|---|---|---|---|---|---|---|---|---|
| | **N392V** | MeP | {14, 19} | 67,859 | **0.68** | **39.7** | 33.03 | **56** | **586000.** | 0.998 |

**Fig. S29.** Example data report for a single chamber after an HT-MEK MeP hydrolysis kinetic experiment (8 individual assays). Each experiment returns 1,568 unique per-chamber reports containing: (A) Mutant identity and image of immobilized enzyme (eGFP channel), experimental parameters (substrate, chamber column and row, eGFP intensity, enzyme concentration, rate compared to the local background (LLoD), and Michaelis-Menten fit parameters ($k_{cat}$, $K_M$, $k_{cat}/K_M$, and $r^2$); (B) Michaelis-Menten fits to scaled initial rates (gray) with initial rate from replicate assay (white); (C) Per-chamber product fluorescence standard curve and fit $r^2$; (D) Comparison of fitted rates (gray points) with interpolated local lower limit of quantification (LLoD, red dashed line) within this chamber's full column (see **Materials and Methods**); (E) Full reaction progress curves (lower panels) and expanded plots (upper panels) for each substrate concentration, with fitted initial rates shown as dashed gray lines. Large gray points were used to determine the linear fit, small gray points were measured but outside the linear region and not used to fit initial rates. Red points and lines denote the progress curves and linear fits from the nearest upstream and downstream background chambers used to determine the local background.

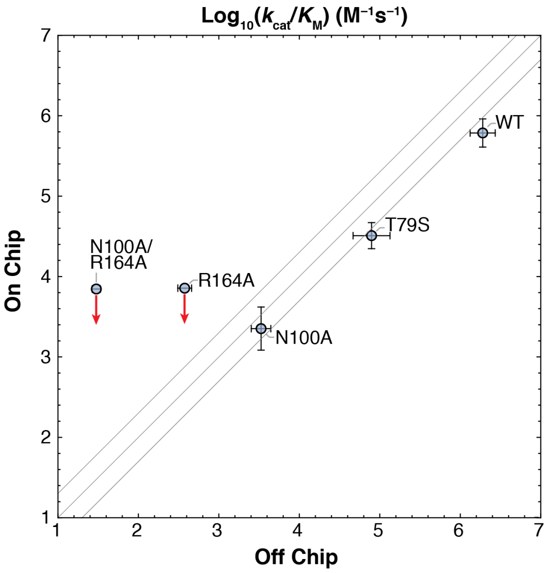

**Fig. S30.** Comparison of on-chip and off-chip MeP $k_{\mathrm{cat}}/K_{\mathrm{M}}$ values for PafA active site mutants. On-chip median values were calculated from measurements of active site variant controls incorporated in the valine and glycine scanning mutant assays. Upper limits for on-chip measurements are denoted by red arrows. Off-chip values are means of two or more replicates and error bars denote standard deviations.

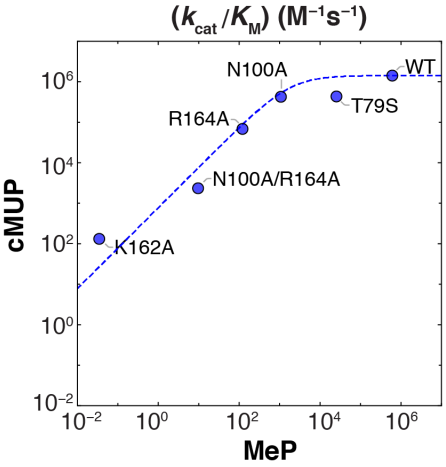

**Fig. S31.** Off-chip measured cMUP $k_{\mathrm{cat}}/K_{\mathrm{M}}$ and MeP $k_{\mathrm{cat}}/K_{\mathrm{M}}$ values for WT PafA and 5 active site mutants with associated non-linear fit to the relative reactivity relationship predicted by their kinetic rate equations (**Eq. 31**). Activity measurements were made using purified enzyme expressed in *E. coli*; the fit to **Eq. 31** yielded $f_{\mathrm{r}}$ = 776±126 (median±SEM), which specifies the relative turnover rates of the substrates when the hydrolysis step is rate limiting for both (as described in **Supplementary Text S2**).

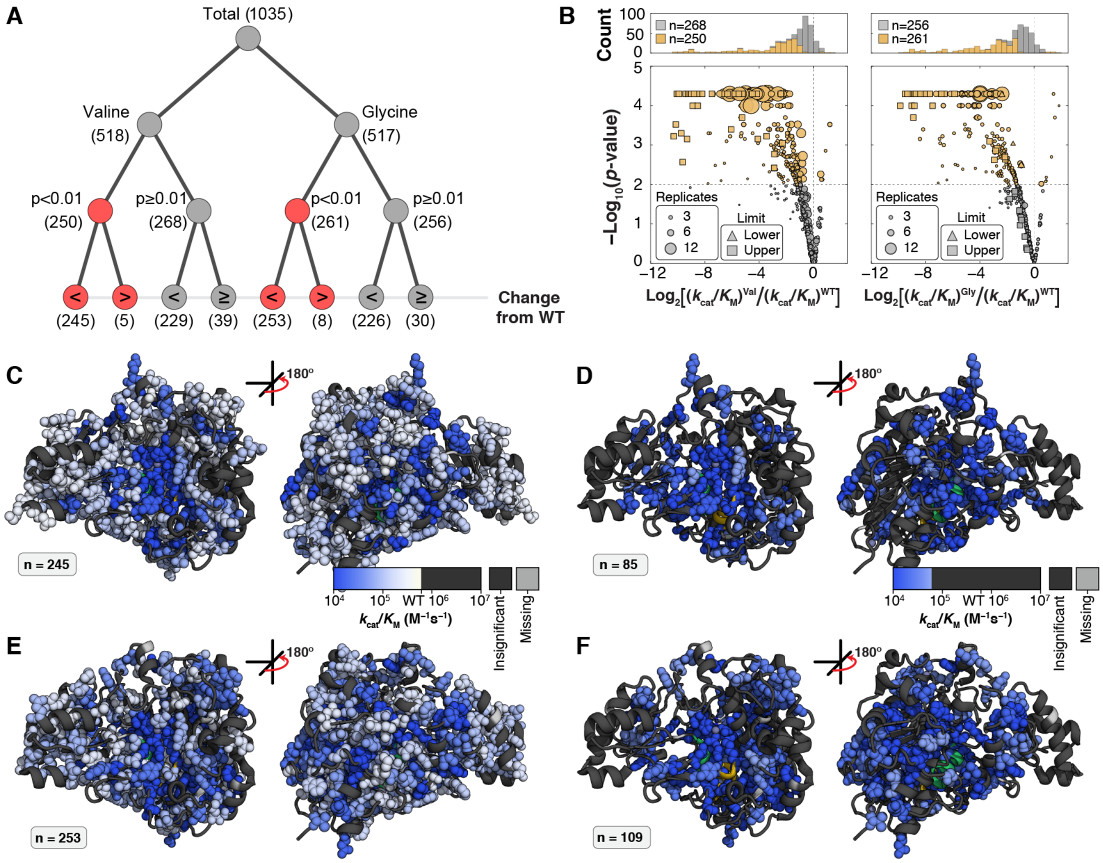

**Fig. S32.** Summary of on-chip measured MeP $k_{\mathrm{cat}}/K_{\mathrm{M}}$ values for glycine and valine library mutants. (A) Tree diagram describing the number of mutants with MeP $k_{\mathrm{cat}}/K_{\mathrm{M}}$ effects ($p<0.01$). (B) Volcano plots showing fold-effects, $p$-value, and number of replicates for the valine scanning mutants (left) and glycine scanning mutants (right). Effects with $p<0.01$ are shown in gold. (C–F) Structural representations of positions with decreases in MeP $k_{\mathrm{kcat}}/K_{\mathrm{M}}$ upon mutation to valine (C, less than WT and $p<0.01$; D, 10-fold or greater decrease from WT and $p<0.01$) or glycine (E, less than WT and $p<0.01$; F, 10-fold or greater decrease from WT and $p<0.01$). “Nucleophile” and “Monoester” helices are shown in green and gold, respectively, but are largely obscured by spheres indicating positions with effects.

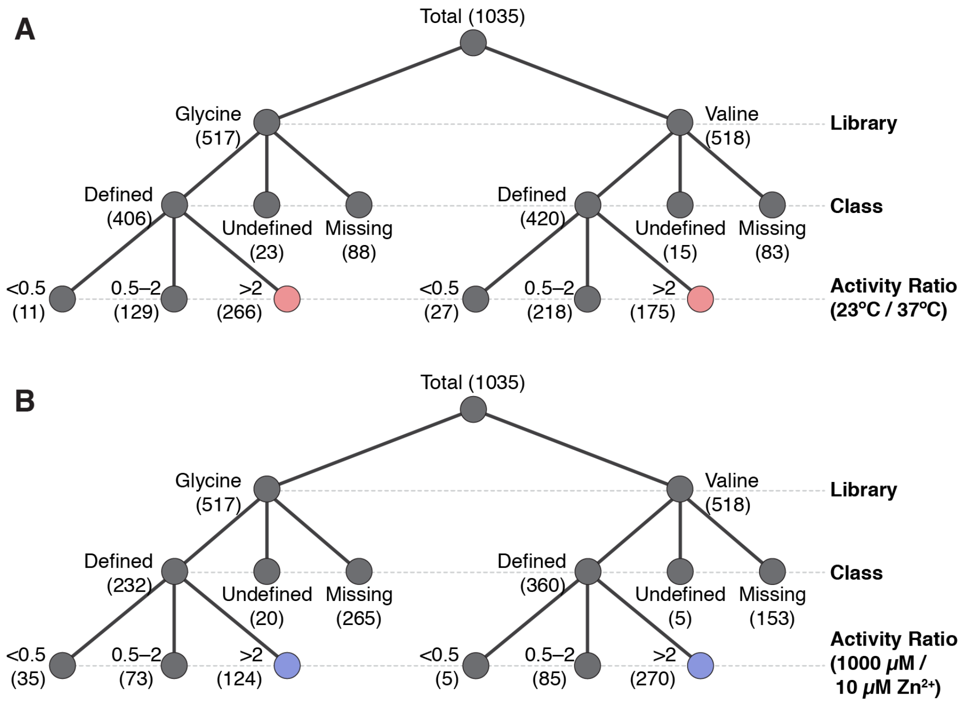

**Fig. S33.** Number of valine and glycine mutants with changes in cMUP $k_{\mathrm{cat}}/K_{\mathrm{M}}$ upon changes in expression (but not assay) conditions. Trees indicating counts of (A) temperature- and (B) Zn-dependent expression effects (T-Effect and Zn-Effect; **Eq. 9** and **10**, respectively) on $k_{\mathrm{cat}}/K_{\mathrm{M}}$ for Val and Gly library mutants. Mutants for which the activity ratio may be less than or greater than unity (as they are limits in one or both direction) are considered "Undefined" and mutants lacking an estimate under either or both expression conditions are "Missing."

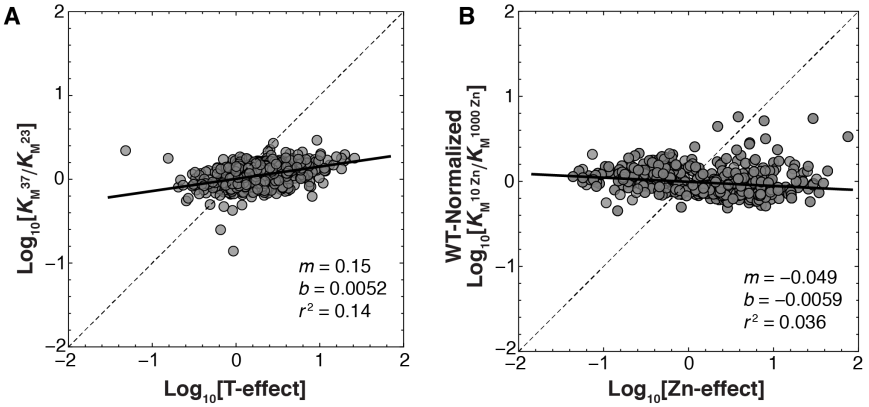

**Fig. S34.** Measured $K_{\mathrm{M}}$ values as a function of temperature- and zinc-effects (T-Effect and Zn-Effect; **Eq. 9** and **10**, respectively) during expression. (A) $\mathrm{Log}_{10}$-transformed ratios of $K_{\mathrm{M}}$ values at 37°C and 23°C (grey markers) as a function of $\log_{10}$-transformed T-effect (**Eq. 9**). (B) WT-normalized $\log_{10}$-transformed ratio of $K_{\mathrm{M}}$ values at 10 µM and 1000 µM $ZnCl_2$ (grey markers) as a function of $\log_{10}$-transformed Zn-effect (**Eq. 10**). Dashed grey lines denote a 1:1 relationship; solid black lines correspond to linear regressions with slopes, intercepts, and $r^2$ values as indicated.

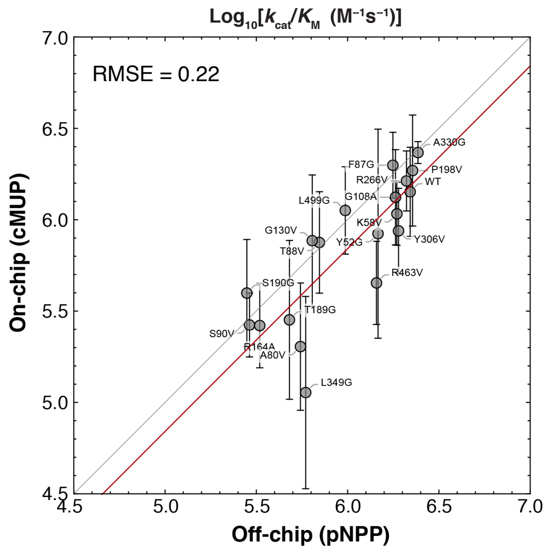

**Fig. S35.** Comparison of measured on- and off-chip measured aryl monoester (cMUP and pNPP, respectively) $k_{\mathrm{cat}}/K_{\mathrm{M}}$ values for WT PafA and 18 library mutants (grey markers, see **Table S3** and **Materials and Methods**). The red line corresponds to a linear least-squares regression with a constrained slope of unity using $\log_{10}$-transformed data (fit intercept $= -0.16$); y-error bars indicate the standard deviations of on-chip measurements. An RMSE of 0.22 in $\log_{10}$-transformed space corresponds to a 1.7-fold effect.

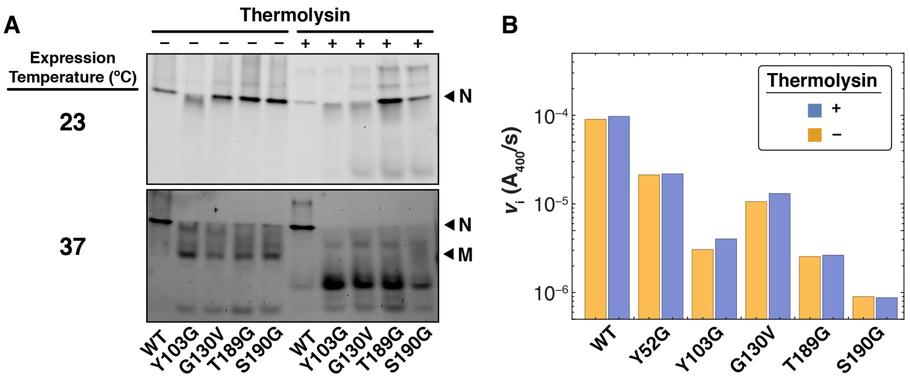

**Fig. S36.** Native gel electrophoresis and turnover kinetics support the existence of a misfolded state among library mutants expressed *in vitro* off-chip. (A) Native gel electrophoresis of a subset of mutants expressed at either 23°C (top) or 37°C (bottom) in the presence (+) or absence (–) of thermolysin. When expressed at 23°C (left), all constructs appeared as a single band with the same electrophoretic mobility as that of WT (the native "N" state). By contrast, expression at 37°C resulted in the appearance of common higher-mobility species among the mutants, but not the WT (the putatively misfolded "M" state). The addition of thermolysin (right) led to near-complete degradation of the M state (37°C panel) but not the N state (23°C panel). (B) Measured initial rates of pNPP turnover for WT PafA and five library mutants expressed off-chip at 37°C, assayed with and without prior thermolysin treatment. Thermolysin treatment did not significantly alter the activities, suggesting that the protease-susceptible M state observed in (A) was catalytically inactive and did not contribute to the observed rate.

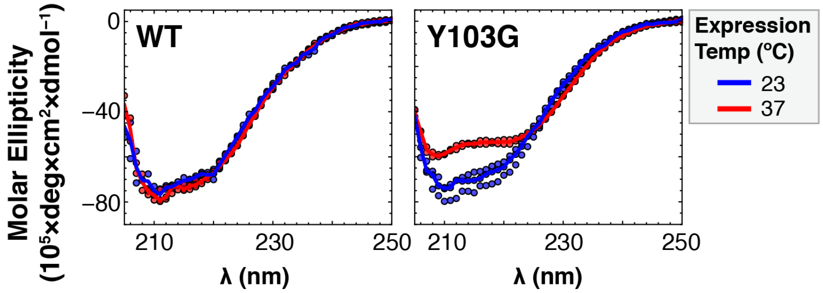

**Fig. S37.** CD spectra for WT PafA and the temperature-sensitive mutant Y103G expressed in *E. coli* at low (23°C, blue markers) and high (37°C, red markers) temperature. Solid lines are the medians of 2 or 3 replicate spectra obtained in different buffers (see **Materials and Methods**).

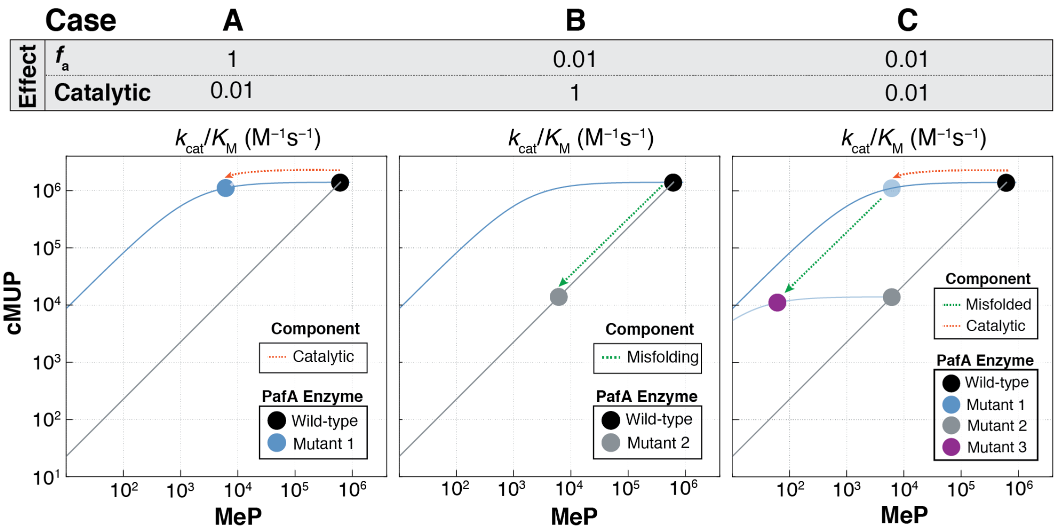

**Fig. S38.** Three examples illustrating how observed mutational effects on $k_{cat}/K_M$ can arise from combinations of catalytic and misfolding effects. (A) Case 1: Mutation has a 100-fold effect on the chemical step of catalysis ($k_{cat}/K_M$) (red dashed arrow) from wild-type (black point). The effect is much greater for MeP hydrolysis $(k_{cat}/K_M)^{MeP}$ as hydrolysis remains diffusion-limited for cMUP $(k_{cat}/K_M)^{cMUP}$ (blue point and curve). (B) Case 2: Mutation results in 99% of the protein misfolding, but the folded 1% has wild-type $(k_{cat}/K_M)^{MeP}$ and $(k_{cat}/K_M)^{cMUP}$, resulting in an observed 100-fold drop in $k_{cat}/K_M$ for both substrates (gray point, and green dashed arrow). (C) Case 3: Mutation results in 99% of the protein misfolding, as in (B) (green dashed arrow), but with an additional 100-fold effect on the chemical step (red dashed arrow). The observed rates (purple point) result from a combination of misfolding (gray point) and catalytic effects (blue point). The observed $(k_{cat}/K_M)^{MeP}$ is therefore decreased $10^4$-fold from the wild-type value, while the observed $(k_{cat}/K_M)^{cMUP}$ is only decreased $10^2$-fold due to the misfolding effect, as hydrolysis remains diffusion-limited for cMUP.

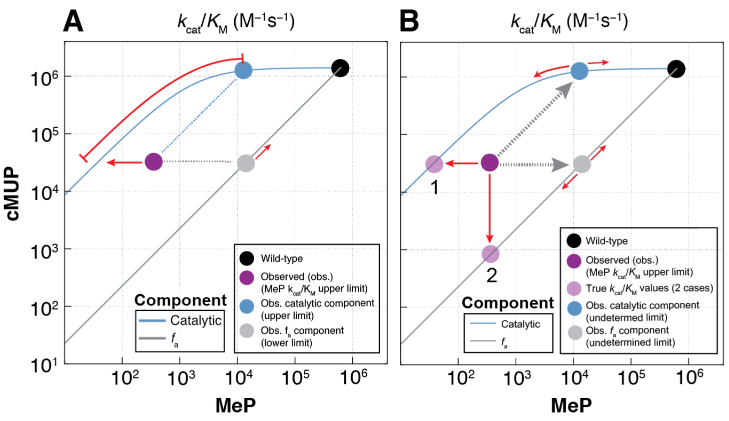

**Fig. S39.** Effect of observed MeP and cMUP $k_{\text{cat}}/K_{\text{M}}$ upper limits on calculated catalytic and fraction active ($f_{\text{a}}$) effects. (A) Effect of a MeP $k_{\text{cat}}/K_{\text{M}}$ upper limit. As determination of catalytic and $f_{\text{a}}$ effects depends on both cMUP and MeP $k_{\text{cat}}/K_{\text{M}}$ , variants for which the observed (obs.) MeP $k_{\text{cat}}/K_{\text{M}}$ is an upper limit (dark purple point) will result in the intrinsic catalytic $k_{\text{cat}}/K_{\text{M}}$ (blue point) and $f_{\text{a}}$ (gray point) effects being upper and lower limits, respectively. The gray arrows illustrate the determination of catalytic and $f_{\text{a}}$ components from the observed point, and the red arrows denote the direction of the limits. It is therefore possible to obtain upper limits for catalytic effects and estimate their significance; however, $p$-values of $f_{\text{a}}$ effects cannot be determined. (B) Effect of both MeP and cMUP $k_{\text{cat}}/K_{\text{M}}$ upper limits on calculated catalytic and $f_{\text{a}}$ components. When both the observed MeP and cMUP $k_{\text{cat}}/K_{\text{M}}$ are upper limits (red arrows and dark purple point), the true effect can be fully catalytic (case 1, light purple), fully due to $f_{\text{a}}$ (case 2, light purple), or any combination of the two components. Therefore, neither the catalytic and $f_{\text{a}}$ effects of the variants nor their $p$-values can be determined.

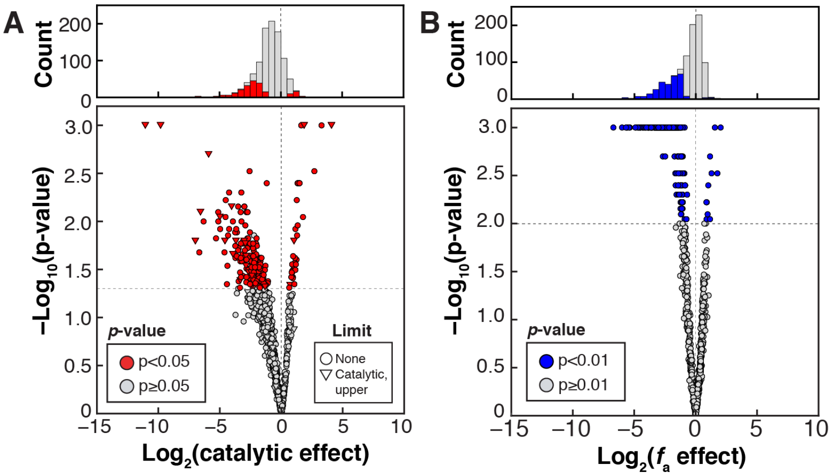

**Fig. S40.** Volcano plots of (A) catalytic effects and (B) fraction active ($f_a$) effects for all measurable variants in the valine and glycine libraries. Upper limits of $p$-values for upper $k_{cat}/K_M^{chem}$ limits (triangles) can only be inferred for catalytic effects (see **Materials and Methods** and **Fig. S39**); therefore, limits are included in (A) but not (B). For both, vertical grid lines denote no change from WT, while horizontal grid lines denote the $p$-value thresholds of 0.05 and 0.01, respectively.

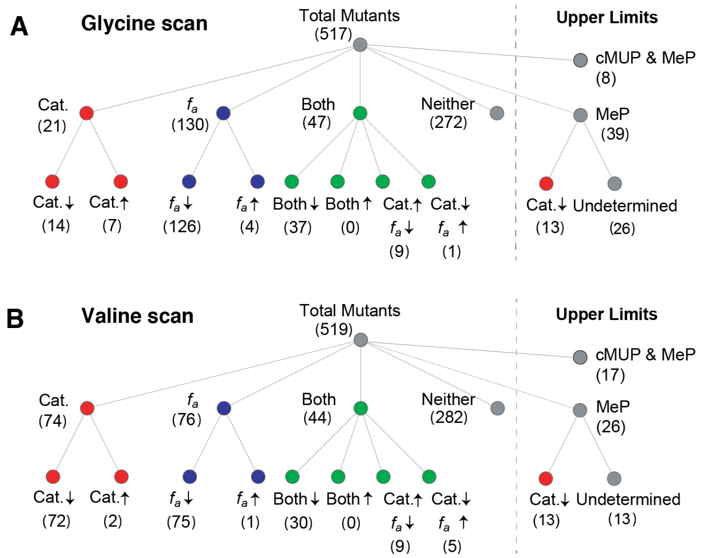

| Effect | *p*-value Threshold | Direction | Total |
|---|---|---|---|
| Catalytic | $p$<0.05 | ↓ | 65 |
| | | ↑ | 16 |
| $f_a$ | $p$<0.01 | ↓ | 172 |
| | | ↑ | 5 |

| Effect | *p*-value Threshold | Direction | Total |
|---|---|---|---|
| Catalytic | $p$<0.05 | ↓ | 120 |
| | | ↑ | 11 |
| $f_a$ | $p$<0.01 | ↓ | 114 |
| | | ↑ | 6 |

**Fig. S41.** Counts of variants within the (A) glycine and (B) valine scan libraries with catalytic (Cat.) effects, fraction active ($f_a$) effects, both catalytic and fraction active effects, and variants without significant effects. Mutants with catalytic effects ($p<0.05$) are colored red, mutants with $f_a$ effects only ($p<0.01$) are colored blue, and mutants with both catalytic and $f_a$ effects are colored green, with the number of mutants within each set given in brackets. Arrows denote the direction of effects relative to WT. $P$-values for limits can only be inferred for catalytic effects, as described in **Materials and Methods** (and see **Fig. S39**). Total (cumulative) numbers of catalytic and $f_a$ effects (including the mutants with both effects and limits) are given in the tables to the right.

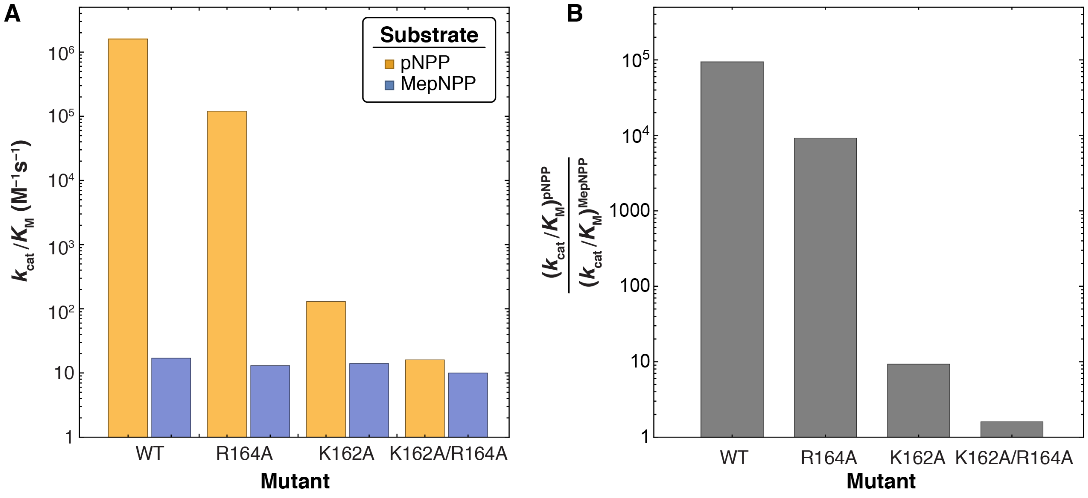

**Fig. S42.** K162 and R164 determine specificity for monoester hydrolysis. (A) Off-chip measurements of $k_{\text{cat}}/K_{\text{M}}$ for pNPP (monoester) and MepNPP (diester) hydrolysis by WT PafA and K162/R164 mutants. (B) Ratio of $k_{\text{cat}}/K_{\text{M}}$ for pNPP and MepNPP hydrolysis ("specificity") for WT PafA and K162/R164 mutants. Data in both panels reproduced from (*16*).

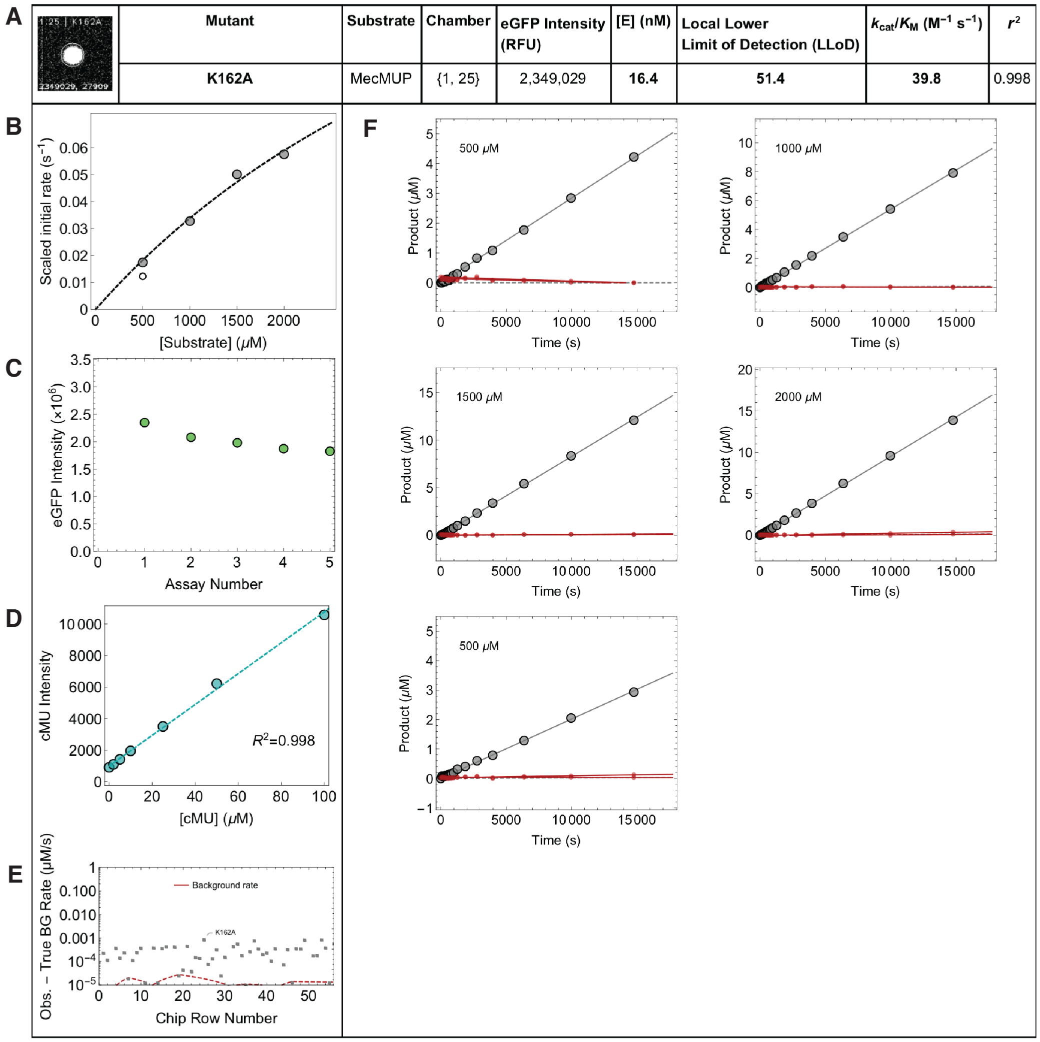

**Fig. S43.** Example data report for a single chamber after an HT-MEK experiment measuring MecMUP hydrolysis (5 individual assays). Each experiment returns 1,568 unique per-chamber reports containing: (A) Identity of the mutant in each chamber, with image of immobilized enzyme (eGFP channel) and experimental parameters including substrate, chamber column and row, eGFP intensity, enzyme concentration, the rate compared to the local background (LLoD, see (E) and **Materials and Methods**), $k_{\mathrm{cat}}/K_{\mathrm{M}}$, and $r^2$; (B) Michaelis-Menten fits to scaled initial rates (gray) with initial rate from replicate assay (white); (C) Measured eGFP intensity for each assay; (D) Per-chamber product fluorescence standard curve and fit $r^2$; (E) Comparison of fitted rates (gray points) with the interpolated local lower limit of quantification (LLoD, red dashed line) within this chamber's full column (see **Materials and Methods**); (F) Full reaction progress curves (lower panels) and expanded plots (upper panels) for each substrate concentration, with fitted initial rates shown as dashed gray lines. Large gray points were used to determine the linear fit, small gray points were measured but outside the linear region and not used to fit initial rates. Red points and lines denote the progress curves and linear fits from the nearest upstream and downstream background chambers used to determine the local background.

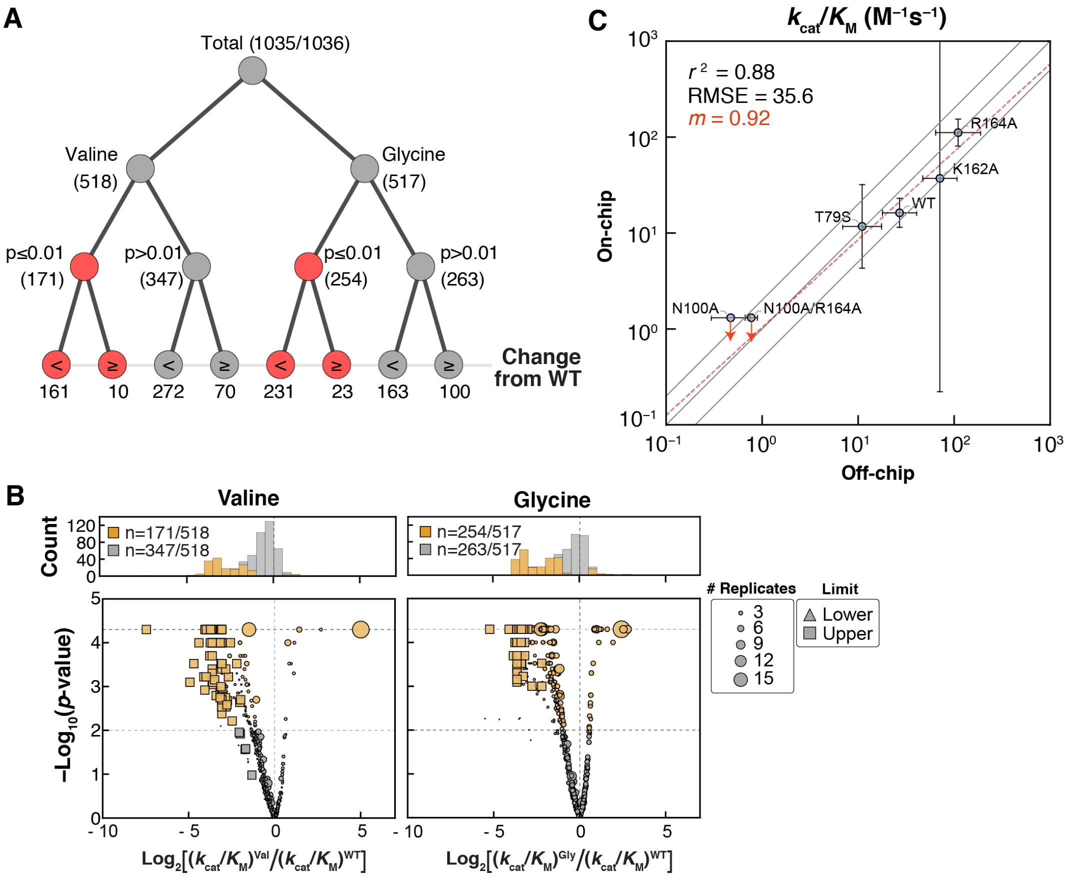

**Fig. S44.** Summary of measured $k_{cat}/K_M$ for MecMUP across all mutants. (A) Tree diagram summarizing fraction of mutations with changes in activity from WT ($p<0.01$) for valine and glycine mutant libraries. The $p$-values were determined from bootstrap hypothesis testing (see **Materials and Methods**). (B) Volcano plots showing effect size vs. $p$-value (as calculated from bootstrap hypothesis testing) for valine (left) and glycine (right) mutants; mutants with MecUP $k_{cat}/K_M$ values of $p<0.01$ are colored gold. (C) Comparison of on- and off-chip MecMUP $k_{cat}/K_M$ measurements for WT PafA and 5 additional mutants. The $r^2$ and RMSE are given with respect to a 1:1 line, and $m$ corresponds to the slope of the best fit line (red).

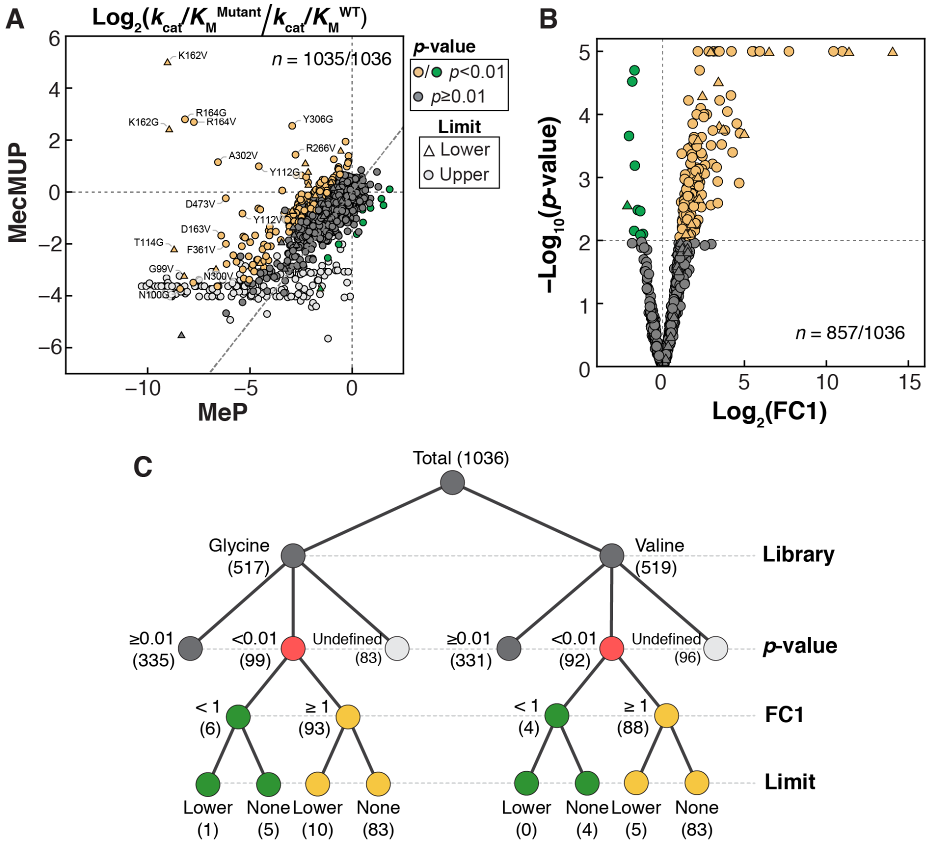

**Fig. S45.** Summary of Functional Component 1 (FC1) effects. (A) Mutational effects on $k_{\mathrm{cat}}/K_{\mathrm{M}}$ of MecMUP and MeP hydrolysis. Mutants with FC1>1 are colored yellow and those with FC1<1 are colored green, if $p<0.01$, determined from bootstrap hypothesis testing (see **Materials and Methods**). Upper FC1 limits (light gray points) arise for mutants whose MecMUP $k_{\mathrm{cat}}/K_{\mathrm{M}}$ was below the dynamic range of the assay; lower FC1 limits are due to MeP $k_{\mathrm{cat}}/K_{\mathrm{M}}$ measurements below the dynamic range (see **Materials and Methods**). (B) Volcano plot of FC1 effects. Upper limits are not shown in the figure as their FC1 effects cannot be determined ("Undefined" $p$-value). (C) Tree diagram summarizing fraction of mutations with FC1 effects ($p<0.01$) and without FC1 effects, including upper limits ("Undefined" $p$-value) and lower limits as described in **Materials and Methods**.

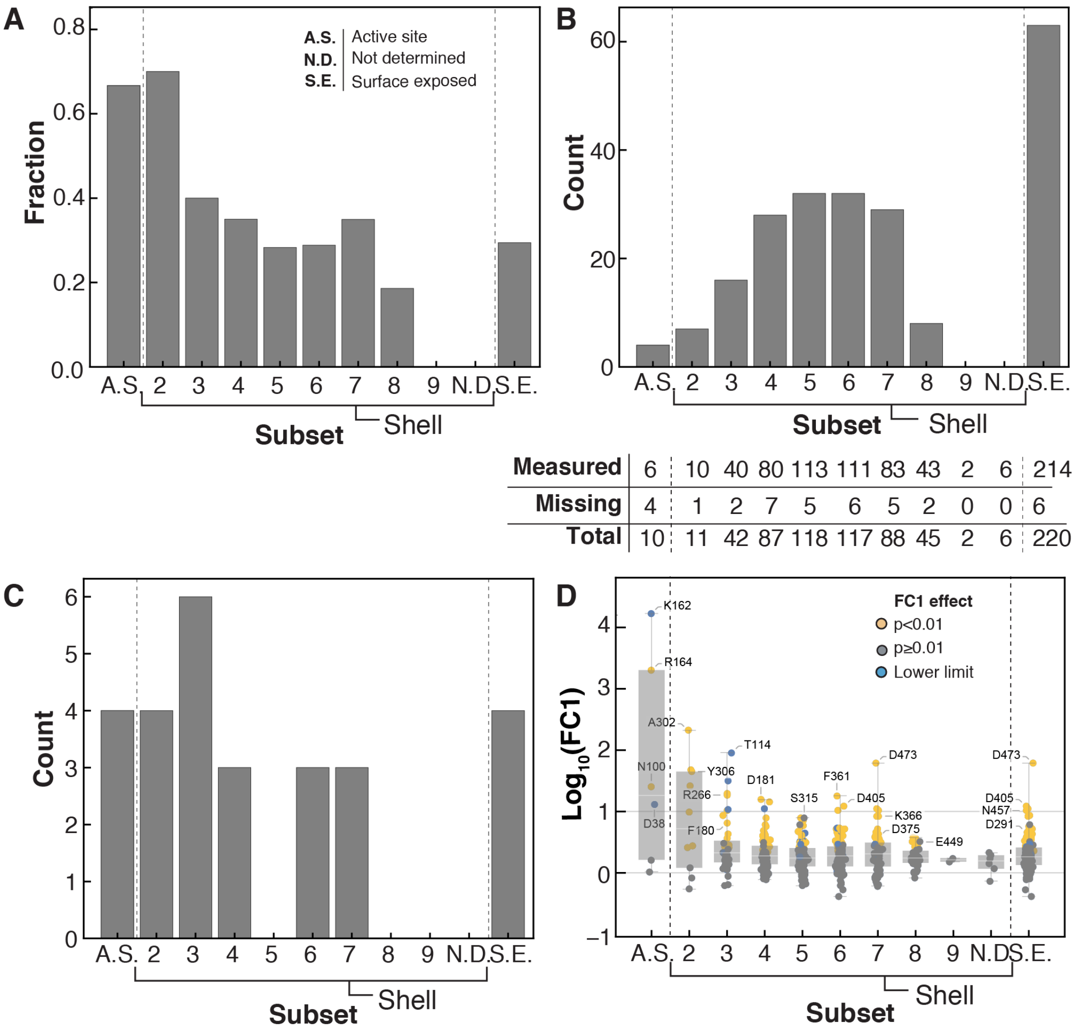

| | | | | | | | | | | | |
|---|---|---|---|---|---|---|---|---|---|---|---|
| **Measured** | 6 | 10 | 40 | 80 | 113 | 111 | 83 | 43 | 2 | 6 | 214 |
| **Missing** | 4 | 1 | 2 | 7 | 5 | 6 | 5 | 2 | 0 | 0 | 6 |
| **Total** | 10 | 11 | 42 | 87 | 118 | 117 | 88 | 45 | 2 | 6 | 220 |

**Fig. S46.** Distributions of FC1 effects within the active site, each K162/R164 interaction shell, and the enzyme surface. (A) Fraction of measurable residues with FC1 effects within the active site (A.S.), each K162/R164 interaction shell, and for surface exposed (S.E.) residues (see **Materials and Methods**). (B) Number of residues with FC1 effects within the A.S., each K162/R164 interaction shell, and for surface-exposed residues. The table below gives the number of residues for which FC1 effects were not determined and the total number of residues within each subset. (C) Number of residues with >10-fold FC1 effects within the A.S., each K162/R164 interaction shell, and for surface-exposed residues. (D) Magnitude of FC1 effects within each shell for all residues that are either measurable or for which limits could be inferred. The horizontal lines denote 10-fold and WT-like FC1 effects, respectively.

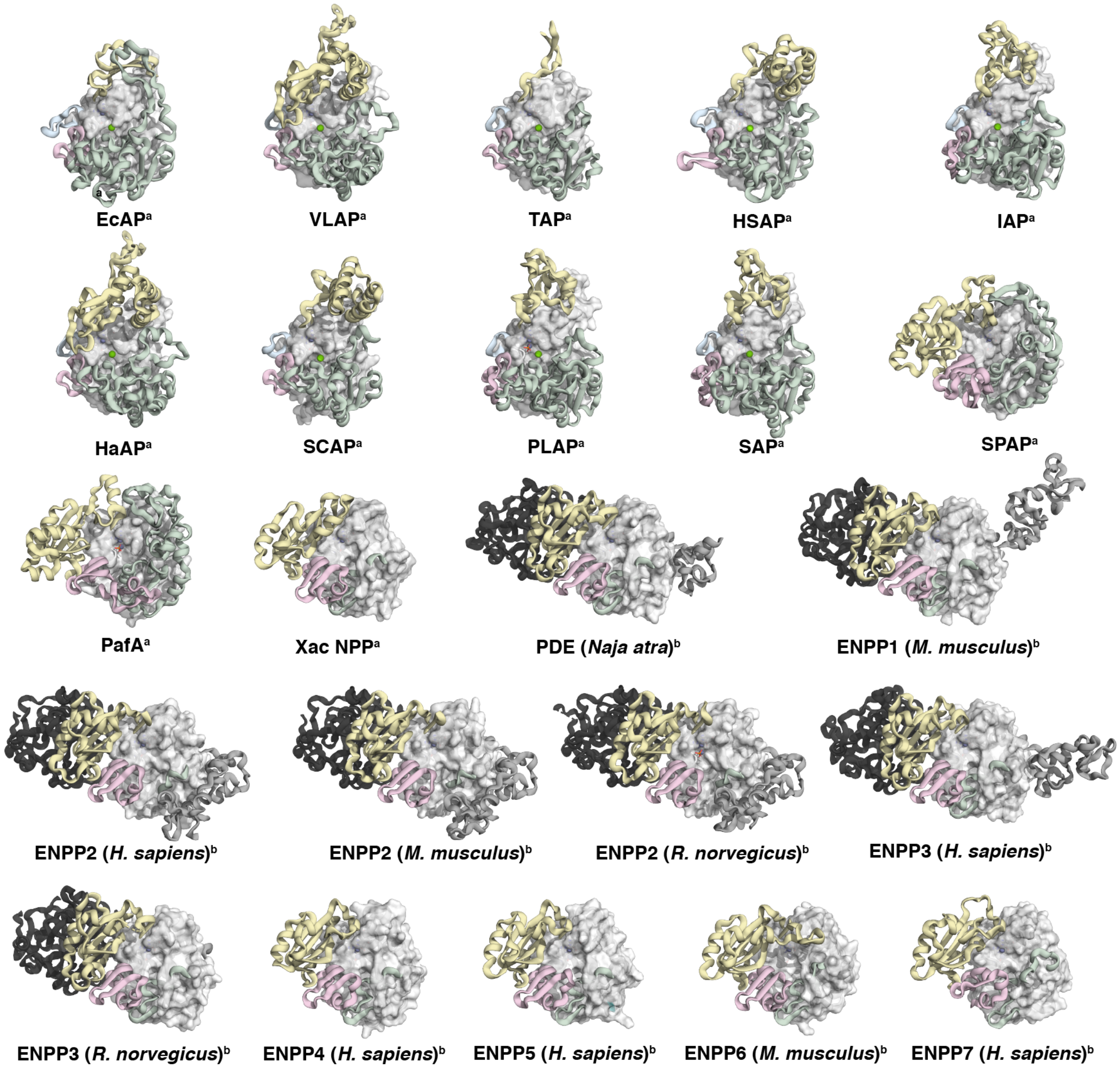

**Cognate reactivity**

[a]Monoesterase

[b]Diesterase

**Structural Domains**

| Color | Name | PafA Range |
|---|---|---|
| (dark gray) | Auxiliary N-term | N-terminus–27 |
| (light blue) | Auxiliary 1 | Not Present |
| (pink) | Auxiliary 2 | 91–137 |
| (light green) | Auxiliary 3 | 159–275 |
| (yellow) | Auxiliary 4 | 356–485 |
| (white) | Rossmann Core | Remainder |
| (black) | Auxiliary C-term | Not Present |

**Fig. S47.** Core and auxiliary domain assignments of unique structurally-characterized alkaline phosphatase (AP) superfamily phosphate monoesterases and diesterases (*cf.* **Table S10**). The Rossmann core is shown as a gray surface and the auxiliary domains are shown as cartoons and colored.

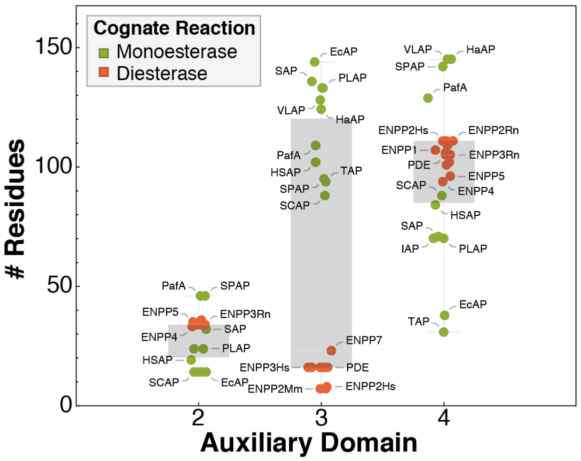

**Fig. S48.** Lengths of auxiliary domains 2–4 for AP superfamily monoesterase and diesterase orthologues.

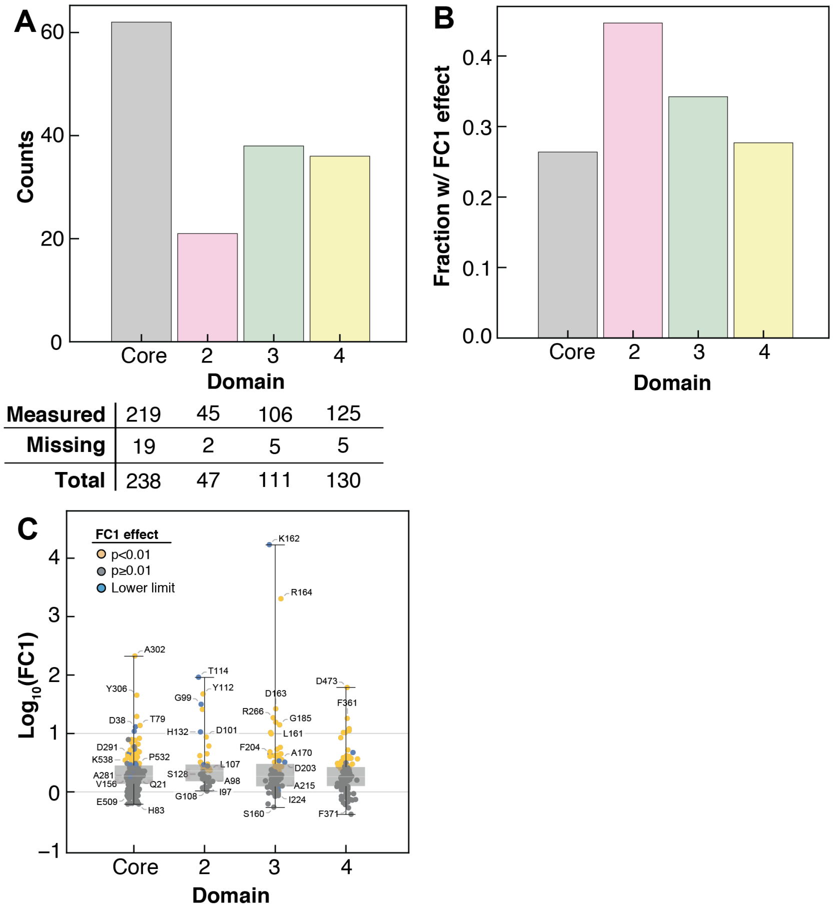

| | Core | 2 | 3 | 4 |
|---|---|---|---|---|
| **Measured** | 219 | 45 | 106 | 125 |
| **Missing** | 19 | 2 | 5 | 5 |
| **Total** | 238 | 47 | 111 | 130 |

**Fig. S49.** Distributions of FC1 effects within the Rossmann fold core and auxiliary domains. (A) Number of residues with FC1 effects ($p$<0.01, Val or Gly mutant) within Rossmann core and auxiliary domains. (B) Fraction of residues with FC1 effects within Rossmann core and auxiliary domains. (C) Magnitude of FC1 effects within the Rossmann core and each auxiliary domain, for all residues either measurable or for which limits could be inferred. The horizontal lines denote 10-fold and WT-like FC1 effects, respectively.

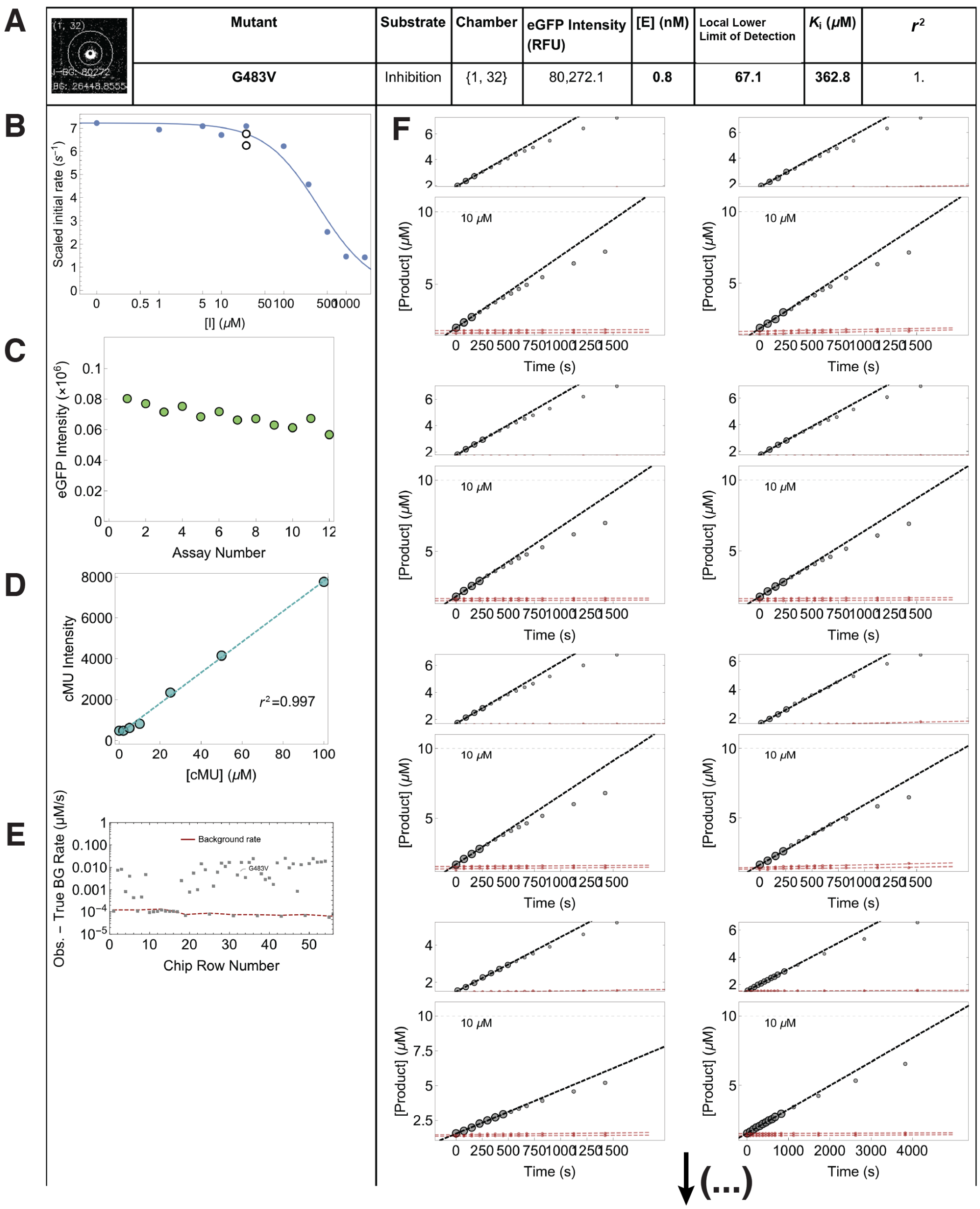

**Fig. S50.** Example data report for a single chamber after an HT-MEK experiment measuring hydrolysis of cMUP in the presence of increasing $P_i$ concentrations to quantify $K_i(P_i)$ (12 assays, with the first 6 shown in this example). Each experiment returns 1,568 unique per-chamber reports containing: (A) Mutant identity, with image of immobilized enzyme in the eGFP channel, experimental parameters (chamber column and row, eGFP intensity, enzyme concentration, the rate compared to the local background (LLoD), $K_i$ and $r^2$); (B) Competitive inhibition fits to scaled initial rates (blue) with initial rates from replicate assays (white); (C) Measured eGFP intensity for each assay; (D) Per-chamber product fluorescence standard curve and fit $r^2$; (E) Comparison of fitted rates (gray points) with the interpolated local lower limit of detection (LLoD, red dashed line) within this chamber's full column; (F) Full reaction progress curves (lower panels) and expanded plots (upper panels) for each $P_i$ concentration, with fitted initial rates shown as dashed black lines. Large gray points were used to determine the linear fit; small gray points were measured but outside the linear region and not used to fit initial rates. Red points and lines denote the progress curves and linear fits from the nearest upstream and downstream background chambers used to determine the local background.

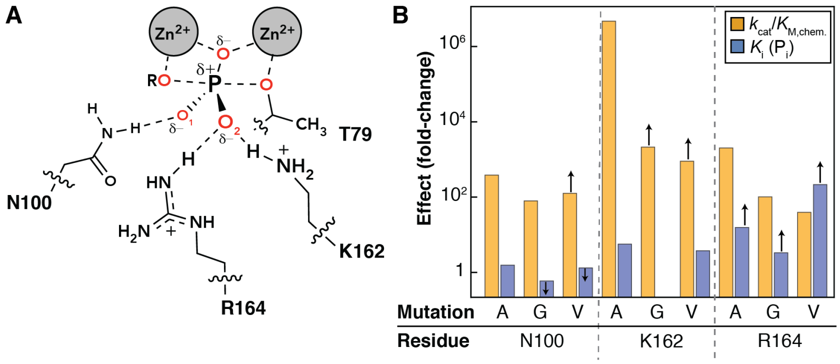

**Fig. S51.** Catalytic and $K_\mathrm{i}(\mathrm{P_i})$ effects for mutations of residues making active site O1 and O2 contacts. (A) Active site schematic of PafA showing the three active site residues making contacts to the phosphoryl O1 and O2 oxygen atoms, the T79 nucleophile, and the catalytic $Zn^{2+}$ ions. (B) Measured $k_\mathrm{cat}/K_\mathrm{M}$ of MeP hydrolysis (assuming full catalytic effect of valine and glycine mutations, as established for the alanine mutants) and $K_\mathrm{i}(\mathrm{P_i})$ effects for mutations to the O1- and O2-contacting residues. Effects are reported as fold-changes in $k_\mathrm{cat}/K_\mathrm{M}$ of MeP hydrolysis and $K_\mathrm{i}(\mathrm{P_i})$ relative to the wild-type values. To facilitate co-plotting, values $>1$ correspond to a deleterious effect on $k_\mathrm{cat}/K_\mathrm{M}$ and an increase in $K_\mathrm{i}(\mathrm{P_i})$. The $k_\mathrm{cat}/K_\mathrm{M}$ and $K_\mathrm{i}(\mathrm{P_i})$ values for the alanine mutants were taken from ref. (*16*) as the on-chip measurements of K162A and R164A $k_\mathrm{cat}/K_\mathrm{M}$ were upper limits. Up and down arrows denote lower and upper limits, respectively.

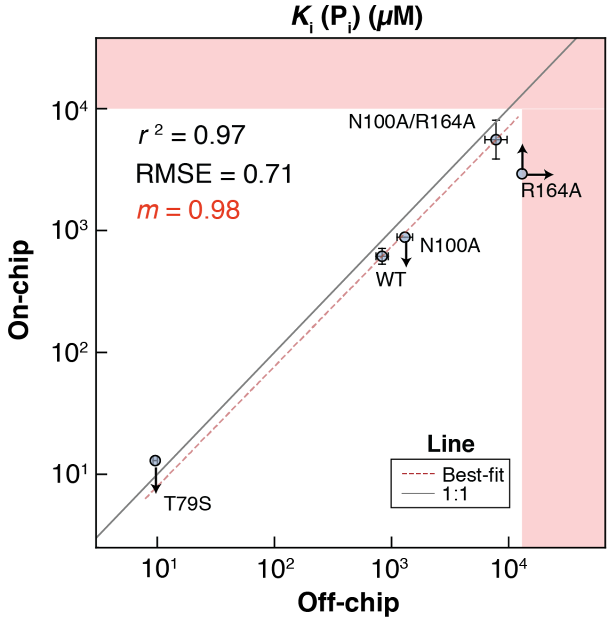

**Fig. S52.** Comparison of on- and off-chip $K_{\mathrm{i}}(\mathrm{P_i})$ measurements for WT PafA and 4 active site mutants. Reported $r^2$ and RMSE ($\log_{10}$-transformed values) are given for the 1:1 line, and $m$ corresponds to the slope of the best fit line fitted to the mutants for which $K_{\mathrm{i}}$ for $\mathrm{P_i}$ was not a limit (WT and N100A/R164A). Pink shaded regions denote the upper limits of dynamic range for $K_{\mathrm{i}}$ for $\mathrm{P_i}$, determined based on the highest $[\mathrm{P_i}]$ concentrations used in the assays; arrows denote upper and lower limits in $K_{\mathrm{i}}(\mathrm{P_i})$.

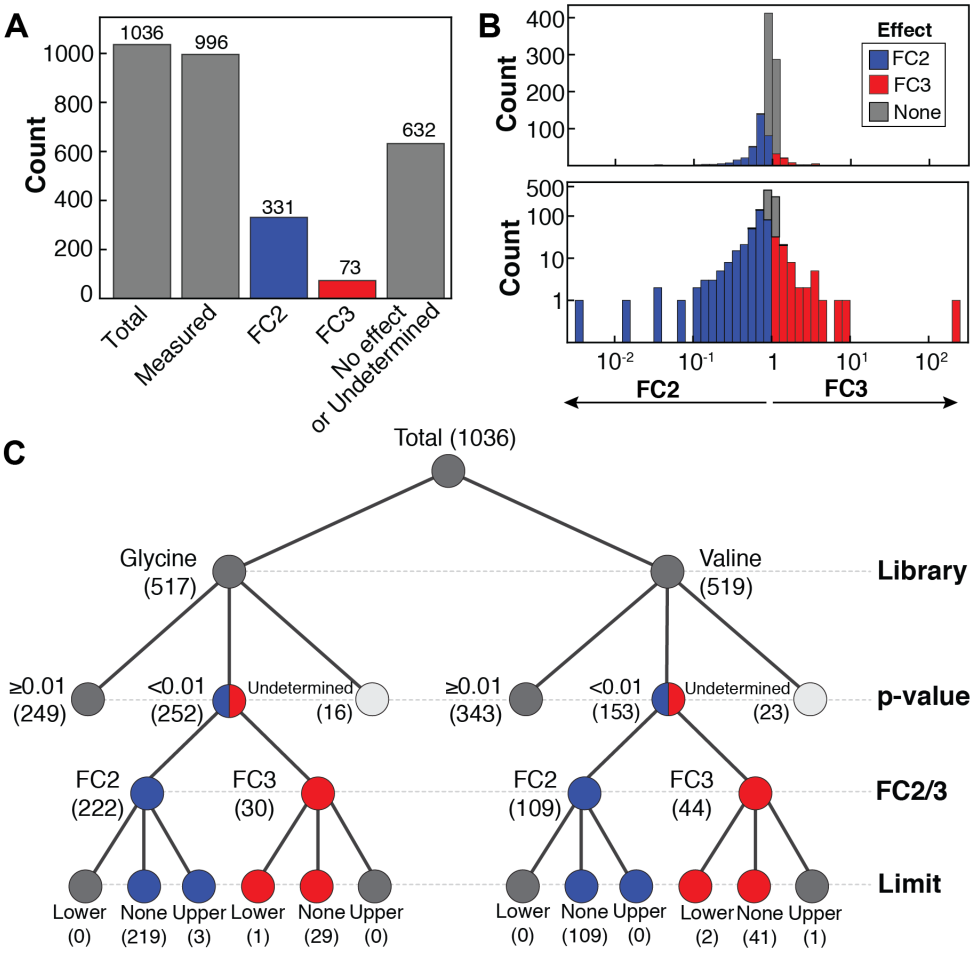

**Fig. S53.** Functional Component (FC) 2 and 3 effects within the glycine and valine scanning libraries. (A) Number of residues found to have FC2 or FC3 effects ($p<0.01$). "No effect or Undetermined" mutants also includes one mutant with an upper limit in FC3 effect (considered "Undefined"). (B) Distribution of measured FC2 and FC3 values, with counts displayed on a linear (top) and logarithmic scale (bottom). (C) Tree diagram summarizing the number of mutations with FC2 or FC3 effects ($p<0.01$) for the glycine and valine mutant libraries, including upper and lower limits as described in **Materials and Methods**. Mutants whose FC2 and FC3 values are lower and upper limits, respectively, cannot be distinguished from WT due to the direction of the limit.

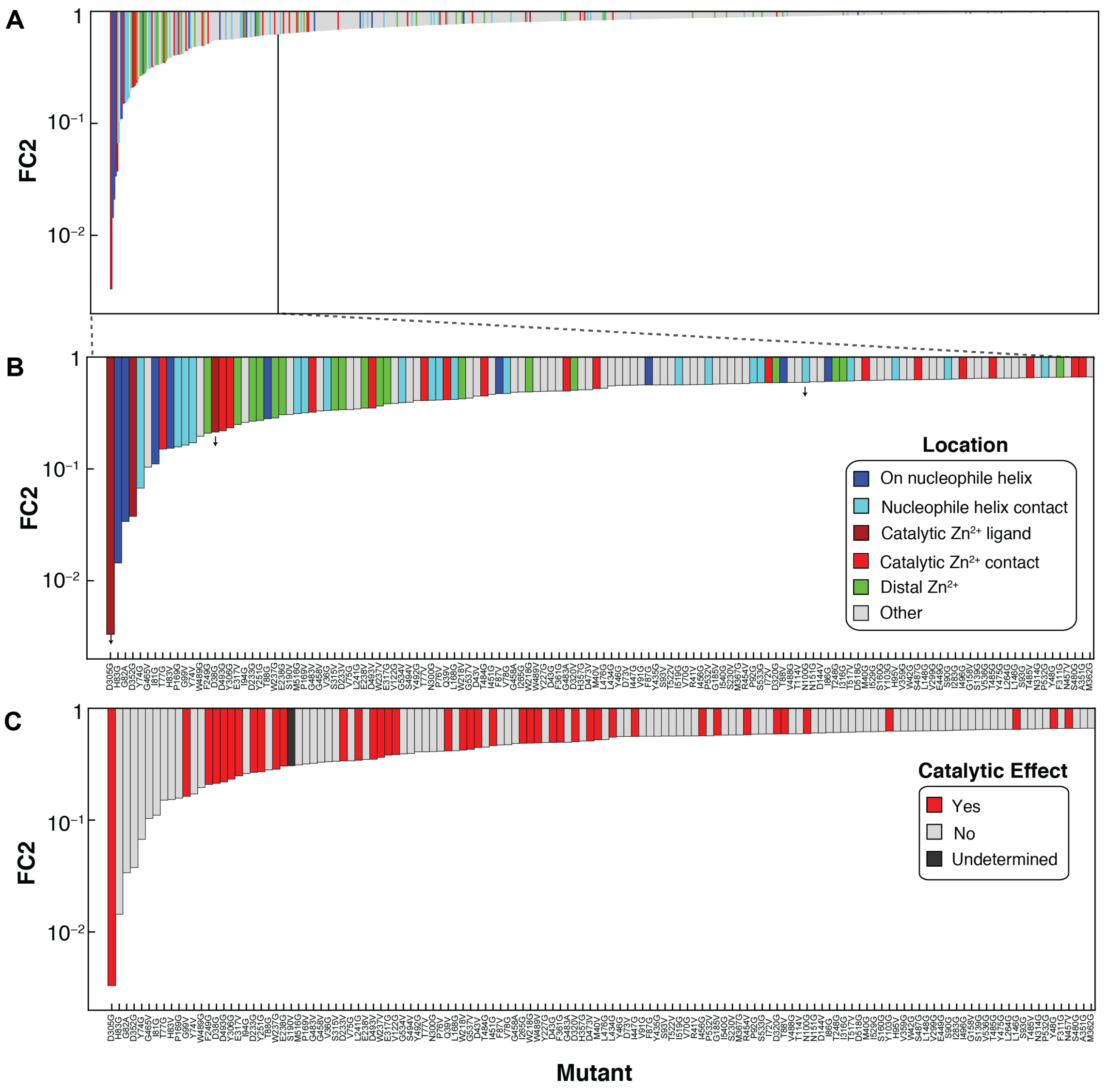

**Fig. S54.** Structural location and magnitudes of the largest FC2 effects. (A) Bar chart of all FC2 effects ($K_i^{\text{mutant}} < K_i^{\text{WT}}$ for $P_i$), sorted by magnitude and colored by their location in PafA. Arrows denote upper limits of FC2 measurements. (B) Same as panel A, but for the mutants with FC2 effects greater than 1.5-fold from wild-type. (C) Same as panel B, with bars colored by the presence of a catalytic effect ($k_{cat}/K_M^{\text{chem}}$) ($p < 0.05$).

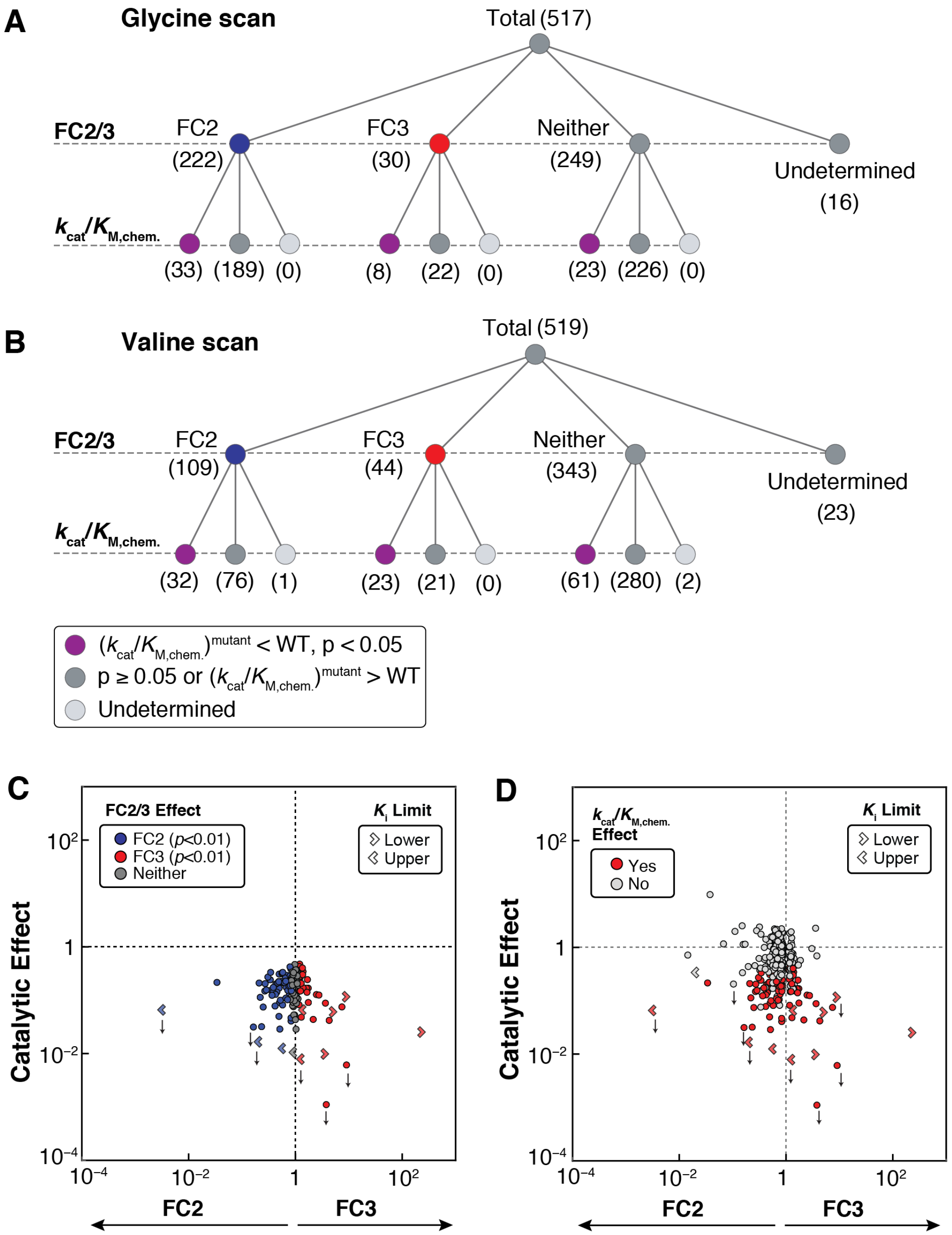

**Fig. S55.** Intersections of FC2/3 and catalytic ($k_{cat}/K_M^{chem}$) effects. (A) Tree with counts of residues with catalytic and FC2/3 effects, for the glycine scanning library. (B) Tree with counts of residues with catalytic and FC2/3 effects, for the valine scanning library. (C) Scatter plot of catalytic and FC2/3 effects for mutants with catalytic effects ($p$<0.05 and $(k_{cat}/K_M^{chem})^{mutant} < (k_{cat}/K_M^{chem})^{WT}$), colored by FC2/3 effects. (D) Scatter plot of catalytic and FC2/3 for mutants with FC2/3 effects ($p$ <0.01), colored by catalytic effect.

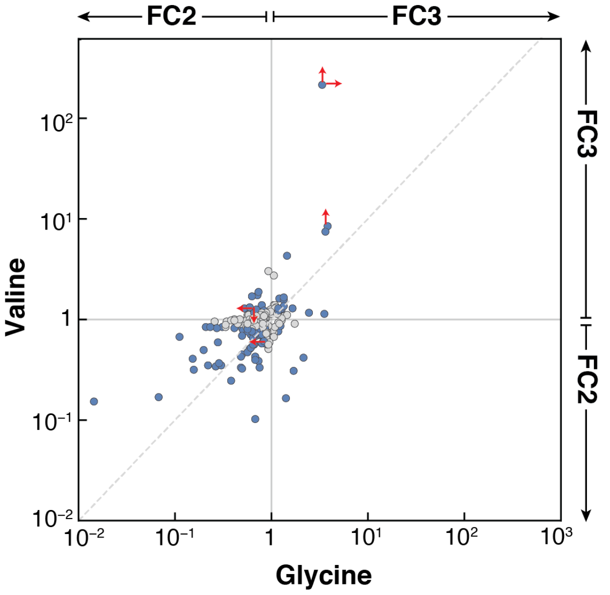

**Fig. S56.** Comparison of FC2/3 effects for valine or glycine substitutions. Residues with an FC2/3 effect ($p$<0.01) when mutated to either valine or glycine are shown in gray, and those with FC2/3 effects for both substitutions are shown in blue. Upper and lower limits are denoted by red arrows.

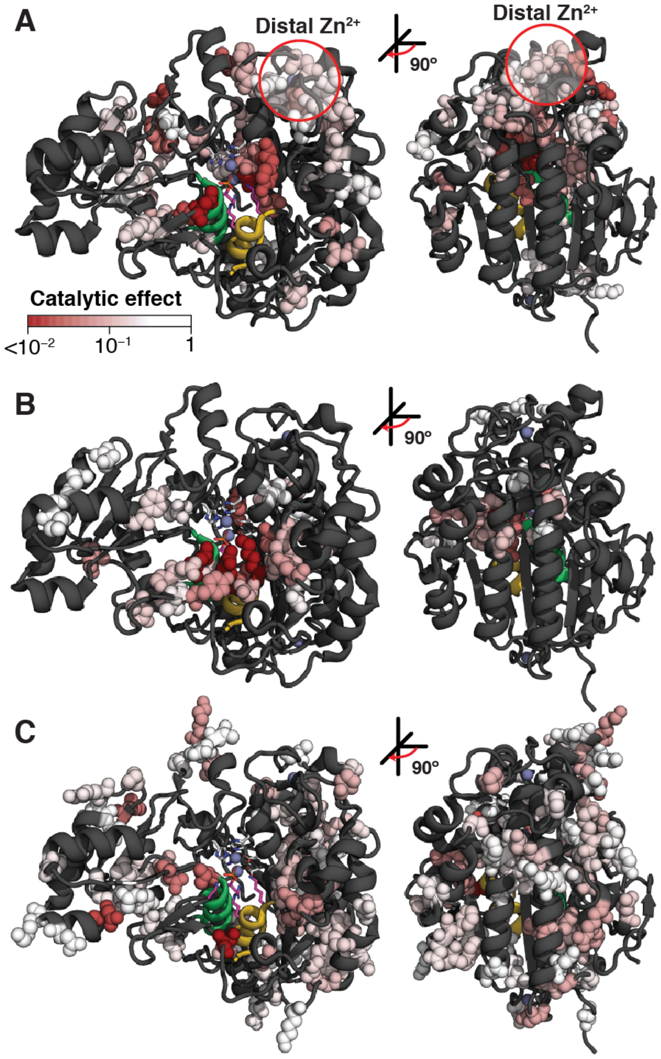

**Fig. S57.** Catalytic effects ($k_{cat}/K_M^{chem}$, fold-change relative to WT) for residues with FC2 or FC3 effects shown as spheres on the PafA structure and colored by the magnitude of the catalytic effect. (A) Positions (55 residues; 65 mutants) with FC2 and catalytic effects ($p<0.01$). (B) Positions (27 positions; 31 mutants) with FC3 and catalytic effects ($p<0.01$). (C) Positions (83 residues; 84 mutants) with catalytic effects ($p<0.01$) and no FC2/3 effects ($p \geq 0.01$).

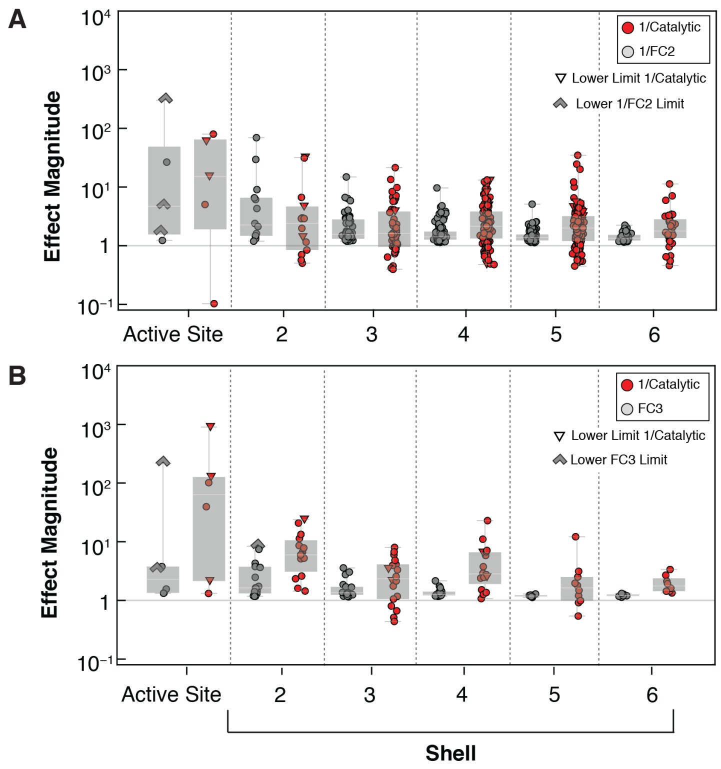

**Fig. S58.** Comparison of the relative magnitudes of catalytic and FC2/3 effects as a function of active site interaction shell. (A) Relative magnitude of FC2 (gray points) and catalytic ($k_{cat}/K_M^{chem}$; red points) effects for residues with FC2 effects ($p$<0.01). (B) Relative magnitude of FC3 (gray points) and catalytic ($k_{cat}/K_M^{chem}$; red points) effects for residues with FC3 effects ($p$<0.01).

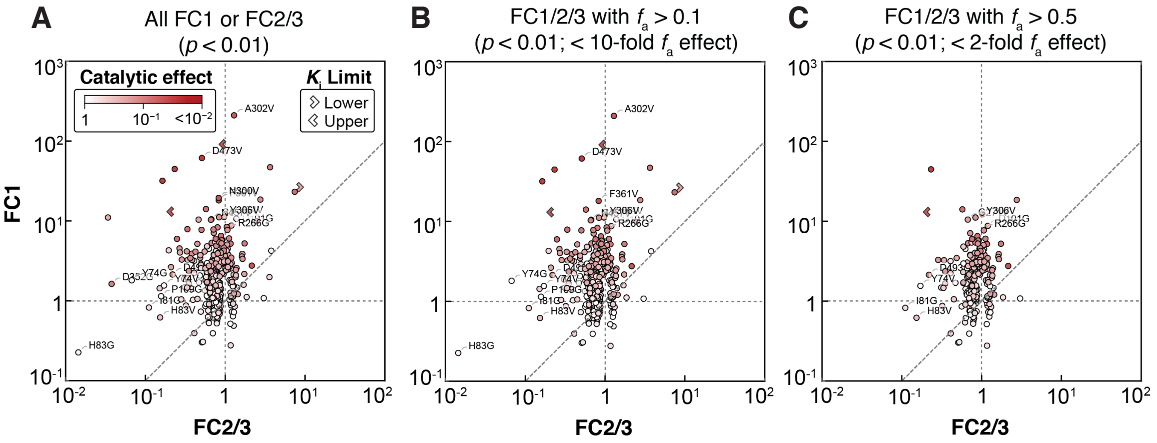

**Fig. S59.** Relative magnitudes of effects of mutations of FC1, FC2, FC3, catalysis ($k_{\mathrm{cat}}/K_{\mathrm{M}}^{\mathrm{chem}}$), and fraction active ($f_{\mathrm{a}}$). (A) Scatter plot showing FC1 effects vs. FC2/3 effects for all mutants for which all FCs were measurable, colored by $k_{\mathrm{cat}}/K_{\mathrm{M}}^{\mathrm{chem}}$. (B) Subset of mutations with $f_{\mathrm{a}}$ effects <10-fold from WT. (C) Subset of mutations with $f_{\mathrm{a}}$ effects <2-fold from WT.

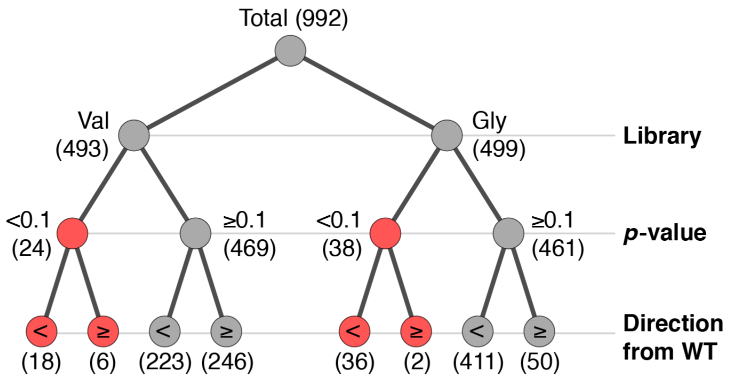

**Fig. S60.** Counts of mutational effects on the rate of phospho-enzyme intermediate hydrolysis ($k_{\mathrm{chem},2}$).

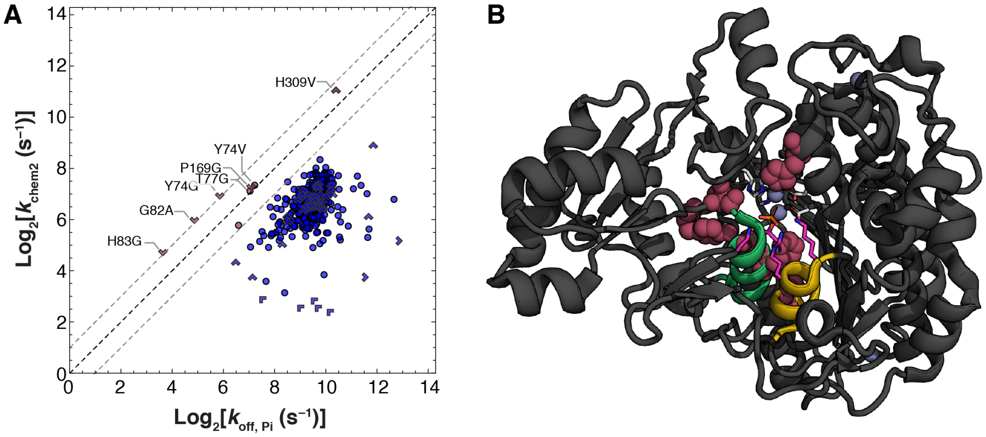

**Fig. S61.** Mutations with phospho-enzyme hydrolysis likely to be partially or fully rate limited by phosphate release. (A) Scatter plot of $k_{\mathrm{chem},2}$ vs. $k_{\mathrm{off,Pi}}$ for Val and Gly scanning mutants. Limits are denoted by chevrons pointing in the direction of or quadrant containing the point. Mutants partially or fully rate-limited by $\mathrm{P_i}$-release are highlighted (burgundy points, $k_{\mathrm{off,Pi}} < 2 \times k_{\mathrm{chem},2}$). (B) Positions of mutants partially or fully rate-limited by $\mathrm{P_i}$-release upon mutation as spheres on the PafA structure.

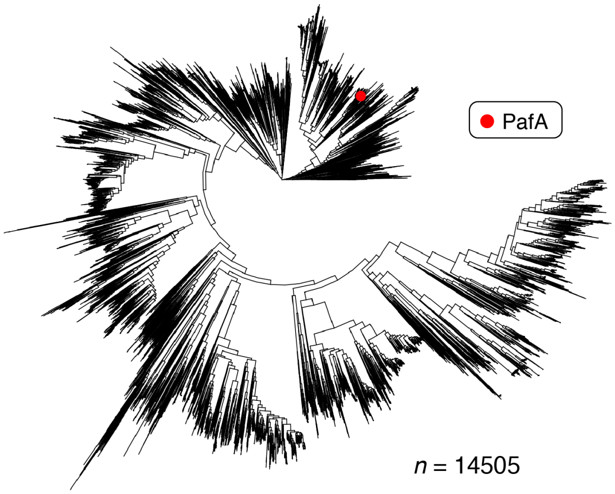

**Fig. S62.** A maximum-likelihood tree inferred from a multiple sequence alignment of 14,505 metagenomic and public database PafA-like AP superfamily members.

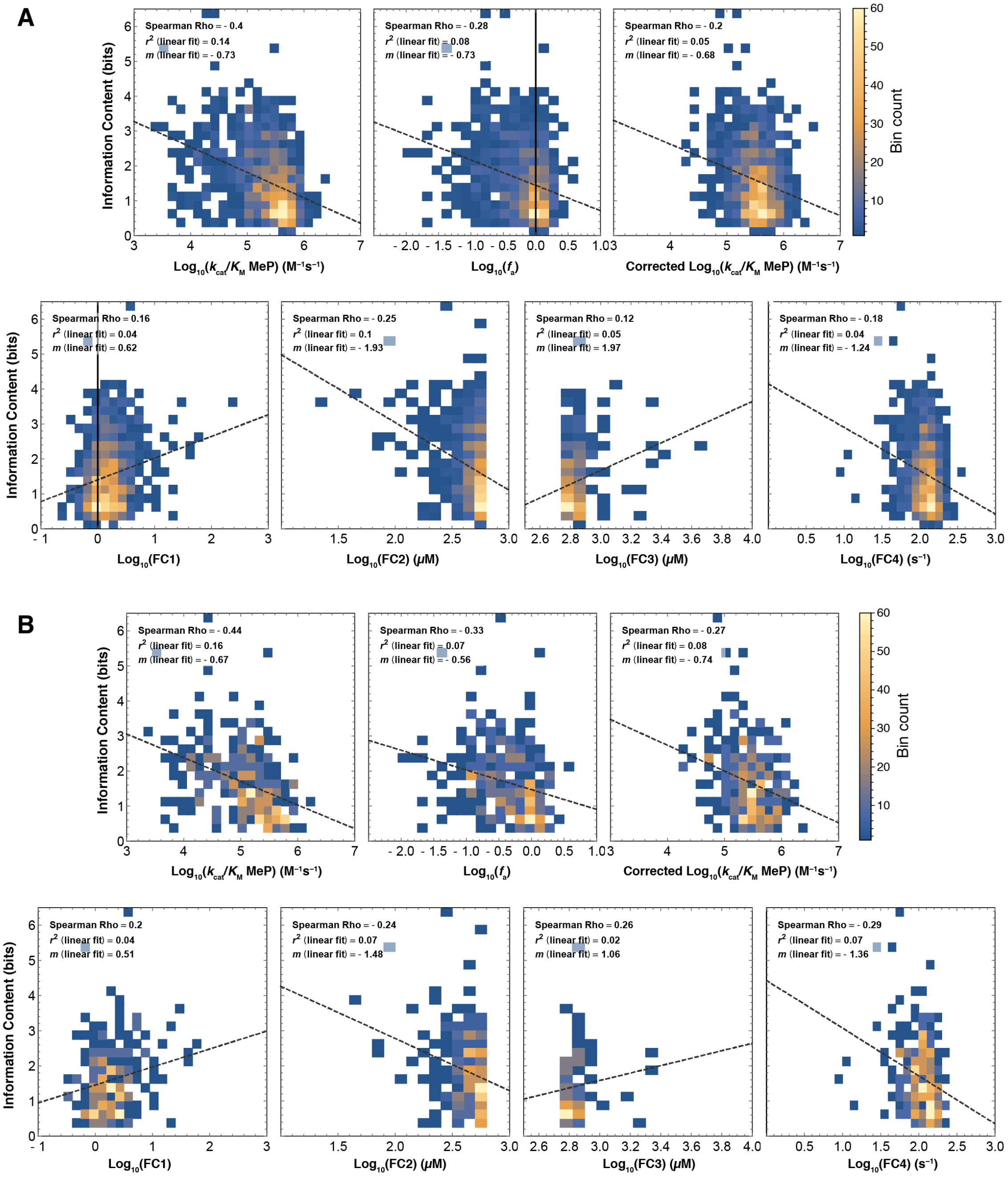

**Fig. S63.** Correlations between phylogenetic conservation and HT-MEK measured parameters and FCs. (A) Correlations (Spearman's rank, $\rho$; linear fit $r^2$) between conservation and $k_{\text{cat}}/K_{\text{M}}^{\text{obs}}$ MeP, $f_{\text{a}}$, $k_{\text{cat}}/K_{\text{M}}^{\text{chem}}$ MeP, and FCs 1–4 for all valine and glycine mutants. (B) Correlations (Spearman rank, $\rho$; linear fit $r^2$) between conservation and $k_{\text{cat}}/K_{\text{M}}^{\text{obs}}$ MeP, $f_{\text{a}}$, $k_{\text{cat}}/K_{\text{M}}^{\text{chem}}$ MeP, and measured FCs 1–4 only for mutations likely to dramatically alter biochemical properties at a given position (with a BLOSUM62 score of substitution <–2 when considering the difference between the WT and mutant residue identities).

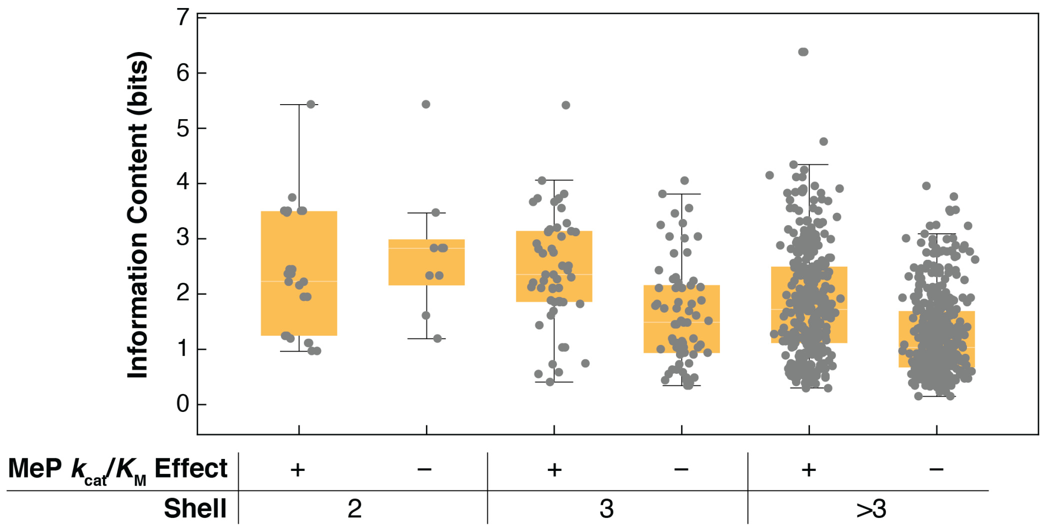

**Fig. S64.** Conservation is not sufficient to accurately predict effects of particular mutations near the active site. Conservation (Information Content) of residues with and without significant observed MeP ($k_{\mathrm{cat}}/K_{\mathrm{M}}$) effects when mutated for residues in the second and third interaction shells around the active site residues K162 and R164, and in all other shells.

### Tables S1 to S16

| Construct | Substrate | $k_{cat}$ ($s^{-1}$) | $K_M$ (M) | $k_{cat}/K_M$ ($M^{-1}s^{-1}$) |
|---|---|---|---|---|
| **WT** | **pNPP** | $3.0 \times 10^2$ | $1.9 \times 10^{-4}$ | $1.6 \times 10^6$ |
| | **cMUP** | $3.2 \times 10^2$ | $1.1 \times 10^{-4}$ | $3.0 \times 10^6$ |
| **R164A** | **pNPP** | 6.6 | $5.1 \times 10^{-5}$ | $1.3 \times 10^5$ |
| | **cMUP** | 4.7 | $3.2 \times 10^{-5}$ | $1.4 \times 10^5$ |
| **T79S** | **pNPP** | 1.8 | $2.9 \times 10^{-6}$ | $6.3 \times 10^5$ |
| | **cMUP** | $9.7 \times 10^{-1}$ | $1.0 \times 10^{-6}$ | $9.2 \times 10^5$ |
| **N100A** | **pNPP** | $3.9 \times 10^{-1}$ | $5.1 \times 10^{-7}$ | $7.6 \times 10^5$ |
| | **cMUP** | $9.9 \times 10^{-1}$ | $7.5 \times 10^{-7}$ | $9.0 \times 10^5$ |
| **N100A/R164A** | **pNPP** | – | – | – |
| | **cMUP** | $9.2 \times 10^{-1}$ | $1.9 \times 10^{-4}$ | $5.0 \times 10^3$ |
| **K162A** | **pNPP** | $3.0 \times 10^{-2}$ | $2.3 \times 10^{-4}$ | $1.3 \times 10^2$ |
| | **cMUP** | $2.8 \times 10^{-2}$ | $4.1 \times 10^{-5}$ | $2.8 \times 10^2$ |

**Table S1.** Measured Michaelis-Menten parameters for hydrolysis of pNPP and cMUP by PafA active site mutants. All measurements were performed with the Strep-tag bearing (non eGFP-fusion) constructs expressed in *E. coli*. pNPP hydrolysis parameters are reproduced from (*16*) and cMUP parameter estimates are means of two or more replicates and were measured herein (see **Materials and Methods**).

| | | | Chambers | Unique Mutants | | | | |
|---|---|---|---|---|---|---|---|---|
| **Tier** | **Print** | **Library** | **Empty & T79G** | **All** | **Val** | **Gly** | **A.S.** | **Median # reps** |
| – | **A.S.** | **A.S.** | 1248 | 7 | 0 | 1 | 6 | 50 |
| **I** | **Val I** | **Val, A.S.** | 68 | 532 | 519 | 7 | 6 | 2 |
| **I** | **Val II** | **Val, A.S.** | 126 | 506 | 497 | 3 | 6 | 2 |
| **I** | **Gly** | **Gly, A.S.** | 298 | 523 | 0 | 517 | 6 | 2 |
| **II** | **Slow** | **All** | 350 | 211 | 65 | 140 | 6 | 4 |
| **II** | **Slow MeP** | **All** | 768 | 43 | 32 | 6 | 5 | 16 |
| **III** | **Slowest** | **All** | 1028 | 61 | 26 | 30 | 5 | 8 |
| **All** | – | **All** | – | 1042 | 519 | 517 | 6 | – |

**Table S2.** Summary of tiered measurement strategy. Skipped chambers were those lacking printed plasmid, and "A.S." denotes Active Site.

| | $k_{cat}/K_M$ ($M^{-1}s^{-1}$) | | |
|---|---|---|---|
| **Mutant** | **pNPP, off-chip** | **cMUP, on-chip** | $k_{rel}$ **(off/on-chip)** |
| **WT** | $2.20 \times 10^6$ | $1.42 \times 10^6$ | 1.5 |
| **Y52G** | $1.47 \times 10^6$ | $8.39 \times 10^5$ | 1.7 |
| **K58V** | $1.86 \times 10^6$ | $1.08 \times 10^6$ | 1.7 |
| **A80V** | $5.52 \times 10^5$ | $2.02 \times 10^5$ | 2.7 |
| **F87G** | $1.77 \times 10^6$ | $1.99 \times 10^6$ | 0.9 |
| **T88V** | $7.01 \times 10^5$ | $7.51 \times 10^5$ | 0.9 |
| **S90V** | $2.90 \times 10^5$ | $2.65 \times 10^5$ | 1.1 |
| **G108A** | $1.83 \times 10^6$ | $1.33 \times 10^6$ | 1.4 |
| **G130V** | $6.40 \times 10^5$ | $7.68 \times 10^5$ | 0.8 |
| **R164A** | $3.30 \times 10^5$ | $2.64 \times 10^5$ | 1.3 |
| **T189G** | $4.80 \times 10^5$ | $2.83 \times 10^5$ | 1.7 |
| **S190G** | $2.80 \times 10^5$ | $3.97 \times 10^5$ | 0.7 |
| **P198V** | $2.27 \times 10^6$ | $1.86 \times 10^6$ | 1.2 |
| **R266V** | $2.10 \times 10^6$ | $1.63 \times 10^6$ | 1.3 |
| **Y306V** | $1.90 \times 10^6$ | $8.69 \times 10^5$ | 2.2 |
| **A330G** | $2.43 \times 10^6$ | $2.33 \times 10^6$ | 1.0 |
| **L349G** | $5.89 \times 10^5$ | $1.13 \times 10^5$ | 5.2 |
| **R463V** | $1.44 \times 10^6$ | $4.51 \times 10^5$ | 3.2 |
| **L499G** | $9.71 \times 10^5$ | $1.12 \times 10^6$ | 0.9 |

**Table S3.** Library variants (18 Val and Gly, 1 Active Site, and the WT) expressed *in vitro* and characterized off-chip to test for chip-specific expression effects.

| | On-chip $k_{cat}/K_M$ ($M^{-1}s^{-1}$) | | | ($k_{cat}/K_M$ $^{37°C}$)/($k_{cat}/K_M$ $^{23°C}$) (WT-normalized) | |
|---|---|---|---|---|---|
| **Mutant** | **cMUP** | **MeP** | **($k_{cat}/K_M^{MeP, WT}$)/($k_{cat}/K_M^{MeP, mutant}$)** | **On-chip** | **Off-chip** |
| **F87G** | $1.99 \times 10^6$ | $6.65 \times 10^5$ | 0.9 | 4.7 | 2.6 |
| **A330G** | $2.33 \times 10^6$ | $4.28 \times 10^5$ | 1.4 | 3.1 | 1.7 |
| **G108A** | $1.33 \times 10^6$ | $2.58 \times 10^5$ | 2.4 | 2.8 | 1.9 |
| **K58V** | $1.08 \times 10^6$ | $2.14 \times 10^5$ | 2.9 | 5.9 | 2.0 |
| **L499G** | $1.12 \times 10^6$ | $1.34 \times 10^5$ | 4.6 | 30 | 8.5 |
| **Y52G** | $8.39 \times 10^5$ | $1.19 \times 10^5$ | 5.1 | 20 | 6.5 |
| **F180V** | $1.91 \times 10^6$ | $1.00 \times 10^5$ | 6.1 | 0.9 | 2.5 |
| **T304V** | $1.19 \times 10^5$ | $9.96 \times 10^4$ | 6.1 | 1.8 | 2.1 |
| **N173V** | $3.78 \times 10^5$ | $9.94 \times 10^4$ | 6.1 | 1.7 | 2.2 |
| **R266V** | $1.63 \times 10^6$ | $8.92 \times 10^4$ | 6.8 | 1.6 | 0.8 |
| **S90V** | $2.65 \times 10^5$ | $8.48 \times 10^4$ | 7.2 | 3.6 | 4.2 |
| **S190G** | $3.97 \times 10^5$ | $7.97 \times 10^4$ | 7.7 | 2.1 | 2.8 |
| **T189G** | $2.83 \times 10^5$ | $7.01 \times 10^4$ | 8.7 | 7.4 | 3.0 |
| **D182G** | $1.48 \times 10^6$ | $5.92 \times 10^4$ | 10.3 | 2.4 | 3.8 |
| **Y306V** | $8.69 \times 10^5$ | $5.70 \times 10^4$ | 10.7 | 4.0 | 1.2 |
| **V433G** | $1.56 \times 10^5$ | $3.42 \times 10^4$ | 17.9 | 9.7 | 1.5 |
| **Y103G** | $2.82 \times 10^5$ | $2.61 \times 10^4$ | 23 | 14 | 4.9 |
| **W218G** | $2.17 \times 10^5$ | $2.09 \times 10^4$ | 29 | 4.3 | 1.9 |
| **A80V** | $2.02 \times 10^5$ | $1.48 \times 10^4$ | 41 | 1.2 | 3.2 |
| **R164A** | $2.64 \times 10^5$ | $7.16 \times 10^3$ | 85 | 0.8 | 0.5 |

**Table S4.** Library variants (19 Val and Gly and 1 Active Site) characterized off chip and expressed *in vivo* to test for expression-temperature-dependent activity changes.

| Technique | PafA Variant | Expression Temperature (°C) | % Content α-helix | % Content β-Sheet |
|---|---|---|---|---|
| X-ray Crystallography (RCSB PDB ID) | WT (5TJ3) | 30 | 34 | 16 |
| | pruned[a] (5TOO) | 23 | 34 | 16 |
| Circular Dichroism | WT | 23 | 32 | 14 |
| | | 37 | 31 | 14 |
| | Y103G | 23 | 32 | 15 |
| | | 37 | 20 | 17 |

[a] "Pruned" PafA bears the following mutations: T79S, N100A, K162A, and R164A (*109*).

**Table S5.** Secondary structure content of PafA variants calculated from previously published crystallographic data and circular dichroism (CD) data collected herein. Content was calculated for CD data with the K2D3 web server and for crystallographic data with DSSP (*110*, *111*).

| Distance from active site (Å) | Catalytic Effect ($k_{cat}/K_M^{MeP, chem. mutant}/k_{cat}/K_M^{MeP, WT}$) | | | | | |
|---|---|---|---|---|---|---|
| | $p < 0.05$ | | | $p \geq 0.05$ | Undefined *p*-value | All *p*-values |
| | < 0.1 | ≥ 0.1 & < 0.2 | ≥ 0.2 & < 1 | Any | Undefined | All |
| **0–5** | 12 | 13 | 8 | 68 | 9 | 110 |
| **5–10** | 6 | 13 | 17 | 140 | 4 | 180 |
| **10–15** | 6 | 20 | 25 | 225 | 10 | 286 |
| **15–20** | 5 | 8 | 26 | 201 | 4 | 244 |
| **20–25** | 0 | 2 | 9 | 107 | 3 | 121 |
| **25–30** | 1 | 0 | 4 | 26 | 1 | 32 |
| **30–35** | 0 | 0 | 1 | 9 | 0 | 10 |
| **35–40** | 0 | 0 | 0 | 2 | 0 | 2 |
| **N.D.**[a] | 0 | 0 | 0 | 12 | 0 | 12 |

[a]N.D. denotes mutations of residues for which the minimum distance from the active site cannot be determined, as these are residues are unmodeled in the PafA crystal structure (PDB ID: 5TJ3, (*16*); see **Materials and Methods**).

**Table S6.** Catalytic effects of all non-active-site variants ($k_{cat}/K_M{}^{MeP, \text{ chem. mutant}}/k_{cat}/K_M{}^{MeP, \text{ WT}}$ $< 1$) as a function of minimum distance from the active site, binned by magnitude of effect and $p$-value.

| A | Scan | | | | | | | |
|---|---|---|---|---|---|---|---|---|
| | Glycine | | Valine | | Contact[a] | | | |
| Residue | FC1 | *p*-value | FC1 | *p*-value | K162 | R164 | $Zn^{2+}$ | $O_n$ |
| **Y112** | $4.7 \times 10^1$ | $<1.0 \times 10^{-5}$ | $2.3 \times 10^1$ | $1.4 \times 10^{-4}$ | – | sc | – | – |
| **C113** | 2.5 | $3.9 \times 10^{-3}$ | 1.4 | 0.28 | – | sc | – | – |
| **S160** | 0.5 | $5.7 \times 10^{-2}$ | – | – | bb | – | – | – |
| **L161** | 9.9 | $6.9 \times 10^{-4}$ | 1.6 | $2.8 \times 10^{-2}$ | bb | – | – | – |
| **D163** | 4.2 | $7.9 \times 10^{-4}$ | $2.6 \times 10^1$ | $2.0 \times 10^{-4}$ | – | bb | – | – |
| **A165** | 1.2 | 0.62 | – | – | sc | – | – | – |
| **S166** | 2.8 | $1.6 \times 10^{-4}$ | 1.8 | $8.9 \times 10^{-2}$ | sc | – | – | – |
| **L168** | – | – | 0.8 | 0.55 | – | bb | – | – |
| **W179** | – | – | – | – | bb | – | – | – |
| **A302** | 3.9 | $1.1 \times 10^{-2}$ | $2.1 \times 10^2$ | $<1.0 \times 10^{-5}$ | sc | – | – | – |
| **Y306** | $4.5 \times 10^1$ | $<1.0 \times 10^{-5}$ | $1.1 \times 10^1$ | $1.0 \times 10^{-5}$ | sc | – | – | – |

| B | Scan | | | | | | | |
|---|---|---|---|---|---|---|---|---|
| | Glycine | | Valine | | Contact[a] | | | |
| Residue | FC1 | *p*-value | FC1 | *p*-value | K162 | R164 | $Zn^{2+}$ | $O_n$ |
| **N100** | $2.6 \times 10^1$ | $1.2 \times 10^{-3}$ | – | – | – | sc | – | 1 |
| **K162** | $>2.7 \times 10^3$ | $<1.0 \times 10^{-5}$ | $>1.7 \times 10^4$ | $<1.0 \times 10^{-5}$ | – | sc | – | 2 |
| **R164** | $2.0 \times 10^3$ | $<1 \times 10^{-5}$ | $1.4 \times 10^3$ | $<1.0 \times 10^{-5}$ | sc | – | – | 2 |
| **D38** | $>1.3 \times 10^1$ | $1.7 \times 10^{-4}$ | – | – | sc | – | – | – |
| **D305** | – | – | – | – | sc | – | 1 | – |
| **H309** | – | – | – | – | – | – | 1 | – |
| **H486** | – | – | – | – | – | – | 1 | – |
| **T79** | – | – | – | – | – | sc | 2 | – |
| **H353** | – | – | 1.0 | 0.9 | – | – | 2 | – |
| **D352** | 1.6 | 0.12 | – | – | – | – | 2 | – |

[a]Denotes interactions between each residue and K162, R164, either active site $Zn^{2+}$ ion (Zn1 denoted by "1" and Zn2 denoted by "2") or a substrate phosphoryl oxygen (O1 denoted by "1" and O2 denoted by "2"). Interactions with the K162 or R164 side chain or backbone atoms are denoted by "sc" and "bb," respectively.

**Table S7.** FC1 effects for glycine and valine mutations of the (A) 2nd shell and (B) active site residues.

| | FC1 | | | |
|---|---|---|---|---|
| | On-chip | | Off-chip | |
| Mutant | Median | *p*-value | MecMUP:MeP[a] | MepNPP:MeP[b] |
| **T79S** | $1.4 \times 10^{1}$ | 0.0005 | 7.6 | 0.9 |
| **N100A** | – | – | 6.2 | 2.5 |
| **K162A** | $>1.1 \times 10^{2}$ | $<1.0 \times 10^{-5}$ | – | $3.6 \times 10^{6}$ |
| **R164A** | $>5.8 \times 10^{2}$ | $<1.0 \times 10^{-5}$ | $1.6 \times 10^{4}$ | $1.7 \times 10^{3}$ |

[a]See **Materials and Methods** for assay conditions. [b]FC1 using MepNPP and MeP calculated from ref. (*16*).

**Table S8.** Comparison of on- and off-chip FC1 measurements for active site mutants.

| | FC1[a] | | | | | |
|---|---|---|---|---|---|---|
| | $p < 0.01$ | | | $p \geq 0.01$ | Undefined *p*-value | All *p*-values |
| **Shell** | **> 10** | **> 5 & < 10** | **< 5** | **Any** | **Undefined** | **Any** |
| **Active Site** | 4 | 0 | 0 | 2 | 4 | 10 |
| **2** | 4 | 1 | 2 | 3 | 1 | 11 |
| **3** | 6 | 2 | 8 | 24 | 2 | 42 |
| **4** | 3 | 3 | 22 | 52 | 7 | 87 |
| **5** | 0 | 4 | 28 | 81 | 5 | 118 |
| **6** | 3 | 5 | 24 | 79 | 6 | 117 |
| **7** | 3 | 3 | 23 | 54 | 5 | 88 |
| **8** | 0 | 0 | 8 | 35 | 2 | 45 |
| **9** | 0 | 0 | 0 | 2 | 0 | 2 |
| **N.D.[b]** | 0 | 0 | 0 | 6 | 0 | 6 |
| **Surface Exposed[c]** | 4 | 6 | 53 | 151 | 6 | 220 |

[a]FC1 and *p*-values for each residue were determined from the larger FC1 effect of the valine and glycine library variants. [b]N.D. denotes mutations of residues for which the interaction shell cannot be determined, as these are residues are unmodeled in the PafA crystal structure (PDB ID: 5TJ3, (*16*); see **Materials and Methods**). [c]See **Materials and Methods** for details of determining surface exposure.

**Table S9.** Distribution of FC1 effect magnitudes for residues within the active site, each K162/R164 interaction shell, and within the subset of residues that are surface exposed.

| | | | | | Auxiliary Domain | | | | | | |
|---|---|---|---|---|---|---|---|---|---|---|---|
| **Name** | **Organism** | **Abbreviation** | **PDB ID** | **Cognate Reaction** | **N-terminal** | **1** | **2** | **3.1** | **3.2** | **4** | **C-terminal** |
| **Alkaline phosphatase** | *Escherichia coli* | EcAP | 3TG0 | Phosphomonoesterase | N-terminus:P41 | A88:V99 | K114:D128 | V146:A200 | G207:P297 | A373:E411 | – |
| **Alkaline phosphatase** | *Halobacterium salinarum* | HSAP | 2X98 | Phosphomonoesterase | – | P105:T112 | K127:V146 | I163:P199 | Q206:N272 | G350:G434 | – |
| **Cold-active alkaline phosphatase** | *Shewanella sp.* AP1 | SCAP | 3A52 | Phosphomonoesterase | – | P48:V55 | K70:P84 | A102:A137 | G144:N197 | G273:G361 | – |
| **Alkaline phosphatase** | *Vibrio sp. G15-21* | VLAP | 3E2D | Phosphomonoesterase | – | P57:V62 | Y77:H91 | V109:A144 | G151:S244 | G319:T464 | – |
| **Alkaline phosphatase** | *Halomonas sp. #593* | HaAP | 3WBH | Phosphomonoesterase | – | P57:V62 | Y77:P91 | V109:P144 | G151:T240 | G315:T460 | – |
| **Intestinal-type alkaline phosphatase 1** | *Rattus norvegicus* | IAP | 4KJG | Phosphomonoesterase | N-terminus:S32 | N84:V89 | K104:E128 | V146:I196 | G203:P286 | V361:T431 | – |
| **Alkaline phosphatase PhoK** | *Sphingomonas sp.* | SPAP | 3Q3Q | Phosphomonoesterase | N-terminus:P39 | – | R101:L147 | V168:A183 | G190:T270 | G348:T490 | – |
| **Alkaline phosphatase PafA** | *Elizabethkingia meningoseptica* | PafA | 5TJ3 | Phosphomonoesterase | N-terminus:P27 | – | V91:L137 | V159:P174 | D181:T275 | A356:T485 | – |
| **Alkaline phosphatase, placental type** | *Homo sapiens* | PLAP | 3MK1 | Phosphomonoesterase | N-terminus:A32 | N84:V89 | K104:E128 | V146:I196 | G203:P286 | V361:T431 | – |
| **Alkaline phosphatase** | *Pandalus borealis* | SAP | 1SHQ | Phosphomonoesterase | N-terminus:Q27 | N78:V83 | A92:F124 | V142:F196 | G203:P285 | T360:T431 | – |
| **Alkaline phosphatase** | Antarctic bacterium TAB5 | TAP | 2IUC | Phosphomonoesterase | – | S76:V81 | K96:A110 | V128:L163 | G170:N229 | G305:G336 | – |
| **Phosphodiesterase-nucleotide pyrophosphatase** | *Xanthomonas axonopodis pv. Citri* | XacNPP | 2RH6 | Phosphodiesterase | N-terminus:A42 | – | R102:G136 | W154:P168 | K176:V178 | A261:S362 | – |
| **Venom phosphodiesterase** | *Naja atra* | PDE | 5GZ5 | Phosphodiesterase | N-terminus:F136 | – | Y197:G231 | Y249:P263 | K271:T273 | E356:T461 | P525:C-terminus |
| **Ectonucleotide pyrophosphatase/ phosphodiesterase family member 1** | *Mus musculus* | ENPP1 | 4B56 | Phosphodiesterase | N-terminus:F189 | – | Y250:K284 | Y302:P316 | G324:V326 | E409:F516 | P580:C-terminus |
| **Ectonucleotide pyrophosphatase/ phosphodiesterase family member 2** | *Homo sapiens* | ENPP2 | 4ZG7 | Phosphodiesterase | N-terminus:F161 | – | Y222:G256 | F275:P281 | V278:I280 | E363:D474 | N538:C-terminus |
| **Ectonucleotide pyrophosphatase/ phosphodiesterase family member 2** | *Rattus norvegicus* | ENPP2 | 5DLT | Phosphodiesterase | N-terminus:F160 | – | Y221:G255 | F274:I279 | V277:I279 | E362:D473 | N537:C-terminus |
| **Ectonucleotide pyrophosphatase/ phosphodiesterase family member 2** | *Mus musculus* | ENPP2 | 3NKR | Phosphodiesterase | N-terminus:F160 | – | Y221:G255 | F274:I279 | V277:I279 | E362:D473 | N537:C-terminus |
| **Ectonucleotide pyrophosphatase/ phosphodiesterase family member 3** | *Homo sapiens* | ENPP3 | 6C01 | Phosphodiesterase | N-terminus:F156 | – | Y217:H251 | Y269:P283 | G291:V293 | D376:N482 | P546:C-terminus |
| **Ectonucleotide pyrophosphatase/ phosphodiesterase family member 3** | *Rattus norvegicus* | ENPP3 | 6F2T | Phosphodiesterase | N-terminus:F157 | – | Y218:S252 | Y270:P284 | N292:V294 | D377:T482 | P546:C-terminus |
| **Bis(5'-adenosyl)-triphosphatase ENPP4** | *Homo sapiens* | ENPP4 | 4LQY | Phosphodiesterase | – | – | Y82:E115 | A134:S148 | S156:V158 | T241:D335 | – |
| **Ectonucleotide pyrophosphatase/ phosphodiesterase family member 5** | *Homo sapiens* | ENPP5 | 5VEM | Phosphodiesterase | N-terminus:L24 | – | F84:E119 | A137:P151 | E159:V161 | T242:N338 | – |
| **Glycerophosphocholine cholinephosphodiesterase ENPP6** | *Mus musculus* | ENPP6 | 5EGH | Phosphodiesterase | – | – | H83:G119 | Y137:P151 | T159:P161 | T244:W353 | – |
| **Ectonucleotide pyrophosphatase/ phosphodiesterase family member 7** | *Homo sapiens* | ENPP7 | 5TCD | Phosphodiesterase | – | – | Y87:G122 | F140:V154 | I162:E171 | T250:E352 | – |

**Table S10.** Unique structurally characterized AP superfamily phosphate monoesterases and diesterases and definitions for their non-conserved "Auxiliary Domains."

| | FC1 | | | | | |
|---|---|---|---|---|---|---|
| | $p < 0.01$ | | | $p \geq 0.01$ | Undefined $p$-value | All $p$-values |
| **Domain** | **>10** | **>5 & <10** | **<5** | **Any** | **Undefined** | **Any** |
| Rossman core | 5 | 11 | 45 | 157 | 20 | 238 |
| Aux. 2 | 5 | 2 | 14 | 24 | 2 | 47 |
| Aux. 3 | 7 | 3 | 28 | 68 | 5 | 111 |
| Aux. 4 | 6 | 2 | 28 | 89 | 5 | 130 |

**Table S11.** Number and magnitudes of FC1 effects within the Rossmann fold core and the three Auxiliary ("Aux.") Domains.

|  | Effect |  |  |  |
|---|---|---|---|---|
| **Region** | **FC2[a]** | **Catalytic[b]** | **FC2[a] & Catalytic[b]** | **None** |
| **Nucleophile helix** | 0.69 | 0.063 | 0.13 | 0.13 |
| **Nucleophile helix contact** | 0.52 | 0.15 | 0.05 | 0.28 |
| **Catalytic $Zn^{2+}$ ligand** | 0.16 | 0 | 0.5 | 0.33 |
| **Catalytic $Zn^{2+}$ contact** | 0.47 | 0.14 | 0.16 | 0.23 |
| **"Distal" $Zn^{2+}$ contact** | 0.27 | 0.02 | 0.27 | 0.44 |
| **Other** | 0.23 | 0.12 | 0.04 | 0.61 |

[a]FC2 effects defined as having $p<0.01$ as determined from bootstrap hypothesis testing.
[b]Catalytic effects defined as having $p<0.05$ as determined from bootstrap hypothesis testing, and $(k_{cat}/K_M^{chem})^{mutant} < (k_{cat}/K_M^{chem})^{WT}$.

**Table S12.** Fraction of measurable mutants with catalytic ($k_{cat}/K_M^{chem}$) effects, FC2 effects, and mutants with both catalytic and FC2 effects within different PafA regions. See **Table S13** for counts of mutants in each category.

| | Effect | | | | | |
|---|---|---|---|---|---|---|
| **Region** | **Total** | **FC2[a]** | **Catalytic[b]** | **FC2[a] & Catalytic[b]** | **None** | **Undetermined** |
| Nucleophile helix | 19 | 11 | 1 | 2 | 2 | 3 |
| Nucleophile helix contact | 65 | 31 | 9 | 3 | 17 | 5 |
| Catalytic $Zn^{2+}$ | 12 | 1 | 0 | 3 | 2 | 6 |
| Catalytic $Zn^{2+}$ contact | 44 | 20 | 6 | 7 | 10 | 1 |
| "Distal" $Zn^{2+}$ contact | 48 | 13 | 1 | 13 | 21 | 0 |
| Other | 848 | 190 | 98 | 36 | 497 | 27 |

[a]FC2 effects defined as having $p<0.01$ as determined from bootstrap hypothesis testing. [b]Catalytic effects defined as having $p<0.05$ as determined from bootstrap hypothesis testing, and $(k_{cat}/K_M^{chem})^{mutant} < (k_{cat}/K_M^{chem})^{WT}$.

**Table S13.** Number of measurable mutants with significant catalytic ($k_{cat}/K_M^{chem}$) effects, FC2 effects, and mutants with both catalytic and FC2 effects within different PafA regions.

| | Glycine | | | Valine | | | Contact [a] | |
|---|---|---|---|---|---|---|---|---|
| Residue | FC2 | FC3 | *p*-value | FC2 | FC3 | *p*-value | $Zn^{2+}$ | $O_n$ |
| N100 | <0.59 | – | $<1 \times 10^{-4}$ | – | <1.3 | $<1 \times 10^{-4}$ | – | 1 |
| K162 | – | – | – | – | 3.8 | $1 \times 10^{-4}$ | – | 2 |
| R164 | – | >3.3 | $<1 \times 10^{-4}$ | – | $>2.2 \times 10^{2}$ | $1 \times 10^{-4}$ | – | 2 |
| D305 | $<3.3 \times 10^{-3}$ | – | $4 \times 10^{-4}$ | – | – | – | 1 | – |
| H309 | – | 1.3 | $<1 \times 10^{-4}$ | – | 1.6 | $<1 \times 10^{-4}$ | 1 | – |
| H486 | – | – | – | – | – | – | 1 | – |
| D38 | <0.2 | – | $<1 \times 10^{-4}$ | – | – | – | 2 | – |
| T79 | – | | – | – | – | – | 2 | – |
| D352 | $3.8 \times 10^{-2}$ | - | $<1 \times 10^{-4}$ | – | – | – | 2 | – |
| H353 | – | – | – | 0.8 | – | 0.007 | 2 | – |

[a]Denotes interactions between each residue and either active site $Zn^{2+}$ ion (Zn1 denoted by "1" and Zn2 denoted by "2") or a substrate phosphoryl oxygen (O1 denoted by "1" and O2 denoted by "2").

**Table S14.** FC2 and FC3 effects for glycine and valine mutations of the PafA active site residues.

| | Overlap of mutants with FC4 effects (*p*<0.1) | | |
|---|---|---|---|
| **Parameter or FC** | ***p*-value Threshold** | **Count** | **FC4 Intersection Count (% of FC4 at *p*<0.1)** |
| $k_{cat}/K_M^{chem}$ | 0.05 | 185 | 38 (70) |
| FC1 | 0.01 | 181 | 29 (54) |
| FC2 | 0.01 | 331 | 36 (67) |
| FC3 | 0.01 | 73 | 8 (15) |
| FC4 | 0.1 | 54 | 54 (100) |

**Table S15.** Overlap of FC4 effects ($p < 0.1$) with those on other parameters and FCs.

| Term | Type | Definition | Description | *p*-value | Units | Eq. |
|---|---|---|---|---|---|---|
| $k_1$ | constant | $1.42 \times 10^6 = (k_{cat}/K_M)^{\text{cMUP, WT}}$ | Second order rate constant for substrate binding to PafA, assumed to be the apparent second order rate constant of cMUP hydrolysis by WT PafA (which is binding-limited) | – | $M^{-1}s^{-1}$ | – |
| $f_r$ | constant | $776 \pm 126$ $(k_{\text{chem},1}^{\text{cMUP}}/k_{\text{chem},1}^{\text{MeP}})$ | Fit relative first order rate constants of MeP vs. cMUP substrate hydrolysis on the enzyme | – | – | 30 |
| T-Effect | parameter | $\frac{[(k_{cat}/K_M)^{\text{mutant}}/(k_{cat}/K_M)^{\text{WT}}]^{23°C}}{[(k_{cat}/K_M)^{\text{mutant}}/(k_{cat}/K_M)^{\text{WT}}]^{37°C}}$ | Fold-change in WT-normalized activity upon expression at 23°C vs. 37°C | – | – | 9 |
| Zn-Effect | parameter | $\frac{[(k_{cat}/K_M)^{\text{mutant}}/(k_{cat}/K_M)^{\text{WT}}]^{1000\ \mu M\ Zn^{2+}}}{[(k_{cat}/K_M)^{\text{mutant}}/(k_{cat}/K_M)^{\text{WT}}]^{10\ \mu M\ Zn^{2+}}}$ | Fold-change in WT-normalized activity upon expression at 1000 μM $Zn^{2+}$ vs. 10 μM $Zn^{2+}$ | – | – | 10 |
| $f_a$ | parameter | $\frac{(f_r-1)(k_{cat}/K_M)^{\text{MeP,obs}}(k_{cat}/K_M)^{\text{cMUP,obs}}}{k_1[f_r(k_{cat}/K_M)^{\text{MeP,obs}}-(k_{cat}/K_M)^{\text{cMUP,obs}}]}$ | The fraction of apparent enzyme that is active (correctly folded) assuming a two-state model | < 0.01 | – | 32 |
| Catalytic Effect | parameter | $\frac{(k_{cat}/K_M)_{\text{mutant}}^{\text{MeP,chem}}}{(k_{cat}/K_M)_{\text{WT}}^{\text{MeP,chem}}}$ | Mutational effect on intrinsic catalytic efficiency (corrected for the $f_a$, the active fraction of enzyme) | < 0.05 | – | 36 |
| FC1 | FC | $\frac{[(k_{cat}/K_M)^{\text{MecMUP, mutant}}/(k_{cat}/K_M)^{\text{MecMUP,WT}}]}{[(k_{cat}/K_M)^{\text{MeP, mutant}}/(k_{cat}/K_M)^{\text{MeP,WT}}]}$ | The ratio of WT-normalized change in activity towards MecMUP to that of MeP. This value is $f_a$ independent. | < 0.01 | – | 12 & 13 |
| FC2 | FC | $(K_i\ P_i)^{\text{mutant}}/(K_i\ P_i)^{\text{WT}}$ where $(K_i\ P_i)^{\text{mutant}} < (K_i\ P_i)^{\text{WT}}$ | The fold-change of $K_i$ for $P_i$ from WT PafA, where $K_i$ for $P_i$ is tighter than that of the WT | < 0.01 | – | – |
| FC3 | FC | $(K_i\ P_i)^{\text{mutant}}/(K_i\ P_i)^{\text{WT}}$ where $(K_i\ P_i)^{\text{mutant}} \geq (K_i\ P_i)^{\text{WT}}$ | The fold-change of $K_i$ for $P_i$ from WT PafA, where $K_i$ for $P_i$ is weaker than that of the WT | < 0.01 | – | – |
| FC4 | FC | $\frac{k_{\text{on,Pi}}K_{\text{i,Pi}}k_{cat}^{\text{cMUP,obs}}}{f_a k_{\text{on,Pi}}K_{\text{i,Pi}}-k_{cat}^{\text{cMUP,obs}}}$ where $k_{\text{on,Pi}} = k_1$ | An estimate for $k_{\text{chem},2}$, the first-order rate constant for E–$P_i$ intermediate hydrolysis (a potential $k_{cat}$ rate limiting step). The definition was made using a rate law obtained by the method of net rate constants. | < 0.1 or < 0.05 | $s^{-1}$ | 40 |

**Table S16.** Definitions and descriptions of a subset constants, parameters, and functional components (FCs) discussed herein. Defined $p$-values thresholds are used to delineate mutants with strong evidence for effects on parameters and FCs.

### Captions for Auxiliary Supplementary Materials, Data S1 and S2

1. **Data S1.** Kinetic and thermodynamic parameter table, CSV (AuxiliarySupplementaryMaterials_S1.csv)

2. **Data S2.** Kinetic and thermodynamic parameter tables, PDF (AuxiliarySupplementaryMaterials_S2.pdf)

#### Captions for OSF Repository Data S1 to S14

The Open Science Foundation repository for this study is located at: https://osf.io/ez9xv/. The repository is organized into three components.

1. **Data S1.** Annotated WT PafA-eGFP PURExpress construct (Sequences/WT_PafA-eGFP_PURExpress.gb)
2. Per-experiment data files for all experiments: CSVs and PDFs of individual experiments measuring hydrolysis of cMUP, MeP, and MecMUP, and inhibition by phosphate. Each experiment has an associated CSV file and a PDF file containing progress curves, Michaelis-Menten curves, and fitted parameters for all chambers (*cf.* **figs. S19, S29, S43, and S50**). CSV and PDF files are located in the Per-Experiment Data and Per-Experiment Reports directories, respectively, within each OSF component.
   (a) **Data S2.** Active Site Mutant Experiments (/Active Site Mutant Experiments/...)
   (b) **Data S3.** cMUP (/cMUP Hydrolysis/...)
   (c) **Data S4.** MecMUP (/MecMUP Hydrolysis/...)
   (d) **Data S5.** MeP (/MeP Hydrolysis/...)
   (e) **Data S6.** Phosphate Inhibition (/Phosphate Inhibition/...)
   (f) **Data S7.** Varied Expression Experiments (/Varied Expression Experiments/...)
   (g) **Data S8.** Urea, Zinc, and pH Experiments (/Urea, Zinc, and pH)
3. Per-assay data files (Kinetic and Thermodynamic Data/Per-Assay Summaries): Aggregate summaries (CSV and PDF files) of kinetic and thermodynamic constants measured for each mutant, based on the replicates that passed all quality control filters (*cf.* **fig. S20**).
   (a) **Data S9.** Active Site Mutants (/Active Site Mutant Experiments/aggregate_KnownMutant_data.*)
   (b) **Data S10.** cMUP (/cMUP/aggregate_cMUP_data.*)
   (c) **Data S11.** MecMUP (/MecMUP/aggregate_MecMUP_data.*)
   (d) **Data S12.** MeP (/MeP/aggregate_MeP_data.*)
   (e) **Data S13.** Phosphate Inhibition (/Phosphate Inhibition/aggregate_KiPi_data.*)
   (f) **Data S14.** Varied Expression Experiments (/Varied Expression Experiments/...)
