## Supplementary Data 2 for "Revealing enzyme functional architecture via high-throughput microfluidic enzyme kinetics"

### Experimental Parameters and Functional Components (FCs)

HT-MEK

### Preface

#### Nomenclature

| Term | Description |
|---|---|
| **N/A** | FC does not apply to the measurement. Applicable to FC2 and FC3 only, which are mutually exclusive. |
| ***** | Parameter has an undefined limit (*i.e.*, the actual parameter value may be great or less than the estimate). These estimates are omitted. |
| **≤** | Estimate is an upper limit (prefix) |
| **≥** | Estimate is a lower limit (prefix) |
| **–** | No estimate was obtained |

#### Definitions

| Term | *p*-value Threshold | Definition |
|---|---|---|
| $f_a$ | 0.01 | $\frac{(f_r-1)\,(k_{cat}/K_M)^{MeP}\,(k_{cat}/K_M)^{cMUP}}{k_1[f_r(k_{cat}/K_M)^{MeP}-(k_{cat}/K_M)^{cMUP}]}$ where $f_r = 776$ |
| $(k_{cat}/K_M)^{MeP}_{chem.}$ | 0.05 | $(k_{cat}/K_M)^{MeP}/f_a$ |
| **FC1** | 0.01 | $\frac{(k_{cat}/K_M)^{MecMUP,mutant}/(k_{cat}/K_M)^{MecMUP,WT}}{(k_{cat}/K_M)^{MeP,mutant}/(k_{cat}/K_M)^{MeP,WT}}$ |
| **FC2** | 0.01 | $(K_i\,P_i)^{mutant}/(K_i\,P_i)^{WT}$ where $(K_i\,P_i)^{mutant} < (K_i\,P_i)^{WT}$ |
| **FC3** | 0.01 | $(K_i\,P_i)^{mutant}/(K_i\,P_i)^{WT}$ where $(K_i\,P_i)^{mutant} \geq (K_i\,P_i)^{WT}$ |
| **FC4** | 0.10 | $\frac{k_{on,Pi}\,K_{i,Pi}\,k_{cat}^{cMUP,obs}}{f_a\,k_{on,Pi}\,K_{i,Pi}-k_{cat}^{cMUP,obs}}$ |

#### Notes

1. **Bold** *p*-values are those below the defined *p*-value threshold for the corresponding parameter or FC
2. FCs without defined units are dimensionless quantities
3. *p*-values of magnitude 0 occur as a result of the bootstrapped hypothesis testing procedure, and are universally considered as below the defined *p*-value threshold for the corresponding parameter or FC
4. Active site variant estimates are derived from replicates on prints corresponding to Tiers I–III (Gly/Val libraries), not from the Active Site print
5. All estimates and *p*-values were rounded to two significant figures in tables below, though more significant figures are given in tables within the OSF Repository Data

### Active site variants

#### Fit parameters

| Variant | $k_{cat}$ cMUP ($s^{-1}$) | $k_{cat}$ cMUP $p$–value | $K_M$ cMUP (μM) | $K_M$ cMUP $p$–value | $k_{cat}/K_M$ cMUP ($M^{-1}s^{-1}$) | $k_{cat}/K_M$ cMUP $p$–value | $k_{cat}/K_M$ MeP ($M^{-1}s^{-1}$) | $k_{cat}/K_M$ MeP $p$–value | $k_{cat}/K_M$ MecMUP ($M^{-1}s^{-1}$) | $k_{cat}/K_M$ MecMUP $p$–value | $K_i$ $PO_4$ (μM) | $K_i$ $PO_4$ $p$–value |
|---|---|---|---|---|---|---|---|---|---|---|---|---|
| T79S | 1.1 | **0** | ≤ 5.0 | **0** | ≥ $2.2\times10^{5}$ | **0** | $3.2\times10^{4}$ | **0** | ≥ $1.2\times10^{1}$ | **$4.1\times10^{-3}$** | ≤ $1.3\times10^{1}$ | **0** |
| N100A | $4.0\times10^{-1}$ | **0** | ≤ 5.0 | **0** | ≥ $7.7\times10^{4}$ | **0** | $2.2\times10^{3}$ | **$1.\times10^{-4}$** | ≤ 1.3 | **0** | ≤ $8.8\times10^{2}$ | **0** |
| K162A | $1.8\times10^{-2}$ | **0** | $5.5\times10^{1}$ | **$1.1\times10^{-3}$** | $3.4\times10^{2}$ | **0** | ≤ $1.2\times10^{4}$ | **0** | $3.7\times10^{1}$ | **0** | – | – |
| R164A | 5.7 | **0** | $2.2\times10^{1}$ | **0** | $2.6\times10^{5}$ | **0** | ≤ $7.2\times10^{3}$ | **0** | $1.1\times10^{2}$ | **0** | ≥ $2.9\times10^{3}$ | **0** |
| N100A/R164A | $6.8\times10^{-1}$ | **0** | $1.8\times10^{2}$ | **0** | $3.7\times10^{3}$ | **0** | ≤ $7.0\times10^{3}$ | **0** | ≤ 1.3 | **0** | $5.6\times10^{3}$ | **0** |
| WT | $1.2\times10^{2}$ | $9.7\times10^{-1}$ | $8.5\times10^{1}$ | 1.0 | $1.4\times10^{6}$ | $9.8\times10^{-1}$ | $6.1\times10^{5}$ | $8.8\times10^{-1}$ | $1.6\times10^{1}$ | $9.8\times10^{-1}$ | $6.1\times10^{2}$ | $8.7\times10^{-1}$ |

#### Additional parameters and FCs

| Variant | $f_a$ | $f_a$ $p$–value | $(k_{cat}/K_M)^{MeP}_{chem.}$ ($M^{-1}s^{-1}$) | $(k_{cat}/K_M)^{MeP}_{chem.}$ $p$–value | FC1 | FC1 $p$–value | FC2 | FC3 | FC2/3 $p$–value | FC4 ($s^{-1}$) | FC4 $p$–value |
|---|---|---|---|---|---|---|---|---|---|---|---|
| T79S | $1.6\times10^{-1}$ | **0** | $2.1\times10^{5}$ | $1.2\times10^{-1}$ | ≥ $1.4\times10^{1}$ | **$5.1\times10^{-4}$** | ≤ $2.1\times10^{-2}$ | N/A | **0** | ≥ $1.1\times10^{1}$ | $6.8\times10^{-1}$ |
| N100A | $5.6\times10^{-2}$ | **0** | $4.0\times10^{4}$ | **$6.0\times10^{-3}$** | – | – | N/A | ≤ 1.4 | **0** | ≥ 7.2 | **$5.0\times10^{-3}$** |
| K162A | ≤ 1.0 | 1.0 | ≤ $3.4\times10^{2}$ | **0** | ≤ $1.1\times10^{2}$ | **0** | – | – | – | – | – |
| R164A | ≤ $1.9\times10^{-1}$ | **0** | ≤ $3.7\times10^{4}$ | **0** | ≤ $5.8\times10^{2}$ | **0** | N/A | ≥ 4.8 | **0** | * | $8.0\times10^{-2}$ |
| N100A/R164A | ≤ 1.0 | 1.0 | ≤ $3.7\times10^{3}$ | **0** | – | – | N/A | 9.1 | **0** | ≥ $2.7\times10^{2}$ | **$5.0\times10^{-3}$** |
| WT | 1.0 | 1.0 | $6.1\times10^{5}$ | 1.0 | 1.0 | 1.0 | N/A | 1.0 | $8.7\times10^{-1}$ | $1.4\times10^{2}$ | 1.0 |

#### Valine and glycine library variants

##### Fit parameters

| Variant | $k_{cat}$ cMUP ($s^{-1}$) | $k_{cat}$ cMUP *p*–value | $K_M$ cMUP (μM) | $K_M$ cMUP *p*–value | $k_{cat}/K_M$ cMUP ($M^{-1}s^{-1}$) | $k_{cat}/K_M$ cMUP *p*–value | $k_{cat}/K_M$ MeP ($M^{-1}s^{-1}$) | $k_{cat}/K_M$ MeP *p*–value | $k_{cat}/K_M$ MecMUP ($M^{-1}s^{-1}$) | $k_{cat}/K_M$ MecMUP *p*–value | $K_i$ $PO_4$ (μM) | $K_i$ $PO_4$ *p*–value |
|---|---|---|---|---|---|---|---|---|---|---|---|---|
| Q21G | $2.2\times10^{2}$ | **$6.0\times10^{-4}$** | $9.0\times10^{1}$ | $6.9\times10^{-1}$ | $2.0\times10^{6}$ | $1.6\times10^{-1}$ | $6.1\times10^{5}$ | $9.8\times10^{-1}$ | $1.5\times10^{1}$ | $7.5\times10^{-1}$ | $6.4\times10^{2}$ | $3.8\times10^{-1}$ |
| Q21V | $2.1\times10^{2}$ | **$1.\times10^{-4}$** | $9.2\times10^{1}$ | $4.4\times10^{-1}$ | $2.2\times10^{6}$ | $1.2\times10^{-2}$ | $4.2\times10^{5}$ | $6.9\times10^{-2}$ | $1.9\times10^{1}$ | $5.2\times10^{-1}$ | $6.6\times10^{2}$ | $2.0\times10^{-1}$ |
| K22G | $7.4\times10^{1}$ | $9.9\times10^{-2}$ | $6.9\times10^{1}$ | $2.3\times10^{-1}$ | $1.6\times10^{6}$ | $6.8\times10^{-1}$ | $5.2\times10^{5}$ | $4.3\times10^{-1}$ | $1.3\times10^{1}$ | $4.1\times10^{-1}$ | $5.3\times10^{2}$ | $1.4\times10^{-2}$ |
| K22V | $1.3\times10^{2}$ | $7.6\times10^{-1}$ | $1.0\times10^{2}$ | $9.4\times10^{-2}$ | $1.2\times10^{6}$ | $3.9\times10^{-1}$ | $4.0\times10^{5}$ | $9.8\times10^{-2}$ | $1.5\times10^{1}$ | $7.2\times10^{-1}$ | $7.1\times10^{2}$ | $1.6\times10^{-2}$ |
| T23G | $1.1\times10^{2}$ | $5.0\times10^{-1}$ | $7.7\times10^{1}$ | $4.1\times10^{-1}$ | $1.4\times10^{6}$ | $9.4\times10^{-1}$ | $6.9\times10^{5}$ | $6.0\times10^{-1}$ | $2.0\times10^{1}$ | $1.4\times10^{-1}$ | $6.3\times10^{2}$ | $5.1\times10^{-1}$ |
| T23V | $1.9\times10^{2}$ | $1.7\times10^{-2}$ | $1.1\times10^{2}$ | $7.0\times10^{-2}$ | $1.9\times10^{6}$ | $1.7\times10^{-1}$ | $3.3\times10^{5}$ | $2.4\times10^{-2}$ | $1.9\times10^{1}$ | $4.5\times10^{-1}$ | $6.4\times10^{2}$ | $5.6\times10^{-1}$ |
| N24G | $1.0\times10^{2}$ | $4.4\times10^{-1}$ | $6.2\times10^{1}$ | $9.4\times10^{-2}$ | $1.8\times10^{6}$ | $4.1\times10^{-1}$ | $5.5\times10^{5}$ | $4.7\times10^{-1}$ | $1.3\times10^{1}$ | $3.4\times10^{-1}$ | $4.8\times10^{2}$ | **$2.0\times10^{-3}$** |
| N24V | $1.3\times10^{2}$ | $3.9\times10^{-1}$ | $9.0\times10^{1}$ | $5.2\times10^{-1}$ | $1.4\times10^{6}$ | $9.4\times10^{-1}$ | $4.2\times10^{5}$ | $1.7\times10^{-1}$ | $2.0\times10^{1}$ | $3.9\times10^{-1}$ | $6.5\times10^{2}$ | $2.3\times10^{-1}$ |
| A25G | $2.1\times10^{2}$ | **$1.9\times10^{-3}$** | $8.6\times10^{1}$ | $9.3\times10^{-1}$ | $2.7\times10^{6}$ | **$2.0\times10^{-3}$** | $8.5\times10^{5}$ | $7.1\times10^{-2}$ | $2.4\times10^{1}$ | $1.6\times10^{-2}$ | $5.8\times10^{2}$ | $1.6\times10^{-1}$ |
| A25V | $2.2\times10^{2}$ | **0** | $9.7\times10^{1}$ | $2.0\times10^{-1}$ | $2.0\times10^{6}$ | $7.3\times10^{-2}$ | $6.7\times10^{5}$ | $6.4\times10^{-1}$ | $1.8\times10^{1}$ | $4.9\times10^{-1}$ | $6.6\times10^{2}$ | $1.6\times10^{-1}$ |
| V26A | $5.1\times10^{1}$ | **$2.9\times10^{-3}$** | $7.9\times10^{1}$ | $4.8\times10^{-1}$ | $7.3\times10^{5}$ | $4.2\times10^{-2}$ | $1.9\times10^{5}$ | **$1.9\times10^{-3}$** | 6.5 | $1.2\times10^{-2}$ | $6.0\times10^{2}$ | $6.3\times10^{-1}$ |
| V26G | $2.1\times10^{1}$ | **0** | $5.7\times10^{1}$ | $1.2\times10^{-2}$ | $2.6\times10^{5}$ | **$1.2\times10^{-3}$** | $3.9\times10^{4}$ | **0** | 3.8 | **$2.0\times10^{-4}$** | $5.4\times10^{2}$ | $1.7\times10^{-2}$ |
| P27G | $5.9\times10^{1}$ | $3.7\times10^{-2}$ | $7.0\times10^{1}$ | $2.5\times10^{-1}$ | $4.3\times10^{5}$ | $1.3\times10^{-2}$ | $3.0\times10^{5}$ | $1.2\times10^{-2}$ | $1.4\times10^{1}$ | $5.0\times10^{-1}$ | $5.5\times10^{2}$ | $4.3\times10^{-2}$ |
| P27V | $1.4\times10^{2}$ | $4.0\times10^{-1}$ | $9.2\times10^{1}$ | $4.3\times10^{-1}$ | $1.5\times10^{6}$ | $7.9\times10^{-1}$ | $2.5\times10^{5}$ | $1.7\times10^{-2}$ | $1.3\times10^{1}$ | $5.2\times10^{-1}$ | $6.2\times10^{2}$ | $6.9\times10^{-1}$ |
| R28G | $4.2\times10^{1}$ | $3.5\times10^{-2}$ | $3.8\times10^{1}$ | $1.6\times10^{-2}$ | $1.1\times10^{6}$ | $5.3\times10^{-1}$ | $3.6\times10^{5}$ | $4.1\times10^{-2}$ | 9.8 | $3.8\times10^{-2}$ | $4.4\times10^{2}$ | **$1.3\times10^{-3}$** |
| R28V | $7.6\times10^{1}$ | $3.0\times10^{-2}$ | $7.8\times10^{1}$ | $4.6\times10^{-1}$ | $9.3\times10^{5}$ | $1.3\times10^{-1}$ | $1.3\times10^{5}$ | **$1.6\times10^{-3}$** | $1.0\times10^{1}$ | $1.1\times10^{-1}$ | $6.2\times10^{2}$ | $9.1\times10^{-1}$ |
| P29G | $1.1\times10^{1}$ | **0** | $6.9\times10^{1}$ | $8.9\times10^{-2}$ | $1.9\times10^{5}$ | **$1.\times10^{-4}$** | $2.3\times10^{4}$ | **$5.0\times10^{-4}$** | ≤ 1.2 | **0** | $4.9\times10^{2}$ | **0** |
| P29V | $1.4\times10^{1}$ | **$3.0\times10^{-4}$** | $7.6\times10^{1}$ | $4.4\times10^{-1}$ | $2.0\times10^{5}$ | **$4.5\times10^{-3}$** | $1.3\times10^{5}$ | **$1.1\times10^{-3}$** | ≤ 1.9 | **$1.\times10^{-4}$** | $4.4\times10^{2}$ | **$1.\times10^{-4}$** |
| K30G | $5.0\times10^{1}$ | **$1.8\times10^{-3}$** | $6.2\times10^{1}$ | $3.0\times10^{-2}$ | $1.2\times10^{6}$ | $5.1\times10^{-1}$ | $2.6\times10^{5}$ | $1.1\times10^{-2}$ | $1.1\times10^{1}$ | $9.8\times10^{-2}$ | $5.3\times10^{2}$ | $1.4\times10^{-2}$ |
| K30V | $8.7\times10^{1}$ | $1.5\times10^{-1}$ | $9.1\times10^{1}$ | $5.7\times10^{-1}$ | $9.5\times10^{5}$ | $2.0\times10^{-1}$ | $1.3\times10^{5}$ | **$1.\times10^{-4}$** | 7.4 | $3.4\times10^{-2}$ | $5.4\times10^{2}$ | $4.5\times10^{-2}$ |
| L31G | $2.3\times10^{-1}$ | **0** | $5.0\times10^{1}$ | **$5.3\times10^{-3}$** | $5.0\times10^{3}$ | **$7.0\times10^{-4}$** | ≤ $1.4\times10^{3}$ | **0** | ≤ $9.9\times10^{-1}$ | **0** | – | – |
| L31V | $1.3\times10^{2}$ | $6.0\times10^{-1}$ | $8.6\times10^{1}$ | $8.7\times10^{-1}$ | $1.4\times10^{6}$ | $8.4\times10^{-1}$ | $2.9\times10^{5}$ | **$6.1\times10^{-3}$** | $1.2\times10^{1}$ | $2.6\times10^{-1}$ | $6.2\times10^{2}$ | $7.4\times10^{-1}$ |
| V32A | $1.6\times10^{2}$ | $6.9\times10^{-2}$ | $1.1\times10^{2}$ | $6.9\times10^{-2}$ | $1.4\times10^{6}$ | $9.3\times10^{-1}$ | $2.5\times10^{5}$ | $3.1\times10^{-2}$ | $1.6\times10^{1}$ | $9.5\times10^{-1}$ | $6.2\times10^{2}$ | $9.7\times10^{-1}$ |
| V32G | $5.1\times10^{1}$ | $5.7\times10^{-2}$ | $6.4\times10^{1}$ | $2.0\times10^{-1}$ | $7.3\times10^{5}$ | $2.0\times10^{-1}$ | $1.7\times10^{5}$ | **$8.2\times10^{-3}$** | 5.4 | **$1.1\times10^{-3}$** | $4.6\times10^{2}$ | **$2.1\times10^{-3}$** |
| V33A | $1.8\times10^{2}$ | $2.6\times10^{-2}$ | $9.8\times10^{1}$ | $2.5\times10^{-1}$ | $1.9\times10^{6}$ | $2.1\times10^{-1}$ | $2.1\times10^{5}$ | **$1.0\times10^{-3}$** | $1.1\times10^{1}$ | $3.7\times10^{-1}$ | $6.0\times10^{2}$ | $4.4\times10^{-1}$ |
| V33G | $3.4\times10^{1}$ | **$1.\times10^{-4}$** | $5.9\times10^{1}$ | $1.6\times10^{-2}$ | $5.5\times10^{5}$ | $1.0\times10^{-2}$ | $2.3\times10^{5}$ | **$5.6\times10^{-3}$** | 6.8 | $1.4\times10^{-2}$ | $5.0\times10^{2}$ | **$2.0\times10^{-4}$** |
| G34A | $9.8\times10^{1}$ | $2.5\times10^{-1}$ | $7.4\times10^{1}$ | $2.2\times10^{-1}$ | $1.2\times10^{6}$ | $4.0\times10^{-1}$ | $7.0\times10^{5}$ | $4.6\times10^{-1}$ | $1.6\times10^{1}$ | $9.6\times10^{-1}$ | $6.4\times10^{2}$ | $4.6\times10^{-1}$ |
| G34V | $1.9\times10^{1}$ | **$2.0\times10^{-4}$** | $8.0\times10^{1}$ | $6.1\times10^{-1}$ | $2.7\times10^{5}$ | **$3.2\times10^{-3}$** | ≤ $3.4\times10^{4}$ | **0** | ≤ 1.9 | **0** | $4.4\times10^{2}$ | **0** |
| L35G | 1.9 | **$2.5\times10^{-3}$** | $8.3\times10^{1}$ | $8.8\times10^{-1}$ | $2.4\times10^{4}$ | **$2.7\times10^{-3}$** | ≤ $1.6\times10^{3}$ | **0** | ≤ 1.3 | **0** | $8.8\times10^{2}$ | **$1.0\times10^{-3}$** |
| L35V | $1.5\times10^{2}$ | $7.5\times10^{-2}$ | $9.1\times10^{1}$ | $5.4\times10^{-1}$ | $2.0\times10^{6}$ | $6.0\times10^{-2}$ | $4.8\times10^{5}$ | $1.2\times10^{-1}$ | $1.7\times10^{1}$ | $8.1\times10^{-1}$ | $6.7\times10^{2}$ | $3.8\times10^{-2}$ |
| V36A | $1.3\times10^{2}$ | $4.9\times10^{-1}$ | $8.0\times10^{1}$ | $4.8\times10^{-1}$ | $1.9\times10^{6}$ | $1.9\times10^{-1}$ | $3.6\times10^{5}$ | $4.3\times10^{-2}$ | $1.3\times10^{1}$ | $5.7\times10^{-1}$ | $5.1\times10^{2}$ | $5.2\times10^{-2}$ |
| V36G | $1.8\times10^{1}$ | **0** | $2.3\times10^{1}$ | **0** | $7.8\times10^{5}$ | $1.8\times10^{-2}$ | $1.4\times10^{5}$ | **$1.4\times10^{-3}$** | 5.3 | **0** | $2.0\times10^{2}$ | **0** |
| V37A | $1.9\times10^{2}$ | **$2.4\times10^{-3}$** | $9.9\times10^{1}$ | $1.7\times10^{-1}$ | $1.8\times10^{6}$ | $2.6\times10^{-1}$ | $3.6\times10^{5}$ | $4.6\times10^{-2}$ | $1.3\times10^{1}$ | $5.2\times10^{-1}$ | $7.2\times10^{2}$ | $1.1\times10^{-2}$ |
| V37G | $9.3\times10^{1}$ | $1.7\times10^{-1}$ | $7.0\times10^{1}$ | $2.3\times10^{-1}$ | $1.2\times10^{6}$ | $5.5\times10^{-1}$ | $2.0\times10^{5}$ | **$4.6\times10^{-3}$** | 6.1 | **$1.7\times10^{-3}$** | $4.8\times10^{2}$ | **$3.0\times10^{-4}$** |
| D38G | 4.2 | **0** | ≤ 5.0 | **0** | ≥ $7.2\times10^{5}$ | **$5.0\times10^{-3}$** | ≤ $6.0\times10^{3}$ | **0** | 2.1 | **$3.0\times10^{-4}$** | ≤ $1.3\times10^{2}$ | **0** |
| D38V | – | – | – | – | ≤ $6.0\times10^{2}$ | **0** | ≤ $9.4\times10^{2}$ | **0** | ≤ 1.3 | **0** | – | – |
| Q39G | $7.0\times10^{1}$ | $4.2\times10^{-2}$ | $3.8\times10^{1}$ | **$5.0\times10^{-4}$** | $1.8\times10^{6}$ | $3.3\times10^{-1}$ | $3.6\times10^{4}$ | **$1.\times10^{-4}$** | 2.6 | **$7.0\times10^{-4}$** | $1.3\times10^{3}$ | **$1.\times10^{-4}$** |
| Q39V | 1.9 | **0** | $4.5\times10^{1}$ | **$3.0\times10^{-4}$** | $4.6\times10^{4}$ | **0** | ≤ $1.6\times10^{3}$ | **0** | ≤ 1.3 | **$1.\times10^{-4}$** | $2.6\times10^{2}$ | **$3.0\times10^{-4}$** |
| M40G | $1.9\times10^{1}$ | **0** | $6.1\times10^{1}$ | $2.4\times10^{-2}$ | $3.1\times10^{5}$ | **$1.0\times10^{-3}$** | $1.2\times10^{5}$ | **$1.9\times10^{-3}$** | 4.9 | **$8.0\times10^{-4}$** | $3.8\times10^{2}$ | **0** |
| M40V | 3.6 | **0** | $5.9\times10^{1}$ | **$2.0\times10^{-3}$** | $7.0\times10^{4}$ | **0** | $3.1\times10^{3}$ | **$5.0\times10^{-4}$** | ≤ 1.3 | **$1.\times10^{-4}$** | $3.2\times10^{2}$ | **$1.\times10^{-4}$** |
| R41G | $2.0\times10^{2}$ | $2.0\times10^{-2}$ | $4.7\times10^{1}$ | $3.1\times10^{-2}$ | $2.9\times10^{6}$ | **$8.9\times10^{-3}$** | $4.3\times10^{5}$ | $1.8\times10^{-1}$ | $1.6\times10^{1}$ | $9.3\times10^{-1}$ | $5.5\times10^{2}$ | $1.1\times10^{-1}$ |
| R41V | $5.6\times10^{1}$ | **$1.7\times10^{-3}$** | $5.0\times10^{1}$ | **$6.0\times10^{-4}$** | $1.2\times10^{6}$ | $4.4\times10^{-1}$ | $2.2\times10^{5}$ | **$5.9\times10^{-3}$** | ≤ 2.3 | **$4.0\times10^{-4}$** | $3.5\times10^{2}$ | **$3.0\times10^{-4}$** |
| W42G | $4.8\times10^{1}$ | **$9.3\times10^{-3}$** | $4.1\times10^{1}$ | **$1.9\times10^{-3}$** | $1.4\times10^{6}$ | $9.4\times10^{-1}$ | $2.6\times10^{5}$ | $1.1\times10^{-2}$ | 6.2 | **$2.3\times10^{-3}$** | $3.8\times10^{2}$ | **$1.\times10^{-4}$** |
| W42V | $1.8\times10^{2}$ | **$7.6\times10^{-3}$** | $7.8\times10^{1}$ | $4.5\times10^{-1}$ | $2.3\times10^{6}$ | **$9.0\times10^{-3}$** | $7.6\times10^{5}$ | $2.6\times10^{-1}$ | $1.1\times10^{1}$ | $1.0\times10^{-1}$ | $5.0\times10^{2}$ | $2.3\times10^{-2}$ |
| D43G | 8.8 | **0** | $3.5\times10^{1}$ | **0** | $2.5\times10^{5}$ | **0** | $1.8\times10^{4}$ | **$4.0\times10^{-4}$** | 1.4 | **0** | $3.0\times10^{2}$ | **0** |
| D43V | 7.8 | **0** | $3.8\times10^{1}$ | **0** | $2.4\times10^{5}$ | **$1.\times10^{-4}$** | $1.5\times10^{4}$ | **$6.0\times10^{-4}$** | 2.0 | **0** | $2.8\times10^{2}$ | **0** |
| Y44G | $6.2\times10^{1}$ | $2.3\times10^{-2}$ | $7.4\times10^{1}$ | $3.1\times10^{-1}$ | $8.6\times10^{5}$ | $1.4\times10^{-1}$ | $2.6\times10^{5}$ | $1.3\times10^{-2}$ | 5.7 | **$5.2\times10^{-3}$** | $6.9\times10^{2}$ | $2.2\times10^{-2}$ |
| Y44V | $1.0\times10^{2}$ | $3.1\times10^{-1}$ | $7.2\times10^{1}$ | $1.9\times10^{-1}$ | $1.6\times10^{6}$ | $5.6\times10^{-1}$ | $2.6\times10^{5}$ | $1.0\times10^{-2}$ | 9.1 | $8.6\times10^{-2}$ | $6.2\times10^{2}$ | $5.8\times10^{-1}$ |
| L45G | $6.5\times10^{1}$ | $6.3\times10^{-2}$ | $6.2\times10^{1}$ | $9.0\times10^{-2}$ | $1.0\times10^{6}$ | $4.0\times10^{-1}$ | $8.3\times10^{4}$ | **$1.4\times10^{-3}$** | 5.8 | **$6.4\times10^{-3}$** | $5.4\times10^{2}$ | $3.8\times10^{-2}$ |
| L45V | $4.9\times10^{1}$ | **$1.7\times10^{-3}$** | $8.9\times10^{1}$ | $6.8\times10^{-1}$ | $6.2\times10^{5}$ | $1.5\times10^{-2}$ | $1.3\times10^{5}$ | **$6.3\times10^{-3}$** | 6.6 | **$2.8\times10^{-3}$** | $5.7\times10^{2}$ | $1.9\times10^{-1}$ |
| Y46G | $2.1\times10^{1}$ | **$3.0\times10^{-4}$** | $4.4\times10^{1}$ | **$1.4\times10^{-3}$** | $4.0\times10^{5}$ | **$6.1\times10^{-3}$** | $2.7\times10^{5}$ | $1.1\times10^{-2}$ | ≤ $3.2\times10^{-1}$ | **0** | $3.4\times10^{2}$ | **$1.\times10^{-4}$** |
| Y46V | $1.2\times10^{2}$ | $9.3\times10^{-1}$ | $8.5\times10^{1}$ | $9.5\times10^{-1}$ | $1.6\times10^{6}$ | $6.5\times10^{-1}$ | $3.4\times10^{5}$ | $1.7\times10^{-2}$ | $1.5\times10^{1}$ | $7.9\times10^{-1}$ | $5.6\times10^{2}$ | $6.6\times10^{-2}$ |
| R47G | $4.0\times10^{1}$ | $1.0\times10^{-2}$ | $5.1\times10^{1}$ | $1.7\times10^{-2}$ | $8.4\times10^{5}$ | $1.8\times10^{-1}$ | $1.8\times10^{5}$ | $2.7\times10^{-2}$ | 6.9 | **$3.1\times10^{-3}$** | $4.8\times10^{2}$ | **0** |
| R47V | $1.3\times10^{2}$ | $5.5\times10^{-1}$ | $8.0\times10^{1}$ | $5.6\times10^{-1}$ | $1.7\times10^{6}$ | $3.9\times10^{-1}$ | $9.8\times10^{4}$ | **$8.0\times10^{-4}$** | $1.2\times10^{1}$ | $2.6\times10^{-1}$ | $5.9\times10^{2}$ | $3.5\times10^{-1}$ |
| Y48G | $1.3\times10^{1}$ | **0** | $4.9\times10^{1}$ | **0** | $3.1\times10^{5}$ | **$2.0\times10^{-4}$** | $3.1\times10^{4}$ | **$2.0\times10^{-4}$** | 1.9 | **$8.0\times10^{-4}$** | $4.1\times10^{2}$ | **0** |
| Y48V | $2.6\times10^{1}$ | **0** | $9.4\times10^{1}$ | $4.2\times10^{-1}$ | $2.7\times10^{5}$ | **$1.6\times10^{-3}$** | ≤ $1.2\times10^{5}$ | **0** | ≤ 1.9 | **0** | $6.0\times10^{2}$ | $5.0\times10^{-1}$ |
| Y49G | $9.9\times10^{1}$ | $3.3\times10^{-1}$ | $6.8\times10^{1}$ | $1.4\times10^{-1}$ | $1.5\times10^{6}$ | $7.6\times10^{-1}$ | $3.0\times10^{5}$ | $2.6\times10^{-2}$ | 5.7 | **$9.0\times10^{-4}$** | $5.1\times10^{2}$ | **$2.3\times10^{-3}$** |
| Y49V | $9.8\times10^{1}$ | $1.7\times10^{-1}$ | $7.6\times10^{1}$ | $3.5\times10^{-1}$ | $1.5\times10^{6}$ | $8.0\times10^{-1}$ | $1.9\times10^{5}$ | **$4.2\times10^{-3}$** | $1.3\times10^{1}$ | $3.6\times10^{-1}$ | $5.6\times10^{2}$ | $1.0\times10^{-1}$ |
| S50G | $1.4\times10^{2}$ | $3.8\times10^{-1}$ | $7.3\times10^{1}$ | $2.5\times10^{-1}$ | $2.1\times10^{6}$ | $4.2\times10^{-2}$ | ≤ $3.0\times10^{5}$ | $1.1\times10^{-1}$ | $2.1\times10^{1}$ | $1.1\times10^{-1}$ | $6.7\times10^{2}$ | $7.8\times10^{-2}$ |
| S50V | $1.7\times10^{2}$ | $1.5\times10^{-2}$ | $9.6\times10^{1}$ | $2.5\times10^{-1}$ | $1.8\times10^{6}$ | $1.9\times10^{-1}$ | $2.3\times10^{5}$ | **$5.8\times10^{-3}$** | $1.6\times10^{1}$ | $9.6\times10^{-1}$ | $6.4\times10^{2}$ | $5.0\times10^{-1}$ |
| K51G | $6.3\times10^{1}$ | $1.0\times10^{-2}$ | $6.7\times10^{1}$ | $9.8\times10^{-2}$ | $8.6\times10^{5}$ | $9.3\times10^{-2}$ | $7.0\times10^{4}$ | **0** | 9.4 | $3.5\times10^{-2}$ | $6.5\times10^{2}$ | $2.6\times10^{-1}$ |
| K51V | $5.6\times10^{1}$ | **$4.4\times10^{-3}$** | $8.3\times10^{1}$ | $7.4\times10^{-1}$ | $8.2\times10^{5}$ | $7.2\times10^{-2}$ | $1.3\times10^{5}$ | **$1.1\times10^{-3}$** | 7.5 | $1.1\times10^{-2}$ | $5.4\times10^{2}$ | $6.9\times10^{-2}$ |
| Y52G | $6.2\times10^{1}$ | **$2.5\times10^{-3}$** | $7.5\times10^{1}$ | $2.1\times10^{-1}$ | $8.4\times10^{5}$ | $4.6\times10^{-2}$ | $1.2\times10^{5}$ | **$4.0\times10^{-4}$** | 3.6 | **$4.0\times10^{-4}$** | $5.7\times10^{2}$ | $4.7\times10^{-2}$ |
| Y52V | $2.5\times10^{2}$ | **0** | $1.0\times10^{2}$ | $4.9\times10^{-2}$ | $2.1\times10^{6}$ | $3.4\times10^{-2}$ | $1.9\times10^{5}$ | **$1.\times10^{-4}$** | $1.5\times10^{1}$ | $8.3\times10^{-1}$ | $6.6\times10^{2}$ | $6.7\times10^{-2}$ |
| G53A | $9.2\times10^{1}$ | $1.3\times10^{-1}$ | $6.7\times10^{1}$ | $9.1\times10^{-2}$ | $1.2\times10^{6}$ | $6.0\times10^{-1}$ | ≤ $3.8\times10^{5}$ | $2.1\times10^{-1}$ | $1.9\times10^{1}$ | $4.5\times10^{-1}$ | $6.3\times10^{2}$ | $4.6\times10^{-1}$ |
| G53V | $6.2\times10^{1}$ | $1.8\times10^{-2}$ | $9.6\times10^{1}$ | $3.4\times10^{-1}$ | $6.1\times10^{5}$ | $3.0\times10^{-2}$ | $4.7\times10^{5}$ | $2.4\times10^{-1}$ | ≤ 1.9 | **$4.0\times10^{-4}$** | $6.0\times10^{2}$ | $6.8\times10^{-1}$ |
| E54G | $1.5\times10^{2}$ | $2.3\times10^{-1}$ | $8.5\times10^{1}$ | $9.6\times10^{-1}$ | $1.9\times10^{6}$ | $2.0\times10^{-1}$ | $4.5\times10^{5}$ | $2.0\times10^{-1}$ | $1.6\times10^{1}$ | $8.3\times10^{-1}$ | $6.0\times10^{2}$ | $6.9\times10^{-1}$ |
| E54V | $2.1\times10^{2}$ | **$3.0\times10^{-4}$** | $1.0\times10^{2}$ | $7.1\times10^{-2}$ | $2.0\times10^{6}$ | $8.6\times10^{-2}$ | $5.6\times10^{5}$ | $5.0\times10^{-1}$ | $1.5\times10^{1}$ | $8.2\times10^{-1}$ | $7.0\times10^{2}$ | $3.5\times10^{-2}$ |
| G55A | $9.0\times10^{1}$ | $1.5\times10^{-1}$ | $6.5\times10^{1}$ | $9.7\times10^{-2}$ | $2.1\times10^{6}$ | $5.2\times10^{-2}$ | $5.3\times10^{5}$ | $5.9\times10^{-1}$ | $2.2\times10^{1}$ | $7.8\times10^{-2}$ | $6.4\times10^{2}$ | $5.3\times10^{-1}$ |
| G55V | $2.1\times10^{2}$ | **$7.3\times10^{-3}$** | $9.7\times10^{1}$ | $4.1\times10^{-1}$ | $2.4\times10^{6}$ | $2.5\times10^{-2}$ | $2.1\times10^{5}$ | $1.7\times10^{-2}$ | $1.9\times10^{1}$ | $4.4\times10^{-1}$ | $6.6\times10^{2}$ | $2.4\times10^{-1}$ |
| G56A | $5.8\times10^{1}$ | $4.0\times10^{-2}$ | $7.9\times10^{1}$ | $6.7\times10^{-1}$ | $7.2\times10^{5}$ | $1.1\times10^{-1}$ | $1.3\times10^{5}$ | **$2.1\times10^{-3}$** | 4.1 | **$2.0\times10^{-4}$** | $6.2\times10^{2}$ | $8.1\times10^{-1}$ |
| G56V | $7.3\times10^{-2}$ | $4.2\times10^{-2}$ | $3.7\times10^{1}$ | $7.4\times10^{-2}$ | $2.0\times10^{3}$ | $6.8\times10^{-2}$ | ≤ $1.8\times10^{3}$ | **0** | ≤ 1.7 | **0** | – | – |
| F57G | $9.1\times10^{-2}$ | **$6.0\times10^{-4}$** | $4.6\times10^{1}$ | **$8.6\times10^{-3}$** | $2.2\times10^{3}$ | **$1.8\times10^{-3}$** | ≤ $1.2\times10^{3}$ | **0** | ≤ 1.4 | **0** | – | – |
| F57V | $1.6\times10^{2}$ | $4.5\times10^{-2}$ | $7.8\times10^{1}$ | $4.1\times10^{-1}$ | $2.0\times10^{6}$ | $8.5\times10^{-2}$ | $1.2\times10^{5}$ | **$7.4\times10^{-3}$** | $1.1\times10^{1}$ | $2.5\times10^{-1}$ | $6.1\times10^{2}$ | $6.8\times10^{-1}$ |
| K58G | $3.9\times10^{1}$ | **$9.2\times10^{-3}$** | $5.1\times10^{1}$ | $1.9\times10^{-2}$ | $8.7\times10^{5}$ | $2.0\times10^{-1}$ | $1.1\times10^{5}$ | **$2.0\times10^{-3}$** | 7.8 | **$5.3\times10^{-3}$** | $4.6\times10^{2}$ | **$3.0\times10^{-4}$** |

| Variant | $k_{cat}$ cMUP ($s^{-1}$) | $k_{cat}$ cMUP $p$–value | $K_M$ cMUP (μM) | $K_M$ cMUP $p$–value | $k_{cat}/K_M$ cMUP ($M^{-1}s^{-1}$) | $k_{cat}/K_M$ cMUP $p$–value | $k_{cat}/K_M$ MeP ($M^{-1}s^{-1}$) | $k_{cat}/K_M$ MeP $p$–value | $k_{cat}/K_M$ MecMUP ($M^{-1}s^{-1}$) | $k_{cat}/K_M$ MecMUP $p$–value | $K_i$ $PO_4$ (μM) | $K_i$ $PO_4$ $p$–value |
|---|---|---|---|---|---|---|---|---|---|---|---|---|
| K58V | $1.2\times10^{2}$ | $8.0\times10^{-1}$ | $8.2\times10^{1}$ | $7.1\times10^{-1}$ | $1.1\times10^{6}$ | $3.4\times10^{-1}$ | $2.1\times10^{5}$ | $1.7\times10^{-2}$ | $1.3\times10^{1}$ | $5.5\times10^{-1}$ | $6.1\times10^{2}$ | $8.9\times10^{-1}$ |
| R59G | $1.5\times10^{1}$ | **$1.0\times10^{-3}$** | $7.3\times10^{1}$ | $3.5\times10^{-1}$ | $2.1\times10^{5}$ | **$4.4\times10^{-3}$** | $3.3\times10^{4}$ | **0** | $\le 9.9\times10^{-1}$ | **0** | $5.8\times10^{2}$ | $2.5\times10^{-1}$ |
| R59V | $2.7\times10^{1}$ | **$1.\times10^{-4}$** | $8.4\times10^{1}$ | $9.0\times10^{-1}$ | $4.3\times10^{5}$ | **$7.9\times10^{-3}$** | $1.9\times10^{4}$ | **0** | 4.2 | **$9.0\times10^{-4}$** | $4.9\times10^{2}$ | $2.4\times10^{-2}$ |
| M60G | $3.4\times10^{1}$ | **$2.8\times10^{-3}$** | $8.7\times10^{1}$ | $8.8\times10^{-1}$ | $3.9\times10^{5}$ | $1.4\times10^{-2}$ | $9.5\times10^{4}$ | **$5.5\times10^{-3}$** | 2.3 | **$2.0\times10^{-4}$** | $6.9\times10^{2}$ | $2.5\times10^{-2}$ |
| M60V | $2.0\times10^{2}$ | **$2.0\times10^{-4}$** | $1.0\times10^{2}$ | $6.1\times10^{-2}$ | $1.7\times10^{6}$ | $3.5\times10^{-1}$ | $3.9\times10^{5}$ | $7.7\times10^{-2}$ | 8.8 | $7.7\times10^{-2}$ | $8.1\times10^{2}$ | **$2.0\times10^{-4}$** |
| L61G | $1.3\times10^{2}$ | $8.5\times10^{-1}$ | $7.0\times10^{1}$ | $2.3\times10^{-1}$ | $1.3\times10^{6}$ | $7.7\times10^{-1}$ | $2.4\times10^{5}$ | **$7.4\times10^{-3}$** | 6.0 | **$1.8\times10^{-3}$** | $4.7\times10^{2}$ | **$4.0\times10^{-4}$** |
| L61V | $6.6\times10^{1}$ | **$3.9\times10^{-3}$** | $8.2\times10^{1}$ | $7.2\times10^{-1}$ | $7.6\times10^{5}$ | $2.1\times10^{-2}$ | $2.7\times10^{5}$ | **$1.9\times10^{-3}$** | 5.6 | **$1.3\times10^{-3}$** | $5.8\times10^{2}$ | $8.2\times10^{-2}$ |
| N62G | $1.0\times10^{2}$ | $3.6\times10^{-1}$ | $6.7\times10^{1}$ | $1.3\times10^{-1}$ | $1.2\times10^{6}$ | $4.9\times10^{-1}$ | $3.3\times10^{5}$ | $2.8\times10^{-2}$ | $1.8\times10^{1}$ | $5.5\times10^{-1}$ | $5.0\times10^{2}$ | $3.0\times10^{-2}$ |
| N62V | $1.6\times10^{2}$ | $2.7\times10^{-2}$ | $1.0\times10^{2}$ | $5.9\times10^{-2}$ | $1.5\times10^{6}$ | $6.6\times10^{-1}$ | $1.1\times10^{5}$ | **$1.0\times10^{-3}$** | $1.2\times10^{1}$ | $1.9\times10^{-1}$ | $6.6\times10^{2}$ | $1.4\times10^{-1}$ |
| T63G | $1.1\times10^{2}$ | $6.4\times10^{-1}$ | $5.8\times10^{1}$ | $5.5\times10^{-2}$ | $2.0\times10^{6}$ | $1.6\times10^{-1}$ | $3.2\times10^{5}$ | $2.9\times10^{-2}$ | $1.8\times10^{1}$ | $5.5\times10^{-1}$ | $5.0\times10^{2}$ | **$1.5\times10^{-3}$** |
| T63V | $1.1\times10^{2}$ | $2.9\times10^{-1}$ | $9.1\times10^{1}$ | $4.7\times10^{-1}$ | $1.2\times10^{6}$ | $3.5\times10^{-1}$ | $3.8\times10^{5}$ | $3.7\times10^{-2}$ | $1.0\times10^{1}$ | $1.4\times10^{-1}$ | $6.5\times10^{2}$ | $2.9\times10^{-1}$ |
| G64A | $1.2\times10^{1}$ | **0** | $7.9\times10^{1}$ | $4.4\times10^{-1}$ | $1.6\times10^{5}$ | **0** | $3.0\times10^{4}$ | **0** | $\le 1.2$ | **0** | $5.9\times10^{2}$ | $3.0\times10^{-1}$ |
| G64V | – | – | – | – | $\le 3.8\times10^{2}$ | **0** | $\le 8.9\times10^{2}$ | **0** | $\le 1.3$ | **0** | – | – |
| Y65G | $4.9\times10^{1}$ | **$1.7\times10^{-3}$** | $6.7\times10^{1}$ | $8.7\times10^{-2}$ | $6.0\times10^{5}$ | $1.4\times10^{-2}$ | $7.8\times10^{4}$ | **$1.2\times10^{-3}$** | 5.6 | **$9.0\times10^{-4}$** | $5.1\times10^{2}$ | **$2.5\times10^{-3}$** |
| Y65V | $1.3\times10^{2}$ | $6.6\times10^{-1}$ | $8.7\times10^{1}$ | $8.4\times10^{-1}$ | $1.4\times10^{6}$ | 1.0 | $2.4\times10^{5}$ | **$1.9\times10^{-3}$** | $1.0\times10^{1}$ | $8.7\times10^{-2}$ | $6.6\times10^{2}$ | $2.2\times10^{-1}$ |
| S66G | $7.5\times10^{1}$ | $3.5\times10^{-2}$ | $6.6\times10^{1}$ | $8.2\times10^{-2}$ | $1.1\times10^{6}$ | $3.7\times10^{-1}$ | $3.1\times10^{5}$ | $1.5\times10^{-2}$ | $1.3\times10^{1}$ | $2.8\times10^{-1}$ | $4.7\times10^{2}$ | **0** |
| S66V | $1.0\times10^{2}$ | $2.0\times10^{-1}$ | $8.9\times10^{1}$ | $6.0\times10^{-1}$ | $1.1\times10^{6}$ | $2.8\times10^{-1}$ | $3.6\times10^{5}$ | $1.9\times10^{-2}$ | 8.8 | $1.6\times10^{-2}$ | $5.9\times10^{2}$ | $2.3\times10^{-1}$ |
| L67G | $3.8\times10^{1}$ | **$3.1\times10^{-3}$** | $5.5\times10^{1}$ | $3.2\times10^{-2}$ | $7.4\times10^{5}$ | $1.2\times10^{-1}$ | $1.1\times10^{5}$ | **$2.1\times10^{-3}$** | 9.0 | $2.7\times10^{-2}$ | $4.7\times10^{2}$ | **$3.0\times10^{-4}$** |
| L67V | $8.3\times10^{1}$ | $7.0\times10^{-2}$ | $8.1\times10^{1}$ | $6.7\times10^{-1}$ | $1.1\times10^{6}$ | $3.1\times10^{-1}$ | $2.1\times10^{5}$ | $5.0\times10^{-2}$ | 8.7 | $5.7\times10^{-2}$ | $6.0\times10^{2}$ | $5.7\times10^{-1}$ |
| N68G | $6.8\times10^{1}$ | $2.1\times10^{-2}$ | $7.7\times10^{1}$ | $4.3\times10^{-1}$ | $8.4\times10^{5}$ | $7.7\times10^{-2}$ | $1.6\times10^{5}$ | $3.1\times10^{-2}$ | 9.5 | $2.9\times10^{-2}$ | $6.3\times10^{2}$ | $6.0\times10^{-1}$ |
| N68V | $1.6\times10^{1}$ | **0** | $7.9\times10^{1}$ | $3.7\times10^{-1}$ | $2.3\times10^{5}$ | **0** | $6.9\times10^{4}$ | $1.1\times10^{-2}$ | $\le 9.7\times10^{-1}$ | **0** | $4.8\times10^{2}$ | **0** |
| N69G | $1.8\times10^{1}$ | **$5.0\times10^{-4}$** | $5.1\times10^{1}$ | $2.0\times10^{-2}$ | $3.4\times10^{5}$ | $1.1\times10^{-2}$ | $1.2\times10^{5}$ | **$1.6\times10^{-3}$** | 6.6 | $1.5\times10^{-2}$ | $4.5\times10^{2}$ | **$2.0\times10^{-4}$** |
| N69V | $5.1\times10^{1}$ | **$2.0\times10^{-3}$** | $7.1\times10^{1}$ | $1.8\times10^{-1}$ | $7.1\times10^{5}$ | $3.5\times10^{-2}$ | $\le 8.2\times10^{4}$ | **$3.9\times10^{-3}$** | 6.3 | **$2.1\times10^{-3}$** | $5.0\times10^{2}$ | **$3.7\times10^{-3}$** |
| V70A | $2.1\times10^{2}$ | **$1.2\times10^{-3}$** | $7.5\times10^{1}$ | $3.7\times10^{-1}$ | $2.4\times10^{6}$ | **$8.3\times10^{-3}$** | $8.4\times10^{5}$ | $1.2\times10^{-1}$ | $1.8\times10^{1}$ | $6.1\times10^{-1}$ | $5.1\times10^{2}$ | $2.8\times10^{-2}$ |
| V70G | 3.2 | **$6.0\times10^{-4}$** | $5.1\times10^{1}$ | $1.8\times10^{-2}$ | $6.2\times10^{4}$ | **$2.3\times10^{-3}$** | $1.0\times10^{4}$ | **$5.0\times10^{-4}$** | $\le 9.9\times10^{-1}$ | **0** | $3.5\times10^{2}$ | **$7.0\times10^{-4}$** |
| H71G | $9.6\times10^{1}$ | $3.0\times10^{-1}$ | $7.4\times10^{1}$ | $2.7\times10^{-1}$ | $1.5\times10^{6}$ | $7.2\times10^{-1}$ | $3.7\times10^{5}$ | $5.8\times10^{-2}$ | 7.5 | **$5.9\times10^{-3}$** | $4.3\times10^{2}$ | **$2.0\times10^{-4}$** |
| H71V | $1.1\times10^{2}$ | $4.5\times10^{-1}$ | $9.0\times10^{1}$ | $7.0\times10^{-1}$ | $1.5\times10^{6}$ | $7.6\times10^{-1}$ | $8.6\times10^{5}$ | $4.7\times10^{-2}$ | $1.0\times10^{1}$ | $8.2\times10^{-2}$ | $6.0\times10^{2}$ | $4.9\times10^{-1}$ |
| I72G | 4.1 | **0** | $9.7\times10^{1}$ | $1.6\times10^{-1}$ | $3.5\times10^{4}$ | **$1.\times10^{-4}$** | $5.9\times10^{3}$ | **$1.\times10^{-4}$** | $\le 1.1$ | **$1.\times10^{-4}$** | $5.0\times10^{2}$ | **$1.\times10^{-4}$** |
| I72V | $8.0\times10^{1}$ | $8.4\times10^{-2}$ | $8.0\times10^{1}$ | $6.7\times10^{-1}$ | $1.1\times10^{6}$ | $3.8\times10^{-1}$ | $2.5\times10^{5}$ | $1.0\times10^{-2}$ | $1.1\times10^{1}$ | $2.1\times10^{-1}$ | $3.6\times10^{2}$ | **0** |
| D73G | $2.1\times10^{1}$ | **$5.0\times10^{-4}$** | $5.7\times10^{1}$ | $2.2\times10^{-2}$ | $3.7\times10^{5}$ | **$5.4\times10^{-3}$** | $1.5\times10^{5}$ | **$1.7\times10^{-3}$** | 4.3 | **0** | $4.1\times10^{2}$ | **$1.\times10^{-4}$** |
| D73V | $2.0\times10^{1}$ | **0** | $7.3\times10^{1}$ | $6.0\times10^{-2}$ | $2.5\times10^{5}$ | **0** | $6.4\times10^{4}$ | **$4.0\times10^{-4}$** | 2.3 | **$4.0\times10^{-4}$** | $3.4\times10^{2}$ | **0** |
| Y74G | 4.9 | **0** | $2.7\times10^{1}$ | **0** | $1.7\times10^{5}$ | **0** | $\ge 1.2\times10^{5}$ | **0** | 5.6 | **$1.\times10^{-4}$** | $4.1\times10^{1}$ | **0** |
| Y74V | $5.6\times10^{1}$ | **$1.9\times10^{-3}$** | $5.1\times10^{1}$ | **$1.8\times10^{-3}$** | $1.0\times10^{6}$ | $1.8\times10^{-1}$ | $5.0\times10^{5}$ | $2.8\times10^{-1}$ | $2.1\times10^{1}$ | $2.7\times10^{-1}$ | $1.1\times10^{2}$ | **0** |
| V75A | $6.6\times10^{1}$ | $1.8\times10^{-2}$ | $7.1\times10^{1}$ | $2.0\times10^{-1}$ | $9.3\times10^{5}$ | $1.4\times10^{-1}$ | $\le 5.6\times10^{4}$ | **$6.0\times10^{-4}$** | 4.3 | **$1.3\times10^{-3}$** | $5.5\times10^{2}$ | $7.8\times10^{-2}$ |
| V75G | $1.0\times10^{1}$ | **$4.0\times10^{-3}$** | $4.3\times10^{1}$ | $1.9\times10^{-2}$ | $2.9\times10^{5}$ | $2.5\times10^{-2}$ | $2.2\times10^{5}$ | **$4.8\times10^{-3}$** | $\le 1.3$ | **$1.\times10^{-4}$** | $2.1\times10^{2}$ | **$1.\times10^{-4}$** |
| P76V | 2.8 | **0** | $5.1\times10^{1}$ | **$8.0\times10^{-4}$** | $7.1\times10^{4}$ | **0** | $5.5\times10^{3}$ | **$8.0\times10^{-3}$** | $\le 1.9$ | **$1.\times10^{-4}$** | $2.5\times10^{2}$ | **$1.\times10^{-4}$** |
| T77G | $1.2\times10^{1}$ | **$3.0\times10^{-4}$** | $3.2\times10^{1}$ | **0** | $2.6\times10^{5}$ | **$1.6\times10^{-3}$** | $2.8\times10^{5}$ | $1.2\times10^{-2}$ | $1.1\times10^{1}$ | $6.2\times10^{-2}$ | $9.3\times10^{1}$ | **$3.0\times10^{-4}$** |
| T77V | $5.3\times10^{1}$ | **$8.3\times10^{-3}$** | $5.9\times10^{1}$ | $2.8\times10^{-2}$ | $9.0\times10^{5}$ | $1.4\times10^{-1}$ | $2.5\times10^{5}$ | **$9.6\times10^{-3}$** | $2.1\times10^{1}$ | $2.8\times10^{-1}$ | $2.5\times10^{2}$ | **$1.9\times10^{-3}$** |
| V78A | $6.5\times10^{1}$ | **$7.1\times10^{-3}$** | $3.0\times10^{1}$ | **0** | $2.7\times10^{6}$ | **$3.0\times10^{-4}$** | $5.4\times10^{4}$ | **$1.\times10^{-4}$** | $\le 1.8$ | **0** | $6.1\times10^{2}$ | $9.8\times10^{-1}$ |
| V78G | $1.7\times10^{1}$ | **$3.0\times10^{-4}$** | $2.4\times10^{1}$ | **$1.\times10^{-4}$** | $6.5\times10^{5}$ | $3.9\times10^{-2}$ | $9.7\times10^{4}$ | **$5.0\times10^{-3}$** | $\le 9.9\times10^{-1}$ | **$4.0\times10^{-4}$** | $2.9\times10^{2}$ | **0** |
| T79G | – | – | – | – | $\le 1.5\times10^{3}$ | **0** | $\le 6.3\times10^{3}$ | **0** | $\le 1.3$ | **0** | – | – |
| T79V | – | – | – | – | $\le 8.0\times10^{2}$ | **0** | $\le 1.4\times10^{3}$ | **0** | $\le 1.3$ | **0** | – | – |
| A80G | $3.5\times10^{1}$ | **$1.\times10^{-4}$** | $6.8\times10^{1}$ | $1.1\times10^{-1}$ | $4.5\times10^{5}$ | **$3.3\times10^{-3}$** | $2.3\times10^{5}$ | **$7.2\times10^{-3}$** | 7.9 | $1.1\times10^{-2}$ | $4.4\times10^{2}$ | **$1.\times10^{-4}$** |
| A80V | 5.2 | **0** | $4.0\times10^{1}$ | **0** | $2.0\times10^{5}$ | **0** | $1.5\times10^{4}$ | **$2.0\times10^{-4}$** | $8.5\times10^{-1}$ | **$1.\times10^{-4}$** | $1.1\times10^{3}$ | **0** |
| I81G | $2.3\times10^{1}$ | **$3.0\times10^{-4}$** | $2.6\times10^{1}$ | **0** | $9.4\times10^{5}$ | $2.0\times10^{-1}$ | $8.0\times10^{5}$ | $1.1\times10^{-1}$ | $1.8\times10^{1}$ | $6.9\times10^{-1}$ | $6.8\times10^{1}$ | **$4.0\times10^{-4}$** |
| I81V | $1.1\times10^{2}$ | $4.1\times10^{-1}$ | $9.3\times10^{1}$ | $4.2\times10^{-1}$ | $1.3\times10^{6}$ | $7.9\times10^{-1}$ | $5.0\times10^{5}$ | $3.6\times10^{-1}$ | $1.8\times10^{1}$ | $8.0\times10^{-1}$ | $4.1\times10^{2}$ | **0** |
| G82A | 1.4 | **0** | $1.6\times10^{1}$ | **0** | $9.7\times10^{4}$ | **0** | $\ge 1.7\times10^{4}$ | **0** | 4.9 | **$5.0\times10^{-4}$** | $2.1\times10^{1}$ | **0** |
| G82V | – | – | – | – | $\le 1.3\times10^{3}$ | **0** | $\le 6.3\times10^{2}$ | **0** | $\le 1.4$ | **0** | – | – |
| H83G | 1.9 | **0** | 5.6 | **0** | $3.1\times10^{5}$ | **0** | $\ge 2.1\times10^{5}$ | **$9.0\times10^{-4}$** | 1.2 | **$1.\times10^{-4}$** | 8.9 | **0** |
| H83V | $3.2\times10^{1}$ | **$3.0\times10^{-4}$** | $1.9\times10^{1}$ | **0** | $1.8\times10^{6}$ | $2.2\times10^{-1}$ | $2.7\times10^{5}$ | $3.6\times10^{-2}$ | 4.4 | **$1.7\times10^{-3}$** | $9.4\times10^{1}$ | **0** |
| T84G | $1.2\times10^{2}$ | $9.5\times10^{-1}$ | $7.9\times10^{1}$ | $5.9\times10^{-1}$ | $1.9\times10^{6}$ | $2.2\times10^{-1}$ | $3.2\times10^{5}$ | $9.5\times10^{-2}$ | $1.7\times10^{1}$ | $8.2\times10^{-1}$ | $8.3\times10^{2}$ | **$6.0\times10^{-4}$** |
| S85G | $7.9\times10^{1}$ | $7.5\times10^{-2}$ | $7.0\times10^{1}$ | $2.1\times10^{-1}$ | $1.1\times10^{6}$ | $4.6\times10^{-1}$ | $7.5\times10^{5}$ | $2.5\times10^{-1}$ | $1.0\times10^{1}$ | $5.2\times10^{-2}$ | $4.8\times10^{2}$ | **$1.\times10^{-4}$** |
| S85V | $1.8\times10^{1}$ | **0** | $7.5\times10^{1}$ | $2.6\times10^{-1}$ | $2.7\times10^{5}$ | **0** | $1.2\times10^{4}$ | **0** | 2.5 | **$5.0\times10^{-4}$** | $4.4\times10^{2}$ | **0** |
| I86G | 7.8 | **$1.\times10^{-4}$** | $7.1\times10^{1}$ | $2.2\times10^{-1}$ | $1.8\times10^{5}$ | **$1.1\times10^{-3}$** | $1.7\times10^{5}$ | **$3.5\times10^{-3}$** | 5.1 | **$7.0\times10^{-4}$** | $3.7\times10^{2}$ | **0** |
| I86V | $1.3\times10^{2}$ | $7.2\times10^{-1}$ | $7.6\times10^{1}$ | $3.7\times10^{-1}$ | $1.6\times10^{6}$ | $5.4\times10^{-1}$ | $3.7\times10^{5}$ | $6.2\times10^{-2}$ | 7.6 | $2.8\times10^{-2}$ | $5.8\times10^{2}$ | $1.7\times10^{-1}$ |
| F87G | $9.9\times10^{1}$ | $3.2\times10^{-1}$ | $6.3\times10^{1}$ | $5.6\times10^{-2}$ | $2.0\times10^{6}$ | $1.2\times10^{-1}$ | $6.7\times10^{5}$ | $6.0\times10^{-1}$ | $1.5\times10^{1}$ | $6.6\times10^{-1}$ | $3.5\times10^{2}$ | **$1.\times10^{-4}$** |
| F87V | $9.8\times10^{1}$ | $2.0\times10^{-1}$ | $6.5\times10^{1}$ | $3.4\times10^{-2}$ | $1.7\times10^{6}$ | $4.4\times10^{-1}$ | $3.7\times10^{5}$ | $5.1\times10^{-2}$ | $1.7\times10^{1}$ | $9.2\times10^{-1}$ | $2.9\times10^{2}$ | **0** |
| T88G | 9.7 | **$3.2\times10^{-3}$** | $4.0\times10^{1}$ | $1.2\times10^{-2}$ | $2.6\times10^{5}$ | $1.7\times10^{-2}$ | $2.2\times10^{5}$ | **$6.6\times10^{-3}$** | 6.2 | **$8.0\times10^{-4}$** | $1.7\times10^{2}$ | **$4.0\times10^{-4}$** |
| T88V | $4.9\times10^{1}$ | **$2.5\times10^{-3}$** | $6.6\times10^{1}$ | $6.6\times10^{-2}$ | $7.5\times10^{5}$ | $4.6\times10^{-2}$ | $6.9\times10^{4}$ | **$1.0\times10^{-3}$** | 3.2 | **$1.8\times10^{-3}$** | $3.6\times10^{2}$ | **0** |
| G89A | $5.3\times10^{1}$ | **$2.6\times10^{-3}$** | $7.8\times10^{1}$ | $4.4\times10^{-1}$ | $7.7\times10^{5}$ | $5.3\times10^{-2}$ | $3.5\times10^{5}$ | $4.9\times10^{-2}$ | $1.4\times10^{1}$ | $4.0\times10^{-1}$ | $6.6\times10^{2}$ | $1.4\times10^{-1}$ |
| G89V | – | – | – | – | $\le 1.3\times10^{3}$ | **0** | $\le 1.3\times10^{3}$ | **0** | $\le 1.3$ | **0** | – | – |
| S90G | $5.2\times10^{1}$ | **$2.0\times10^{-3}$** | $6.1\times10^{1}$ | $2.2\times10^{-2}$ | $1.1\times10^{6}$ | $3.5\times10^{-1}$ | $5.1\times10^{5}$ | $3.8\times10^{-1}$ | 8.3 | $1.8\times10^{-2}$ | $3.9\times10^{2}$ | **0** |
| S90V | $2.0\times10^{1}$ | **$5.0\times10^{-3}$** | $7.9\times10^{1}$ | $6.7\times10^{-1}$ | $2.7\times10^{5}$ | $2.0\times10^{-2}$ | $8.5\times10^{4}$ | $1.3\times10^{-2}$ | $\le 1.9$ | **$2.9\times10^{-3}$** | $4.7\times10^{2}$ | $3.8\times10^{-2}$ |
| V91A | $6.4\times10^{1}$ | **$5.4\times10^{-3}$** | $7.0\times10^{1}$ | $1.0\times10^{-1}$ | $1.0\times10^{6}$ | $1.9\times10^{-1}$ | $\le 8.2\times10^{4}$ | **$3.8\times10^{-3}$** | 6.4 | $1.3\times10^{-2}$ | $4.7\times10^{2}$ | **$3.0\times10^{-4}$** |
| V91G | 4.6 | **0** | $4.9\times10^{1}$ | **$1.\times10^{-4}$** | $9.5\times10^{4}$ | **0** | $1.9\times10^{4}$ | **0** | $\le 1.2$ | **0** | $3.5\times10^{2}$ | **0** |
| P92G | $1.2\times10^{1}$ | **0** | $5.2\times10^{1}$ | **$1.\times10^{-4}$** | $2.3\times10^{5}$ | **0** | $9.3\times10^{4}$ | **$3.0\times10^{-4}$** | 2.9 | **0** | $3.6\times10^{2}$ | **0** |
| P92V | 7.1 | **0** | $7.7\times10^{1}$ | $4.4\times10^{-1}$ | $1.2\times10^{5}$ | **$6.0\times10^{-4}$** | $\le 1.4\times10^{4}$ | **0** | $\le 1.6$ | **0** | $5.2\times10^{2}$ | $2.3\times10^{-2}$ |
| S93G | $3.0\times10^{1}$ | **$4.0\times10^{-4}$** | $5.4\times10^{1}$ | **$9.0\times10^{-3}$** | $5.4\times10^{5}$ | $1.7\times10^{-2}$ | $2.5\times10^{5}$ | **$8.0\times10^{-3}$** | 8.0 | **$4.4\times10^{-3}$** | $4.0\times10^{2}$ | **0** |
| S93V | 2.5 | **0** | $4.6\times10^{1}$ | **$2.4\times10^{-3}$** | $5.5\times10^{4}$ | **$3.0\times10^{-4}$** | $4.0\times10^{3}$ | **$3.3\times10^{-3}$** | $\le 1.3$ | **$1.\times10^{-4}$** | $3.5\times10^{2}$ | **$2.0\times10^{-4}$** |
| I94G | 1.3 | **0** | $4.0\times10^{1}$ | **$1.\times10^{-4}$** | $2.9\times10^{4}$ | **0** | $5.7\times10^{3}$ | **$6.0\times10^{-4}$** | $\le 1.1$ | **$1.\times10^{-4}$** | $1.6\times10^{2}$ | **0** |
| I94V | $1.4\times10^{2}$ | $3.7\times10^{-1}$ | $8.8\times10^{1}$ | $8.3\times10^{-1}$ | $1.5\times10^{6}$ | $6.9\times10^{-1}$ | $3.6\times10^{5}$ | $2.1\times10^{-2}$ | $1.7\times10^{1}$ | $9.7\times10^{-1}$ | $5.9\times10^{2}$ | $4.1\times10^{-1}$ |
| H95G | 3.0 | **$4.6\times10^{-3}$** | $6.8\times10^{1}$ | $2.9\times10^{-1}$ | $4.7\times10^{4}$ | $1.2\times10^{-2}$ | $1.3\times10^{5}$ | **$3.7\times10^{-3}$** | $\le 9.9\times10^{-1}$ | **$2.0\times10^{-4}$** | – | – |
| H95V | $3.8\times10^{1}$ | **$3.0\times10^{-4}$** | $8.1\times10^{1}$ | $6.1\times10^{-1}$ | $4.8\times10^{5}$ | **$3.7\times10^{-3}$** | $9.7\times10^{4}$ | **$6.0\times10^{-4}$** | $1.1\times10^{1}$ | $2.9\times10^{-1}$ | $3.8\times10^{2}$ | **$6.0\times10^{-4}$** |
| G96A | $2.9\times10^{1}$ | **$2.0\times10^{-3}$** | $6.2\times10^{1}$ | $9.0\times10^{-2}$ | $6.7\times10^{5}$ | $8.7\times10^{-2}$ | $1.1\times10^{5}$ | **$1.8\times10^{-3}$** | 4.8 | **$1.\times10^{-4}$** | $4.9\times10^{2}$ | **$1.1\times10^{-3}$** |
| G96V | – | – | – | – | $\le 2.2\times10^{3}$ | **0** | $\le 1.0\times10^{3}$ | **0** | $\le 1.0$ | **0** | – | – |

| Variant | $k_{cat}$ cMUP ($s^{-1}$) | $k_{cat}$ cMUP $p$–value | $K_M$ cMUP (μM) | $K_M$ cMUP $p$–value | $k_{cat}/K_M$ cMUP ($M^{-1}s^{-1}$) | $k_{cat}/K_M$ cMUP $p$–value | $k_{cat}/K_M$ MeP ($M^{-1}s^{-1}$) | $k_{cat}/K_M$ MeP $p$–value | $k_{cat}/K_M$ MecMUP ($M^{-1}s^{-1}$) | $k_{cat}/K_M$ MecMUP $p$–value | $K_i$ $PO_4$ (μM) | $K_i$ $PO_4$ $p$–value |
|---|---|---|---|---|---|---|---|---|---|---|---|---|
| I97G | – | – | – | – | $\leq 1.4\times10^{3}$ | **0** | $\leq 6.5\times10^{2}$ | **$1.\times10^{-4}$** | ≤ 1.2 | **0** | – | – |
| I97V | $1.5\times10^{2}$ | $1.4\times10^{-1}$ | $8.9\times10^{1}$ | $6.0\times10^{-1}$ | $1.7\times10^{6}$ | $3.8\times10^{-1}$ | $4.3\times10^{5}$ | $7.2\times10^{-2}$ | $1.4\times10^{1}$ | $5.7\times10^{-1}$ | $5.7\times10^{2}$ | $2.3\times10^{-1}$ |
| A98G | $5.2\times10^{1}$ | $2.2\times10^{-2}$ | $6.0\times10^{1}$ | $6.1\times10^{-2}$ | $8.1\times10^{5}$ | $1.7\times10^{-1}$ | $2.5\times10^{5}$ | $3.0\times10^{-2}$ | $1.0\times10^{1}$ | $3.9\times10^{-2}$ | $5.3\times10^{2}$ | $1.2\times10^{-2}$ |
| A98V | $1.2\times10^{2}$ | $6.5\times10^{-1}$ | $8.7\times10^{1}$ | $8.2\times10^{-1}$ | $1.3\times10^{6}$ | $5.9\times10^{-1}$ | $4.4\times10^{5}$ | $8.3\times10^{-2}$ | $1.3\times10^{1}$ | $5.4\times10^{-1}$ | $5.0\times10^{2}$ | **$7.0\times10^{-4}$** |
| G99A | $1.1\times10^{2}$ | $5.3\times10^{-1}$ | $7.4\times10^{1}$ | $3.0\times10^{-1}$ | $1.0\times10^{6}$ | $3.0\times10^{-1}$ | $2.7\times10^{5}$ | $1.8\times10^{-2}$ | $1.1\times10^{1}$ | $6.7\times10^{-2}$ | $8.7\times10^{2}$ | **$2.0\times10^{-4}$** |
| G99V | 1.3 | **0** | 5.5 | **0** | $1.4\times10^{5}$ | **0** | $\leq 2.1\times10^{3}$ | **0** | 1.7 | **$1.\times10^{-4}$** | $1.0\times10^{2}$ | **0** |
| N100G | 1.3 | **0** | ≤ 5.0 | **0** | $\geq 2.7\times10^{5}$ | **0** | $1.8\times10^{3}$ | **$7.0\times10^{-4}$** | 1.2 | **$1.\times10^{-4}$** | $\leq 3.7\times10^{2}$ | **0** |
| N100V | 1.8 | **0** | ≤ 5.0 | **0** | $\geq 3.5\times10^{5}$ | **$2.0\times10^{-4}$** | $\leq 1.7\times10^{3}$ | **$1.\times10^{-4}$** | ≤ 1.3 | **0** | $\leq 8.1\times10^{2}$ | **0** |
| D101G | $4.0\times10^{1}$ | **$4.6\times10^{-3}$** | $2.9\times10^{1}$ | **$7.0\times10^{-4}$** | $1.3\times10^{6}$ | $7.9\times10^{-1}$ | $1.1\times10^{5}$ | **$1.9\times10^{-3}$** | $2.5\times10^{1}$ | **$1.1\times10^{-3}$** | $7.3\times10^{2}$ | **$6.8\times10^{-3}$** |
| D101V | $1.4\times10^{2}$ | $2.0\times10^{-1}$ | $6.2\times10^{1}$ | **$2.8\times10^{-3}$** | $2.3\times10^{6}$ | **$1.6\times10^{-3}$** | $1.7\times10^{5}$ | **$1.\times10^{-4}$** | $1.6\times10^{1}$ | $8.9\times10^{-1}$ | $9.9\times10^{2}$ | **0** |
| W102G | – | – | – | – | $\leq 1.3\times10^{3}$ | **0** | $\leq 1.3\times10^{3}$ | **$1.\times10^{-4}$** | ≤ 1.0 | **0** | – | – |
| W102V | 1.8 | **0** | $2.5\times10^{1}$ | **0** | $5.7\times10^{4}$ | **0** | $3.7\times10^{3}$ | **0** | ≤ 1.3 | **$1.\times10^{-4}$** | $7.6\times10^{2}$ | **$4.0\times10^{-4}$** |
| Y103G | 9.7 | **0** | $2.9\times10^{1}$ | **0** | $2.8\times10^{5}$ | **$5.0\times10^{-4}$** | $2.6\times10^{4}$ | **0** | 1.9 | **$1.\times10^{-4}$** | $3.8\times10^{2}$ | **$1.\times10^{-4}$** |
| Y103V | $9.0\times10^{1}$ | $1.3\times10^{-1}$ | $7.5\times10^{1}$ | $3.2\times10^{-1}$ | $1.1\times10^{6}$ | $3.0\times10^{-1}$ | $1.3\times10^{5}$ | **$1.\times10^{-4}$** | 5.4 | **$4.8\times10^{-3}$** | $4.7\times10^{2}$ | **0** |
| D104G | $4.4\times10^{1}$ | **$2.0\times10^{-3}$** | $6.9\times10^{1}$ | $1.7\times10^{-1}$ | $6.7\times10^{5}$ | $5.0\times10^{-2}$ | $1.8\times10^{5}$ | **$4.2\times10^{-3}$** | $1.0\times10^{1}$ | $4.3\times10^{-2}$ | $4.9\times10^{2}$ | **$1.\times10^{-4}$** |
| D104V | $1.4\times10^{2}$ | $4.5\times10^{-1}$ | $9.6\times10^{1}$ | $3.5\times10^{-1}$ | $1.5\times10^{6}$ | $8.2\times10^{-1}$ | $1.1\times10^{5}$ | **$5.8\times10^{-3}$** | ≤ 4.1 | $1.2\times10^{-2}$ | $8.5\times10^{2}$ | **0** |
| K105G | $1.8\times10^{2}$ | $1.7\times10^{-2}$ | $7.6\times10^{1}$ | $3.9\times10^{-1}$ | $2.1\times10^{6}$ | $7.8\times10^{-2}$ | $5.3\times10^{5}$ | $5.2\times10^{-1}$ | $2.5\times10^{1}$ | **$1.8\times10^{-3}$** | $6.5\times10^{2}$ | $3.4\times10^{-1}$ |
| K105V | $2.2\times10^{2}$ | **$8.0\times10^{-4}$** | $8.6\times10^{1}$ | $9.8\times10^{-1}$ | $2.7\times10^{6}$ | **$1.0\times10^{-3}$** | $6.9\times10^{5}$ | $5.3\times10^{-1}$ | $1.8\times10^{1}$ | $6.8\times10^{-1}$ | $6.2\times10^{2}$ | $7.9\times10^{-1}$ |
| E106G | $1.1\times10^{2}$ | $6.8\times10^{-1}$ | $6.5\times10^{1}$ | $4.3\times10^{-2}$ | $1.3\times10^{6}$ | $8.1\times10^{-1}$ | $2.5\times10^{5}$ | $1.1\times10^{-2}$ | $1.8\times10^{1}$ | $4.9\times10^{-1}$ | $5.4\times10^{2}$ | $4.2\times10^{-2}$ |
| E106V | $5.1\times10^{1}$ | **$2.4\times10^{-3}$** | $7.8\times10^{1}$ | $4.6\times10^{-1}$ | $6.0\times10^{5}$ | $1.2\times10^{-2}$ | $4.8\times10^{5}$ | $2.8\times10^{-1}$ | $1.1\times10^{1}$ | $2.1\times10^{-1}$ | $5.4\times10^{2}$ | **$7.1\times10^{-3}$** |
| L107G | $1.1\times10^{2}$ | $4.8\times10^{-1}$ | $5.5\times10^{1}$ | $1.8\times10^{-2}$ | $2.1\times10^{6}$ | $1.0\times10^{-1}$ | $4.3\times10^{5}$ | $1.3\times10^{-1}$ | $2.6\times10^{1}$ | **$9.0\times10^{-4}$** | $4.5\times10^{2}$ | **$2.0\times10^{-4}$** |
| L107V | $1.5\times10^{2}$ | $1.7\times10^{-1}$ | $1.0\times10^{2}$ | $6.2\times10^{-2}$ | $1.5\times10^{6}$ | $8.8\times10^{-1}$ | $6.3\times10^{5}$ | $8.5\times10^{-1}$ | $1.4\times10^{1}$ | $5.4\times10^{-1}$ | $6.6\times10^{2}$ | $1.5\times10^{-1}$ |
| G108A | $7.2\times10^{1}$ | $4.0\times10^{-2}$ | $6.3\times10^{1}$ | $4.7\times10^{-2}$ | $1.3\times10^{6}$ | $7.7\times10^{-1}$ | $2.6\times10^{5}$ | $1.1\times10^{-2}$ | 7.3 | **$1.7\times10^{-3}$** | $5.4\times10^{2}$ | $1.5\times10^{-2}$ |
| G108V | $1.4\times10^{2}$ | $3.2\times10^{-1}$ | $1.0\times10^{2}$ | $1.2\times10^{-1}$ | $1.4\times10^{6}$ | $9.4\times10^{-1}$ | $3.9\times10^{5}$ | $4.4\times10^{-2}$ | 9.2 | $1.4\times10^{-1}$ | $6.6\times10^{2}$ | $2.2\times10^{-1}$ |
| K109G | $2.4\times10^{2}$ | **$3.0\times10^{-4}$** | $1.0\times10^{2}$ | $1.5\times10^{-1}$ | $2.6\times10^{6}$ | **$1.2\times10^{-3}$** | $7.3\times10^{5}$ | $3.2\times10^{-1}$ | $2.2\times10^{1}$ | $6.5\times10^{-2}$ | $7.7\times10^{2}$ | **$7.0\times10^{-4}$** |
| K109V | $1.5\times10^{2}$ | $1.1\times10^{-1}$ | $8.9\times10^{1}$ | $7.2\times10^{-1}$ | $1.9\times10^{6}$ | $7.7\times10^{-2}$ | $5.7\times10^{5}$ | $8.0\times10^{-1}$ | $1.6\times10^{1}$ | $9.2\times10^{-1}$ | $5.9\times10^{2}$ | $4.4\times10^{-1}$ |
| S110G | $1.4\times10^{2}$ | $5.2\times10^{-1}$ | $7.3\times10^{1}$ | $2.9\times10^{-1}$ | $1.7\times10^{6}$ | $4.1\times10^{-1}$ | $6.5\times10^{5}$ | $7.7\times10^{-1}$ | $2.0\times10^{1}$ | $1.9\times10^{-1}$ | $4.5\times10^{2}$ | **$1.\times10^{-4}$** |
| S110V | $9.3\times10^{1}$ | $1.2\times10^{-1}$ | $8.6\times10^{1}$ | $9.1\times10^{-1}$ | $1.7\times10^{6}$ | $4.9\times10^{-1}$ | $4.0\times10^{5}$ | $8.6\times10^{-2}$ | $1.2\times10^{1}$ | $2.6\times10^{-1}$ | $1.1\times10^{3}$ | **0** |
| V111A | $1.4\times10^{2}$ | $3.1\times10^{-1}$ | $6.6\times10^{1}$ | $2.9\times10^{-2}$ | $1.6\times10^{6}$ | $4.7\times10^{-1}$ | $1.8\times10^{5}$ | **$6.0\times10^{-4}$** | $1.3\times10^{1}$ | $3.0\times10^{-1}$ | $7.1\times10^{2}$ | **$2.1\times10^{-3}$** |
| V111G | 9.2 | **$2.3\times10^{-3}$** | $8.5\times10^{1}$ | $9.7\times10^{-1}$ | $1.1\times10^{5}$ | **$3.7\times10^{-3}$** | $1.0\times10^{5}$ | **$7.0\times10^{-4}$** | 5.4 | **$9.0\times10^{-4}$** | $2.2\times10^{3}$ | **0** |
| Y112G | $1.9\times10^{1}$ | **0** | $2.6\times10^{1}$ | **0** | $6.6\times10^{5}$ | **$3.4\times10^{-3}$** | $2.6\times10^{4}$ | **0** | $3.2\times10^{1}$ | **0** | $2.2\times10^{3}$ | **0** |
| Y112V | $3.5\times10^{1}$ | **$1.\times10^{-4}$** | $1.3\times10^{2}$ | **$2.0\times10^{-4}$** | $4.4\times10^{5}$ | **$2.0\times10^{-3}$** | $1.5\times10^{4}$ | **$1.\times10^{-4}$** | 9.1 | $3.8\times10^{-2}$ | $4.6\times10^{3}$ | **0** |
| C113G | $8.9\times10^{1}$ | $2.4\times10^{-1}$ | $6.4\times10^{1}$ | $1.3\times10^{-1}$ | $1.3\times10^{6}$ | $7.9\times10^{-1}$ | $2.1\times10^{5}$ | **$4.5\times10^{-3}$** | $1.4\times10^{1}$ | $5.7\times10^{-1}$ | $7.2\times10^{2}$ | **$8.4\times10^{-3}$** |
| C113V | $2.9\times10^{2}$ | **0** | $9.8\times10^{1}$ | $1.7\times10^{-1}$ | $2.5\times10^{6}$ | **$1.4\times10^{-3}$** | $5.5\times10^{5}$ | $6.3\times10^{-1}$ | $2.0\times10^{1}$ | $3.9\times10^{-1}$ | $4.7\times10^{2}$ | **$4.8\times10^{-3}$** |
| T114G | 1.3 | **0** | ≤ 5.0 | **0** | $\geq 2.5\times10^{5}$ | **0** | $\leq 1.5\times10^{3}$ | **0** | 3.5 | **0** | $\leq 5.9\times10^{2}$ | $2.9\times10^{-1}$ |
| T114V | $9.8\times10^{1}$ | $2.4\times10^{-1}$ | $8.6\times10^{1}$ | $9.7\times10^{-1}$ | $1.4\times10^{6}$ | $8.3\times10^{-1}$ | $5.4\times10^{5}$ | $5.5\times10^{-1}$ | $4.2\times10^{1}$ | **0** | $3.6\times10^{2}$ | **$6.0\times10^{-4}$** |
| S115G | $8.7\times10^{1}$ | $1.4\times10^{-1}$ | $7.0\times10^{1}$ | $2.2\times10^{-1}$ | $1.5\times10^{6}$ | $7.4\times10^{-1}$ | $4.5\times10^{5}$ | $1.6\times10^{-1}$ | $2.0\times10^{1}$ | $1.9\times10^{-1}$ | $5.8\times10^{2}$ | $1.9\times10^{-1}$ |
| S115V | $1.1\times10^{2}$ | $6.5\times10^{-1}$ | $8.4\times10^{1}$ | $9.1\times10^{-1}$ | $1.1\times10^{6}$ | $2.6\times10^{-1}$ | $4.3\times10^{5}$ | $8.6\times10^{-2}$ | $1.2\times10^{1}$ | $2.6\times10^{-1}$ | $1.9\times10^{3}$ | **0** |
| D116G | $8.9\times10^{-2}$ | **$4.0\times10^{-4}$** | ≤ 5.0 | **0** | $\geq 1.8\times10^{4}$ | **$1.8\times10^{-3}$** | $\leq 1.2\times10^{3}$ | **0** | ≤ 1.3 | **0** | $\leq 5.3\times10^{2}$ | $2.1\times10^{-1}$ |
| D116V | 7.1 | **0** | $4.1\times10^{1}$ | **$8.0\times10^{-4}$** | $2.1\times10^{5}$ | **$8.0\times10^{-4}$** | $\leq 9.1\times10^{3}$ | **$4.0\times10^{-4}$** | ≤ 1.3 | **$2.0\times10^{-4}$** | – | – |
| E117G | $6.9\times10^{1}$ | $2.5\times10^{-2}$ | $6.1\times10^{1}$ | $2.1\times10^{-2}$ | $1.8\times10^{6}$ | $2.2\times10^{-1}$ | $3.8\times10^{5}$ | $7.0\times10^{-2}$ | $1.9\times10^{1}$ | $3.6\times10^{-1}$ | $5.6\times10^{2}$ | $6.8\times10^{-2}$ |
| E117V | $1.6\times10^{2}$ | $5.2\times10^{-2}$ | $9.4\times10^{1}$ | $3.3\times10^{-1}$ | $1.7\times10^{6}$ | $4.0\times10^{-1}$ | $5.3\times10^{5}$ | $4.8\times10^{-1}$ | $1.3\times10^{1}$ | $4.9\times10^{-1}$ | $6.5\times10^{2}$ | $2.4\times10^{-1}$ |
| T118G | $1.0\times10^{2}$ | $4.0\times10^{-1}$ | $6.4\times10^{1}$ | $7.0\times10^{-2}$ | $1.9\times10^{6}$ | $1.9\times10^{-1}$ | $4.7\times10^{5}$ | $1.6\times10^{-1}$ | $1.3\times10^{1}$ | $3.7\times10^{-1}$ | $5.4\times10^{2}$ | $1.2\times10^{-1}$ |
| T118V | $1.5\times10^{2}$ | $1.2\times10^{-1}$ | $9.1\times10^{1}$ | $5.4\times10^{-1}$ | $1.6\times10^{6}$ | $6.2\times10^{-1}$ | $4.9\times10^{5}$ | $3.1\times10^{-1}$ | $1.4\times10^{1}$ | $5.5\times10^{-1}$ | $6.0\times10^{2}$ | $6.6\times10^{-1}$ |
| V119A | $2.3\times10^{2}$ | **0** | $9.3\times10^{1}$ | $4.2\times10^{-1}$ | $1.8\times10^{6}$ | $2.1\times10^{-1}$ | $6.1\times10^{5}$ | $9.6\times10^{-1}$ | $2.2\times10^{1}$ | $8.7\times10^{-2}$ | $7.2\times10^{2}$ | $1.4\times10^{-2}$ |
| V119G | 7.7 | **$3.2\times10^{-3}$** | $7.8\times10^{1}$ | $6.0\times10^{-1}$ | $1.0\times10^{5}$ | **$4.5\times10^{-3}$** | $8.2\times10^{4}$ | **$8.0\times10^{-4}$** | 6.5 | **$2.9\times10^{-3}$** | $7.4\times10^{2}$ | $1.5\times10^{-2}$ |
| Q120G | $1.1\times10^{2}$ | $5.5\times10^{-1}$ | $7.1\times10^{1}$ | $2.7\times10^{-1}$ | $2.0\times10^{6}$ | $1.9\times10^{-1}$ | $5.2\times10^{5}$ | $3.6\times10^{-1}$ | $2.1\times10^{1}$ | $1.6\times10^{-1}$ | $5.6\times10^{2}$ | $1.1\times10^{-1}$ |
| Q120V | $1.6\times10^{2}$ | $3.2\times10^{-2}$ | $9.7\times10^{1}$ | $2.1\times10^{-1}$ | $1.5\times10^{6}$ | $7.2\times10^{-1}$ | $5.3\times10^{5}$ | $2.9\times10^{-1}$ | $1.2\times10^{1}$ | $2.0\times10^{-1}$ | $6.8\times10^{2}$ | $1.4\times10^{-2}$ |
| P121G | $8.9\times10^{1}$ | $2.5\times10^{-1}$ | $4.8\times10^{1}$ | **$9.8\times10^{-3}$** | $1.7\times10^{6}$ | $5.3\times10^{-1}$ | $1.1\times10^{6}$ | **$6.6\times10^{-3}$** | $1.4\times10^{1}$ | $5.0\times10^{-1}$ | $5.6\times10^{2}$ | $2.0\times10^{-1}$ |
| P121V | $8.9\times10^{1}$ | $8.0\times10^{-2}$ | $8.0\times10^{1}$ | $5.6\times10^{-1}$ | $1.0\times10^{6}$ | $2.0\times10^{-1}$ | $5.4\times10^{5}$ | $4.2\times10^{-1}$ | $1.5\times10^{1}$ | $7.1\times10^{-1}$ | $6.2\times10^{2}$ | $5.9\times10^{-1}$ |
| V122A | $1.5\times10^{2}$ | $2.6\times10^{-1}$ | $9.4\times10^{1}$ | $4.8\times10^{-1}$ | $1.6\times10^{6}$ | $7.3\times10^{-1}$ | $2.3\times10^{5}$ | **$6.8\times10^{-3}$** | $1.2\times10^{1}$ | $2.8\times10^{-1}$ | $6.0\times10^{2}$ | $6.2\times10^{-1}$ |
| V122G | 2.4 | **0** | 5.1 | **0** | $3.9\times10^{5}$ | **$2.0\times10^{-4}$** | $2.9\times10^{4}$ | **$1.\times10^{-4}$** | 2.5 | **0** | $2.4\times10^{2}$ | **0** |
| G123A | $9.1\times10^{1}$ | $3.1\times10^{-1}$ | $6.2\times10^{1}$ | $1.3\times10^{-1}$ | $1.3\times10^{6}$ | $8.2\times10^{-1}$ | $2.9\times10^{5}$ | $1.9\times10^{-2}$ | $1.8\times10^{1}$ | $6.5\times10^{-1}$ | $6.0\times10^{2}$ | $4.7\times10^{-1}$ |
| G123V | $9.5\times10^{1}$ | $3.1\times10^{-1}$ | $6.9\times10^{1}$ | $1.6\times10^{-1}$ | $1.9\times10^{6}$ | $2.4\times10^{-1}$ | $1.6\times10^{5}$ | **$2.9\times10^{-3}$** | 9.5 | $1.5\times10^{-1}$ | $5.7\times10^{2}$ | $1.4\times10^{-1}$ |
| T124G | $8.4\times10^{1}$ | $1.2\times10^{-1}$ | $5.7\times10^{1}$ | $2.9\times10^{-2}$ | $1.7\times10^{6}$ | $4.7\times10^{-1}$ | $3.0\times10^{5}$ | $2.1\times10^{-2}$ | $2.0\times10^{1}$ | $2.0\times10^{-1}$ | $6.3\times10^{2}$ | $6.7\times10^{-1}$ |
| T124V | $1.3\times10^{2}$ | $6.1\times10^{-1}$ | $7.4\times10^{1}$ | $3.6\times10^{-1}$ | $2.0\times10^{6}$ | $1.3\times10^{-1}$ | $3.8\times10^{5}$ | $2.3\times10^{-2}$ | $1.3\times10^{1}$ | $4.1\times10^{-1}$ | $5.2\times10^{2}$ | $1.5\times10^{-2}$ |
| T125G | $1.2\times10^{2}$ | $9.0\times10^{-1}$ | $5.4\times10^{1}$ | $3.2\times10^{-2}$ | $2.3\times10^{6}$ | $5.0\times10^{-2}$ | $4.3\times10^{5}$ | $1.4\times10^{-1}$ | $2.3\times10^{1}$ | $3.5\times10^{-2}$ | $5.6\times10^{2}$ | $8.4\times10^{-2}$ |
| T125V | $1.3\times10^{2}$ | $5.8\times10^{-1}$ | $9.7\times10^{1}$ | $2.9\times10^{-1}$ | $1.3\times10^{6}$ | $6.4\times10^{-1}$ | $6.4\times10^{5}$ | $7.9\times10^{-1}$ | $1.0\times10^{1}$ | $1.8\times10^{-1}$ | $6.5\times10^{2}$ | $2.4\times10^{-1}$ |
| S126G | $7.6\times10^{1}$ | $6.3\times10^{-2}$ | $6.5\times10^{1}$ | $8.9\times10^{-2}$ | $8.2\times10^{5}$ | $1.6\times10^{-1}$ | $3.8\times10^{5}$ | $8.2\times10^{-2}$ | $2.4\times10^{1}$ | $1.1\times10^{-2}$ | $5.9\times10^{2}$ | $4.8\times10^{-1}$ |
| S126V | $1.6\times10^{2}$ | $7.7\times10^{-2}$ | $1.1\times10^{2}$ | $5.7\times10^{-2}$ | $1.4\times10^{6}$ | $9.5\times10^{-1}$ | $7.8\times10^{5}$ | $2.9\times10^{-1}$ | $1.6\times10^{1}$ | $9.4\times10^{-1}$ | $6.9\times10^{2}$ | $1.3\times10^{-1}$ |
| N127G | $1.0\times10^{2}$ | $3.9\times10^{-1}$ | $8.9\times10^{1}$ | $7.9\times10^{-1}$ | $1.5\times10^{6}$ | $7.3\times10^{-1}$ | $5.2\times10^{5}$ | $5.1\times10^{-1}$ | $2.1\times10^{1}$ | $1.0\times10^{-1}$ | $7.9\times10^{2}$ | **$7.0\times10^{-4}$** |
| N127V | $1.5\times10^{2}$ | $1.3\times10^{-1}$ | $8.8\times10^{1}$ | $8.4\times10^{-1}$ | $1.5\times10^{6}$ | $7.7\times10^{-1}$ | $6.7\times10^{5}$ | $7.0\times10^{-1}$ | $1.3\times10^{1}$ | $4.9\times10^{-1}$ | $6.3\times10^{2}$ | $5.9\times10^{-1}$ |
| S128G | $1.4\times10^{2}$ | $2.9\times10^{-1}$ | $8.6\times10^{1}$ | 1.0 | $1.9\times10^{6}$ | $1.9\times10^{-1}$ | $4.0\times10^{5}$ | $9.7\times10^{-2}$ | $2.2\times10^{1}$ | $8.3\times10^{-2}$ | $6.2\times10^{2}$ | $6.7\times10^{-1}$ |
| S128V | $8.4\times10^{1}$ | $5.7\times10^{-2}$ | $9.3\times10^{1}$ | $4.5\times10^{-1}$ | $1.2\times10^{6}$ | $5.3\times10^{-1}$ | $3.8\times10^{5}$ | $8.0\times10^{-2}$ | $1.1\times10^{1}$ | $2.5\times10^{-1}$ | $7.0\times10^{2}$ | $3.0\times10^{-2}$ |
| V129A | $1.9\times10^{2}$ | **$3.0\times10^{-3}$** | $1.1\times10^{2}$ | **$7.9\times10^{-3}$** | $1.7\times10^{6}$ | $4.7\times10^{-1}$ | $3.5\times10^{5}$ | $4.7\times10^{-2}$ | $2.5\times10^{1}$ | **$5.5\times10^{-3}$** | $6.9\times10^{2}$ | $3.7\times10^{-2}$ |
| V129G | $7.5\times10^{1}$ | $6.7\times10^{-2}$ | $8.3\times10^{1}$ | $8.0\times10^{-1}$ | $6.8\times10^{5}$ | $6.6\times10^{-2}$ | $4.8\times10^{5}$ | $2.7\times10^{-1}$ | $1.3\times10^{1}$ | $3.7\times10^{-1}$ | $5.9\times10^{2}$ | $4.6\times10^{-1}$ |
| G130A | $1.4\times10^{2}$ | $3.0\times10^{-1}$ | $6.1\times10^{1}$ | $5.7\times10^{-2}$ | $2.0\times10^{6}$ | $1.3\times10^{-1}$ | $3.1\times10^{5}$ | $5.6\times10^{-2}$ | $2.1\times10^{1}$ | $2.0\times10^{-1}$ | $6.3\times10^{2}$ | $5.7\times10^{-1}$ |
| G130V | 7.9 | **0** | 6.4 | **0** | $7.7\times10^{5}$ | $1.2\times10^{-2}$ | $3.6\times10^{4}$ | **$9.0\times10^{-4}$** | 5.8 | **0** | $6.9\times10^{2}$ | $1.3\times10^{-2}$ |
| Q131G | $7.1\times10^{1}$ | $2.3\times10^{-2}$ | $5.4\times10^{1}$ | **$6.5\times10^{-3}$** | $1.5\times10^{6}$ | $8.1\times10^{-1}$ | $5.1\times10^{5}$ | $3.0\times10^{-1}$ | $3.2\times10^{1}$ | **$1.\times10^{-4}$** | $6.1\times10^{2}$ | $7.0\times10^{-1}$ |
| H132G | $4.5\times10^{1}$ | **$9.0\times10^{-4}$** | $4.2\times10^{1}$ | **$2.0\times10^{-4}$** | $1.0\times10^{6}$ | $2.4\times10^{-1}$ | $3.3\times10^{5}$ | $2.7\times10^{-2}$ | $3.9\times10^{1}$ | **$1.\times10^{-4}$** | $6.6\times10^{2}$ | $1.2\times10^{-1}$ |
| H132V | $6.0\times10^{1}$ | $2.1\times10^{-2}$ | $1.1\times10^{2}$ | $6.8\times10^{-2}$ | $6.2\times10^{5}$ | $2.9\times10^{-2}$ | $\leq 1.2\times10^{5}$ | **$9.0\times10^{-4}$** | $3.5\times10^{1}$ | **$5.0\times10^{-4}$** | $8.5\times10^{2}$ | **$1.9\times10^{-3}$** |
| S133G | $7.3\times10^{1}$ | $8.7\times10^{-2}$ | $5.4\times10^{1}$ | $3.3\times10^{-2}$ | $1.5\times10^{6}$ | $8.1\times10^{-1}$ | $3.0\times10^{5}$ | $1.9\times10^{-2}$ | $1.3\times10^{1}$ | $4.3\times10^{-1}$ | $4.4\times10^{2}$ | **$1.4\times10^{-3}$** |
| S133V | – | – | – | – | $\leq 1.3\times10^{3}$ | **0** | $\leq 1.2\times10^{3}$ | **0** | ≤ 1.3 | **0** | – | – |
| P134G | $1.3\times10^{1}$ | **0** | $6.5\times10^{1}$ | $2.5\times10^{-2}$ | $1.8\times10^{5}$ | **0** | $6.8\times10^{4}$ | **0** | 2.6 | **0** | $5.7\times10^{2}$ | $1.3\times10^{-1}$ |
| P134V | $1.2\times10^{2}$ | $8.0\times10^{-1}$ | $8.0\times10^{1}$ | $5.5\times10^{-1}$ | $1.6\times10^{6}$ | $5.7\times10^{-1}$ | $2.6\times10^{5}$ | **$4.1\times10^{-3}$** | 7.6 | $2.1\times10^{-2}$ | $5.5\times10^{2}$ | $5.8\times10^{-2}$ |

| Variant | $k_{cat}$ cMUP ($s^{-1}$) | $k_{cat}$ cMUP *p*–value | $K_M$ cMUP (μM) | $K_M$ cMUP *p*–value | $k_{cat}/K_M$ cMUP ($M^{-1}s^{-1}$) | $k_{cat}/K_M$ cMUP *p*–value | $k_{cat}/K_M$ MeP ($M^{-1}s^{-1}$) | $k_{cat}/K_M$ MeP *p*–value | $k_{cat}/K_M$ MecMUP ($M^{-1}s^{-1}$) | $k_{cat}/K_M$ MecMUP *p*–value | $K_i$ $PO_4$ (μM) | $K_i$ $PO_4$ *p*–value |
|---|---|---|---|---|---|---|---|---|---|---|---|---|
| R135G | $2.5\times10^{2}$ | **0** | $7.7\times10^{1}$ | $4.3\times10^{-1}$ | $3.2\times10^{6}$ | **0** | $5.4\times10^{5}$ | $5.6\times10^{-1}$ | $3.6\times10^{1}$ | **0** | $5.9\times10^{2}$ | $3.7\times10^{-1}$ |
| R135V | $2.3\times10^{2}$ | **0** | $1.1\times10^{2}$ | $5.3\times10^{-2}$ | $1.9\times10^{6}$ | $1.4\times10^{-1}$ | $8.4\times10^{5}$ | $4.1\times10^{-2}$ | $2.3\times10^{1}$ | $5.8\times10^{-2}$ | $7.0\times10^{2}$ | $3.4\times10^{-2}$ |
| N136G | $4.7\times10^{1}$ | **$1.0\times10^{-3}$** | $6.1\times10^{1}$ | $2.3\times10^{-2}$ | $8.6\times10^{5}$ | $9.8\times10^{-2}$ | $3.4\times10^{5}$ | $3.6\times10^{-2}$ | $1.1\times10^{1}$ | $1.9\times10^{-1}$ | $5.8\times10^{2}$ | $3.2\times10^{-1}$ |
| N136V | $2.3\times10^{1}$ | **$1.\times10^{-4}$** | $7.5\times10^{1}$ | $3.7\times10^{-1}$ | $4.8\times10^{5}$ | $1.4\times10^{-2}$ | $1.8\times10^{5}$ | **$2.0\times10^{-3}$** | 5.6 | **$4.9\times10^{-3}$** | $7.0\times10^{2}$ | $4.3\times10^{-2}$ |
| L137G | $2.1\times10^{1}$ | **$4.1\times10^{-3}$** | $4.8\times10^{1}$ | $1.1\times10^{-2}$ | $3.6\times10^{5}$ | $1.3\times10^{-2}$ | $7.3\times10^{4}$ | **$4.8\times10^{-3}$** | 8.1 | $1.2\times10^{-2}$ | $4.7\times10^{2}$ | **$3.8\times10^{-3}$** |
| L137V | $1.5\times10^{2}$ | $7.7\times10^{-2}$ | $1.1\times10^{2}$ | $1.6\times10^{-2}$ | $1.5\times10^{6}$ | $7.6\times10^{-1}$ | $5.0\times10^{5}$ | $2.7\times10^{-1}$ | $1.6\times10^{1}$ | $9.7\times10^{-1}$ | $7.0\times10^{2}$ | $1.1\times10^{-2}$ |
| W138G | $1.6\times10^{1}$ | **0** | $5.0\times10^{1}$ | **$1.\times10^{-4}$** | $4.7\times10^{5}$ | **$1.2\times10^{-3}$** | $1.0\times10^{5}$ | **0** | 4.7 | **0** | $4.3\times10^{2}$ | **0** |
| W138V | $3.5\times10^{1}$ | **$1.\times10^{-4}$** | $7.2\times10^{1}$ | $1.9\times10^{-1}$ | $5.1\times10^{5}$ | **$6.3\times10^{-3}$** | $9.5\times10^{4}$ | **$4.0\times10^{-4}$** | 4.7 | **$2.4\times10^{-3}$** | $5.3\times10^{2}$ | $1.3\times10^{-2}$ |
| S139G | $3.7\times10^{1}$ | **$4.0\times10^{-3}$** | $4.3\times10^{1}$ | **$4.2\times10^{-3}$** | $4.9\times10^{5}$ | $2.5\times10^{-2}$ | $2.5\times10^{5}$ | **$9.1\times10^{-3}$** | $1.0\times10^{1}$ | $6.8\times10^{-2}$ | $4.0\times10^{2}$ | **$1.0\times10^{-3}$** |
| S139V | $1.4\times10^{2}$ | $4.9\times10^{-1}$ | $8.4\times10^{1}$ | $8.9\times10^{-1}$ | $1.7\times10^{6}$ | $4.9\times10^{-1}$ | $4.0\times10^{5}$ | $1.2\times10^{-1}$ | $1.2\times10^{1}$ | $3.0\times10^{-1}$ | $5.9\times10^{2}$ | $3.5\times10^{-1}$ |
| T140G | $2.0\times10^{1}$ | **$4.0\times10^{-4}$** | $5.0\times10^{1}$ | **$5.0\times10^{-3}$** | $5.4\times10^{5}$ | $1.7\times10^{-2}$ | $1.3\times10^{5}$ | **$1.8\times10^{-3}$** | 6.8 | **$9.0\times10^{-4}$** | $4.4\times10^{2}$ | **$2.0\times10^{-4}$** |
| T140V | $4.4\times10^{1}$ | **$5.0\times10^{-4}$** | $7.8\times10^{1}$ | $3.9\times10^{-1}$ | $5.2\times10^{5}$ | **$4.0\times10^{-3}$** | $2.2\times10^{5}$ | **$4.8\times10^{-3}$** | $1.0\times10^{1}$ | $1.7\times10^{-1}$ | $5.0\times10^{2}$ | **$1.9\times10^{-3}$** |
| T141G | $2.6\times10^{1}$ | **0** | $5.2\times10^{1}$ | **$2.6\times10^{-3}$** | $7.8\times10^{5}$ | $5.6\times10^{-2}$ | $2.0\times10^{5}$ | **$5.4\times10^{-3}$** | 8.3 | **$6.9\times10^{-3}$** | $4.6\times10^{2}$ | **$3.0\times10^{-4}$** |
| T141V | $1.1\times10^{2}$ | $5.4\times10^{-1}$ | $7.9\times10^{1}$ | $4.5\times10^{-1}$ | $1.5\times10^{6}$ | $6.9\times10^{-1}$ | $9.3\times10^{4}$ | **$4.2\times10^{-3}$** | 7.6 | $2.0\times10^{-2}$ | $5.5\times10^{2}$ | $8.9\times10^{-2}$ |
| V142A | $1.4\times10^{2}$ | $3.9\times10^{-1}$ | $9.7\times10^{1}$ | $2.2\times10^{-1}$ | $1.5\times10^{6}$ | $8.8\times10^{-1}$ | $2.0\times10^{5}$ | **$3.0\times10^{-4}$** | $1.1\times10^{1}$ | $1.3\times10^{-1}$ | $5.6\times10^{2}$ | $2.9\times10^{-2}$ |
| V142G | $1.2\times10^{1}$ | **0** | $5.6\times10^{1}$ | **$7.4\times10^{-3}$** | $2.3\times10^{5}$ | **$3.0\times10^{-4}$** | $1.0\times10^{5}$ | **$1.\times10^{-4}$** | 2.7 | **$2.0\times10^{-4}$** | $4.7\times10^{2}$ | **0** |
| T143G | $6.3\times10^{1}$ | $2.6\times10^{-2}$ | $7.6\times10^{1}$ | $3.7\times10^{-1}$ | $8.5\times10^{5}$ | $1.3\times10^{-1}$ | $3.5\times10^{5}$ | $3.5\times10^{-2}$ | $1.2\times10^{1}$ | $3.3\times10^{-1}$ | $6.2\times10^{2}$ | $8.4\times10^{-1}$ |
| T143V | $8.6\times10^{1}$ | $5.4\times10^{-2}$ | $7.7\times10^{1}$ | $3.8\times10^{-1}$ | $1.2\times10^{6}$ | $5.4\times10^{-1}$ | $2.8\times10^{5}$ | **$3.6\times10^{-3}$** | 8.1 | $3.1\times10^{-2}$ | $6.3\times10^{2}$ | $5.5\times10^{-1}$ |
| D144G | $1.5\times10^{1}$ | **0** | $6.7\times10^{1}$ | $3.0\times10^{-2}$ | $2.4\times10^{5}$ | **0** | $4.2\times10^{4}$ | **0** | ≤ 1.2 | **0** | $5.3\times10^{2}$ | **$1.8\times10^{-3}$** |
| D144V | $6.2\times10^{-1}$ | **0** | $6.2\times10^{1}$ | $4.4\times10^{-2}$ | $9.5\times10^{3}$ | **$6.0\times10^{-4}$** | ≤ $1.3\times10^{3}$ | **$1.\times10^{-4}$** | ≤ 1.3 | **$1.\times10^{-4}$** | $3.7\times10^{2}$ | **$1.\times10^{-4}$** |
| Q145G | $7.4\times10^{1}$ | $1.0\times10^{-1}$ | $3.9\times10^{1}$ | **$1.9\times10^{-3}$** | $1.9\times10^{6}$ | $2.5\times10^{-1}$ | $4.3\times10^{5}$ | $1.4\times10^{-1}$ | $1.1\times10^{1}$ | $9.6\times10^{-2}$ | $3.4\times10^{2}$ | $1.4\times10^{-2}$ |
| Q145V | $1.2\times10^{2}$ | $9.3\times10^{-1}$ | $7.4\times10^{1}$ | $4.7\times10^{-1}$ | $1.6\times10^{6}$ | $7.6\times10^{-1}$ | $3.3\times10^{5}$ | $2.9\times10^{-2}$ | 8.9 | $3.3\times10^{-2}$ | $5.1\times10^{2}$ | **$2.9\times10^{-3}$** |
| L146G | 6.7 | **0** | $5.6\times10^{1}$ | **$2.0\times10^{-4}$** | $1.4\times10^{5}$ | **0** | $1.5\times10^{4}$ | **0** | ≤ 1.3 | **0** | $4.0\times10^{2}$ | **0** |
| L146V | $1.8\times10^{2}$ | $1.9\times10^{-2}$ | $8.7\times10^{1}$ | $8.6\times10^{-1}$ | $2.3\times10^{6}$ | **$8.4\times10^{-3}$** | $3.9\times10^{5}$ | $6.2\times10^{-2}$ | $1.1\times10^{1}$ | $1.5\times10^{-1}$ | $6.6\times10^{2}$ | $1.2\times10^{-1}$ |
| G147A | $9.1\times10^{1}$ | $1.9\times10^{-1}$ | $7.1\times10^{1}$ | $2.8\times10^{-1}$ | $9.8\times10^{5}$ | $2.2\times10^{-1}$ | $3.5\times10^{5}$ | $5.6\times10^{-2}$ | $1.7\times10^{1}$ | $8.4\times10^{-1}$ | $5.1\times10^{2}$ | $3.2\times10^{-2}$ |
| G147V | $1.1\times10^{2}$ | $5.3\times10^{-1}$ | $9.7\times10^{1}$ | $2.4\times10^{-1}$ | $1.2\times10^{6}$ | $4.3\times10^{-1}$ | $3.1\times10^{5}$ | $1.1\times10^{-2}$ | 8.4 | $2.0\times10^{-2}$ | $6.2\times10^{2}$ | $6.5\times10^{-1}$ |
| L148G | $1.5\times10^{1}$ | **$1.\times10^{-4}$** | $5.8\times10^{1}$ | $2.9\times10^{-2}$ | $2.0\times10^{5}$ | **$1.5\times10^{-3}$** | $1.1\times10^{5}$ | $1.5\times10^{-2}$ | ≤ $8.9\times10^{-1}$ | **0** | $3.9\times10^{2}$ | **0** |
| L148V | $6.1\times10^{1}$ | **$8.5\times10^{-3}$** | $8.5\times10^{1}$ | $9.5\times10^{-1}$ | $7.7\times10^{5}$ | $5.1\times10^{-2}$ | $1.6\times10^{5}$ | **$2.7\times10^{-3}$** | 7.1 | $1.6\times10^{-2}$ | $6.4\times10^{2}$ | $4.4\times10^{-1}$ |
| A149G | $1.2\times10^{2}$ | $6.8\times10^{-1}$ | $6.4\times10^{1}$ | $6.7\times10^{-2}$ | $1.7\times10^{6}$ | $4.9\times10^{-1}$ | $2.7\times10^{5}$ | $4.1\times10^{-2}$ | $1.9\times10^{1}$ | $4.7\times10^{-1}$ | $6.2\times10^{2}$ | $6.4\times10^{-1}$ |
| A149V | $3.7\times10^{1}$ | **$8.0\times10^{-4}$** | $7.2\times10^{1}$ | $2.3\times10^{-1}$ | $5.2\times10^{5}$ | $1.1\times10^{-2}$ | $3.3\times10^{5}$ | $1.4\times10^{-2}$ | 6.8 | $1.1\times10^{-2}$ | $4.5\times10^{2}$ | **$7.0\times10^{-4}$** |
| T150G | $6.4\times10^{1}$ | $4.9\times10^{-2}$ | $6.3\times10^{1}$ | $8.3\times10^{-2}$ | $1.4\times10^{6}$ | $9.1\times10^{-1}$ | $3.2\times10^{5}$ | $2.5\times10^{-2}$ | $1.0\times10^{1}$ | $9.0\times10^{-2}$ | $5.0\times10^{2}$ | **$1.3\times10^{-3}$** |
| T150V | $9.2\times10^{1}$ | $1.1\times10^{-1}$ | $8.1\times10^{1}$ | $5.6\times10^{-1}$ | $1.2\times10^{6}$ | $3.8\times10^{-1}$ | $2.6\times10^{5}$ | **$3.1\times10^{-3}$** | 8.8 | $5.4\times10^{-2}$ | $5.6\times10^{2}$ | $5.8\times10^{-2}$ |
| N151G | 9.4 | **$4.0\times10^{-4}$** | $6.4\times10^{1}$ | $1.1\times10^{-1}$ | $1.7\times10^{5}$ | **$3.4\times10^{-3}$** | $1.8\times10^{4}$ | **0** | ≤ 1.2 | **0** | $3.7\times10^{2}$ | **$1.6\times10^{-3}$** |
| N151V | $1.2\times10^{2}$ | $8.3\times10^{-1}$ | $9.0\times10^{1}$ | $6.0\times10^{-1}$ | $1.3\times10^{6}$ | $8.0\times10^{-1}$ | $1.5\times10^{5}$ | **$3.1\times10^{-3}$** | $1.2\times10^{1}$ | $3.5\times10^{-1}$ | $6.1\times10^{2}$ | $9.1\times10^{-1}$ |
| F152G | $3.7\times10^{1}$ | **$3.0\times10^{-4}$** | $6.0\times10^{1}$ | $2.1\times10^{-2}$ | $4.3\times10^{5}$ | **$3.4\times10^{-3}$** | $1.5\times10^{5}$ | **$8.6\times10^{-3}$** | 8.9 | $3.8\times10^{-2}$ | $5.4\times10^{2}$ | $2.2\times10^{-2}$ |
| F152V | 7.5 | **$1.5\times10^{-3}$** | $7.6\times10^{1}$ | $4.9\times10^{-1}$ | $9.9\times10^{4}$ | **$2.6\times10^{-3}$** | $1.0\times10^{5}$ | **$4.0\times10^{-4}$** | ≤ 1.3 | **$4.0\times10^{-4}$** | $5.1\times10^{2}$ | $1.5\times10^{-2}$ |
| T153G | $9.2\times10^{1}$ | $1.3\times10^{-1}$ | $6.5\times10^{1}$ | $5.5\times10^{-2}$ | $1.4\times10^{6}$ | $8.5\times10^{-1}$ | $3.5\times10^{5}$ | $4.2\times10^{-2}$ | $2.0\times10^{1}$ | $3.6\times10^{-1}$ | $5.3\times10^{2}$ | $1.5\times10^{-2}$ |
| T153V | $1.9\times10^{2}$ | **$1.4\times10^{-3}$** | $9.2\times10^{1}$ | $4.4\times10^{-1}$ | $2.0\times10^{6}$ | $6.6\times10^{-2}$ | $4.0\times10^{5}$ | $1.0\times10^{-1}$ | $1.8\times10^{1}$ | $6.7\times10^{-1}$ | $6.7\times10^{2}$ | $1.2\times10^{-1}$ |
| S154G | $8.9\times10^{1}$ | $1.9\times10^{-1}$ | $7.5\times10^{1}$ | $3.5\times10^{-1}$ | $1.6\times10^{6}$ | $7.6\times10^{-1}$ | $5.2\times10^{5}$ | $4.4\times10^{-1}$ | $1.4\times10^{1}$ | $4.8\times10^{-1}$ | $5.8\times10^{2}$ | $3.8\times10^{-1}$ |
| S154V | $3.5\times10^{1}$ | **$8.0\times10^{-4}$** | $7.9\times10^{1}$ | $5.8\times10^{-1}$ | $4.7\times10^{5}$ | $1.2\times10^{-2}$ | $8.6\times10^{4}$ | **$1.0\times10^{-3}$** | 3.9 | **$1.9\times10^{-3}$** | $5.3\times10^{2}$ | **$8.5\times10^{-3}$** |
| K155G | $4.2\times10^{1}$ | **$1.5\times10^{-3}$** | $7.3\times10^{1}$ | $3.0\times10^{-1}$ | $6.3\times10^{5}$ | $3.4\times10^{-2}$ | $2.2\times10^{5}$ | **$5.5\times10^{-3}$** | 6.5 | **$1.8\times10^{-3}$** | $5.4\times10^{2}$ | $3.0\times10^{-2}$ |
| K155V | $5.9\times10^{1}$ | $1.2\times10^{-2}$ | $9.0\times10^{1}$ | $6.9\times10^{-1}$ | $5.3\times10^{5}$ | $1.6\times10^{-2}$ | $3.6\times10^{5}$ | $2.4\times10^{-2}$ | ≤ 2.3 | **$2.9\times10^{-3}$** | $5.9\times10^{2}$ | $3.7\times10^{-1}$ |
| V156A | $1.3\times10^{2}$ | $5.7\times10^{-1}$ | $8.3\times10^{1}$ | $7.8\times10^{-1}$ | $1.5\times10^{6}$ | $6.9\times10^{-1}$ | $2.5\times10^{5}$ | $1.0\times10^{-2}$ | $1.2\times10^{1}$ | $2.5\times10^{-1}$ | $5.9\times10^{2}$ | $5.4\times10^{-1}$ |
| V156G | $1.0\times10^{2}$ | $4.2\times10^{-1}$ | $6.9\times10^{1}$ | $2.7\times10^{-1}$ | $1.3\times10^{6}$ | $7.8\times10^{-1}$ | $1.6\times10^{5}$ | **$2.2\times10^{-3}$** | ≤ 2.8 | **0** | $5.0\times10^{2}$ | **$4.5\times10^{-3}$** |
| V157A | $5.7\times10^{1}$ | $1.4\times10^{-2}$ | $9.0\times10^{1}$ | $6.5\times10^{-1}$ | $5.5\times10^{5}$ | $1.9\times10^{-2}$ | $3.2\times10^{5}$ | $1.2\times10^{-2}$ | $1.0\times10^{1}$ | $1.2\times10^{-1}$ | $5.7\times10^{2}$ | $1.4\times10^{-1}$ |
| V157G | $1.6\times10^{1}$ | **0** | $6.0\times10^{1}$ | **$6.2\times10^{-3}$** | $2.7\times10^{5}$ | **$3.0\times10^{-4}$** | $3.6\times10^{4}$ | **0** | 1.9 | **$4.0\times10^{-4}$** | $5.7\times10^{2}$ | $8.3\times10^{-2}$ |
| G158A | $1.0\times10^{2}$ | $3.3\times10^{-1}$ | $6.4\times10^{1}$ | $8.3\times10^{-2}$ | $8.1\times10^{5}$ | $1.1\times10^{-1}$ | $4.5\times10^{5}$ | $2.9\times10^{-1}$ | $1.6\times10^{1}$ | $9.4\times10^{-1}$ | $5.5\times10^{2}$ | $8.1\times10^{-2}$ |
| G158V | $1.6\times10^{1}$ | **0** | $6.6\times10^{1}$ | $4.4\times10^{-2}$ | $2.3\times10^{5}$ | **$1.\times10^{-4}$** | $6.0\times10^{4}$ | **$4.0\times10^{-4}$** | 1.1 | **$1.\times10^{-4}$** | $3.9\times10^{2}$ | **0** |
| V159A | $1.4\times10^{2}$ | $3.6\times10^{-1}$ | $8.7\times10^{1}$ | $8.8\times10^{-1}$ | $1.7\times10^{6}$ | $4.5\times10^{-1}$ | $9.1\times10^{4}$ | **0** | $1.0\times10^{1}$ | $1.6\times10^{-1}$ | $5.9\times10^{2}$ | $5.5\times10^{-1}$ |
| V159G | $6.8\times10^{1}$ | $7.9\times10^{-2}$ | $6.7\times10^{1}$ | $1.9\times10^{-1}$ | $9.7\times10^{5}$ | $3.1\times10^{-1}$ | $3.3\times10^{5}$ | $3.7\times10^{-2}$ | $1.4\times10^{1}$ | $4.9\times10^{-1}$ | $5.0\times10^{2}$ | $2.4\times10^{-2}$ |
| S160G | $4.1\times10^{1}$ | **$3.0\times10^{-4}$** | $4.3\times10^{1}$ | **$4.0\times10^{-4}$** | $1.1\times10^{6}$ | $4.2\times10^{-1}$ | $8.6\times10^{5}$ | $1.0\times10^{-1}$ | $1.2\times10^{1}$ | $1.6\times10^{-1}$ | $3.8\times10^{2}$ | **0** |
| S160V | – | – | – | – | ≤ $1.3\times10^{3}$ | **0** | ≤ $7.1\times10^{2}$ | **$5.0\times10^{-4}$** | ≤ 1.0 | **0** | – | – |
| L161G | $2.6\times10^{1}$ | **0** | $1.1\times10^{2}$ | **$6.0\times10^{-3}$** | $2.6\times10^{5}$ | **0** | $1.4\times10^{4}$ | **$1.\times10^{-4}$** | 3.8 | **$6.0\times10^{-4}$** | $1.5\times10^{3}$ | **0** |
| L161V | $2.7\times10^{2}$ | **0** | $1.0\times10^{2}$ | $8.3\times10^{-2}$ | $2.6\times10^{6}$ | **$6.0\times10^{-4}$** | $4.8\times10^{5}$ | $1.1\times10^{-1}$ | $2.1\times10^{1}$ | $1.3\times10^{-1}$ | $7.2\times10^{2}$ | **$3.3\times10^{-3}$** |
| K162G | $2.8\times10^{-2}$ | **0** | $1.0\times10^{2}$ | $2.2\times10^{-1}$ | $2.8\times10^{2}$ | **$1.\times10^{-4}$** | ≤ $1.2\times10^{3}$ | **0** | $8.8\times10^{1}$ | **0** | – | – |
| K162V | $3.1\times10^{-2}$ | **0** | $4.6\times10^{1}$ | **$2.3\times10^{-3}$** | $6.7\times10^{2}$ | **$4.0\times10^{-4}$** | ≤ $1.2\times10^{3}$ | **0** | $5.2\times10^{2}$ | **0** | $2.3\times10^{3}$ | **$1.\times10^{-4}$** |
| D163G | $1.1\times10^{1}$ | **$2.0\times10^{-4}$** | $3.6\times10^{1}$ | **$1.8\times10^{-3}$** | $3.1\times10^{5}$ | $1.0\times10^{-2}$ | $9.3\times10^{4}$ | **$1.5\times10^{-3}$** | $1.1\times10^{1}$ | $6.4\times10^{-2}$ | $2.3\times10^{3}$ | **$1.\times10^{-4}$** |
| D163V | 3.6 | **0** | $2.6\times10^{1}$ | **0** | $1.4\times10^{5}$ | **$1.\times10^{-4}$** | $7.2\times10^{3}$ | **$8.0\times10^{-4}$** | 5.1 | **$4.6\times10^{-3}$** | ≥ $5.1\times10^{3}$ | **0** |
| R164G | 4.6 | **0** | $1.2\times10^{1}$ | **0** | $3.8\times10^{5}$ | **$1.\times10^{-4}$** | $2.1\times10^{3}$ | **$4.0\times10^{-4}$** | $1.1\times10^{2}$ | **0** | ≥ $2.1\times10^{3}$ | **0** |
| R164V | 9.9 | **0** | $4.4\times10^{1}$ | **0** | $2.4\times10^{5}$ | **$1.\times10^{-4}$** | $2.9\times10^{3}$ | **$5.0\times10^{-4}$** | $1.0\times10^{2}$ | **0** | ≥ $1.3\times10^{5}$ | **$1.\times10^{-4}$** |
| A165G | $2.6\times10^{2}$ | **$6.0\times10^{-4}$** | $1.2\times10^{2}$ | $3.1\times10^{-2}$ | $2.4\times10^{6}$ | $1.9\times10^{-2}$ | $2.1\times10^{5}$ | **$4.7\times10^{-3}$** | 6.6 | **$2.5\times10^{-3}$** | $8.8\times10^{2}$ | **$3.0\times10^{-4}$** |
| A165V | $1.8\times10^{1}$ | **0** | $3.4\times10^{2}$ | **0** | $5.8\times10^{4}$ | **0** | ≤ $1.1\times10^{3}$ | **0** | ≤ 1.2 | **$2.0\times10^{-4}$** | $2.6\times10^{3}$ | **$1.\times10^{-4}$** |
| S166G | $1.6\times10^{2}$ | $1.6\times10^{-1}$ | $7.8\times10^{1}$ | $6.2\times10^{-1}$ | $2.3\times10^{6}$ | $5.7\times10^{-2}$ | $3.2\times10^{5}$ | $3.6\times10^{-2}$ | $2.4\times10^{1}$ | **$9.6\times10^{-3}$** | $6.6\times10^{2}$ | $2.4\times10^{-1}$ |
| S166V | $1.8\times10^{2}$ | $1.6\times10^{-2}$ | $5.8\times10^{1}$ | $2.3\times10^{-2}$ | $2.5\times10^{6}$ | **$6.5\times10^{-3}$** | $3.7\times10^{5}$ | $5.5\times10^{-2}$ | $1.8\times10^{1}$ | $8.5\times10^{-1}$ | $4.2\times10^{2}$ | **$6.0\times10^{-4}$** |
| I167G | – | – | – | – | ≤ $1.2\times10^{3}$ | **0** | ≤ $3.4\times10^{3}$ | **$1.\times10^{-4}$** | ≤ $9.9\times10^{-1}$ | **0** | – | – |
| I167V | $1.3\times10^{2}$ | $5.9\times10^{-1}$ | $9.8\times10^{1}$ | $1.9\times10^{-1}$ | $1.4\times10^{6}$ | $9.8\times10^{-1}$ | $5.2\times10^{5}$ | $4.7\times10^{-1}$ | $1.2\times10^{1}$ | $3.3\times10^{-1}$ | $7.3\times10^{2}$ | **$2.7\times10^{-3}$** |
| L168G | 8.4 | **$1.\times10^{-4}$** | $7.7\times10^{1}$ | $4.3\times10^{-1}$ | $1.0\times10^{5}$ | **$6.0\times10^{-4}$** | $6.1\times10^{4}$ | **0** | ≤ 1.2 | **0** | $2.6\times10^{2}$ | **0** |
| L168V | $4.9\times10^{1}$ | **$1.0\times10^{-3}$** | $8.0\times10^{1}$ | $5.3\times10^{-1}$ | $6.5\times10^{5}$ | $2.0\times10^{-2}$ | $2.1\times10^{5}$ | **$3.7\times10^{-3}$** | 4.6 | **$2.0\times10^{-4}$** | $6.1\times10^{2}$ | $9.5\times10^{-1}$ |
| P169G | $2.1\times10^{1}$ | **0** | $5.3\times10^{1}$ | **$3.0\times10^{-4}$** | $4.1\times10^{5}$ | **$2.0\times10^{-4}$** | ≥ $3.2\times10^{5}$ | **$3.1\times10^{-3}$** | 9.6 | $1.4\times10^{-2}$ | $9.7\times10^{1}$ | **0** |
| P169V | $1.5\times10^{1}$ | **0** | $5.5\times10^{1}$ | **$6.0\times10^{-3}$** | $2.7\times10^{5}$ | **$1.0\times10^{-3}$** | $2.7\times10^{5}$ | $1.2\times10^{-2}$ | ≤ 2.5 | **$6.0\times10^{-4}$** | $2.0\times10^{2}$ | **$3.0\times10^{-4}$** |
| A170G | $1.0\times10^{2}$ | $3.7\times10^{-1}$ | $7.4\times10^{1}$ | $4.8\times10^{-1}$ | $1.5\times10^{6}$ | $9.3\times10^{-1}$ | $1.6\times10^{5}$ | **$2.4\times10^{-3}$** | $1.5\times10^{1}$ | $7.4\times10^{-1}$ | $5.8\times10^{2}$ | $2.9\times10^{-1}$ |
| A170V | $1.0\times10^{2}$ | $3.7\times10^{-1}$ | $8.7\times10^{1}$ | $8.8\times10^{-1}$ | $1.2\times10^{6}$ | $4.7\times10^{-1}$ | $2.7\times10^{5}$ | $1.5\times10^{-2}$ | $1.0\times10^{1}$ | $8.1\times10^{-2}$ | $6.4\times10^{2}$ | $4.4\times10^{-1}$ |
| G171A | 8.4 | **$4.3\times10^{-3}$** | $5.1\times10^{1}$ | $5.1\times10^{-2}$ | $1.7\times10^{5}$ | $1.6\times10^{-2}$ | $9.8\times10^{4}$ | **$2.1\times10^{-3}$** | 4.6 | **$2.0\times10^{-4}$** | $4.9\times10^{2}$ | **$3.8\times10^{-3}$** |
| G171V | – | – | – | – | ≤ $3.8\times10^{2}$ | **0** | ≤ $4.9\times10^{2}$ | **$6.0\times10^{-4}$** | ≤ 1.3 | **0** | – | – |
| H172V | $2.6\times10^{1}$ | **0** | $7.6\times10^{1}$ | $2.7\times10^{-1}$ | $3.6\times10^{5}$ | **$1.1\times10^{-3}$** | $7.1\times10^{4}$ | **$1.\times10^{-4}$** | ≤ 2.2 | **$2.5\times10^{-3}$** | $5.0\times10^{2}$ | **$9.0\times10^{-4}$** |

| Variant | $k_{cat}$ cMUP ($s^{-1}$) | $k_{cat}$ cMUP *p*–value | $K_M$ cMUP (μM) | $K_M$ cMUP *p*–value | $k_{cat}/K_M$ cMUP ($M^{-1}s^{-1}$) | $k_{cat}/K_M$ cMUP *p*–value | $k_{cat}/K_M$ MeP ($M^{-1}s^{-1}$) | $k_{cat}/K_M$ MeP *p*–value | $k_{cat}/K_M$ MecMUP ($M^{-1}s^{-1}$) | $k_{cat}/K_M$ MecMUP *p*–value | $K_i$ $PO_4$ (μM) | $K_i$ $PO_4$ *p*–value |
|---|---|---|---|---|---|---|---|---|---|---|---|---|
| N173G | $4.4\times10^{1}$ | **$1.0\times10^{-3}$** | $5.1\times10^{1}$ | **$3.5\times10^{-3}$** | $7.3\times10^{5}$ | $4.0\times10^{-2}$ | $2.3\times10^{5}$ | **$6.7\times10^{-3}$** | $1.3\times10^{1}$ | $2.6\times10^{-1}$ | $4.6\times10^{2}$ | **$1.\times10^{-4}$** |
| N173V | $2.9\times10^{1}$ | **0** | $8.2\times10^{1}$ | $7.0\times10^{-1}$ | $3.8\times10^{5}$ | **$8.0\times10^{-4}$** | $9.9\times10^{4}$ | **$3.0\times10^{-4}$** | $\le 1.9$ | **$1.9\times10^{-3}$** | $5.4\times10^{2}$ | $3.1\times10^{-2}$ |
| P174G | $1.1\times10^{1}$ | **0** | $5.3\times10^{1}$ | **0** | $2.3\times10^{5}$ | **0** | $7.4\times10^{4}$ | **0** | 3.9 | **0** | $4.4\times10^{2}$ | **0** |
| P174V | $1.1\times10^{1}$ | **0** | $6.7\times10^{1}$ | $4.4\times10^{-2}$ | $1.9\times10^{5}$ | **0** | $3.7\times10^{4}$ | **$3.0\times10^{-4}$** | 1.7 | **$3.0\times10^{-4}$** | $4.5\times10^{2}$ | **0** |
| T175G | $5.1\times10^{1}$ | **$2.0\times10^{-3}$** | $7.4\times10^{1}$ | $2.6\times10^{-1}$ | $6.8\times10^{5}$ | $2.5\times10^{-2}$ | $2.1\times10^{5}$ | **$5.0\times10^{-3}$** | $1.4\times10^{1}$ | $5.6\times10^{-1}$ | $5.5\times10^{2}$ | $2.6\times10^{-2}$ |
| T175V | $1.8\times10^{2}$ | **$7.2\times10^{-3}$** | $1.1\times10^{2}$ | $2.8\times10^{-2}$ | $1.8\times10^{6}$ | $2.4\times10^{-1}$ | $6.2\times10^{5}$ | $8.7\times10^{-1}$ | $1.4\times10^{1}$ | $3.7\times10^{-1}$ | $6.7\times10^{2}$ | $4.5\times10^{-2}$ |
| G176A | $8.4\times10^{1}$ | $1.0\times10^{-1}$ | $5.7\times10^{1}$ | $1.9\times10^{-2}$ | $1.6\times10^{6}$ | $7.4\times10^{-1}$ | $\le 3.1\times10^{5}$ | $6.9\times10^{-2}$ | $1.8\times10^{1}$ | $4.1\times10^{-1}$ | $5.2\times10^{2}$ | $1.5\times10^{-2}$ |
| G176V | $3.6\times10^{1}$ | **$9.0\times10^{-4}$** | $8.6\times10^{1}$ | $9.3\times10^{-1}$ | $4.6\times10^{5}$ | **$7.7\times10^{-3}$** | $2.4\times10^{5}$ | $1.0\times10^{-2}$ | $\le 1.9$ | **$1.4\times10^{-3}$** | $5.0\times10^{2}$ | **$6.7\times10^{-3}$** |
| A177G | $2.8\times10^{1}$ | **$8.0\times10^{-4}$** | $5.6\times10^{1}$ | $1.7\times10^{-2}$ | $4.0\times10^{5}$ | **$6.5\times10^{-3}$** | $1.0\times10^{5}$ | **$1.6\times10^{-3}$** | 5.6 | **$3.0\times10^{-4}$** | $6.0\times10^{2}$ | $5.7\times10^{-1}$ |
| A177V | $1.1\times10^{2}$ | $5.9\times10^{-1}$ | $9.1\times10^{1}$ | $4.4\times10^{-1}$ | $1.3\times10^{6}$ | $6.0\times10^{-1}$ | $3.5\times10^{5}$ | $2.0\times10^{-2}$ | 7.6 | $1.2\times10^{-2}$ | $6.5\times10^{2}$ | $2.0\times10^{-1}$ |
| F178G | 1.7 | **0** | $6.7\times10^{1}$ | $8.1\times10^{-2}$ | $1.8\times10^{4}$ | **$1.\times10^{-4}$** | $\le 5.7\times10^{3}$ | **0** | $\le 1.1$ | **$3.0\times10^{-4}$** | $5.7\times10^{2}$ | $1.2\times10^{-1}$ |
| F178V | $1.0\times10^{2}$ | $2.6\times10^{-1}$ | $9.1\times10^{1}$ | $5.9\times10^{-1}$ | $1.2\times10^{6}$ | $5.7\times10^{-1}$ | $3.6\times10^{5}$ | $3.1\times10^{-2}$ | $1.2\times10^{1}$ | $3.4\times10^{-1}$ | $5.7\times10^{2}$ | $1.4\times10^{-1}$ |
| W179G | – | – | – | – | $\le 4.2\times10^{2}$ | **0** | $\le 8.4\times10^{2}$ | **0** | $\le 1.3$ | **0** | – | – |
| W179V | 2.7 | $3.8\times10^{-2}$ | $5.9\times10^{1}$ | $2.9\times10^{-1}$ | $4.6\times10^{4}$ | $6.0\times10^{-2}$ | $\le 1.4\times10^{4}$ | **0** | $\le 1.8$ | **$1.8\times10^{-3}$** | $5.7\times10^{2}$ | $5.1\times10^{-1}$ |
| F180G | $3.9\times10^{1}$ | **$5.0\times10^{-4}$** | $4.5\times10^{1}$ | **$9.0\times10^{-4}$** | $1.1\times10^{6}$ | $2.9\times10^{-1}$ | $3.7\times10^{4}$ | **0** | 6.5 | **$1.5\times10^{-3}$** | $5.4\times10^{2}$ | $6.7\times10^{-2}$ |
| F180V | $1.3\times10^{2}$ | $6.4\times10^{-1}$ | $4.8\times10^{1}$ | $4.3\times10^{-2}$ | $1.9\times10^{6}$ | $3.7\times10^{-1}$ | $1.0\times10^{5}$ | $1.7\times10^{-2}$ | $1.4\times10^{1}$ | $6.8\times10^{-1}$ | $4.2\times10^{2}$ | **$4.0\times10^{-4}$** |
| D181G | $3.0\times10^{1}$ | **0** | $7.9\times10^{1}$ | $5.0\times10^{-1}$ | $3.4\times10^{5}$ | **$4.0\times10^{-4}$** | $4.0\times10^{4}$ | **0** | 2.0 | **$1.0\times10^{-3}$** | $7.5\times10^{2}$ | **$5.0\times10^{-4}$** |
| D181V | $5.4\times10^{1}$ | $1.1\times10^{-2}$ | $6.2\times10^{1}$ | $4.6\times10^{-2}$ | $8.6\times10^{5}$ | $1.3\times10^{-1}$ | $2.5\times10^{4}$ | **0** | $1.0\times10^{1}$ | $1.8\times10^{-1}$ | – | – |
| D182G | $6.2\times10^{1}$ | $5.9\times10^{-2}$ | $6.5\times10^{1}$ | $1.0\times10^{-1}$ | $1.5\times10^{6}$ | $8.4\times10^{-1}$ | $5.9\times10^{4}$ | **0** | 8.2 | **$6.7\times10^{-3}$** | $1.1\times10^{3}$ | **0** |
| D182V | $2.6\times10^{2}$ | **0** | $5.3\times10^{1}$ | **$4.1\times10^{-3}$** | $4.1\times10^{6}$ | **0** | $1.4\times10^{5}$ | **$1.5\times10^{-3}$** | $2.1\times10^{1}$ | $2.7\times10^{-1}$ | $5.6\times10^{2}$ | $1.3\times10^{-1}$ |
| T183G | $8.6\times10^{1}$ | $1.1\times10^{-1}$ | $7.0\times10^{1}$ | $2.3\times10^{-1}$ | $1.3\times10^{6}$ | $7.4\times10^{-1}$ | $4.8\times10^{5}$ | $2.1\times10^{-1}$ | $1.6\times10^{1}$ | $9.7\times10^{-1}$ | $4.3\times10^{2}$ | **$1.3\times10^{-3}$** |
| T183V | $2.0\times10^{2}$ | **$7.0\times10^{-4}$** | $7.7\times10^{1}$ | $3.6\times10^{-1}$ | $2.1\times10^{6}$ | $4.4\times10^{-2}$ | $5.4\times10^{5}$ | $5.5\times10^{-1}$ | $1.9\times10^{1}$ | $4.3\times10^{-1}$ | $5.6\times10^{2}$ | $1.6\times10^{-1}$ |
| T184G | $7.1\times10^{1}$ | $4.0\times10^{-2}$ | $6.3\times10^{1}$ | $6.0\times10^{-2}$ | $1.4\times10^{6}$ | $9.0\times10^{-1}$ | $3.0\times10^{5}$ | $2.1\times10^{-2}$ | $1.1\times10^{1}$ | $6.3\times10^{-2}$ | $4.5\times10^{2}$ | **0** |
| T184V | $7.8\times10^{1}$ | $4.1\times10^{-2}$ | $7.2\times10^{1}$ | $1.8\times10^{-1}$ | $1.0\times10^{6}$ | $1.9\times10^{-1}$ | $3.7\times10^{5}$ | $3.7\times10^{-2}$ | $1.1\times10^{1}$ | $2.2\times10^{-1}$ | $5.8\times10^{2}$ | $1.7\times10^{-1}$ |
| G185A | $1.8\times10^{2}$ | $4.9\times10^{-2}$ | $8.5\times10^{1}$ | $9.8\times10^{-1}$ | $2.2\times10^{6}$ | $6.1\times10^{-2}$ | $6.4\times10^{5}$ | $8.3\times10^{-1}$ | $2.4\times10^{1}$ | $1.0\times10^{-2}$ | $5.2\times10^{2}$ | $1.8\times10^{-2}$ |
| G185V | $5.3\times10^{1}$ | **$7.3\times10^{-3}$** | $6.8\times10^{1}$ | $1.5\times10^{-1}$ | $7.5\times10^{5}$ | $7.2\times10^{-2}$ | $2.7\times10^{4}$ | **0** | $1.0\times10^{1}$ | $1.4\times10^{-1}$ | $3.5\times10^{2}$ | **$5.0\times10^{-4}$** |
| K186G | $1.1\times10^{2}$ | $4.2\times10^{-1}$ | $6.8\times10^{1}$ | $9.7\times10^{-2}$ | $1.7\times10^{6}$ | $4.2\times10^{-1}$ | $3.0\times10^{5}$ | $2.2\times10^{-2}$ | $1.7\times10^{1}$ | $8.4\times10^{-1}$ | $5.9\times10^{2}$ | $3.0\times10^{-1}$ |
| K186V | $1.8\times10^{2}$ | $1.3\times10^{-2}$ | $8.4\times10^{1}$ | $9.0\times10^{-1}$ | $1.9\times10^{6}$ | $1.2\times10^{-1}$ | $1.6\times10^{5}$ | **$3.0\times10^{-4}$** | $1.2\times10^{1}$ | $3.3\times10^{-1}$ | $6.4\times10^{2}$ | $4.3\times10^{-1}$ |
| F187G | $1.0\times10^{1}$ | **$1.\times10^{-4}$** | $5.2\times10^{1}$ | **$1.5\times10^{-3}$** | $1.7\times10^{5}$ | **$3.0\times10^{-4}$** | $1.6\times10^{4}$ | **$4.1\times10^{-3}$** | $\le 1.1$ | **0** | $6.2\times10^{2}$ | $8.6\times10^{-1}$ |
| F187V | $3.5\times10^{1}$ | **0** | $7.1\times10^{1}$ | $1.7\times10^{-1}$ | $5.0\times10^{5}$ | **$6.1\times10^{-3}$** | $1.0\times10^{5}$ | **$1.4\times10^{-3}$** | $\le 2.4$ | **$2.1\times10^{-3}$** | $6.0\times10^{2}$ | $4.5\times10^{-1}$ |
| I188V | $2.1\times10^{2}$ | **0** | $9.3\times10^{1}$ | $4.4\times10^{-1}$ | $1.9\times10^{6}$ | $1.7\times10^{-1}$ | $3.8\times10^{5}$ | $5.2\times10^{-2}$ | $1.9\times10^{1}$ | $5.1\times10^{-1}$ | $6.2\times10^{2}$ | $8.1\times10^{-1}$ |
| T189G | $1.9\times10^{1}$ | **$1.\times10^{-4}$** | $6.6\times10^{1}$ | $6.4\times10^{-2}$ | $2.8\times10^{5}$ | **$1.0\times10^{-3}$** | $7.0\times10^{4}$ | **0** | 2.1 | **$5.0\times10^{-4}$** | $5.4\times10^{2}$ | $4.2\times10^{-2}$ |
| S190G | $1.1\times10^{1}$ | **0** | $3.3\times10^{1}$ | **$1.\times10^{-4}$** | $4.0\times10^{5}$ | **$2.7\times10^{-3}$** | $8.0\times10^{4}$ | **$6.0\times10^{-4}$** | 3.4 | **0** | $1.0\times10^{3}$ | **0** |
| S190V | – | – | – | – | $\le 4.7\times10^{2}$ | **0** | $\le 7.7\times10^{2}$ | **$2.7\times10^{-3}$** | $\le 1.3$ | **0** | $1.9\times10^{2}$ | **$9.0\times10^{-4}$** |
| T191G | $1.6\times10^{2}$ | $1.8\times10^{-1}$ | $7.3\times10^{1}$ | $3.5\times10^{-1}$ | $2.1\times10^{6}$ | $9.7\times10^{-2}$ | $5.4\times10^{5}$ | $6.9\times10^{-1}$ | $2.3\times10^{1}$ | $6.5\times10^{-2}$ | $5.6\times10^{2}$ | $8.9\times10^{-2}$ |
| T191V | $1.9\times10^{1}$ | **$7.0\times10^{-4}$** | $7.0\times10^{1}$ | $1.9\times10^{-1}$ | $3.0\times10^{5}$ | **$3.1\times10^{-3}$** | $7.5\times10^{4}$ | **$3.9\times10^{-3}$** | $\le 1.9$ | **$2.0\times10^{-3}$** | $6.1\times10^{2}$ | $8.2\times10^{-1}$ |
| Y192V | 8.7 | **$5.1\times10^{-3}$** | $7.8\times10^{1}$ | $6.5\times10^{-1}$ | $1.1\times10^{5}$ | $1.2\times10^{-2}$ | $\le 3.6\times10^{4}$ | **$7.0\times10^{-4}$** | $\le 1.9$ | **$1.0\times10^{-3}$** | – | – |
| Y193G | – | – | – | – | $\le 6.0\times10^{2}$ | **0** | $\le 9.5\times10^{2}$ | **0** | $\le 1.3$ | **0** | – | – |
| Y193V | $2.6\times10^{1}$ | **$2.0\times10^{-4}$** | $7.6\times10^{1}$ | $3.9\times10^{-1}$ | $3.9\times10^{5}$ | **$4.8\times10^{-3}$** | $3.9\times10^{4}$ | **0** | $\le 1.9$ | **$3.0\times10^{-4}$** | $5.2\times10^{2}$ | $4.2\times10^{-2}$ |
| T194G | $1.3\times10^{2}$ | $7.2\times10^{-1}$ | $7.9\times10^{1}$ | $5.4\times10^{-1}$ | $2.1\times10^{6}$ | $4.6\times10^{-2}$ | $7.0\times10^{5}$ | $4.6\times10^{-1}$ | $1.9\times10^{1}$ | $5.5\times10^{-1}$ | $6.0\times10^{2}$ | $6.4\times10^{-1}$ |
| T194V | $1.3\times10^{2}$ | $5.5\times10^{-1}$ | $8.7\times10^{1}$ | $7.8\times10^{-1}$ | $1.1\times10^{6}$ | $3.7\times10^{-1}$ | $7.3\times10^{5}$ | $2.4\times10^{-1}$ | $1.3\times10^{1}$ | $3.9\times10^{-1}$ | $6.4\times10^{2}$ | $3.0\times10^{-1}$ |
| K195G | $1.7\times10^{2}$ | $3.9\times10^{-2}$ | $7.5\times10^{1}$ | $3.9\times10^{-1}$ | $2.2\times10^{6}$ | $4.6\times10^{-2}$ | $5.7\times10^{5}$ | $7.7\times10^{-1}$ | $1.9\times10^{1}$ | $5.5\times10^{-1}$ | $6.0\times10^{2}$ | $7.2\times10^{-1}$ |
| K195V | $1.8\times10^{2}$ | **$4.5\times10^{-3}$** | $8.7\times10^{1}$ | $8.4\times10^{-1}$ | $2.0\times10^{6}$ | $5.7\times10^{-2}$ | $3.8\times10^{5}$ | $3.3\times10^{-2}$ | $1.6\times10^{1}$ | $9.6\times10^{-1}$ | $6.6\times10^{2}$ | $1.4\times10^{-1}$ |
| E196G | $1.4\times10^{2}$ | $4.3\times10^{-1}$ | $6.7\times10^{1}$ | $1.3\times10^{-1}$ | $2.0\times10^{6}$ | $9.5\times10^{-2}$ | $4.6\times10^{5}$ | $2.0\times10^{-1}$ | $1.4\times10^{1}$ | $4.7\times10^{-1}$ | $5.4\times10^{2}$ | $4.3\times10^{-2}$ |
| E196V | $2.1\times10^{2}$ | **$8.0\times10^{-4}$** | $8.7\times10^{1}$ | $8.7\times10^{-1}$ | $2.1\times10^{6}$ | $5.8\times10^{-2}$ | $4.1\times10^{5}$ | $4.8\times10^{-2}$ | $1.4\times10^{1}$ | $6.4\times10^{-1}$ | $6.6\times10^{2}$ | $1.0\times10^{-1}$ |
| L197G | $3.0\times10^{1}$ | **$4.0\times10^{-4}$** | $7.5\times10^{1}$ | $3.7\times10^{-1}$ | $3.1\times10^{5}$ | **$2.2\times10^{-3}$** | $1.4\times10^{5}$ | **$2.4\times10^{-3}$** | 6.0 | **$8.1\times10^{-3}$** | $4.9\times10^{2}$ | **$1.3\times10^{-3}$** |
| L197V | $1.2\times10^{2}$ | $7.7\times10^{-1}$ | $1.0\times10^{2}$ | $1.0\times10^{-1}$ | $1.3\times10^{6}$ | $6.9\times10^{-1}$ | $4.8\times10^{5}$ | $1.3\times10^{-1}$ | $1.1\times10^{1}$ | $1.4\times10^{-1}$ | $6.0\times10^{2}$ | $7.3\times10^{-1}$ |
| P198G | $6.1\times10^{1}$ | $1.4\times10^{-2}$ | $7.0\times10^{1}$ | $2.1\times10^{-1}$ | $7.5\times10^{5}$ | $6.8\times10^{-2}$ | $1.1\times10^{5}$ | **$1.5\times10^{-3}$** | 8.9 | $2.3\times10^{-2}$ | $5.4\times10^{2}$ | $3.4\times10^{-2}$ |
| P198V | $1.7\times10^{2}$ | $4.5\times10^{-2}$ | $8.5\times10^{1}$ | $9.4\times10^{-1}$ | $1.9\times10^{6}$ | $2.3\times10^{-1}$ | $1.2\times10^{5}$ | **$1.7\times10^{-3}$** | 9.5 | $3.7\times10^{-2}$ | $6.0\times10^{2}$ | $6.9\times10^{-1}$ |
| K199G | $1.5\times10^{2}$ | $1.3\times10^{-1}$ | $8.1\times10^{1}$ | $7.0\times10^{-1}$ | $1.9\times10^{6}$ | $1.2\times10^{-1}$ | $6.6\times10^{5}$ | $6.6\times10^{-1}$ | $1.7\times10^{1}$ | $9.3\times10^{-1}$ | $5.8\times10^{2}$ | $3.2\times10^{-1}$ |
| K199V | $1.5\times10^{2}$ | $1.9\times10^{-1}$ | $8.4\times10^{1}$ | $8.8\times10^{-1}$ | $1.7\times10^{6}$ | $5.4\times10^{-1}$ | $5.4\times10^{5}$ | $6.6\times10^{-1}$ | $1.8\times10^{1}$ | $6.7\times10^{-1}$ | $6.5\times10^{2}$ | $2.2\times10^{-1}$ |
| W200G | 3.8 | **0** | $6.3\times10^{1}$ | $3.5\times10^{-2}$ | $6.1\times10^{4}$ | **0** | $\le 5.3\times10^{3}$ | **0** | $\le 1.2$ | **$1.\times10^{-4}$** | $5.5\times10^{2}$ | $1.4\times10^{-2}$ |
| W200V | $1.7\times10^{1}$ | **$2.4\times10^{-3}$** | $8.1\times10^{1}$ | $7.2\times10^{-1}$ | $2.1\times10^{5}$ | **$6.7\times10^{-3}$** | $2.3\times10^{4}$ | **0** | $\le 1.2$ | **0** | $4.4\times10^{2}$ | **$9.0\times10^{-4}$** |
| V201A | $2.1\times10^{2}$ | **$5.5\times10^{-3}$** | $1.0\times10^{2}$ | $2.8\times10^{-1}$ | $1.9\times10^{6}$ | $3.1\times10^{-1}$ | $3.4\times10^{5}$ | $3.6\times10^{-2}$ | $1.5\times10^{1}$ | $7.5\times10^{-1}$ | $6.7\times10^{2}$ | $1.7\times10^{-1}$ |
| V201G | $2.9\times10^{1}$ | **0** | $7.3\times10^{1}$ | $1.4\times10^{-1}$ | $3.7\times10^{5}$ | **$5.0\times10^{-4}$** | $7.1\times10^{4}$ | **$1.\times10^{-4}$** | 3.1 | **$1.4\times10^{-3}$** | $5.2\times10^{2}$ | **$4.1\times10^{-3}$** |
| N202G | $1.3\times10^{2}$ | $7.9\times10^{-1}$ | $7.7\times10^{1}$ | $3.9\times10^{-1}$ | $1.8\times10^{6}$ | $2.8\times10^{-1}$ | $3.1\times10^{5}$ | $2.7\times10^{-2}$ | $1.9\times10^{1}$ | $4.2\times10^{-1}$ | $5.8\times10^{2}$ | $1.5\times10^{-1}$ |
| N202V | $1.4\times10^{2}$ | $2.0\times10^{-1}$ | $9.6\times10^{1}$ | $2.5\times10^{-1}$ | $1.7\times10^{6}$ | $3.6\times10^{-1}$ | $5.1\times10^{5}$ | $3.9\times10^{-1}$ | $1.1\times10^{1}$ | $2.9\times10^{-1}$ | $6.6\times10^{2}$ | $1.3\times10^{-1}$ |
| D203G | $9.6\times10^{1}$ | $3.8\times10^{-1}$ | $7.3\times10^{1}$ | $2.5\times10^{-1}$ | $1.7\times10^{6}$ | $4.2\times10^{-1}$ | $\le 1.7\times10^{5}$ | $2.4\times10^{-2}$ | $1.5\times10^{1}$ | $6.6\times10^{-1}$ | $6.9\times10^{2}$ | $3.6\times10^{-2}$ |
| D203V | $1.4\times10^{2}$ | $2.7\times10^{-1}$ | $9.5\times10^{1}$ | $2.5\times10^{-1}$ | $1.3\times10^{6}$ | $6.8\times10^{-1}$ | $4.2\times10^{5}$ | $1.1\times10^{-1}$ | $1.4\times10^{1}$ | $6.0\times10^{-1}$ | $6.7\times10^{2}$ | $6.7\times10^{-2}$ |
| F204G | 2.2 | **$2.3\times10^{-3}$** | $7.0\times10^{1}$ | $2.6\times10^{-1}$ | $3.0\times10^{4}$ | **$2.5\times10^{-3}$** | $4.6\times10^{3}$ | **$9.2\times10^{-3}$** | $\le 9.9\times10^{-1}$ | **0** | $5.8\times10^{2}$ | $3.0\times10^{-1}$ |
| F204V | $8.4\times10^{1}$ | $6.5\times10^{-2}$ | $8.6\times10^{1}$ | $9.0\times10^{-1}$ | $9.7\times10^{5}$ | $1.7\times10^{-1}$ | $\le 8.3\times10^{4}$ | **$1.7\times10^{-3}$** | 9.8 | $1.7\times10^{-1}$ | $6.0\times10^{2}$ | $5.0\times10^{-1}$ |
| N205V | $4.2\times10^{1}$ | **$1.9\times10^{-3}$** | $7.7\times10^{1}$ | $4.7\times10^{-1}$ | $5.3\times10^{5}$ | $1.8\times10^{-2}$ | $3.1\times10^{4}$ | **0** | 4.0 | **$2.5\times10^{-3}$** | $4.7\times10^{2}$ | **$2.0\times10^{-3}$** |
| N206G | $7.8\times10^{1}$ | $3.1\times10^{-2}$ | $7.1\times10^{1}$ | $1.3\times10^{-1}$ | $1.1\times10^{6}$ | $3.4\times10^{-1}$ | $1.5\times10^{6}$ | **$2.0\times10^{-3}$** | $1.9\times10^{1}$ | $4.4\times10^{-1}$ | $5.4\times10^{2}$ | $3.1\times10^{-2}$ |
| N206V | $1.3\times10^{2}$ | $5.0\times10^{-1}$ | $8.6\times10^{1}$ | $8.7\times10^{-1}$ | $1.4\times10^{6}$ | $9.3\times10^{-1}$ | $4.3\times10^{5}$ | $2.0\times10^{-2}$ | $1.2\times10^{1}$ | $1.3\times10^{-1}$ | $6.4\times10^{2}$ | $2.3\times10^{-1}$ |
| K207G | $1.4\times10^{2}$ | $4.5\times10^{-1}$ | $7.2\times10^{1}$ | $2.5\times10^{-1}$ | $2.0\times10^{6}$ | $1.4\times10^{-1}$ | $4.7\times10^{5}$ | $2.5\times10^{-1}$ | $1.5\times10^{1}$ | $8.4\times10^{-1}$ | $5.9\times10^{2}$ | $3.7\times10^{-1}$ |
| K207V | $1.1\times10^{2}$ | $6.3\times10^{-1}$ | $9.2\times10^{1}$ | $5.6\times10^{-1}$ | $1.3\times10^{6}$ | $8.3\times10^{-1}$ | $4.1\times10^{5}$ | $3.4\times10^{-2}$ | $\le 3.5$ | **$3.0\times10^{-4}$** | $6.4\times10^{2}$ | $4.3\times10^{-1}$ |
| N208G | $1.4\times10^{2}$ | $4.1\times10^{-1}$ | $6.7\times10^{1}$ | $1.6\times10^{-1}$ | $1.9\times10^{6}$ | $2.7\times10^{-1}$ | $3.9\times10^{5}$ | $6.2\times10^{-2}$ | $1.9\times10^{1}$ | $3.5\times10^{-1}$ | $5.5\times10^{2}$ | $3.3\times10^{-1}$ |
| N208V | $2.3\times10^{2}$ | **$2.0\times10^{-4}$** | $8.4\times10^{1}$ | $8.9\times10^{-1}$ | $2.5\times10^{6}$ | **$2.2\times10^{-3}$** | $2.7\times10^{5}$ | $1.7\times10^{-2}$ | $1.7\times10^{1}$ | $7.6\times10^{-1}$ | $6.3\times10^{2}$ | $6.0\times10^{-1}$ |
| V209A | $2.5\times10^{2}$ | **$4.0\times10^{-4}$** | $1.1\times10^{2}$ | $8.2\times10^{-2}$ | $2.3\times10^{6}$ | $1.6\times10^{-2}$ | $4.6\times10^{5}$ | $1.2\times10^{-1}$ | $1.7\times10^{1}$ | $8.8\times10^{-1}$ | $7.0\times10^{2}$ | $3.6\times10^{-2}$ |
| V209G | $8.4\times10^{1}$ | $2.3\times10^{-1}$ | $6.5\times10^{1}$ | $1.9\times10^{-1}$ | $1.6\times10^{6}$ | $7.5\times10^{-1}$ | $3.5\times10^{5}$ | $2.8\times10^{-2}$ | $1.5\times10^{1}$ | $6.2\times10^{-1}$ | $5.3\times10^{2}$ | **$8.6\times10^{-3}$** |
| P210G | $8.7\times10^{1}$ | $1.2\times10^{-1}$ | $7.3\times10^{1}$ | $2.4\times10^{-1}$ | $1.2\times10^{6}$ | $5.7\times10^{-1}$ | $3.9\times10^{5}$ | $7.4\times10^{-2}$ | $1.9\times10^{1}$ | $3.0\times10^{-1}$ | $5.6\times10^{2}$ | $1.2\times10^{-1}$ |
| P210V | $5.4\times10^{1}$ | **$9.0\times10^{-3}$** | $7.5\times10^{1}$ | $3.6\times10^{-1}$ | $7.3\times10^{5}$ | $6.8\times10^{-2}$ | $4.3\times10^{5}$ | $1.1\times10^{-1}$ | 7.2 | $1.6\times10^{-2}$ | $4.8\times10^{2}$ | **$1.0\times10^{-3}$** |
| A211G | $1.2\times10^{2}$ | $8.2\times10^{-1}$ | $5.9\times10^{1}$ | $7.9\times10^{-2}$ | $2.2\times10^{6}$ | $6.8\times10^{-2}$ | $6.4\times10^{5}$ | $8.5\times10^{-1}$ | $1.8\times10^{1}$ | $6.9\times10^{-1}$ | $6.1\times10^{2}$ | $7.6\times10^{-1}$ |
| A211V | $1.7\times10^{2}$ | $5.0\times10^{-2}$ | $9.3\times10^{1}$ | $5.2\times10^{-1}$ | $2.0\times10^{6}$ | $1.3\times10^{-1}$ | $4.5\times10^{5}$ | $1.2\times10^{-1}$ | $1.5\times10^{1}$ | $7.1\times10^{-1}$ | $6.2\times10^{2}$ | $6.9\times10^{-1}$ |
| Q212G | $1.6\times10^{2}$ | $1.1\times10^{-1}$ | $7.4\times10^{1}$ | $2.5\times10^{-1}$ | $2.0\times10^{6}$ | $8.3\times10^{-2}$ | $7.1\times10^{5}$ | $4.3\times10^{-1}$ | $1.8\times10^{1}$ | $5.8\times10^{-1}$ | $5.6\times10^{2}$ | $8.8\times10^{-2}$ |

| Variant | $k_{cat}$ cMUP ($s^{-1}$) | $k_{cat}$ cMUP *p*–value | $K_M$ cMUP (μM) | $K_M$ cMUP *p*–value | $k_{cat}/K_M$ cMUP ($M^{-1}s^{-1}$) | $k_{cat}/K_M$ cMUP *p*–value | $k_{cat}/K_M$ MeP ($M^{-1}s^{-1}$) | $k_{cat}/K_M$ MeP *p*–value | $k_{cat}/K_M$ MecMUP ($M^{-1}s^{-1}$) | $k_{cat}/K_M$ MecMUP *p*–value | $K_i$ $PO_4$ (μM) | $K_i$ $PO_4$ *p*–value |
|---|---|---|---|---|---|---|---|---|---|---|---|---|
| Q212V | $1.7\times10^{2}$ | $2.5\times10^{-2}$ | $8.6\times10^{1}$ | $9.8\times10^{-1}$ | $1.9\times10^{6}$ | $1.5\times10^{-1}$ | $4.2\times10^{5}$ | $7.1\times10^{-2}$ | $1.2\times10^{1}$ | $3.6\times10^{-1}$ | $6.4\times10^{2}$ | $4.4\times10^{-1}$ |
| L213G | $1.6\times10^{2}$ | $1.4\times10^{-1}$ | $6.7\times10^{1}$ | $1.7\times10^{-1}$ | $2.7\times10^{6}$ | **$5.1\times10^{-3}$** | $3.5\times10^{5}$ | $4.0\times10^{-2}$ | $1.6\times10^{1}$ | $9.2\times10^{-1}$ | $5.6\times10^{2}$ | $8.7\times10^{-2}$ |
| L213V | $1.3\times10^{2}$ | $5.2\times10^{-1}$ | $7.2\times10^{1}$ | $1.7\times10^{-1}$ | $1.5\times10^{6}$ | $6.5\times10^{-1}$ | $3.7\times10^{5}$ | $6.3\times10^{-2}$ | $1.4\times10^{1}$ | $5.6\times10^{-1}$ | $5.6\times10^{2}$ | $8.0\times10^{-2}$ |
| V214A | $1.4\times10^{2}$ | $1.8\times10^{-1}$ | $8.0\times10^{1}$ | $5.1\times10^{-1}$ | $1.6\times10^{6}$ | $6.1\times10^{-1}$ | $5.0\times10^{5}$ | $1.7\times10^{-1}$ | $1.4\times10^{1}$ | $4.3\times10^{-1}$ | $5.7\times10^{2}$ | $7.1\times10^{-2}$ |
| V214G | $7.2\times10^{1}$ | $4.4\times10^{-2}$ | $4.9\times10^{1}$ | **$4.9\times10^{-3}$** | $1.6\times10^{6}$ | $6.3\times10^{-1}$ | $2.2\times10^{5}$ | **$5.9\times10^{-3}$** | $1.6\times10^{1}$ | $9.7\times10^{-1}$ | $4.2\times10^{2}$ | **0** |
| A215G | $1.4\times10^{2}$ | $5.2\times10^{-1}$ | $6.4\times10^{1}$ | $1.2\times10^{-1}$ | $2.2\times10^{6}$ | $7.6\times10^{-2}$ | $5.2\times10^{5}$ | $5.7\times10^{-1}$ | $1.8\times10^{1}$ | $6.3\times10^{-1}$ | $5.5\times10^{2}$ | $1.1\times10^{-1}$ |
| A215V | $1.8\times10^{2}$ | $2.4\times10^{-2}$ | $8.8\times10^{1}$ | $7.8\times10^{-1}$ | $2.4\times10^{6}$ | $1.2\times10^{-2}$ | $5.1\times10^{5}$ | $4.1\times10^{-1}$ | $1.5\times10^{1}$ | $6.4\times10^{-1}$ | $6.1\times10^{2}$ | $7.3\times10^{-1}$ |
| N216G | $1.9\times10^{2}$ | $1.1\times10^{-2}$ | $9.4\times10^{1}$ | $4.6\times10^{-1}$ | $2.2\times10^{6}$ | $4.8\times10^{-2}$ | $6.0\times10^{5}$ | $9.2\times10^{-1}$ | $1.9\times10^{1}$ | $4.3\times10^{-1}$ | $5.4\times10^{2}$ | $2.3\times10^{-2}$ |
| N216V | $1.1\times10^{2}$ | $8.0\times10^{-1}$ | $9.1\times10^{1}$ | $7.6\times10^{-1}$ | $9.9\times10^{5}$ | $4.5\times10^{-1}$ | $1.2\times10^{5}$ | **0** | ≤ 3.9 | $1.1\times10^{-2}$ | – | – |
| G217A | $1.7\times10^{1}$ | **$1.\times10^{-4}$** | $7.1\times10^{1}$ | $1.7\times10^{-1}$ | $2.2\times10^{5}$ | **$6.0\times10^{-4}$** | $2.1\times10^{4}$ | **$4.0\times10^{-4}$** | 1.4 | **$6.0\times10^{-3}$** | $5.8\times10^{2}$ | $4.9\times10^{-1}$ |
| G217V | $1.7\times10^{2}$ | $2.0\times10^{-2}$ | $7.9\times10^{1}$ | $5.4\times10^{-1}$ | $1.9\times10^{6}$ | $1.9\times10^{-1}$ | $3.0\times10^{5}$ | $2.5\times10^{-2}$ | $2.0\times10^{1}$ | $4.0\times10^{-1}$ | $6.3\times10^{2}$ | $6.8\times10^{-1}$ |
| W218G | 7.5 | **0** | $3.2\times10^{1}$ | **0** | $2.2\times10^{5}$ | **$1.\times10^{-4}$** | $2.1\times10^{4}$ | **0** | 2.1 | **$1.1\times10^{-3}$** | $3.0\times10^{2}$ | **0** |
| W218V | $1.7\times10^{1}$ | **0** | $3.5\times10^{1}$ | **0** | $4.3\times10^{5}$ | **$8.0\times10^{-4}$** | $2.4\times10^{4}$ | **$1.\times10^{-4}$** | 2.6 | **$2.0\times10^{-4}$** | $2.6\times10^{2}$ | **0** |
| N219G | $9.2\times10^{1}$ | $1.4\times10^{-1}$ | $6.9\times10^{1}$ | $1.3\times10^{-1}$ | $1.4\times10^{6}$ | $8.8\times10^{-1}$ | $3.4\times10^{5}$ | $3.3\times10^{-2}$ | $1.7\times10^{1}$ | $7.1\times10^{-1}$ | $5.9\times10^{2}$ | $3.0\times10^{-1}$ |
| N219V | $1.5\times10^{2}$ | $1.1\times10^{-1}$ | $9.3\times10^{1}$ | $4.7\times10^{-1}$ | $1.4\times10^{6}$ | $8.1\times10^{-1}$ | $4.5\times10^{5}$ | $1.6\times10^{-1}$ | $1.7\times10^{1}$ | $8.7\times10^{-1}$ | $6.1\times10^{2}$ | $8.5\times10^{-1}$ |
| T220G | $5.4\times10^{1}$ | **$3.2\times10^{-3}$** | $7.0\times10^{1}$ | $1.6\times10^{-1}$ | $6.2\times10^{5}$ | $1.5\times10^{-2}$ | $2.4\times10^{5}$ | **$7.2\times10^{-3}$** | 5.4 | **$1.0\times10^{-3}$** | $4.9\times10^{2}$ | **$3.0\times10^{-4}$** |
| T220V | $1.3\times10^{2}$ | $7.8\times10^{-1}$ | $9.6\times10^{1}$ | $2.7\times10^{-1}$ | $1.4\times10^{6}$ | $8.6\times10^{-1}$ | $3.6\times10^{5}$ | $2.4\times10^{-2}$ | $1.3\times10^{1}$ | $4.2\times10^{-1}$ | $6.4\times10^{2}$ | $4.3\times10^{-1}$ |
| L221G | $8.3\times10^{1}$ | $1.9\times10^{-1}$ | $6.3\times10^{1}$ | $1.2\times10^{-1}$ | $1.4\times10^{6}$ | $9.3\times10^{-1}$ | $1.9\times10^{5}$ | $3.2\times10^{-2}$ | $1.1\times10^{1}$ | $1.3\times10^{-1}$ | $5.4\times10^{2}$ | $5.2\times10^{-2}$ |
| L221V | $1.5\times10^{2}$ | $3.5\times10^{-1}$ | $8.1\times10^{1}$ | $7.0\times10^{-1}$ | $1.6\times10^{6}$ | $6.8\times10^{-1}$ | $4.5\times10^{5}$ | $1.5\times10^{-1}$ | $1.9\times10^{1}$ | $5.5\times10^{-1}$ | $6.3\times10^{2}$ | $7.7\times10^{-1}$ |
| L222G | $7.7\times10^{1}$ | $6.7\times10^{-2}$ | $5.7\times10^{1}$ | $2.0\times10^{-2}$ | $1.2\times10^{6}$ | $4.7\times10^{-1}$ | $2.7\times10^{5}$ | $1.2\times10^{-2}$ | $1.2\times10^{1}$ | $2.1\times10^{-1}$ | $5.1\times10^{2}$ | $1.6\times10^{-2}$ |
| L222V | $1.1\times10^{2}$ | $4.7\times10^{-1}$ | $9.9\times10^{1}$ | $1.2\times10^{-1}$ | $1.1\times10^{6}$ | $3.3\times10^{-1}$ | $3.6\times10^{5}$ | **$7.2\times10^{-3}$** | 9.3 | $1.4\times10^{-2}$ | $6.5\times10^{2}$ | $1.4\times10^{-1}$ |
| P223G | $9.4\times10^{1}$ | $1.8\times10^{-1}$ | $6.6\times10^{1}$ | $6.4\times10^{-2}$ | $1.4\times10^{6}$ | $9.4\times10^{-1}$ | $3.7\times10^{5}$ | $5.5\times10^{-2}$ | $1.4\times10^{1}$ | $6.4\times10^{-1}$ | $5.0\times10^{2}$ | **$2.2\times10^{-3}$** |
| P223V | $1.2\times10^{2}$ | $7.9\times10^{-1}$ | $7.4\times10^{1}$ | $2.1\times10^{-1}$ | $1.6\times10^{6}$ | $6.8\times10^{-1}$ | $1.6\times10^{5}$ | **$6.0\times10^{-4}$** | $1.1\times10^{1}$ | $1.5\times10^{-1}$ | $6.0\times10^{2}$ | $5.7\times10^{-1}$ |
| I224G | $1.0\times10^{2}$ | $2.7\times10^{-1}$ | $7.6\times10^{1}$ | $4.2\times10^{-1}$ | $1.4\times10^{6}$ | $9.7\times10^{-1}$ | $5.2\times10^{5}$ | $4.5\times10^{-1}$ | $1.2\times10^{1}$ | $2.7\times10^{-1}$ | $5.4\times10^{2}$ | $4.8\times10^{-2}$ |
| I224V | $5.9\times10^{1}$ | $1.5\times10^{-2}$ | $9.7\times10^{1}$ | $2.9\times10^{-1}$ | $7.4\times10^{5}$ | $6.8\times10^{-2}$ | $2.9\times10^{5}$ | **$8.3\times10^{-3}$** | 7.4 | **$7.7\times10^{-3}$** | $7.2\times10^{2}$ | **$1.1\times10^{-3}$** |
| N225G | $2.0\times10^{2}$ | **$4.7\times10^{-3}$** | $6.8\times10^{1}$ | $1.9\times10^{-1}$ | $2.5\times10^{6}$ | $1.3\times10^{-2}$ | $4.0\times10^{5}$ | $1.3\times10^{-1}$ | $1.9\times10^{1}$ | $4.7\times10^{-1}$ | $6.1\times10^{2}$ | $7.9\times10^{-1}$ |
| N225V | $1.6\times10^{2}$ | $1.7\times10^{-2}$ | $1.1\times10^{2}$ | $1.6\times10^{-2}$ | $1.5\times10^{6}$ | $7.1\times10^{-1}$ | $5.4\times10^{5}$ | $5.8\times10^{-1}$ | $1.5\times10^{1}$ | $6.8\times10^{-1}$ | $6.9\times10^{2}$ | **$5.7\times10^{-3}$** |
| Q226G | $7.4\times10^{1}$ | $1.1\times10^{-1}$ | $4.9\times10^{1}$ | $1.4\times10^{-2}$ | $1.3\times10^{6}$ | $7.8\times10^{-1}$ | $2.8\times10^{5}$ | $5.7\times10^{-2}$ | $1.4\times10^{1}$ | $5.2\times10^{-1}$ | $4.4\times10^{2}$ | **$2.0\times10^{-4}$** |
| Q226V | $1.6\times10^{2}$ | $2.1\times10^{-2}$ | $9.3\times10^{1}$ | $3.1\times10^{-1}$ | $1.9\times10^{6}$ | $9.9\times10^{-2}$ | $4.8\times10^{5}$ | $1.4\times10^{-1}$ | $1.3\times10^{1}$ | $3.5\times10^{-1}$ | $6.5\times10^{2}$ | $1.5\times10^{-1}$ |
| Y227G | $2.5\times10^{1}$ | **0** | $4.0\times10^{1}$ | **$1.\times10^{-4}$** | $4.9\times10^{5}$ | **$4.5\times10^{-3}$** | $1.4\times10^{5}$ | **$8.0\times10^{-3}$** | 2.7 | **$5.0\times10^{-4}$** | $3.0\times10^{2}$ | **$2.0\times10^{-4}$** |
| Y227V | $4.5\times10^{1}$ | **$2.7\times10^{-3}$** | $7.3\times10^{1}$ | $2.7\times10^{-1}$ | $7.3\times10^{5}$ | $6.5\times10^{-2}$ | $1.9\times10^{5}$ | **$2.9\times10^{-3}$** | 5.6 | $5.4\times10^{-2}$ | $4.3\times10^{2}$ | **$4.0\times10^{-4}$** |
| T228G | $7.4\times10^{1}$ | $5.6\times10^{-2}$ | $6.9\times10^{1}$ | $1.5\times10^{-1}$ | $7.4\times10^{5}$ | $8.8\times10^{-2}$ | $1.2\times10^{5}$ | **0** | $1.3\times10^{1}$ | $4.4\times10^{-1}$ | $6.4\times10^{2}$ | $4.8\times10^{-1}$ |
| T228V | $1.1\times10^{2}$ | $3.8\times10^{-1}$ | $9.3\times10^{1}$ | $4.0\times10^{-1}$ | $1.1\times10^{6}$ | $4.4\times10^{-1}$ | $4.3\times10^{5}$ | $1.1\times10^{-1}$ | $1.0\times10^{1}$ | $1.7\times10^{-1}$ | $6.7\times10^{2}$ | $7.7\times10^{-2}$ |
| E229G | $4.9\times10^{1}$ | **$1.4\times10^{-3}$** | $5.1\times10^{1}$ | **$3.1\times10^{-3}$** | $8.9\times10^{5}$ | $1.0\times10^{-1}$ | $1.6\times10^{5}$ | **$4.0\times10^{-4}$** | 6.1 | **$1.\times10^{-4}$** | $5.3\times10^{2}$ | **$8.7\times10^{-3}$** |
| E229V | $5.8\times10^{1}$ | **$3.6\times10^{-3}$** | $7.7\times10^{1}$ | $3.5\times10^{-1}$ | $8.1\times10^{5}$ | $5.3\times10^{-2}$ | $1.4\times10^{5}$ | $1.5\times10^{-2}$ | 7.9 | $1.1\times10^{-2}$ | $5.8\times10^{2}$ | $2.7\times10^{-1}$ |
| S230G | $3.9\times10^{1}$ | **$3.0\times10^{-4}$** | $5.6\times10^{1}$ | **$7.9\times10^{-3}$** | $7.4\times10^{5}$ | $4.1\times10^{-2}$ | ≤ $2.9\times10^{5}$ | $4.6\times10^{-2}$ | 8.1 | $1.2\times10^{-2}$ | $4.9\times10^{2}$ | **$1.0\times10^{-3}$** |
| S230V | $3.3\times10^{1}$ | **0** | $5.8\times10^{1}$ | **$6.7\times10^{-3}$** | $5.3\times10^{5}$ | **$4.3\times10^{-3}$** | $2.1\times10^{5}$ | **$7.0\times10^{-4}$** | 5.0 | **$3.7\times10^{-3}$** | $3.5\times10^{2}$ | **$1.\times10^{-4}$** |
| S231G | $5.0\times10^{1}$ | $1.1\times10^{-2}$ | $6.0\times10^{1}$ | $7.4\times10^{-2}$ | $1.3\times10^{6}$ | $8.0\times10^{-1}$ | $3.9\times10^{5}$ | $6.8\times10^{-2}$ | 9.2 | $4.3\times10^{-2}$ | $4.9\times10^{2}$ | **$7.0\times10^{-4}$** |
| S231V | $7.2\times10^{1}$ | $2.4\times10^{-2}$ | $7.8\times10^{1}$ | $4.4\times10^{-1}$ | $1.0\times10^{6}$ | $2.0\times10^{-1}$ | $3.4\times10^{5}$ | $4.4\times10^{-2}$ | $1.1\times10^{1}$ | $1.8\times10^{-1}$ | $5.8\times10^{2}$ | $2.4\times10^{-1}$ |
| E232G | $1.1\times10^{2}$ | $6.0\times10^{-1}$ | $7.9\times10^{1}$ | $5.6\times10^{-1}$ | $1.2\times10^{6}$ | $5.9\times10^{-1}$ | $3.5\times10^{5}$ | $5.1\times10^{-2}$ | $1.5\times10^{1}$ | $7.9\times10^{-1}$ | $6.2\times10^{2}$ | $8.9\times10^{-1}$ |
| E232V | $7.7\times10^{1}$ | $5.5\times10^{-2}$ | $8.9\times10^{1}$ | $7.1\times10^{-1}$ | $9.2\times10^{5}$ | $1.6\times10^{-1}$ | $5.1\times10^{5}$ | $3.9\times10^{-1}$ | $1.1\times10^{1}$ | $2.1\times10^{-1}$ | $5.7\times10^{2}$ | $2.2\times10^{-1}$ |
| D233G | $1.1\times10^{1}$ | **0** | $2.5\times10^{1}$ | **0** | $4.5\times10^{5}$ | **$7.0\times10^{-4}$** | $2.9\times10^{4}$ | **$1.\times10^{-4}$** | 3.5 | **$1.\times10^{-4}$** | $1.6\times10^{2}$ | **0** |
| D233V | $2.9\times10^{1}$ | **0** | $3.6\times10^{1}$ | **0** | $8.2\times10^{5}$ | $5.0\times10^{-2}$ | $8.6\times10^{4}$ | **$1.4\times10^{-3}$** | 7.4 | $2.3\times10^{-2}$ | $2.1\times10^{2}$ | **0** |
| N234G | $8.9\times10^{1}$ | $1.7\times10^{-1}$ | $6.5\times10^{1}$ | $9.1\times10^{-2}$ | $1.4\times10^{6}$ | $8.8\times10^{-1}$ | $4.3\times10^{5}$ | $1.3\times10^{-1}$ | $1.5\times10^{1}$ | $6.9\times10^{-1}$ | $5.6\times10^{2}$ | $1.3\times10^{-1}$ |
| N234V | $1.1\times10^{2}$ | $4.4\times10^{-1}$ | $8.3\times10^{1}$ | $8.3\times10^{-1}$ | $1.3\times10^{6}$ | $8.2\times10^{-1}$ | $2.1\times10^{5}$ | **$3.8\times10^{-3}$** | $1.7\times10^{1}$ | $9.0\times10^{-1}$ | $6.1\times10^{2}$ | $6.7\times10^{-1}$ |
| V235A | $1.8\times10^{2}$ | $1.1\times10^{-2}$ | $9.5\times10^{1}$ | $3.2\times10^{-1}$ | $2.2\times10^{6}$ | $2.6\times10^{-2}$ | $4.2\times10^{5}$ | $1.5\times10^{-1}$ | $2.3\times10^{1}$ | $7.3\times10^{-2}$ | $6.2\times10^{2}$ | $7.3\times10^{-1}$ |
| V235G | $5.9\times10^{1}$ | **$6.4\times10^{-3}$** | $7.2\times10^{1}$ | $1.9\times10^{-1}$ | $6.2\times10^{5}$ | $1.7\times10^{-2}$ | $2.8\times10^{5}$ | $5.9\times10^{-2}$ | $1.3\times10^{1}$ | $3.1\times10^{-1}$ | $6.0\times10^{2}$ | $4.5\times10^{-1}$ |
| E236G | $7.9\times10^{1}$ | $7.9\times10^{-2}$ | $6.1\times10^{1}$ | $3.3\times10^{-2}$ | $1.1\times10^{6}$ | $4.1\times10^{-1}$ | $4.0\times10^{5}$ | $1.2\times10^{-1}$ | $1.3\times10^{1}$ | $2.7\times10^{-1}$ | $5.1\times10^{2}$ | **$8.9\times10^{-3}$** |
| E236V | $1.3\times10^{2}$ | $7.1\times10^{-1}$ | $9.4\times10^{1}$ | $3.8\times10^{-1}$ | $1.2\times10^{6}$ | $4.4\times10^{-1}$ | $5.8\times10^{5}$ | $7.1\times10^{-1}$ | $1.4\times10^{1}$ | $4.9\times10^{-1}$ | $6.4\times10^{2}$ | $6.2\times10^{-1}$ |
| W237G | $2.9\times10^{1}$ | **0** | $2.3\times10^{1}$ | **0** | $1.1\times10^{6}$ | $2.7\times10^{-1}$ | $7.3\times10^{4}$ | **0** | 6.4 | **$6.0\times10^{-4}$** | $1.8\times10^{2}$ | **0** |
| W237V | $1.0\times10^{1}$ | **0** | $4.1\times10^{1}$ | **0** | $5.0\times10^{5}$ | **$1.3\times10^{-3}$** | ≤ $2.7\times10^{4}$ | **0** | 2.4 | **$7.0\times10^{-4}$** | $2.2\times10^{2}$ | **0** |
| E238G | $1.1\times10^{1}$ | **0** | $2.7\times10^{1}$ | **0** | $4.1\times10^{5}$ | **$7.0\times10^{-4}$** | $4.8\times10^{4}$ | **0** | 2.4 | **$1.\times10^{-4}$** | $1.9\times10^{2}$ | **0** |
| E238V | $3.5\times10^{1}$ | **$2.4\times10^{-3}$** | $3.3\times10^{1}$ | **$7.0\times10^{-4}$** | $1.0\times10^{6}$ | $3.6\times10^{-1}$ | $7.1\times10^{4}$ | $1.8\times10^{-2}$ | 4.4 | **$6.0\times10^{-4}$** | $2.1\times10^{2}$ | **$2.0\times10^{-4}$** |
| G239A | $5.8\times10^{1}$ | **$5.2\times10^{-3}$** | $6.4\times10^{1}$ | $5.2\times10^{-2}$ | $9.3\times10^{5}$ | $1.4\times10^{-1}$ | $4.2\times10^{5}$ | $1.5\times10^{-1}$ | $1.9\times10^{1}$ | $2.9\times10^{-1}$ | $5.5\times10^{2}$ | $1.4\times10^{-1}$ |
| G239V | $9.6\times10^{1}$ | $1.8\times10^{-1}$ | $8.9\times10^{1}$ | $7.0\times10^{-1}$ | $1.1\times10^{6}$ | $2.7\times10^{-1}$ | $3.4\times10^{5}$ | $3.2\times10^{-2}$ | $1.0\times10^{1}$ | $1.1\times10^{-1}$ | $6.0\times10^{2}$ | $6.4\times10^{-1}$ |
| L240G | $1.2\times10^{2}$ | $8.9\times10^{-1}$ | $5.2\times10^{1}$ | $2.0\times10^{-2}$ | $2.4\times10^{6}$ | $2.9\times10^{-2}$ | $7.5\times10^{5}$ | $2.5\times10^{-1}$ | $1.8\times10^{1}$ | $6.4\times10^{-1}$ | $4.8\times10^{2}$ | **$6.0\times10^{-3}$** |
| L240V | $6.5\times10^{1}$ | $3.1\times10^{-2}$ | $9.4\times10^{1}$ | $4.3\times10^{-1}$ | $7.9\times10^{5}$ | $9.3\times10^{-2}$ | $3.7\times10^{5}$ | $1.5\times10^{-2}$ | 9.5 | $1.2\times10^{-1}$ | $6.9\times10^{2}$ | $1.3\times10^{-2}$ |
| L241G | $1.3\times10^{1}$ | **0** | $3.1\times10^{1}$ | **0** | $5.4\times10^{5}$ | **$3.0\times10^{-3}$** | $7.7\times10^{4}$ | **$1.\times10^{-4}$** | 3.9 | **0** | $2.1\times10^{2}$ | **0** |
| L241V | $9.7\times10^{1}$ | $1.6\times10^{-1}$ | $7.4\times10^{1}$ | $2.2\times10^{-1}$ | $1.4\times10^{6}$ | $9.4\times10^{-1}$ | $3.5\times10^{5}$ | $4.3\times10^{-2}$ | $1.2\times10^{1}$ | $4.0\times10^{-1}$ | $5.8\times10^{2}$ | $4.3\times10^{-1}$ |
| G242A | $7.2\times10^{1}$ | $5.3\times10^{-2}$ | $5.1\times10^{1}$ | **$5.9\times10^{-3}$** | $9.2\times10^{5}$ | $1.6\times10^{-1}$ | $3.7\times10^{5}$ | $6.1\times10^{-2}$ | 8.8 | $2.9\times10^{-2}$ | $4.2\times10^{2}$ | **$6.0\times10^{-4}$** |
| G242V | $6.9\times10^{1}$ | $1.0\times10^{-2}$ | $7.8\times10^{1}$ | $3.8\times10^{-1}$ | $1.0\times10^{6}$ | $2.1\times10^{-1}$ | $1.9\times10^{5}$ | **$1.9\times10^{-3}$** | 9.7 | $1.3\times10^{-1}$ | $4.9\times10^{2}$ | **$6.0\times10^{-4}$** |
| S243G | $7.7\times10^{1}$ | $4.8\times10^{-2}$ | $6.6\times10^{1}$ | $7.1\times10^{-2}$ | $1.3\times10^{6}$ | $7.4\times10^{-1}$ | $3.9\times10^{5}$ | $7.9\times10^{-2}$ | $1.1\times10^{1}$ | $9.7\times10^{-2}$ | $5.3\times10^{2}$ | **$8.2\times10^{-3}$** |
| S243V | $9.0\times10^{1}$ | $1.1\times10^{-1}$ | $8.0\times10^{1}$ | $6.4\times10^{-1}$ | $1.1\times10^{6}$ | $4.6\times10^{-1}$ | $3.5\times10^{5}$ | $2.4\times10^{-2}$ | 8.8 | $2.8\times10^{-2}$ | $5.7\times10^{2}$ | $1.5\times10^{-1}$ |
| K244G | $6.5\times10^{1}$ | $1.5\times10^{-2}$ | $6.7\times10^{1}$ | $9.4\times10^{-2}$ | $7.9\times10^{5}$ | $6.8\times10^{-2}$ | $5.2\times10^{5}$ | $5.4\times10^{-1}$ | $1.6\times10^{1}$ | $9.3\times10^{-1}$ | $5.4\times10^{2}$ | $3.4\times10^{-2}$ |
| K244V | $1.1\times10^{2}$ | $2.9\times10^{-1}$ | $7.6\times10^{1}$ | $3.8\times10^{-1}$ | $1.2\times10^{6}$ | $5.2\times10^{-1}$ | $3.5\times10^{5}$ | $1.4\times10^{-2}$ | $1.1\times10^{1}$ | $2.4\times10^{-1}$ | $6.1\times10^{2}$ | $6.5\times10^{-1}$ |
| K245G | $1.0\times10^{2}$ | $4.4\times10^{-1}$ | $7.6\times10^{1}$ | $2.9\times10^{-1}$ | $1.3\times10^{6}$ | $7.1\times10^{-1}$ | $4.0\times10^{5}$ | $9.1\times10^{-2}$ | $1.9\times10^{1}$ | $4.0\times10^{-1}$ | $6.2\times10^{2}$ | $7.7\times10^{-1}$ |
| K245V | $1.8\times10^{2}$ | $9.3\times10^{-2}$ | $6.8\times10^{1}$ | $2.8\times10^{-1}$ | $2.7\times10^{6}$ | $1.3\times10^{-2}$ | $5.4\times10^{5}$ | $5.4\times10^{-1}$ | $2.2\times10^{1}$ | $1.6\times10^{-1}$ | $6.1\times10^{2}$ | $6.5\times10^{-1}$ |
| T246G | $3.7\times10^{1}$ | $1.7\times10^{-2}$ | $8.0\times10^{1}$ | $7.0\times10^{-1}$ | $4.6\times10^{5}$ | $7.2\times10^{-2}$ | $2.0\times10^{5}$ | $1.5\times10^{-2}$ | ≤ 1.1 | **0** | $5.3\times10^{2}$ | $1.1\times10^{-1}$ |
| T246V | $1.6\times10^{2}$ | $1.1\times10^{-1}$ | $9.3\times10^{1}$ | $4.3\times10^{-1}$ | $1.6\times10^{6}$ | $5.6\times10^{-1}$ | $2.7\times10^{5}$ | **$3.9\times10^{-3}$** | $1.5\times10^{1}$ | $7.4\times10^{-1}$ | $6.8\times10^{2}$ | $1.5\times10^{-1}$ |
| P247G | $4.8\times10^{1}$ | **$1.1\times10^{-3}$** | $5.6\times10^{1}$ | **$9.3\times10^{-3}$** | $5.9\times10^{5}$ | $1.4\times10^{-2}$ | $2.1\times10^{5}$ | **$5.3\times10^{-3}$** | 8.3 | **$6.4\times10^{-3}$** | $5.6\times10^{2}$ | $5.1\times10^{-2}$ |
| P247V | $3.7\times10^{1}$ | **0** | $6.9\times10^{1}$ | $9.7\times10^{-2}$ | $5.4\times10^{5}$ | **$3.9\times10^{-3}$** | $2.9\times10^{5}$ | $1.6\times10^{-2}$ | 6.8 | $1.4\times10^{-2}$ | $4.5\times10^{2}$ | **$8.0\times10^{-4}$** |
| T248G | $2.3\times10^{1}$ | **$4.0\times10^{-4}$** | $3.9\times10^{1}$ | **$7.0\times10^{-4}$** | $4.4\times10^{5}$ | **$6.7\times10^{-3}$** | $2.7\times10^{5}$ | $1.3\times10^{-2}$ | 4.5 | **0** | $3.7\times10^{2}$ | **0** |
| T248V | $5.4\times10^{1}$ | $1.3\times10^{-2}$ | $8.2\times10^{1}$ | $7.4\times10^{-1}$ | $5.4\times10^{5}$ | $2.2\times10^{-2}$ | $6.5\times10^{5}$ | $8.0\times10^{-1}$ | 9.3 | $1.3\times10^{-1}$ | $5.5\times10^{2}$ | $1.2\times10^{-1}$ |
| F249G | 8.9 | **0** | $2.2\times10^{1}$ | **0** | $3.4\times10^{5}$ | **$1.\times10^{-4}$** | $3.1\times10^{4}$ | **0** | 2.2 | **0** | $1.3\times10^{2}$ | **0** |
| F249V | $3.8\times10^{1}$ | **$3.0\times10^{-4}$** | $7.6\times10^{1}$ | $3.5\times10^{-1}$ | $4.7\times10^{5}$ | **$5.3\times10^{-3}$** | $2.5\times10^{5}$ | **$3.2\times10^{-3}$** | 6.3 | **$9.2\times10^{-3}$** | $5.2\times10^{2}$ | **$3.9\times10^{-3}$** |

| Variant | $k_{cat}$ cMUP ($s^{-1}$) | $k_{cat}$ cMUP $p$–value | $K_M$ cMUP (μM) | $K_M$ cMUP $p$–value | $k_{cat}/K_M$ cMUP ($M^{-1}s^{-1}$) | $k_{cat}/K_M$ cMUP $p$–value | $k_{cat}/K_M$ MeP ($M^{-1}s^{-1}$) | $k_{cat}/K_M$ MeP $p$–value | $k_{cat}/K_M$ MecMUP ($M^{-1}s^{-1}$) | $k_{cat}/K_M$ MecMUP $p$–value | $K_i$ $PO_4$ (μM) | $K_i$ $PO_4$ $p$–value |
|---|---|---|---|---|---|---|---|---|---|---|---|---|
| P250G | $3.8\times10^{1}$ | **$1.4\times10^{-3}$** | $5.7\times10^{1}$ | $2.4\times10^{-2}$ | $7.5\times10^{5}$ | $7.7\times10^{-2}$ | $3.9\times10^{4}$ | **0** | 5.5 | **$9.0\times10^{-4}$** | $5.1\times10^{2}$ | $1.8\times10^{-2}$ |
| P250V | $6.0\times10^{1}$ | $2.6\times10^{-2}$ | $8.8\times10^{1}$ | $8.2\times10^{-1}$ | $7.5\times10^{5}$ | $6.2\times10^{-2}$ | $4.2\times10^{5}$ | $9.0\times10^{-2}$ | 7.5 | $1.4\times10^{-2}$ | $5.7\times10^{2}$ | $1.7\times10^{-1}$ |
| Y251G | 7.4 | **0** | $2.3\times10^{1}$ | **0** | $2.8\times10^{5}$ | **0** | $2.5\times10^{4}$ | **0** | 2.7 | **$2.0\times10^{-4}$** | $1.7\times10^{2}$ | **0** |
| Y251V | $4.1\times10^{1}$ | **$1.9\times10^{-3}$** | $7.1\times10^{1}$ | $2.2\times10^{-1}$ | $5.6\times10^{5}$ | $2.2\times10^{-2}$ | $2.6\times10^{5}$ | $1.0\times10^{-2}$ | 5.1 | **$4.8\times10^{-3}$** | $5.1\times10^{2}$ | **$9.6\times10^{-3}$** |
| T252G | $1.7\times10^{2}$ | $4.5\times10^{-2}$ | $8.4\times10^{1}$ | $9.4\times10^{-1}$ | $2.0\times10^{6}$ | $1.3\times10^{-1}$ | $5.5\times10^{5}$ | $6.5\times10^{-1}$ | $1.4\times10^{1}$ | $6.2\times10^{-1}$ | $5.9\times10^{2}$ | $4.9\times10^{-1}$ |
| T252V | $1.5\times10^{2}$ | $1.5\times10^{-1}$ | $9.5\times10^{1}$ | $3.7\times10^{-1}$ | $1.5\times10^{6}$ | $8.7\times10^{-1}$ | $4.4\times10^{5}$ | $1.2\times10^{-1}$ | $1.0\times10^{1}$ | $1.4\times10^{-1}$ | $6.7\times10^{2}$ | $3.4\times10^{-1}$ |
| D253G | $8.8\times10^{1}$ | $7.2\times10^{-2}$ | $7.4\times10^{1}$ | $2.6\times10^{-1}$ | $1.2\times10^{6}$ | $5.2\times10^{-1}$ | $3.7\times10^{5}$ | $5.6\times10^{-2}$ | $1.6\times10^{1}$ | $9.6\times10^{-1}$ | $5.3\times10^{2}$ | $1.0\times10^{-2}$ |
| D253V | $8.0\times10^{1}$ | $4.6\times10^{-2}$ | $9.1\times10^{1}$ | $6.0\times10^{-1}$ | $1.3\times10^{6}$ | $7.0\times10^{-1}$ | $4.7\times10^{5}$ | $2.4\times10^{-1}$ | $1.1\times10^{1}$ | $2.3\times10^{-1}$ | $6.0\times10^{2}$ | $5.3\times10^{-1}$ |
| L254G | $5.2\times10^{1}$ | $1.6\times10^{-2}$ | $5.5\times10^{1}$ | $2.7\times10^{-2}$ | $1.4\times10^{6}$ | $9.7\times10^{-1}$ | $3.8\times10^{5}$ | $7.1\times10^{-2}$ | 8.7 | $3.1\times10^{-2}$ | $4.3\times10^{2}$ | **$1.\times10^{-4}$** |
| L254V | $1.3\times10^{2}$ | $8.5\times10^{-1}$ | $8.6\times10^{1}$ | $9.3\times10^{-1}$ | $1.6\times10^{6}$ | $4.7\times10^{-1}$ | $8.6\times10^{5}$ | $1.0\times10^{-1}$ | $1.4\times10^{1}$ | $6.2\times10^{-1}$ | $6.1\times10^{2}$ | $7.7\times10^{-1}$ |
| A255G | $9.2\times10^{1}$ | $1.3\times10^{-1}$ | $6.4\times10^{1}$ | $4.4\times10^{-2}$ | $1.5\times10^{6}$ | $8.5\times10^{-1}$ | $2.4\times10^{5}$ | $2.1\times10^{-2}$ | $2.1\times10^{1}$ | $1.9\times10^{-1}$ | $5.2\times10^{2}$ | **$5.6\times10^{-3}$** |
| A255V | $9.9\times10^{1}$ | $1.9\times10^{-1}$ | $7.7\times10^{1}$ | $2.9\times10^{-1}$ | $1.3\times10^{6}$ | $5.1\times10^{-1}$ | $4.4\times10^{5}$ | $4.5\times10^{-2}$ | $1.2\times10^{1}$ | $1.6\times10^{-1}$ | $5.5\times10^{2}$ | $1.4\times10^{-2}$ |
| K256G | $1.0\times10^{2}$ | $3.8\times10^{-1}$ | $6.6\times10^{1}$ | $1.1\times10^{-1}$ | $1.6\times10^{6}$ | $7.0\times10^{-1}$ | $4.3\times10^{5}$ | $2.2\times10^{-1}$ | $1.9\times10^{1}$ | $3.6\times10^{-1}$ | $5.3\times10^{2}$ | $1.7\times10^{-2}$ |
| K256V | $4.0\times10^{1}$ | **$8.6\times10^{-3}$** | $1.0\times10^{2}$ | $2.1\times10^{-1}$ | $4.2\times10^{5}$ | $2.0\times10^{-2}$ | $1.1\times10^{5}$ | **0** | ≤ 1.9 | **$3.0\times10^{-4}$** | $6.5\times10^{2}$ | $4.7\times10^{-1}$ |
| D257G | $6.2\times10^{1}$ | $4.1\times10^{-2}$ | $3.2\times10^{1}$ | **$6.0\times10^{-4}$** | $2.3\times10^{6}$ | $4.4\times10^{-2}$ | $3.4\times10^{5}$ | $8.7\times10^{-2}$ | $1.9\times10^{1}$ | $4.3\times10^{-1}$ | $4.2\times10^{2}$ | **$1.9\times10^{-3}$** |
| D257V | $1.2\times10^{2}$ | $9.6\times10^{-1}$ | $7.5\times10^{1}$ | $3.4\times10^{-1}$ | $1.7\times10^{6}$ | $3.5\times10^{-1}$ | $1.5\times10^{5}$ | $2.3\times10^{-2}$ | $1.4\times10^{1}$ | $6.8\times10^{-1}$ | $5.1\times10^{2}$ | **$7.3\times10^{-3}$** |
| Y258G | $1.3\times10^{2}$ | $6.2\times10^{-1}$ | $7.0\times10^{1}$ | $2.1\times10^{-1}$ | $1.9\times10^{6}$ | $2.0\times10^{-1}$ | $3.9\times10^{5}$ | $7.6\times10^{-2}$ | $1.4\times10^{1}$ | $5.3\times10^{-1}$ | $5.0\times10^{2}$ | **$6.0\times10^{-3}$** |
| Y258V | $1.5\times10^{2}$ | $1.2\times10^{-1}$ | $7.4\times10^{1}$ | $3.2\times10^{-1}$ | $1.9\times10^{6}$ | $2.0\times10^{-1}$ | $8.7\times10^{5}$ | $4.0\times10^{-2}$ | $1.8\times10^{1}$ | $6.4\times10^{-1}$ | $5.1\times10^{2}$ | $1.1\times10^{-2}$ |
| E259G | $1.3\times10^{2}$ | $6.9\times10^{-1}$ | $6.7\times10^{1}$ | $1.0\times10^{-1}$ | $1.8\times10^{6}$ | $2.1\times10^{-1}$ | $2.2\times10^{5}$ | **$6.0\times10^{-3}$** | $2.2\times10^{1}$ | $4.0\times10^{-2}$ | $4.4\times10^{2}$ | **0** |
| E259V | $1.7\times10^{2}$ | $5.2\times10^{-2}$ | $9.6\times10^{1}$ | $3.3\times10^{-1}$ | $1.7\times10^{6}$ | $5.5\times10^{-1}$ | $5.0\times10^{5}$ | $3.4\times10^{-1}$ | $1.8\times10^{1}$ | $6.7\times10^{-1}$ | $5.8\times10^{2}$ | $2.7\times10^{-1}$ |
| A260G | $8.5\times10^{1}$ | $2.0\times10^{-1}$ | $5.8\times10^{1}$ | $5.4\times10^{-2}$ | $2.0\times10^{6}$ | $1.5\times10^{-1}$ | $4.2\times10^{5}$ | $1.2\times10^{-1}$ | $2.1\times10^{1}$ | $1.2\times10^{-1}$ | $5.7\times10^{2}$ | $1.8\times10^{-1}$ |
| A260V | $1.7\times10^{2}$ | $1.4\times10^{-2}$ | $9.4\times10^{1}$ | $3.4\times10^{-1}$ | $2.0\times10^{6}$ | $8.2\times10^{-2}$ | $4.7\times10^{5}$ | $1.8\times10^{-1}$ | $1.4\times10^{1}$ | $6.5\times10^{-1}$ | $6.7\times10^{2}$ | $9.2\times10^{-2}$ |
| K261G | $6.8\times10^{1}$ | $8.0\times10^{-2}$ | $1.0\times10^{2}$ | $2.7\times10^{-1}$ | $8.8\times10^{5}$ | $2.1\times10^{-1}$ | $6.2\times10^{5}$ | $9.5\times10^{-1}$ | $1.7\times10^{1}$ | $7.5\times10^{-1}$ | $6.3\times10^{2}$ | $5.1\times10^{-1}$ |
| K261V | $1.0\times10^{2}$ | $4.5\times10^{-1}$ | $9.4\times10^{1}$ | $4.8\times10^{-1}$ | $1.4\times10^{6}$ | $8.6\times10^{-1}$ | $4.3\times10^{5}$ | $9.8\times10^{-2}$ | $1.1\times10^{1}$ | $1.6\times10^{-1}$ | $6.2\times10^{2}$ | $7.9\times10^{-1}$ |
| K262G | $1.0\times10^{2}$ | $3.7\times10^{-1}$ | $7.6\times10^{1}$ | $3.7\times10^{-1}$ | $1.6\times10^{6}$ | $6.5\times10^{-1}$ | $4.5\times10^{5}$ | $2.1\times10^{-1}$ | $2.5\times10^{1}$ | **$1.9\times10^{-3}$** | $4.6\times10^{2}$ | **0** |
| K262V | $1.0\times10^{2}$ | $3.1\times10^{-1}$ | $7.1\times10^{1}$ | $1.2\times10^{-1}$ | $1.3\times10^{6}$ | $7.2\times10^{-1}$ | $4.3\times10^{5}$ | $1.1\times10^{-1}$ | $1.8\times10^{1}$ | $6.1\times10^{-1}$ | $5.3\times10^{2}$ | $1.6\times10^{-2}$ |
| G263A | $8.5\times10^{1}$ | $7.9\times10^{-2}$ | $7.0\times10^{1}$ | $1.5\times10^{-1}$ | $1.1\times10^{6}$ | $3.0\times10^{-1}$ | $4.0\times10^{5}$ | $1.8\times10^{-1}$ | $2.0\times10^{1}$ | $2.0\times10^{-1}$ | $5.6\times10^{2}$ | $7.9\times10^{-2}$ |
| G263V | $9.7\times10^{1}$ | $1.4\times10^{-1}$ | $8.2\times10^{1}$ | $6.6\times10^{-1}$ | $1.1\times10^{6}$ | $2.1\times10^{-1}$ | $6.3\times10^{5}$ | $9.5\times10^{-1}$ | $1.6\times10^{1}$ | $9.8\times10^{-1}$ | $5.3\times10^{2}$ | $2.4\times10^{-2}$ |
| L264G | $6.5\times10^{1}$ | $1.3\times10^{-2}$ | $6.3\times10^{1}$ | $4.2\times10^{-2}$ | $1.0\times10^{6}$ | $2.1\times10^{-1}$ | $3.3\times10^{5}$ | $4.3\times10^{-2}$ | $1.1\times10^{1}$ | $1.2\times10^{-1}$ | $4.0\times10^{2}$ | **0** |
| L264V | $6.0\times10^{1}$ | $3.6\times10^{-2}$ | $1.0\times10^{2}$ | $1.5\times10^{-1}$ | $5.8\times10^{5}$ | $5.0\times10^{-2}$ | $5.3\times10^{5}$ | $5.4\times10^{-1}$ | ≤ 1.9 | **$1.7\times10^{-3}$** | $6.9\times10^{2}$ | $5.1\times10^{-2}$ |
| I265G | $2.9\times10^{1}$ | **0** | $4.2\times10^{1}$ | **$2.0\times10^{-4}$** | $7.4\times10^{5}$ | $2.6\times10^{-2}$ | $7.2\times10^{4}$ | **$5.0\times10^{-4}$** | 6.6 | **$9.0\times10^{-4}$** | $3.0\times10^{2}$ | **0** |
| I265V | $5.8\times10^{1}$ | $2.9\times10^{-2}$ | $8.8\times10^{1}$ | $8.3\times10^{-1}$ | $9.3\times10^{5}$ | $2.4\times10^{-1}$ | $7.0\times10^{5}$ | $5.4\times10^{-1}$ | 8.0 | $2.9\times10^{-2}$ | $5.5\times10^{2}$ | $5.1\times10^{-2}$ |
| R266G | $1.3\times10^{2}$ | $7.8\times10^{-1}$ | $6.1\times10^{1}$ | $3.3\times10^{-2}$ | $2.3\times10^{6}$ | $1.2\times10^{-2}$ | ≤ $1.4\times10^{5}$ | **$2.0\times10^{-3}$** | $2.8\times10^{1}$ | **$1.\times10^{-4}$** | $6.5\times10^{2}$ | $2.6\times10^{-1}$ |
| R266V | $2.2\times10^{2}$ | **$1.\times10^{-4}$** | $1.2\times10^{2}$ | **$4.0\times10^{-4}$** | $1.6\times10^{6}$ | $4.8\times10^{-1}$ | $8.9\times10^{4}$ | **0** | $4.4\times10^{1}$ | **0** | $1.7\times10^{3}$ | **0** |
| T267G | $1.0\times10^{2}$ | $4.3\times10^{-1}$ | $5.1\times10^{1}$ | **$5.5\times10^{-3}$** | $2.6\times10^{6}$ | **$1.3\times10^{-3}$** | $5.6\times10^{5}$ | $6.6\times10^{-1}$ | $2.4\times10^{1}$ | $1.2\times10^{-2}$ | $4.5\times10^{2}$ | **$3.0\times10^{-4}$** |
| T267V | $1.8\times10^{2}$ | **$3.8\times10^{-3}$** | $8.3\times10^{1}$ | $7.2\times10^{-1}$ | $2.1\times10^{6}$ | $2.9\times10^{-2}$ | $4.7\times10^{5}$ | $1.6\times10^{-1}$ | $1.2\times10^{1}$ | $3.2\times10^{-1}$ | $6.0\times10^{2}$ | $6.4\times10^{-1}$ |
| T268V | $1.4\times10^{2}$ | $1.9\times10^{-1}$ | $7.0\times10^{1}$ | $7.9\times10^{-2}$ | $2.1\times10^{6}$ | $2.2\times10^{-2}$ | $4.6\times10^{5}$ | $5.6\times10^{-2}$ | $1.4\times10^{1}$ | $4.5\times10^{-1}$ | $5.4\times10^{2}$ | **$6.0\times10^{-3}$** |
| P269G | $1.2\times10^{2}$ | $8.9\times10^{-1}$ | $9.1\times10^{1}$ | $5.7\times10^{-1}$ | $1.4\times10^{6}$ | $9.7\times10^{-1}$ | $2.2\times10^{5}$ | **$7.1\times10^{-3}$** | 6.6 | $1.3\times10^{-2}$ | $7.3\times10^{2}$ | **$4.0\times10^{-3}$** |
| P269V | $1.7\times10^{2}$ | $2.1\times10^{-2}$ | $8.1\times10^{1}$ | $6.3\times10^{-1}$ | $2.3\times10^{6}$ | $1.1\times10^{-2}$ | $2.4\times10^{5}$ | **$7.5\times10^{-3}$** | 9.9 | $6.3\times10^{-2}$ | $5.7\times10^{2}$ | $1.2\times10^{-1}$ |
| F270G | $5.2\times10^{1}$ | $1.6\times10^{-2}$ | $5.1\times10^{1}$ | $1.8\times10^{-2}$ | $1.4\times10^{6}$ | $9.5\times10^{-1}$ | $3.7\times10^{5}$ | $5.4\times10^{-2}$ | $1.4\times10^{1}$ | $4.4\times10^{-1}$ | $4.8\times10^{2}$ | **$2.0\times10^{-4}$** |
| F270V | $2.1\times10^{2}$ | **$2.0\times10^{-4}$** | $8.8\times10^{1}$ | $8.0\times10^{-1}$ | $2.3\times10^{6}$ | **$3.2\times10^{-3}$** | $3.0\times10^{5}$ | **$6.6\times10^{-3}$** | $1.4\times10^{1}$ | $6.7\times10^{-1}$ | $6.0\times10^{2}$ | $6.5\times10^{-1}$ |
| G271A | $8.6\times10^{1}$ | $1.2\times10^{-1}$ | $5.4\times10^{1}$ | $1.3\times10^{-2}$ | $2.0\times10^{6}$ | $1.5\times10^{-1}$ | $3.0\times10^{5}$ | $2.1\times10^{-2}$ | 9.4 | $2.2\times10^{-2}$ | $5.0\times10^{2}$ | **$5.0\times10^{-4}$** |
| G271V | $3.8\times10^{1}$ | **$1.\times10^{-4}$** | $4.2\times10^{1}$ | **$4.0\times10^{-4}$** | $1.2\times10^{6}$ | $5.4\times10^{-1}$ | $9.1\times10^{4}$ | **$9.0\times10^{-4}$** | $2.5\times10^{1}$ | $1.2\times10^{-2}$ | $4.1\times10^{2}$ | **$5.0\times10^{-4}$** |
| N272G | $7.9\times10^{1}$ | $1.3\times10^{-1}$ | $6.9\times10^{1}$ | $1.3\times10^{-1}$ | $1.1\times10^{6}$ | $3.7\times10^{-1}$ | $1.1\times10^{5}$ | **$1.1\times10^{-3}$** | 8.2 | $1.6\times10^{-2}$ | $6.6\times10^{2}$ | $1.5\times10^{-1}$ |
| N272V | $4.1\times10^{1}$ | $1.0\times10^{-2}$ | $8.1\times10^{1}$ | $7.7\times10^{-1}$ | $4.5\times10^{5}$ | $2.3\times10^{-2}$ | $2.0\times10^{5}$ | **$4.0\times10^{-3}$** | ≤ 1.8 | **0** | $4.6\times10^{2}$ | **0** |
| T273G | $5.5\times10^{1}$ | **$4.8\times10^{-3}$** | $6.4\times10^{1}$ | $5.2\times10^{-2}$ | $1.1\times10^{6}$ | $3.1\times10^{-1}$ | $1.6\times10^{5}$ | $1.2\times10^{-2}$ | $1.3\times10^{1}$ | $3.6\times10^{-1}$ | $5.3\times10^{2}$ | **$3.0\times10^{-3}$** |
| T273V | $2.4\times10^{2}$ | **0** | $9.2\times10^{1}$ | $4.7\times10^{-1}$ | $2.6\times10^{6}$ | **$1.\times10^{-4}$** | $1.7\times10^{5}$ | **$3.0\times10^{-4}$** | $1.5\times10^{1}$ | $7.4\times10^{-1}$ | $7.3\times10^{2}$ | **$2.4\times10^{-3}$** |
| L274G | $1.2\times10^{1}$ | **$2.0\times10^{-3}$** | $7.4\times10^{1}$ | $3.9\times10^{-1}$ | $1.7\times10^{5}$ | **$5.4\times10^{-3}$** | $2.2\times10^{4}$ | **$4.0\times10^{-4}$** | ≤ $9.9\times10^{-1}$ | **$1.\times10^{-4}$** | $6.0\times10^{2}$ | $7.5\times10^{-1}$ |
| L274V | $1.6\times10^{2}$ | $7.9\times10^{-2}$ | $1.1\times10^{2}$ | $9.1\times10^{-2}$ | $1.4\times10^{6}$ | $9.9\times10^{-1}$ | $4.7\times10^{5}$ | $1.6\times10^{-1}$ | $1.1\times10^{1}$ | $2.4\times10^{-1}$ | $6.5\times10^{2}$ | $3.1\times10^{-1}$ |
| T275G | $5.3\times10^{1}$ | **$1.3\times10^{-3}$** | $6.4\times10^{1}$ | $2.6\times10^{-2}$ | $8.3\times10^{5}$ | $5.6\times10^{-2}$ | $1.0\times10^{5}$ | **0** | 6.8 | **$4.0\times10^{-4}$** | $6.2\times10^{2}$ | $7.5\times10^{-1}$ |
| T275V | $1.2\times10^{2}$ | $9.0\times10^{-1}$ | $7.4\times10^{1}$ | $3.0\times10^{-1}$ | $1.8\times10^{6}$ | $3.4\times10^{-1}$ | $2.9\times10^{5}$ | **$5.3\times10^{-3}$** | $1.7\times10^{1}$ | $8.9\times10^{-1}$ | $6.1\times10^{2}$ | $8.7\times10^{-1}$ |
| L276G | 2.7 | **$3.0\times10^{-4}$** | $7.9\times10^{1}$ | $6.2\times10^{-1}$ | $3.3\times10^{4}$ | **$1.4\times10^{-3}$** | $6.0\times10^{3}$ | **$1.0\times10^{-3}$** | ≤ 1.1 | **0** | $3.8\times10^{2}$ | $2.2\times10^{-2}$ |
| L276V | $1.4\times10^{2}$ | $4.2\times10^{-1}$ | $9.1\times10^{1}$ | $4.6\times10^{-1}$ | $1.4\times10^{6}$ | $9.0\times10^{-1}$ | $2.7\times10^{5}$ | $1.4\times10^{-2}$ | $1.2\times10^{1}$ | $2.4\times10^{-1}$ | $6.1\times10^{2}$ | $8.1\times10^{-1}$ |
| Q277G | $1.5\times10^{2}$ | $1.6\times10^{-1}$ | $8.5\times10^{1}$ | $9.9\times10^{-1}$ | $1.7\times10^{6}$ | $4.9\times10^{-1}$ | $1.7\times10^{5}$ | **$3.5\times10^{-3}$** | $1.6\times10^{1}$ | $9.9\times10^{-1}$ | $6.1\times10^{2}$ | $6.0\times10^{-1}$ |
| Q277V | $1.4\times10^{2}$ | $3.4\times10^{-1}$ | $9.4\times10^{1}$ | $4.4\times10^{-1}$ | $1.4\times10^{6}$ | $8.2\times10^{-1}$ | $3.5\times10^{5}$ | $5.5\times10^{-2}$ | $1.1\times10^{1}$ | $2.8\times10^{-1}$ | $6.4\times10^{2}$ | $4.3\times10^{-1}$ |
| M278G | $1.5\times10^{1}$ | **$1.\times10^{-4}$** | $5.8\times10^{1}$ | $4.5\times10^{-2}$ | $3.3\times10^{5}$ | $1.0\times10^{-2}$ | $6.4\times10^{4}$ | **$1.\times10^{-4}$** | ≤ 1.1 | **0** | $4.8\times10^{2}$ | **$2.1\times10^{-3}$** |
| M278V | $1.0\times10^{2}$ | $3.6\times10^{-1}$ | $9.4\times10^{1}$ | $4.9\times10^{-1}$ | $1.2\times10^{6}$ | $4.6\times10^{-1}$ | $3.3\times10^{5}$ | $1.7\times10^{-2}$ | $1.0\times10^{1}$ | $2.0\times10^{-1}$ | $6.6\times10^{2}$ | $1.4\times10^{-1}$ |
| A279G | $6.5\times10^{1}$ | $5.9\times10^{-2}$ | $5.9\times10^{1}$ | $7.2\times10^{-2}$ | $9.0\times10^{5}$ | $2.3\times10^{-1}$ | $1.7\times10^{5}$ | **$3.9\times10^{-3}$** | 8.2 | **$7.5\times10^{-3}$** | $4.7\times10^{2}$ | **$1.1\times10^{-3}$** |
| A279V | $1.3\times10^{2}$ | $7.6\times10^{-1}$ | $9.0\times10^{1}$ | $5.8\times10^{-1}$ | $1.4\times10^{6}$ | $9.0\times10^{-1}$ | $2.9\times10^{5}$ | **$6.1\times10^{-3}$** | $1.1\times10^{1}$ | $1.7\times10^{-1}$ | $6.1\times10^{2}$ | $8.8\times10^{-1}$ |
| D280G | $1.2\times10^{2}$ | $9.5\times10^{-1}$ | $8.4\times10^{1}$ | $9.5\times10^{-1}$ | $1.9\times10^{6}$ | $2.1\times10^{-1}$ | $6.1\times10^{5}$ | $9.8\times10^{-1}$ | $1.8\times10^{1}$ | $5.2\times10^{-1}$ | $7.2\times10^{2}$ | $4.2\times10^{-2}$ |
| D280V | $6.2\times10^{1}$ | $1.6\times10^{-2}$ | $8.3\times10^{1}$ | $8.3\times10^{-1}$ | $6.7\times10^{5}$ | $3.2\times10^{-2}$ | $4.1\times10^{5}$ | $2.0\times10^{-2}$ | 6.9 | $1.8\times10^{-2}$ | $5.7\times10^{2}$ | $1.1\times10^{-1}$ |
| A281G | $1.7\times10^{2}$ | $6.4\times10^{-2}$ | $9.0\times10^{1}$ | $6.7\times10^{-1}$ | $1.7\times10^{6}$ | $4.7\times10^{-1}$ | $2.3\times10^{5}$ | **$6.7\times10^{-3}$** | 9.3 | $2.5\times10^{-2}$ | $6.3\times10^{2}$ | $4.8\times10^{-1}$ |
| A281V | $7.7\times10^{1}$ | $1.1\times10^{-1}$ | $8.1\times10^{1}$ | $6.7\times10^{-1}$ | $9.1\times10^{5}$ | $1.5\times10^{-1}$ | $2.1\times10^{5}$ | **$5.3\times10^{-3}$** | $1.0\times10^{1}$ | $1.8\times10^{-1}$ | $6.1\times10^{2}$ | $8.2\times10^{-1}$ |
| A282G | $4.5\times10^{1}$ | $2.5\times10^{-2}$ | $6.2\times10^{1}$ | $1.8\times10^{-1}$ | $8.6\times10^{5}$ | $2.9\times10^{-1}$ | $1.7\times10^{5}$ | **$9.8\times10^{-3}$** | 7.8 | **$3.9\times10^{-3}$** | $5.4\times10^{2}$ | $6.8\times10^{-2}$ |
| A282V | $7.3\times10^{1}$ | $1.7\times10^{-2}$ | $7.7\times10^{1}$ | $3.9\times10^{-1}$ | $8.6\times10^{5}$ | $5.5\times10^{-2}$ | $4.2\times10^{5}$ | $2.3\times10^{-2}$ | 8.4 | $2.0\times10^{-2}$ | $5.3\times10^{2}$ | **$1.7\times10^{-3}$** |
| I283G | 3.4 | **0** | $7.2\times10^{1}$ | $2.5\times10^{-1}$ | $4.9\times10^{4}$ | **$3.0\times10^{-4}$** | $5.6\times10^{3}$ | **$3.4\times10^{-3}$** | ≤ 1.1 | **0** | $3.9\times10^{2}$ | **0** |
| I283V | $1.4\times10^{2}$ | $3.4\times10^{-1}$ | $7.7\times10^{1}$ | $4.0\times10^{-1}$ | $1.4\times10^{6}$ | $9.4\times10^{-1}$ | $5.3\times10^{5}$ | $6.8\times10^{-1}$ | $1.5\times10^{1}$ | $6.8\times10^{-1}$ | $5.8\times10^{2}$ | $2.3\times10^{-1}$ |
| D284G | $9.2\times10^{1}$ | $4.4\times10^{-2}$ | $7.4\times10^{1}$ | $1.6\times10^{-1}$ | $1.7\times10^{6}$ | $3.4\times10^{-1}$ | $3.0\times10^{5}$ | **$2.9\times10^{-3}$** | $1.8\times10^{1}$ | $4.8\times10^{-1}$ | $5.8\times10^{2}$ | $1.6\times10^{-1}$ |
| D284V | $2.1\times10^{2}$ | **$2.0\times10^{-4}$** | $1.0\times10^{2}$ | $6.4\times10^{-2}$ | $2.0\times10^{6}$ | $5.0\times10^{-2}$ | $4.5\times10^{5}$ | $1.2\times10^{-1}$ | $1.7\times10^{1}$ | $7.3\times10^{-1}$ | $6.5\times10^{2}$ | $1.1\times10^{-1}$ |
| G285A | $1.7\times10^{2}$ | $6.3\times10^{-2}$ | $8.0\times10^{1}$ | $6.4\times10^{-1}$ | $2.4\times10^{6}$ | **$8.1\times10^{-3}$** | $5.8\times10^{5}$ | $8.2\times10^{-1}$ | $1.7\times10^{1}$ | $9.1\times10^{-1}$ | $6.3\times10^{2}$ | $6.4\times10^{-1}$ |
| G285V | $6.0\times10^{1}$ | **$4.1\times10^{-3}$** | $9.6\times10^{1}$ | $2.4\times10^{-1}$ | $6.4\times10^{5}$ | $1.2\times10^{-2}$ | $1.1\times10^{5}$ | **$1.\times10^{-4}$** | 5.0 | **$2.0\times10^{-4}$** | $6.1\times10^{2}$ | $7.3\times10^{-1}$ |
| N286G | $1.5\times10^{2}$ | $2.0\times10^{-1}$ | $7.0\times10^{1}$ | $1.9\times10^{-1}$ | $2.0\times10^{6}$ | $1.4\times10^{-1}$ | $2.9\times10^{5}$ | $1.8\times10^{-2}$ | $1.3\times10^{1}$ | $2.6\times10^{-1}$ | $6.0\times10^{2}$ | $5.2\times10^{-1}$ |
| N286V | $1.9\times10^{2}$ | **$2.7\times10^{-3}$** | $9.4\times10^{1}$ | $4.0\times10^{-1}$ | $2.6\times10^{6}$ | **$7.0\times10^{-4}$** | $4.3\times10^{5}$ | $2.0\times10^{-1}$ | $1.6\times10^{1}$ | $8.9\times10^{-1}$ | $6.6\times10^{2}$ | $1.5\times10^{-1}$ |
| Q287G | $9.4\times10^{1}$ | $2.5\times10^{-1}$ | $7.9\times10^{1}$ | $5.2\times10^{-1}$ | $1.5\times10^{6}$ | $7.5\times10^{-1}$ | $4.9\times10^{5}$ | $3.2\times10^{-1}$ | $1.8\times10^{1}$ | $5.3\times10^{-1}$ | $5.7\times10^{2}$ | $1.7\times10^{-1}$ |
| Q287V | $2.0\times10^{2}$ | **$8.0\times10^{-4}$** | $9.8\times10^{1}$ | $2.2\times10^{-1}$ | $1.8\times10^{6}$ | $3.3\times10^{-1}$ | $3.3\times10^{5}$ | $2.5\times10^{-2}$ | $1.4\times10^{1}$ | $5.7\times10^{-1}$ | $7.0\times10^{2}$ | $3.9\times10^{-2}$ |

| Variant | $k_{cat}$ cMUP ($s^{-1}$) | $k_{cat}$ cMUP *p*–value | $K_M$ cMUP (μM) | $K_M$ cMUP *p*–value | $k_{cat}/K_M$ cMUP ($M^{-1}s^{-1}$) | $k_{cat}/K_M$ cMUP *p*–value | $k_{cat}/K_M$ MeP ($M^{-1}s^{-1}$) | $k_{cat}/K_M$ MeP *p*–value | $k_{cat}/K_M$ MecMUP ($M^{-1}s^{-1}$) | $k_{cat}/K_M$ MecMUP *p*–value | $K_i$ $PO_4$ (μM) | $K_i$ $PO_4$ *p*–value |
|---|---|---|---|---|---|---|---|---|---|---|---|---|
| M288G | $1.3\times10^{1}$ | **0** | $6.4\times10^{1}$ | $3.9\times10^{-2}$ | $2.0\times10^{5}$ | **$7.0\times10^{-4}$** | $2.3\times10^{4}$ | **0** | ≤ 1.3 | **$2.0\times10^{-4}$** | $6.1\times10^{2}$ | $4.9\times10^{-1}$ |
| M288V | $4.0\times10^{1}$ | **$7.5\times10^{-3}$** | $8.7\times10^{1}$ | $8.9\times10^{-1}$ | $4.7\times10^{5}$ | $2.5\times10^{-2}$ | – | – | – | – | $5.7\times10^{2}$ | $3.4\times10^{-1}$ |
| G289A | $6.4\times10^{1}$ | $6.3\times10^{-2}$ | $6.2\times10^{1}$ | $9.7\times10^{-2}$ | $1.1\times10^{6}$ | $5.2\times10^{-1}$ | $1.8\times10^{5}$ | **$3.2\times10^{-3}$** | $1.2\times10^{1}$ | $2.1\times10^{-1}$ | $5.9\times10^{2}$ | $4.7\times10^{-1}$ |
| V290A | $1.4\times10^{2}$ | $2.6\times10^{-1}$ | $9.8\times10^{1}$ | $1.9\times10^{-1}$ | $1.6\times10^{6}$ | $6.0\times10^{-1}$ | $5.5\times10^{5}$ | $5.8\times10^{-1}$ | $1.6\times10^{1}$ | $9.9\times10^{-1}$ | $6.6\times10^{2}$ | $7.5\times10^{-2}$ |
| V290G | $1.1\times10^{2}$ | $6.0\times10^{-1}$ | $7.1\times10^{1}$ | $3.4\times10^{-1}$ | $1.6\times10^{6}$ | $7.5\times10^{-1}$ | $1.6\times10^{6}$ | **0** | $1.4\times10^{1}$ | $4.8\times10^{-1}$ | $6.3\times10^{2}$ | $6.7\times10^{-1}$ |
| D291G | $1.3\times10^{2}$ | $6.2\times10^{-1}$ | $5.9\times10^{1}$ | $6.6\times10^{-2}$ | $1.9\times10^{6}$ | $3.2\times10^{-1}$ | $4.1\times10^{5}$ | $1.5\times10^{-1}$ | $2.1\times10^{1}$ | $1.4\times10^{-1}$ | $5.5\times10^{2}$ | $2.6\times10^{-1}$ |
| D291V | $1.3\times10^{2}$ | $6.3\times10^{-1}$ | $8.1\times10^{1}$ | $6.7\times10^{-1}$ | $1.5\times10^{6}$ | $7.6\times10^{-1}$ | $6.4\times10^{4}$ | **$4.8\times10^{-3}$** | 7.7 | $1.8\times10^{-2}$ | $5.8\times10^{2}$ | $3.4\times10^{-1}$ |
| D292G | $1.4\times10^{2}$ | $4.3\times10^{-1}$ | $8.6\times10^{1}$ | $9.5\times10^{-1}$ | $1.9\times10^{6}$ | $1.5\times10^{-1}$ | $8.2\times10^{5}$ | $9.3\times10^{-2}$ | $1.9\times10^{1}$ | $4.3\times10^{-1}$ | $6.3\times10^{2}$ | $7.3\times10^{-1}$ |
| D292V | $1.0\times10^{2}$ | $3.3\times10^{-1}$ | $8.9\times10^{1}$ | $6.4\times10^{-1}$ | $1.3\times10^{6}$ | $5.3\times10^{-1}$ | $3.1\times10^{5}$ | $7.0\times10^{-2}$ | ≤ 4.9 | $2.7\times10^{-2}$ | $6.5\times10^{2}$ | $2.1\times10^{-1}$ |
| I293G | $1.1\times10^{2}$ | $7.6\times10^{-1}$ | $7.4\times10^{1}$ | $2.8\times10^{-1}$ | $1.7\times10^{6}$ | $5.4\times10^{-1}$ | $4.5\times10^{5}$ | $1.8\times10^{-1}$ | $1.9\times10^{1}$ | $5.0\times10^{-1}$ | $6.1\times10^{2}$ | $7.9\times10^{-1}$ |
| I293V | $8.4\times10^{1}$ | $7.0\times10^{-2}$ | $8.5\times10^{1}$ | $9.7\times10^{-1}$ | $9.0\times10^{5}$ | $1.1\times10^{-1}$ | $6.0\times10^{5}$ | $9.7\times10^{-1}$ | 9.8 | $5.4\times10^{-2}$ | $6.2\times10^{2}$ | $8.9\times10^{-1}$ |
| T294G | $3.8\times10^{1}$ | $2.0\times10^{-2}$ | $4.3\times10^{1}$ | $2.6\times10^{-2}$ | $6.5\times10^{5}$ | $1.4\times10^{-1}$ | $1.2\times10^{5}$ | **$6.2\times10^{-3}$** | 6.3 | **$1.9\times10^{-3}$** | $5.3\times10^{2}$ | $2.4\times10^{-2}$ |
| T294V | $1.8\times10^{2}$ | **$5.9\times10^{-3}$** | $9.6\times10^{1}$ | $2.4\times10^{-1}$ | $1.8\times10^{6}$ | $1.8\times10^{-1}$ | $3.8\times10^{5}$ | $1.3\times10^{-1}$ | $1.9\times10^{1}$ | $4.3\times10^{-1}$ | $6.8\times10^{2}$ | $2.9\times10^{-2}$ |
| D295G | $3.5\times10^{1}$ | **$6.8\times10^{-3}$** | $5.1\times10^{1}$ | $1.7\times10^{-2}$ | $6.6\times10^{5}$ | $7.4\times10^{-2}$ | $1.7\times10^{5}$ | $1.5\times10^{-2}$ | 6.3 | **$2.0\times10^{-3}$** | $4.9\times10^{2}$ | **$5.3\times10^{-3}$** |
| D295V | $1.1\times10^{1}$ | **0** | $6.0\times10^{1}$ | **$8.2\times10^{-3}$** | $1.6\times10^{5}$ | **$2.0\times10^{-4}$** | $1.7\times10^{4}$ | **$3.0\times10^{-4}$** | ≤ 1.3 | **0** | $5.0\times10^{2}$ | **$1.\times10^{-4}$** |
| F296G | 9.7 | **0** | $6.1\times10^{1}$ | $1.2\times10^{-2}$ | $1.6\times10^{5}$ | **$1.\times10^{-4}$** | $1.8\times10^{4}$ | **$3.0\times10^{-4}$** | 1.5 | **$8.2\times10^{-3}$** | $4.3\times10^{2}$ | **0** |
| F296V | $6.6\times10^{1}$ | $5.1\times10^{-2}$ | $7.6\times10^{1}$ | $4.7\times10^{-1}$ | $9.8\times10^{5}$ | $3.0\times10^{-1}$ | $4.4\times10^{5}$ | $1.7\times10^{-1}$ | ≤ 4.0 | **$2.3\times10^{-3}$** | $4.8\times10^{2}$ | **$9.0\times10^{-4}$** |
| L297G | $4.0\times10^{-1}$ | **$1.1\times10^{-3}$** | $4.8\times10^{1}$ | $1.1\times10^{-2}$ | $7.6\times10^{3}$ | **$1.8\times10^{-3}$** | ≤ $2.1\times10^{3}$ | **0** | ≤ 1.1 | **0** | $4.6\times10^{2}$ | $2.8\times10^{-2}$ |
| L297V | $1.7\times10^{2}$ | $1.9\times10^{-2}$ | $8.6\times10^{1}$ | $9.8\times10^{-1}$ | $1.6\times10^{6}$ | $5.0\times10^{-1}$ | $3.4\times10^{5}$ | $1.7\times10^{-2}$ | $1.4\times10^{1}$ | $4.2\times10^{-1}$ | $6.2\times10^{2}$ | $7.2\times10^{-1}$ |
| T298G | $7.8\times10^{1}$ | $4.9\times10^{-2}$ | $6.3\times10^{1}$ | $3.4\times10^{-2}$ | $1.2\times10^{6}$ | $4.7\times10^{-1}$ | $3.2\times10^{5}$ | $3.6\times10^{-2}$ | $1.2\times10^{1}$ | $2.1\times10^{-1}$ | $4.9\times10^{2}$ | **$6.0\times10^{-4}$** |
| T298V | $1.9\times10^{2}$ | **$2.3\times10^{-3}$** | $7.5\times10^{1}$ | $2.4\times10^{-1}$ | $2.6\times10^{6}$ | **$4.0\times10^{-4}$** | $9.6\times10^{5}$ | **$7.5\times10^{-3}$** | $1.9\times10^{1}$ | $5.3\times10^{-1}$ | $5.8\times10^{2}$ | $2.8\times10^{-1}$ |
| V299A | $8.3\times10^{1}$ | $1.2\times10^{-1}$ | $8.1\times10^{1}$ | $7.1\times10^{-1}$ | $1.0\times10^{6}$ | $1.9\times10^{-1}$ | $1.0\times10^{5}$ | **$5.9\times10^{-3}$** | $1.1\times10^{1}$ | $2.6\times10^{-1}$ | $5.1\times10^{2}$ | $4.6\times10^{-2}$ |
| V299G | $9.3\times10^{-1}$ | **$2.0\times10^{-4}$** | $6.7\times10^{1}$ | $1.1\times10^{-1}$ | $1.4\times10^{4}$ | **$5.0\times10^{-4}$** | $1.2\times10^{4}$ | **$3.3\times10^{-3}$** | ≤ 1.3 | **$1.\times10^{-4}$** | $3.9\times10^{2}$ | **0** |
| N300G | $1.3\times10^{2}$ | $5.7\times10^{-1}$ | $3.5\times10^{1}$ | **0** | $3.5\times10^{6}$ | **0** | $6.5\times10^{5}$ | $8.3\times10^{-1}$ | 6.8 | **$3.6\times10^{-3}$** | $2.5\times10^{2}$ | **$2.0\times10^{-4}$** |
| N300V | 2.8 | **$1.\times10^{-4}$** | $1.9\times10^{1}$ | **0** | $1.3\times10^{5}$ | **$2.0\times10^{-4}$** | $2.8\times10^{3}$ | **$9.5\times10^{-3}$** | 1.4 | **$4.0\times10^{-4}$** | $5.1\times10^{2}$ | **$3.6\times10^{-3}$** |
| L301G | 9.3 | **0** | $6.2\times10^{1}$ | $1.0\times10^{-2}$ | $1.5\times10^{5}$ | **0** | $1.5\times10^{4}$ | **0** | ≤ 1.1 | **0** | $6.1\times10^{2}$ | $7.2\times10^{-1}$ |
| L301V | $8.0\times10^{1}$ | $1.4\times10^{-1}$ | $1.5\times10^{2}$ | **$1.\times10^{-4}$** | $6.1\times10^{5}$ | $5.7\times10^{-2}$ | $3.2\times10^{5}$ | $2.8\times10^{-2}$ | 5.5 | $4.5\times10^{-2}$ | $7.7\times10^{2}$ | **$3.2\times10^{-3}$** |
| A302G | $8.3\times10^{1}$ | $1.5\times10^{-1}$ | $5.2\times10^{1}$ | $1.8\times10^{-2}$ | $1.7\times10^{6}$ | $5.6\times10^{-1}$ | $1.1\times10^{5}$ | **$4.5\times10^{-3}$** | $1.1\times10^{1}$ | $1.3\times10^{-1}$ | $1.0\times10^{3}$ | **$4.0\times10^{-4}$** |
| A302V | $2.5\times10^{1}$ | **0** | $9.7\times10^{1}$ | $1.6\times10^{-1}$ | $2.9\times10^{5}$ | **0** | $6.4\times10^{3}$ | **$2.9\times10^{-3}$** | $3.6\times10^{1}$ | **$1.\times10^{-4}$** | $7.9\times10^{2}$ | **0** |
| S303G | $1.6\times10^{2}$ | $1.5\times10^{-1}$ | $9.3\times10^{1}$ | $5.7\times10^{-1}$ | $1.7\times10^{6}$ | $5.4\times10^{-1}$ | $3.5\times10^{5}$ | $4.3\times10^{-2}$ | $1.5\times10^{1}$ | $7.7\times10^{-1}$ | $7.5\times10^{2}$ | $3.2\times10^{-2}$ |
| S303V | $5.2\times10^{1}$ | **$1.3\times10^{-3}$** | $5.8\times10^{1}$ | **$5.8\times10^{-3}$** | $8.9\times10^{5}$ | $8.5\times10^{-2}$ | $8.1\times10^{4}$ | **$1.2\times10^{-3}$** | 2.9 | **$6.0\times10^{-4}$** | $6.9\times10^{2}$ | $8.2\times10^{-2}$ |
| T304G | $1.6\times10^{2}$ | $1.3\times10^{-1}$ | $5.7\times10^{1}$ | $2.1\times10^{-2}$ | $2.8\times10^{6}$ | **$7.0\times10^{-4}$** | ≤ $3.2\times10^{5}$ | $8.0\times10^{-2}$ | $1.8\times10^{1}$ | $4.7\times10^{-1}$ | $4.3\times10^{2}$ | **$4.0\times10^{-4}$** |
| T304V | $1.0\times10^{1}$ | **$2.0\times10^{-4}$** | $7.9\times10^{1}$ | $5.2\times10^{-1}$ | $1.2\times10^{5}$ | **$7.0\times10^{-4}$** | $1.0\times10^{5}$ | **0** | ≤ 1.8 | **$1.\times10^{-4}$** | $7.7\times10^{2}$ | **$1.\times10^{-4}$** |
| D305G | $7.4\times10^{-1}$ | **0** | ≤ 7.1 | **0** | ≥ $1.0\times10^{5}$ | **$1.\times10^{-4}$** | ≤ $2.9\times10^{3}$ | **0** | ≤ 1.3 | **0** | ≤ 2.0 | **$4.0\times10^{-4}$** |
| D305V | – | – | – | – | ≤ $2.1\times10^{2}$ | **0** | ≤ $5.4\times10^{2}$ | **$3.0\times10^{-4}$** | ≤ 1.3 | **0** | – | – |
| Y306G | $4.6\times10^{1}$ | **$2.6\times10^{-3}$** | 9.9 | **0** | $5.3\times10^{6}$ | **0** | $8.0\times10^{4}$ | **0** | $9.5\times10^{1}$ | **0** | $1.4\times10^{2}$ | **0** |
| Y306V | $5.9\times10^{1}$ | **$5.3\times10^{-3}$** | $6.0\times10^{1}$ | $1.9\times10^{-2}$ | $8.7\times10^{5}$ | $9.8\times10^{-2}$ | $5.7\times10^{4}$ | **0** | $1.7\times10^{1}$ | $8.4\times10^{-1}$ | $5.2\times10^{2}$ | **$6.6\times10^{-3}$** |
| V307A | $2.2\times10^{2}$ | **$1.\times10^{-4}$** | $1.1\times10^{2}$ | $4.2\times10^{-2}$ | $2.2\times10^{6}$ | $2.8\times10^{-2}$ | $2.8\times10^{5}$ | **$6.5\times10^{-3}$** | $1.6\times10^{1}$ | $8.8\times10^{-1}$ | $8.4\times10^{2}$ | **$2.0\times10^{-4}$** |
| V307G | $1.1\times10^{2}$ | $5.9\times10^{-1}$ | $8.4\times10^{1}$ | $8.9\times10^{-1}$ | $2.0\times10^{6}$ | $1.6\times10^{-1}$ | $1.4\times10^{5}$ | $2.3\times10^{-2}$ | 9.3 | $4.4\times10^{-2}$ | $8.3\times10^{2}$ | **$8.0\times10^{-4}$** |
| G308A | $4.7\times10^{1}$ | **$2.8\times10^{-3}$** | $8.1\times10^{1}$ | $6.7\times10^{-1}$ | $6.4\times10^{5}$ | $3.6\times10^{-2}$ | $2.8\times10^{5}$ | $1.4\times10^{-2}$ | $1.2\times10^{1}$ | $3.0\times10^{-1}$ | $5.1\times10^{2}$ | **$1.2\times10^{-3}$** |
| G308V | $1.6\times10^{1}$ | **0** | $7.9\times10^{1}$ | $3.5\times10^{-1}$ | $2.3\times10^{5}$ | **0** | $7.4\times10^{4}$ | **0** | ≤ 1.8 | **0** | $4.7\times10^{2}$ | **0** |
| H309G | $4.3\times10^{1}$ | **0** | $1.2\times10^{2}$ | **$3.0\times10^{-4}$** | $3.2\times10^{5}$ | **$3.0\times10^{-4}$** | $1.0\times10^{5}$ | **$5.3\times10^{-3}$** | ≤ 1.1 | **0** | $8.3\times10^{2}$ | **0** |
| H309V | 3.0 | **0** | $5.8\times10^{2}$ | **0** | $5.2\times10^{3}$ | **0** | ≤ $1.1\times10^{3}$ | **0** | ≤ 1.3 | **$1.\times10^{-4}$** | $9.6\times10^{2}$ | **0** |
| N310G | $9.9\times10^{1}$ | $3.5\times10^{-1}$ | $6.7\times10^{1}$ | $1.3\times10^{-1}$ | $1.3\times10^{6}$ | $6.8\times10^{-1}$ | $2.4\times10^{5}$ | **$8.2\times10^{-3}$** | $1.7\times10^{1}$ | $8.9\times10^{-1}$ | $6.0\times10^{2}$ | $5.0\times10^{-1}$ |
| N310V | $8.0\times10^{1}$ | $2.7\times10^{-2}$ | $6.3\times10^{1}$ | $3.4\times10^{-2}$ | $1.4\times10^{6}$ | $8.7\times10^{-1}$ | $4.3\times10^{5}$ | $8.9\times10^{-2}$ | 9.8 | $1.2\times10^{-1}$ | $4.7\times10^{2}$ | **$2.0\times10^{-4}$** |
| F311G | $3.2\times10^{1}$ | **$1.2\times10^{-3}$** | $6.3\times10^{1}$ | $6.5\times10^{-2}$ | $5.2\times10^{5}$ | $1.6\times10^{-2}$ | $3.0\times10^{5}$ | $1.9\times10^{-2}$ | 5.4 | **$6.0\times10^{-3}$** | $4.1\times10^{2}$ | **$1.\times10^{-4}$** |
| F311V | $6.7\times10^{1}$ | $2.7\times10^{-2}$ | $9.1\times10^{1}$ | $5.3\times10^{-1}$ | $7.9\times10^{5}$ | $6.9\times10^{-2}$ | $3.1\times10^{5}$ | $1.1\times10^{-2}$ | 6.7 | $1.2\times10^{-2}$ | $5.9\times10^{2}$ | $3.8\times10^{-1}$ |
| G312A | $5.7\times10^{1}$ | **$9.5\times10^{-3}$** | $5.3\times10^{1}$ | $1.1\times10^{-2}$ | $1.0\times10^{6}$ | $3.2\times10^{-1}$ | $3.0\times10^{5}$ | $5.0\times10^{-2}$ | $1.2\times10^{1}$ | $2.0\times10^{-1}$ | $5.6\times10^{2}$ | $6.6\times10^{-2}$ |
| G312V | – | – | – | – | ≤ $2.8\times10^{4}$ | **0** | $1.1\times10^{3}$ | $1.2\times10^{-2}$ | ≤ 1.3 | **$9.0\times10^{-4}$** | – | $8.1\times10^{-2}$ |
| P313G | $4.6\times10^{1}$ | **$3.4\times10^{-3}$** | $5.2\times10^{1}$ | **$6.6\times10^{-3}$** | $1.0\times10^{6}$ | $2.8\times10^{-1}$ | $3.2\times10^{5}$ | $2.3\times10^{-2}$ | $1.5\times10^{1}$ | $8.1\times10^{-1}$ | $5.4\times10^{2}$ | $2.3\times10^{-2}$ |
| P313V | $1.7\times10^{2}$ | $4.5\times10^{-2}$ | $9.6\times10^{1}$ | $3.3\times10^{-1}$ | $1.7\times10^{6}$ | $4.5\times10^{-1}$ | $7.1\times10^{5}$ | $3.3\times10^{-1}$ | $1.2\times10^{1}$ | $4.7\times10^{-1}$ | $6.9\times10^{2}$ | $4.2\times10^{-2}$ |
| N314G | $4.4\times10^{1}$ | **$3.0\times10^{-4}$** | $4.3\times10^{1}$ | **$1.\times10^{-4}$** | $1.0\times10^{6}$ | $2.6\times10^{-1}$ | $2.8\times10^{5}$ | $2.0\times10^{-2}$ | 9.1 | $1.8\times10^{-2}$ | $4.0\times10^{2}$ | **$1.\times10^{-4}$** |
| N314V | $9.9\times10^{1}$ | $2.5\times10^{-1}$ | $5.7\times10^{1}$ | **$7.0\times10^{-3}$** | $1.9\times10^{6}$ | $1.4\times10^{-1}$ | $1.3\times10^{5}$ | **$1.8\times10^{-3}$** | $2.4\times10^{1}$ | $1.5\times10^{-2}$ | $4.4\times10^{2}$ | **$2.0\times10^{-4}$** |
| S315G | $5.0\times10^{1}$ | **$1.6\times10^{-3}$** | $5.8\times10^{1}$ | $1.3\times10^{-2}$ | $7.7\times10^{5}$ | $5.2\times10^{-2}$ | $1.8\times10^{5}$ | $1.1\times10^{-2}$ | 4.9 | **$5.0\times10^{-4}$** | $4.6\times10^{2}$ | **0** |
| S315V | $1.5\times10^{1}$ | **$9.0\times10^{-4}$** | $3.2\times10^{1}$ | **$2.0\times10^{-4}$** | $4.1\times10^{5}$ | **$7.7\times10^{-3}$** | $1.4\times10^{4}$ | $3.3\times10^{-2}$ | 2.9 | **$5.0\times10^{-4}$** | $2.1\times10^{2}$ | **0** |
| I316G | $3.6\times10^{1}$ | **$7.0\times10^{-4}$** | $5.0\times10^{1}$ | **$4.1\times10^{-3}$** | $6.6\times10^{5}$ | $4.0\times10^{-2}$ | $1.6\times10^{5}$ | **$2.9\times10^{-3}$** | ≤ $7.5\times10^{-1}$ | **0** | $3.7\times10^{2}$ | **$1.\times10^{-4}$** |
| I316V | $1.5\times10^{2}$ | $1.1\times10^{-1}$ | $9.4\times10^{1}$ | $4.0\times10^{-1}$ | $1.7\times10^{6}$ | $3.5\times10^{-1}$ | $5.0\times10^{5}$ | $3.1\times10^{-1}$ | $1.2\times10^{1}$ | $3.0\times10^{-1}$ | $6.6\times10^{2}$ | $2.1\times10^{-1}$ |
| E317G | 9.6 | **0** | $3.1\times10^{1}$ | **0** | $3.3\times10^{5}$ | **$3.0\times10^{-4}$** | $2.0\times10^{4}$ | **0** | 2.1 | **0** | $2.3\times10^{2}$ | **0** |
| E317V | $1.1\times10^{1}$ | **0** | $2.5\times10^{1}$ | **0** | $5.5\times10^{5}$ | **$2.3\times10^{-3}$** | $1.8\times10^{4}$ | **$7.0\times10^{-4}$** | 2.9 | **$1.\times10^{-4}$** | $1.5\times10^{2}$ | **0** |
| V318A | $2.1\times10^{2}$ | **$2.0\times10^{-4}$** | $1.0\times10^{2}$ | $1.4\times10^{-1}$ | $2.0\times10^{6}$ | $7.1\times10^{-2}$ | $2.0\times10^{5}$ | **$4.4\times10^{-3}$** | $1.5\times10^{1}$ | $7.3\times10^{-1}$ | $7.1\times10^{2}$ | $1.5\times10^{-2}$ |
| V318G | $8.2\times10^{1}$ | $1.2\times10^{-1}$ | $6.8\times10^{1}$ | $1.5\times10^{-1}$ | $1.2\times10^{6}$ | $6.0\times10^{-1}$ | $2.5\times10^{5}$ | $1.4\times10^{-2}$ | 5.7 | **$1.6\times10^{-3}$** | $5.4\times10^{2}$ | $3.6\times10^{-2}$ |
| E319G | $3.7\times10^{1}$ | **$1.1\times10^{-3}$** | $6.5\times10^{1}$ | $8.2\times10^{-2}$ | $5.3\times10^{5}$ | $1.8\times10^{-2}$ | $1.4\times10^{5}$ | $1.3\times10^{-2}$ | 5.7 | **$5.0\times10^{-4}$** | $4.8\times10^{2}$ | **$2.0\times10^{-4}$** |
| E319V | $5.5\times10^{1}$ | $1.1\times10^{-2}$ | $7.6\times10^{1}$ | $3.9\times10^{-1}$ | $6.6\times10^{5}$ | $4.8\times10^{-2}$ | $2.0\times10^{5}$ | **$4.0\times10^{-3}$** | 8.5 | $5.6\times10^{-2}$ | $5.2\times10^{2}$ | $2.0\times10^{-2}$ |
| D320G | $1.0\times10^{1}$ | **0** | $4.8\times10^{1}$ | **0** | $2.1\times10^{5}$ | **0** | $2.9\times10^{4}$ | **0** | 1.7 | **$9.0\times10^{-4}$** | $3.6\times10^{2}$ | **0** |
| D320V | 6.1 | **0** | $3.6\times10^{1}$ | **0** | $1.6\times10^{5}$ | **$1.\times10^{-4}$** | $6.3\times10^{3}$ | **0** | 1.3 | **$5.0\times10^{-4}$** | $3.1\times10^{2}$ | **0** |
| T321G | $1.0\times10^{2}$ | $5.2\times10^{-1}$ | $5.3\times10^{1}$ | $2.6\times10^{-2}$ | $1.9\times10^{6}$ | $2.8\times10^{-1}$ | $2.7\times10^{5}$ | $1.7\times10^{-2}$ | $1.3\times10^{1}$ | $3.9\times10^{-1}$ | $4.0\times10^{2}$ | $1.9\times10^{-2}$ |
| T321V | $1.5\times10^{2}$ | $2.0\times10^{-1}$ | $8.8\times10^{1}$ | $7.7\times10^{-1}$ | $1.7\times10^{6}$ | $3.6\times10^{-1}$ | $5.5\times10^{5}$ | $6.7\times10^{-1}$ | $1.6\times10^{1}$ | $9.0\times10^{-1}$ | $6.7\times10^{2}$ | $7.8\times10^{-2}$ |
| Y322G | 5.2 | **0** | $8.7\times10^{1}$ | $8.3\times10^{-1}$ | $6.1\times10^{4}$ | **0** | ≤ $1.1\times10^{4}$ | **0** | ≤ 1.4 | **0** | $8.3\times10^{2}$ | **0** |
| Y322V | $3.2\times10^{2}$ | **0** | $1.3\times10^{2}$ | **$5.0\times10^{-4}$** | $2.4\times10^{6}$ | **$4.8\times10^{-3}$** | $1.5\times10^{5}$ | **$2.0\times10^{-3}$** | $1.3\times10^{1}$ | $5.4\times10^{-1}$ | $1.0\times10^{3}$ | **$6.0\times10^{-4}$** |
| L323G | $4.6\times10^{1}$ | $2.4\times10^{-2}$ | $6.9\times10^{1}$ | $3.1\times10^{-1}$ | $5.9\times10^{5}$ | $8.7\times10^{-2}$ | $1.2\times10^{5}$ | **$5.2\times10^{-3}$** | ≤ $9.9\times10^{-1}$ | **0** | $5.2\times10^{2}$ | $1.2\times10^{-2}$ |
| L323V | $1.5\times10^{2}$ | $9.1\times10^{-2}$ | $8.6\times10^{1}$ | $9.1\times10^{-1}$ | $1.8\times10^{6}$ | $2.5\times10^{-1}$ | $4.7\times10^{5}$ | $3.6\times10^{-1}$ | $1.2\times10^{1}$ | $2.8\times10^{-1}$ | $6.6\times10^{2}$ | $1.0\times10^{-1}$ |
| R324G | $6.1\times10^{1}$ | $1.9\times10^{-2}$ | $6.1\times10^{1}$ | $4.3\times10^{-2}$ | $9.8\times10^{5}$ | $2.3\times10^{-1}$ | $1.4\times10^{5}$ | $2.8\times10^{-2}$ | 8.5 | **$8.9\times10^{-3}$** | $4.4\times10^{2}$ | **$8.0\times10^{-4}$** |
| R324V | $1.3\times10^{2}$ | $6.0\times10^{-1}$ | $8.2\times10^{1}$ | $7.6\times10^{-1}$ | $1.6\times10^{6}$ | $5.6\times10^{-1}$ | $1.1\times10^{5}$ | **$1.0\times10^{-3}$** | $1.3\times10^{1}$ | $3.0\times10^{-1}$ | $5.9\times10^{2}$ | $3.6\times10^{-1}$ |
| L325G | 1.8 | **0** | $8.5\times10^{1}$ | $9.6\times10^{-1}$ | $2.2\times10^{4}$ | **0** | $4.1\times10^{3}$ | **0** | ≤ 1.3 | **0** | $6.7\times10^{2}$ | $1.3\times10^{-1}$ |
| L325V | $1.4\times10^{2}$ | $4.6\times10^{-1}$ | $1.2\times10^{2}$ | **$2.0\times10^{-3}$** | $1.1\times10^{6}$ | $4.1\times10^{-1}$ | $1.7\times10^{5}$ | **$1.9\times10^{-3}$** | $1.2\times10^{1}$ | $2.1\times10^{-1}$ | $8.5\times10^{2}$ | **$1.\times10^{-4}$** |

| Variant | $k_{cat}$ cMUP ($s^{-1}$) | $k_{cat}$ cMUP *p*–value | $K_M$ cMUP (μM) | $K_M$ cMUP *p*–value | $k_{cat}/K_M$ cMUP ($M^{-1}s^{-1}$) | $k_{cat}/K_M$ cMUP *p*–value | $k_{cat}/K_M$ MeP ($M^{-1}s^{-1}$) | $k_{cat}/K_M$ MeP *p*–value | $k_{cat}/K_M$ MecMUP ($M^{-1}s^{-1}$) | $k_{cat}/K_M$ MecMUP *p*–value | $K_i$ $PO_4$ (μM) | $K_i$ $PO_4$ *p*–value |
|---|---|---|---|---|---|---|---|---|---|---|---|---|
| D326G | 1.1 | **0** | $5.7\times10^{1}$ | $1.0\times10^{-2}$ | $1.7\times10^{4}$ | **$1.\times10^{-4}$** | $\le 2.1\times10^{3}$ | **0** | ≤ 1.3 | **0** | $6.1\times10^{2}$ | $6.6\times10^{-1}$ |
| D326V | – | – | – | – | $\le 7.0\times10^{2}$ | **0** | $\le 6.6\times10^{3}$ | **0** | ≤ 1.3 | **0** | – | – |
| R327G | $1.1\times10^{2}$ | $6.4\times10^{-1}$ | $7.4\times10^{1}$ | $4.1\times10^{-1}$ | $1.9\times10^{6}$ | $2.5\times10^{-1}$ | $2.2\times10^{5}$ | **$5.3\times10^{-3}$** | $1.9\times10^{1}$ | $3.8\times10^{-1}$ | $5.8\times10^{2}$ | $2.5\times10^{-1}$ |
| R327V | $2.3\times10^{2}$ | **0** | $9.1\times10^{1}$ | $5.5\times10^{-1}$ | $2.1\times10^{6}$ | $2.2\times10^{-2}$ | $3.2\times10^{5}$ | $2.8\times10^{-2}$ | $1.9\times10^{1}$ | $5.9\times10^{-1}$ | $6.6\times10^{2}$ | $1.8\times10^{-1}$ |
| D328G | $6.8\times10^{1}$ | $1.9\times10^{-2}$ | $7.2\times10^{1}$ | $1.9\times10^{-1}$ | $1.0\times10^{6}$ | $2.3\times10^{-1}$ | $1.6\times10^{5}$ | **$2.1\times10^{-3}$** | $1.2\times10^{1}$ | $2.1\times10^{-1}$ | $5.9\times10^{2}$ | $5.2\times10^{-1}$ |
| D328V | $5.2\times10^{1}$ | **$5.8\times10^{-3}$** | $9.8\times10^{1}$ | $2.5\times10^{-1}$ | $6.3\times10^{5}$ | $3.3\times10^{-2}$ | $1.4\times10^{5}$ | **$1.5\times10^{-3}$** | $1.0\times10^{1}$ | $1.9\times10^{-1}$ | $6.2\times10^{2}$ | $7.6\times10^{-1}$ |
| L329G | $9.9\times10^{-2}$ | $1.5\times10^{-2}$ | $3.5\times10^{1}$ | **$7.3\times10^{-3}$** | $2.8\times10^{3}$ | $1.2\times10^{-2}$ | $\le 2.0\times10^{3}$ | **0** | ≤ 1.1 | **0** | – | – |
| L329V | $1.3\times10^{2}$ | $4.3\times10^{-1}$ | $8.4\times10^{1}$ | $9.0\times10^{-1}$ | $1.6\times10^{6}$ | $4.7\times10^{-1}$ | $3.6\times10^{5}$ | $2.4\times10^{-2}$ | $1.5\times10^{1}$ | $7.5\times10^{-1}$ | $6.3\times10^{2}$ | $5.8\times10^{-1}$ |
| A330G | $1.9\times10^{2}$ | **$8.9\times10^{-3}$** | $7.9\times10^{1}$ | $5.6\times10^{-1}$ | $2.3\times10^{6}$ | $1.5\times10^{-2}$ | $4.3\times10^{5}$ | $1.5\times10^{-1}$ | $1.5\times10^{1}$ | $8.0\times10^{-1}$ | $5.5\times10^{2}$ | $4.9\times10^{-2}$ |
| A330V | $1.7\times10^{2}$ | $3.6\times10^{-2}$ | $9.8\times10^{1}$ | $2.0\times10^{-1}$ | $1.7\times10^{6}$ | $5.0\times10^{-1}$ | $3.0\times10^{5}$ | **$6.6\times10^{-3}$** | $1.2\times10^{1}$ | $3.8\times10^{-1}$ | $6.7\times10^{2}$ | $1.4\times10^{-1}$ |
| D331G | $1.1\times10^{2}$ | $7.5\times10^{-1}$ | $7.3\times10^{1}$ | $3.5\times10^{-1}$ | $1.5\times10^{6}$ | $7.9\times10^{-1}$ | $2.9\times10^{5}$ | $1.8\times10^{-2}$ | $1.8\times10^{1}$ | $6.5\times10^{-1}$ | $5.9\times10^{2}$ | $6.0\times10^{-1}$ |
| D331V | $9.1\times10^{1}$ | $1.8\times10^{-1}$ | $9.0\times10^{1}$ | $7.0\times10^{-1}$ | $9.2\times10^{5}$ | $1.4\times10^{-1}$ | $5.3\times10^{5}$ | $5.7\times10^{-1}$ | $1.0\times10^{1}$ | $8.0\times10^{-2}$ | $6.0\times10^{2}$ | $5.9\times10^{-1}$ |
| F332G | $4.7\times10^{-1}$ | **$1.\times10^{-4}$** | $6.4\times10^{1}$ | $3.8\times10^{-2}$ | $7.5\times10^{3}$ | **$1.\times10^{-4}$** | $\le 5.3\times10^{3}$ | **0** | $\le 9.9\times10^{-1}$ | **0** | $5.7\times10^{2}$ | $1.1\times10^{-1}$ |
| F332V | $9.9\times10^{1}$ | $4.6\times10^{-1}$ | $8.1\times10^{1}$ | $7.6\times10^{-1}$ | $1.5\times10^{6}$ | $8.1\times10^{-1}$ | $6.1\times10^{4}$ | **0** | 9.3 | $1.3\times10^{-1}$ | $5.5\times10^{2}$ | $1.0\times10^{-1}$ |
| F333G | 4.7 | **$4.0\times10^{-4}$** | $7.3\times10^{1}$ | $2.8\times10^{-1}$ | $6.5\times10^{4}$ | **$6.0\times10^{-4}$** | $\le 6.2\times10^{3}$ | **0** | ≤ 1.1 | **0** | $4.5\times10^{2}$ | **0** |
| F333V | $1.7\times10^{2}$ | $4.6\times10^{-2}$ | $9.7\times10^{1}$ | $3.1\times10^{-1}$ | $1.9\times10^{6}$ | $2.1\times10^{-1}$ | $9.6\times10^{4}$ | **$1.8\times10^{-3}$** | 9.9 | $5.6\times10^{-2}$ | $6.3\times10^{2}$ | $5.7\times10^{-1}$ |
| N334G | $1.0\times10^{2}$ | $2.1\times10^{-1}$ | $6.7\times10^{1}$ | $5.3\times10^{-2}$ | $1.2\times10^{6}$ | $5.3\times10^{-1}$ | $4.8\times10^{5}$ | $2.8\times10^{-1}$ | $1.7\times10^{1}$ | $7.0\times10^{-1}$ | $5.2\times10^{2}$ | **$7.3\times10^{-3}$** |
| N334V | $1.4\times10^{2}$ | $4.1\times10^{-1}$ | $1.0\times10^{2}$ | $1.0\times10^{-1}$ | $1.4\times10^{6}$ | $7.8\times10^{-1}$ | $8.4\times10^{5}$ | $7.2\times10^{-2}$ | $1.1\times10^{1}$ | $1.3\times10^{-1}$ | $6.3\times10^{2}$ | $6.2\times10^{-1}$ |
| N335G | $8.7\times10^{1}$ | $1.3\times10^{-1}$ | $7.1\times10^{1}$ | $2.3\times10^{-1}$ | $1.5\times10^{6}$ | $8.8\times10^{-1}$ | $6.1\times10^{5}$ | $9.4\times10^{-1}$ | $1.8\times10^{1}$ | $4.7\times10^{-1}$ | $6.1\times10^{2}$ | $9.2\times10^{-1}$ |
| N335V | $2.0\times10^{2}$ | **$2.0\times10^{-4}$** | $9.7\times10^{1}$ | $2.2\times10^{-1}$ | $2.1\times10^{6}$ | $3.3\times10^{-2}$ | $6.0\times10^{5}$ | $9.2\times10^{-1}$ | $1.6\times10^{1}$ | $9.3\times10^{-1}$ | $6.8\times10^{2}$ | $7.4\times10^{-2}$ |
| L336G | $2.4\times10^{-1}$ | **0** | $4.0\times10^{1}$ | **$5.0\times10^{-4}$** | $4.6\times10^{3}$ | **$5.0\times10^{-4}$** | $\le 2.3\times10^{3}$ | **0** | ≤ 1.1 | **0** | $6.7\times10^{2}$ | $4.5\times10^{-1}$ |
| L336V | $9.5\times10^{1}$ | $2.9\times10^{-1}$ | $8.5\times10^{1}$ | $9.8\times10^{-1}$ | $7.8\times10^{5}$ | $1.4\times10^{-1}$ | $4.6\times10^{5}$ | $1.4\times10^{-1}$ | $1.1\times10^{1}$ | $2.4\times10^{-1}$ | $5.6\times10^{2}$ | $1.4\times10^{-1}$ |
| D337G | $1.0\times10^{1}$ | **0** | $7.4\times10^{1}$ | $1.6\times10^{-1}$ | $1.4\times10^{5}$ | **0** | $2.3\times10^{4}$ | **0** | ≤ 1.3 | **0** | $5.8\times10^{2}$ | $2.0\times10^{-1}$ |
| D337V | $2.1\times10^{1}$ | **$6.0\times10^{-4}$** | $8.0\times10^{1}$ | $6.6\times10^{-1}$ | $2.5\times10^{5}$ | **$4.8\times10^{-3}$** | $9.8\times10^{3}$ | **0** | $\le 5.3\times10^{-1}$ | **$8.0\times10^{-4}$** | $4.5\times10^{2}$ | **$8.0\times10^{-4}$** |
| K338G | $7.9\times10^{1}$ | $4.7\times10^{-2}$ | $6.9\times10^{1}$ | $1.0\times10^{-1}$ | $1.3\times10^{6}$ | $7.3\times10^{-1}$ | $2.2\times10^{5}$ | $2.3\times10^{-2}$ | $1.5\times10^{1}$ | $7.0\times10^{-1}$ | $5.7\times10^{2}$ | $9.0\times10^{-2}$ |
| K338V | $2.2\times10^{2}$ | **$7.0\times10^{-4}$** | $9.8\times10^{1}$ | $2.8\times10^{-1}$ | $1.8\times10^{6}$ | $2.6\times10^{-1}$ | $3.6\times10^{5}$ | $3.3\times10^{-2}$ | $1.8\times10^{1}$ | $7.3\times10^{-1}$ | $6.5\times10^{2}$ | $2.6\times10^{-1}$ |
| K339G | $5.3\times10^{1}$ | $2.4\times10^{-2}$ | $5.4\times10^{1}$ | $4.1\times10^{-2}$ | $9.5\times10^{5}$ | $2.7\times10^{-1}$ | $2.2\times10^{5}$ | **$7.7\times10^{-3}$** | 8.5 | $1.1\times10^{-2}$ | $5.1\times10^{2}$ | **$7.2\times10^{-3}$** |
| K339V | $9.6\times10^{1}$ | $2.2\times10^{-1}$ | $8.0\times10^{1}$ | $6.5\times10^{-1}$ | $1.1\times10^{6}$ | $4.1\times10^{-1}$ | $3.9\times10^{5}$ | $6.8\times10^{-2}$ | $1.1\times10^{1}$ | $1.9\times10^{-1}$ | $5.9\times10^{2}$ | $3.9\times10^{-1}$ |
| V340A | $1.7\times10^{2}$ | $5.8\times10^{-2}$ | $8.2\times10^{1}$ | $7.5\times10^{-1}$ | $1.8\times10^{6}$ | $2.7\times10^{-1}$ | $1.1\times10^{5}$ | **$5.7\times10^{-3}$** | $1.5\times10^{1}$ | $7.9\times10^{-1}$ | $5.5\times10^{2}$ | $1.3\times10^{-1}$ |
| V340G | $1.1\times10^{1}$ | **0** | $9.0\times10^{1}$ | $6.9\times10^{-1}$ | $1.2\times10^{5}$ | **$3.0\times10^{-4}$** | $2.1\times10^{4}$ | **$1.\times10^{-4}$** | ≤ 1.3 | **0** | $7.5\times10^{2}$ | **0** |
| G341A | $3.7\times10^{1}$ | **$7.0\times10^{-4}$** | $5.9\times10^{1}$ | $3.3\times10^{-2}$ | $6.2\times10^{5}$ | $3.2\times10^{-2}$ | $2.4\times10^{5}$ | **$9.1\times10^{-3}$** | 5.3 | **$7.0\times10^{-4}$** | $5.8\times10^{2}$ | $3.0\times10^{-1}$ |
| G341V | $8.5\times10^{1}$ | $7.6\times10^{-2}$ | $8.6\times10^{1}$ | $9.6\times10^{-1}$ | $1.0\times10^{6}$ | $2.1\times10^{-1}$ | $4.6\times10^{4}$ | **0** | 5.6 | **$9.0\times10^{-4}$** | $6.1\times10^{2}$ | $8.7\times10^{-1}$ |
| K342G | $7.9\times10^{1}$ | $1.4\times10^{-1}$ | $6.9\times10^{1}$ | $2.4\times10^{-1}$ | $1.4\times10^{6}$ | $9.9\times10^{-1}$ | $3.3\times10^{5}$ | $3.7\times10^{-2}$ | $1.3\times10^{1}$ | $3.7\times10^{-1}$ | $5.4\times10^{2}$ | $1.1\times10^{-1}$ |
| K342V | $1.7\times10^{2}$ | $1.1\times10^{-2}$ | $8.6\times10^{1}$ | $9.4\times10^{-1}$ | $1.9\times10^{6}$ | $8.7\times10^{-2}$ | $4.8\times10^{5}$ | $2.2\times10^{-1}$ | $1.7\times10^{1}$ | $8.5\times10^{-1}$ | $6.6\times10^{2}$ | $1.9\times10^{-1}$ |
| G343A | $1.2\times10^{2}$ | $8.7\times10^{-1}$ | $7.4\times10^{1}$ | $4.6\times10^{-1}$ | $1.9\times10^{6}$ | $2.5\times10^{-1}$ | $8.6\times10^{5}$ | $1.0\times10^{-1}$ | $1.8\times10^{1}$ | $6.4\times10^{-1}$ | $6.1\times10^{2}$ | $9.8\times10^{-1}$ |
| G343V | $1.9\times10^{2}$ | **$4.9\times10^{-3}$** | $8.1\times10^{1}$ | $6.3\times10^{-1}$ | $2.0\times10^{6}$ | $1.1\times10^{-1}$ | $2.1\times10^{5}$ | **$4.4\times10^{-3}$** | $1.4\times10^{1}$ | $5.5\times10^{-1}$ | $6.3\times10^{2}$ | $6.8\times10^{-1}$ |
| N344G | $7.5\times10^{1}$ | $5.6\times10^{-2}$ | $5.9\times10^{1}$ | $3.6\times10^{-2}$ | $1.3\times10^{6}$ | $7.8\times10^{-1}$ | $3.1\times10^{5}$ | $2.7\times10^{-2}$ | $1.2\times10^{1}$ | $1.3\times10^{-1}$ | $5.1\times10^{2}$ | $1.2\times10^{-2}$ |
| N344V | $2.8\times10^{1}$ | **$3.7\times10^{-3}$** | $8.0\times10^{1}$ | $6.9\times10^{-1}$ | $3.5\times10^{5}$ | $1.2\times10^{-2}$ | $1.7\times10^{5}$ | **$5.0\times10^{-4}$** | ≤ 1.9 | **$1.\times10^{-4}$** | $5.3\times10^{2}$ | $2.0\times10^{-2}$ |
| Y345G | 5.6 | **0** | $6.7\times10^{1}$ | $9.6\times10^{-2}$ | $8.8\times10^{4}$ | **0** | $3.0\times10^{4}$ | **$3.9\times10^{-3}$** | ≤ 1.3 | **0** | $5.1\times10^{2}$ | **$8.0\times10^{-4}$** |
| Y345V | $1.4\times10^{2}$ | $3.3\times10^{-1}$ | $1.0\times10^{2}$ | $1.1\times10^{-1}$ | $1.4\times10^{6}$ | $9.4\times10^{-1}$ | $2.4\times10^{5}$ | **$1.9\times10^{-3}$** | $1.3\times10^{1}$ | $4.3\times10^{-1}$ | $6.6\times10^{2}$ | $1.8\times10^{-1}$ |
| L346G | $3.3\times10^{1}$ | **$7.0\times10^{-4}$** | $6.1\times10^{1}$ | $4.7\times10^{-2}$ | $4.7\times10^{5}$ | **$8.0\times10^{-3}$** | $1.2\times10^{5}$ | $3.0\times10^{-2}$ | 5.1 | **$3.0\times10^{-4}$** | $4.9\times10^{2}$ | **$2.6\times10^{-3}$** |
| L346V | $1.8\times10^{2}$ | **$5.4\times10^{-3}$** | $8.5\times10^{1}$ | 1.0 | $1.9\times10^{6}$ | $1.1\times10^{-1}$ | $5.2\times10^{5}$ | $4.5\times10^{-1}$ | $1.7\times10^{1}$ | $9.0\times10^{-1}$ | $6.3\times10^{2}$ | $4.0\times10^{-1}$ |
| V347A | $2.0\times10^{2}$ | **$9.0\times10^{-4}$** | $9.2\times10^{1}$ | $5.4\times10^{-1}$ | $2.2\times10^{6}$ | $2.6\times10^{-2}$ | $6.7\times10^{5}$ | $5.9\times10^{-1}$ | $1.3\times10^{1}$ | $4.2\times10^{-1}$ | $6.3\times10^{2}$ | $6.0\times10^{-1}$ |
| V347G | $8.8\times10^{1}$ | $1.3\times10^{-1}$ | $6.6\times10^{1}$ | $1.4\times10^{-1}$ | $1.2\times10^{6}$ | $5.5\times10^{-1}$ | $\le 8.5\times10^{4}$ | **$1.4\times10^{-3}$** | 5.3 | **$3.0\times10^{-4}$** | $5.4\times10^{2}$ | $2.8\times10^{-2}$ |
| F348G | 8.2 | **0** | $7.7\times10^{1}$ | $3.2\times10^{-1}$ | $1.4\times10^{5}$ | **0** | $2.8\times10^{4}$ | **0** | ≤ 1.1 | **0** | $5.8\times10^{2}$ | $1.8\times10^{-1}$ |
| F348V | $5.7\times10^{1}$ | **$5.1\times10^{-3}$** | $8.7\times10^{1}$ | $8.4\times10^{-1}$ | $6.8\times10^{5}$ | $2.6\times10^{-2}$ | $2.2\times10^{5}$ | **$2.0\times10^{-3}$** | 3.9 | **$7.0\times10^{-4}$** | $6.1\times10^{2}$ | $9.7\times10^{-1}$ |
| L349G | 9.4 | **0** | $8.8\times10^{1}$ | $7.2\times10^{-1}$ | $1.1\times10^{5}$ | **0** | $1.9\times10^{4}$ | **0** | $\le 9.9\times10^{-1}$ | **$1.\times10^{-4}$** | $7.5\times10^{2}$ | **$1.\times10^{-4}$** |
| L349V | $6.5\times10^{1}$ | **$6.4\times10^{-3}$** | $8.3\times10^{1}$ | $8.2\times10^{-1}$ | $6.7\times10^{5}$ | $1.7\times10^{-2}$ | $2.6\times10^{5}$ | **$9.8\times10^{-3}$** | ≤ 1.9 | **$2.9\times10^{-3}$** | $5.3\times10^{2}$ | **$7.6\times10^{-3}$** |
| S350G | $4.3\times10^{1}$ | **$3.0\times10^{-4}$** | $8.3\times10^{1}$ | $8.1\times10^{-1}$ | $5.1\times10^{5}$ | **$6.1\times10^{-3}$** | $1.6\times10^{5}$ | $1.7\times10^{-2}$ | 6.5 | **$5.0\times10^{-4}$** | $7.6\times10^{2}$ | **$1.\times10^{-4}$** |
| S350V | $8.9\times10^{1}$ | $7.7\times10^{-2}$ | $1.1\times10^{2}$ | **$5.0\times10^{-3}$** | $8.9\times10^{5}$ | $8.1\times10^{-2}$ | $1.6\times10^{5}$ | **$8.0\times10^{-4}$** | 3.9 | **$2.6\times10^{-3}$** | $7.7\times10^{2}$ | **$9.0\times10^{-4}$** |
| A351G | $8.9\times10^{1}$ | $2.2\times10^{-1}$ | $7.9\times10^{1}$ | $6.1\times10^{-1}$ | $1.1\times10^{6}$ | $3.5\times10^{-1}$ | $1.6\times10^{5}$ | $1.5\times10^{-2}$ | $\le 8.4\times10^{-1}$ | **0** | $4.1\times10^{2}$ | **$1.\times10^{-4}$** |
| A351V | $5.4\times10^{1}$ | **$2.4\times10^{-3}$** | $6.0\times10^{1}$ | **$9.7\times10^{-3}$** | $1.0\times10^{6}$ | $1.4\times10^{-1}$ | $4.6\times10^{4}$ | **0** | ≤ 1.6 | **$1.\times10^{-4}$** | $4.6\times10^{2}$ | **$1.\times10^{-4}$** |
| D352G | 4.0 | **0** | $4.6\times10^{1}$ | **$3.0\times10^{-4}$** | $1.2\times10^{5}$ | **0** | $5.1\times10^{5}$ | $4.5\times10^{-1}$ | $2.2\times10^{1}$ | $7.6\times10^{-2}$ | $2.3\times10^{1}$ | **0** |
| D352V | – | – | – | – | $\le 4.6\times10^{2}$ | **0** | $\le 1.2\times10^{3}$ | **0** | $\le 9.2\times10^{-2}$ | **0** | – | – |
| H353G | – | – | – | – | $\le 1.1\times10^{3}$ | **0** | $\le 1.9\times10^{3}$ | **0** | $3.6\times10^{-1}$ | **$3.9\times10^{-3}$** | – | – |
| H353V | $5.0\times10^{1}$ | **$2.2\times10^{-3}$** | $6.3\times10^{1}$ | $3.3\times10^{-2}$ | $9.3\times10^{5}$ | $1.3\times10^{-1}$ | $8.0\times10^{4}$ | **$1.1\times10^{-3}$** | 2.2 | **$5.0\times10^{-4}$** | $5.0\times10^{2}$ | **$6.9\times10^{-3}$** |
| G354A | $1.6\times10^{1}$ | **$1.\times10^{-4}$** | $4.4\times10^{1}$ | **$1.5\times10^{-3}$** | $3.4\times10^{5}$ | **$4.2\times10^{-3}$** | $1.1\times10^{5}$ | $2.9\times10^{-2}$ | $1.3\times10^{1}$ | $4.5\times10^{-1}$ | $5.9\times10^{2}$ | $4.6\times10^{-1}$ |
| G354V | – | – | – | – | $\le 1.3\times10^{3}$ | **0** | $\le 1.0\times10^{3}$ | **0** | ≤ 1.3 | **0** | – | – |
| A355G | $1.3\times10^{2}$ | $6.2\times10^{-1}$ | $6.7\times10^{1}$ | $1.3\times10^{-1}$ | $1.9\times10^{6}$ | $2.3\times10^{-1}$ | $4.5\times10^{5}$ | $1.8\times10^{-1}$ | $2.5\times10^{1}$ | **$1.5\times10^{-3}$** | $5.8\times10^{2}$ | $3.7\times10^{-1}$ |
| A355V | $4.0\times10^{1}$ | **0** | $8.9\times10^{1}$ | $6.6\times10^{-1}$ | $4.3\times10^{5}$ | **$3.0\times10^{-4}$** | $1.9\times10^{5}$ | **$6.0\times10^{-4}$** | ≤ 1.5 | **$1.6\times10^{-3}$** | $5.3\times10^{2}$ | $1.5\times10^{-2}$ |
| A356G | $1.2\times10^{2}$ | $8.6\times10^{-1}$ | $6.8\times10^{1}$ | $1.4\times10^{-1}$ | $1.8\times10^{6}$ | $2.5\times10^{-1}$ | $5.3\times10^{5}$ | $5.0\times10^{-1}$ | $3.5\times10^{1}$ | **0** | $4.7\times10^{2}$ | **0** |
| A356V | $6.8\times10^{1}$ | **$2.9\times10^{-3}$** | $6.9\times10^{1}$ | $5.3\times10^{-2}$ | $1.0\times10^{6}$ | $1.6\times10^{-1}$ | $3.6\times10^{5}$ | **$4.6\times10^{-3}$** | 9.0 | $2.4\times10^{-2}$ | $4.6\times10^{2}$ | **0** |
| H357G | $1.6\times10^{2}$ | $2.3\times10^{-1}$ | $3.7\times10^{1}$ | **$8.2\times10^{-3}$** | $4.8\times10^{6}$ | **0** | $2.2\times10^{6}$ | **$1.\times10^{-4}$** | $1.7\times10^{1}$ | $6.8\times10^{-1}$ | $3.1\times10^{2}$ | **$9.0\times10^{-4}$** |
| H357V | $3.5\times10^{1}$ | **$1.\times10^{-4}$** | $5.8\times10^{1}$ | $1.4\times10^{-2}$ | $8.0\times10^{5}$ | $6.7\times10^{-2}$ | $2.0\times10^{5}$ | **$6.0\times10^{-4}$** | 8.1 | $1.1\times10^{-2}$ | $4.8\times10^{2}$ | **$1.9\times10^{-3}$** |
| S358G | $3.2\times10^{1}$ | **$2.0\times10^{-4}$** | $4.6\times10^{1}$ | **$3.0\times10^{-4}$** | $9.5\times10^{5}$ | $1.5\times10^{-1}$ | $3.5\times10^{5}$ | $7.9\times10^{-2}$ | 9.2 | $3.9\times10^{-2}$ | $4.6\times10^{2}$ | **0** |
| S358V | $4.8\times10^{1}$ | **$3.9\times10^{-3}$** | $8.0\times10^{1}$ | $6.2\times10^{-1}$ | $6.8\times10^{5}$ | $5.3\times10^{-2}$ | $1.2\times10^{5}$ | **$8.0\times10^{-4}$** | 5.8 | **$6.5\times10^{-3}$** | $5.0\times10^{2}$ | **$5.7\times10^{-3}$** |
| V359A | $8.7\times10^{1}$ | $6.4\times10^{-2}$ | $8.9\times10^{1}$ | $7.0\times10^{-1}$ | $9.3\times10^{5}$ | $1.1\times10^{-1}$ | $1.6\times10^{5}$ | **$3.5\times10^{-3}$** | 8.8 | $4.9\times10^{-2}$ | $5.7\times10^{2}$ | $1.6\times10^{-1}$ |
| V359G | $4.0\times10^{1}$ | **$5.0\times10^{-4}$** | $6.8\times10^{1}$ | $9.7\times10^{-2}$ | $6.1\times10^{5}$ | $1.5\times10^{-2}$ | $3.8\times10^{5}$ | $6.1\times10^{-2}$ | 5.3 | **0** | $3.8\times10^{2}$ | **$2.0\times10^{-4}$** |
| G360A | $9.8\times10^{1}$ | $2.9\times10^{-1}$ | $7.3\times10^{1}$ | $2.1\times10^{-1}$ | $1.2\times10^{6}$ | $4.5\times10^{-1}$ | $5.5\times10^{5}$ | $6.6\times10^{-1}$ | $1.4\times10^{1}$ | $3.6\times10^{-1}$ | $5.2\times10^{2}$ | **$4.2\times10^{-3}$** |
| G360V | $5.4\times10^{1}$ | **$1.3\times10^{-3}$** | $8.0\times10^{1}$ | $5.1\times10^{-1}$ | $6.9\times10^{5}$ | $1.8\times10^{-2}$ | $3.5\times10^{5}$ | **$2.9\times10^{-3}$** | 5.8 | **$7.0\times10^{-3}$** | $5.9\times10^{2}$ | $1.7\times10^{-1}$ |
| F361G | 2.2 | **0** | $2.9\times10^{1}$ | **0** | $6.4\times10^{4}$ | **0** | $8.5\times10^{3}$ | **0** | $6.4\times10^{-1}$ | **$4.5\times10^{-3}$** | $3.0\times10^{2}$ | **0** |
| F361V | $3.2\times10^{1}$ | **$1.\times10^{-4}$** | $7.1\times10^{1}$ | $2.2\times10^{-1}$ | $4.5\times10^{5}$ | **$7.9\times10^{-3}$** | $8.4\times10^{3}$ | **0** | 4.1 | **$2.0\times10^{-4}$** | $5.1\times10^{2}$ | **$8.0\times10^{-3}$** |
| M362G | 7.4 | **0** | $5.1\times10^{1}$ | **$4.9\times10^{-3}$** | $2.1\times10^{5}$ | **$1.1\times10^{-3}$** | $1.9\times10^{5}$ | **$4.9\times10^{-3}$** | ≤ 1.5 | **0** | $4.1\times10^{2}$ | **0** |
| M362V | $2.1\times10^{1}$ | **0** | $9.4\times10^{1}$ | $3.6\times10^{-1}$ | $2.2\times10^{5}$ | **0** | $1.1\times10^{4}$ | **0** | ≤ 1.5 | **0** | $6.4\times10^{2}$ | $4.4\times10^{-1}$ |
| Q363G | $5.3\times10^{1}$ | **$7.0\times10^{-3}$** | $5.2\times10^{1}$ | **$8.5\times10^{-3}$** | $8.9\times10^{5}$ | $1.4\times10^{-1}$ | $3.1\times10^{5}$ | $1.7\times10^{-2}$ | 6.2 | **$6.0\times10^{-4}$** | $4.4\times10^{2}$ | **$1.\times10^{-4}$** |

| Variant | $k_{cat}$ cMUP ($s^{-1}$) | $k_{cat}$ cMUP *p*–value | $K_M$ cMUP (μM) | $K_M$ cMUP *p*–value | $k_{cat}/K_M$ cMUP ($M^{-1}s^{-1}$) | $k_{cat}/K_M$ cMUP *p*–value | $k_{cat}/K_M$ MeP ($M^{-1}s^{-1}$) | $k_{cat}/K_M$ MeP *p*–value | $k_{cat}/K_M$ MecMUP ($M^{-1}s^{-1}$) | $k_{cat}/K_M$ MecMUP *p*–value | $K_i$ $PO_4$ (μM) | $K_i$ $PO_4$ *p*–value |
|---|---|---|---|---|---|---|---|---|---|---|---|---|
| Q363V | $1.7\times10^{2}$ | $2.3\times10^{-2}$ | $8.8\times10^{1}$ | $7.1\times10^{-1}$ | $2.2\times10^{6}$ | $1.1\times10^{-2}$ | $3.2\times10^{5}$ | $1.1\times10^{-2}$ | $1.8\times10^{1}$ | $6.7\times10^{-1}$ | $6.4\times10^{2}$ | $4.0\times10^{-1}$ |
| A364V | $1.6\times10^{2}$ | $5.2\times10^{-2}$ | $9.3\times10^{1}$ | $3.8\times10^{-1}$ | $1.6\times10^{6}$ | $4.7\times10^{-1}$ | $2.1\times10^{5}$ | **$1.8\times10^{-3}$** | $1.5\times10^{1}$ | $9.1\times10^{-1}$ | $6.8\times10^{2}$ | $4.9\times10^{-2}$ |
| H365G | $3.8\times10^{1}$ | **$5.0\times10^{-4}$** | $7.1\times10^{1}$ | $1.6\times10^{-1}$ | $5.4\times10^{5}$ | **$9.0\times10^{-3}$** | $2.9\times10^{5}$ | $1.6\times10^{-2}$ | 5.6 | **$3.0\times10^{-4}$** | $5.8\times10^{2}$ | $3.1\times10^{-1}$ |
| H365V | $1.2\times10^{2}$ | $8.7\times10^{-1}$ | $8.6\times10^{1}$ | $9.8\times10^{-1}$ | $1.3\times10^{6}$ | $7.2\times10^{-1}$ | $2.1\times10^{5}$ | **$4.2\times10^{-3}$** | $1.6\times10^{1}$ | $9.2\times10^{-1}$ | $6.4\times10^{2}$ | $2.5\times10^{-1}$ |
| K366G | $3.2\times10^{1}$ | **$5.0\times10^{-4}$** | $4.9\times10^{1}$ | **$4.8\times10^{-3}$** | $5.4\times10^{5}$ | $1.7\times10^{-2}$ | $\leq 2.3\times10^{5}$ | $1.7\times10^{-2}$ | 8.6 | $2.1\times10^{-2}$ | $5.0\times10^{2}$ | **$6.0\times10^{-4}$** |
| K366V | $5.7\times10^{1}$ | **$5.9\times10^{-3}$** | $9.8\times10^{1}$ | $2.1\times10^{-1}$ | $5.9\times10^{5}$ | $1.3\times10^{-2}$ | $2.3\times10^{4}$ | **0** | 5.1 | **$4.4\times10^{-3}$** | $6.4\times10^{2}$ | $3.2\times10^{-1}$ |
| M367G | 6.6 | **0** | $4.9\times10^{1}$ | **$3.0\times10^{-4}$** | $1.4\times10^{5}$ | **0** | $3.7\times10^{4}$ | **$1.\times10^{-4}$** | 1.9 | **$2.0\times10^{-4}$** | $3.6\times10^{2}$ | **0** |
| M367V | $6.4\times10^{1}$ | $1.7\times10^{-2}$ | $8.3\times10^{1}$ | $7.8\times10^{-1}$ | $6.8\times10^{5}$ | $3.7\times10^{-2}$ | $9.6\times10^{4}$ | **$1.\times10^{-4}$** | 6.4 | **$9.5\times10^{-3}$** | $6.8\times10^{2}$ | $4.0\times10^{-2}$ |
| P368G | $2.1\times10^{1}$ | **0** | $6.0\times10^{1}$ | $1.8\times10^{-2}$ | $3.4\times10^{5}$ | **$1.3\times10^{-3}$** | $5.4\times10^{4}$ | **$6.0\times10^{-4}$** | 4.5 | **$2.0\times10^{-4}$** | $5.0\times10^{2}$ | **$3.0\times10^{-4}$** |
| P368V | $1.0\times10^{2}$ | $4.0\times10^{-1}$ | $8.0\times10^{1}$ | $5.6\times10^{-1}$ | $1.2\times10^{6}$ | $4.9\times10^{-1}$ | $1.4\times10^{5}$ | **$2.0\times10^{-4}$** | $1.4\times10^{1}$ | $6.0\times10^{-1}$ | $5.8\times10^{2}$ | $1.6\times10^{-1}$ |
| T369G | $1.8\times10^{2}$ | $2.7\times10^{-2}$ | $7.3\times10^{1}$ | $2.8\times10^{-1}$ | $2.4\times10^{6}$ | **$7.1\times10^{-3}$** | $4.7\times10^{5}$ | $2.7\times10^{-1}$ | $1.4\times10^{1}$ | $5.2\times10^{-1}$ | $5.3\times10^{2}$ | **$8.9\times10^{-3}$** |
| T369V | $1.4\times10^{2}$ | $3.8\times10^{-1}$ | $8.1\times10^{1}$ | $6.4\times10^{-1}$ | $1.6\times10^{6}$ | $5.4\times10^{-1}$ | $3.7\times10^{5}$ | $5.3\times10^{-2}$ | $1.0\times10^{1}$ | $1.9\times10^{-1}$ | $5.5\times10^{2}$ | $1.1\times10^{-1}$ |
| G370A | $2.8\times10^{1}$ | **0** | $7.3\times10^{1}$ | $2.3\times10^{-1}$ | $4.2\times10^{5}$ | **$3.8\times10^{-3}$** | $2.5\times10^{5}$ | $1.0\times10^{-2}$ | 4.5 | **$2.0\times10^{-4}$** | $5.7\times10^{2}$ | $8.1\times10^{-2}$ |
| G370V | $1.1\times10^{1}$ | **$2.0\times10^{-4}$** | $9.9\times10^{1}$ | $2.2\times10^{-1}$ | $1.1\times10^{5}$ | **$3.0\times10^{-4}$** | $5.1\times10^{3}$ | **0** | $\leq 1.9$ | **$4.2\times10^{-3}$** | – | – |
| F371G | $2.9\times10^{1}$ | **$2.0\times10^{-4}$** | $1.1\times10^{2}$ | $2.3\times10^{-2}$ | $2.5\times10^{5}$ | **$4.0\times10^{-4}$** | $2.7\times10^{5}$ | $1.5\times10^{-2}$ | 2.8 | **$1.\times10^{-4}$** | $5.5\times10^{2}$ | $5.3\times10^{-2}$ |
| F371V | $2.2\times10^{2}$ | **0** | $1.5\times10^{2}$ | **0** | $1.3\times10^{6}$ | $7.3\times10^{-1}$ | $5.7\times10^{5}$ | $7.6\times10^{-1}$ | 8.1 | $3.7\times10^{-2}$ | $6.2\times10^{2}$ | $9.4\times10^{-1}$ |
| F372G | $8.7\times10^{1}$ | $9.4\times10^{-2}$ | $8.2\times10^{1}$ | $7.4\times10^{-1}$ | $9.3\times10^{5}$ | $1.3\times10^{-1}$ | $5.2\times10^{5}$ | $4.8\times10^{-1}$ | $1.8\times10^{1}$ | $7.6\times10^{-1}$ | $5.9\times10^{2}$ | $4.1\times10^{-1}$ |
| F372V | $1.5\times10^{2}$ | $1.2\times10^{-1}$ | $1.2\times10^{2}$ | **$2.2\times10^{-3}$** | $1.4\times10^{6}$ | $9.0\times10^{-1}$ | $5.1\times10^{5}$ | $3.9\times10^{-1}$ | $1.1\times10^{1}$ | $1.3\times10^{-1}$ | $6.8\times10^{2}$ | $4.4\times10^{-2}$ |
| V373A | $1.2\times10^{2}$ | $9.0\times10^{-1}$ | $1.0\times10^{2}$ | $1.9\times10^{-1}$ | $1.2\times10^{6}$ | $5.3\times10^{-1}$ | $5.3\times10^{5}$ | $4.8\times10^{-1}$ | 7.3 | **$7.3\times10^{-3}$** | $7.1\times10^{2}$ | **$6.6\times10^{-3}$** |
| V373G | $6.0\times10^{1}$ | $3.6\times10^{-2}$ | $5.2\times10^{1}$ | $1.7\times10^{-2}$ | $1.0\times10^{6}$ | $3.3\times10^{-1}$ | $6.5\times10^{5}$ | $7.3\times10^{-1}$ | $1.3\times10^{1}$ | $3.8\times10^{-1}$ | $5.4\times10^{2}$ | $3.4\times10^{-2}$ |
| E374G | $1.5\times10^{2}$ | $2.0\times10^{-1}$ | $7.1\times10^{1}$ | $2.2\times10^{-1}$ | $2.2\times10^{6}$ | $4.6\times10^{-2}$ | $4.8\times10^{5}$ | $2.6\times10^{-1}$ | $2.4\times10^{1}$ | $1.1\times10^{-2}$ | $5.0\times10^{2}$ | **$6.7\times10^{-3}$** |
| E374V | $2.1\times10^{2}$ | **$2.0\times10^{-4}$** | $9.3\times10^{1}$ | $4.6\times10^{-1}$ | $2.1\times10^{6}$ | $4.2\times10^{-2}$ | $5.6\times10^{5}$ | $7.1\times10^{-1}$ | $2.4\times10^{1}$ | $1.3\times10^{-2}$ | $5.6\times10^{2}$ | $1.1\times10^{-1}$ |
| D375G | $5.9\times10^{1}$ | $2.1\times10^{-2}$ | $5.5\times10^{1}$ | $1.5\times10^{-2}$ | $8.8\times10^{5}$ | $1.6\times10^{-1}$ | $2.2\times10^{5}$ | **$7.8\times10^{-3}$** | $1.2\times10^{1}$ | $2.5\times10^{-1}$ | $4.9\times10^{2}$ | **$3.0\times10^{-4}$** |
| D375V | $9.9\times10^{1}$ | $2.1\times10^{-1}$ | $9.7\times10^{1}$ | $2.4\times10^{-1}$ | $1.1\times10^{6}$ | $2.6\times10^{-1}$ | $1.0\times10^{5}$ | **$5.3\times10^{-3}$** | $1.0\times10^{1}$ | $1.7\times10^{-1}$ | $6.1\times10^{2}$ | $8.7\times10^{-1}$ |
| M376G | $4.7\times10^{1}$ | **$9.7\times10^{-3}$** | $6.5\times10^{1}$ | $1.4\times10^{-1}$ | $4.6\times10^{5}$ | $2.1\times10^{-2}$ | $2.9\times10^{5}$ | $2.5\times10^{-2}$ | 8.7 | $1.4\times10^{-2}$ | $5.7\times10^{2}$ | $1.7\times10^{-1}$ |
| M376V | $1.2\times10^{2}$ | $9.7\times10^{-1}$ | $8.8\times10^{1}$ | $8.0\times10^{-1}$ | $1.6\times10^{6}$ | $7.0\times10^{-1}$ | $7.9\times10^{5}$ | $1.1\times10^{-1}$ | $1.3\times10^{1}$ | $3.6\times10^{-1}$ | $6.2\times10^{2}$ | $6.8\times10^{-1}$ |
| K377G | $1.5\times10^{2}$ | $3.4\times10^{-1}$ | $8.1\times10^{1}$ | $7.2\times10^{-1}$ | $1.8\times10^{6}$ | $3.8\times10^{-1}$ | $5.8\times10^{5}$ | $7.9\times10^{-1}$ | $1.4\times10^{1}$ | $5.7\times10^{-1}$ | $6.1\times10^{2}$ | $7.8\times10^{-1}$ |
| K377V | $1.5\times10^{2}$ | $8.3\times10^{-2}$ | $1.0\times10^{2}$ | $3.8\times10^{-2}$ | $1.4\times10^{6}$ | $8.7\times10^{-1}$ | $6.0\times10^{5}$ | $9.4\times10^{-1}$ | $1.0\times10^{1}$ | $1.6\times10^{-1}$ | $5.7\times10^{2}$ | $1.5\times10^{-1}$ |
| K378G | $9.0\times10^{1}$ | $1.4\times10^{-1}$ | $7.2\times10^{1}$ | $2.2\times10^{-1}$ | $1.3\times10^{6}$ | $8.5\times10^{-1}$ | $7.1\times10^{5}$ | $4.2\times10^{-1}$ | $2.4\times10^{1}$ | **$8.0\times10^{-3}$** | $5.9\times10^{2}$ | $3.5\times10^{-1}$ |
| K378V | $2.2\times10^{2}$ | **$1.2\times10^{-3}$** | $1.1\times10^{2}$ | $7.0\times10^{-2}$ | $1.7\times10^{6}$ | $4.9\times10^{-1}$ | $3.1\times10^{5}$ | $2.4\times10^{-2}$ | $1.7\times10^{1}$ | $8.0\times10^{-1}$ | $7.0\times10^{2}$ | $2.5\times10^{-2}$ |
| E379G | $1.2\times10^{2}$ | $7.8\times10^{-1}$ | $6.8\times10^{1}$ | $1.8\times10^{-1}$ | $1.9\times10^{6}$ | $1.4\times10^{-1}$ | $4.1\times10^{5}$ | $1.0\times10^{-1}$ | $1.5\times10^{1}$ | $8.7\times10^{-1}$ | $5.6\times10^{2}$ | $7.4\times10^{-2}$ |
| E379V | $2.3\times10^{2}$ | **$1.\times10^{-4}$** | $9.6\times10^{1}$ | $3.1\times10^{-1}$ | $2.1\times10^{6}$ | $3.6\times10^{-2}$ | $6.8\times10^{5}$ | $6.4\times10^{-1}$ | $2.3\times10^{1}$ | $5.8\times10^{-2}$ | $6.5\times10^{2}$ | $3.7\times10^{-1}$ |
| M380G | $1.1\times10^{2}$ | $6.3\times10^{-1}$ | $7.6\times10^{1}$ | $2.1\times10^{-1}$ | $1.4\times10^{6}$ | 1.0 | $3.2\times10^{5}$ | **$3.3\times10^{-3}$** | $1.2\times10^{1}$ | $9.4\times10^{-2}$ | $5.5\times10^{2}$ | **$3.3\times10^{-3}$** |
| M380V | $1.4\times10^{2}$ | $4.0\times10^{-1}$ | $9.1\times10^{1}$ | $6.1\times10^{-1}$ | $1.4\times10^{6}$ | $9.9\times10^{-1}$ | $3.7\times10^{5}$ | $4.7\times10^{-2}$ | $1.9\times10^{1}$ | $6.1\times10^{-1}$ | $6.3\times10^{2}$ | $5.8\times10^{-1}$ |
| N381G | $7.3\times10^{1}$ | $5.0\times10^{-2}$ | $7.4\times10^{1}$ | $3.0\times10^{-1}$ | $1.4\times10^{6}$ | $8.9\times10^{-1}$ | $5.5\times10^{5}$ | $6.2\times10^{-1}$ | $1.4\times10^{1}$ | $6.4\times10^{-1}$ | $6.4\times10^{2}$ | $4.5\times10^{-1}$ |
| N381V | $5.3\times10^{1}$ | **$2.6\times10^{-3}$** | $8.5\times10^{1}$ | $9.4\times10^{-1}$ | $5.8\times10^{5}$ | $1.2\times10^{-2}$ | $1.7\times10^{5}$ | **$3.3\times10^{-3}$** | 5.4 | **$6.0\times10^{-3}$** | $5.4\times10^{2}$ | $3.1\times10^{-2}$ |
| A382G | $1.1\times10^{2}$ | $5.9\times10^{-1}$ | $7.9\times10^{1}$ | $6.1\times10^{-1}$ | $1.4\times10^{6}$ | $8.7\times10^{-1}$ | $1.1\times10^{6}$ | **$8.1\times10^{-3}$** | $2.0\times10^{1}$ | $2.7\times10^{-1}$ | $6.3\times10^{2}$ | $7.0\times10^{-1}$ |
| A382V | $2.0\times10^{2}$ | **$5.0\times10^{-4}$** | $1.1\times10^{2}$ | $1.2\times10^{-2}$ | $1.9\times10^{6}$ | $1.1\times10^{-1}$ | $5.6\times10^{5}$ | $7.1\times10^{-1}$ | $1.5\times10^{1}$ | $8.0\times10^{-1}$ | $6.9\times10^{2}$ | $3.8\times10^{-2}$ |
| K383G | $6.4\times10^{1}$ | $5.2\times10^{-2}$ | $6.2\times10^{1}$ | $8.3\times10^{-2}$ | $1.1\times10^{6}$ | $5.7\times10^{-1}$ | $5.5\times10^{5}$ | $7.5\times10^{-1}$ | 9.3 | $1.5\times10^{-2}$ | $5.2\times10^{2}$ | $1.3\times10^{-2}$ |
| K383V | $1.7\times10^{2}$ | $1.6\times10^{-2}$ | $1.0\times10^{2}$ | $7.6\times10^{-2}$ | $1.6\times10^{6}$ | $5.6\times10^{-1}$ | $2.6\times10^{5}$ | **$3.8\times10^{-3}$** | $1.2\times10^{1}$ | $2.0\times10^{-1}$ | $7.0\times10^{2}$ | **$7.0\times10^{-3}$** |
| L384G | $5.7\times10^{1}$ | $1.4\times10^{-2}$ | $5.5\times10^{1}$ | $1.5\times10^{-2}$ | $1.2\times10^{6}$ | $4.8\times10^{-1}$ | $2.1\times10^{5}$ | **$4.6\times10^{-3}$** | $1.5\times10^{1}$ | $6.8\times10^{-1}$ | $4.9\times10^{2}$ | **$4.7\times10^{-3}$** |
| L384V | $1.1\times10^{2}$ | $7.3\times10^{-1}$ | $8.7\times10^{1}$ | $8.2\times10^{-1}$ | $1.2\times10^{6}$ | $7.0\times10^{-1}$ | $4.1\times10^{5}$ | $8.5\times10^{-2}$ | $1.3\times10^{1}$ | $2.8\times10^{-1}$ | $6.5\times10^{2}$ | $4.7\times10^{-1}$ |
| K385G | $1.6\times10^{2}$ | $9.0\times10^{-2}$ | $1.1\times10^{2}$ | $5.1\times10^{-2}$ | $1.5\times10^{6}$ | $8.3\times10^{-1}$ | $4.5\times10^{5}$ | $2.0\times10^{-1}$ | $1.2\times10^{1}$ | $2.6\times10^{-1}$ | $7.2\times10^{2}$ | **$3.9\times10^{-3}$** |
| K385V | $1.9\times10^{2}$ | **$7.0\times10^{-4}$** | $9.6\times10^{1}$ | $2.3\times10^{-1}$ | $2.1\times10^{6}$ | $2.1\times10^{-2}$ | $4.4\times10^{5}$ | $1.1\times10^{-1}$ | $2.1\times10^{1}$ | $2.8\times10^{-1}$ | $7.4\times10^{2}$ | **$1.7\times10^{-3}$** |
| Q386G | 7.6 | **$1.8\times10^{-3}$** | $6.9\times10^{1}$ | $2.5\times10^{-1}$ | $1.1\times10^{5}$ | **$3.0\times10^{-3}$** | $8.9\times10^{4}$ | **$3.4\times10^{-3}$** | $\leq 9.9\times10^{-1}$ | **0** | $4.5\times10^{2}$ | **$3.0\times10^{-4}$** |
| Q386V | $1.8\times10^{2}$ | $3.7\times10^{-2}$ | $1.0\times10^{2}$ | $2.1\times10^{-1}$ | $2.2\times10^{6}$ | $9.4\times10^{-2}$ | $4.7\times10^{5}$ | $3.1\times10^{-1}$ | $2.3\times10^{1}$ | $4.0\times10^{-2}$ | $6.3\times10^{2}$ | $6.3\times10^{-1}$ |
| K387G | $6.6\times10^{1}$ | $1.1\times10^{-2}$ | $6.5\times10^{1}$ | $6.9\times10^{-2}$ | $8.4\times10^{5}$ | $7.9\times10^{-2}$ | $4.0\times10^{5}$ | $9.7\times10^{-2}$ | $1.2\times10^{1}$ | $2.7\times10^{-1}$ | $5.5\times10^{2}$ | $6.6\times10^{-2}$ |
| K387V | $2.3\times10^{2}$ | **$1.\times10^{-4}$** | $1.2\times10^{2}$ | **$1.7\times10^{-3}$** | $1.8\times10^{6}$ | $1.9\times10^{-1}$ | $4.2\times10^{5}$ | $5.7\times10^{-2}$ | $1.6\times10^{1}$ | $9.2\times10^{-1}$ | $7.1\times10^{2}$ | $2.4\times10^{-2}$ |
| F388G | $3.2\times10^{1}$ | **$1.0\times10^{-3}$** | $6.5\times10^{1}$ | $8.5\times10^{-2}$ | $4.1\times10^{5}$ | **$5.7\times10^{-3}$** | $1.5\times10^{5}$ | **$2.5\times10^{-3}$** | $\leq 2.7$ | **$1.\times10^{-4}$** | $5.9\times10^{2}$ | $4.1\times10^{-1}$ |
| F388V | $1.6\times10^{2}$ | $1.2\times10^{-1}$ | $9.1\times10^{1}$ | $5.9\times10^{-1}$ | $1.7\times10^{6}$ | $4.3\times10^{-1}$ | $1.8\times10^{5}$ | **$7.0\times10^{-4}$** | $1.7\times10^{1}$ | $8.4\times10^{-1}$ | $6.6\times10^{2}$ | $1.7\times10^{-1}$ |
| G389A | $1.3\times10^{2}$ | $6.5\times10^{-1}$ | $7.7\times10^{1}$ | $3.5\times10^{-1}$ | $1.8\times10^{6}$ | $2.8\times10^{-1}$ | $8.2\times10^{5}$ | $9.9\times10^{-2}$ | $1.5\times10^{1}$ | $7.4\times10^{-1}$ | $6.3\times10^{2}$ | $6.3\times10^{-1}$ |
| G389V | $1.0\times10^{2}$ | $2.3\times10^{-1}$ | $8.9\times10^{1}$ | $7.7\times10^{-1}$ | $1.1\times10^{6}$ | $3.3\times10^{-1}$ | $3.4\times10^{5}$ | $8.2\times10^{-2}$ | $1.4\times10^{1}$ | $5.9\times10^{-1}$ | $6.5\times10^{2}$ | $2.5\times10^{-1}$ |
| A390G | $1.0\times10^{2}$ | $3.3\times10^{-1}$ | $7.1\times10^{1}$ | $2.4\times10^{-1}$ | $1.5\times10^{6}$ | $6.8\times10^{-1}$ | $5.6\times10^{5}$ | $7.6\times10^{-1}$ | $1.6\times10^{1}$ | $9.5\times10^{-1}$ | $5.8\times10^{2}$ | $2.3\times10^{-1}$ |
| A390V | $2.0\times10^{2}$ | **$9.0\times10^{-4}$** | $9.7\times10^{1}$ | $2.5\times10^{-1}$ | $2.1\times10^{6}$ | $5.4\times10^{-2}$ | $3.9\times10^{5}$ | $1.3\times10^{-1}$ | $2.1\times10^{1}$ | $3.4\times10^{-1}$ | $7.3\times10^{2}$ | $1.6\times10^{-2}$ |
| D391G | $1.0\times10^{2}$ | $5.1\times10^{-1}$ | $5.9\times10^{1}$ | $5.5\times10^{-2}$ | $1.2\times10^{6}$ | $6.7\times10^{-1}$ | $4.2\times10^{5}$ | $1.3\times10^{-1}$ | $1.2\times10^{1}$ | $3.7\times10^{-1}$ | $5.0\times10^{2}$ | $3.0\times10^{-2}$ |
| D391V | $1.2\times10^{2}$ | $9.3\times10^{-1}$ | $8.5\times10^{1}$ | $9.8\times10^{-1}$ | $1.3\times10^{6}$ | $7.0\times10^{-1}$ | $2.5\times10^{5}$ | **$3.3\times10^{-3}$** | $1.0\times10^{1}$ | $2.1\times10^{-1}$ | $6.1\times10^{2}$ | $7.3\times10^{-1}$ |
| N392G | $9.5\times10^{1}$ | $2.1\times10^{-1}$ | $6.9\times10^{1}$ | $2.1\times10^{-1}$ | $1.5\times10^{6}$ | $9.3\times10^{-1}$ | $1.8\times10^{6}$ | **$2.0\times10^{-4}$** | $1.1\times10^{1}$ | $2.0\times10^{-1}$ | $6.7\times10^{2}$ | $1.6\times10^{-1}$ |
| N392V | $4.6\times10^{1}$ | **0** | $9.0\times10^{1}$ | $5.7\times10^{-1}$ | $5.3\times10^{5}$ | **$2.2\times10^{-3}$** | $2.7\times10^{5}$ | **$3.0\times10^{-4}$** | 7.9 | $1.2\times10^{-2}$ | $6.3\times10^{2}$ | $4.5\times10^{-1}$ |
| I393G | $7.2\times10^{1}$ | $8.7\times10^{-2}$ | $4.3\times10^{1}$ | **$4.6\times10^{-3}$** | $1.5\times10^{6}$ | $8.9\times10^{-1}$ | $3.3\times10^{5}$ | $6.2\times10^{-2}$ | $1.2\times10^{1}$ | $2.1\times10^{-1}$ | $5.5\times10^{2}$ | $6.7\times10^{-2}$ |
| I394G | $3.1\times10^{1}$ | **$1.0\times10^{-3}$** | $6.6\times10^{1}$ | $1.1\times10^{-1}$ | $3.4\times10^{5}$ | **$4.7\times10^{-3}$** | $\leq 8.0\times10^{4}$ | **$2.4\times10^{-3}$** | 10. | $6.5\times10^{-2}$ | $5.7\times10^{2}$ | $1.6\times10^{-1}$ |
| I394V | $1.8\times10^{2}$ | **$1.6\times10^{-3}$** | $9.3\times10^{1}$ | $3.4\times10^{-1}$ | $2.0\times10^{6}$ | $4.3\times10^{-2}$ | $4.6\times10^{5}$ | $1.1\times10^{-1}$ | $2.0\times10^{1}$ | $3.9\times10^{-1}$ | $6.7\times10^{2}$ | $7.5\times10^{-2}$ |
| A395G | $7.4\times10^{1}$ | $1.5\times10^{-1}$ | $6.7\times10^{1}$ | $1.6\times10^{-1}$ | $1.4\times10^{6}$ | $9.9\times10^{-1}$ | $9.5\times10^{5}$ | $1.5\times10^{-2}$ | $1.4\times10^{1}$ | $6.8\times10^{-1}$ | $5.8\times10^{2}$ | $2.3\times10^{-1}$ |
| A395V | $2.4\times10^{2}$ | **0** | $8.4\times10^{1}$ | $8.6\times10^{-1}$ | $2.6\times10^{6}$ | **$1.\times10^{-4}$** | $3.8\times10^{5}$ | $5.8\times10^{-2}$ | $3.2\times10^{1}$ | **$3.0\times10^{-4}$** | $7.0\times10^{2}$ | $2.5\times10^{-2}$ |
| A396G | $8.0\times10^{1}$ | $1.4\times10^{-1}$ | $7.1\times10^{1}$ | $2.9\times10^{-1}$ | $1.2\times10^{6}$ | $6.8\times10^{-1}$ | $4.9\times10^{5}$ | $3.0\times10^{-1}$ | 7.3 | **$5.2\times10^{-3}$** | $4.9\times10^{2}$ | **$8.2\times10^{-3}$** |
| A396V | $1.0\times10^{2}$ | $3.1\times10^{-1}$ | $8.7\times10^{1}$ | $8.5\times10^{-1}$ | $1.3\times10^{6}$ | $5.6\times10^{-1}$ | $4.7\times10^{5}$ | $2.1\times10^{-1}$ | $1.0\times10^{1}$ | $9.0\times10^{-2}$ | $5.8\times10^{2}$ | $3.8\times10^{-1}$ |
| A397G | $6.2\times10^{1}$ | $1.0\times10^{-1}$ | $5.1\times10^{1}$ | $6.7\times10^{-2}$ | $1.2\times10^{6}$ | $7.0\times10^{-1}$ | $5.1\times10^{5}$ | $3.8\times10^{-1}$ | $1.5\times10^{1}$ | $7.2\times10^{-1}$ | $5.8\times10^{2}$ | $3.5\times10^{-1}$ |
| A397V | $2.2\times10^{2}$ | **0** | $9.0\times10^{1}$ | $6.3\times10^{-1}$ | $2.3\times10^{6}$ | **$3.2\times10^{-3}$** | $1.4\times10^{6}$ | **$3.0\times10^{-4}$** | $2.8\times10^{1}$ | **$1.\times10^{-4}$** | $5.6\times10^{2}$ | $2.5\times10^{-2}$ |
| M398G | $4.4\times10^{1}$ | **$2.6\times10^{-3}$** | $4.4\times10^{1}$ | **$1.4\times10^{-3}$** | $1.1\times10^{6}$ | $3.3\times10^{-1}$ | $2.6\times10^{5}$ | $1.2\times10^{-2}$ | 4.9 | **$6.0\times10^{-4}$** | $6.6\times10^{2}$ | $6.0\times10^{-2}$ |
| M398V | $9.1\times10^{1}$ | $1.2\times10^{-1}$ | $8.6\times10^{1}$ | $9.9\times10^{-1}$ | $1.2\times10^{6}$ | $4.4\times10^{-1}$ | $3.5\times10^{5}$ | $3.4\times10^{-2}$ | $1.2\times10^{1}$ | $3.8\times10^{-1}$ | $6.2\times10^{2}$ | $8.2\times10^{-1}$ |
| N399G | $1.5\times10^{1}$ | **$1.\times10^{-4}$** | $4.4\times10^{1}$ | **$9.0\times10^{-4}$** | $4.8\times10^{5}$ | **$5.0\times10^{-3}$** | $2.2\times10^{5}$ | **$5.9\times10^{-3}$** | 4.6 | **$2.0\times10^{-4}$** | $5.8\times10^{2}$ | $2.0\times10^{-1}$ |
| N399V | $4.3\times10^{1}$ | **$2.2\times10^{-3}$** | $5.3\times10^{1}$ | **$9.3\times10^{-3}$** | $8.3\times10^{5}$ | $1.2\times10^{-1}$ | $2.0\times10^{4}$ | **$1.\times10^{-4}$** | $\leq 1.0$ | **$6.0\times10^{-4}$** | $5.7\times10^{2}$ | $1.4\times10^{-1}$ |
| Y400G | $2.5\times10^{2}$ | **$1.3\times10^{-3}$** | $7.3\times10^{1}$ | $3.6\times10^{-1}$ | $3.4\times10^{6}$ | **0** | $\leq 4.3\times10^{5}$ | $4.5\times10^{-1}$ | $1.3\times10^{1}$ | $3.2\times10^{-1}$ | $7.2\times10^{2}$ | **$8.1\times10^{-3}$** |
| Y400V | $1.9\times10^{2}$ | **$7.0\times10^{-4}$** | $9.6\times10^{1}$ | $2.6\times10^{-1}$ | $2.2\times10^{6}$ | **$8.0\times10^{-3}$** | $2.0\times10^{5}$ | **$6.0\times10^{-4}$** | 6.6 | $1.1\times10^{-2}$ | $7.9\times10^{2}$ | **$9.0\times10^{-4}$** |
| Q401G | $3.2\times10^{1}$ | **$2.0\times10^{-4}$** | $5.1\times10^{1}$ | **$5.1\times10^{-3}$** | $1.1\times10^{6}$ | $3.1\times10^{-1}$ | $4.3\times10^{5}$ | $1.2\times10^{-1}$ | 8.5 | $1.9\times10^{-2}$ | $5.6\times10^{2}$ | $8.2\times10^{-2}$ |
| Q401V | $6.0\times10^{1}$ | **$6.4\times10^{-3}$** | $8.9\times10^{1}$ | $6.8\times10^{-1}$ | $8.3\times10^{5}$ | $7.9\times10^{-2}$ | $1.2\times10^{5}$ | **$1.1\times10^{-3}$** | $\leq 1.2$ | **$1.\times10^{-4}$** | $7.0\times10^{2}$ | $1.4\times10^{-2}$ |

| Variant | $k_{cat}$ cMUP ($s^{-1}$) | $k_{cat}$ cMUP *p*–value | $K_M$ cMUP (μM) | $K_M$ cMUP *p*–value | $k_{cat}/K_M$ cMUP ($M^{-1}s^{-1}$) | $k_{cat}/K_M$ cMUP *p*–value | $k_{cat}/K_M$ MeP ($M^{-1}s^{-1}$) | $k_{cat}/K_M$ MeP *p*–value | $k_{cat}/K_M$ MecMUP ($M^{-1}s^{-1}$) | $k_{cat}/K_M$ MecMUP *p*–value | $K_i$ $PO_4$ (μM) | $K_i$ $PO_4$ *p*–value |
|---|---|---|---|---|---|---|---|---|---|---|---|---|
| V402A | $1.9\times10^{2}$ | **$2.3\times10^{-3}$** | $8.8\times10^{1}$ | $8.1\times10^{-1}$ | $2.2\times10^{6}$ | $1.3\times10^{-2}$ | $2.3\times10^{5}$ | **$1.0\times10^{-3}$** | $1.4\times10^{1}$ | $6.8\times10^{-1}$ | $6.3\times10^{2}$ | $4.4\times10^{-1}$ |
| V402G | $5.0\times10^{1}$ | **$2.1\times10^{-3}$** | $6.8\times10^{1}$ | $9.9\times10^{-2}$ | $8.5\times10^{5}$ | $8.8\times10^{-2}$ | $1.7\times10^{5}$ | **$2.7\times10^{-3}$** | 9.2 | $1.9\times10^{-2}$ | $5.4\times10^{2}$ | $1.3\times10^{-2}$ |
| Y403G | $1.4\times10^{1}$ | **$9.0\times10^{-4}$** | $4.9\times10^{1}$ | $1.1\times10^{-2}$ | $4.3\times10^{5}$ | $1.6\times10^{-2}$ | $2.4\times10^{5}$ | **$9.3\times10^{-3}$** | ≤ 1.7 | **$5.0\times10^{-4}$** | $4.3\times10^{2}$ | **$1.\times10^{-4}$** |
| Y403V | $1.0\times10^{2}$ | $4.5\times10^{-1}$ | $8.4\times10^{1}$ | $8.4\times10^{-1}$ | $1.3\times10^{6}$ | $8.4\times10^{-1}$ | $1.7\times10^{5}$ | **$3.2\times10^{-3}$** | $1.2\times10^{1}$ | $1.8\times10^{-1}$ | $6.6\times10^{2}$ | $1.4\times10^{-1}$ |
| F404G | $5.4\times10^{1}$ | $1.9\times10^{-2}$ | $7.1\times10^{1}$ | $2.6\times10^{-1}$ | $7.1\times10^{5}$ | $8.1\times10^{-2}$ | $1.1\times10^{5}$ | **$1.3\times10^{-3}$** | 3.9 | **$2.4\times10^{-3}$** | $5.3\times10^{2}$ | $8.2\times10^{-2}$ |
| F404V | $1.9\times10^{2}$ | **$1.6\times10^{-3}$** | $9.0\times10^{1}$ | $5.7\times10^{-1}$ | $2.2\times10^{6}$ | **$6.7\times10^{-3}$** | $2.4\times10^{5}$ | **$7.9\times10^{-3}$** | $1.9\times10^{1}$ | $4.2\times10^{-1}$ | $7.1\times10^{2}$ | $1.4\times10^{-2}$ |
| D405G | $4.6\times10^{1}$ | **$7.5\times10^{-3}$** | $5.7\times10^{1}$ | $4.6\times10^{-2}$ | $8.0\times10^{5}$ | $1.6\times10^{-1}$ | $9.1\times10^{4}$ | **$5.3\times10^{-3}$** | 9.3 | $4.4\times10^{-2}$ | $4.8\times10^{2}$ | **$2.8\times10^{-3}$** |
| D405V | $9.7\times10^{1}$ | $1.8\times10^{-1}$ | $8.6\times10^{1}$ | $9.2\times10^{-1}$ | $1.1\times10^{6}$ | $3.2\times10^{-1}$ | $1.5\times10^{4}$ | **0** | 4.9 | **$2.8\times10^{-3}$** | $6.3\times10^{2}$ | $6.0\times10^{-1}$ |
| R406G | $1.3\times10^{2}$ | $8.1\times10^{-1}$ | $8.6\times10^{1}$ | $9.7\times10^{-1}$ | $1.4\times10^{6}$ | $8.8\times10^{-1}$ | $2.7\times10^{5}$ | $6.2\times10^{-2}$ | $1.4\times10^{1}$ | $6.3\times10^{-1}$ | $7.3\times10^{2}$ | $2.2\times10^{-2}$ |
| R406V | $2.6\times10^{2}$ | **0** | $1.1\times10^{2}$ | $2.8\times10^{-2}$ | $2.1\times10^{6}$ | $4.9\times10^{-2}$ | $4.0\times10^{5}$ | $6.5\times10^{-2}$ | $1.7\times10^{1}$ | $7.3\times10^{-1}$ | $7.3\times10^{2}$ | $2.2\times10^{-2}$ |
| K407G | $1.3\times10^{2}$ | $6.3\times10^{-1}$ | $7.3\times10^{1}$ | $3.4\times10^{-1}$ | $1.6\times10^{6}$ | $6.1\times10^{-1}$ | $8.5\times10^{5}$ | $6.0\times10^{-2}$ | $1.9\times10^{1}$ | $4.5\times10^{-1}$ | $5.8\times10^{2}$ | $1.9\times10^{-1}$ |
| K407V | $1.9\times10^{2}$ | **$3.4\times10^{-3}$** | $9.8\times10^{1}$ | $1.7\times10^{-1}$ | $1.9\times10^{6}$ | $1.5\times10^{-1}$ | $4.4\times10^{5}$ | $8.8\times10^{-2}$ | $1.4\times10^{1}$ | $6.3\times10^{-1}$ | $6.8\times10^{2}$ | $5.6\times10^{-2}$ |
| V408A | $2.1\times10^{2}$ | **$6.0\times10^{-4}$** | $9.7\times10^{1}$ | $2.4\times10^{-1}$ | $2.1\times10^{6}$ | $4.1\times10^{-2}$ | $8.8\times10^{5}$ | $1.2\times10^{-1}$ | $2.2\times10^{1}$ | $1.0\times10^{-1}$ | $6.7\times10^{2}$ | $1.1\times10^{-1}$ |
| V408G | $9.4\times10^{1}$ | $1.7\times10^{-1}$ | $7.8\times10^{1}$ | $4.0\times10^{-1}$ | $1.4\times10^{6}$ | $8.3\times10^{-1}$ | $2.0\times10^{5}$ | $1.5\times10^{-2}$ | $1.4\times10^{1}$ | $4.2\times10^{-1}$ | $6.6\times10^{2}$ | $8.7\times10^{-2}$ |
| L409G | $2.4\times10^{1}$ | **0** | $6.8\times10^{1}$ | $8.0\times10^{-2}$ | $3.9\times10^{5}$ | **$1.0\times10^{-3}$** | $9.2\times10^{4}$ | **$6.0\times10^{-4}$** | 5.1 | **0** | $5.9\times10^{2}$ | $4.2\times10^{-1}$ |
| L409V | $1.3\times10^{2}$ | $7.0\times10^{-1}$ | $9.3\times10^{1}$ | $3.5\times10^{-1}$ | $1.3\times10^{6}$ | $6.4\times10^{-1}$ | $1.3\times10^{5}$ | **$8.9\times10^{-3}$** | 9.6 | $1.1\times10^{-1}$ | $7.2\times10^{2}$ | $1.2\times10^{-2}$ |
| A410G | $1.3\times10^{2}$ | $6.5\times10^{-1}$ | $7.3\times10^{1}$ | $3.4\times10^{-1}$ | $1.8\times10^{6}$ | $2.9\times10^{-1}$ | $8.3\times10^{5}$ | $7.0\times10^{-2}$ | $1.6\times10^{1}$ | $9.5\times10^{-1}$ | $6.0\times10^{2}$ | $6.4\times10^{-1}$ |
| A410V | $1.4\times10^{2}$ | $3.3\times10^{-1}$ | $9.6\times10^{1}$ | $2.6\times10^{-1}$ | $1.7\times10^{6}$ | $3.5\times10^{-1}$ | $3.7\times10^{5}$ | $3.6\times10^{-2}$ | $1.4\times10^{1}$ | $6.7\times10^{-1}$ | $6.5\times10^{2}$ | $2.7\times10^{-1}$ |
| D411G | $1.2\times10^{2}$ | $8.7\times10^{-1}$ | $8.5\times10^{1}$ | $9.8\times10^{-1}$ | $1.4\times10^{6}$ | $9.6\times10^{-1}$ | $4.2\times10^{5}$ | $1.1\times10^{-1}$ | $1.4\times10^{1}$ | $5.0\times10^{-1}$ | $6.5\times10^{2}$ | $2.5\times10^{-1}$ |
| D411V | $1.6\times10^{2}$ | **$4.5\times10^{-3}$** | $9.3\times10^{1}$ | $2.5\times10^{-1}$ | $1.7\times10^{6}$ | $2.0\times10^{-1}$ | $3.2\times10^{5}$ | **$5.0\times10^{-4}$** | $1.3\times10^{1}$ | $1.6\times10^{-1}$ | $6.3\times10^{2}$ | $2.4\times10^{-1}$ |
| S412G | $9.3\times10^{1}$ | $1.6\times10^{-1}$ | $8.0\times10^{1}$ | $5.6\times10^{-1}$ | $1.3\times10^{6}$ | $7.9\times10^{-1}$ | $3.0\times10^{5}$ | $1.8\times10^{-2}$ | $1.3\times10^{1}$ | $2.8\times10^{-1}$ | $5.8\times10^{2}$ | $2.4\times10^{-1}$ |
| S412V | $2.4\times10^{2}$ | **$1.\times10^{-4}$** | $1.1\times10^{2}$ | $1.1\times10^{-2}$ | $1.9\times10^{6}$ | $1.2\times10^{-1}$ | $5.5\times10^{5}$ | $5.5\times10^{-1}$ | $1.8\times10^{1}$ | $6.9\times10^{-1}$ | $6.8\times10^{2}$ | $6.9\times10^{-2}$ |
| K413G | $1.2\times10^{2}$ | $9.4\times10^{-1}$ | $8.3\times10^{1}$ | $7.6\times10^{-1}$ | $1.6\times10^{6}$ | $6.9\times10^{-1}$ | $3.9\times10^{5}$ | $1.8\times10^{-1}$ | $2.4\times10^{1}$ | **$5.1\times10^{-3}$** | $5.8\times10^{2}$ | $1.4\times10^{-1}$ |
| K413V | $2.0\times10^{2}$ | **$7.0\times10^{-4}$** | $8.8\times10^{1}$ | $7.7\times10^{-1}$ | $2.1\times10^{6}$ | $3.0\times10^{-2}$ | $3.1\times10^{5}$ | **$9.6\times10^{-3}$** | $1.6\times10^{1}$ | $8.2\times10^{-1}$ | $6.5\times10^{2}$ | $2.1\times10^{-1}$ |
| L414G | $3.3\times10^{1}$ | **$6.0\times10^{-4}$** | $7.4\times10^{1}$ | $3.4\times10^{-1}$ | $5.4\times10^{5}$ | $1.8\times10^{-2}$ | $9.1\times10^{4}$ | **$1.5\times10^{-3}$** | 6.5 | **$2.1\times10^{-3}$** | $6.0\times10^{2}$ | $5.3\times10^{-1}$ |
| L414V | $9.4\times10^{1}$ | $2.0\times10^{-1}$ | $8.8\times10^{1}$ | $7.9\times10^{-1}$ | $9.2\times10^{5}$ | $1.8\times10^{-1}$ | $2.7\times10^{5}$ | **$3.7\times10^{-3}$** | $1.2\times10^{1}$ | $1.7\times10^{-1}$ | $6.4\times10^{2}$ | $5.5\times10^{-1}$ |
| E415G | $8.1\times10^{1}$ | $7.6\times10^{-2}$ | $7.3\times10^{1}$ | $2.9\times10^{-1}$ | $6.9\times10^{5}$ | $6.4\times10^{-2}$ | $2.0\times10^{5}$ | **$5.0\times10^{-3}$** | $1.7\times10^{1}$ | $6.4\times10^{-1}$ | $5.9\times10^{2}$ | $3.6\times10^{-1}$ |
| E415V | $1.2\times10^{2}$ | $9.7\times10^{-1}$ | $8.5\times10^{1}$ | $9.7\times10^{-1}$ | $1.3\times10^{6}$ | $7.4\times10^{-1}$ | $6.2\times10^{5}$ | $9.6\times10^{-1}$ | 9.9 | $5.8\times10^{-2}$ | $6.2\times10^{2}$ | $5.8\times10^{-1}$ |
| L416G | $3.2\times10^{1}$ | $1.0\times10^{-2}$ | $8.1\times10^{1}$ | $7.8\times10^{-1}$ | $3.9\times10^{5}$ | $4.1\times10^{-2}$ | $5.7\times10^{5}$ | $7.1\times10^{-1}$ | ≤ $9.9\times10^{-1}$ | **0** | $6.6\times10^{2}$ | $1.8\times10^{-1}$ |
| L416V | $5.0\times10^{1}$ | **$4.1\times10^{-3}$** | $9.6\times10^{1}$ | $3.6\times10^{-1}$ | $5.1\times10^{5}$ | $1.2\times10^{-2}$ | $9.9\times10^{4}$ | **0** | 7.3 | $2.0\times10^{-2}$ | $6.7\times10^{2}$ | $1.1\times10^{-1}$ |
| D417G | $7.2\times10^{1}$ | $3.6\times10^{-2}$ | $7.1\times10^{1}$ | $2.2\times10^{-1}$ | $1.1\times10^{6}$ | $4.8\times10^{-1}$ | $3.1\times10^{5}$ | $2.6\times10^{-2}$ | $1.2\times10^{1}$ | $2.6\times10^{-1}$ | $6.4\times10^{2}$ | $5.8\times10^{-1}$ |
| D417V | $1.2\times10^{2}$ | $7.6\times10^{-1}$ | $1.0\times10^{2}$ | $1.5\times10^{-1}$ | $1.2\times10^{6}$ | $6.1\times10^{-1}$ | $2.4\times10^{5}$ | **$2.7\times10^{-3}$** | $1.1\times10^{1}$ | $1.6\times10^{-1}$ | $7.1\times10^{2}$ | $2.1\times10^{-2}$ |
| D418G | $9.2\times10^{1}$ | $2.5\times10^{-1}$ | $7.8\times10^{1}$ | $6.3\times10^{-1}$ | $1.3\times10^{6}$ | $7.7\times10^{-1}$ | $2.6\times10^{5}$ | $1.4\times10^{-2}$ | $1.7\times10^{1}$ | $8.5\times10^{-1}$ | $6.7\times10^{2}$ | $1.6\times10^{-1}$ |
| D418V | $2.1\times10^{2}$ | **$1.0\times10^{-3}$** | $1.1\times10^{2}$ | $7.5\times10^{-2}$ | $2.2\times10^{6}$ | $4.1\times10^{-2}$ | $3.1\times10^{5}$ | $1.3\times10^{-2}$ | $2.2\times10^{1}$ | $1.5\times10^{-1}$ | $6.7\times10^{2}$ | $1.2\times10^{-1}$ |
| V419A | $1.4\times10^{2}$ | $3.1\times10^{-1}$ | $9.3\times10^{1}$ | $3.7\times10^{-1}$ | $1.5\times10^{6}$ | $7.8\times10^{-1}$ | $3.3\times10^{5}$ | $1.9\times10^{-2}$ | $1.4\times10^{1}$ | $6.6\times10^{-1}$ | $6.5\times10^{2}$ | $1.6\times10^{-1}$ |
| V419G | $2.7\times10^{1}$ | **$4.7\times10^{-3}$** | $5.4\times10^{1}$ | $3.4\times10^{-2}$ | $5.3\times10^{5}$ | $3.6\times10^{-2}$ | $7.3\times10^{4}$ | **$8.0\times10^{-4}$** | 6.4 | **$2.3\times10^{-3}$** | $4.8\times10^{2}$ | **$8.5\times10^{-3}$** |
| R420G | $2.7\times10^{1}$ | **$3.0\times10^{-4}$** | $7.2\times10^{1}$ | $1.9\times10^{-1}$ | $4.7\times10^{5}$ | **$4.1\times10^{-3}$** | $9.8\times10^{4}$ | **$4.9\times10^{-3}$** | 4.9 | **$2.0\times10^{-4}$** | $6.0\times10^{2}$ | $4.1\times10^{-1}$ |
| R420V | $9.1\times10^{1}$ | $8.8\times10^{-2}$ | $9.1\times10^{1}$ | $5.5\times10^{-1}$ | $9.2\times10^{5}$ | $7.5\times10^{-2}$ | $2.2\times10^{5}$ | **$1.4\times10^{-3}$** | ≤ 4.1 | **$2.0\times10^{-3}$** | $7.4\times10^{2}$ | **$1.0\times10^{-3}$** |
| D421G | $1.2\times10^{2}$ | $9.0\times10^{-1}$ | $8.9\times10^{1}$ | $6.7\times10^{-1}$ | $1.4\times10^{6}$ | $8.3\times10^{-1}$ | $3.3\times10^{5}$ | $2.7\times10^{-2}$ | $1.5\times10^{1}$ | $7.9\times10^{-1}$ | $6.2\times10^{2}$ | $7.9\times10^{-1}$ |
| D421V | $7.9\times10^{1}$ | $4.2\times10^{-2}$ | $7.9\times10^{1}$ | $5.0\times10^{-1}$ | $9.6\times10^{5}$ | $1.6\times10^{-1}$ | $2.1\times10^{5}$ | **$4.3\times10^{-3}$** | 9.1 | $7.3\times10^{-2}$ | $5.9\times10^{2}$ | $5.1\times10^{-1}$ |
| Y422G | $4.0\times10^{1}$ | **$2.1\times10^{-3}$** | $6.9\times10^{1}$ | $1.6\times10^{-1}$ | $8.7\times10^{5}$ | $1.4\times10^{-1}$ | $1.2\times10^{5}$ | **$1.2\times10^{-3}$** | 9.9 | $7.3\times10^{-2}$ | $5.9\times10^{2}$ | $3.3\times10^{-1}$ |
| Y422V | $1.6\times10^{2}$ | $1.3\times10^{-1}$ | $9.5\times10^{1}$ | $3.9\times10^{-1}$ | $1.6\times10^{6}$ | $7.1\times10^{-1}$ | $5.0\times10^{5}$ | $3.2\times10^{-1}$ | $1.9\times10^{1}$ | $5.5\times10^{-1}$ | $6.5\times10^{2}$ | $3.8\times10^{-1}$ |
| V423A | $1.3\times10^{2}$ | $4.6\times10^{-1}$ | $8.8\times10^{1}$ | $7.8\times10^{-1}$ | $1.6\times10^{6}$ | $6.9\times10^{-1}$ | $3.7\times10^{5}$ | $5.7\times10^{-2}$ | $1.2\times10^{1}$ | $4.4\times10^{-1}$ | $6.3\times10^{2}$ | $4.6\times10^{-1}$ |
| V423G | $4.1\times10^{1}$ | **$1.7\times10^{-3}$** | $6.3\times10^{1}$ | $5.5\times10^{-2}$ | $6.3\times10^{5}$ | $3.1\times10^{-2}$ | $1.5\times10^{5}$ | **$7.6\times10^{-3}$** | 8.9 | $1.2\times10^{-2}$ | $5.7\times10^{2}$ | $1.9\times10^{-1}$ |
| M424G | $2.3\times10^{1}$ | **$2.0\times10^{-4}$** | $5.3\times10^{1}$ | **$9.6\times10^{-3}$** | $4.3\times10^{5}$ | **$7.9\times10^{-3}$** | $2.5\times10^{5}$ | $1.1\times10^{-2}$ | 5.6 | **$1.3\times10^{-3}$** | $4.5\times10^{2}$ | **$2.0\times10^{-4}$** |
| M424V | $1.1\times10^{2}$ | $6.9\times10^{-1}$ | $8.6\times10^{1}$ | $9.0\times10^{-1}$ | $1.2\times10^{6}$ | $4.6\times10^{-1}$ | $7.9\times10^{5}$ | $9.5\times10^{-2}$ | $1.2\times10^{1}$ | $3.9\times10^{-1}$ | $6.0\times10^{2}$ | $5.5\times10^{-1}$ |
| T425V | $1.5\times10^{2}$ | $2.3\times10^{-1}$ | $8.0\times10^{1}$ | $6.7\times10^{-1}$ | $1.3\times10^{6}$ | $6.9\times10^{-1}$ | $5.4\times10^{5}$ | $5.7\times10^{-1}$ | $1.2\times10^{1}$ | $3.6\times10^{-1}$ | $6.3\times10^{2}$ | $5.4\times10^{-1}$ |
| E426G | $9.5\times10^{1}$ | $2.5\times10^{-1}$ | $7.6\times10^{1}$ | $3.8\times10^{-1}$ | $1.3\times10^{6}$ | $6.7\times10^{-1}$ | $2.1\times10^{5}$ | **$4.9\times10^{-3}$** | 8.3 | $1.6\times10^{-2}$ | $5.3\times10^{2}$ | **$6.2\times10^{-3}$** |
| E426V | $1.6\times10^{2}$ | $5.5\times10^{-2}$ | $9.5\times10^{1}$ | $3.5\times10^{-1}$ | $1.8\times10^{6}$ | $2.8\times10^{-1}$ | $3.0\times10^{5}$ | **$7.4\times10^{-3}$** | $1.3\times10^{1}$ | $4.3\times10^{-1}$ | $6.5\times10^{2}$ | $3.2\times10^{-1}$ |
| L427G | $1.4\times10^{1}$ | **$2.0\times10^{-4}$** | $6.1\times10^{1}$ | $4.5\times10^{-2}$ | $2.5\times10^{5}$ | **$2.4\times10^{-3}$** | ≤ $9.2\times10^{4}$ | **$1.2\times10^{-3}$** | 4.0 | **$4.0\times10^{-4}$** | $5.0\times10^{2}$ | **$2.4\times10^{-3}$** |
| L427V | $1.2\times10^{2}$ | $9.6\times10^{-1}$ | $8.8\times10^{1}$ | $8.0\times10^{-1}$ | $1.4\times10^{6}$ | $9.2\times10^{-1}$ | $2.2\times10^{5}$ | **$9.0\times10^{-4}$** | $1.1\times10^{1}$ | $2.0\times10^{-1}$ | $6.6\times10^{2}$ | $1.5\times10^{-1}$ |
| K428G | $6.5\times10^{1}$ | $2.6\times10^{-2}$ | $6.8\times10^{1}$ | $1.4\times10^{-1}$ | $5.7\times10^{5}$ | $1.9\times10^{-2}$ | $2.6\times10^{5}$ | $2.7\times10^{-2}$ | $1.2\times10^{1}$ | $3.4\times10^{-1}$ | $6.3\times10^{2}$ | $6.6\times10^{-1}$ |
| K428V | $7.3\times10^{1}$ | **$7.6\times10^{-3}$** | $7.9\times10^{1}$ | $4.5\times10^{-1}$ | $8.7\times10^{5}$ | $3.6\times10^{-2}$ | $1.5\times10^{5}$ | **0** | $1.0\times10^{1}$ | $4.6\times10^{-2}$ | $5.6\times10^{2}$ | $3.3\times10^{-2}$ |
| K429G | $1.3\times10^{2}$ | $4.4\times10^{-1}$ | $8.7\times10^{1}$ | $8.4\times10^{-1}$ | $1.4\times10^{6}$ | $9.9\times10^{-1}$ | $4.3\times10^{5}$ | $1.8\times10^{-1}$ | $1.5\times10^{1}$ | $7.8\times10^{-1}$ | $5.7\times10^{2}$ | $1.2\times10^{-1}$ |
| K429V | $1.8\times10^{2}$ | **$4.8\times10^{-3}$** | $9.4\times10^{1}$ | $3.3\times10^{-1}$ | $2.0\times10^{6}$ | $8.3\times10^{-2}$ | $2.1\times10^{5}$ | **$5.6\times10^{-3}$** | $1.7\times10^{1}$ | $7.7\times10^{-1}$ | $7.1\times10^{2}$ | $2.7\times10^{-2}$ |
| E430G | $7.4\times10^{1}$ | $1.2\times10^{-2}$ | $9.2\times10^{1}$ | $4.6\times10^{-1}$ | $9.3\times10^{5}$ | $7.6\times10^{-2}$ | $1.2\times10^{5}$ | **$7.0\times10^{-4}$** | 6.7 | **$1.\times10^{-4}$** | $7.8\times10^{2}$ | **0** |
| E430V | $1.2\times10^{2}$ | $9.3\times10^{-1}$ | $1.0\times10^{2}$ | $2.5\times10^{-1}$ | $1.4\times10^{6}$ | $8.5\times10^{-1}$ | $4.4\times10^{5}$ | $1.1\times10^{-1}$ | $1.3\times10^{1}$ | $3.7\times10^{-1}$ | $7.7\times10^{2}$ | **$5.0\times10^{-4}$** |
| P431G | $7.1\times10^{1}$ | $4.2\times10^{-2}$ | $6.8\times10^{1}$ | $1.4\times10^{-1}$ | $1.1\times10^{6}$ | $4.2\times10^{-1}$ | $1.9\times10^{5}$ | **$4.0\times10^{-3}$** | 9.8 | $3.8\times10^{-2}$ | $6.3\times10^{2}$ | $4.9\times10^{-1}$ |
| P431V | $2.0\times10^{2}$ | **$1.6\times10^{-3}$** | $8.4\times10^{1}$ | $9.3\times10^{-1}$ | $2.1\times10^{6}$ | $4.9\times10^{-2}$ | $2.0\times10^{5}$ | $1.6\times10^{-2}$ | $2.0\times10^{1}$ | $3.4\times10^{-1}$ | $6.7\times10^{2}$ | $7.2\times10^{-2}$ |
| S432G | $9.4\times10^{1}$ | $1.7\times10^{-1}$ | $9.8\times10^{1}$ | $2.7\times10^{-1}$ | $1.2\times10^{6}$ | $4.7\times10^{-1}$ | $4.7\times10^{5}$ | $1.8\times10^{-1}$ | 8.2 | $1.8\times10^{-2}$ | $7.1\times10^{2}$ | $1.0\times10^{-2}$ |
| S432V | $3.5\times10^{1}$ | **0** | $9.4\times10^{1}$ | $3.2\times10^{-1}$ | $3.7\times10^{5}$ | **$1.4\times10^{-3}$** | $1.6\times10^{5}$ | **0** | ≤ 2.7 | **$1.\times10^{-4}$** | $7.0\times10^{2}$ | **$4.3\times10^{-3}$** |
| V433A | $1.1\times10^{2}$ | $6.9\times10^{-1}$ | $1.1\times10^{2}$ | $8.1\times10^{-2}$ | $1.1\times10^{6}$ | $3.4\times10^{-1}$ | $3.1\times10^{5}$ | $2.3\times10^{-2}$ | 9.2 | $8.3\times10^{-2}$ | $7.6\times10^{2}$ | **$4.6\times10^{-3}$** |
| V433G | $1.3\times10^{1}$ | **0** | $8.2\times10^{1}$ | $5.6\times10^{-1}$ | $1.6\times10^{5}$ | **0** | $3.4\times10^{4}$ | **$2.0\times10^{-4}$** | 1.1 | **$1.\times10^{-4}$** | $7.1\times10^{2}$ | **$4.0\times10^{-4}$** |
| L434G | $1.4\times10^{1}$ | **0** | $4.0\times10^{1}$ | **0** | $3.5\times10^{5}$ | **0** | $3.5\times10^{4}$ | **0** | 3.8 | **0** | $3.4\times10^{2}$ | **0** |
| L434V | $4.9\times10^{1}$ | **$1.2\times10^{-3}$** | $8.0\times10^{1}$ | $5.0\times10^{-1}$ | $6.7\times10^{5}$ | $1.8\times10^{-2}$ | $1.6\times10^{4}$ | **0** | 4.8 | **$4.0\times10^{-4}$** | $5.9\times10^{2}$ | $3.3\times10^{-1}$ |
| Y435G | 4.0 | **$4.0\times10^{-4}$** | $6.3\times10^{1}$ | $1.0\times10^{-1}$ | $5.4\times10^{4}$ | **$2.1\times10^{-3}$** | $5.0\times10^{3}$ | **$9.0\times10^{-4}$** | ≤ 1.2 | **0** | $3.5\times10^{2}$ | **0** |
| Y435V | $1.4\times10^{1}$ | **0** | $7.2\times10^{1}$ | $1.0\times10^{-1}$ | $1.9\times10^{5}$ | **$1.\times10^{-4}$** | $1.5\times10^{4}$ | **$5.0\times10^{-4}$** | 1.6 | **$7.0\times10^{-4}$** | $4.9\times10^{2}$ | **$3.0\times10^{-4}$** |
| V436A | $1.4\times10^{2}$ | $3.2\times10^{-1}$ | $1.1\times10^{2}$ | $2.0\times10^{-2}$ | $1.3\times10^{6}$ | $7.6\times10^{-1}$ | $4.3\times10^{5}$ | $1.1\times10^{-1}$ | $1.2\times10^{1}$ | $3.4\times10^{-1}$ | $6.8\times10^{2}$ | $1.5\times10^{-1}$ |
| V436G | $2.3\times10^{1}$ | **$4.0\times10^{-4}$** | $5.4\times10^{1}$ | $1.2\times10^{-2}$ | $4.5\times10^{5}$ | $1.1\times10^{-2}$ | $1.1\times10^{5}$ | **$6.1\times10^{-3}$** | 6.0 | **$2.0\times10^{-3}$** | $4.7\times10^{2}$ | **$1.2\times10^{-3}$** |
| L437G | $2.3\times10^{1}$ | **0** | $6.8\times10^{1}$ | $7.3\times10^{-2}$ | $4.1\times10^{5}$ | **$1.1\times10^{-3}$** | $1.0\times10^{5}$ | **$1.7\times10^{-3}$** | 5.0 | **$1.0\times10^{-3}$** | $6.3\times10^{2}$ | $6.6\times10^{-1}$ |
| L437V | $2.1\times10^{2}$ | **$1.\times10^{-4}$** | $9.0\times10^{1}$ | $5.5\times10^{-1}$ | $2.1\times10^{6}$ | $2.7\times10^{-2}$ | $8.8\times10^{5}$ | **$5.5\times10^{-3}$** | $1.7\times10^{1}$ | $8.7\times10^{-1}$ | $6.6\times10^{2}$ | $4.7\times10^{-2}$ |
| S438G | $5.0\times10^{1}$ | **$2.6\times10^{-3}$** | $6.9\times10^{1}$ | $1.3\times10^{-1}$ | $7.6\times10^{5}$ | $4.4\times10^{-2}$ | $2.4\times10^{5}$ | **$7.1\times10^{-3}$** | $1.0\times10^{1}$ | $1.0\times10^{-1}$ | $6.4\times10^{2}$ | $2.3\times10^{-1}$ |
| S438V | $9.1\times10^{1}$ | $1.8\times10^{-1}$ | $9.0\times10^{1}$ | $5.4\times10^{-1}$ | $1.1\times10^{6}$ | $3.8\times10^{-1}$ | $1.8\times10^{5}$ | **$3.6\times10^{-3}$** | 8.2 | $1.8\times10^{-2}$ | $6.4\times10^{2}$ | $5.3\times10^{-1}$ |
| T439G | $2.0\times10^{1}$ | **0** | $5.5\times10^{1}$ | **$2.4\times10^{-3}$** | $3.7\times10^{5}$ | **$7.0\times10^{-4}$** | $9.5\times10^{4}$ | **0** | 2.3 | **$9.8\times10^{-3}$** | $5.1\times10^{2}$ | **$1.4\times10^{-3}$** |
| T439V | $1.6\times10^{2}$ | $3.7\times10^{-2}$ | $9.4\times10^{1}$ | $3.5\times10^{-1}$ | $2.0\times10^{6}$ | $9.7\times10^{-2}$ | $2.7\times10^{5}$ | $4.4\times10^{-2}$ | $1.2\times10^{1}$ | $2.7\times10^{-1}$ | $7.5\times10^{2}$ | **$2.9\times10^{-3}$** |

| Variant | $k_{cat}$ cMUP ($s^{-1}$) | $k_{cat}$ cMUP $p$–value | $K_M$ cMUP (μM) | $K_M$ cMUP $p$–value | $k_{cat}/K_M$ cMUP ($M^{-1}s^{-1}$) | $k_{cat}/K_M$ cMUP $p$–value | $k_{cat}/K_M$ MeP ($M^{-1}s^{-1}$) | $k_{cat}/K_M$ MeP $p$–value | $k_{cat}/K_M$ MecMUP ($M^{-1}s^{-1}$) | $k_{cat}/K_M$ MecMUP $p$–value | $K_i$ $PO_4$ (μM) | $K_i$ $PO_4$ $p$–value |
|---|---|---|---|---|---|---|---|---|---|---|---|---|
| D440G | $1.1\times10^{2}$ | $4.5\times10^{-1}$ | $7.7\times10^{1}$ | $6.0\times10^{-1}$ | $1.6\times10^{6}$ | $5.1\times10^{-1}$ | $3.1\times10^{5}$ | $2.1\times10^{-2}$ | $1.0\times10^{1}$ | $4.5\times10^{-2}$ | $5.4\times10^{2}$ | $1.7\times10^{-2}$ |
| D440V | $2.2\times10^{1}$ | **$2.0\times10^{-4}$** | $8.9\times10^{1}$ | $7.5\times10^{-1}$ | $2.9\times10^{5}$ | **$2.1\times10^{-3}$** | $1.1\times10^{5}$ | **$6.6\times10^{-3}$** | 4.8 | $3.4\times10^{-2}$ | $6.2\times10^{2}$ | $8.3\times10^{-1}$ |
| E441G | $8.4\times10^{1}$ | $1.1\times10^{-1}$ | $5.9\times10^{1}$ | $2.5\times10^{-2}$ | $1.2\times10^{6}$ | $6.1\times10^{-1}$ | $3.4\times10^{5}$ | $3.3\times10^{-2}$ | $1.6\times10^{1}$ | $9.3\times10^{-1}$ | $5.3\times10^{2}$ | $2.5\times10^{-2}$ |
| E441V | $1.1\times10^{2}$ | $5.4\times10^{-1}$ | $9.9\times10^{1}$ | $1.1\times10^{-1}$ | $1.4\times10^{6}$ | $7.2\times10^{-1}$ | $4.0\times10^{5}$ | $2.2\times10^{-2}$ | $1.1\times10^{1}$ | $8.6\times10^{-2}$ | $6.8\times10^{2}$ | $2.6\times10^{-2}$ |
| I442G | 7.0 | **0** | $6.6\times10^{1}$ | $4.8\times10^{-2}$ | $1.6\times10^{5}$ | **0** | $4.6\times10^{4}$ | **$1.1\times10^{-3}$** | $\leq 9.9\times10^{-1}$ | **0** | $4.8\times10^{2}$ | **0** |
| I442V | $1.0\times10^{2}$ | $3.9\times10^{-1}$ | $9.4\times10^{1}$ | $3.4\times10^{-1}$ | $1.1\times10^{6}$ | $4.7\times10^{-1}$ | $3.3\times10^{5}$ | $7.7\times10^{-2}$ | $\leq 6.5$ | $1.1\times10^{-1}$ | $6.5\times10^{2}$ | $4.6\times10^{-1}$ |
| W443G | $1.3\times10^{2}$ | $7.9\times10^{-1}$ | $6.9\times10^{1}$ | $2.1\times10^{-1}$ | $1.5\times10^{6}$ | $8.9\times10^{-1}$ | $5.1\times10^{5}$ | $4.8\times10^{-1}$ | $2.1\times10^{1}$ | $2.2\times10^{-1}$ | $6.6\times10^{2}$ | $1.8\times10^{-1}$ |
| W443V | $2.1\times10^{2}$ | **$4.0\times10^{-4}$** | $9.8\times10^{1}$ | $2.1\times10^{-1}$ | $2.0\times10^{6}$ | $8.4\times10^{-2}$ | $3.3\times10^{5}$ | $3.2\times10^{-2}$ | $1.5\times10^{1}$ | $6.8\times10^{-1}$ | $6.7\times10^{2}$ | $1.3\times10^{-1}$ |
| E444G | $1.4\times10^{2}$ | $4.3\times10^{-1}$ | $8.1\times10^{1}$ | $6.7\times10^{-1}$ | $1.5\times10^{6}$ | $9.3\times10^{-1}$ | $4.3\times10^{5}$ | $1.8\times10^{-1}$ | $2.0\times10^{1}$ | $3.0\times10^{-1}$ | $7.4\times10^{2}$ | **$7.0\times10^{-4}$** |
| E444V | $1.5\times10^{2}$ | $2.3\times10^{-1}$ | $9.7\times10^{1}$ | $2.5\times10^{-1}$ | $1.5\times10^{6}$ | $9.3\times10^{-1}$ | $3.9\times10^{5}$ | $6.0\times10^{-2}$ | 7.6 | $3.2\times10^{-2}$ | $7.4\times10^{2}$ | **$2.1\times10^{-3}$** |
| S445G | $1.8\times10^{1}$ | **0** | $8.0\times10^{1}$ | $6.2\times10^{-1}$ | $2.3\times10^{5}$ | **$2.2\times10^{-3}$** | $9.9\times10^{4}$ | **$6.1\times10^{-3}$** | 3.7 | **$1.\times10^{-4}$** | $6.2\times10^{2}$ | $8.2\times10^{-1}$ |
| S445V | $1.3\times10^{2}$ | $7.1\times10^{-1}$ | $9.5\times10^{1}$ | $4.6\times10^{-1}$ | $1.5\times10^{6}$ | $9.1\times10^{-1}$ | $3.5\times10^{5}$ | $1.4\times10^{-1}$ | $1.4\times10^{1}$ | $5.8\times10^{-1}$ | $6.4\times10^{2}$ | $4.5\times10^{-1}$ |
| S446G | $3.4\times10^{1}$ | **0** | $6.2\times10^{1}$ | $3.1\times10^{-2}$ | $5.1\times10^{5}$ | **$6.9\times10^{-3}$** | $1.5\times10^{5}$ | **$3.1\times10^{-3}$** | 4.5 | **$3.6\times10^{-3}$** | $4.6\times10^{2}$ | **0** |
| S446V | $2.0\times10^{2}$ | **$1.\times10^{-4}$** | $8.2\times10^{1}$ | $6.6\times10^{-1}$ | $2.1\times10^{6}$ | $1.9\times10^{-2}$ | $4.5\times10^{5}$ | $1.8\times10^{-1}$ | $2.2\times10^{1}$ | $2.0\times10^{-1}$ | $6.0\times10^{2}$ | $5.4\times10^{-1}$ |
| I447G | $7.0\times10^{-1}$ | **0** | $4.7\times10^{1}$ | **$9.0\times10^{-4}$** | $1.1\times10^{4}$ | **$1.\times10^{-4}$** | $\leq 1.6\times10^{3}$ | **0** | $\leq 1.1$ | **0** | $3.4\times10^{2}$ | **0** |
| I447V | $1.2\times10^{2}$ | $9.9\times10^{-1}$ | $9.0\times10^{1}$ | $6.2\times10^{-1}$ | $1.5\times10^{6}$ | $7.7\times10^{-1}$ | $5.9\times10^{5}$ | $9.4\times10^{-1}$ | $1.1\times10^{1}$ | $3.2\times10^{-1}$ | $6.8\times10^{2}$ | $1.4\times10^{-1}$ |
| P448G | 5.1 | **0** | $5.6\times10^{1}$ | **$5.0\times10^{-4}$** | $7.6\times10^{4}$ | **0** | $1.6\times10^{4}$ | **$1.\times10^{-4}$** | $\leq 1.1$ | **0** | $4.8\times10^{2}$ | **0** |
| E449G | $6.9\times10^{1}$ | $2.3\times10^{-2}$ | $6.6\times10^{1}$ | $7.6\times10^{-2}$ | $1.1\times10^{6}$ | $2.9\times10^{-1}$ | $3.3\times10^{5}$ | $3.1\times10^{-2}$ | $1.2\times10^{1}$ | $2.4\times10^{-1}$ | $3.9\times10^{2}$ | **$1.\times10^{-4}$** |
| E449V | $8.2\times10^{1}$ | $3.3\times10^{-2}$ | $8.5\times10^{1}$ | $9.7\times10^{-1}$ | $1.1\times10^{6}$ | $2.8\times10^{-1}$ | $1.7\times10^{5}$ | **$7.8\times10^{-3}$** | $1.4\times10^{1}$ | $6.3\times10^{-1}$ | $5.1\times10^{2}$ | **$5.0\times10^{-3}$** |
| P450G | $1.1\times10^{2}$ | $6.0\times10^{-1}$ | $6.9\times10^{1}$ | $1.6\times10^{-1}$ | $1.9\times10^{6}$ | $1.9\times10^{-1}$ | $3.7\times10^{5}$ | $6.7\times10^{-2}$ | $1.6\times10^{1}$ | $9.6\times10^{-1}$ | $6.4\times10^{2}$ | $4.7\times10^{-1}$ |
| P450V | $1.6\times10^{2}$ | $1.2\times10^{-1}$ | $8.2\times10^{1}$ | $7.2\times10^{-1}$ | $1.9\times10^{6}$ | $2.4\times10^{-1}$ | $6.4\times10^{5}$ | $7.6\times10^{-1}$ | $1.5\times10^{1}$ | $7.2\times10^{-1}$ | $5.6\times10^{2}$ | $8.7\times10^{-2}$ |
| I451G | 5.5 | **0** | $4.6\times10^{1}$ | **0** | $1.2\times10^{5}$ | **0** | $1.1\times10^{4}$ | **$3.1\times10^{-3}$** | 3.0 | **$3.0\times10^{-4}$** | $2.8\times10^{2}$ | **0** |
| I451V | $8.3\times10^{1}$ | $3.5\times10^{-2}$ | $9.0\times10^{1}$ | $5.9\times10^{-1}$ | $1.0\times10^{6}$ | $1.6\times10^{-1}$ | $4.7\times10^{5}$ | $8.5\times10^{-2}$ | 8.5 | $1.1\times10^{-2}$ | $5.6\times10^{2}$ | $5.6\times10^{-2}$ |
| K452G | $1.3\times10^{1}$ | **0** | $6.0\times10^{1}$ | $2.0\times10^{-2}$ | $2.0\times10^{5}$ | **$5.0\times10^{-4}$** | $3.1\times10^{4}$ | **0** | 1.9 | **$2.0\times10^{-4}$** | $4.7\times10^{2}$ | **0** |
| K452V | $1.2\times10^{1}$ | **0** | $8.2\times10^{1}$ | $7.0\times10^{-1}$ | $1.5\times10^{5}$ | **$5.0\times10^{-4}$** | $2.5\times10^{4}$ | **$1.\times10^{-4}$** | $\leq 9.3\times10^{-1}$ | **$6.0\times10^{-4}$** | $5.0\times10^{2}$ | **$6.6\times10^{-3}$** |
| S453G | $8.9\times10^{1}$ | $1.4\times10^{-1}$ | $6.3\times10^{1}$ | $4.3\times10^{-2}$ | $1.5\times10^{6}$ | $9.1\times10^{-1}$ | $7.3\times10^{5}$ | $3.2\times10^{-1}$ | $1.3\times10^{1}$ | $4.0\times10^{-1}$ | $4.4\times10^{2}$ | **$1.\times10^{-4}$** |
| S453V | $1.2\times10^{2}$ | $7.4\times10^{-1}$ | $8.0\times10^{1}$ | $6.4\times10^{-1}$ | $1.5\times10^{6}$ | $8.2\times10^{-1}$ | $1.4\times10^{5}$ | **$1.9\times10^{-3}$** | $1.5\times10^{1}$ | $7.7\times10^{-1}$ | $5.6\times10^{2}$ | $1.2\times10^{-1}$ |
| R454G | 5.0 | **0** | $6.5\times10^{1}$ | $1.6\times10^{-2}$ | $8.3\times10^{4}$ | **0** | $6.7\times10^{3}$ | **0** | $\leq 9.9\times10^{-1}$ | **0** | $5.2\times10^{2}$ | **$7.0\times10^{-4}$** |
| R454V | 5.8 | **0** | $6.4\times10^{1}$ | **$3.1\times10^{-3}$** | $8.1\times10^{4}$ | **0** | $1.2\times10^{4}$ | **0** | $\leq 1.3$ | **$1.\times10^{-4}$** | $3.6\times10^{2}$ | **0** |
| V455A | $1.3\times10^{2}$ | $5.6\times10^{-1}$ | $8.2\times10^{1}$ | $7.7\times10^{-1}$ | $1.7\times10^{6}$ | $4.1\times10^{-1}$ | $1.6\times10^{5}$ | **$5.0\times10^{-4}$** | 8.6 | $6.4\times10^{-2}$ | $6.8\times10^{2}$ | $5.3\times10^{-2}$ |
| V455G | $1.3\times10^{1}$ | **0** | $5.9\times10^{1}$ | **$4.1\times10^{-3}$** | $2.0\times10^{5}$ | **$1.\times10^{-4}$** | $2.1\times10^{4}$ | **0** | 1.1 | **$1.1\times10^{-3}$** | $5.2\times10^{2}$ | **$2.0\times10^{-4}$** |
| I456G | $1.2\times10^{1}$ | **0** | $4.7\times10^{1}$ | **0** | $2.4\times10^{5}$ | **0** | $3.4\times10^{4}$ | **0** | 1.6 | **0** | $3.5\times10^{2}$ | **0** |
| I456V | $1.2\times10^{2}$ | $7.0\times10^{-1}$ | $8.9\times10^{1}$ | $5.4\times10^{-1}$ | $1.2\times10^{6}$ | $3.0\times10^{-1}$ | $5.2\times10^{5}$ | $2.2\times10^{-1}$ | 8.8 | $2.6\times10^{-2}$ | $6.4\times10^{2}$ | $2.6\times10^{-1}$ |
| N457G | $3.3\times10^{1}$ | **0** | $6.6\times10^{1}$ | $5.1\times10^{-2}$ | $5.4\times10^{5}$ | **$4.4\times10^{-3}$** | $1.2\times10^{5}$ | **$1.0\times10^{-3}$** | 4.9 | **$2.0\times10^{-4}$** | $5.5\times10^{2}$ | $1.8\times10^{-2}$ |
| N457V | $1.3\times10^{1}$ | **0** | $6.0\times10^{1}$ | **$1.7\times10^{-3}$** | $1.8\times10^{5}$ | **0** | $8.6\times10^{3}$ | **$6.0\times10^{-4}$** | 2.4 | **$1.0\times10^{-3}$** | $4.1\times10^{2}$ | **0** |
| G458A | $1.4\times10^{1}$ | **$5.0\times10^{-4}$** | $5.4\times10^{1}$ | **$9.8\times10^{-3}$** | $2.6\times10^{5}$ | **$2.2\times10^{-3}$** | $1.4\times10^{5}$ | $1.5\times10^{-2}$ | 7.8 | **$3.1\times10^{-3}$** | $3.0\times10^{2}$ | **$1.\times10^{-4}$** |
| G458V | 9.4 | **0** | $2.9\times10^{1}$ | **0** | $3.2\times10^{5}$ | **$1.\times10^{-4}$** | $2.1\times10^{4}$ | **$7.2\times10^{-3}$** | 2.3 | **$5.0\times10^{-4}$** | $2.0\times10^{2}$ | **0** |
| Y459G | 1.4 | **0** | $6.2\times10^{1}$ | $2.2\times10^{-2}$ | $2.2\times10^{4}$ | **0** | $\leq 1.8\times10^{3}$ | **0** | $\leq 1.1$ | **0** | $5.4\times10^{2}$ | $1.7\times10^{-2}$ |
| Y459V | $3.8\times10^{1}$ | **$1.2\times10^{-3}$** | $8.8\times10^{1}$ | $8.3\times10^{-1}$ | $5.6\times10^{5}$ | $2.1\times10^{-2}$ | $9.0\times10^{4}$ | **$1.3\times10^{-3}$** | 3.9 | **$1.4\times10^{-3}$** | $6.0\times10^{2}$ | $6.0\times10^{-1}$ |
| N460V | $3.1\times10^{1}$ | **$5.3\times10^{-3}$** | $9.4\times10^{1}$ | $4.9\times10^{-1}$ | $3.2\times10^{5}$ | $1.1\times10^{-2}$ | $1.4\times10^{5}$ | $2.5\times10^{-2}$ | $\leq 1.9$ | **$3.0\times10^{-4}$** | $5.8\times10^{2}$ | $2.1\times10^{-1}$ |
| W461G | $1.0\times10^{1}$ | **$1.\times10^{-4}$** | $4.4\times10^{1}$ | **$1.7\times10^{-3}$** | $1.8\times10^{5}$ | **$1.4\times10^{-3}$** | $1.3\times10^{5}$ | **$1.7\times10^{-3}$** | 5.0 | **$9.0\times10^{-4}$** | $4.2\times10^{2}$ | **$6.0\times10^{-4}$** |
| W461V | $1.1\times10^{1}$ | **$2.9\times10^{-3}$** | $6.4\times10^{1}$ | $1.4\times10^{-1}$ | $1.7\times10^{5}$ | **$5.5\times10^{-3}$** | $1.3\times10^{5}$ | **0** | 3.8 | $2.2\times10^{-2}$ | $4.4\times10^{2}$ | **0** |
| K462G | $1.1\times10^{2}$ | $6.3\times10^{-1}$ | $8.0\times10^{1}$ | $5.9\times10^{-1}$ | $1.8\times10^{6}$ | $2.2\times10^{-1}$ | $3.1\times10^{5}$ | $2.8\times10^{-2}$ | $2.1\times10^{1}$ | $1.8\times10^{-1}$ | $6.4\times10^{2}$ | $2.4\times10^{-1}$ |
| K462V | $7.2\times10^{1}$ | $9.3\times10^{-2}$ | $7.8\times10^{1}$ | $5.8\times10^{-1}$ | $9.2\times10^{5}$ | $2.3\times10^{-1}$ | $4.4\times10^{5}$ | $1.0\times10^{-1}$ | 8.1 | $4.5\times10^{-2}$ | $6.2\times10^{2}$ | $8.6\times10^{-1}$ |
| R463G | 6.1 | **0** | $3.2\times10^{1}$ | **0** | $2.5\times10^{5}$ | **0** | $2.6\times10^{4}$ | **0** | 1.5 | **$1.\times10^{-4}$** | $6.7\times10^{2}$ | $9.9\times10^{-2}$ |
| R463V | $3.5\times10^{1}$ | **0** | $8.4\times10^{1}$ | $8.5\times10^{-1}$ | $4.5\times10^{5}$ | **$1.7\times10^{-3}$** | $1.2\times10^{5}$ | **$1.2\times10^{-3}$** | 3.3 | **$9.0\times10^{-4}$** | $5.2\times10^{2}$ | **$3.8\times10^{-3}$** |
| S464G | $7.9\times10^{1}$ | $2.0\times10^{-1}$ | $5.0\times10^{1}$ | $3.5\times10^{-2}$ | $1.6\times10^{6}$ | $7.7\times10^{-1}$ | $5.7\times10^{5}$ | $7.4\times10^{-1}$ | $1.4\times10^{1}$ | $6.7\times10^{-1}$ | $4.5\times10^{2}$ | **$1.7\times10^{-3}$** |
| S464V | – | – | – | – | $\leq 3.0\times10^{4}$ | **0** | $\leq 1.6\times10^{4}$ | **0** | $\leq 2.2$ | **$1.8\times10^{-3}$** | – | – |
| G465A | $1.2\times10^{1}$ | **0** | $3.7\times10^{1}$ | **$3.0\times10^{-4}$** | $2.4\times10^{5}$ | **$1.5\times10^{-3}$** | $8.8\times10^{4}$ | **0** | 2.9 | **$1.7\times10^{-3}$** | $4.1\times10^{2}$ | **0** |
| G465V | $7.2\times10^{-2}$ | **$1.\times10^{-4}$** | 8.9 | **0** | $6.4\times10^{3}$ | **$6.0\times10^{-4}$** | $\leq 5.7\times10^{2}$ | **0** | $\leq 1.3$ | **0** | $6.4\times10^{1}$ | **$4.0\times10^{-4}$** |
| D466G | $3.2\times10^{1}$ | **0** | $7.7\times10^{1}$ | $4.1\times10^{-1}$ | $5.2\times10^{5}$ | **$7.3\times10^{-3}$** | $7.2\times10^{4}$ | **$1.0\times10^{-3}$** | 5.6 | **$6.0\times10^{-4}$** | $6.7\times10^{2}$ | $5.7\times10^{-2}$ |
| D466V | $8.5\times10^{1}$ | $8.5\times10^{-2}$ | $9.4\times10^{1}$ | $3.9\times10^{-1}$ | $1.0\times10^{6}$ | $2.3\times10^{-1}$ | $3.7\times10^{4}$ | **0** | 5.1 | **$3.0\times10^{-3}$** | $7.2\times10^{2}$ | **$5.1\times10^{-3}$** |
| I467G | $2.2\times10^{1}$ | **$2.0\times10^{-4}$** | $7.1\times10^{1}$ | $1.7\times10^{-1}$ | $3.1\times10^{5}$ | **$1.3\times10^{-3}$** | $4.7\times10^{4}$ | **0** | 1.7 | **$4.0\times10^{-4}$** | $6.2\times10^{2}$ | $7.9\times10^{-1}$ |
| I467V | $7.3\times10^{1}$ | $2.1\times10^{-2}$ | $8.7\times10^{1}$ | $9.0\times10^{-1}$ | $9.4\times10^{5}$ | $1.4\times10^{-1}$ | $4.9\times10^{5}$ | $2.7\times10^{-1}$ | 7.7 | $1.5\times10^{-2}$ | $6.2\times10^{2}$ | $9.6\times10^{-1}$ |
| Q468G | $2.1\times10^{2}$ | **$1.3\times10^{-3}$** | $8.2\times10^{1}$ | $7.1\times10^{-1}$ | $2.3\times10^{6}$ | $1.7\times10^{-2}$ | $4.0\times10^{5}$ | $8.6\times10^{-2}$ | $1.7\times10^{1}$ | $7.4\times10^{-1}$ | $6.6\times10^{2}$ | $1.2\times10^{-1}$ |
| Q468V | $5.0\times10^{1}$ | **$9.0\times10^{-4}$** | $7.2\times10^{1}$ | $1.6\times10^{-1}$ | $6.0\times10^{5}$ | **$5.6\times10^{-3}$** | $3.9\times10^{5}$ | $3.6\times10^{-2}$ | 5.0 | **$3.4\times10^{-3}$** | $7.8\times10^{2}$ | **$2.8\times10^{-3}$** |
| I469G | $1.0\times10^{2}$ | $4.0\times10^{-1}$ | $7.7\times10^{1}$ | $5.0\times10^{-1}$ | $1.3\times10^{6}$ | $8.1\times10^{-1}$ | $1.1\times10^{5}$ | **$4.6\times10^{-3}$** | 7.5 | **$7.4\times10^{-3}$** | $6.4\times10^{2}$ | $3.7\times10^{-1}$ |
| I469V | $7.1\times10^{1}$ | $2.9\times10^{-2}$ | $8.5\times10^{1}$ | $9.3\times10^{-1}$ | $8.1\times10^{5}$ | $6.9\times10^{-2}$ | $5.6\times10^{5}$ | $7.0\times10^{-1}$ | $1.1\times10^{1}$ | $2.1\times10^{-1}$ | $6.1\times10^{2}$ | $9.5\times10^{-1}$ |
| I470G | 6.7 | **$2.0\times10^{-4}$** | $5.1\times10^{1}$ | **$5.1\times10^{-3}$** | $1.8\times10^{5}$ | **$1.5\times10^{-3}$** | $2.2\times10^{4}$ | **$6.0\times10^{-4}$** | 1.9 | **$2.0\times10^{-4}$** | $4.3\times10^{2}$ | **$4.0\times10^{-4}$** |
| I470V | $4.4\times10^{1}$ | **$9.0\times10^{-4}$** | $9.5\times10^{1}$ | $3.2\times10^{-1}$ | $4.8\times10^{5}$ | **$5.6\times10^{-3}$** | $5.2\times10^{5}$ | $5.1\times10^{-1}$ | 7.1 | $1.5\times10^{-2}$ | $6.0\times10^{2}$ | $4.4\times10^{-1}$ |
| S471G | $1.7\times10^{2}$ | $4.3\times10^{-2}$ | $8.4\times10^{1}$ | $9.0\times10^{-1}$ | $2.3\times10^{6}$ | $1.6\times10^{-2}$ | $6.7\times10^{5}$ | $6.2\times10^{-1}$ | $1.9\times10^{1}$ | $3.6\times10^{-1}$ | $5.0\times10^{2}$ | **$1.3\times10^{-3}$** |
| S471V | $1.3\times10^{2}$ | $6.9\times10^{-1}$ | $1.1\times10^{2}$ | $1.0\times10^{-1}$ | $1.5\times10^{6}$ | $8.0\times10^{-1}$ | $1.1\times10^{6}$ | **$1.4\times10^{-3}$** | $1.1\times10^{1}$ | $2.1\times10^{-1}$ | $6.1\times10^{2}$ | $7.1\times10^{-1}$ |
| K472G | $2.9\times10^{1}$ | **$3.0\times10^{-4}$** | $5.7\times10^{1}$ | $2.2\times10^{-2}$ | $6.0\times10^{5}$ | $2.4\times10^{-2}$ | $2.9\times10^{5}$ | $1.7\times10^{-2}$ | 7.3 | **$2.4\times10^{-3}$** | $4.7\times10^{2}$ | **$1.\times10^{-4}$** |
| K472V | $2.6\times10^{1}$ | **0** | $7.9\times10^{1}$ | $5.1\times10^{-1}$ | $3.7\times10^{5}$ | **$1.6\times10^{-3}$** | $3.5\times10^{5}$ | $4.9\times10^{-2}$ | $\leq 2.0$ | **$1.3\times10^{-3}$** | $5.1\times10^{2}$ | **$3.0\times10^{-4}$** |
| D473G | $1.3\times10^{2}$ | $7.1\times10^{-1}$ | $7.4\times10^{1}$ | $2.8\times10^{-1}$ | $1.8\times10^{6}$ | $2.6\times10^{-1}$ | $5.6\times10^{5}$ | $6.7\times10^{-1}$ | $2.2\times10^{1}$ | $6.4\times10^{-2}$ | $5.7\times10^{2}$ | $1.4\times10^{-1}$ |
| D473V | $4.4\times10^{1}$ | **$5.0\times10^{-4}$** | $6.2\times10^{1}$ | $2.5\times10^{-2}$ | $6.2\times10^{5}$ | $1.6\times10^{-2}$ | $8.4\times10^{3}$ | **0** | $1.4\times10^{1}$ | $4.6\times10^{-1}$ | $3.1\times10^{2}$ | **$3.0\times10^{-4}$** |
| G474A | $3.5\times10^{1}$ | **$1.\times10^{-4}$** | $5.7\times10^{1}$ | $2.2\times10^{-2}$ | $7.9\times10^{5}$ | $9.5\times10^{-2}$ | $4.9\times10^{5}$ | $3.8\times10^{-1}$ | 7.9 | **$3.6\times10^{-3}$** | $4.9\times10^{2}$ | **$2.0\times10^{-4}$** |
| G474V | $1.8\times10^{1}$ | **$3.0\times10^{-4}$** | $6.3\times10^{1}$ | $9.6\times10^{-2}$ | $2.9\times10^{5}$ | **$7.0\times10^{-3}$** | $4.8\times10^{4}$ | **$3.2\times10^{-3}$** | $\leq 2.9$ | **$6.1\times10^{-3}$** | $4.3\times10^{2}$ | **0** |
| Y475G | $1.2\times10^{1}$ | **0** | $5.0\times10^{1}$ | **$2.3\times10^{-3}$** | $2.2\times10^{5}$ | **$7.0\times10^{-4}$** | $4.3\times10^{4}$ | **$3.0\times10^{-4}$** | 1.7 | **$1.\times10^{-4}$** | $4.0\times10^{2}$ | **0** |
| Y475V | $7.0\times10^{1}$ | $8.3\times10^{-2}$ | $1.0\times10^{2}$ | $2.5\times10^{-1}$ | $7.2\times10^{5}$ | $1.2\times10^{-1}$ | $5.2\times10^{5}$ | $4.8\times10^{-1}$ | 5.7 | **$6.6\times10^{-3}$** | $6.3\times10^{2}$ | $6.3\times10^{-1}$ |
| L476G | $7.8\times10^{1}$ | $1.6\times10^{-2}$ | $7.0\times10^{1}$ | $1.1\times10^{-1}$ | $1.3\times10^{6}$ | $7.0\times10^{-1}$ | $8.8\times10^{5}$ | **$9.6\times10^{-3}$** | 7.2 | **0** | $3.2\times10^{2}$ | **0** |
| L476V | $1.3\times10^{2}$ | $5.5\times10^{-1}$ | $1.1\times10^{2}$ | $3.3\times10^{-2}$ | $1.3\times10^{6}$ | $7.5\times10^{-1}$ | $6.1\times10^{5}$ | $9.5\times10^{-1}$ | $1.2\times10^{1}$ | $2.2\times10^{-1}$ | $7.9\times10^{2}$ | **0** |
| S477G | $1.2\times10^{2}$ | $8.7\times10^{-1}$ | $6.8\times10^{1}$ | $1.3\times10^{-1}$ | $1.5\times10^{6}$ | $7.3\times10^{-1}$ | $3.8\times10^{5}$ | $3.4\times10^{-2}$ | $3.0\times10^{1}$ | **0** | $5.1\times10^{2}$ | $1.3\times10^{-2}$ |
| S477V | $7.4\times10^{1}$ | $9.9\times10^{-2}$ | $1.0\times10^{2}$ | $1.6\times10^{-1}$ | $7.0\times10^{5}$ | $9.1\times10^{-2}$ | $4.7\times10^{5}$ | $1.7\times10^{-1}$ | $\leq 9.7\times10^{-1}$ | **$1.2\times10^{-3}$** | $5.5\times10^{2}$ | $4.5\times10^{-2}$ |
| A478G | $8.9\times10^{1}$ | $1.1\times10^{-1}$ | $8.3\times10^{1}$ | $8.6\times10^{-1}$ | $1.3\times10^{6}$ | $7.7\times10^{-1}$ | $4.7\times10^{5}$ | $2.5\times10^{-1}$ | $1.7\times10^{1}$ | $6.8\times10^{-1}$ | $6.1\times10^{2}$ | $8.2\times10^{-1}$ |

| Variant | $k_{cat}$ cMUP ($s^{-1}$) | $k_{cat}$ cMUP *p*–value | $K_M$ cMUP (μM) | $K_M$ cMUP *p*–value | $k_{cat}/K_M$ cMUP ($M^{-1}s^{-1}$) | $k_{cat}/K_M$ cMUP *p*–value | $k_{cat}/K_M$ MeP ($M^{-1}s^{-1}$) | $k_{cat}/K_M$ MeP *p*–value | $k_{cat}/K_M$ MecMUP ($M^{-1}s^{-1}$) | $k_{cat}/K_M$ MecMUP *p*–value | $K_i$ $PO_4$ (μM) | $K_i$ $PO_4$ *p*–value |
|---|---|---|---|---|---|---|---|---|---|---|---|---|
| A478V | $1.0\times10^{2}$ | $3.0\times10^{-1}$ | $9.0\times10^{1}$ | $6.3\times10^{-1}$ | $1.1\times10^{6}$ | $3.8\times10^{-1}$ | $3.1\times10^{5}$ | $3.0\times10^{-2}$ | 6.2 | $1.1\times10^{-2}$ | $6.7\times10^{2}$ | $1.2\times10^{-1}$ |
| Y479G | $1.7\times10^{2}$ | $2.1\times10^{-1}$ | $7.3\times10^{1}$ | $4.4\times10^{-1}$ | $2.4\times10^{6}$ | $6.6\times10^{-2}$ | $1.1\times10^{6}$ | **$8.4\times10^{-3}$** | $1.2\times10^{1}$ | $1.6\times10^{-1}$ | $5.3\times10^{2}$ | $1.4\times10^{-2}$ |
| Y479V | $1.3\times10^{2}$ | $4.7\times10^{-1}$ | $1.2\times10^{2}$ | **$5.0\times10^{-4}$** | $1.1\times10^{6}$ | $3.4\times10^{-1}$ | $7.1\times10^{5}$ | $2.7\times10^{-1}$ | 5.3 | **$3.0\times10^{-4}$** | $6.4\times10^{2}$ | $2.8\times10^{-1}$ |
| S480G | $1.1\times10^{2}$ | $4.6\times10^{-1}$ | $6.3\times10^{1}$ | $6.3\times10^{-2}$ | $1.6\times10^{6}$ | $5.6\times10^{-1}$ | $4.8\times10^{5}$ | $2.4\times10^{-1}$ | $2.5\times10^{1}$ | **$4.4\times10^{-3}$** | $4.1\times10^{2}$ | **$3.0\times10^{-4}$** |
| S480V | $1.4\times10^{2}$ | $3.3\times10^{-1}$ | $1.3\times10^{2}$ | **$7.0\times10^{-4}$** | $9.6\times10^{5}$ | $1.6\times10^{-1}$ | $7.6\times10^{5}$ | $2.2\times10^{-1}$ | 5.6 | $5.1\times10^{-2}$ | $7.2\times10^{2}$ | **$8.7\times10^{-3}$** |
| K481G | $1.1\times10^{2}$ | $5.9\times10^{-1}$ | $8.3\times10^{1}$ | $8.7\times10^{-1}$ | $1.8\times10^{6}$ | $2.8\times10^{-1}$ | $6.3\times10^{5}$ | $8.5\times10^{-1}$ | $1.6\times10^{1}$ | $8.6\times10^{-1}$ | $6.0\times10^{2}$ | $5.7\times10^{-1}$ |
| K481V | $1.3\times10^{2}$ | $7.9\times10^{-1}$ | $1.1\times10^{2}$ | $8.1\times10^{-2}$ | $1.3\times10^{6}$ | $7.5\times10^{-1}$ | $5.1\times10^{5}$ | $4.2\times10^{-1}$ | $1.1\times10^{1}$ | $2.6\times10^{-1}$ | $6.8\times10^{2}$ | $7.4\times10^{-2}$ |
| K482G | $7.8\times10^{1}$ | $4.6\times10^{-2}$ | $7.8\times10^{1}$ | $4.2\times10^{-1}$ | $1.1\times10^{6}$ | $3.6\times10^{-1}$ | $3.5\times10^{5}$ | $4.7\times10^{-2}$ | 7.2 | **$4.6\times10^{-3}$** | $4.8\times10^{2}$ | **0** |
| K482V | $1.3\times10^{2}$ | $6.2\times10^{-1}$ | $9.4\times10^{1}$ | $4.4\times10^{-1}$ | $1.4\times10^{6}$ | $9.4\times10^{-1}$ | $5.7\times10^{5}$ | $7.3\times10^{-1}$ | $1.1\times10^{1}$ | $1.3\times10^{-1}$ | $6.3\times10^{2}$ | $4.7\times10^{-1}$ |
| G483A | $3.6\times10^{1}$ | **$3.0\times10^{-4}$** | $6.0\times10^{1}$ | $1.7\times10^{-2}$ | $4.7\times10^{5}$ | **$5.5\times10^{-3}$** | $1.1\times10^{5}$ | **0** | 7.4 | **$9.0\times10^{-4}$** | $3.1\times10^{2}$ | **$1.\times10^{-4}$** |
| G483V | $3.1\times10^{1}$ | **$9.8\times10^{-3}$** | $3.7\times10^{1}$ | **$9.5\times10^{-3}$** | $9.4\times10^{5}$ | $3.5\times10^{-1}$ | $1.5\times10^{5}$ | **$2.6\times10^{-3}$** | 3.5 | **$5.0\times10^{-4}$** | $2.0\times10^{2}$ | **$2.0\times10^{-4}$** |
| T484G | $6.9\times10^{1}$ | $4.0\times10^{-2}$ | $5.0\times10^{1}$ | **$4.4\times10^{-3}$** | $1.6\times10^{6}$ | $6.7\times10^{-1}$ | $2.7\times10^{5}$ | $1.2\times10^{-2}$ | 7.5 | **$2.0\times10^{-3}$** | $2.8\times10^{2}$ | **0** |
| T484V | $8.7\times10^{1}$ | $4.5\times10^{-2}$ | $1.0\times10^{2}$ | $9.3\times10^{-2}$ | $8.4\times10^{5}$ | $5.0\times10^{-2}$ | $3.5\times10^{5}$ | $4.1\times10^{-2}$ | 9.1 | $2.0\times10^{-2}$ | $5.6\times10^{2}$ | $6.7\times10^{-2}$ |
| T485G | $9.6\times10^{1}$ | $1.9\times10^{-1}$ | $6.8\times10^{1}$ | $1.0\times10^{-1}$ | $1.3\times10^{6}$ | $6.2\times10^{-1}$ | $4.4\times10^{5}$ | $2.1\times10^{-1}$ | $2.0\times10^{1}$ | $3.0\times10^{-1}$ | $4.0\times10^{2}$ | **$1.1\times10^{-4}$** |
| T485V | $7.2\times10^{1}$ | $2.6\times10^{-2}$ | $1.2\times10^{2}$ | **$6.4\times10^{-3}$** | $6.6\times10^{5}$ | $2.2\times10^{-2}$ | $5.1\times10^{5}$ | $4.4\times10^{-1}$ | ≤ 1.4 | **$7.0\times10^{-4}$** | $4.0\times10^{2}$ | **0** |
| H486G | 6.6 | **0** | $1.6\times10^{2}$ | **0** | $4.3\times10^{4}$ | **0** | ≤ $1.2\times10^{3}$ | **$1.\times10^{-4}$** | ≤ 1.1 | **0** | – | – |
| H486V | $6.9\times10^{-1}$ | **$4.7\times10^{-3}$** | $9.9\times10^{1}$ | $4.0\times10^{-1}$ | $6.9\times10^{3}$ | **$9.7\times10^{-3}$** | ≤ $9.6\times10^{2}$ | **$7.0\times10^{-4}$** | ≤ 1.3 | **$5.0\times10^{-4}$** | – | – |
| S487G | $9.0\times10^{1}$ | $2.1\times10^{-1}$ | $4.8\times10^{1}$ | $1.1\times10^{-2}$ | $1.3\times10^{6}$ | $7.5\times10^{-1}$ | ≤ $4.1\times10^{5}$ | $4.1\times10^{-1}$ | $4.9\times10^{1}$ | **0** | $3.9\times10^{2}$ | **$2.0\times10^{-4}$** |
| S487V | $1.6\times10^{2}$ | $2.9\times10^{-2}$ | $9.4\times10^{1}$ | $2.9\times10^{-1}$ | $1.7\times10^{6}$ | $3.5\times10^{-1}$ | $1.8\times10^{5}$ | **$4.0\times10^{-4}$** | ≤ $7.7\times10^{-1}$ | **$1.\times10^{-4}$** | $1.0\times10^{3}$ | **$1.1\times10^{-4}$** |
| V488A | $2.5\times10^{2}$ | **$4.0\times10^{-4}$** | $8.6\times10^{1}$ | $9.2\times10^{-1}$ | $3.2\times10^{6}$ | **$3.0\times10^{-4}$** | $8.3\times10^{5}$ | $7.4\times10^{-2}$ | $2.9\times10^{1}$ | **$3.0\times10^{-4}$** | $5.7\times10^{2}$ | $3.3\times10^{-1}$ |
| V488G | $1.5\times10^{2}$ | $1.0\times10^{-1}$ | $5.8\times10^{1}$ | $1.2\times10^{-2}$ | $2.8\times10^{6}$ | **$1.\times10^{-4}$** | $4.9\times10^{5}$ | $3.2\times10^{-1}$ | $6.2\times10^{1}$ | **$1.\times10^{-4}$** | $3.6\times10^{2}$ | **0** |
| W489G | 7.5 | **0** | $2.5\times10^{1}$ | **0** | $3.6\times10^{5}$ | **$2.0\times10^{-4}$** | $6.8\times10^{4}$ | **0** | 7.7 | **$1.7\times10^{-3}$** | $1.2\times10^{2}$ | **0** |
| W489V | $3.5\times10^{1}$ | **$2.0\times10^{-4}$** | $5.9\times10^{1}$ | $1.8\times10^{-2}$ | $6.2\times10^{5}$ | $1.8\times10^{-2}$ | $1.4\times10^{4}$ | **0** | ≤ 2.4 | **$2.6\times10^{-3}$** | $3.0\times10^{2}$ | **$2.0\times10^{-4}$** |
| N490G | $1.0\times10^{2}$ | $4.3\times10^{-1}$ | $6.0\times10^{1}$ | $2.4\times10^{-2}$ | $1.8\times10^{6}$ | $2.7\times10^{-1}$ | $2.1\times10^{5}$ | **$4.8\times10^{-3}$** | $2.1\times10^{1}$ | $1.6\times10^{-1}$ | $4.4\times10^{2}$ | **$1.1\times10^{-4}$** |
| N490V | $6.3\times10^{1}$ | **$2.3\times10^{-3}$** | $7.6\times10^{1}$ | $2.5\times10^{-1}$ | $1.0\times10^{6}$ | $1.2\times10^{-1}$ | $1.8\times10^{5}$ | **0** | 9.9 | $7.6\times10^{-2}$ | $4.9\times10^{2}$ | **0** |
| S491G | $5.6\times10^{1}$ | $1.1\times10^{-2}$ | $6.3\times10^{1}$ | $4.8\times10^{-2}$ | $8.9\times10^{5}$ | $1.4\times10^{-1}$ | $2.7\times10^{5}$ | $1.1\times10^{-2}$ | 8.0 | **$4.4\times10^{-3}$** | $5.3\times10^{2}$ | **$6.5\times10^{-3}$** |
| S491V | $5.5\times10^{1}$ | **$1.9\times10^{-3}$** | $6.4\times10^{1}$ | $2.6\times10^{-2}$ | $1.0\times10^{6}$ | $1.6\times10^{-1}$ | $2.8\times10^{5}$ | **$5.7\times10^{-3}$** | 5.3 | **$9.0\times10^{-4}$** | $5.0\times10^{2}$ | **$1.1\times10^{-4}$** |
| Y492G | $4.2\times10^{1}$ | **$2.2\times10^{-3}$** | $3.6\times10^{1}$ | **$5.0\times10^{-4}$** | $1.3\times10^{6}$ | $7.2\times10^{-1}$ | $2.2\times10^{5}$ | **$6.1\times10^{-3}$** | $1.3\times10^{1}$ | $2.9\times10^{-1}$ | $2.5\times10^{2}$ | **$1.1\times10^{-4}$** |
| Y492V | $4.4\times10^{1}$ | **$9.0\times10^{-4}$** | $6.8\times10^{1}$ | $9.6\times10^{-2}$ | $7.5\times10^{5}$ | $4.9\times10^{-2}$ | $2.7\times10^{5}$ | **$2.2\times10^{-3}$** | ≤ 1.9 | **0** | $6.0\times10^{2}$ | $5.7\times10^{-1}$ |
| D493G | $1.8\times10^{1}$ | **0** | $3.8\times10^{1}$ | **0** | $7.5\times10^{5}$ | $4.8\times10^{-2}$ | $3.7\times10^{4}$ | **$2.0\times10^{-4}$** | 2.1 | $1.0\times10^{-2}$ | $1.3\times10^{2}$ | **$1.1\times10^{-4}$** |
| D493V | $5.8\times10^{1}$ | **$2.6\times10^{-3}$** | $5.4\times10^{1}$ | **$2.2\times10^{-3}$** | $1.2\times10^{6}$ | $5.1\times10^{-1}$ | $8.2\times10^{4}$ | **$3.0\times10^{-4}$** | 3.6 | **$1.4\times10^{-3}$** | $2.1\times10^{2}$ | **0** |
| S494G | $5.2\times10^{1}$ | **$6.2\times10^{-3}$** | $6.2\times10^{1}$ | $5.3\times10^{-2}$ | $8.3\times10^{5}$ | $1.1\times10^{-1}$ | $2.0\times10^{5}$ | **$3.8\times10^{-3}$** | $1.1\times10^{1}$ | $1.2\times10^{-1}$ | $4.2\times10^{2}$ | **$1.1\times10^{-4}$** |
| S494V | $1.7\times10^{1}$ | **0** | $5.4\times10^{1}$ | **$3.0\times10^{-4}$** | $2.8\times10^{5}$ | **0** | $7.1\times10^{4}$ | **0** | 1.9 | **$5.0\times10^{-4}$** | $2.4\times10^{2}$ | **0** |
| H495G | – | – | – | – | ≤ $1.1\times10^{3}$ | **0** | ≤ $1.2\times10^{3}$ | **0** | ≤ 1.3 | **0** | – | – |
| H495V | $1.9\times10^{1}$ | **$5.1\times10^{-3}$** | $8.5\times10^{1}$ | 1.0 | $2.1\times10^{5}$ | $1.6\times10^{-2}$ | $1.5\times10^{5}$ | $1.1\times10^{-2}$ | ≤ 1.9 | **$3.0\times10^{-4}$** | $5.1\times10^{2}$ | $1.9\times10^{-2}$ |
| I496G | $5.8\times10^{-1}$ | **$1.\times10^{-4}$** | $7.4\times10^{1}$ | $3.0\times10^{-1}$ | $7.5\times10^{3}$ | **$3.0\times10^{-4}$** | ≤ $1.3\times10^{3}$ | **0** | ≤ 1.1 | **0** | $3.9\times10^{2}$ | **$1.1\times10^{-3}$** |
| I496V | $3.6\times10^{1}$ | **$4.0\times10^{-4}$** | $8.9\times10^{1}$ | $7.7\times10^{-1}$ | $4.1\times10^{5}$ | **$4.9\times10^{-3}$** | $3.4\times10^{5}$ | $2.0\times10^{-2}$ | 6.2 | **$7.3\times10^{-3}$** | $5.2\times10^{2}$ | **$8.1\times10^{-3}$** |
| P497G | $1.8\times10^{1}$ | **$3.6\times10^{-3}$** | $6.0\times10^{1}$ | $8.0\times10^{-2}$ | $2.2\times10^{5}$ | **$6.5\times10^{-3}$** | $9.8\times10^{4}$ | **$1.0\times10^{-3}$** | ≤ 1.1 | **$2.0\times10^{-4}$** | $5.7\times10^{2}$ | $2.8\times10^{-1}$ |
| L498G | $4.7\times10^{1}$ | $1.3\times10^{-2}$ | $4.1\times10^{1}$ | **$3.7\times10^{-3}$** | $8.5\times10^{5}$ | $1.9\times10^{-1}$ | ≤ $1.4\times10^{5}$ | **$2.3\times10^{-3}$** | $2.0\times10^{1}$ | $3.3\times10^{-1}$ | $4.6\times10^{2}$ | **$4.0\times10^{-4}$** |
| L498V | $5.6\times10^{1}$ | $1.2\times10^{-2}$ | $1.0\times10^{2}$ | $2.0\times10^{-1}$ | $5.6\times10^{5}$ | $2.3\times10^{-2}$ | $2.8\times10^{5}$ | **$4.8\times10^{-3}$** | 5.0 | **$7.0\times10^{-4}$** | $6.7\times10^{2}$ | $1.1\times10^{-1}$ |
| L499G | $9.1\times10^{1}$ | $2.6\times10^{-1}$ | $7.5\times10^{1}$ | $5.1\times10^{-1}$ | $1.1\times10^{6}$ | $5.1\times10^{-1}$ | $1.3\times10^{5}$ | **$9.0\times10^{-4}$** | 7.7 | $2.4\times10^{-2}$ | $6.0\times10^{2}$ | $6.3\times10^{-1}$ |
| L499V | $9.2\times10^{1}$ | $1.1\times10^{-1}$ | $9.8\times10^{1}$ | $1.8\times10^{-1}$ | $9.8\times10^{5}$ | $1.6\times10^{-1}$ | $4.9\times10^{5}$ | $2.8\times10^{-1}$ | 9.5 | $5.1\times10^{-2}$ | $6.8\times10^{2}$ | $6.9\times10^{-2}$ |
| F500G | $1.6\times10^{-2}$ | **$1.4\times10^{-3}$** | $2.7\times10^{1}$ | **$2.0\times10^{-4}$** | $7.3\times10^{2}$ | **$3.0\times10^{-3}$** | ≤ $1.1\times10^{3}$ | **0** | ≤ 1.6 | **0** | – | – |
| F500V | $3.2\times10^{1}$ | **$2.1\times10^{-3}$** | $7.3\times10^{1}$ | $3.4\times10^{-1}$ | $4.6\times10^{5}$ | $2.1\times10^{-2}$ | $1.1\times10^{5}$ | **$1.0\times10^{-3}$** | ≤ 1.8 | **$1.3\times10^{-3}$** | $4.2\times10^{2}$ | **0** |
| M501G | $5.6\times10^{1}$ | $2.9\times10^{-2}$ | $7.3\times10^{1}$ | $3.4\times10^{-1}$ | $7.6\times10^{5}$ | $1.3\times10^{-1}$ | $3.3\times10^{4}$ | **0** | 2.9 | **$1.2\times10^{-3}$** | $5.9\times10^{2}$ | $3.5\times10^{-1}$ |
| M501V | $9.4\times10^{1}$ | $2.6\times10^{-1}$ | $7.4\times10^{1}$ | $2.6\times10^{-1}$ | $1.1\times10^{6}$ | $3.2\times10^{-1}$ | $1.9\times10^{5}$ | $7.5\times10^{-2}$ | ≤ 5.0 | $2.6\times10^{-2}$ | $5.9\times10^{2}$ | $3.8\times10^{-1}$ |
| G502A | $1.8\times10^{1}$ | **$4.7\times10^{-3}$** | $9.1\times10^{1}$ | $7.0\times10^{-1}$ | $2.2\times10^{5}$ | $1.6\times10^{-2}$ | $7.7\times10^{4}$ | **$1.4\times10^{-3}$** | ≤ $9.9\times10^{-1}$ | **$2.0\times10^{-4}$** | $5.5\times10^{2}$ | $2.1\times10^{-1}$ |
| G502V | $8.3\times10^{-2}$ | **$5.0\times10^{-4}$** | $5.2\times10^{1}$ | $1.9\times10^{-2}$ | $1.5\times10^{3}$ | **$1.9\times10^{-3}$** | ≤ $1.1\times10^{4}$ | **0** | ≤ 1.1 | **0** | – | – |
| W503G | $3.0\times10^{1}$ | **$4.0\times10^{-4}$** | $6.9\times10^{1}$ | $1.5\times10^{-1}$ | $5.6\times10^{5}$ | $1.7\times10^{-2}$ | $1.4\times10^{5}$ | **$1.3\times10^{-3}$** | 5.6 | **$9.0\times10^{-4}$** | $6.0\times10^{2}$ | $5.6\times10^{-1}$ |
| W503V | $4.5\times10^{1}$ | **$1.0\times10^{-3}$** | $8.4\times10^{1}$ | $8.5\times10^{-1}$ | $4.2\times10^{5}$ | **$3.0\times10^{-3}$** | $9.9\times10^{4}$ | $2.3\times10^{-2}$ | 6.4 | $1.2\times10^{-2}$ | $6.0\times10^{2}$ | $5.4\times10^{-1}$ |
| G504A | $9.1\times10^{1}$ | $1.1\times10^{-1}$ | $6.9\times10^{1}$ | $1.1\times10^{-1}$ | $1.2\times10^{6}$ | $5.7\times10^{-1}$ | $2.7\times10^{5}$ | $5.7\times10^{-2}$ | 9.6 | $2.0\times10^{-2}$ | $6.1\times10^{2}$ | $6.7\times10^{-1}$ |
| G504V | $3.5\times10^{1}$ | **$2.0\times10^{-4}$** | $8.4\times10^{1}$ | $9.1\times10^{-1}$ | $3.7\times10^{5}$ | **$6.0\times10^{-4}$** | $2.1\times10^{5}$ | **$4.0\times10^{-3}$** | ≤ 1.9 | **$1.\times10^{-4}$** | $6.4\times10^{2}$ | $3.7\times10^{-1}$ |
| I505G | $1.2\times10^{1}$ | **0** | $6.3\times10^{1}$ | $2.3\times10^{-2}$ | $1.8\times10^{5}$ | **$3.0\times10^{-4}$** | $2.5\times10^{4}$ | **0** | ≤ 1.2 | **0** | $6.4\times10^{2}$ | $3.6\times10^{-1}$ |
| I505V | $1.6\times10^{2}$ | $5.2\times10^{-2}$ | $9.4\times10^{1}$ | $4.1\times10^{-1}$ | $1.9\times10^{6}$ | $1.8\times10^{-1}$ | $6.1\times10^{5}$ | 1.0 | $1.3\times10^{1}$ | $5.9\times10^{-1}$ | $6.8\times10^{2}$ | $6.2\times10^{-2}$ |
| K506G | $1.7\times10^{2}$ | $3.6\times10^{-2}$ | $7.3\times10^{1}$ | $2.8\times10^{-1}$ | $2.3\times10^{6}$ | $2.8\times10^{-2}$ | $4.3\times10^{5}$ | $1.2\times10^{-1}$ | $1.8\times10^{1}$ | $5.2\times10^{-1}$ | $5.9\times10^{2}$ | $3.9\times10^{-1}$ |
| K506V | $1.2\times10^{2}$ | $7.9\times10^{-1}$ | $9.3\times10^{1}$ | $3.3\times10^{-1}$ | $1.2\times10^{6}$ | $4.6\times10^{-1}$ | $3.6\times10^{5}$ | $5.0\times10^{-2}$ | 9.9 | $1.1\times10^{-1}$ | $6.9\times10^{2}$ | $1.7\times10^{-2}$ |
| Q507G | $1.1\times10^{2}$ | $5.5\times10^{-1}$ | $6.8\times10^{1}$ | $2.2\times10^{-1}$ | $1.6\times10^{6}$ | $6.7\times10^{-1}$ | $5.1\times10^{5}$ | $4.0\times10^{-1}$ | $1.4\times10^{1}$ | $5.2\times10^{-1}$ | $5.3\times10^{2}$ | **$9.2\times10^{-3}$** |
| Q507V | $1.8\times10^{2}$ | $7.1\times10^{-2}$ | $8.6\times10^{1}$ | $9.3\times10^{-1}$ | $1.7\times10^{6}$ | $5.6\times10^{-1}$ | $6.9\times10^{5}$ | $5.2\times10^{-1}$ | $1.3\times10^{1}$ | $4.2\times10^{-1}$ | $6.3\times10^{2}$ | $6.7\times10^{-1}$ |
| G508A | $4.9\times10^{1}$ | **$1.7\times10^{-3}$** | $6.6\times10^{1}$ | $7.5\times10^{-2}$ | $6.9\times10^{5}$ | $2.9\times10^{-2}$ | $2.5\times10^{5}$ | **$9.7\times10^{-3}$** | 8.9 | $1.3\times10^{-2}$ | $5.2\times10^{2}$ | **$6.3\times10^{-3}$** |
| G508V | $1.7\times10^{1}$ | **0** | $8.0\times10^{1}$ | $5.6\times10^{-1}$ | $2.4\times10^{5}$ | **$2.0\times10^{-4}$** | $1.2\times10^{5}$ | **$4.0\times10^{-4}$** | 2.2 | **$9.0\times10^{-4}$** | $5.0\times10^{2}$ | **0** |
| E509G | $1.3\times10^{1}$ | **$2.0\times10^{-4}$** | $6.0\times10^{1}$ | $4.2\times10^{-2}$ | $2.2\times10^{5}$ | **$1.1\times10^{-3}$** | $1.6\times10^{5}$ | $3.9\times10^{-2}$ | 2.7 | **$2.0\times10^{-4}$** | $4.3\times10^{2}$ | **0** |
| E509V | $4.2\times10^{1}$ | **$1.4\times10^{-3}$** | $8.5\times10^{1}$ | $9.7\times10^{-1}$ | $5.7\times10^{5}$ | $2.2\times10^{-2}$ | $2.1\times10^{5}$ | **$1.1\times10^{-3}$** | 4.7 | **$4.5\times10^{-3}$** | $5.5\times10^{2}$ | $4.5\times10^{-2}$ |
| S510G | $7.3\times10^{1}$ | $3.2\times10^{-2}$ | $6.6\times10^{1}$ | $7.0\times10^{-2}$ | $1.2\times10^{6}$ | $6.2\times10^{-1}$ | $2.7\times10^{5}$ | $1.1\times10^{-2}$ | $1.0\times10^{1}$ | $5.9\times10^{-2}$ | $5.4\times10^{2}$ | $3.4\times10^{-2}$ |
| S510V | $1.5\times10^{1}$ | **$4.0\times10^{-4}$** | $8.0\times10^{1}$ | $7.0\times10^{-1}$ | $1.8\times10^{5}$ | **$4.1\times10^{-3}$** | $3.4\times10^{4}$ | **0** | ≤ $6.2\times10^{-1}$ | **$3.0\times10^{-4}$** | $4.5\times10^{2}$ | **$3.2\times10^{-3}$** |
| N511G | $1.2\times10^{2}$ | $9.7\times10^{-1}$ | $7.0\times10^{1}$ | $1.7\times10^{-1}$ | $1.7\times10^{6}$ | $3.9\times10^{-1}$ | $2.6\times10^{5}$ | $5.7\times10^{-2}$ | $1.5\times10^{1}$ | $8.1\times10^{-1}$ | $6.1\times10^{2}$ | $7.7\times10^{-1}$ |
| N511V | $2.6\times10^{2}$ | **$1.\times10^{-4}$** | $9.1\times10^{1}$ | $6.2\times10^{-1}$ | $2.3\times10^{6}$ | $2.4\times10^{-2}$ | $9.5\times10^{5}$ | $7.5\times10^{-2}$ | $1.7\times10^{1}$ | $8.9\times10^{-1}$ | $6.1\times10^{2}$ | $5.7\times10^{-1}$ |
| Q512G | $6.5\times10^{1}$ | $2.0\times10^{-2}$ | $6.1\times10^{1}$ | $3.9\times10^{-2}$ | $9.8\times10^{5}$ | $2.5\times10^{-1}$ | $2.1\times10^{5}$ | **$4.5\times10^{-3}$** | $1.6\times10^{1}$ | 1.0 | $5.4\times10^{2}$ | $3.3\times10^{-2}$ |
| Q512V | $8.9\times10^{1}$ | $1.2\times10^{-1}$ | $9.7\times10^{1}$ | $2.6\times10^{-1}$ | $1.1\times10^{6}$ | $3.9\times10^{-1}$ | $6.6\times10^{5}$ | $6.5\times10^{-1}$ | $1.0\times10^{1}$ | $1.6\times10^{-1}$ | $6.2\times10^{2}$ | $7.1\times10^{-1}$ |
| P513G | $9.4\times10^{1}$ | $1.4\times10^{-1}$ | $7.6\times10^{1}$ | $3.9\times10^{-1}$ | $1.3\times10^{6}$ | $7.8\times10^{-1}$ | $2.1\times10^{5}$ | **$5.6\times10^{-3}$** | $1.2\times10^{1}$ | $1.8\times10^{-1}$ | $5.9\times10^{2}$ | $3.5\times10^{-1}$ |
| P513V | $8.7\times10^{1}$ | $1.3\times10^{-2}$ | $9.1\times10^{1}$ | $5.1\times10^{-1}$ | $8.8\times10^{5}$ | $2.4\times10^{-2}$ | $3.5\times10^{5}$ | **$1.9\times10^{-3}$** | 7.7 | **$2.0\times10^{-3}$** | $6.2\times10^{2}$ | $5.3\times10^{-1}$ |
| Y514G | 9.3 | **0** | $6.8\times10^{1}$ | $8.4\times10^{-2}$ | $1.8\times10^{5}$ | **0** | $5.6\times10^{4}$ | **0** | ≤ 1.2 | **0** | $5.5\times10^{2}$ | $1.3\times10^{-2}$ |
| Y514V | $6.7\times10^{1}$ | $2.9\times10^{-2}$ | $9.0\times10^{1}$ | $7.1\times10^{-1}$ | $7.3\times10^{5}$ | $6.8\times10^{-2}$ | $2.8\times10^{5}$ | $1.4\times10^{-2}$ | 6.7 | $1.7\times10^{-2}$ | $5.7\times10^{2}$ | $2.8\times10^{-1}$ |
| H515G | $7.9\times10^{1}$ | $1.3\times10^{-1}$ | $5.1\times10^{1}$ | $2.5\times10^{-2}$ | $1.8\times10^{6}$ | $3.8\times10^{-1}$ | $8.7\times10^{5}$ | $6.0\times10^{-2}$ | $1.1\times10^{1}$ | $8.0\times10^{-2}$ | $5.2\times10^{2}$ | $1.9\times10^{-2}$ |
| H515V | $1.3\times10^{2}$ | $5.2\times10^{-1}$ | $8.6\times10^{1}$ | $8.7\times10^{-1}$ | $1.5\times10^{6}$ | $8.3\times10^{-1}$ | $2.0\times10^{5}$ | $1.4\times10^{-2}$ | $1.5\times10^{1}$ | $7.5\times10^{-1}$ | $5.6\times10^{2}$ | $1.2\times10^{-1}$ |
| M516G | $7.2\times10^{1}$ | $4.4\times10^{-2}$ | $4.7\times10^{1}$ | **$2.6\times10^{-3}$** | $1.6\times10^{6}$ | $7.2\times10^{-1}$ | $4.1\times10^{5}$ | $1.3\times10^{-1}$ | $1.6\times10^{1}$ | 1.0 | $1.9\times10^{2}$ | **$2.0\times10^{-4}$** |

| Variant | $k_{cat}$ cMUP ($s^{-1}$) | $k_{cat}$ cMUP *p*–value | $K_M$ cMUP (μM) | $K_M$ cMUP *p*–value | $k_{cat}/K_M$ cMUP ($M^{-1}s^{-1}$) | $k_{cat}/K_M$ cMUP *p*–value | $k_{cat}/K_M$ MeP ($M^{-1}s^{-1}$) | $k_{cat}/K_M$ MeP *p*–value | $k_{cat}/K_M$ MecMUP ($M^{-1}s^{-1}$) | $k_{cat}/K_M$ MecMUP *p*–value | $K_i$ $PO_4$ (μM) | $K_i$ $PO_4$ *p*–value |
|---|---|---|---|---|---|---|---|---|---|---|---|---|
| M516V | $5.2\times10^{1}$ | **$7.5\times10^{-3}$** | $9.4\times10^{1}$ | $4.5\times10^{-1}$ | $6.4\times10^{5}$ | $3.6\times10^{-2}$ | $3.1\times10^{5}$ | $1.0\times10^{-2}$ | 6.5 | $1.3\times10^{-2}$ | $5.2\times10^{2}$ | **$4.4\times10^{-3}$** |
| T517G | $2.8\times10^{1}$ | **$5.0\times10^{-4}$** | $6.0\times10^{1}$ | $3.5\times10^{-2}$ | $4.7\times10^{5}$ | **$9.9\times10^{-3}$** | $1.5\times10^{5}$ | **$2.7\times10^{-3}$** | ≤ 1.1 | **0** | $5.2\times10^{2}$ | **$5.3\times10^{-3}$** |
| T517V | $1.2\times10^{1}$ | **0** | $7.4\times10^{1}$ | $1.8\times10^{-1}$ | $1.8\times10^{5}$ | **$1.\times10^{-4}$** | $1.2\times10^{5}$ | **0** | 2.3 | **$1.3\times10^{-3}$** | $3.8\times10^{2}$ | **0** |
| D518G | $1.9\times10^{1}$ | **$6.0\times10^{-4}$** | $6.1\times10^{1}$ | $4.8\times10^{-2}$ | $2.7\times10^{5}$ | **$3.2\times10^{-3}$** | $8.2\times10^{4}$ | **$1.6\times10^{-3}$** | 3.4 | **0** | $3.8\times10^{2}$ | **0** |
| D518V | $9.2\times10^{-1}$ | **0** | $5.4\times10^{1}$ | $1.2\times10^{-2}$ | $1.8\times10^{4}$ | **$5.0\times10^{-4}$** | ≤ $1.6\times10^{4}$ | **$2.0\times10^{-4}$** | ≤ 1.3 | **0** | $4.3\times10^{2}$ | **$6.0\times10^{-4}$** |
| I519G | 9.9 | **$5.0\times10^{-4}$** | $6.2\times10^{1}$ | $5.3\times10^{-2}$ | $1.6\times10^{5}$ | **$1.3\times10^{-3}$** | $1.3\times10^{4}$ | **$1.\times10^{-4}$** | ≤ 1.1 | **$1.\times10^{-4}$** | $3.5\times10^{2}$ | **$7.0\times10^{-4}$** |
| I519V | $7.8\times10^{1}$ | $7.2\times10^{-2}$ | $8.8\times10^{1}$ | $7.7\times10^{-1}$ | $9.0\times10^{5}$ | $1.6\times10^{-1}$ | $3.4\times10^{5}$ | $4.2\times10^{-2}$ | 8.6 | $2.2\times10^{-2}$ | $5.9\times10^{2}$ | $2.3\times10^{-1}$ |
| A520G | $1.2\times10^{2}$ | $8.8\times10^{-1}$ | $6.3\times10^{1}$ | $6.2\times10^{-2}$ | $1.8\times10^{6}$ | $2.9\times10^{-1}$ | $3.6\times10^{5}$ | $4.6\times10^{-2}$ | $1.2\times10^{1}$ | $3.0\times10^{-1}$ | $4.4\times10^{2}$ | **$1.\times10^{-4}$** |
| A520V | $5.6\times10^{1}$ | $1.1\times10^{-2}$ | $8.1\times10^{1}$ | $6.8\times10^{-1}$ | $6.8\times10^{5}$ | $4.8\times10^{-2}$ | $3.7\times10^{5}$ | $5.3\times10^{-2}$ | ≤ 2.0 | **$1.8\times10^{-3}$** | $6.3\times10^{2}$ | $5.6\times10^{-1}$ |
| P521G | $4.2\times10^{1}$ | **$4.5\times10^{-3}$** | $5.7\times10^{1}$ | $4.3\times10^{-2}$ | $8.1\times10^{5}$ | $1.7\times10^{-1}$ | $2.0\times10^{5}$ | **$4.4\times10^{-3}$** | 9.9 | $4.2\times10^{-2}$ | $4.6\times10^{2}$ | **$7.0\times10^{-4}$** |
| P521V | $1.6\times10^{2}$ | $3.2\times10^{-2}$ | $8.8\times10^{1}$ | $8.2\times10^{-1}$ | $2.1\times10^{6}$ | $6.3\times10^{-2}$ | $5.0\times10^{5}$ | $3.7\times10^{-1}$ | $1.4\times10^{1}$ | $5.7\times10^{-1}$ | $6.0\times10^{2}$ | $7.0\times10^{-1}$ |
| T522G | 2.0 | **0** | $7.3\times10^{1}$ | $2.4\times10^{-1}$ | $2.8\times10^{4}$ | **0** | $4.9\times10^{3}$ | **$7.0\times10^{-4}$** | ≤ 1.2 | **0** | $5.8\times10^{2}$ | $2.4\times10^{-1}$ |
| T522V | $1.0\times10^{1}$ | **$1.0\times10^{-3}$** | $5.8\times10^{1}$ | $4.9\times10^{-2}$ | $2.3\times10^{5}$ | **$4.4\times10^{-3}$** | $2.7\times10^{4}$ | **$5.0\times10^{-4}$** | 1.9 | **$8.0\times10^{-3}$** | $3.5\times10^{2}$ | **$1.3\times10^{-3}$** |
| V523A | $1.0\times10^{2}$ | $3.1\times10^{-1}$ | $7.9\times10^{1}$ | $4.8\times10^{-1}$ | $1.1\times10^{6}$ | $3.9\times10^{-1}$ | $8.4\times10^{4}$ | **$3.9\times10^{-3}$** | 8.9 | $6.8\times10^{-2}$ | $5.4\times10^{2}$ | $3.3\times10^{-2}$ |
| V523G | $6.0\times10^{-2}$ | **0** | $3.7\times10^{1}$ | **$4.0\times10^{-4}$** | $1.5\times10^{3}$ | **$4.0\times10^{-4}$** | ≤ $1.3\times10^{3}$ | **$2.0\times10^{-4}$** | ≤ $9.9\times10^{-1}$ | **0** | – | – |
| S524G | $7.9\times10^{1}$ | $4.0\times10^{-2}$ | $5.9\times10^{1}$ | $1.4\times10^{-2}$ | $1.4\times10^{6}$ | $8.9\times10^{-1}$ | $4.1\times10^{5}$ | $8.1\times10^{-2}$ | $1.6\times10^{1}$ | $9.4\times10^{-1}$ | $5.2\times10^{2}$ | **$5.5\times10^{-3}$** |
| S524V | $8.5\times10^{1}$ | $5.6\times10^{-2}$ | $8.8\times10^{1}$ | $7.7\times10^{-1}$ | $1.0\times10^{6}$ | $1.9\times10^{-1}$ | $4.5\times10^{5}$ | $1.8\times10^{-1}$ | $1.3\times10^{1}$ | $3.3\times10^{-1}$ | $6.0\times10^{2}$ | $5.4\times10^{-1}$ |
| S525G | $7.5\times10^{1}$ | $2.9\times10^{-2}$ | $6.5\times10^{1}$ | $6.1\times10^{-2}$ | $8.5\times10^{5}$ | $8.7\times10^{-2}$ | ≤ $2.3\times10^{5}$ | $1.5\times10^{-2}$ | $1.2\times10^{1}$ | $1.9\times10^{-1}$ | $5.8\times10^{2}$ | $2.7\times10^{-1}$ |
| S525V | $9.1\times10^{1}$ | $1.2\times10^{-1}$ | $9.1\times10^{1}$ | $5.5\times10^{-1}$ | $9.5\times10^{5}$ | $1.4\times10^{-1}$ | $6.2\times10^{5}$ | $9.2\times10^{-1}$ | 9.8 | $1.4\times10^{-1}$ | $6.3\times10^{2}$ | $5.0\times10^{-1}$ |
| L526G | $1.2\times10^{1}$ | **0** | $5.9\times10^{1}$ | **$6.6\times10^{-3}$** | $2.2\times10^{5}$ | **$1.\times10^{-4}$** | $5.4\times10^{4}$ | **0** | 2.2 | **$4.0\times10^{-4}$** | $5.2\times10^{2}$ | **$5.0\times10^{-4}$** |
| L526V | $1.7\times10^{2}$ | $6.1\times10^{-2}$ | $1.0\times10^{2}$ | $1.0\times10^{-1}$ | $1.6\times10^{6}$ | $6.0\times10^{-1}$ | $7.1\times10^{5}$ | $4.9\times10^{-1}$ | $1.6\times10^{1}$ | $9.4\times10^{-1}$ | $6.2\times10^{2}$ | $7.8\times10^{-1}$ |
| L527G | 8.9 | **0** | $6.5\times10^{1}$ | $3.0\times10^{-2}$ | $1.4\times10^{5}$ | **0** | $1.2\times10^{4}$ | **0** | ≤ 1.3 | **$2.0\times10^{-4}$** | $5.3\times10^{2}$ | **$3.6\times10^{-3}$** |
| L527V | $1.2\times10^{2}$ | $7.2\times10^{-1}$ | $8.1\times10^{1}$ | $5.8\times10^{-1}$ | $1.5\times10^{6}$ | $6.2\times10^{-1}$ | $3.1\times10^{5}$ | **$5.4\times10^{-3}$** | 7.6 | $2.7\times10^{-2}$ | $6.0\times10^{2}$ | $4.8\times10^{-1}$ |
| K528G | 7.1 | **0** | $5.2\times10^{1}$ | **$2.0\times10^{-4}$** | $1.5\times10^{5}$ | **0** | $1.9\times10^{4}$ | **0** | 1.6 | **$2.0\times10^{-4}$** | $4.3\times10^{2}$ | **0** |
| K528V | $1.8\times10^{1}$ | **0** | $8.1\times10^{1}$ | $6.1\times10^{-1}$ | $2.5\times10^{5}$ | **$1.\times10^{-4}$** | $1.6\times10^{5}$ | **$2.0\times10^{-3}$** | ≤ 1.3 | **$8.0\times10^{-4}$** | $4.2\times10^{2}$ | **$6.0\times10^{-4}$** |
| I529G | $1.1\times10^{1}$ | **0** | $5.2\times10^{1}$ | **$6.0\times10^{-4}$** | $2.2\times10^{5}$ | **0** | $6.3\times10^{4}$ | **0** | 1.5 | **$1.4\times10^{-3}$** | $3.8\times10^{2}$ | **0** |
| I529V | $8.2\times10^{1}$ | $1.0\times10^{-1}$ | $8.4\times10^{1}$ | $9.2\times10^{-1}$ | $1.3\times10^{6}$ | $7.8\times10^{-1}$ | $5.8\times10^{5}$ | $8.3\times10^{-1}$ | $1.2\times10^{1}$ | $4.0\times10^{-1}$ | $6.1\times10^{2}$ | $6.9\times10^{-1}$ |
| Q530G | $9.7\times10^{1}$ | $2.1\times10^{-1}$ | $4.8\times10^{1}$ | **$8.0\times10^{-4}$** | $1.7\times10^{6}$ | $3.9\times10^{-1}$ | $6.2\times10^{4}$ | $3.9\times10^{-2}$ | 9.9 | $1.7\times10^{-2}$ | $4.7\times10^{2}$ | **$1.\times10^{-4}$** |
| Q530V | $5.2\times10^{1}$ | **$6.0\times10^{-4}$** | $6.6\times10^{1}$ | $3.4\times10^{-2}$ | $8.0\times10^{5}$ | $3.0\times10^{-2}$ | $3.9\times10^{5}$ | $1.3\times10^{-2}$ | 4.0 | **$1.\times10^{-4}$** | $4.3\times10^{2}$ | **0** |
| F531G | $1.2\times10^{2}$ | $9.1\times10^{-1}$ | $6.6\times10^{1}$ | $1.2\times10^{-1}$ | $2.1\times10^{6}$ | $6.8\times10^{-2}$ | $3.9\times10^{5}$ | $1.2\times10^{-1}$ | $1.5\times10^{1}$ | $7.5\times10^{-1}$ | $5.0\times10^{2}$ | **$2.0\times10^{-3}$** |
| F531V | $7.6\times10^{1}$ | $3.1\times10^{-2}$ | $8.2\times10^{1}$ | $7.0\times10^{-1}$ | $9.7\times10^{5}$ | $1.6\times10^{-1}$ | $3.6\times10^{5}$ | $4.4\times10^{-2}$ | 7.4 | $1.9\times10^{-2}$ | $5.8\times10^{2}$ | $2.3\times10^{-1}$ |
| P532G | $8.7\times10^{1}$ | $1.2\times10^{-1}$ | $5.3\times10^{1}$ | $1.5\times10^{-2}$ | $1.8\times10^{6}$ | $3.8\times10^{-1}$ | $2.0\times10^{5}$ | **$5.0\times10^{-3}$** | $1.7\times10^{1}$ | $9.4\times10^{-1}$ | $4.0\times10^{2}$ | **$1.\times10^{-4}$** |
| P532V | $2.7\times10^{1}$ | **$5.0\times10^{-4}$** | $6.7\times10^{1}$ | $1.3\times10^{-1}$ | $3.7\times10^{5}$ | **$4.5\times10^{-3}$** | $8.9\times10^{4}$ | $1.7\times10^{-2}$ | ≤ 1.6 | **$1.\times10^{-4}$** | $3.5\times10^{2}$ | **0** |
| S533G | $8.4\times10^{1}$ | $1.1\times10^{-1}$ | $4.2\times10^{1}$ | **$1.5\times10^{-3}$** | $1.5\times10^{6}$ | $8.2\times10^{-1}$ | $2.6\times10^{5}$ | $1.3\times10^{-2}$ | $1.6\times10^{1}$ | $8.7\times10^{-1}$ | $3.6\times10^{2}$ | **$3.0\times10^{-4}$** |
| S533V | 2.2 | **0** | $3.6\times10^{1}$ | **0** | $5.9\times10^{4}$ | **0** | ≤ $3.4\times10^{3}$ | **0** | ≤ 1.6 | **$1.\times10^{-4}$** | – | – |
| G534A | $1.3\times10^{2}$ | $5.9\times10^{-1}$ | $6.8\times10^{1}$ | $8.9\times10^{-2}$ | $2.2\times10^{6}$ | $2.9\times10^{-2}$ | $3.9\times10^{5}$ | $1.1\times10^{-1}$ | $2.9\times10^{1}$ | **0** | $4.5\times10^{2}$ | **0** |
| G534V | $2.5\times10^{1}$ | **$3.0\times10^{-4}$** | $5.4\times10^{1}$ | **$4.0\times10^{-3}$** | $4.6\times10^{5}$ | **$3.4\times10^{-3}$** | $2.0\times10^{5}$ | **$3.2\times10^{-3}$** | 7.4 | $2.1\times10^{-2}$ | $2.4\times10^{2}$ | **$1.\times10^{-4}$** |
| A535G | $1.2\times10^{2}$ | $7.9\times10^{-1}$ | $6.3\times10^{1}$ | $6.4\times10^{-2}$ | $2.1\times10^{6}$ | $5.4\times10^{-2}$ | $4.0\times10^{5}$ | $7.3\times10^{-2}$ | $2.6\times10^{1}$ | **$4.0\times10^{-4}$** | $6.0\times10^{2}$ | $6.6\times10^{-1}$ |
| A535V | $1.4\times10^{2}$ | $5.1\times10^{-1}$ | $9.5\times10^{1}$ | $3.4\times10^{-1}$ | $1.1\times10^{6}$ | $4.2\times10^{-1}$ | $6.8\times10^{5}$ | $4.3\times10^{-1}$ | $1.3\times10^{1}$ | $4.3\times10^{-1}$ | $6.4\times10^{2}$ | $3.6\times10^{-1}$ |
| V536A | $1.0\times10^{2}$ | $2.7\times10^{-1}$ | $7.7\times10^{1}$ | $3.1\times10^{-1}$ | $1.2\times10^{6}$ | $6.8\times10^{-1}$ | $1.1\times10^{5}$ | **$6.3\times10^{-3}$** | 7.3 | **$4.9\times10^{-3}$** | $5.3\times10^{2}$ | $3.5\times10^{-2}$ |
| V536G | $1.5\times10^{1}$ | **$1.\times10^{-4}$** | $6.3\times10^{1}$ | $4.9\times10^{-2}$ | $3.2\times10^{5}$ | **$4.3\times10^{-3}$** | ≤ $6.2\times10^{4}$ | **$8.0\times10^{-4}$** | 2.1 | **0** | $4.0\times10^{2}$ | **$1.\times10^{-4}$** |
| G537A | 9.4 | **0** | $6.1\times10^{1}$ | $2.0\times10^{-2}$ | $1.2\times10^{5}$ | **0** | $5.2\times10^{4}$ | **$4.8\times10^{-3}$** | ≤ $9.9\times10^{-1}$ | **0** | $4.1\times10^{2}$ | **0** |
| G537V | 3.1 | **0** | $4.2\times10^{1}$ | **0** | $8.7\times10^{4}$ | **$1.\times10^{-4}$** | ≤ $7.6\times10^{3}$ | **0** | ≤ 1.6 | **0** | $2.6\times10^{2}$ | **0** |
| K538G | $1.2\times10^{2}$ | $7.4\times10^{-1}$ | $6.4\times10^{1}$ | $9.7\times10^{-2}$ | $2.3\times10^{6}$ | $4.0\times10^{-2}$ | $2.4\times10^{5}$ | **$8.0\times10^{-3}$** | $2.2\times10^{1}$ | $1.1\times10^{-1}$ | $4.7\times10^{2}$ | **$5.4\times10^{-3}$** |
| K538V | $1.4\times10^{2}$ | $5.6\times10^{-1}$ | $8.2\times10^{1}$ | $7.3\times10^{-1}$ | $1.9\times10^{6}$ | $2.5\times10^{-1}$ | $4.3\times10^{5}$ | $1.2\times10^{-1}$ | $1.1\times10^{1}$ | $2.1\times10^{-1}$ | $5.9\times10^{2}$ | $3.4\times10^{-1}$ |
| P539G | $1.4\times10^{2}$ | $4.5\times10^{-1}$ | $8.1\times10^{1}$ | $6.6\times10^{-1}$ | $1.6\times10^{6}$ | $5.7\times10^{-1}$ | $5.5\times10^{5}$ | $6.2\times10^{-1}$ | $1.2\times10^{1}$ | $1.4\times10^{-1}$ | $6.3\times10^{2}$ | $4.3\times10^{-1}$ |
| P539V | $1.5\times10^{2}$ | $9.1\times10^{-2}$ | $9.4\times10^{1}$ | $4.1\times10^{-1}$ | $1.8\times10^{6}$ | $2.4\times10^{-1}$ | $9.1\times10^{5}$ | **$7.5\times10^{-3}$** | $1.4\times10^{1}$ | $5.0\times10^{-1}$ | $6.5\times10^{2}$ | $2.7\times10^{-1}$ |
| I540G | 3.8 | **0** | $5.2\times10^{1}$ | **$2.6\times10^{-3}$** | $6.1\times10^{4}$ | **0** | $7.1\times10^{3}$ | **$7.7\times10^{-3}$** | ≤ $9.9\times10^{-1}$ | **0** | $3.5\times10^{2}$ | **$1.0\times10^{-3}$** |
| I540V | $7.4\times10^{1}$ | $2.9\times10^{-2}$ | $8.7\times10^{1}$ | $8.9\times10^{-1}$ | $7.9\times10^{5}$ | $4.5\times10^{-2}$ | $4.0\times10^{5}$ | $1.0\times10^{-1}$ | 6.9 | $1.5\times10^{-2}$ | $6.0\times10^{2}$ | $5.2\times10^{-1}$ |
| T541G | $3.9\times10^{1}$ | **0** | $6.1\times10^{1}$ | $2.2\times10^{-2}$ | $6.6\times10^{5}$ | $2.7\times10^{-2}$ | $2.4\times10^{5}$ | $2.2\times10^{-2}$ | 8.9 | $3.2\times10^{-2}$ | $5.2\times10^{2}$ | **$1.9\times10^{-3}$** |
| T541V | $1.2\times10^{2}$ | $9.3\times10^{-1}$ | $8.9\times10^{1}$ | $7.3\times10^{-1}$ | $1.4\times10^{6}$ | $6.7\times10^{-1}$ | $5.5\times10^{5}$ | $5.5\times10^{-1}$ | $1.1\times10^{1}$ | $2.9\times10^{-1}$ | $6.6\times10^{2}$ | $1.2\times10^{-1}$ |
| E542G | $3.2\times10^{1}$ | **$9.0\times10^{-4}$** | $7.0\times10^{1}$ | $2.1\times10^{-1}$ | $5.0\times10^{5}$ | $1.5\times10^{-2}$ | $2.1\times10^{5}$ | $1.8\times10^{-2}$ | 8.1 | $1.4\times10^{-2}$ | $6.7\times10^{2}$ | $1.4\times10^{-1}$ |
| E542V | $6.1\times10^{1}$ | $2.1\times10^{-2}$ | $7.7\times10^{1}$ | $4.7\times10^{-1}$ | $8.1\times10^{5}$ | $9.2\times10^{-2}$ | $2.5\times10^{5}$ | **$8.7\times10^{-3}$** | 9.2 | $7.3\times10^{-2}$ | $5.3\times10^{2}$ | $2.5\times10^{-2}$ |
| V543A | $2.5\times10^{2}$ | **$1.\times10^{-4}$** | $9.6\times10^{1}$ | $3.3\times10^{-1}$ | $2.2\times10^{6}$ | $1.5\times10^{-2}$ | $5.7\times10^{5}$ | $7.1\times10^{-1}$ | $1.7\times10^{1}$ | $8.5\times10^{-1}$ | $6.6\times10^{2}$ | $1.6\times10^{-1}$ |
| V543G | $6.4\times10^{1}$ | $1.5\times10^{-2}$ | $7.1\times10^{1}$ | $1.9\times10^{-1}$ | $8.8\times10^{5}$ | $1.0\times10^{-1}$ | $1.3\times10^{5}$ | **$1.4\times10^{-3}$** | 7.3 | **$1.4\times10^{-3}$** | $5.9\times10^{2}$ | $2.7\times10^{-1}$ |
| I544G | $4.1\times10^{1}$ | **$5.8\times10^{-3}$** | $4.8\times10^{1}$ | $1.2\times10^{-2}$ | $8.5\times10^{5}$ | $1.9\times10^{-1}$ | $3.0\times10^{5}$ | $1.6\times10^{-2}$ | 8.1 | $1.4\times10^{-2}$ | $4.2\times10^{2}$ | **$8.0\times10^{-4}$** |
| I544V | $5.2\times10^{1}$ | **$1.5\times10^{-3}$** | $8.8\times10^{1}$ | $7.9\times10^{-1}$ | $6.2\times10^{5}$ | **$9.4\times10^{-3}$** | $3.6\times10^{5}$ | $4.9\times10^{-2}$ | 7.4 | **$6.3\times10^{-3}$** | $5.9\times10^{2}$ | $5.8\times10^{-1}$ |
| G545A | $1.2\times10^{2}$ | $9.6\times10^{-1}$ | $7.5\times10^{1}$ | $3.8\times10^{-1}$ | $2.2\times10^{6}$ | $3.9\times10^{-2}$ | $5.2\times10^{5}$ | $4.8\times10^{-1}$ | $2.1\times10^{1}$ | $1.3\times10^{-1}$ | $5.6\times10^{2}$ | $2.0\times10^{-1}$ |
| G545V | $2.0\times10^{2}$ | **0** | $9.2\times10^{1}$ | $4.0\times10^{-1}$ | $2.0\times10^{6}$ | $3.3\times10^{-2}$ | $6.2\times10^{5}$ | $9.2\times10^{-1}$ | $1.4\times10^{1}$ | $6.9\times10^{-1}$ | $6.5\times10^{2}$ | $2.5\times10^{-1}$ |
| R546G | $1.7\times10^{2}$ | $4.3\times10^{-2}$ | $8.1\times10^{1}$ | $7.4\times10^{-1}$ | $1.9\times10^{6}$ | $1.8\times10^{-1}$ | $4.2\times10^{5}$ | $1.3\times10^{-1}$ | $1.3\times10^{1}$ | $4.6\times10^{-1}$ | $5.9\times10^{2}$ | $3.5\times10^{-1}$ |
| R546V | $1.2\times10^{2}$ | $8.8\times10^{-1}$ | $9.6\times10^{1}$ | $2.8\times10^{-1}$ | $1.3\times10^{6}$ | $7.9\times10^{-1}$ | $5.3\times10^{5}$ | $3.8\times10^{-1}$ | $1.2\times10^{1}$ | $2.8\times10^{-1}$ | $6.4\times10^{2}$ | $4.3\times10^{-1}$ |

#### Additional parameters and FCs

| Variant | $f_a$ | $f_a$ *p*–value | $(k_{cat}/K_M)^{MeP}_{chem.}$ ($M^{-1}s^{-1}$) | $(k_{cat}/K_M)^{MeP}_{chem.}$ *p*–value | FC1 | FC1 *p*–value | FC2 | FC3 | FC2/3 *p*–value | FC4 ($s^{-1}$) | FC4 *p*–value |
|---|---|---|---|---|---|---|---|---|---|---|---|
| Q21G | 1.4 | $1.2\times10^{-1}$ | $4.4\times10^{5}$ | $2.7\times10^{-1}$ | $9.5\times10^{-1}$ | $7.4\times10^{-1}$ | N/A | 1.0 | $3.8\times10^{-1}$ | $1.9\times10^{2}$ | $3.4\times10^{-1}$ |
| Q21V | 1.5 | $3.8\times10^{-2}$ | $2.8\times10^{5}$ | $5.6\times10^{-2}$ | 1.7 | $1.1\times10^{-1}$ | N/A | 1.1 | $2.0\times10^{-1}$ | $1.6\times10^{2}$ | $6.1\times10^{-1}$ |
| K22G | 1.1 | $7.3\times10^{-1}$ | $4.6\times10^{5}$ | $5.1\times10^{-1}$ | $9.7\times10^{-1}$ | $9.0\times10^{-1}$ | $8.7\times10^{-1}$ | N/A | $1.4\times10^{-2}$ | $7.2\times10^{1}$ | $2.6\times10^{-1}$ |
| K22V | $8.2\times10^{-1}$ | $3.9\times10^{-1}$ | $4.9\times10^{5}$ | $4.7\times10^{-1}$ | 1.4 | $2.6\times10^{-1}$ | N/A | 1.2 | $1.6\times10^{-2}$ | $1.8\times10^{2}$ | $3.0\times10^{-1}$ |
| T23G | 1.0 | $9.3\times10^{-1}$ | $6.8\times10^{5}$ | $8.2\times10^{-1}$ | 1.1 | $7.5\times10^{-1}$ | N/A | 1.0 | $5.1\times10^{-1}$ | $1.3\times10^{2}$ | $7.3\times10^{-1}$ |
| T23V | 1.4 | $2.3\times10^{-1}$ | $2.4\times10^{5}$ | $8.7\times10^{-2}$ | 2.1 | $1.2\times10^{-2}$ | N/A | 1.0 | $5.6\times10^{-1}$ | $1.6\times10^{2}$ | $7.1\times10^{-1}$ |
| N24G | 1.3 | $4.6\times10^{-1}$ | $4.4\times10^{5}$ | $4.4\times10^{-1}$ | $8.7\times10^{-1}$ | $6.8\times10^{-1}$ | $7.9\times10^{-1}$ | N/A | **$2.0\times10^{-3}$** | $9.2\times10^{1}$ | $3.7\times10^{-1}$ |
| N24V | $9.9\times10^{-1}$ | $9.1\times10^{-1}$ | $4.3\times10^{5}$ | $3.3\times10^{-1}$ | 1.7 | $6.8\times10^{-2}$ | N/A | 1.1 | $2.3\times10^{-1}$ | $1.6\times10^{2}$ | $6.8\times10^{-1}$ |
| A25G | 1.9 | $1.9\times10^{-2}$ | $4.5\times10^{5}$ | $4.2\times10^{-1}$ | 1.1 | $7.8\times10^{-1}$ | $9.4\times10^{-1}$ | N/A | $1.6\times10^{-1}$ | $1.3\times10^{2}$ | $8.1\times10^{-1}$ |
| A25V | 1.4 | $1.1\times10^{-1}$ | $4.8\times10^{5}$ | $5.0\times10^{-1}$ | 1.0 | $9.4\times10^{-1}$ | N/A | 1.1 | $1.6\times10^{-1}$ | $1.9\times10^{2}$ | $2.8\times10^{-1}$ |
| V26A | $5.2\times10^{-1}$ | $1.2\times10^{-2}$ | $3.6\times10^{5}$ | $2.3\times10^{-1}$ | 1.3 | $4.3\times10^{-1}$ | $9.7\times10^{-1}$ | N/A | $6.3\times10^{-1}$ | $1.1\times10^{2}$ | $5.2\times10^{-1}$ |
| V26G | $1.9\times10^{-1}$ | **0** | $2.1\times10^{5}$ | **$4.0\times10^{-2}$** | 3.7 | **$1.8\times10^{-3}$** | $8.8\times10^{-1}$ | N/A | $1.7\times10^{-2}$ | $1.3\times10^{2}$ | $8.4\times10^{-1}$ |
| P27G | $3.0\times10^{-1}$ | **$2.0\times10^{-3}$** | $9.7\times10^{5}$ | $1.2\times10^{-1}$ | 1.8 | $5.4\times10^{-2}$ | $9.0\times10^{-1}$ | N/A | $4.3\times10^{-2}$ | $2.6\times10^{2}$ | $9.5\times10^{-2}$ |
| P27V | 1.1 | $7.5\times10^{-1}$ | $2.4\times10^{5}$ | $7.1\times10^{-2}$ | 1.9 | $5.5\times10^{-2}$ | N/A | 1.0 | $6.9\times10^{-1}$ | $1.5\times10^{2}$ | $8.8\times10^{-1}$ |
| R28G | $7.6\times10^{-1}$ | $5.1\times10^{-1}$ | $4.7\times10^{5}$ | $7.1\times10^{-1}$ | 1.0 | $9.0\times10^{-1}$ | $7.1\times10^{-1}$ | N/A | **$1.3\times10^{-3}$** | $6.0\times10^{1}$ | $2.8\times10^{-1}$ |
| R28V | $6.6\times10^{-1}$ | $8.7\times10^{-2}$ | $2.0\times10^{5}$ | $6.8\times10^{-2}$ | 2.9 | **$5.7\times10^{-3}$** | N/A | 1.0 | $9.1\times10^{-1}$ | $1.3\times10^{2}$ | $8.7\times10^{-1}$ |
| P29G | $1.3\times10^{-1}$ | **0** | $1.7\times10^{5}$ | **$4.9\times10^{-2}$** | – | – | $8.0\times10^{-1}$ | N/A | **0** | $9.1\times10^{1}$ | $2.7\times10^{-1}$ |
| P29V | $1.4\times10^{-1}$ | **0** | $9.1\times10^{5}$ | $1.8\times10^{-1}$ | – | – | $7.1\times10^{-1}$ | N/A | **$1.\times10^{-4}$** | $1.2\times10^{2}$ | $7.5\times10^{-1}$ |
| K30G | $8.5\times10^{-1}$ | $5.8\times10^{-1}$ | $3.1\times10^{5}$ | $1.5\times10^{-1}$ | 1.6 | $1.3\times10^{-1}$ | $8.7\times10^{-1}$ | N/A | $1.4\times10^{-2}$ | $6.4\times10^{1}$ | $1.1\times10^{-1}$ |
| K30V | $6.7\times10^{-1}$ | $1.3\times10^{-1}$ | $1.9\times10^{5}$ | $5.9\times10^{-2}$ | 2.2 | $2.3\times10^{-2}$ | $8.7\times10^{-1}$ | N/A | $4.5\times10^{-2}$ | $1.6\times10^{2}$ | $7.7\times10^{-1}$ |
| L31G | $\leq 3.6\times10^{-3}$ | **0** | $\leq 3.8\times10^{5}$ | $2.1\times10^{-1}$ | – | – | – | – | – | – | – |
| L31V | $9.8\times10^{-1}$ | $8.9\times10^{-1}$ | $3.0\times10^{5}$ | $6.7\times10^{-2}$ | 1.5 | $1.6\times10^{-1}$ | N/A | 1.0 | $7.4\times10^{-1}$ | $1.6\times10^{2}$ | $6.6\times10^{-1}$ |
| V32A | $9.9\times10^{-1}$ | $9.4\times10^{-1}$ | $2.6\times10^{5}$ | $1.2\times10^{-1}$ | 2.4 | $1.5\times10^{-2}$ | N/A | 1.0 | $9.7\times10^{-1}$ | $2.0\times10^{2}$ | $2.0\times10^{-1}$ |
| V32G | $5.2\times10^{-1}$ | $9.7\times10^{-2}$ | $3.2\times10^{5}$ | $2.9\times10^{-1}$ | 1.2 | $6.8\times10^{-1}$ | $7.5\times10^{-1}$ | N/A | **$2.1\times10^{-3}$** | $1.2\times10^{2}$ | $7.5\times10^{-1}$ |
| V33A | 1.3 | $2.8\times10^{-1}$ | $1.6\times10^{5}$ | $5.5\times10^{-2}$ | 1.9 | $1.4\times10^{-1}$ | $9.7\times10^{-1}$ | N/A | $4.4\times10^{-1}$ | $1.6\times10^{2}$ | $7.0\times10^{-1}$ |
| V33G | $3.9\times10^{-1}$ | **0** | $5.9\times10^{5}$ | $9.1\times10^{-1}$ | 1.1 | $7.7\times10^{-1}$ | $8.1\times10^{-1}$ | N/A | **$2.0\times10^{-4}$** | $1.0\times10^{2}$ | $4.3\times10^{-1}$ |
| G34A | $8.3\times10^{-1}$ | $4.3\times10^{-1}$ | $8.4\times10^{5}$ | $2.8\times10^{-1}$ | $8.8\times10^{-1}$ | $6.4\times10^{-1}$ | N/A | 1.0 | $4.6\times10^{-1}$ | $1.4\times10^{2}$ | $9.1\times10^{-1}$ |
| G34V | $\leq 1.9\times10^{-1}$ | **0** | $\leq 1.7\times10^{5}$ | $6.4\times10^{-2}$ | – | – | $7.1\times10^{-1}$ | N/A | **0** | $\geq 1.2\times10^{2}$ | $6.8\times10^{-1}$ |
| L35G | $\leq 1.7\times10^{-2}$ | **0** | $\leq 9.4\times10^{4}$ | $6.5\times10^{-2}$ | – | – | N/A | 1.4 | **$1.0\times10^{-3}$** | $\geq 1.2\times10^{2}$ | $7.5\times10^{-1}$ |
| L35V | 1.4 | $9.2\times10^{-2}$ | $3.4\times10^{5}$ | $9.5\times10^{-2}$ | 1.3 | $2.2\times10^{-1}$ | N/A | 1.1 | $3.8\times10^{-2}$ | $1.2\times10^{2}$ | $6.2\times10^{-1}$ |
| V36A | 1.3 | $2.6\times10^{-1}$ | $2.7\times10^{5}$ | $9.4\times10^{-2}$ | 1.4 | $4.1\times10^{-1}$ | $8.4\times10^{-1}$ | N/A | $5.2\times10^{-2}$ | $1.2\times10^{2}$ | $6.5\times10^{-1}$ |
| V36G | $5.5\times10^{-1}$ | **$4.0\times10^{-3}$** | $2.5\times10^{5}$ | $7.1\times10^{-2}$ | 1.5 | $1.7\times10^{-1}$ | $3.3\times10^{-1}$ | N/A | **0** | $3.6\times10^{1}$ | **$2.6\times10^{-2}$** |
| V37A | 1.3 | $3.1\times10^{-1}$ | $2.8\times10^{5}$ | $8.4\times10^{-2}$ | 1.3 | $3.9\times10^{-1}$ | N/A | 1.2 | $1.1\times10^{-2}$ | $1.8\times10^{2}$ | $4.3\times10^{-1}$ |
| V37G | $8.4\times10^{-1}$ | $5.4\times10^{-1}$ | $2.4\times10^{5}$ | $1.0\times10^{-1}$ | 1.1 | $7.9\times10^{-1}$ | $7.9\times10^{-1}$ | N/A | **$3.0\times10^{-4}$** | $1.3\times10^{2}$ | $8.7\times10^{-1}$ |
| D38G | $\leq 6.0\times10^{-1}$ | $1.1\times10^{-2}$ | $\leq 1.0\times10^{4}$ | **$2.0\times10^{-3}$** | $\leq 1.3\times10^{1}$ | **$1.7\times10^{-4}$** | $\leq 2.1\times10^{-1}$ | N/A | **0** | $\geq 7.3$ | **$2.0\times10^{-3}$** |
| D38V | – | – | – | – | – | – | – | – | – | – | – |
| Q39G | 1.3 | $2.5\times10^{-1}$ | $2.7\times10^{4}$ | **$6.0\times10^{-3}$** | 2.8 | **$8.5\times10^{-3}$** | N/A | 2.2 | **$1.\times10^{-4}$** | $5.4\times10^{1}$ | $9.6\times10^{-2}$ |
| Q39V | $\leq 3.3\times10^{-2}$ | **0** | $\leq 4.7\times10^{4}$ | **$1.6\times10^{-2}$** | – | – | $4.2\times10^{-1}$ | N/A | **$3.0\times10^{-4}$** | $\geq 6.7\times10^{1}$ | $1.2\times10^{-1}$ |
| M40G | $2.2\times10^{-1}$ | **0** | $5.7\times10^{5}$ | $8.4\times10^{-1}$ | 1.5 | $2.5\times10^{-1}$ | $6.2\times10^{-1}$ | N/A | **0** | $1.1\times10^{2}$ | $5.1\times10^{-1}$ |
| M40V | $5.1\times10^{-2}$ | **0** | $6.1\times10^{4}$ | **$2.6\times10^{-2}$** | – | – | $5.2\times10^{-1}$ | N/A | **$1.\times10^{-4}$** | $8.5\times10^{1}$ | $2.1\times10^{-1}$ |
| R41G | 2.1 | $5.6\times10^{-2}$ | $2.1\times10^{5}$ | $1.4\times10^{-1}$ | 1.4 | $2.2\times10^{-1}$ | $8.9\times10^{-1}$ | N/A | $1.1\times10^{-1}$ | $1.1\times10^{2}$ | $7.2\times10^{-1}$ |
| R41V | $8.4\times10^{-1}$ | $4.1\times10^{-1}$ | $2.6\times10^{5}$ | $8.4\times10^{-2}$ | – | – | $5.7\times10^{-1}$ | N/A | **$3.0\times10^{-4}$** | $7.6\times10^{1}$ | $1.0\times10^{-1}$ |
| W42G | 1.0 | $9.5\times10^{-1}$ | $2.5\times10^{5}$ | $1.4\times10^{-1}$ | $9.1\times10^{-1}$ | $6.7\times10^{-1}$ | $6.3\times10^{-1}$ | N/A | **$1.\times10^{-4}$** | $5.2\times10^{1}$ | $1.6\times10^{-1}$ |
| W42V | 1.6 | $4.2\times10^{-2}$ | $4.7\times10^{5}$ | $5.2\times10^{-1}$ | $5.2\times10^{-1}$ | $4.3\times10^{-2}$ | $8.1\times10^{-1}$ | N/A | $2.3\times10^{-2}$ | $1.4\times10^{2}$ | $9.3\times10^{-1}$ |
| D43G | $1.8\times10^{-1}$ | **0** | $1.0\times10^{5}$ | **$3.9\times10^{-2}$** | 2.9 | $1.4\times10^{-2}$ | $4.9\times10^{-1}$ | N/A | **0** | $5.6\times10^{1}$ | $5.4\times10^{-2}$ |
| D43V | $1.7\times10^{-1}$ | **0** | $8.6\times10^{4}$ | **$3.1\times10^{-2}$** | 5.2 | **$3.0\times10^{-3}$** | $4.5\times10^{-1}$ | N/A | **0** | $5.0\times10^{1}$ | $6.2\times10^{-2}$ |
| Y44G | $6.0\times10^{-1}$ | $5.8\times10^{-2}$ | $4.3\times10^{5}$ | $3.7\times10^{-1}$ | $8.3\times10^{-1}$ | $5.7\times10^{-1}$ | N/A | 1.1 | $2.2\times10^{-2}$ | $1.1\times10^{2}$ | $6.3\times10^{-1}$ |
| Y44V | 1.2 | $4.7\times10^{-1}$ | $2.3\times10^{5}$ | **$3.6\times10^{-2}$** | 1.3 | $4.3\times10^{-1}$ | N/A | 1.0 | $5.8\times10^{-1}$ | $9.8\times10^{1}$ | $2.9\times10^{-1}$ |
| L45G | $7.5\times10^{-1}$ | $3.4\times10^{-1}$ | $1.1\times10^{5}$ | $6.9\times10^{-2}$ | 2.6 | $1.8\times10^{-2}$ | $8.7\times10^{-1}$ | N/A | $3.8\times10^{-2}$ | $9.9\times10^{1}$ | $5.1\times10^{-1}$ |
| L45V | $4.4\times10^{-1}$ | **0** | $3.1\times10^{5}$ | $1.9\times10^{-1}$ | 1.9 | $7.3\times10^{-2}$ | $9.3\times10^{-1}$ | N/A | $1.9\times10^{-1}$ | $1.3\times10^{2}$ | $8.5\times10^{-1}$ |
| Y46G | $2.8\times10^{-1}$ | **0** | $9.4\times10^{5}$ | $1.4\times10^{-1}$ | – | – | $5.6\times10^{-1}$ | N/A | **$1.\times10^{-4}$** | $8.8\times10^{1}$ | $3.3\times10^{-1}$ |
| Y46V | 1.1 | $7.0\times10^{-1}$ | $3.1\times10^{5}$ | $9.6\times10^{-2}$ | 1.7 | $1.0\times10^{-1}$ | $9.1\times10^{-1}$ | N/A | $6.6\times10^{-2}$ | $1.3\times10^{2}$ | $8.1\times10^{-1}$ |
| R47G | $5.9\times10^{-1}$ | $8.7\times10^{-2}$ | $3.1\times10^{5}$ | $2.5\times10^{-1}$ | 1.4 | $2.6\times10^{-1}$ | $7.8\times10^{-1}$ | N/A | **0** | $7.5\times10^{1}$ | $2.6\times10^{-1}$ |
| R47V | 1.2 | $3.9\times10^{-1}$ | $8.1\times10^{4}$ | **$1.1\times10^{-2}$** | 4.6 | **$1.1\times10^{-3}$** | $9.6\times10^{-1}$ | N/A | $3.5\times10^{-1}$ | $1.3\times10^{2}$ | $6.8\times10^{-1}$ |
| Y48G | $2.2\times10^{-1}$ | **0** | $1.4\times10^{5}$ | **$3.0\times10^{-2}$** | 2.3 | $2.7\times10^{-2}$ | $6.6\times10^{-1}$ | N/A | **0** | $6.7\times10^{1}$ | $1.1\times10^{-1}$ |
| Y48V | $\leq 1.9\times10^{-1}$ | **0** | $\leq 6.0\times10^{5}$ | $9.5\times10^{-1}$ | – | – | $9.7\times10^{-1}$ | N/A | $5.0\times10^{-1}$ | $\geq 1.6\times10^{2}$ | $7.0\times10^{-1}$ |
| Y49G | 1.1 | $7.6\times10^{-1}$ | $2.8\times10^{5}$ | $1.1\times10^{-1}$ | $7.1\times10^{-1}$ | $2.6\times10^{-1}$ | $8.2\times10^{-1}$ | N/A | **$2.3\times10^{-3}$** | $1.0\times10^{2}$ | $4.9\times10^{-1}$ |
| Y49V | 1.0 | $8.2\times10^{-1}$ | $1.8\times10^{5}$ | **$4.9\times10^{-2}$** | 2.5 | $1.1\times10^{-2}$ | $9.1\times10^{-1}$ | N/A | $1.0\times10^{-1}$ | $1.1\times10^{2}$ | $4.3\times10^{-1}$ |
| S50G | $\leq 1.5$ | $9.4\times10^{-2}$ | $\leq 2.0\times10^{5}$ | $6.5\times10^{-2}$ | $\leq 2.7$ | **$7.7\times10^{-3}$** | N/A | 1.1 | $7.8\times10^{-2}$ | $\geq 1.0\times10^{2}$ | $4.1\times10^{-1}$ |
| S50V | 1.3 | $2.4\times10^{-1}$ | $1.7\times10^{5}$ | **$3.9\times10^{-2}$** | 2.6 | **$3.0\times10^{-3}$** | N/A | 1.0 | $5.0\times10^{-1}$ | $1.5\times10^{2}$ | $7.8\times10^{-1}$ |
| K51G | $6.1\times10^{-1}$ | $4.1\times10^{-2}$ | $1.2\times10^{5}$ | **$1.8\times10^{-2}$** | 5.0 | **$8.0\times10^{-5}$** | N/A | 1.1 | $2.6\times10^{-1}$ | $1.2\times10^{2}$ | $6.2\times10^{-1}$ |
| K51V | $5.8\times10^{-1}$ | $3.4\times10^{-2}$ | $2.3\times10^{5}$ | $7.3\times10^{-2}$ | 2.1 | $2.0\times10^{-2}$ | $8.9\times10^{-1}$ | N/A | $6.9\times10^{-2}$ | $1.1\times10^{2}$ | $5.1\times10^{-1}$ |
| Y52G | $5.9\times10^{-1}$ | **$9.0\times10^{-3}$** | $2.0\times10^{5}$ | **$2.4\times10^{-2}$** | 1.1 | $7.1\times10^{-1}$ | $9.2\times10^{-1}$ | N/A | $4.7\times10^{-2}$ | $1.2\times10^{2}$ | $5.9\times10^{-1}$ |
| Y52V | 1.5 | $7.2\times10^{-2}$ | $1.3\times10^{5}$ | **$1.7\times10^{-2}$** | 3.1 | **$3.4\times10^{-4}$** | N/A | 1.1 | $6.7\times10^{-2}$ | $2.0\times10^{2}$ | $1.2\times10^{-1}$ |
| G53A | $\leq 8.6\times10^{-1}$ | $5.0\times10^{-1}$ | $\leq 4.4\times10^{5}$ | $3.4\times10^{-1}$ | $\leq 1.9$ | $2.5\times10^{-2}$ | N/A | 1.0 | $4.6\times10^{-1}$ | $\geq 1.2\times10^{2}$ | $6.9\times10^{-1}$ |
| G53V | $4.3\times10^{-1}$ | **$6.0\times10^{-3}$** | $1.1\times10^{6}$ | $6.1\times10^{-2}$ | – | – | $9.7\times10^{-1}$ | N/A | $6.8\times10^{-1}$ | $1.7\times10^{2}$ | $5.0\times10^{-1}$ |
| E54G | 1.3 | $2.6\times10^{-1}$ | $3.4\times10^{5}$ | $2.2\times10^{-1}$ | 1.3 | $3.5\times10^{-1}$ | $9.8\times10^{-1}$ | N/A | $6.9\times10^{-1}$ | $1.3\times10^{2}$ | $8.3\times10^{-1}$ |
| E54V | 1.4 | $1.2\times10^{-1}$ | $4.0\times10^{5}$ | $1.8\times10^{-1}$ | 1.0 | $8.4\times10^{-1}$ | N/A | 1.1 | $3.5\times10^{-2}$ | $1.8\times10^{2}$ | $3.7\times10^{-1}$ |
| G55A | 1.5 | $1.3\times10^{-1}$ | $3.5\times10^{5}$ | $2.5\times10^{-1}$ | 1.6 | $1.7\times10^{-1}$ | N/A | 1.0 | $5.3\times10^{-1}$ | $6.4\times10^{1}$ | $1.3\times10^{-1}$ |
| G55V | 1.7 | $9.1\times10^{-2}$ | $1.3\times10^{5}$ | $9.8\times10^{-2}$ | 3.3 | **$6.5\times10^{-3}$** | N/A | 1.1 | $2.4\times10^{-1}$ | $1.4\times10^{2}$ | $9.7\times10^{-1}$ |
| G56A | $5.1\times10^{-1}$ | $3.3\times10^{-2}$ | $2.6\times10^{5}$ | $1.5\times10^{-1}$ | 1.2 | $5.7\times10^{-1}$ | N/A | 1.0 | $8.1\times10^{-1}$ | $1.3\times10^{2}$ | $8.9\times10^{-1}$ |
| G56V | $\leq 1.4\times10^{-3}$ | **$3.0\times10^{-3}$** | $\leq 1.3\times10^{6}$ | $1.3\times10^{-1}$ | – | – | – | – | – | – | – |
| F57G | $\leq 1.6\times10^{-3}$ | **0** | $\leq 7.5\times10^{5}$ | $4.5\times10^{-1}$ | – | – | – | – | – | – | – |
| F57V | 1.4 | $1.2\times10^{-1}$ | $8.6\times10^{4}$ | **$3.8\times10^{-2}$** | 3.3 | **$9.8\times10^{-3}$** | $9.9\times10^{-1}$ | N/A | $6.8\times10^{-1}$ | $1.3\times10^{2}$ | $7.9\times10^{-1}$ |
| K58G | $6.2\times10^{-1}$ | $1.1\times10^{-1}$ | $1.8\times10^{5}$ | $9.3\times10^{-2}$ | 2.7 | **$4.3\times10^{-3}$** | $7.5\times10^{-1}$ | N/A | **$3.0\times10^{-4}$** | $7.0\times10^{1}$ | $2.6\times10^{-1}$ |

| Variant | $f_a$ | $f_a$ $p$–value | $(k_{cat}/K_M)^{MeP}_{chem.}$ ($M^{-1}s^{-1}$) | $(k_{cat}/K_M)^{MeP}_{chem.}$ $p$–value | FC1 | FC1 $p$–value | FC2 | FC3 | FC2/3 $p$–value | FC4 ($s^{-1}$) | FC4 $p$–value |
|---|---|---|---|---|---|---|---|---|---|---|---|
| K58V | $7.6\times10^{-1}$ | $2.9\times10^{-1}$ | $2.8\times10^{5}$ | $1.5\times10^{-1}$ | 2.3 | $2.9\times10^{-2}$ | $9.9\times10^{-1}$ | N/A | $8.9\times10^{-1}$ | $1.9\times10^{2}$ | $3.7\times10^{-1}$ |
| R59G | $1.5\times10^{-1}$ | **$1.0\times10^{-3}$** | $2.3\times10^{5}$ | $1.1\times10^{-1}$ | – | – | $9.4\times10^{-1}$ | N/A | $2.5\times10^{-1}$ | $1.2\times10^{2}$ | $7.5\times10^{-1}$ |
| R59V | $3.1\times10^{-1}$ | **$1.0\times10^{-3}$** | $6.2\times10^{4}$ | **$1.7\times10^{-2}$** | 8.3 | **$6.1\times10^{-4}$** | $8.0\times10^{-1}$ | N/A | $2.4\times10^{-2}$ | $9.9\times10^{1}$ | $4.4\times10^{-1}$ |
| M60G | $2.8\times10^{-1}$ | **0** | $3.4\times10^{5}$ | $2.5\times10^{-1}$ | $9.1\times10^{-1}$ | $7.7\times10^{-1}$ | N/A | 1.1 | $2.5\times10^{-2}$ | $1.4\times10^{2}$ | $9.8\times10^{-1}$ |
| M60V | 1.2 | $5.3\times10^{-1}$ | $3.2\times10^{5}$ | $1.5\times10^{-1}$ | $8.5\times10^{-1}$ | $6.3\times10^{-1}$ | N/A | 1.3 | **$2.0\times10^{-4}$** | $1.9\times10^{2}$ | $2.4\times10^{-1}$ |
| L61G | $9.0\times10^{-1}$ | $7.2\times10^{-1}$ | $2.7\times10^{5}$ | $1.5\times10^{-1}$ | $9.3\times10^{-1}$ | $8.5\times10^{-1}$ | $7.6\times10^{-1}$ | N/A | **$4.0\times10^{-4}$** | $1.8\times10^{2}$ | $5.6\times10^{-1}$ |
| L61V | $5.4\times10^{-1}$ | **$1.0\times10^{-3}$** | $4.9\times10^{5}$ | $4.6\times10^{-1}$ | $8.0\times10^{-1}$ | $4.8\times10^{-1}$ | $9.4\times10^{-1}$ | N/A | $8.2\times10^{-2}$ | $1.4\times10^{2}$ | $9.4\times10^{-1}$ |
| N62G | $8.4\times10^{-1}$ | $4.8\times10^{-1}$ | $3.9\times10^{5}$ | $2.7\times10^{-1}$ | 2.0 | $1.1\times10^{-2}$ | $8.1\times10^{-1}$ | N/A | $3.0\times10^{-2}$ | $1.5\times10^{2}$ | $8.7\times10^{-1}$ |
| N62V | 1.1 | $6.4\times10^{-1}$ | $1.0\times10^{5}$ | **$1.3\times10^{-2}$** | 3.8 | **$1.1\times10^{-3}$** | N/A | 1.1 | $1.4\times10^{-1}$ | $1.8\times10^{2}$ | $3.4\times10^{-1}$ |
| T63G | 1.4 | $3.9\times10^{-1}$ | $2.2\times10^{5}$ | $1.4\times10^{-1}$ | 2.1 | $1.2\times10^{-2}$ | $8.2\times10^{-1}$ | N/A | **$1.5\times10^{-3}$** | $8.8\times10^{1}$ | $3.5\times10^{-1}$ |
| T63V | $8.3\times10^{-1}$ | $4.0\times10^{-1}$ | $4.6\times10^{5}$ | $4.1\times10^{-1}$ | 1.0 | $9.8\times10^{-1}$ | N/A | 1.1 | $2.9\times10^{-1}$ | $1.5\times10^{2}$ | $8.8\times10^{-1}$ |
| G64A | $1.1\times10^{-1}$ | **0** | $2.7\times10^{5}$ | $8.2\times10^{-2}$ | – | – | $9.6\times10^{-1}$ | N/A | $3.0\times10^{-1}$ | $1.3\times10^{2}$ | $8.0\times10^{-1}$ |
| G64V | – | – | – | – | – | – | – | – | – | – | – |
| Y65G | $4.2\times10^{-1}$ | **0** | $1.8\times10^{5}$ | $6.9\times10^{-2}$ | 2.7 | **$7.2\times10^{-3}$** | $8.3\times10^{-1}$ | N/A | **$2.5\times10^{-3}$** | $1.4\times10^{2}$ | $9.7\times10^{-1}$ |
| Y65V | 1.0 | $9.6\times10^{-1}$ | $2.4\times10^{5}$ | $5.9\times10^{-2}$ | 1.6 | $1.2\times10^{-1}$ | N/A | 1.1 | $2.2\times10^{-1}$ | $1.5\times10^{2}$ | $8.6\times10^{-1}$ |
| S66G | $7.9\times10^{-1}$ | $3.2\times10^{-1}$ | $3.9\times10^{5}$ | $2.2\times10^{-1}$ | 1.6 | $9.6\times10^{-2}$ | $7.7\times10^{-1}$ | N/A | **0** | $1.1\times10^{2}$ | $5.2\times10^{-1}$ |
| S66V | $8.1\times10^{-1}$ | $3.3\times10^{-1}$ | $4.4\times10^{5}$ | $2.7\times10^{-1}$ | $9.2\times10^{-1}$ | $7.6\times10^{-1}$ | $9.6\times10^{-1}$ | N/A | $2.3\times10^{-1}$ | $1.5\times10^{2}$ | $7.4\times10^{-1}$ |
| L67G | $5.2\times10^{-1}$ | $5.0\times10^{-2}$ | $2.2\times10^{5}$ | $1.4\times10^{-1}$ | 3.0 | **$3.6\times10^{-3}$** | $7.6\times10^{-1}$ | N/A | **$3.0\times10^{-4}$** | $8.1\times10^{1}$ | $3.5\times10^{-1}$ |
| L67V | $7.6\times10^{-1}$ | $2.0\times10^{-1}$ | $2.8\times10^{5}$ | $1.2\times10^{-1}$ | 1.5 | $2.8\times10^{-1}$ | $9.8\times10^{-1}$ | N/A | $5.7\times10^{-1}$ | $1.3\times10^{2}$ | $7.3\times10^{-1}$ |
| N68G | $5.9\times10^{-1}$ | $3.1\times10^{-2}$ | $2.7\times10^{5}$ | $1.4\times10^{-1}$ | 2.2 | $2.5\times10^{-2}$ | N/A | 1.0 | $6.0\times10^{-1}$ | $1.3\times10^{2}$ | $8.2\times10^{-1}$ |
| N68V | $1.6\times10^{-1}$ | **0** | $4.2\times10^{5}$ | $4.1\times10^{-1}$ | – | – | $7.9\times10^{-1}$ | N/A | **0** | $1.1\times10^{2}$ | $5.5\times10^{-1}$ |
| N69G | $2.4\times10^{-1}$ | **0** | $4.9\times10^{5}$ | $6.2\times10^{-1}$ | 2.1 | $4.8\times10^{-2}$ | $7.4\times10^{-1}$ | N/A | **$2.0\times10^{-4}$** | $8.5\times10^{1}$ | $3.5\times10^{-1}$ |
| N69V | ≤ $5.0\times10^{-1}$ | **$7.0\times10^{-3}$** | ≤ $1.6\times10^{5}$ | $6.2\times10^{-2}$ | ≤ 2.9 | $1.4\times10^{-2}$ | $8.1\times10^{-1}$ | N/A | **$3.7\times10^{-3}$** | ≥ $1.2\times10^{2}$ | $6.1\times10^{-1}$ |
| V70A | 1.7 | $5.8\times10^{-2}$ | $4.9\times10^{5}$ | $5.8\times10^{-1}$ | $8.2\times10^{-1}$ | $5.6\times10^{-1}$ | $8.2\times10^{-1}$ | N/A | $2.8\times10^{-2}$ | $1.5\times10^{2}$ | $9.1\times10^{-1}$ |
| V70G | $4.4\times10^{-2}$ | **0** | $2.3\times10^{5}$ | $1.3\times10^{-1}$ | – | – | $5.7\times10^{-1}$ | N/A | **$7.0\times10^{-4}$** | $8.5\times10^{1}$ | $3.7\times10^{-1}$ |
| H71G | 1.1 | $7.7\times10^{-1}$ | $3.5\times10^{5}$ | $1.9\times10^{-1}$ | $7.6\times10^{-1}$ | $4.2\times10^{-1}$ | $7.1\times10^{-1}$ | N/A | **$2.0\times10^{-4}$** | $1.0\times10^{2}$ | $4.7\times10^{-1}$ |
| H71V | 1.1 | $7.7\times10^{-1}$ | $8.1\times10^{5}$ | $3.2\times10^{-1}$ | $4.4\times10^{-1}$ | **$7.9\times10^{-3}$** | $9.7\times10^{-1}$ | N/A | $4.9\times10^{-1}$ | $1.2\times10^{2}$ | $6.5\times10^{-1}$ |
| I72G | $2.5\times10^{-2}$ | **0** | $2.4\times10^{5}$ | $7.1\times10^{-2}$ | – | – | $8.1\times10^{-1}$ | N/A | **$1.\times10^{-4}$** | $2.2\times10^{2}$ | $1.0\times10^{-1}$ |
| I72V | $7.8\times10^{-1}$ | $2.9\times10^{-1}$ | $3.2\times10^{5}$ | $1.2\times10^{-1}$ | 1.7 | $1.3\times10^{-1}$ | $5.9\times10^{-1}$ | N/A | **0** | $1.3\times10^{2}$ | $8.2\times10^{-1}$ |
| D73G | $2.6\times10^{-1}$ | **0** | $5.5\times10^{5}$ | $8.0\times10^{-1}$ | 1.1 | $7.4\times10^{-1}$ | $6.7\times10^{-1}$ | N/A | **$1.\times10^{-4}$** | $9.0\times10^{1}$ | $3.4\times10^{-1}$ |
| D73V | $1.8\times10^{-1}$ | **0** | $3.5\times10^{5}$ | $1.4\times10^{-1}$ | 1.3 | $2.6\times10^{-1}$ | $5.6\times10^{-1}$ | N/A | **0** | $1.4\times10^{2}$ | $9.1\times10^{-1}$ |
| Y74G | ≥ $1.2\times10^{-1}$ | **0** | ≥ $7.1\times10^{5}$ | $5.5\times10^{-1}$ | 1.8 | $1.5\times10^{-2}$ | $6.7\times10^{-2}$ | N/A | **0** | ≤ $1.2\times10^{2}$ | $6.9\times10^{-1}$ |
| Y74V | $7.2\times10^{-1}$ | $1.1\times10^{-1}$ | $7.0\times10^{5}$ | $6.2\times10^{-1}$ | 1.6 | $1.4\times10^{-1}$ | $1.7\times10^{-1}$ | N/A | **0** | $1.6\times10^{2}$ | $6.1\times10^{-1}$ |
| V75A | ≤ $6.7\times10^{-1}$ | $7.6\times10^{-2}$ | ≤ $8.3\times10^{4}$ | **$2.3\times10^{-2}$** | ≤ 2.9 | $1.3\times10^{-2}$ | $9.0\times10^{-1}$ | N/A | $7.8\times10^{-2}$ | ≥ $1.1\times10^{2}$ | $5.7\times10^{-1}$ |
| V75G | $2.0\times10^{-1}$ | **$1.0\times10^{-3}$** | $1.1\times10^{6}$ | $1.5\times10^{-1}$ | – | – | $3.4\times10^{-1}$ | N/A | **$1.\times10^{-4}$** | $6.0\times10^{1}$ | $2.5\times10^{-1}$ |
| P76V | $5.1\times10^{-2}$ | **0** | $1.1\times10^{5}$ | $7.1\times10^{-2}$ | – | – | $4.1\times10^{-1}$ | N/A | **$1.\times10^{-4}$** | $6.6\times10^{1}$ | $1.5\times10^{-1}$ |
| T77G | $1.8\times10^{-1}$ | **0** | $1.5\times10^{6}$ | **$1.1\times10^{-2}$** | 1.4 | $2.4\times10^{-1}$ | $1.5\times10^{-1}$ | N/A | **$3.0\times10^{-4}$** | $1.4\times10^{2}$ | $9.8\times10^{-1}$ |
| T77V | $6.4\times10^{-1}$ | $7.3\times10^{-2}$ | $3.9\times10^{5}$ | $2.7\times10^{-1}$ | 3.2 | **$1.9\times10^{-3}$** | $4.1\times10^{-1}$ | N/A | **$1.9\times10^{-3}$** | $1.1\times10^{2}$ | $5.7\times10^{-1}$ |
| V78A | 2.0 | **$4.0\times10^{-3}$** | $2.6\times10^{4}$ | **$9.0\times10^{-3}$** | – | – | 1.0 | N/A | $9.8\times10^{-1}$ | $3.3\times10^{1}$ | **$3.1\times10^{-2}$** |
| V78G | $4.6\times10^{-1}$ | **$5.0\times10^{-3}$** | $2.1\times10^{5}$ | $9.3\times10^{-2}$ | – | – | $4.7\times10^{-1}$ | N/A | **0** | $4.0\times10^{1}$ | $7.3\times10^{-2}$ |
| T79G | – | – | – | – | – | – | – | – | – | – | – |
| T79V | – | – | – | – | – | – | – | – | – | – | – |
| A80G | $3.2\times10^{-1}$ | **$1.0\times10^{-3}$** | $7.2\times10^{5}$ | $6.0\times10^{-1}$ | 1.3 | $3.8\times10^{-1}$ | $7.2\times10^{-1}$ | N/A | **$1.\times10^{-4}$** | $1.3\times10^{2}$ | $8.5\times10^{-1}$ |
| A80V | $1.4\times10^{-1}$ | **0** | $1.0\times10^{5}$ | **$1.8\times10^{-2}$** | 2.2 | $2.1\times10^{-2}$ | N/A | 1.7 | **0** | $3.7\times10^{1}$ | **$3.2\times10^{-2}$** |
| I81G | $6.6\times10^{-1}$ | $1.1\times10^{-1}$ | $1.2\times10^{6}$ | **$2.9\times10^{-2}$** | $8.2\times10^{-1}$ | $5.0\times10^{-1}$ | $1.1\times10^{-1}$ | N/A | **$4.0\times10^{-4}$** | $5.5\times10^{1}$ | $1.1\times10^{-1}$ |
| I81V | $9.5\times10^{-1}$ | $7.7\times10^{-1}$ | $5.3\times10^{5}$ | $6.3\times10^{-1}$ | 1.3 | $3.6\times10^{-1}$ | $6.7\times10^{-1}$ | N/A | **0** | $1.4\times10^{2}$ | $9.9\times10^{-1}$ |
| G82A | ≥ $6.9\times10^{-2}$ | **0** | ≥ $1.3\times10^{5}$ | **$1.4\times10^{-2}$** | $1.1\times10^{1}$ | **$1.5\times10^{-4}$** | $3.4\times10^{-2}$ | N/A | **0** | ≤ $6.4\times10^{1}$ | $1.0\times10^{-1}$ |
| G82V | – | – | – | – | – | – | – | – | – | – | – |
| H83G | ≥ $2.2\times10^{-1}$ | **0** | ≥ $4.4\times10^{5}$ | $2.8\times10^{-1}$ | $2.3\times10^{-1}$ | **$2.7\times10^{-3}$** | $1.4\times10^{-2}$ | N/A | **0** | ≤ $2.7\times10^{1}$ | **$2.9\times10^{-2}$** |
| H83V | 1.3 | $2.7\times10^{-1}$ | $2.0\times10^{5}$ | $8.3\times10^{-2}$ | $6.2\times10^{-1}$ | $1.7\times10^{-1}$ | $1.5\times10^{-1}$ | N/A | **0** | $3.0\times10^{1}$ | **$4.1\times10^{-2}$** |
| T84G | 1.4 | $2.8\times10^{-1}$ | $2.4\times10^{5}$ | $1.5\times10^{-1}$ | 2.0 | $5.1\times10^{-2}$ | N/A | 1.3 | **$6.0\times10^{-4}$** | $9.8\times10^{1}$ | $4.2\times10^{-1}$ |
| S85G | $7.9\times10^{-1}$ | $4.2\times10^{-1}$ | $9.5\times10^{5}$ | $1.2\times10^{-1}$ | $5.2\times10^{-1}$ | $1.4\times10^{-2}$ | $7.8\times10^{-1}$ | N/A | **$1.\times10^{-4}$** | $1.2\times10^{2}$ | $6.3\times10^{-1}$ |
| S85V | $1.9\times10^{-1}$ | **0** | $6.2\times10^{4}$ | **$2.4\times10^{-2}$** | 7.8 | **$3.1\times10^{-4}$** | $7.1\times10^{-1}$ | N/A | **0** | $1.1\times10^{2}$ | $5.3\times10^{-1}$ |
| I86G | $1.3\times10^{-1}$ | **0** | $1.4\times10^{6}$ | **$3.2\times10^{-2}$** | 1.1 | $7.3\times10^{-1}$ | $6.1\times10^{-1}$ | N/A | **0** | $6.8\times10^{1}$ | $2.0\times10^{-1}$ |
| I86V | 1.2 | $4.8\times10^{-1}$ | $3.1\times10^{5}$ | $1.4\times10^{-1}$ | $7.8\times10^{-1}$ | $5.0\times10^{-1}$ | $9.4\times10^{-1}$ | N/A | $1.7\times10^{-1}$ | $1.3\times10^{2}$ | $7.4\times10^{-1}$ |
| F87G | 1.4 | $2.4\times10^{-1}$ | $4.7\times10^{5}$ | $5.6\times10^{-1}$ | $8.6\times10^{-1}$ | $5.2\times10^{-1}$ | $5.6\times10^{-1}$ | N/A | **$1.\times10^{-4}$** | $8.2\times10^{1}$ | $2.2\times10^{-1}$ |
| F87V | 1.2 | $5.3\times10^{-1}$ | $3.1\times10^{5}$ | $1.6\times10^{-1}$ | 1.7 | $1.3\times10^{-1}$ | $4.7\times10^{-1}$ | N/A | **0** | $1.0\times10^{2}$ | $3.6\times10^{-1}$ |
| T88G | $1.8\times10^{-1}$ | **$2.0\times10^{-3}$** | $1.2\times10^{6}$ | $8.8\times10^{-2}$ | 1.0 | $9.0\times10^{-1}$ | $2.8\times10^{-1}$ | N/A | **$4.0\times10^{-4}$** | $6.9\times10^{1}$ | $3.5\times10^{-1}$ |
| T88V | $5.4\times10^{-1}$ | $2.2\times10^{-2}$ | $1.3\times10^{5}$ | **$4.4\times10^{-2}$** | 1.7 | $1.5\times10^{-1}$ | $5.9\times10^{-1}$ | N/A | **0** | $1.1\times10^{2}$ | $5.4\times10^{-1}$ |
| G89A | $5.4\times10^{-1}$ | **$9.0\times10^{-3}$** | $6.5\times10^{5}$ | $8.5\times10^{-1}$ | 1.5 | $2.3\times10^{-1}$ | N/A | 1.1 | $1.4\times10^{-1}$ | $1.1\times10^{2}$ | $5.1\times10^{-1}$ |
| G89V | – | – | – | – | – | – | – | – | – | – | – |
| S90G | $7.8\times10^{-1}$ | $2.3\times10^{-1}$ | $6.5\times10^{5}$ | $7.8\times10^{-1}$ | $6.2\times10^{-1}$ | $1.2\times10^{-1}$ | $6.3\times10^{-1}$ | N/A | **0** | $7.6\times10^{1}$ | $1.6\times10^{-1}$ |
| S90V | $1.9\times10^{-1}$ | **0** | $4.5\times10^{5}$ | $5.9\times10^{-1}$ | – | – | $7.7\times10^{-1}$ | N/A | $3.8\times10^{-2}$ | $1.3\times10^{2}$ | $8.3\times10^{-1}$ |
| V91A | ≤ $7.4\times10^{-1}$ | $1.6\times10^{-1}$ | ≤ $1.1\times10^{5}$ | **$3.4\times10^{-2}$** | ≤ 2.9 | $1.5\times10^{-2}$ | $7.6\times10^{-1}$ | N/A | **$3.0\times10^{-4}$** | ≥ $9.9\times10^{1}$ | $3.1\times10^{-1}$ |
| V91G | $6.7\times10^{-2}$ | **0** | $2.8\times10^{5}$ | $8.2\times10^{-2}$ | – | – | $5.6\times10^{-1}$ | N/A | **0** | $8.0\times10^{1}$ | $1.7\times10^{-1}$ |
| P92G | $1.6\times10^{-1}$ | **0** | $5.8\times10^{5}$ | $8.7\times10^{-1}$ | 1.2 | $6.1\times10^{-1}$ | $5.9\times10^{-1}$ | N/A | **0** | $8.3\times10^{1}$ | $1.8\times10^{-1}$ |
| P92V | ≤ $8.6\times10^{-2}$ | **0** | ≤ $1.6\times10^{5}$ | **$4.8\times10^{-2}$** | – | – | $8.4\times10^{-1}$ | N/A | $2.3\times10^{-2}$ | ≥ $9.3\times10^{1}$ | $3.7\times10^{-1}$ |
| S93G | $3.8\times10^{-1}$ | **$5.0\times10^{-3}$** | $6.5\times10^{5}$ | $8.3\times10^{-1}$ | 1.2 | $5.7\times10^{-1}$ | $6.5\times10^{-1}$ | N/A | **0** | $9.2\times10^{1}$ | $3.5\times10^{-1}$ |
| S93V | $3.9\times10^{-2}$ | **0** | $1.0\times10^{5}$ | $5.8\times10^{-2}$ | – | – | $5.6\times10^{-1}$ | N/A | **$2.0\times10^{-4}$** | $7.3\times10^{1}$ | $2.3\times10^{-1}$ |
| I94G | $2.0\times10^{-2}$ | **0** | $2.8\times10^{5}$ | $1.0\times10^{-1}$ | – | – | $2.6\times10^{-1}$ | N/A | **0** | $9.0\times10^{1}$ | $3.2\times10^{-1}$ |
| I94V | 1.1 | $7.3\times10^{-1}$ | $3.3\times10^{5}$ | $1.4\times10^{-1}$ | 1.7 | $7.5\times10^{-2}$ | $9.6\times10^{-1}$ | N/A | $4.1\times10^{-1}$ | $1.5\times10^{2}$ | $8.8\times10^{-1}$ |
| H95G | $3.3\times10^{-2}$ | **0** | $3.9\times10^{6}$ | **$3.0\times10^{-3}$** | – | – | – | – | – | – | – |
| H95V | $3.4\times10^{-1}$ | **$1.0\times10^{-3}$** | $2.8\times10^{5}$ | $1.4\times10^{-1}$ | 4.5 | **$2.1\times10^{-3}$** | $6.3\times10^{-1}$ | N/A | **$6.0\times10^{-4}$** | $1.4\times10^{2}$ | $9.8\times10^{-1}$ |
| G96A | $4.8\times10^{-1}$ | $2.8\times10^{-2}$ | $2.2\times10^{5}$ | $9.5\times10^{-2}$ | 1.7 | $1.3\times10^{-1}$ | $7.9\times10^{-1}$ | N/A | **$1.1\times10^{-3}$** | $6.7\times10^{1}$ | $2.1\times10^{-1}$ |
| G96V | – | – | – | – | – | – | – | – | – | – | – |

| Variant | $f_a$ | $f_a$ $p$–value | $(k_{cat}/K_M)^{MeP}_{chem.}$ ($M^{-1}s^{-1}$) | $(k_{cat}/K_M)^{MeP}_{chem.}$ $p$–value | FC1 | FC1 $p$–value | FC2 | FC3 | FC2/3 $p$–value | FC4 ($s^{-1}$) | FC4 $p$–value |
|---|---|---|---|---|---|---|---|---|---|---|---|
| I97G | – | – | – | – | – | – | – | – | – | – | – |
| I97V | 1.2 | $4.2\times10^{-1}$ | $3.6\times10^{5}$ | $1.2\times10^{-1}$ | 1.3 | $4.2\times10^{-1}$ | $9.3\times10^{-1}$ | N/A | $2.3\times10^{-1}$ | $1.5\times10^{2}$ | $8.1\times10^{-1}$ |
| A98G | $5.7\times10^{-1}$ | $8.2\times10^{-2}$ | $4.3\times10^{5}$ | $4.8\times10^{-1}$ | 1.5 | $1.2\times10^{-1}$ | $8.6\times10^{-1}$ | N/A | $1.2\times10^{-2}$ | $1.0\times10^{2}$ | $5.4\times10^{-1}$ |
| A98V | $8.8\times10^{-1}$ | $6.7\times10^{-1}$ | $5.0\times10^{5}$ | $5.5\times10^{-1}$ | 1.1 | $7.8\times10^{-1}$ | $8.1\times10^{-1}$ | N/A | **$7.0\times10^{-4}$** | $1.6\times10^{2}$ | $6.5\times10^{-1}$ |
| G99A | $7.2\times10^{-1}$ | $2.4\times10^{-1}$ | $3.8\times10^{5}$ | $3.4\times10^{-1}$ | 1.5 | $1.8\times10^{-1}$ | N/A | 1.4 | **$2.0\times10^{-4}$** | $1.7\times10^{2}$ | $5.5\times10^{-1}$ |
| G99V | $\leq 1.1\times10^{-1}$ | **0** | $\leq 1.9\times10^{4}$ | **$9.0\times10^{-3}$** | $\leq 3.2\times10^{1}$ | **$2.0\times10^{-4}$** | $1.6\times10^{-1}$ | N/A | **0** | $\geq 1.3\times10^{1}$ | **$3.6\times10^{-2}$** |
| N100G | $2.3\times10^{-1}$ | **0** | $7.6\times10^{3}$ | **$1.0\times10^{-2}$** | $2.6\times10^{1}$ | **$1.2\times10^{-3}$** | $\leq 5.9\times10^{-1}$ | N/A | **0** | $\geq 5.8$ | **$1.8\times10^{-2}$** |
| N100V | $\leq 3.4\times10^{-1}$ | **$1.0\times10^{-3}$** | $\leq 4.8\times10^{3}$ | **$1.6\times10^{-2}$** | – | – | N/A | $\leq 1.3$ | **0** | $\geq 5.2$ | **$2.1\times10^{-2}$** |
| D101G | $9.4\times10^{-1}$ | $8.0\times10^{-1}$ | $1.2\times10^{5}$ | $7.8\times10^{-2}$ | 8.7 | **$1.0\times10^{-5}$** | N/A | 1.2 | **$6.8\times10^{-3}$** | $4.5\times10^{1}$ | $1.2\times10^{-1}$ |
| D101V | 1.6 | $1.0\times10^{-2}$ | $1.0\times10^{5}$ | **$3.0\times10^{-3}$** | 3.6 | **$1.4\times10^{-4}$** | N/A | 1.6 | **0** | $9.1\times10^{1}$ | $1.3\times10^{-1}$ |
| W102G | – | – | – | – | – | – | – | – | – | – | – |
| W102V | $4.1\times10^{-2}$ | **0** | $9.1\times10^{4}$ | **$1.9\times10^{-2}$** | – | – | N/A | 1.2 | **$4.0\times10^{-4}$** | $4.5\times10^{1}$ | $5.8\times10^{-2}$ |
| Y103G | $2.0\times10^{-1}$ | **0** | $1.3\times10^{5}$ | **$3.4\times10^{-2}$** | 2.7 | $1.1\times10^{-2}$ | $6.2\times10^{-1}$ | N/A | **$1.\times10^{-4}$** | $5.3\times10^{1}$ | $7.2\times10^{-2}$ |
| Y103V | $7.6\times10^{-1}$ | $2.6\times10^{-1}$ | $1.7\times10^{5}$ | $5.3\times10^{-2}$ | 1.6 | $1.9\times10^{-1}$ | $7.6\times10^{-1}$ | N/A | **0** | $1.5\times10^{2}$ | $9.3\times10^{-1}$ |
| D104G | $4.8\times10^{-1}$ | $1.5\times10^{-2}$ | $3.7\times10^{5}$ | $2.7\times10^{-1}$ | 2.2 | $2.3\times10^{-2}$ | $8.0\times10^{-1}$ | N/A | **$1.\times10^{-4}$** | $1.1\times10^{2}$ | $5.1\times10^{-1}$ |
| D104V | 1.1 | $7.9\times10^{-1}$ | $1.0\times10^{5}$ | **$3.1\times10^{-2}$** | – | – | N/A | 1.4 | **0** | $1.4\times10^{2}$ | $9.2\times10^{-1}$ |
| K105G | 1.5 | $1.1\times10^{-1}$ | $3.6\times10^{5}$ | $1.5\times10^{-1}$ | 1.8 | $4.5\times10^{-2}$ | N/A | 1.1 | $3.4\times10^{-1}$ | $1.4\times10^{2}$ | $9.7\times10^{-1}$ |
| K105V | 1.9 | $1.4\times10^{-2}$ | $3.6\times10^{5}$ | $1.8\times10^{-1}$ | $9.7\times10^{-1}$ | $9.5\times10^{-1}$ | N/A | 1.0 | $7.9\times10^{-1}$ | $1.3\times10^{2}$ | $9.1\times10^{-1}$ |
| E106G | $9.5\times10^{-1}$ | $8.2\times10^{-1}$ | $2.6\times10^{5}$ | $8.9\times10^{-2}$ | 2.8 | **$1.5\times10^{-3}$** | $8.7\times10^{-1}$ | N/A | $4.2\times10^{-2}$ | $1.4\times10^{2}$ | $9.6\times10^{-1}$ |
| E106V | $4.2\times10^{-1}$ | **$2.0\times10^{-3}$** | $1.1\times10^{6}$ | **$3.3\times10^{-2}$** | $8.5\times10^{-1}$ | $6.8\times10^{-1}$ | $8.7\times10^{-1}$ | N/A | **$7.1\times10^{-3}$** | $1.4\times10^{2}$ | $9.9\times10^{-1}$ |
| L107G | 1.5 | $1.4\times10^{-1}$ | $2.9\times10^{5}$ | $1.2\times10^{-1}$ | 2.3 | **$4.9\times10^{-3}$** | $7.3\times10^{-1}$ | N/A | **$2.0\times10^{-4}$** | $8.2\times10^{1}$ | $2.6\times10^{-1}$ |
| L107V | 1.0 | $8.6\times10^{-1}$ | $6.1\times10^{5}$ | $9.9\times10^{-1}$ | $8.1\times10^{-1}$ | $4.5\times10^{-1}$ | N/A | 1.1 | $1.5\times10^{-1}$ | $1.7\times10^{2}$ | $5.5\times10^{-1}$ |
| G108A | $9.4\times10^{-1}$ | $7.5\times10^{-1}$ | $2.8\times10^{5}$ | $1.7\times10^{-1}$ | 1.1 | $8.3\times10^{-1}$ | $8.7\times10^{-1}$ | N/A | $1.5\times10^{-2}$ | $8.5\times10^{1}$ | $2.9\times10^{-1}$ |
| G108V | $9.9\times10^{-1}$ | $9.0\times10^{-1}$ | $3.9\times10^{5}$ | $1.6\times10^{-1}$ | $8.8\times10^{-1}$ | $7.1\times10^{-1}$ | N/A | 1.1 | $2.2\times10^{-1}$ | $1.6\times10^{2}$ | $6.0\times10^{-1}$ |
| K109G | 1.8 | $2.5\times10^{-2}$ | $4.0\times10^{5}$ | $2.8\times10^{-1}$ | 1.1 | $5.4\times10^{-1}$ | N/A | 1.3 | **$7.0\times10^{-4}$** | $1.5\times10^{2}$ | $8.9\times10^{-1}$ |
| K109V | 1.4 | $1.4\times10^{-1}$ | $4.2\times10^{5}$ | $2.5\times10^{-1}$ | 1.0 | $9.2\times10^{-1}$ | $9.7\times10^{-1}$ | N/A | $4.4\times10^{-1}$ | $1.3\times10^{2}$ | $7.3\times10^{-1}$ |
| S110G | 1.2 | $4.2\times10^{-1}$ | $5.4\times10^{5}$ | $6.8\times10^{-1}$ | 1.2 | $5.5\times10^{-1}$ | $7.4\times10^{-1}$ | N/A | **$1.\times10^{-4}$** | $1.4\times10^{2}$ | $9.4\times10^{-1}$ |
| S110V | 1.2 | $5.3\times10^{-1}$ | $3.5\times10^{5}$ | $1.9\times10^{-1}$ | 1.1 | $6.6\times10^{-1}$ | N/A | 1.9 | **0** | $8.4\times10^{1}$ | $1.5\times10^{-1}$ |
| V111A | 1.2 | $4.7\times10^{-1}$ | $1.5\times10^{5}$ | **$1.8\times10^{-2}$** | 2.8 | **$1.1\times10^{-3}$** | N/A | 1.2 | **$2.1\times10^{-3}$** | $1.4\times10^{2}$ | $9.0\times10^{-1}$ |
| V111G | $7.4\times10^{-2}$ | **0** | $1.4\times10^{6}$ | **$2.8\times10^{-2}$** | 2.0 | $4.7\times10^{-2}$ | N/A | 3.6 | **0** | $1.3\times10^{2}$ | $8.7\times10^{-1}$ |
| Y112G | $4.8\times10^{-1}$ | **0** | $5.4\times10^{4}$ | **$7.0\times10^{-3}$** | $4.7\times10^{1}$ | **0** | N/A | 3.7 | **0** | $3.9\times10^{1}$ | **$2.7\times10^{-2}$** |
| Y112V | $3.2\times10^{-1}$ | **0** | $4.6\times10^{4}$ | **$1.8\times10^{-2}$** | $2.3\times10^{1}$ | **$1.4\times10^{-4}$** | N/A | 7.4 | **0** | $1.1\times10^{2}$ | $5.3\times10^{-1}$ |
| C113G | $9.1\times10^{-1}$ | $7.7\times10^{-1}$ | $2.3\times10^{5}$ | $1.2\times10^{-1}$ | 2.5 | **$3.9\times10^{-3}$** | N/A | 1.2 | **$8.4\times10^{-3}$** | $1.1\times10^{2}$ | $5.8\times10^{-1}$ |
| C113V | 1.8 | $1.0\times10^{-2}$ | $3.1\times10^{5}$ | $9.2\times10^{-2}$ | 1.4 | $2.8\times10^{-1}$ | $7.6\times10^{-1}$ | N/A | **$4.8\times10^{-3}$** | $2.1\times10^{2}$ | $1.0\times10^{-1}$ |
| T114G | $\leq 2.3\times10^{-1}$ | **0** | $\leq 6.4\times10^{3}$ | **$8.0\times10^{-3}$** | $\leq 9.1\times10^{1}$ | **0** | $\leq 9.6\times10^{-1}$ | N/A | $2.9\times10^{-1}$ | $\geq 5.8$ | **$2.1\times10^{-2}$** |
| T114V | $9.7\times10^{-1}$ | $8.4\times10^{-1}$ | $5.5\times10^{5}$ | $7.4\times10^{-1}$ | 3.0 | **$3.5\times10^{-4}$** | $5.9\times10^{-1}$ | N/A | **$6.0\times10^{-4}$** | $1.3\times10^{2}$ | $7.4\times10^{-1}$ |
| S115G | 1.1 | $7.3\times10^{-1}$ | $4.1\times10^{5}$ | $3.1\times10^{-1}$ | 1.7 | $1.2\times10^{-1}$ | $9.4\times10^{-1}$ | N/A | $1.9\times10^{-1}$ | $9.0\times10^{1}$ | $2.7\times10^{-1}$ |
| S115V | $7.4\times10^{-1}$ | $2.4\times10^{-1}$ | $5.8\times10^{5}$ | $8.6\times10^{-1}$ | 1.1 | $7.6\times10^{-1}$ | N/A | 3.0 | **0** | $1.6\times10^{2}$ | $6.4\times10^{-1}$ |
| D116G | $\leq 1.3\times10^{-2}$ | **0** | $\leq 9.2\times10^{4}$ | $5.5\times10^{-2}$ | – | – | $\leq 8.6\times10^{-1}$ | N/A | $2.1\times10^{-1}$ | $\geq 7.0$ | $8.5\times10^{-2}$ |
| D116V | $\leq 1.5\times10^{-1}$ | **0** | $\leq 6.1\times10^{4}$ | **$4.1\times10^{-2}$** | – | – | – | – | – | – | – |
| E117G | 1.3 | $2.7\times10^{-1}$ | $2.9\times10^{5}$ | $9.6\times10^{-2}$ | 1.8 | $3.9\times10^{-2}$ | $9.2\times10^{-1}$ | N/A | $6.8\times10^{-2}$ | $5.7\times10^{1}$ | $9.2\times10^{-2}$ |
| E117V | 1.2 | $4.5\times10^{-1}$ | $4.4\times10^{5}$ | $3.3\times10^{-1}$ | $9.0\times10^{-1}$ | $8.2\times10^{-1}$ | N/A | 1.1 | $2.4\times10^{-1}$ | $1.5\times10^{2}$ | $7.7\times10^{-1}$ |
| T118G | 1.4 | $1.8\times10^{-1}$ | $3.5\times10^{5}$ | $1.3\times10^{-1}$ | 1.0 | $8.5\times10^{-1}$ | $8.8\times10^{-1}$ | N/A | $1.2\times10^{-1}$ | $8.3\times10^{1}$ | $2.2\times10^{-1}$ |
| T118V | 1.1 | $5.9\times10^{-1}$ | $4.4\times10^{5}$ | $3.3\times10^{-1}$ | 1.1 | $8.4\times10^{-1}$ | $9.8\times10^{-1}$ | N/A | $6.6\times10^{-1}$ | $1.6\times10^{2}$ | $6.5\times10^{-1}$ |
| V119A | 1.3 | $3.1\times10^{-1}$ | $4.7\times10^{5}$ | $4.6\times10^{-1}$ | 1.4 | $3.3\times10^{-1}$ | N/A | 1.2 | $1.4\times10^{-2}$ | $2.1\times10^{2}$ | $1.3\times10^{-1}$ |
| V119G | $7.1\times10^{-2}$ | **0** | $1.2\times10^{6}$ | $5.7\times10^{-2}$ | 3.0 | **$4.0\times10^{-3}$** | N/A | 1.2 | $1.5\times10^{-2}$ | $1.2\times10^{2}$ | $7.6\times10^{-1}$ |
| Q120G | 1.4 | $1.4\times10^{-1}$ | $3.7\times10^{5}$ | $2.3\times10^{-1}$ | 1.5 | $1.5\times10^{-1}$ | $9.2\times10^{-1}$ | N/A | $1.1\times10^{-1}$ | $8.8\times10^{1}$ | $2.8\times10^{-1}$ |
| Q120V | 1.1 | $7.0\times10^{-1}$ | $4.9\times10^{5}$ | $3.3\times10^{-1}$ | $8.5\times10^{-1}$ | $5.5\times10^{-1}$ | N/A | 1.1 | $1.4\times10^{-2}$ | $1.8\times10^{2}$ | $2.9\times10^{-1}$ |
| P121G | 1.2 | $5.5\times10^{-1}$ | $9.5\times10^{5}$ | $1.4\times10^{-1}$ | $4.5\times10^{-1}$ | $2.0\times10^{-2}$ | $9.0\times10^{-1}$ | N/A | $2.0\times10^{-1}$ | $8.2\times10^{1}$ | $3.4\times10^{-1}$ |
| P121V | $7.3\times10^{-1}$ | $1.1\times10^{-1}$ | $7.4\times10^{5}$ | $3.7\times10^{-1}$ | 1.0 | $8.9\times10^{-1}$ | N/A | 1.0 | $5.9\times10^{-1}$ | $1.4\times10^{2}$ | $9.8\times10^{-1}$ |
| V122A | 1.1 | $6.7\times10^{-1}$ | $2.0\times10^{5}$ | $6.7\times10^{-2}$ | 1.9 | $5.7\times10^{-2}$ | $9.8\times10^{-1}$ | N/A | $6.2\times10^{-1}$ | $1.6\times10^{2}$ | $7.5\times10^{-1}$ |
| V122G | $2.8\times10^{-1}$ | **0** | $1.1\times10^{5}$ | **$1.7\times10^{-2}$** | 3.2 | **$1.6\times10^{-3}$** | $3.9\times10^{-1}$ | N/A | **0** | 8.8 | **$1.9\times10^{-2}$** |
| G123A | $9.3\times10^{-1}$ | $7.9\times10^{-1}$ | $3.1\times10^{5}$ | $1.6\times10^{-1}$ | 2.3 | **$5.2\times10^{-3}$** | $9.7\times10^{-1}$ | N/A | $4.7\times10^{-1}$ | $1.1\times10^{2}$ | $6.2\times10^{-1}$ |
| G123V | 1.4 | $2.3\times10^{-1}$ | $1.2\times10^{5}$ | **$3.2\times10^{-2}$** | 2.3 | $2.9\times10^{-2}$ | $9.3\times10^{-1}$ | N/A | $1.4\times10^{-1}$ | $7.7\times10^{1}$ | $2.1\times10^{-1}$ |
| T124G | 1.2 | $4.3\times10^{-1}$ | $2.5\times10^{5}$ | $1.3\times10^{-1}$ | 2.6 | **$2.3\times10^{-3}$** | N/A | 1.0 | $6.7\times10^{-1}$ | $7.6\times10^{1}$ | $2.0\times10^{-1}$ |
| T124V | 1.4 | $1.8\times10^{-1}$ | $2.6\times10^{5}$ | $7.8\times10^{-2}$ | 1.3 | $3.8\times10^{-1}$ | $8.5\times10^{-1}$ | N/A | $1.5\times10^{-2}$ | $1.1\times10^{2}$ | $4.9\times10^{-1}$ |
| T125G | 1.6 | $1.2\times10^{-1}$ | $2.7\times10^{5}$ | $1.2\times10^{-1}$ | 2.0 | $2.1\times10^{-2}$ | $9.0\times10^{-1}$ | N/A | $8.4\times10^{-2}$ | $8.2\times10^{1}$ | $3.2\times10^{-1}$ |
| T125V | $8.9\times10^{-1}$ | $6.7\times10^{-1}$ | $7.2\times10^{5}$ | $5.3\times10^{-1}$ | $6.0\times10^{-1}$ | $1.4\times10^{-1}$ | N/A | 1.1 | $2.4\times10^{-1}$ | $1.7\times10^{2}$ | $5.1\times10^{-1}$ |
| S126G | $5.8\times10^{-1}$ | $4.1\times10^{-2}$ | $6.6\times10^{5}$ | $7.8\times10^{-1}$ | 2.3 | **$4.6\times10^{-3}$** | $9.6\times10^{-1}$ | N/A | $4.8\times10^{-1}$ | $1.5\times10^{2}$ | $7.9\times10^{-1}$ |
| S126V | $9.9\times10^{-1}$ | $9.5\times10^{-1}$ | $7.9\times10^{5}$ | $3.9\times10^{-1}$ | $7.9\times10^{-1}$ | $5.4\times10^{-1}$ | N/A | 1.1 | $1.3\times10^{-1}$ | $1.9\times10^{2}$ | $2.7\times10^{-1}$ |
| N127G | 1.1 | $7.6\times10^{-1}$ | $4.8\times10^{5}$ | $5.2\times10^{-1}$ | 1.5 | $1.1\times10^{-1}$ | N/A | 1.3 | **$7.0\times10^{-4}$** | $1.0\times10^{2}$ | $4.5\times10^{-1}$ |
| N127V | 1.1 | $8.5\times10^{-1}$ | $6.4\times10^{5}$ | $9.1\times10^{-1}$ | $7.5\times10^{-1}$ | $4.2\times10^{-1}$ | N/A | 1.0 | $5.9\times10^{-1}$ | $1.7\times10^{2}$ | $5.2\times10^{-1}$ |
| S128G | 1.4 | $2.4\times10^{-1}$ | $2.9\times10^{5}$ | $1.3\times10^{-1}$ | 2.0 | $1.5\times10^{-2}$ | N/A | 1.0 | $6.7\times10^{-1}$ | $1.2\times10^{2}$ | $7.0\times10^{-1}$ |
| S128V | $8.6\times10^{-1}$ | $5.6\times10^{-1}$ | $4.4\times10^{5}$ | $4.1\times10^{-1}$ | 1.1 | $8.3\times10^{-1}$ | N/A | 1.1 | $3.0\times10^{-2}$ | $1.1\times10^{2}$ | $4.8\times10^{-1}$ |
| V129A | 1.2 | $4.9\times10^{-1}$ | $3.0\times10^{5}$ | $1.0\times10^{-1}$ | 2.7 | **$1.9\times10^{-3}$** | N/A | 1.1 | $3.7\times10^{-2}$ | $1.9\times10^{2}$ | $2.4\times10^{-1}$ |
| V129G | $4.8\times10^{-1}$ | $1.8\times10^{-2}$ | $1.0\times10^{6}$ | $9.4\times10^{-2}$ | 1.0 | $8.7\times10^{-1}$ | $9.6\times10^{-1}$ | N/A | $4.6\times10^{-1}$ | $1.9\times10^{2}$ | $3.5\times10^{-1}$ |
| G130A | 1.4 | $2.0\times10^{-1}$ | $2.2\times10^{5}$ | $1.1\times10^{-1}$ | 2.6 | $1.1\times10^{-2}$ | N/A | 1.0 | $5.7\times10^{-1}$ | $1.1\times10^{2}$ | $6.1\times10^{-1}$ |
| G130V | $5.5\times10^{-1}$ | **$4.0\times10^{-3}$** | $6.5\times10^{4}$ | **$1.0\times10^{-2}$** | 6.0 | **$6.4\times10^{-4}$** | N/A | 1.1 | $1.3\times10^{-2}$ | $1.4\times10^{1}$ | **$1.6\times10^{-2}$** |
| Q131G | 1.0 | $8.3\times10^{-1}$ | $4.9\times10^{5}$ | $5.6\times10^{-1}$ | 2.4 | **$3.4\times10^{-3}$** | $9.9\times10^{-1}$ | N/A | $7.0\times10^{-1}$ | $7.4\times10^{1}$ | $1.5\times10^{-1}$ |
| H132G | $7.3\times10^{-1}$ | $2.0\times10^{-1}$ | $4.5\times10^{5}$ | $3.5\times10^{-1}$ | 4.5 | **$1.0\times10^{-5}$** | N/A | 1.1 | $1.2\times10^{-1}$ | $6.6\times10^{1}$ | $1.4\times10^{-1}$ |
| H132V | $\leq 4.4\times10^{-1}$ | **$6.0\times10^{-3}$** | $\leq 2.8\times10^{5}$ | $1.5\times10^{-1}$ | $\leq 1.1\times10^{1}$ | **$3.0\times10^{-5}$** | N/A | 1.4 | **$1.9\times10^{-3}$** | $\geq 1.6\times10^{2}$ | $7.8\times10^{-1}$ |
| S133G | 1.1 | $8.3\times10^{-1}$ | $2.7\times10^{5}$ | $1.1\times10^{-1}$ | 1.7 | $7.6\times10^{-2}$ | $7.1\times10^{-1}$ | N/A | **$1.4\times10^{-3}$** | $7.6\times10^{1}$ | $2.7\times10^{-1}$ |
| S133V | – | – | – | – | – | – | – | – | – | – | – |
| P134G | $1.3\times10^{-1}$ | **0** | $5.2\times10^{5}$ | $5.9\times10^{-1}$ | 1.4 | $2.4\times10^{-1}$ | $9.4\times10^{-1}$ | N/A | $1.3\times10^{-1}$ | $1.1\times10^{2}$ | $5.7\times10^{-1}$ |
| P134V | 1.2 | $5.4\times10^{-1}$ | $2.2\times10^{5}$ | $1.1\times10^{-1}$ | 1.1 | $7.5\times10^{-1}$ | $9.0\times10^{-1}$ | N/A | $5.8\times10^{-2}$ | $1.2\times10^{2}$ | $6.2\times10^{-1}$ |

| Variant | $f_a$ | $f_a$ *p*–value | $(k_{cat}/K_M)^{MeP}_{chem.}$ ($M^{-1}s^{-1}$) | $(k_{cat}/K_M)^{MeP}_{chem.}$ *p*–value | FC1 | FC1 *p*–value | FC2 | FC3 | FC2/3 *p*–value | FC4 ($s^{-1}$) | FC4 *p*–value |
|---|---|---|---|---|---|---|---|---|---|---|---|
| R135G | 2.3 | **$2.0\times10^{-3}$** | $2.4\times10^{5}$ | $7.4\times10^{-2}$ | 2.6 | **$1.8\times10^{-3}$** | $9.6\times10^{-1}$ | N/A | $3.7\times10^{-1}$ | $1.2\times10^{2}$ | $7.5\times10^{-1}$ |
| R135V | 1.4 | $2.1\times10^{-1}$ | $6.2\times10^{5}$ | $9.6\times10^{-1}$ | 1.0 | $8.9\times10^{-1}$ | N/A | 1.1 | $3.4\times10^{-2}$ | $2.1\times10^{2}$ | $1.6\times10^{-1}$ |
| N136G | $6.1\times10^{-1}$ | $3.5\times10^{-2}$ | $5.6\times10^{5}$ | $7.9\times10^{-1}$ | 1.2 | $5.5\times10^{-1}$ | $9.4\times10^{-1}$ | N/A | $3.2\times10^{-1}$ | $8.5\times10^{1}$ | $2.5\times10^{-1}$ |
| N136V | $3.4\times10^{-1}$ | **$5.0\times10^{-3}$** | $5.4\times10^{5}$ | $7.0\times10^{-1}$ | 1.2 | $7.6\times10^{-1}$ | N/A | 1.1 | $4.3\times10^{-2}$ | $7.3\times10^{1}$ | $2.0\times10^{-1}$ |
| L137G | $2.6\times10^{-1}$ | **0** | $2.8\times10^{5}$ | $1.6\times10^{-1}$ | 4.2 | **$4.8\times10^{-3}$** | $7.7\times10^{-1}$ | N/A | **$3.8\times10^{-3}$** | $9.3\times10^{1}$ | $4.5\times10^{-1}$ |
| L137V | 1.1 | $7.5\times10^{-1}$ | $4.6\times10^{5}$ | $4.2\times10^{-1}$ | 1.2 | $4.8\times10^{-1}$ | N/A | 1.1 | $1.1\times10^{-2}$ | $1.7\times10^{2}$ | $5.1\times10^{-1}$ |
| W138G | $3.3\times10^{-1}$ | **0** | $3.1\times10^{5}$ | $1.1\times10^{-1}$ | 1.7 | $4.5\times10^{-2}$ | $7.0\times10^{-1}$ | N/A | **0** | $5.1\times10^{1}$ | $5.9\times10^{-2}$ |
| W138V | $3.6\times10^{-1}$ | **0** | $2.7\times10^{5}$ | $9.9\times10^{-2}$ | 1.8 | $7.7\times10^{-2}$ | $8.6\times10^{-1}$ | N/A | $1.3\times10^{-2}$ | $1.1\times10^{2}$ | $5.8\times10^{-1}$ |
| S139G | $3.4\times10^{-1}$ | **$3.0\times10^{-3}$** | $7.2\times10^{5}$ | $6.4\times10^{-1}$ | 1.5 | $1.4\times10^{-1}$ | $6.4\times10^{-1}$ | N/A | **$1.0\times10^{-3}$** | $1.4\times10^{2}$ | $9.3\times10^{-1}$ |
| S139V | 1.2 | $4.7\times10^{-1}$ | $3.4\times10^{5}$ | $1.1\times10^{-1}$ | 1.1 | $7.7\times10^{-1}$ | $9.6\times10^{-1}$ | N/A | $3.5\times10^{-1}$ | $1.4\times10^{2}$ | $9.0\times10^{-1}$ |
| T140G | $3.8\times10^{-1}$ | **$4.0\times10^{-3}$** | $3.5\times10^{5}$ | $2.1\times10^{-1}$ | 1.9 | $4.0\times10^{-2}$ | $7.2\times10^{-1}$ | N/A | **$2.0\times10^{-4}$** | $5.7\times10^{1}$ | $1.4\times10^{-1}$ |
| T140V | $3.7\times10^{-1}$ | **0** | $6.0\times10^{5}$ | $9.5\times10^{-1}$ | 1.8 | $1.2\times10^{-1}$ | $8.1\times10^{-1}$ | N/A | **$1.9\times10^{-3}$** | $1.4\times10^{2}$ | $9.8\times10^{-1}$ |
| T141G | $5.5\times10^{-1}$ | $1.1\times10^{-2}$ | $3.7\times10^{5}$ | $2.2\times10^{-1}$ | 1.5 | $1.7\times10^{-1}$ | $7.5\times10^{-1}$ | N/A | **$3.0\times10^{-4}$** | $5.0\times10^{1}$ | $9.7\times10^{-2}$ |
| T141V | 1.1 | $6.4\times10^{-1}$ | $8.4\times10^{4}$ | **$2.5\times10^{-2}$** | 3.1 | $1.7\times10^{-2}$ | $9.0\times10^{-1}$ | N/A | $8.9\times10^{-2}$ | $1.2\times10^{2}$ | $6.1\times10^{-1}$ |
| V142A | 1.0 | $8.7\times10^{-1}$ | $1.9\times10^{5}$ | **$2.4\times10^{-2}$** | 2.2 | **$2.5\times10^{-3}$** | $9.1\times10^{-1}$ | N/A | $2.9\times10^{-2}$ | $1.6\times10^{2}$ | $6.6\times10^{-1}$ |
| V142G | $1.6\times10^{-1}$ | **0** | $6.5\times10^{5}$ | $8.6\times10^{-1}$ | $9.6\times10^{-1}$ | $8.4\times10^{-1}$ | $7.6\times10^{-1}$ | N/A | **0** | $8.0\times10^{1}$ | $2.1\times10^{-1}$ |
| T143G | $6.0\times10^{-1}$ | $5.1\times10^{-2}$ | $5.9\times10^{5}$ | $9.5\times10^{-1}$ | 1.3 | $2.7\times10^{-1}$ | N/A | 1.0 | $8.4\times10^{-1}$ | $1.2\times10^{2}$ | $6.8\times10^{-1}$ |
| T143V | $8.5\times10^{-1}$ | $5.1\times10^{-1}$ | $3.3\times10^{5}$ | $1.5\times10^{-1}$ | 1.1 | $7.9\times10^{-1}$ | N/A | 1.0 | $5.5\times10^{-1}$ | $1.1\times10^{2}$ | $5.9\times10^{-1}$ |
| D144G | $1.7\times10^{-1}$ | **0** | $2.5\times10^{5}$ | $5.4\times10^{-2}$ | – | – | $8.7\times10^{-1}$ | N/A | **$1.8\times10^{-3}$** | $1.0\times10^{2}$ | $3.5\times10^{-1}$ |
| D144V | $\leq 6.7\times10^{-3}$ | **0** | $\leq 1.9\times10^{5}$ | $7.6\times10^{-2}$ | – | – | $6.0\times10^{-1}$ | N/A | **$1.\times10^{-4}$** | $\geq 1.1\times10^{2}$ | $6.3\times10^{-1}$ |
| Q145G | 1.3 | $3.9\times10^{-1}$ | $3.2\times10^{5}$ | $2.3\times10^{-1}$ | $9.5\times10^{-1}$ | $8.8\times10^{-1}$ | $5.5\times10^{-1}$ | N/A | $1.4\times10^{-2}$ | $6.2\times10^{1}$ | $2.0\times10^{-1}$ |
| Q145V | 1.1 | $6.8\times10^{-1}$ | $2.9\times10^{5}$ | $1.8\times10^{-1}$ | 1.0 | $9.4\times10^{-1}$ | $8.3\times10^{-1}$ | N/A | **$2.9\times10^{-3}$** | $1.2\times10^{2}$ | $8.0\times10^{-1}$ |
| L146G | $9.7\times10^{-2}$ | **0** | $1.6\times10^{5}$ | **$2.8\times10^{-2}$** | – | – | $6.5\times10^{-1}$ | N/A | **0** | $7.8\times10^{1}$ | $1.5\times10^{-1}$ |
| L146V | 1.6 | $3.9\times10^{-2}$ | $2.4\times10^{5}$ | $5.1\times10^{-2}$ | 1.1 | $7.8\times10^{-1}$ | N/A | 1.1 | $1.2\times10^{-1}$ | $1.2\times10^{2}$ | $6.6\times10^{-1}$ |
| G147A | $6.9\times10^{-1}$ | $1.8\times10^{-1}$ | $5.1\times10^{5}$ | $6.6\times10^{-1}$ | 1.8 | $3.1\times10^{-2}$ | $8.3\times10^{-1}$ | N/A | $3.2\times10^{-2}$ | $1.6\times10^{2}$ | $7.3\times10^{-1}$ |
| G147V | $8.1\times10^{-1}$ | $4.9\times10^{-1}$ | $3.8\times10^{5}$ | $2.6\times10^{-1}$ | 1.0 | $9.8\times10^{-1}$ | N/A | 1.0 | $6.5\times10^{-1}$ | $1.6\times10^{2}$ | $6.9\times10^{-1}$ |
| L148G | $1.4\times10^{-1}$ | **0** | $7.4\times10^{5}$ | $5.6\times10^{-1}$ | – | – | $6.3\times10^{-1}$ | N/A | **0** | $1.3\times10^{2}$ | $8.0\times10^{-1}$ |
| L148V | $5.4\times10^{-1}$ | $1.4\times10^{-2}$ | $2.9\times10^{5}$ | $1.2\times10^{-1}$ | 1.7 | $1.5\times10^{-1}$ | N/A | 1.0 | $4.4\times10^{-1}$ | $1.3\times10^{2}$ | $8.0\times10^{-1}$ |
| A149G | 1.2 | $4.8\times10^{-1}$ | $2.3\times10^{5}$ | $9.6\times10^{-2}$ | 2.5 | $1.1\times10^{-2}$ | N/A | 1.0 | $6.4\times10^{-1}$ | $1.1\times10^{2}$ | $5.2\times10^{-1}$ |
| A149V | $3.7\times10^{-1}$ | **$1.0\times10^{-3}$** | $9.1\times10^{5}$ | $1.4\times10^{-1}$ | $7.7\times10^{-1}$ | $4.5\times10^{-1}$ | $7.4\times10^{-1}$ | N/A | **$7.0\times10^{-4}$** | $1.2\times10^{2}$ | $7.0\times10^{-1}$ |
| T150G | 1.0 | $9.1\times10^{-1}$ | $3.1\times10^{5}$ | $2.1\times10^{-1}$ | 1.2 | $5.9\times10^{-1}$ | $8.1\times10^{-1}$ | N/A | **$1.3\times10^{-3}$** | $6.8\times10^{1}$ | $2.1\times10^{-1}$ |
| T150V | $8.1\times10^{-1}$ | $3.5\times10^{-1}$ | $3.1\times10^{5}$ | $1.5\times10^{-1}$ | 1.3 | $4.4\times10^{-1}$ | $9.0\times10^{-1}$ | N/A | $5.8\times10^{-2}$ | $1.3\times10^{2}$ | $8.2\times10^{-1}$ |
| N151G | $1.2\times10^{-1}$ | **0** | $1.5\times10^{5}$ | $8.1\times10^{-2}$ | – | – | $6.0\times10^{-1}$ | N/A | **$1.6\times10^{-3}$** | $9.5\times10^{1}$ | $4.5\times10^{-1}$ |
| N151V | $9.6\times10^{-1}$ | $8.3\times10^{-1}$ | $1.6\times10^{5}$ | **$2.4\times10^{-2}$** | 3.0 | **$3.8\times10^{-3}$** | $9.9\times10^{-1}$ | N/A | $9.1\times10^{-1}$ | $1.5\times10^{2}$ | $7.7\times10^{-1}$ |
| F152G | $3.1\times10^{-1}$ | **0** | $4.9\times10^{5}$ | $5.6\times10^{-1}$ | 2.2 | $2.6\times10^{-2}$ | $8.8\times10^{-1}$ | N/A | $2.2\times10^{-2}$ | $1.4\times10^{2}$ | $9.7\times10^{-1}$ |
| F152V | $6.9\times10^{-2}$ | **0** | $1.4\times10^{6}$ | **$2.5\times10^{-2}$** | – | – | $8.3\times10^{-1}$ | N/A | $1.5\times10^{-2}$ | $1.3\times10^{2}$ | $8.0\times10^{-1}$ |
| T153G | $9.7\times10^{-1}$ | $8.3\times10^{-1}$ | $3.6\times10^{5}$ | $2.1\times10^{-1}$ | 2.1 | $1.1\times10^{-2}$ | $8.6\times10^{-1}$ | N/A | $1.5\times10^{-2}$ | $1.1\times10^{2}$ | $4.8\times10^{-1}$ |
| T153V | 1.4 | $1.1\times10^{-1}$ | $2.8\times10^{5}$ | $7.5\times10^{-2}$ | 1.7 | $8.7\times10^{-2}$ | N/A | 1.1 | $1.2\times10^{-1}$ | $1.6\times10^{2}$ | $6.9\times10^{-1}$ |
| S154G | 1.1 | $7.5\times10^{-1}$ | $4.7\times10^{5}$ | $4.7\times10^{-1}$ | 1.0 | $9.2\times10^{-1}$ | $9.4\times10^{-1}$ | N/A | $3.8\times10^{-1}$ | $8.9\times10^{1}$ | $3.5\times10^{-1}$ |
| S154V | $3.3\times10^{-1}$ | **0** | $2.6\times10^{5}$ | $1.1\times10^{-1}$ | 1.7 | $2.0\times10^{-1}$ | $8.6\times10^{-1}$ | N/A | **$8.5\times10^{-3}$** | $1.2\times10^{2}$ | $7.8\times10^{-1}$ |
| K155G | $4.4\times10^{-1}$ | **$2.0\times10^{-3}$** | $4.9\times10^{5}$ | $5.6\times10^{-1}$ | 1.1 | $7.4\times10^{-1}$ | $8.7\times10^{-1}$ | N/A | $3.0\times10^{-2}$ | $1.1\times10^{2}$ | $6.1\times10^{-1}$ |
| K155V | $3.7\times10^{-1}$ | **$1.0\times10^{-3}$** | $9.5\times10^{5}$ | $9.0\times10^{-2}$ | – | – | $9.6\times10^{-1}$ | N/A | $3.7\times10^{-1}$ | $1.9\times10^{2}$ | $2.9\times10^{-1}$ |
| V156A | 1.1 | $7.3\times10^{-1}$ | $2.3\times10^{5}$ | $1.1\times10^{-1}$ | 1.8 | $1.5\times10^{-1}$ | $9.7\times10^{-1}$ | N/A | $5.4\times10^{-1}$ | $1.4\times10^{2}$ | $9.6\times10^{-1}$ |
| V156G | $9.2\times10^{-1}$ | $7.8\times10^{-1}$ | $1.7\times10^{5}$ | $1.0\times10^{-1}$ | – | – | $8.1\times10^{-1}$ | N/A | **$4.5\times10^{-3}$** | $1.3\times10^{2}$ | $8.6\times10^{-1}$ |
| V157A | $3.9\times10^{-1}$ | **$2.0\times10^{-3}$** | $8.2\times10^{5}$ | $2.7\times10^{-1}$ | 1.2 | $6.6\times10^{-1}$ | $9.3\times10^{-1}$ | N/A | $1.4\times10^{-1}$ | $1.8\times10^{2}$ | $4.6\times10^{-1}$ |
| V157G | $1.9\times10^{-1}$ | **0** | $1.9\times10^{5}$ | **$2.8\times10^{-2}$** | 2.0 | $3.1\times10^{-2}$ | $9.2\times10^{-1}$ | N/A | $8.3\times10^{-2}$ | $9.5\times10^{1}$ | $2.8\times10^{-1}$ |
| G158A | $5.7\times10^{-1}$ | $3.0\times10^{-2}$ | $8.0\times10^{5}$ | $3.9\times10^{-1}$ | 1.4 | $3.0\times10^{-1}$ | $8.9\times10^{-1}$ | N/A | $8.1\times10^{-2}$ | $2.3\times10^{2}$ | $1.3\times10^{-1}$ |
| G158V | $1.6\times10^{-1}$ | **0** | $3.8\times10^{5}$ | $2.3\times10^{-1}$ | $6.8\times10^{-1}$ | $2.5\times10^{-1}$ | $6.4\times10^{-1}$ | N/A | **0** | $1.2\times10^{2}$ | $6.5\times10^{-1}$ |
| V159A | 1.2 | $4.1\times10^{-1}$ | $7.6\times10^{4}$ | **$7.0\times10^{-3}$** | 4.2 | **$7.0\times10^{-4}$** | $9.7\times10^{-1}$ | N/A | $5.5\times10^{-1}$ | $1.3\times10^{2}$ | $8.9\times10^{-1}$ |
| V159G | $6.9\times10^{-1}$ | $1.7\times10^{-1}$ | $4.8\times10^{5}$ | $5.0\times10^{-1}$ | 1.6 | $1.5\times10^{-1}$ | $8.2\times10^{-1}$ | N/A | $2.4\times10^{-2}$ | $1.1\times10^{2}$ | $6.5\times10^{-1}$ |
| S160G | $8.0\times10^{-1}$ | $3.7\times10^{-1}$ | $1.1\times10^{6}$ | **$3.8\times10^{-2}$** | $5.4\times10^{-1}$ | $5.7\times10^{-2}$ | $6.2\times10^{-1}$ | N/A | **0** | $5.7\times10^{1}$ | $9.2\times10^{-2}$ |
| S160V | – | – | – | – | – | – | – | – | – | – | – |
| L161G | $1.9\times10^{-1}$ | **0** | $7.8\times10^{4}$ | **$1.7\times10^{-2}$** | 9.9 | **$6.9\times10^{-4}$** | N/A | 2.5 | **0** | $1.5\times10^{2}$ | $8.8\times10^{-1}$ |
| L161V | 1.9 | **$9.0\times10^{-3}$** | $2.6\times10^{5}$ | **$4.5\times10^{-2}$** | 1.6 | $2.8\times10^{-2}$ | N/A | 1.2 | **$3.3\times10^{-3}$** | $1.7\times10^{2}$ | $4.6\times10^{-1}$ |
| K162G | $\leq 1.0$ | 1.0 | $\leq 2.8\times10^{2}$ | **0** | $\leq 2.7\times10^{3}$ | **0** | – | – | – | – | – |
| K162V | $\leq 1.0$ | 1.0 | $\leq 6.7\times10^{2}$ | **0** | $\leq 1.7\times10^{4}$ | **0** | N/A | 3.8 | **$1.\times10^{-4}$** | $\geq 6.6\times10^{1}$ | $1.8\times10^{-1}$ |
| D163G | $2.2\times10^{-1}$ | **0** | $4.2\times10^{5}$ | $4.2\times10^{-1}$ | 4.2 | **$7.9\times10^{-4}$** | N/A | 3.8 | **$1.\times10^{-4}$** | $4.9\times10^{1}$ | $1.5\times10^{-1}$ |
| D163V | $1.0\times10^{-1}$ | **0** | $7.1\times10^{4}$ | **$3.7\times10^{-2}$** | $2.6\times10^{1}$ | **$2.0\times10^{-4}$** | N/A | $\geq 8.4$ | **0** | $3.5\times10^{1}$ | $8.0\times10^{-2}$ |
| R164G | $3.5\times10^{-1}$ | **0** | $6.0\times10^{3}$ | **$2.1\times10^{-2}$** | $2.0\times10^{3}$ | **0** | N/A | $\geq 3.3$ | **0** | $1.3\times10^{1}$ | **$2.4\times10^{-2}$** |
| R164V | $1.9\times10^{-1}$ | **0** | $1.5\times10^{4}$ | **$1.5\times10^{-2}$** | $1.4\times10^{3}$ | **0** | N/A | $\geq 2.2\times10^{2}$ | **$1.\times10^{-4}$** | $5.3\times10^{1}$ | $6.7\times10^{-2}$ |
| A165G | 1.7 | $8.3\times10^{-2}$ | $1.2\times10^{5}$ | $6.2\times10^{-2}$ | 1.2 | $6.2\times10^{-1}$ | N/A | 1.4 | **$3.0\times10^{-4}$** | $1.7\times10^{2}$ | $6.1\times10^{-1}$ |
| A165V | $\leq 4.4\times10^{-2}$ | **0** | $\leq 2.6\times10^{4}$ | **$1.6\times10^{-2}$** | – | – | N/A | 4.3 | **$1.\times10^{-4}$** | $\geq 4.6\times10^{2}$ | **0** |
| S166G | 1.6 | $1.3\times10^{-1}$ | $2.0\times10^{5}$ | $1.0\times10^{-1}$ | 2.8 | **$1.6\times10^{-3}$** | N/A | 1.1 | $2.4\times10^{-1}$ | $1.1\times10^{2}$ | $6.6\times10^{-1}$ |
| S166V | 1.8 | $2.8\times10^{-2}$ | $2.1\times10^{5}$ | $5.9\times10^{-2}$ | 1.8 | $8.9\times10^{-2}$ | $6.8\times10^{-1}$ | N/A | **$6.0\times10^{-4}$** | $1.2\times10^{2}$ | $7.9\times10^{-1}$ |
| I167G | – | – | – | – | – | – | – | – | – | – | – |
| I167V | 1.0 | $9.7\times10^{-1}$ | $5.2\times10^{5}$ | $5.9\times10^{-1}$ | $8.8\times10^{-1}$ | $6.5\times10^{-1}$ | N/A | 1.2 | **$2.7\times10^{-3}$** | $1.5\times10^{2}$ | $8.7\times10^{-1}$ |
| L168G | $7.1\times10^{-2}$ | **0** | $8.6\times10^{5}$ | $1.8\times10^{-1}$ | – | – | $4.2\times10^{-1}$ | N/A | **0** | $1.8\times10^{2}$ | $4.9\times10^{-1}$ |
| L168V | $4.6\times10^{-1}$ | **$5.0\times10^{-3}$** | $4.5\times10^{5}$ | $4.1\times10^{-1}$ | $8.4\times10^{-1}$ | $5.5\times10^{-1}$ | 1.0 | N/A | $9.5\times10^{-1}$ | $1.2\times10^{2}$ | $7.1\times10^{-1}$ |
| P169G | $\geq 2.9\times10^{-1}$ | **0** | $\geq 6.9\times10^{5}$ | $6.5\times10^{-1}$ | 1.1 | $5.3\times10^{-1}$ | $1.6\times10^{-1}$ | N/A | **0** | $\leq 1.6\times10^{2}$ | $7.2\times10^{-1}$ |
| P169V | $1.9\times10^{-1}$ | **0** | $1.5\times10^{6}$ | **$6.0\times10^{-3}$** | – | – | $3.2\times10^{-1}$ | N/A | **$3.0\times10^{-4}$** | $1.1\times10^{2}$ | $6.1\times10^{-1}$ |
| A170G | 1.0 | $8.4\times10^{-1}$ | $1.5\times10^{5}$ | $8.6\times10^{-2}$ | 3.6 | **$1.8\times10^{-3}$** | $9.5\times10^{-1}$ | N/A | $2.9\times10^{-1}$ | $1.1\times10^{2}$ | $5.9\times10^{-1}$ |
| A170V | $8.1\times10^{-1}$ | $3.3\times10^{-1}$ | $3.3\times10^{5}$ | $1.1\times10^{-1}$ | 1.4 | $2.3\times10^{-1}$ | N/A | 1.0 | $4.4\times10^{-1}$ | $1.4\times10^{2}$ | $9.3\times10^{-1}$ |
| G171A | $1.2\times10^{-1}$ | **0** | $8.4\times10^{5}$ | $3.7\times10^{-1}$ | 1.8 | $1.3\times10^{-1}$ | $7.9\times10^{-1}$ | N/A | **$3.8\times10^{-3}$** | $8.0\times10^{1}$ | $4.1\times10^{-1}$ |
| G171V | – | – | – | – | – | – | – | – | – | – | – |
| H172V | $2.5\times10^{-1}$ | **0** | $2.8\times10^{5}$ | $1.1\times10^{-1}$ | – | – | $8.1\times10^{-1}$ | N/A | **$9.0\times10^{-4}$** | $1.2\times10^{2}$ | $6.3\times10^{-1}$ |

| Variant | $f_a$ | $f_a$ $p$–value | $(k_{cat}/K_M)^{MeP}_{chem.}$ ($M^{-1}s^{-1}$) | $(k_{cat}/K_M)^{MeP}_{chem.}$ $p$–value | FC1 | FC1 $p$–value | FC2 | FC3 | FC2/3 $p$–value | FC4 ($s^{-1}$) | FC4 $p$–value |
|---|---|---|---|---|---|---|---|---|---|---|---|
| N173G | $5.1\times10^{-1}$ | $1.5\times10^{-2}$ | $4.5\times10^{5}$ | $4.4\times10^{-1}$ | 2.2 | **$8.6\times10^{-3}$** | $7.6\times10^{-1}$ | N/A | **$1.\times10^{-4}$** | $9.7\times10^{1}$ | $3.4\times10^{-1}$ |
| N173V | $2.7\times10^{-1}$ | **0** | $3.7\times10^{5}$ | $2.0\times10^{-1}$ | – | – | $8.8\times10^{-1}$ | N/A | $3.1\times10^{-2}$ | $1.2\times10^{2}$ | $7.4\times10^{-1}$ |
| P174G | $1.6\times10^{-1}$ | **0** | $4.6\times10^{5}$ | $3.9\times10^{-1}$ | 2.0 | $1.1\times10^{-2}$ | $7.1\times10^{-1}$ | N/A | **0** | $7.8\times10^{1}$ | $1.7\times10^{-1}$ |
| P174V | $1.4\times10^{-1}$ | **0** | $2.7\times10^{5}$ | $8.5\times10^{-2}$ | 1.8 | $9.8\times10^{-2}$ | $7.3\times10^{-1}$ | N/A | **0** | $8.9\times10^{1}$ | $2.5\times10^{-1}$ |
| T175G | $4.8\times10^{-1}$ | $1.0\times10^{-2}$ | $4.4\times10^{5}$ | $4.1\times10^{-1}$ | 2.5 | **$4.3\times10^{-3}$** | $8.9\times10^{-1}$ | N/A | $2.6\times10^{-2}$ | $1.2\times10^{2}$ | $7.0\times10^{-1}$ |
| T175V | 1.3 | $3.2\times10^{-1}$ | $4.9\times10^{5}$ | $5.0\times10^{-1}$ | $8.4\times10^{-1}$ | $4.3\times10^{-1}$ | N/A | 1.1 | $4.5\times10^{-2}$ | $1.7\times10^{2}$ | $5.4\times10^{-1}$ |
| G176A | ≤ 1.1 | $7.4\times10^{-1}$ | ≤ $2.8\times10^{5}$ | $1.4\times10^{-1}$ | ≤ 2.2 | $1.5\times10^{-2}$ | $8.4\times10^{-1}$ | N/A | $1.5\times10^{-2}$ | ≥ $8.6\times10^{1}$ | $2.7\times10^{-1}$ |
| G176V | $3.2\times10^{-1}$ | **0** | $7.4\times10^{5}$ | $5.3\times10^{-1}$ | – | – | $8.1\times10^{-1}$ | N/A | **$6.7\times10^{-3}$** | $1.3\times10^{2}$ | $8.6\times10^{-1}$ |
| A177G | $2.8\times10^{-1}$ | **$1.0\times10^{-3}$** | $3.6\times10^{5}$ | $2.8\times10^{-1}$ | 2.1 | $2.3\times10^{-2}$ | $9.8\times10^{-1}$ | N/A | $5.7\times10^{-1}$ | $1.1\times10^{2}$ | $5.7\times10^{-1}$ |
| A177V | $8.9\times10^{-1}$ | $5.4\times10^{-1}$ | $3.9\times10^{5}$ | $1.4\times10^{-1}$ | $8.3\times10^{-1}$ | $5.8\times10^{-1}$ | N/A | 1.1 | $2.0\times10^{-1}$ | $1.4\times10^{2}$ | $9.4\times10^{-1}$ |
| F178G | ≤ $1.3\times10^{-2}$ | **0** | ≤ $4.4\times10^{5}$ | $3.4\times10^{-1}$ | – | – | $9.3\times10^{-1}$ | N/A | $1.2\times10^{-1}$ | ≥ $1.6\times10^{2}$ | $7.5\times10^{-1}$ |
| F178V | $8.3\times10^{-1}$ | $3.5\times10^{-1}$ | $4.4\times10^{5}$ | $3.3\times10^{-1}$ | 1.3 | $4.2\times10^{-1}$ | $9.3\times10^{-1}$ | N/A | $1.4\times10^{-1}$ | $1.4\times10^{2}$ | $9.8\times10^{-1}$ |
| W179G | – | – | – | – | – | – | – | – | – | – | – |
| W179V | ≤ $3.2\times10^{-2}$ | **$5.0\times10^{-3}$** | ≤ $4.2\times10^{5}$ | $5.5\times10^{-1}$ | – | – | $9.3\times10^{-1}$ | N/A | $5.1\times10^{-1}$ | ≥ $9.4\times10^{1}$ | $6.9\times10^{-1}$ |
| F180G | $7.7\times10^{-1}$ | $2.9\times10^{-1}$ | $4.8\times10^{4}$ | **$1.2\times10^{-2}$** | 6.5 | **$1.\times10^{-4}$** | $8.9\times10^{-1}$ | N/A | $6.7\times10^{-2}$ | $5.4\times10^{1}$ | $1.0\times10^{-1}$ |
| F180V | 1.4 | $4.2\times10^{-1}$ | $7.3\times10^{4}$ | $1.1\times10^{-1}$ | 5.3 | **$7.4\times10^{-4}$** | $6.8\times10^{-1}$ | N/A | **$4.0\times10^{-4}$** | $1.2\times10^{2}$ | $7.5\times10^{-1}$ |
| D181G | $2.4\times10^{-1}$ | **0** | $1.6\times10^{5}$ | **$3.1\times10^{-2}$** | 1.9 | $7.3\times10^{-2}$ | N/A | 1.2 | **$5.0\times10^{-4}$** | $1.4\times10^{2}$ | 1.0 |
| D181V | $6.3\times10^{-1}$ | $7.4\times10^{-2}$ | $4.0\times10^{4}$ | **$1.0\times10^{-2}$** | $1.6\times10^{1}$ | **$9.0\times10^{-5}$** | – | – | – | – | – |
| D182G | 1.1 | $7.9\times10^{-1}$ | $5.5\times10^{4}$ | **$4.2\times10^{-2}$** | 5.2 | **$2.3\times10^{-4}$** | N/A | 1.7 | **0** | $6.0\times10^{1}$ | $1.9\times10^{-1}$ |
| D182V | 3.0 | **0** | $4.6\times10^{4}$ | **$2.4\times10^{-2}$** | 5.8 | **$1.\times10^{-4}$** | $9.2\times10^{-1}$ | N/A | $1.3\times10^{-1}$ | $9.9\times10^{1}$ | $3.9\times10^{-1}$ |
| T183G | $9.5\times10^{-1}$ | $7.5\times10^{-1}$ | $5.0\times10^{5}$ | $6.3\times10^{-1}$ | 1.3 | $5.0\times10^{-1}$ | $7.0\times10^{-1}$ | N/A | **$1.3\times10^{-3}$** | $1.1\times10^{2}$ | $5.7\times10^{-1}$ |
| T183V | 1.5 | $1.1\times10^{-1}$ | $3.7\times10^{5}$ | $1.1\times10^{-1}$ | 1.3 | $4.4\times10^{-1}$ | $9.1\times10^{-1}$ | N/A | $1.6\times10^{-1}$ | $1.7\times10^{2}$ | $5.0\times10^{-1}$ |
| T184G | $9.9\times10^{-1}$ | $9.0\times10^{-1}$ | $3.0\times10^{5}$ | $1.2\times10^{-1}$ | 1.3 | $3.0\times10^{-1}$ | $7.3\times10^{-1}$ | N/A | **0** | $8.1\times10^{1}$ | $2.3\times10^{-1}$ |
| T184V | $7.1\times10^{-1}$ | $1.7\times10^{-1}$ | $5.1\times10^{5}$ | $6.3\times10^{-1}$ | 1.1 | $6.7\times10^{-1}$ | $9.4\times10^{-1}$ | N/A | $1.7\times10^{-1}$ | $1.3\times10^{2}$ | $7.6\times10^{-1}$ |
| G185A | 1.5 | $1.5\times10^{-1}$ | $4.1\times10^{5}$ | $3.1\times10^{-1}$ | 1.4 | $2.2\times10^{-1}$ | $8.5\times10^{-1}$ | N/A | $1.8\times10^{-2}$ | $1.3\times10^{2}$ | $9.1\times10^{-1}$ |
| G185V | $5.5\times10^{-1}$ | $3.1\times10^{-2}$ | $5.0\times10^{4}$ | **$2.3\times10^{-2}$** | $1.4\times10^{1}$ | **$1.8\times10^{-4}$** | $5.7\times10^{-1}$ | N/A | **$5.0\times10^{-4}$** | $1.2\times10^{2}$ | $7.0\times10^{-1}$ |
| K186G | 1.2 | $4.3\times10^{-1}$ | $2.4\times10^{5}$ | $7.2\times10^{-2}$ | 2.2 | **$9.3\times10^{-3}$** | $9.5\times10^{-1}$ | N/A | $3.0\times10^{-1}$ | $9.8\times10^{1}$ | $3.2\times10^{-1}$ |
| K186V | 1.4 | $1.6\times10^{-1}$ | $1.2\times10^{5}$ | **$1.7\times10^{-2}$** | 2.7 | **$4.4\times10^{-3}$** | N/A | 1.0 | $4.3\times10^{-1}$ | $1.5\times10^{2}$ | $8.2\times10^{-1}$ |
| F187G | $1.2\times10^{-1}$ | **0** | $1.3\times10^{5}$ | $6.7\times10^{-2}$ | – | – | N/A | 1.0 | $8.6\times10^{-1}$ | $9.5\times10^{1}$ | $3.3\times10^{-1}$ |
| F187V | $3.5\times10^{-1}$ | **0** | $2.9\times10^{5}$ | $1.4\times10^{-1}$ | – | – | $9.8\times10^{-1}$ | N/A | $4.5\times10^{-1}$ | $1.1\times10^{2}$ | $5.9\times10^{-1}$ |
| I188V | 1.3 | $2.1\times10^{-1}$ | $2.9\times10^{5}$ | $8.3\times10^{-2}$ | 1.9 | $4.2\times10^{-2}$ | N/A | 1.0 | $8.1\times10^{-1}$ | $2.0\times10^{2}$ | $2.2\times10^{-1}$ |
| T189G | $2.0\times10^{-1}$ | **0** | $3.5\times10^{5}$ | $1.8\times10^{-1}$ | 1.1 | $7.3\times10^{-1}$ | $8.7\times10^{-1}$ | N/A | $4.2\times10^{-2}$ | $1.1\times10^{2}$ | $5.3\times10^{-1}$ |
| S190G | $2.8\times10^{-1}$ | **0** | $2.8\times10^{5}$ | $1.4\times10^{-1}$ | 1.6 | $1.1\times10^{-1}$ | N/A | 1.7 | **0** | $4.0\times10^{1}$ | $7.3\times10^{-2}$ |
| S190V | – | – | – | – | – | – | $3.1\times10^{-1}$ | N/A | **$9.0\times10^{-4}$** | – | – |
| T191G | 1.5 | $1.8\times10^{-1}$ | $3.6\times10^{5}$ | $2.7\times10^{-1}$ | 1.6 | $1.5\times10^{-1}$ | $9.1\times10^{-1}$ | N/A | $8.9\times10^{-2}$ | $1.2\times10^{2}$ | $7.1\times10^{-1}$ |
| T191V | $2.1\times10^{-1}$ | **0** | $3.5\times10^{5}$ | $2.6\times10^{-1}$ | – | – | $9.9\times10^{-1}$ | N/A | $8.2\times10^{-1}$ | $1.0\times10^{2}$ | $4.6\times10^{-1}$ |
| Y192V | ≤ $7.8\times10^{-2}$ | **0** | ≤ $4.5\times10^{5}$ | $5.3\times10^{-1}$ | – | – | – | – | – | – | – |
| Y193G | – | – | – | – | – | – | – | – | – | – | – |
| Y193V | $2.8\times10^{-1}$ | **0** | $1.4\times10^{5}$ | **$4.4\times10^{-2}$** | – | – | $8.4\times10^{-1}$ | N/A | $4.2\times10^{-2}$ | $1.1\times10^{2}$ | $5.2\times10^{-1}$ |
| T194G | 1.5 | $1.8\times10^{-1}$ | $4.6\times10^{5}$ | $4.9\times10^{-1}$ | 1.0 | $9.9\times10^{-1}$ | $9.8\times10^{-1}$ | N/A | $6.4\times10^{-1}$ | $9.3\times10^{1}$ | $3.5\times10^{-1}$ |
| T194V | $7.9\times10^{-1}$ | $2.8\times10^{-1}$ | $9.2\times10^{5}$ | $6.8\times10^{-2}$ | $6.8\times10^{-1}$ | $1.7\times10^{-1}$ | N/A | 1.0 | $3.0\times10^{-1}$ | $2.0\times10^{2}$ | $1.5\times10^{-1}$ |
| K195G | 1.5 | $1.2\times10^{-1}$ | $3.7\times10^{5}$ | $2.2\times10^{-1}$ | 1.2 | $4.5\times10^{-1}$ | $9.8\times10^{-1}$ | N/A | $7.2\times10^{-1}$ | $1.3\times10^{2}$ | $8.4\times10^{-1}$ |
| K195V | 1.4 | $9.4\times10^{-2}$ | $2.6\times10^{5}$ | **$4.3\times10^{-2}$** | 1.6 | $1.3\times10^{-1}$ | N/A | 1.1 | $1.4\times10^{-1}$ | $1.5\times10^{2}$ | $8.6\times10^{-1}$ |
| E196G | 1.4 | $2.1\times10^{-1}$ | $3.2\times10^{5}$ | $1.8\times10^{-1}$ | 1.1 | $7.6\times10^{-1}$ | $8.9\times10^{-1}$ | N/A | $4.3\times10^{-2}$ | $1.1\times10^{2}$ | $5.1\times10^{-1}$ |
| E196V | 1.5 | $1.1\times10^{-1}$ | $2.8\times10^{5}$ | $7.5\times10^{-2}$ | 1.3 | $3.0\times10^{-1}$ | N/A | 1.1 | $1.0\times10^{-1}$ | $1.6\times10^{2}$ | $6.1\times10^{-1}$ |
| L197G | $2.2\times10^{-1}$ | **0** | $6.4\times10^{5}$ | $9.0\times10^{-1}$ | 1.6 | $1.7\times10^{-1}$ | $8.0\times10^{-1}$ | N/A | **$1.3\times10^{-3}$** | $1.7\times10^{2}$ | $5.9\times10^{-1}$ |
| L197V | $9.2\times10^{-1}$ | $6.6\times10^{-1}$ | $5.2\times10^{5}$ | $5.2\times10^{-1}$ | $8.4\times10^{-1}$ | $6.0\times10^{-1}$ | $9.8\times10^{-1}$ | N/A | $7.3\times10^{-1}$ | $1.5\times10^{2}$ | $8.1\times10^{-1}$ |
| P198G | $5.3\times10^{-1}$ | $2.4\times10^{-2}$ | $2.0\times10^{5}$ | $8.9\times10^{-2}$ | 3.2 | **$2.5\times10^{-3}$** | $8.9\times10^{-1}$ | N/A | $3.4\times10^{-2}$ | $1.4\times10^{2}$ | $9.3\times10^{-1}$ |
| P198V | 1.3 | $2.9\times10^{-1}$ | $8.7\times10^{4}$ | **$4.3\times10^{-2}$** | 3.1 | **$2.4\times10^{-3}$** | $9.8\times10^{-1}$ | N/A | $6.9\times10^{-1}$ | $1.5\times10^{2}$ | $8.1\times10^{-1}$ |
| K199G | 1.4 | $2.4\times10^{-1}$ | $4.8\times10^{5}$ | $5.6\times10^{-1}$ | $9.4\times10^{-1}$ | $8.1\times10^{-1}$ | $9.5\times10^{-1}$ | N/A | $3.2\times10^{-1}$ | $1.3\times10^{2}$ | $8.4\times10^{-1}$ |
| K199V | 1.2 | $5.0\times10^{-1}$ | $4.6\times10^{5}$ | $4.1\times10^{-1}$ | 1.2 | $5.9\times10^{-1}$ | N/A | 1.1 | $2.2\times10^{-1}$ | $1.5\times10^{2}$ | $8.5\times10^{-1}$ |
| W200G | ≤ $4.4\times10^{-2}$ | **0** | ≤ $1.2\times10^{5}$ | **$4.1\times10^{-2}$** | – | – | $8.9\times10^{-1}$ | N/A | $1.4\times10^{-2}$ | ≥ $9.7\times10^{1}$ | $3.8\times10^{-1}$ |
| W200V | $1.5\times10^{-1}$ | **0** | $1.6\times10^{5}$ | $5.9\times10^{-2}$ | – | – | $7.2\times10^{-1}$ | N/A | **$9.0\times10^{-4}$** | $1.4\times10^{2}$ | 1.0 |
| V201A | 1.3 | $3.6\times10^{-1}$ | $2.6\times10^{5}$ | $1.5\times10^{-1}$ | 1.6 | $1.6\times10^{-1}$ | N/A | 1.1 | $1.7\times10^{-1}$ | $1.9\times10^{2}$ | $3.9\times10^{-1}$ |
| V201G | $2.6\times10^{-1}$ | **0** | $2.7\times10^{5}$ | $6.1\times10^{-2}$ | 1.6 | $1.5\times10^{-1}$ | $8.4\times10^{-1}$ | N/A | **$4.1\times10^{-3}$** | $1.3\times10^{2}$ | $8.4\times10^{-1}$ |
| N202G | 1.2 | $3.5\times10^{-1}$ | $2.5\times10^{5}$ | $6.6\times10^{-2}$ | 2.3 | **$6.1\times10^{-3}$** | $9.4\times10^{-1}$ | N/A | $1.5\times10^{-1}$ | $1.1\times10^{2}$ | $5.7\times10^{-1}$ |
| N202V | 1.2 | $3.9\times10^{-1}$ | $4.2\times10^{5}$ | $2.5\times10^{-1}$ | $8.3\times10^{-1}$ | $6.6\times10^{-1}$ | N/A | 1.1 | $1.3\times10^{-1}$ | $1.4\times10^{2}$ | $9.2\times10^{-1}$ |
| D203G | ≤ 1.2 | $4.1\times10^{-1}$ | ≤ $1.4\times10^{5}$ | **$4.2\times10^{-2}$** | ≤ 3.2 | **$7.6\times10^{-3}$** | N/A | 1.1 | $3.6\times10^{-2}$ | ≥ $8.5\times10^{1}$ | $2.1\times10^{-1}$ |
| D203V | $9.3\times10^{-1}$ | $6.9\times10^{-1}$ | $4.5\times10^{5}$ | $3.6\times10^{-1}$ | 1.3 | $4.7\times10^{-1}$ | N/A | 1.1 | $6.7\times10^{-2}$ | $1.8\times10^{2}$ | $3.8\times10^{-1}$ |
| F204G | $2.1\times10^{-2}$ | **0** | $2.1\times10^{5}$ | $1.4\times10^{-1}$ | – | – | $9.4\times10^{-1}$ | N/A | $3.0\times10^{-1}$ | $1.2\times10^{2}$ | $6.9\times10^{-1}$ |
| F204V | ≤ $6.9\times10^{-1}$ | $9.9\times10^{-2}$ | ≤ $1.2\times10^{5}$ | **$2.3\times10^{-2}$** | ≤ 4.5 | **$9.7\times10^{-4}$** | $9.7\times10^{-1}$ | N/A | $5.0\times10^{-1}$ | ≥ $1.4\times10^{2}$ | $9.9\times10^{-1}$ |
| N205V | $3.8\times10^{-1}$ | **0** | $8.1\times10^{4}$ | **$2.3\times10^{-2}$** | 4.9 | **$1.2\times10^{-3}$** | $7.7\times10^{-1}$ | N/A | **$2.0\times10^{-3}$** | $1.3\times10^{2}$ | $8.5\times10^{-1}$ |
| N206G | $7.8\times10^{-1}$ | $2.6\times10^{-1}$ | $1.9\times10^{6}$ | **$1.0\times10^{-3}$** | $4.7\times10^{-1}$ | $3.0\times10^{-2}$ | $8.9\times10^{-1}$ | N/A | $3.1\times10^{-2}$ | $1.1\times10^{2}$ | $5.7\times10^{-1}$ |
| N206V | 1.0 | $9.8\times10^{-1}$ | $4.3\times10^{5}$ | $1.4\times10^{-1}$ | 1.0 | $8.6\times10^{-1}$ | N/A | 1.0 | $2.3\times10^{-1}$ | $1.5\times10^{2}$ | $7.7\times10^{-1}$ |
| K207G | 1.4 | $1.2\times10^{-1}$ | $3.3\times10^{5}$ | $2.0\times10^{-1}$ | 1.2 | $4.9\times10^{-1}$ | $9.6\times10^{-1}$ | N/A | $3.7\times10^{-1}$ | $1.1\times10^{2}$ | $5.8\times10^{-1}$ |
| K207V | $9.5\times10^{-1}$ | $8.3\times10^{-1}$ | $4.3\times10^{5}$ | $3.2\times10^{-1}$ | – | – | N/A | 1.0 | $4.3\times10^{-1}$ | $1.4\times10^{2}$ | $9.4\times10^{-1}$ |
| N208G | 1.3 | $3.5\times10^{-1}$ | $2.9\times10^{5}$ | $1.8\times10^{-1}$ | 1.8 | $2.7\times10^{-2}$ | $9.0\times10^{-1}$ | N/A | $3.3\times10^{-1}$ | $1.2\times10^{2}$ | $7.8\times10^{-1}$ |
| N208V | 1.7 | $2.5\times10^{-2}$ | $1.5\times10^{5}$ | **$3.9\times10^{-2}$** | 2.4 | **$6.5\times10^{-3}$** | N/A | 1.0 | $6.0\times10^{-1}$ | $1.5\times10^{2}$ | $8.3\times10^{-1}$ |
| V209A | 1.6 | $5.9\times10^{-2}$ | $2.8\times10^{5}$ | $1.1\times10^{-1}$ | 1.4 | $2.6\times10^{-1}$ | N/A | 1.1 | $3.6\times10^{-2}$ | $1.8\times10^{2}$ | $4.0\times10^{-1}$ |
| V209G | 1.1 | $7.5\times10^{-1}$ | $3.2\times10^{5}$ | $1.3\times10^{-1}$ | 1.6 | $1.4\times10^{-1}$ | $8.5\times10^{-1}$ | N/A | **$8.6\times10^{-3}$** | $8.5\times10^{1}$ | $3.4\times10^{-1}$ |
| P210G | $8.3\times10^{-1}$ | $5.1\times10^{-1}$ | $4.7\times10^{5}$ | $4.8\times10^{-1}$ | 1.9 | $2.4\times10^{-2}$ | $9.1\times10^{-1}$ | N/A | $1.2\times10^{-1}$ | $1.2\times10^{2}$ | $7.1\times10^{-1}$ |
| P210V | $5.1\times10^{-1}$ | $1.7\times10^{-2}$ | $8.3\times10^{5}$ | $2.5\times10^{-1}$ | $6.3\times10^{-1}$ | $1.7\times10^{-1}$ | $7.9\times10^{-1}$ | N/A | **$1.0\times10^{-3}$** | $1.2\times10^{2}$ | $7.6\times10^{-1}$ |
| A211G | 1.5 | $1.7\times10^{-1}$ | $4.1\times10^{5}$ | $3.8\times10^{-1}$ | 1.0 | $8.4\times10^{-1}$ | $9.8\times10^{-1}$ | N/A | $7.6\times10^{-1}$ | $8.2\times10^{1}$ | $3.0\times10^{-1}$ |
| A211V | 1.4 | $2.0\times10^{-1}$ | $3.2\times10^{5}$ | $1.3\times10^{-1}$ | 1.3 | $4.5\times10^{-1}$ | N/A | 1.0 | $6.9\times10^{-1}$ | $1.4\times10^{2}$ | $9.9\times10^{-1}$ |
| Q212G | 1.4 | $1.6\times10^{-1}$ | $4.9\times10^{5}$ | $6.1\times10^{-1}$ | $9.5\times10^{-1}$ | $8.7\times10^{-1}$ | $9.0\times10^{-1}$ | N/A | $8.8\times10^{-2}$ | $1.3\times10^{2}$ | $7.2\times10^{-1}$ |

| Variant | $f_a$ | $f_a$ $p$–value | $(k_{cat}/K_M)^{MeP}_{chem.}$ ($M^{-1}s^{-1}$) | $(k_{cat}/K_M)^{MeP}_{chem.}$ $p$–value | FC1 | FC1 $p$–value | FC2 | FC3 | FC2/3 $p$–value | FC4 ($s^{-1}$) | FC4 $p$–value |
|---|---|---|---|---|---|---|---|---|---|---|---|
| Q212V | 1.3 | $1.7\times10^{-1}$ | $3.2\times10^{5}$ | $1.1\times10^{-1}$ | 1.1 | $8.6\times10^{-1}$ | N/A | 1.0 | $4.4\times10^{-1}$ | $1.4\times10^{2}$ | $9.0\times10^{-1}$ |
| L213G | 1.9 | $3.6\times10^{-2}$ | $1.8\times10^{5}$ | $8.2\times10^{-2}$ | 1.7 | $5.4\times10^{-2}$ | $9.1\times10^{-1}$ | N/A | $8.7\times10^{-2}$ | $9.5\times10^{1}$ | $4.2\times10^{-1}$ |
| L213V | 1.1 | $7.2\times10^{-1}$ | $3.4\times10^{5}$ | $1.1\times10^{-1}$ | 1.5 | $1.9\times10^{-1}$ | $9.1\times10^{-1}$ | N/A | $8.0\times10^{-2}$ | $1.4\times10^{2}$ | $9.1\times10^{-1}$ |
| V214A | 1.1 | $5.4\times10^{-1}$ | $4.5\times10^{5}$ | $1.9\times10^{-1}$ | 1.1 | $7.9\times10^{-1}$ | $9.3\times10^{-1}$ | N/A | $7.1\times10^{-2}$ | $1.5\times10^{2}$ | $7.6\times10^{-1}$ |
| V214G | 1.1 | $6.2\times10^{-1}$ | $1.9\times10^{5}$ | $6.1\times10^{-2}$ | 2.7 | **$2.0\times10^{-3}$** | $6.8\times10^{-1}$ | N/A | **0** | $7.1\times10^{1}$ | $1.6\times10^{-1}$ |
| A215G | 1.6 | $2.1\times10^{-1}$ | $3.3\times10^{5}$ | $3.4\times10^{-1}$ | 1.3 | $3.4\times10^{-1}$ | $9.0\times10^{-1}$ | N/A | $1.1\times10^{-1}$ | $9.9\times10^{1}$ | $4.8\times10^{-1}$ |
| A215V | 1.7 | $4.2\times10^{-2}$ | $3.0\times10^{5}$ | $1.2\times10^{-1}$ | 1.1 | $7.7\times10^{-1}$ | $9.9\times10^{-1}$ | N/A | $7.3\times10^{-1}$ | $1.2\times10^{2}$ | $6.6\times10^{-1}$ |
| N216G | 1.5 | $9.6\times10^{-2}$ | $3.9\times10^{5}$ | $2.5\times10^{-1}$ | 1.2 | $5.3\times10^{-1}$ | $8.7\times10^{-1}$ | N/A | $2.3\times10^{-2}$ | $1.5\times10^{2}$ | $9.2\times10^{-1}$ |
| N216V | $7.0\times10^{-1}$ | $4.3\times10^{-1}$ | $1.8\times10^{5}$ | $1.5\times10^{-1}$ | – | – | – | – | – | – | – |
| G217A | $1.6\times10^{-1}$ | **0** | $1.3\times10^{5}$ | **$4.7\times10^{-2}$** | 2.6 | $4.7\times10^{-2}$ | $9.5\times10^{-1}$ | N/A | $4.9\times10^{-1}$ | $1.2\times10^{2}$ | $6.7\times10^{-1}$ |
| G217V | 1.3 | $2.2\times10^{-1}$ | $2.3\times10^{5}$ | $5.8\times10^{-2}$ | 2.5 | **$7.5\times10^{-3}$** | N/A | 1.0 | $6.8\times10^{-1}$ | $1.5\times10^{2}$ | $7.7\times10^{-1}$ |
| W218G | $1.5\times10^{-1}$ | **0** | $1.4\times10^{5}$ | **$3.1\times10^{-2}$** | 3.7 | **$5.0\times10^{-3}$** | $4.9\times10^{-1}$ | N/A | **0** | $5.5\times10^{1}$ | $8.4\times10^{-2}$ |
| W218V | $3.1\times10^{-1}$ | **0** | $7.8\times10^{4}$ | **$2.0\times10^{-2}$** | 4.1 | **$2.3\times10^{-3}$** | $4.2\times10^{-1}$ | N/A | **0** | $6.3\times10^{1}$ | $9.9\times10^{-2}$ |
| N219G | $9.7\times10^{-1}$ | $8.8\times10^{-1}$ | $3.5\times10^{5}$ | $1.4\times10^{-1}$ | 2.0 | $1.4\times10^{-2}$ | $9.6\times10^{-1}$ | N/A | $3.0\times10^{-1}$ | $1.1\times10^{2}$ | $4.4\times10^{-1}$ |
| N219V | $9.6\times10^{-1}$ | $7.3\times10^{-1}$ | $4.7\times10^{5}$ | $3.8\times10^{-1}$ | 1.4 | $3.0\times10^{-1}$ | $9.9\times10^{-1}$ | N/A | $8.5\times10^{-1}$ | $1.9\times10^{2}$ | $2.0\times10^{-1}$ |
| T220G | $4.4\times10^{-1}$ | **$8.0\times10^{-3}$** | $5.4\times10^{5}$ | $7.7\times10^{-1}$ | $8.5\times10^{-1}$ | $6.8\times10^{-1}$ | $7.9\times10^{-1}$ | N/A | **$3.0\times10^{-4}$** | $1.5\times10^{2}$ | $8.5\times10^{-1}$ |
| T220V | 1.0 | $9.0\times10^{-1}$ | $3.6\times10^{5}$ | $1.5\times10^{-1}$ | 1.4 | $3.1\times10^{-1}$ | N/A | 1.0 | $4.3\times10^{-1}$ | $1.4\times10^{2}$ | $9.7\times10^{-1}$ |
| L221G | $9.8\times10^{-1}$ | $9.5\times10^{-1}$ | $1.9\times10^{5}$ | $1.1\times10^{-1}$ | 2.3 | $2.4\times10^{-2}$ | $8.9\times10^{-1}$ | N/A | $5.2\times10^{-2}$ | $9.5\times10^{1}$ | $4.4\times10^{-1}$ |
| L221V | 1.1 | $7.3\times10^{-1}$ | $4.0\times10^{5}$ | $3.4\times10^{-1}$ | 1.6 | $1.5\times10^{-1}$ | N/A | 1.0 | $7.7\times10^{-1}$ | $1.6\times10^{2}$ | $8.3\times10^{-1}$ |
| L222G | $8.2\times10^{-1}$ | $4.7\times10^{-1}$ | $3.3\times10^{5}$ | $2.3\times10^{-1}$ | 1.6 | $6.3\times10^{-2}$ | $8.4\times10^{-1}$ | N/A | $1.6\times10^{-2}$ | $1.1\times10^{2}$ | $5.0\times10^{-1}$ |
| L222V | $8.0\times10^{-1}$ | $3.3\times10^{-1}$ | $4.5\times10^{5}$ | $3.4\times10^{-1}$ | $9.7\times10^{-1}$ | $8.5\times10^{-1}$ | N/A | 1.1 | $1.4\times10^{-1}$ | $1.7\times10^{2}$ | $5.1\times10^{-1}$ |
| P223G | $9.9\times10^{-1}$ | $9.6\times10^{-1}$ | $3.7\times10^{5}$ | $2.8\times10^{-1}$ | 1.5 | $1.9\times10^{-1}$ | $8.2\times10^{-1}$ | N/A | **$2.2\times10^{-3}$** | $1.1\times10^{2}$ | $4.8\times10^{-1}$ |
| P223V | 1.1 | $6.7\times10^{-1}$ | $1.4\times10^{5}$ | **$3.4\times10^{-2}$** | 2.5 | **$6.7\times10^{-3}$** | $9.7\times10^{-1}$ | N/A | $5.7\times10^{-1}$ | $1.2\times10^{2}$ | $6.3\times10^{-1}$ |
| I224G | 1.0 | $9.8\times10^{-1}$ | $5.1\times10^{5}$ | $6.5\times10^{-1}$ | $8.6\times10^{-1}$ | $6.4\times10^{-1}$ | $8.8\times10^{-1}$ | N/A | $4.8\times10^{-2}$ | $1.2\times10^{2}$ | $6.1\times10^{-1}$ |
| I224V | $5.2\times10^{-1}$ | $2.4\times10^{-2}$ | $5.7\times10^{5}$ | $8.3\times10^{-1}$ | $9.4\times10^{-1}$ | $7.9\times10^{-1}$ | N/A | 1.2 | **$1.1\times10^{-3}$** | $1.3\times10^{2}$ | $8.2\times10^{-1}$ |
| N225G | 1.8 | $7.2\times10^{-2}$ | $2.2\times10^{5}$ | $1.2\times10^{-1}$ | 1.8 | $8.6\times10^{-2}$ | $9.9\times10^{-1}$ | N/A | $7.9\times10^{-1}$ | $1.3\times10^{2}$ | $8.8\times10^{-1}$ |
| N225V | 1.1 | $6.3\times10^{-1}$ | $5.0\times10^{5}$ | $5.2\times10^{-1}$ | 1.0 | $8.9\times10^{-1}$ | N/A | 1.1 | **$5.7\times10^{-3}$** | $1.8\times10^{2}$ | $2.9\times10^{-1}$ |
| Q226G | $9.2\times10^{-1}$ | $8.0\times10^{-1}$ | $3.0\times10^{5}$ | $2.1\times10^{-1}$ | 1.9 | $6.8\times10^{-2}$ | $7.2\times10^{-1}$ | N/A | **$2.0\times10^{-4}$** | $9.3\times10^{1}$ | $4.0\times10^{-1}$ |
| Q226V | 1.3 | $1.2\times10^{-1}$ | $3.7\times10^{5}$ | $6.5\times10^{-2}$ | 1.0 | $9.2\times10^{-1}$ | N/A | 1.1 | $1.5\times10^{-1}$ | $1.4\times10^{2}$ | $9.8\times10^{-1}$ |
| Y227G | $3.5\times10^{-1}$ | **$1.0\times10^{-3}$** | $4.0\times10^{5}$ | $3.6\times10^{-1}$ | $7.3\times10^{-1}$ | $5.2\times10^{-1}$ | $4.9\times10^{-1}$ | N/A | **$2.0\times10^{-4}$** | $8.7\times10^{1}$ | $2.9\times10^{-1}$ |
| Y227V | $5.2\times10^{-1}$ | $2.2\times10^{-2}$ | $3.7\times10^{5}$ | $2.4\times10^{-1}$ | 1.1 | $9.0\times10^{-1}$ | $6.9\times10^{-1}$ | N/A | **$4.0\times10^{-4}$** | $1.0\times10^{2}$ | $4.8\times10^{-1}$ |
| T228G | $5.3\times10^{-1}$ | $1.8\times10^{-2}$ | $2.3\times10^{5}$ | **$4.4\times10^{-2}$** | 4.2 | **$1.\times10^{-4}$** | N/A | 1.0 | $4.8\times10^{-1}$ | $1.7\times10^{2}$ | $6.0\times10^{-1}$ |
| T228V | $8.1\times10^{-1}$ | $2.7\times10^{-1}$ | $5.3\times10^{5}$ | $5.7\times10^{-1}$ | $9.2\times10^{-1}$ | $8.3\times10^{-1}$ | N/A | 1.1 | $7.7\times10^{-2}$ | $1.5\times10^{2}$ | $7.5\times10^{-1}$ |
| E229G | $6.3\times10^{-1}$ | $5.0\times10^{-2}$ | $2.6\times10^{5}$ | $8.0\times10^{-2}$ | 1.4 | $2.1\times10^{-1}$ | $8.6\times10^{-1}$ | N/A | **$8.7\times10^{-3}$** | $8.7\times10^{1}$ | $2.4\times10^{-1}$ |
| E229V | $5.8\times10^{-1}$ | $1.5\times10^{-2}$ | $2.4\times10^{5}$ | $1.2\times10^{-1}$ | 2.2 | $3.7\times10^{-2}$ | $9.5\times10^{-1}$ | N/A | $2.7\times10^{-1}$ | $1.1\times10^{2}$ | $5.5\times10^{-1}$ |
| S230G | $\leq 5.2\times10^{-1}$ | $1.4\times10^{-2}$ | $\leq 5.5\times10^{5}$ | $7.6\times10^{-1}$ | $\leq 1.1$ | $8.2\times10^{-1}$ | $8.0\times10^{-1}$ | N/A | **$1.0\times10^{-3}$** | $\geq 8.4\times10^{1}$ | $2.3\times10^{-1}$ |
| S230V | $3.7\times10^{-1}$ | **$1.0\times10^{-3}$** | $5.6\times10^{5}$ | $7.8\times10^{-1}$ | $9.1\times10^{-1}$ | $7.9\times10^{-1}$ | $5.8\times10^{-1}$ | N/A | **$1.\times10^{-4}$** | $1.1\times10^{2}$ | $4.3\times10^{-1}$ |
| S231G | $9.3\times10^{-1}$ | $7.5\times10^{-1}$ | $4.2\times10^{5}$ | $3.6\times10^{-1}$ | $8.8\times10^{-1}$ | $6.9\times10^{-1}$ | $7.9\times10^{-1}$ | N/A | **$7.0\times10^{-4}$** | $5.9\times10^{1}$ | $1.9\times10^{-1}$ |
| S231V | $7.0\times10^{-1}$ | $1.4\times10^{-1}$ | $4.9\times10^{5}$ | $5.4\times10^{-1}$ | 1.2 | $5.9\times10^{-1}$ | $9.4\times10^{-1}$ | N/A | $2.4\times10^{-1}$ | $1.2\times10^{2}$ | $6.3\times10^{-1}$ |
| E232G | $8.6\times10^{-1}$ | $5.5\times10^{-1}$ | $4.1\times10^{5}$ | $3.4\times10^{-1}$ | 1.6 | $7.9\times10^{-2}$ | N/A | 1.0 | $8.9\times10^{-1}$ | $1.5\times10^{2}$ | $8.7\times10^{-1}$ |
| E232V | $6.5\times10^{-1}$ | $1.1\times10^{-1}$ | $7.9\times10^{5}$ | $3.9\times10^{-1}$ | $8.2\times10^{-1}$ | $5.7\times10^{-1}$ | $9.3\times10^{-1}$ | N/A | $2.2\times10^{-1}$ | $1.4\times10^{2}$ | $9.7\times10^{-1}$ |
| D233G | $3.2\times10^{-1}$ | **0** | $9.1\times10^{4}$ | **$1.9\times10^{-2}$** | 4.5 | **$1.1\times10^{-3}$** | $2.7\times10^{-1}$ | N/A | **0** | $4.1\times10^{1}$ | $5.0\times10^{-2}$ |
| D233V | $5.8\times10^{-1}$ | $1.3\times10^{-2}$ | $1.5\times10^{5}$ | **$2.8\times10^{-2}$** | 3.3 | **$3.7\times10^{-3}$** | $3.4\times10^{-1}$ | N/A | **0** | $5.9\times10^{1}$ | $9.2\times10^{-2}$ |
| N234G | $9.6\times10^{-1}$ | $8.9\times10^{-1}$ | $4.4\times10^{5}$ | $4.4\times10^{-1}$ | 1.3 | $3.9\times10^{-1}$ | $9.1\times10^{-1}$ | N/A | $1.3\times10^{-1}$ | $1.0\times10^{2}$ | $5.1\times10^{-1}$ |
| N234V | $9.4\times10^{-1}$ | $8.0\times10^{-1}$ | $2.2\times10^{5}$ | $6.0\times10^{-2}$ | 3.0 | **$2.3\times10^{-3}$** | $9.9\times10^{-1}$ | N/A | $6.7\times10^{-1}$ | $1.3\times10^{2}$ | $8.8\times10^{-1}$ |
| V235A | 1.5 | $7.1\times10^{-2}$ | $2.7\times10^{5}$ | $9.7\times10^{-2}$ | 2.0 | $3.8\times10^{-2}$ | N/A | 1.0 | $7.3\times10^{-1}$ | $1.3\times10^{2}$ | $8.9\times10^{-1}$ |
| V235G | $4.4\times10^{-1}$ | **$4.0\times10^{-3}$** | $6.5\times10^{5}$ | $8.5\times10^{-1}$ | 1.7 | $1.1\times10^{-1}$ | $9.7\times10^{-1}$ | N/A | $4.5\times10^{-1}$ | $1.6\times10^{2}$ | $6.7\times10^{-1}$ |
| E236G | $7.7\times10^{-1}$ | $3.1\times10^{-1}$ | $5.2\times10^{5}$ | $5.3\times10^{-1}$ | 1.2 | $5.0\times10^{-1}$ | $8.3\times10^{-1}$ | N/A | **$8.9\times10^{-3}$** | $1.2\times10^{2}$ | $6.9\times10^{-1}$ |
| E236V | $8.5\times10^{-1}$ | $4.4\times10^{-1}$ | $6.8\times10^{5}$ | $6.6\times10^{-1}$ | $9.1\times10^{-1}$ | $7.0\times10^{-1}$ | N/A | 1.0 | $6.2\times10^{-1}$ | $1.8\times10^{2}$ | $3.7\times10^{-1}$ |
| W237G | $7.7\times10^{-1}$ | $2.4\times10^{-1}$ | $9.5\times10^{4}$ | **$1.5\times10^{-2}$** | 3.3 | **$6.1\times10^{-4}$** | $2.9\times10^{-1}$ | N/A | **0** | $4.4\times10^{1}$ | $5.5\times10^{-2}$ |
| W237V | $\leq 3.6\times10^{-1}$ | **0** | $\leq 7.5\times10^{4}$ | **$1.6\times10^{-2}$** | $\leq 3.4$ | **$5.4\times10^{-3}$** | $3.6\times10^{-1}$ | N/A | **0** | $\geq 3.1\times10^{1}$ | $5.3\times10^{-2}$ |
| E238G | $2.9\times10^{-1}$ | **0** | $1.7\times10^{5}$ | **$3.2\times10^{-2}$** | 1.9 | $1.7\times10^{-2}$ | $3.1\times10^{-1}$ | N/A | **0** | $4.5\times10^{1}$ | $5.8\times10^{-2}$ |
| E238V | $7.4\times10^{-1}$ | $3.2\times10^{-1}$ | $9.5\times10^{4}$ | $9.5\times10^{-2}$ | 2.4 | $2.9\times10^{-2}$ | $3.5\times10^{-1}$ | N/A | **$2.0\times10^{-4}$** | $5.5\times10^{1}$ | $1.7\times10^{-1}$ |
| G239A | $6.6\times10^{-1}$ | $9.4\times10^{-2}$ | $6.4\times10^{5}$ | $8.8\times10^{-1}$ | 1.7 | $7.2\times10^{-2}$ | $9.0\times10^{-1}$ | N/A | $1.4\times10^{-1}$ | $9.9\times10^{1}$ | $3.6\times10^{-1}$ |
| G239V | $7.8\times10^{-1}$ | $2.7\times10^{-1}$ | $4.3\times10^{5}$ | $2.7\times10^{-1}$ | 1.1 | $6.2\times10^{-1}$ | $9.8\times10^{-1}$ | N/A | $6.4\times10^{-1}$ | $1.4\times10^{2}$ | $9.3\times10^{-1}$ |
| L240G | 1.7 | $9.4\times10^{-2}$ | $4.5\times10^{5}$ | $4.6\times10^{-1}$ | $8.9\times10^{-1}$ | $6.8\times10^{-1}$ | $7.9\times10^{-1}$ | N/A | **$6.0\times10^{-3}$** | $7.9\times10^{1}$ | $3.0\times10^{-1}$ |
| L240V | $5.6\times10^{-1}$ | $3.1\times10^{-2}$ | $6.7\times10^{5}$ | $7.3\times10^{-1}$ | $9.6\times10^{-1}$ | $8.6\times10^{-1}$ | N/A | 1.1 | $1.3\times10^{-2}$ | $1.3\times10^{2}$ | $9.0\times10^{-1}$ |
| L241G | $3.8\times10^{-1}$ | **0** | $2.0\times10^{5}$ | **$4.0\times10^{-2}$** | 1.9 | $1.7\times10^{-2}$ | $3.4\times10^{-1}$ | N/A | **0** | $3.9\times10^{1}$ | **$4.8\times10^{-2}$** |
| L241V | 1.0 | $9.9\times10^{-1}$ | $3.5\times10^{5}$ | $1.4\times10^{-1}$ | 1.3 | $4.2\times10^{-1}$ | $9.5\times10^{-1}$ | N/A | $4.3\times10^{-1}$ | $1.1\times10^{2}$ | $4.7\times10^{-1}$ |
| G242A | $6.5\times10^{-1}$ | $1.3\times10^{-1}$ | $5.8\times10^{5}$ | $8.9\times10^{-1}$ | $8.9\times10^{-1}$ | $7.3\times10^{-1}$ | $6.9\times10^{-1}$ | N/A | **$6.0\times10^{-4}$** | $1.4\times10^{2}$ | $9.4\times10^{-1}$ |
| G242V | $7.3\times10^{-1}$ | $1.6\times10^{-1}$ | $2.6\times10^{5}$ | $7.2\times10^{-2}$ | 1.9 | $6.5\times10^{-2}$ | $7.9\times10^{-1}$ | N/A | **$6.0\times10^{-4}$** | $1.1\times10^{2}$ | $4.8\times10^{-1}$ |
| S243G | $9.2\times10^{-1}$ | $8.2\times10^{-1}$ | $4.3\times10^{5}$ | $3.2\times10^{-1}$ | 1.1 | $8.7\times10^{-1}$ | $8.6\times10^{-1}$ | N/A | **$8.2\times10^{-3}$** | $9.5\times10^{1}$ | $3.2\times10^{-1}$ |
| S243V | $7.8\times10^{-1}$ | $4.7\times10^{-1}$ | $4.4\times10^{5}$ | $5.0\times10^{-1}$ | $9.5\times10^{-1}$ | $8.8\times10^{-1}$ | $9.3\times10^{-1}$ | N/A | $1.5\times10^{-1}$ | $1.3\times10^{2}$ | $8.9\times10^{-1}$ |
| K244G | $5.5\times10^{-1}$ | $1.8\times10^{-2}$ | $9.4\times10^{5}$ | $1.8\times10^{-1}$ | 1.1 | $6.6\times10^{-1}$ | $8.9\times10^{-1}$ | N/A | $3.4\times10^{-2}$ | $1.4\times10^{2}$ | $9.6\times10^{-1}$ |
| K244V | $8.6\times10^{-1}$ | $5.2\times10^{-1}$ | $4.1\times10^{5}$ | $2.2\times10^{-1}$ | 1.2 | $5.3\times10^{-1}$ | $9.9\times10^{-1}$ | N/A | $6.5\times10^{-1}$ | $1.4\times10^{2}$ | $9.2\times10^{-1}$ |
| K245G | $9.3\times10^{-1}$ | $7.6\times10^{-1}$ | $4.3\times10^{5}$ | $3.6\times10^{-1}$ | 1.8 | $4.8\times10^{-2}$ | N/A | 1.0 | $7.7\times10^{-1}$ | $1.3\times10^{2}$ | $7.9\times10^{-1}$ |
| K245V | 1.9 | $9.4\times10^{-2}$ | $2.8\times10^{5}$ | $1.9\times10^{-1}$ | 1.5 | $2.1\times10^{-1}$ | $9.9\times10^{-1}$ | N/A | $6.5\times10^{-1}$ | $1.1\times10^{2}$ | $6.6\times10^{-1}$ |
| T246G | $3.3\times10^{-1}$ | $1.1\times10^{-2}$ | $6.3\times10^{5}$ | $9.6\times10^{-1}$ | – | – | $8.6\times10^{-1}$ | N/A | $1.1\times10^{-1}$ | $1.3\times10^{2}$ | $9.3\times10^{-1}$ |
| T246V | 1.1 | $6.0\times10^{-1}$ | $2.4\times10^{5}$ | $7.0\times10^{-2}$ | 2.0 | $4.0\times10^{-2}$ | N/A | 1.1 | $1.5\times10^{-1}$ | $1.6\times10^{2}$ | $6.6\times10^{-1}$ |
| P247G | $4.2\times10^{-1}$ | **$3.0\times10^{-3}$** | $5.1\times10^{5}$ | $6.1\times10^{-1}$ | 1.5 | $2.2\times10^{-1}$ | $9.0\times10^{-1}$ | N/A | $5.1\times10^{-2}$ | $1.3\times10^{2}$ | $8.6\times10^{-1}$ |
| P247V | $3.8\times10^{-1}$ | **0** | $7.6\times10^{5}$ | $4.0\times10^{-1}$ | $8.9\times10^{-1}$ | $7.8\times10^{-1}$ | $7.4\times10^{-1}$ | N/A | **$8.0\times10^{-4}$** | $1.1\times10^{2}$ | $5.6\times10^{-1}$ |
| T248G | $3.1\times10^{-1}$ | **0** | $8.6\times10^{5}$ | $2.5\times10^{-1}$ | $6.3\times10^{-1}$ | $1.6\times10^{-1}$ | $6.1\times10^{-1}$ | N/A | **0** | $8.6\times10^{1}$ | $3.1\times10^{-1}$ |
| T248V | $3.8\times10^{-1}$ | **$3.0\times10^{-3}$** | $1.7\times10^{6}$ | **$4.0\times10^{-3}$** | $5.4\times10^{-1}$ | $5.8\times10^{-2}$ | $8.9\times10^{-1}$ | N/A | $1.2\times10^{-1}$ | $1.7\times10^{2}$ | $5.1\times10^{-1}$ |
| F249G | $2.4\times10^{-1}$ | **0** | $1.3\times10^{5}$ | **$3.5\times10^{-2}$** | 2.6 | **$7.9\times10^{-3}$** | $2.1\times10^{-1}$ | N/A | **0** | $4.6\times10^{1}$ | $5.6\times10^{-2}$ |
| F249V | $3.3\times10^{-1}$ | **$1.0\times10^{-3}$** | $7.6\times10^{5}$ | $4.2\times10^{-1}$ | $9.4\times10^{-1}$ | $8.8\times10^{-1}$ | $8.4\times10^{-1}$ | N/A | **$3.9\times10^{-3}$** | $1.4\times10^{2}$ | $9.4\times10^{-1}$ |

| Variant | $f_a$ | $f_a$ $p$–value | $(k_{cat}/K_M)^{MeP}_{chem.}$ ($M^{-1}s^{-1}$) | $(k_{cat}/K_M)^{MeP}_{chem.}$ $p$–value | FC1 | FC1 $p$–value | FC2 | FC3 | FC2/3 $p$–value | FC4 ($s^{-1}$) | FC4 $p$–value |
|---|---|---|---|---|---|---|---|---|---|---|---|
| P250G | $5.4\times10^{-1}$ | $2.8\times10^{-2}$ | $7.1\times10^{4}$ | **$1.6\times10^{-2}$** | 5.4 | **$4.6\times10^{-4}$** | $8.4\times10^{-1}$ | N/A | $1.8\times10^{-2}$ | $7.6\times10^{1}$ | $2.1\times10^{-1}$ |
| P250V | $5.3\times10^{-1}$ | $2.2\times10^{-2}$ | $7.9\times10^{5}$ | $4.2\times10^{-1}$ | $6.8\times10^{-1}$ | $3.6\times10^{-1}$ | $9.2\times10^{-1}$ | N/A | $1.7\times10^{-1}$ | $1.3\times10^{2}$ | $8.9\times10^{-1}$ |
| Y251G | $2.0\times10^{-1}$ | **0** | $1.2\times10^{5}$ | **$1.9\times10^{-2}$** | 4.0 | **$1.1\times10^{-3}$** | $2.7\times10^{-1}$ | N/A | **0** | $4.4\times10^{1}$ | **$4.4\times10^{-2}$** |
| Y251V | $4.0\times10^{-1}$ | **$1.0\times10^{-3}$** | $6.5\times10^{5}$ | $8.4\times10^{-1}$ | $7.4\times10^{-1}$ | $3.9\times10^{-1}$ | $8.3\times10^{-1}$ | N/A | **$9.6\times10^{-3}$** | $1.2\times10^{2}$ | $7.5\times10^{-1}$ |
| T252G | 1.4 | $2.0\times10^{-1}$ | $3.9\times10^{5}$ | $2.4\times10^{-1}$ | $9.6\times10^{-1}$ | $8.7\times10^{-1}$ | $9.6\times10^{-1}$ | N/A | $4.9\times10^{-1}$ | $1.4\times10^{2}$ | 1.0 |
| T252V | 1.0 | $8.4\times10^{-1}$ | $4.2\times10^{5}$ | $3.9\times10^{-1}$ | $8.9\times10^{-1}$ | $7.6\times10^{-1}$ | N/A | 1.1 | $3.4\times10^{-1}$ | $1.7\times10^{2}$ | $5.2\times10^{-1}$ |
| D253G | $8.7\times10^{-1}$ | $4.9\times10^{-1}$ | $4.3\times10^{5}$ | $2.5\times10^{-1}$ | 1.7 | $7.4\times10^{-2}$ | $8.7\times10^{-1}$ | N/A | $1.0\times10^{-2}$ | $1.2\times10^{2}$ | $5.5\times10^{-1}$ |
| D253V | $9.2\times10^{-1}$ | $6.8\times10^{-1}$ | $5.1\times10^{5}$ | $6.2\times10^{-1}$ | $8.4\times10^{-1}$ | $5.1\times10^{-1}$ | $9.7\times10^{-1}$ | N/A | $5.3\times10^{-1}$ | $9.6\times10^{1}$ | $3.3\times10^{-1}$ |
| L254G | 1.0 | $9.8\times10^{-1}$ | $3.8\times10^{5}$ | $2.5\times10^{-1}$ | $8.6\times10^{-1}$ | $6.1\times10^{-1}$ | $7.0\times10^{-1}$ | N/A | **$1.\times10^{-4}$** | $5.7\times10^{1}$ | $1.6\times10^{-1}$ |
| L254V | 1.2 | $4.6\times10^{-1}$ | $7.4\times10^{5}$ | $3.8\times10^{-1}$ | $6.2\times10^{-1}$ | $2.3\times10^{-1}$ | $9.9\times10^{-1}$ | N/A | $7.7\times10^{-1}$ | $1.2\times10^{2}$ | $7.3\times10^{-1}$ |
| A255G | 1.0 | $8.2\times10^{-1}$ | $2.3\times10^{5}$ | $8.2\times10^{-2}$ | 3.3 | **$3.3\times10^{-3}$** | $8.5\times10^{-1}$ | N/A | **$5.6\times10^{-3}$** | $1.0\times10^{2}$ | $3.8\times10^{-1}$ |
| A255V | $9.0\times10^{-1}$ | $4.3\times10^{-1}$ | $4.9\times10^{5}$ | $3.9\times10^{-1}$ | 1.0 | $9.4\times10^{-1}$ | $8.9\times10^{-1}$ | N/A | $1.4\times10^{-2}$ | $1.3\times10^{2}$ | $7.6\times10^{-1}$ |
| K256G | 1.1 | $6.3\times10^{-1}$ | $3.9\times10^{5}$ | $3.4\times10^{-1}$ | 1.6 | $1.1\times10^{-1}$ | $8.7\times10^{-1}$ | N/A | $1.7\times10^{-2}$ | $1.1\times10^{2}$ | $4.8\times10^{-1}$ |
| K256V | $3.0\times10^{-1}$ | **$1.0\times10^{-3}$** | $3.7\times10^{5}$ | $2.1\times10^{-1}$ | – | – | N/A | 1.1 | $4.7\times10^{-1}$ | $1.5\times10^{2}$ | $8.3\times10^{-1}$ |
| D257G | 1.6 | $1.3\times10^{-1}$ | $2.1\times10^{5}$ | $1.2\times10^{-1}$ | 2.1 | $2.4\times10^{-2}$ | $6.8\times10^{-1}$ | N/A | **$1.9\times10^{-3}$** | $4.1\times10^{1}$ | $1.3\times10^{-1}$ |
| D257V | 1.2 | $3.6\times10^{-1}$ | $1.2\times10^{5}$ | **$4.9\times10^{-2}$** | 3.5 | **$2.7\times10^{-3}$** | $8.3\times10^{-1}$ | N/A | **$7.3\times10^{-3}$** | $1.1\times10^{2}$ | $5.2\times10^{-1}$ |
| Y258G | 1.3 | $2.7\times10^{-1}$ | $2.9\times10^{5}$ | $1.2\times10^{-1}$ | 1.3 | $3.1\times10^{-1}$ | $8.1\times10^{-1}$ | N/A | **$6.0\times10^{-3}$** | $1.1\times10^{2}$ | $5.8\times10^{-1}$ |
| Y258V | 1.3 | $1.8\times10^{-1}$ | $6.4\times10^{5}$ | $8.3\times10^{-1}$ | $8.0\times10^{-1}$ | $6.3\times10^{-1}$ | $8.4\times10^{-1}$ | N/A | $1.1\times10^{-2}$ | $1.3\times10^{2}$ | $8.8\times10^{-1}$ |
| E259G | 1.3 | $2.8\times10^{-1}$ | $1.7\times10^{5}$ | **$4.3\times10^{-2}$** | 3.8 | **$2.3\times10^{-4}$** | $7.2\times10^{-1}$ | N/A | **0** | $1.2\times10^{2}$ | $5.7\times10^{-1}$ |
| E259V | 1.2 | $5.6\times10^{-1}$ | $4.3\times10^{5}$ | $3.4\times10^{-1}$ | 1.4 | $3.4\times10^{-1}$ | $9.4\times10^{-1}$ | N/A | $2.7\times10^{-1}$ | $1.7\times10^{2}$ | $5.0\times10^{-1}$ |
| A260G | 1.4 | $3.0\times10^{-1}$ | $3.0\times10^{5}$ | $1.5\times10^{-1}$ | 1.9 | $2.2\times10^{-2}$ | $9.4\times10^{-1}$ | N/A | $1.8\times10^{-1}$ | $6.5\times10^{1}$ | $2.1\times10^{-1}$ |
| A260V | 1.4 | $9.3\times10^{-2}$ | $3.3\times10^{5}$ | $9.9\times10^{-2}$ | 1.1 | $6.8\times10^{-1}$ | N/A | 1.1 | $9.2\times10^{-2}$ | $1.4\times10^{2}$ | 1.0 |
| K261G | $6.2\times10^{-1}$ | $1.0\times10^{-1}$ | $1.0\times10^{6}$ | $1.1\times10^{-1}$ | 1.1 | $8.9\times10^{-1}$ | N/A | 1.0 | $5.1\times10^{-1}$ | $1.3\times10^{2}$ | $8.1\times10^{-1}$ |
| K261V | $9.6\times10^{-1}$ | $8.5\times10^{-1}$ | $4.5\times10^{5}$ | $4.2\times10^{-1}$ | 1.0 | $9.9\times10^{-1}$ | N/A | 1.0 | $7.9\times10^{-1}$ | $1.2\times10^{2}$ | $7.1\times10^{-1}$ |
| K262G | 1.1 | $7.0\times10^{-1}$ | $4.1\times10^{5}$ | $2.8\times10^{-1}$ | 2.1 | $1.4\times10^{-2}$ | $7.5\times10^{-1}$ | N/A | **0** | $1.1\times10^{2}$ | $4.4\times10^{-1}$ |
| K262V | $9.2\times10^{-1}$ | $6.6\times10^{-1}$ | $4.7\times10^{5}$ | $3.9\times10^{-1}$ | 1.6 | $1.1\times10^{-1}$ | $8.7\times10^{-1}$ | N/A | $1.6\times10^{-2}$ | $1.3\times10^{2}$ | $8.3\times10^{-1}$ |
| G263A | $7.6\times10^{-1}$ | $2.2\times10^{-1}$ | $5.2\times10^{5}$ | $6.6\times10^{-1}$ | 1.9 | $5.9\times10^{-2}$ | $9.2\times10^{-1}$ | N/A | $7.9\times10^{-2}$ | $1.3\times10^{2}$ | $8.1\times10^{-1}$ |
| G263V | $7.5\times10^{-1}$ | $1.2\times10^{-1}$ | $8.3\times10^{5}$ | $1.5\times10^{-1}$ | $9.7\times10^{-1}$ | $9.5\times10^{-1}$ | $8.7\times10^{-1}$ | N/A | $2.4\times10^{-2}$ | $1.6\times10^{2}$ | $7.3\times10^{-1}$ |
| L264G | $7.2\times10^{-1}$ | $1.3\times10^{-1}$ | $4.6\times10^{5}$ | $3.9\times10^{-1}$ | 1.2 | $5.0\times10^{-1}$ | $6.5\times10^{-1}$ | N/A | **0** | $1.1\times10^{2}$ | $4.6\times10^{-1}$ |
| L264V | $4.1\times10^{-1}$ | $1.0\times10^{-2}$ | $1.3\times10^{6}$ | **$2.7\times10^{-2}$** | – | – | N/A | 1.1 | $5.1\times10^{-2}$ | $1.8\times10^{2}$ | $5.7\times10^{-1}$ |
| I265G | $5.3\times10^{-1}$ | **$3.0\times10^{-3}$** | $1.4\times10^{5}$ | **$3.6\times10^{-2}$** | 3.5 | **$1.8\times10^{-3}$** | $4.9\times10^{-1}$ | N/A | **0** | $6.3\times10^{1}$ | $1.1\times10^{-1}$ |
| I265V | $6.6\times10^{-1}$ | $1.9\times10^{-1}$ | $1.1\times10^{6}$ | $1.2\times10^{-1}$ | $4.3\times10^{-1}$ | $1.2\times10^{-2}$ | $9.0\times10^{-1}$ | N/A | $5.1\times10^{-2}$ | $1.0\times10^{2}$ | $4.8\times10^{-1}$ |
| R266G | ≤ 1.6 | $3.3\times10^{-2}$ | ≤ $8.4\times10^{4}$ | **$2.9\times10^{-2}$** | ≤ 7.7 | **0** | N/A | 1.1 | $2.6\times10^{-1}$ | ≥ $8.4\times10^{1}$ | $2.2\times10^{-1}$ |
| R266V | 1.2 | $4.3\times10^{-1}$ | $7.6\times10^{4}$ | **$7.0\times10^{-3}$** | $1.8\times10^{1}$ | **0** | N/A | 2.8 | **0** | $2.1\times10^{2}$ | $1.2\times10^{-1}$ |
| T267G | 1.9 | $2.3\times10^{-2}$ | $3.0\times10^{5}$ | $1.6\times10^{-1}$ | 1.6 | $1.3\times10^{-1}$ | $7.3\times10^{-1}$ | N/A | **$3.0\times10^{-4}$** | $6.1\times10^{1}$ | $1.3\times10^{-1}$ |
| T267V | 1.5 | $6.6\times10^{-2}$ | $3.1\times10^{5}$ | $7.1\times10^{-2}$ | $9.6\times10^{-1}$ | $9.3\times10^{-1}$ | $9.8\times10^{-1}$ | N/A | $6.4\times10^{-1}$ | $1.4\times10^{2}$ | $9.3\times10^{-1}$ |
| T268V | 1.5 | $6.1\times10^{-2}$ | $3.1\times10^{5}$ | $5.6\times10^{-2}$ | 1.2 | $4.6\times10^{-1}$ | $8.8\times10^{-1}$ | N/A | **$6.0\times10^{-3}$** | $1.1\times10^{2}$ | $4.4\times10^{-1}$ |
| P269G | 1.0 | $9.6\times10^{-1}$ | $2.2\times10^{5}$ | $7.7\times10^{-2}$ | 1.1 | $7.5\times10^{-1}$ | N/A | 1.2 | **$4.0\times10^{-3}$** | $1.3\times10^{2}$ | $8.3\times10^{-1}$ |
| P269V | 1.6 | $4.3\times10^{-2}$ | $1.5\times10^{5}$ | **$4.2\times10^{-2}$** | 1.5 | $1.1\times10^{-1}$ | $9.3\times10^{-1}$ | N/A | $1.2\times10^{-1}$ | $1.2\times10^{2}$ | $6.6\times10^{-1}$ |
| F270G | 1.0 | $9.4\times10^{-1}$ | $3.6\times10^{5}$ | $2.1\times10^{-1}$ | 1.4 | $1.1\times10^{-1}$ | $7.8\times10^{-1}$ | N/A | **$2.0\times10^{-4}$** | $5.6\times10^{1}$ | $1.8\times10^{-1}$ |
| F270V | 1.7 | $1.5\times10^{-2}$ | $1.8\times10^{5}$ | **$2.4\times10^{-2}$** | 1.8 | $5.8\times10^{-2}$ | $9.8\times10^{-1}$ | N/A | $6.5\times10^{-1}$ | $1.5\times10^{2}$ | $9.3\times10^{-1}$ |
| G271A | 1.4 | $1.9\times10^{-1}$ | $2.1\times10^{5}$ | $6.7\times10^{-2}$ | 1.2 | $5.9\times10^{-1}$ | $8.1\times10^{-1}$ | N/A | **$5.0\times10^{-4}$** | $6.7\times10^{1}$ | $1.3\times10^{-1}$ |
| G271V | $8.6\times10^{-1}$ | $5.4\times10^{-1}$ | $1.1\times10^{5}$ | **$2.5\times10^{-2}$** | $1.0\times10^{1}$ | **$1.0\times10^{-5}$** | $6.7\times10^{-1}$ | N/A | **$5.0\times10^{-4}$** | $4.8\times10^{1}$ | $6.1\times10^{-2}$ |
| N272G | $7.8\times10^{-1}$ | $3.3\times10^{-1}$ | $1.4\times10^{5}$ | **$4.6\times10^{-2}$** | 2.8 | **$4.5\times10^{-3}$** | N/A | 1.1 | $1.5\times10^{-1}$ | $1.1\times10^{2}$ | $6.0\times10^{-1}$ |
| N272V | $3.2\times10^{-1}$ | **$3.0\times10^{-3}$** | $6.4\times10^{5}$ | $9.0\times10^{-1}$ | – | – | $7.4\times10^{-1}$ | N/A | **0** | $1.6\times10^{2}$ | $7.8\times10^{-1}$ |
| T273G | $7.7\times10^{-1}$ | $2.8\times10^{-1}$ | $2.1\times10^{5}$ | $9.1\times10^{-2}$ | 3.1 | $1.0\times10^{-2}$ | $8.6\times10^{-1}$ | N/A | **$3.0\times10^{-3}$** | $7.9\times10^{1}$ | $1.9\times10^{-1}$ |
| T273V | 1.8 | **$6.0\times10^{-3}$** | $9.0\times10^{4}$ | **$1.5\times10^{-2}$** | 3.5 | **$4.3\times10^{-4}$** | N/A | 1.2 | **$2.4\times10^{-3}$** | $1.5\times10^{2}$ | $8.6\times10^{-1}$ |
| L274G | $1.2\times10^{-1}$ | **0** | $1.8\times10^{5}$ | $1.0\times10^{-1}$ | – | – | $9.7\times10^{-1}$ | N/A | $7.5\times10^{-1}$ | $1.1\times10^{2}$ | $6.6\times10^{-1}$ |
| L274V | 1.0 | 1.0 | $4.7\times10^{5}$ | $4.4\times10^{-1}$ | $9.1\times10^{-1}$ | $7.4\times10^{-1}$ | N/A | 1.1 | $3.1\times10^{-1}$ | $2.0\times10^{2}$ | $2.7\times10^{-1}$ |
| T275G | $5.9\times10^{-1}$ | $2.4\times10^{-2}$ | $1.7\times10^{5}$ | **$3.3\times10^{-2}$** | 2.5 | **$2.3\times10^{-3}$** | N/A | 1.0 | $7.5\times10^{-1}$ | $1.0\times10^{2}$ | $3.6\times10^{-1}$ |
| T275V | 1.3 | $3.4\times10^{-1}$ | $2.3\times10^{5}$ | $6.4\times10^{-2}$ | 2.2 | $2.3\times10^{-2}$ | 1.0 | N/A | $8.7\times10^{-1}$ | $1.1\times10^{2}$ | $5.2\times10^{-1}$ |
| L276G | $2.3\times10^{-2}$ | **0** | $2.6\times10^{5}$ | $1.3\times10^{-1}$ | – | – | $6.2\times10^{-1}$ | N/A | $2.2\times10^{-2}$ | $1.5\times10^{2}$ | $9.2\times10^{-1}$ |
| L276V | $9.9\times10^{-1}$ | $9.3\times10^{-1}$ | $2.7\times10^{5}$ | $8.0\times10^{-2}$ | 1.7 | $1.0\times10^{-1}$ | $9.9\times10^{-1}$ | N/A | $8.1\times10^{-1}$ | $1.6\times10^{2}$ | $5.5\times10^{-1}$ |
| Q277G | 1.2 | $5.0\times10^{-1}$ | $1.4\times10^{5}$ | $5.4\times10^{-2}$ | 3.6 | **$7.1\times10^{-4}$** | $9.9\times10^{-1}$ | N/A | $6.0\times10^{-1}$ | $1.5\times10^{2}$ | $8.9\times10^{-1}$ |
| Q277V | $9.7\times10^{-1}$ | $8.2\times10^{-1}$ | $3.6\times10^{5}$ | $1.9\times10^{-1}$ | 1.2 | $6.6\times10^{-1}$ | N/A | 1.0 | $4.3\times10^{-1}$ | $1.7\times10^{2}$ | $5.5\times10^{-1}$ |
| M278G | $2.3\times10^{-1}$ | **0** | $2.8\times10^{5}$ | $1.6\times10^{-1}$ | – | – | $7.9\times10^{-1}$ | N/A | **$2.1\times10^{-3}$** | $7.4\times10^{1}$ | $2.8\times10^{-1}$ |
| M278V | $8.4\times10^{-1}$ | $3.0\times10^{-1}$ | $4.0\times10^{5}$ | $1.6\times10^{-1}$ | 1.2 | $5.9\times10^{-1}$ | N/A | 1.1 | $1.4\times10^{-1}$ | $1.4\times10^{2}$ | $9.6\times10^{-1}$ |
| A279G | $6.4\times10^{-1}$ | $1.1\times10^{-1}$ | $2.6\times10^{5}$ | $1.4\times10^{-1}$ | 1.9 | $4.0\times10^{-2}$ | $7.7\times10^{-1}$ | N/A | **$1.1\times10^{-3}$** | $1.2\times10^{2}$ | $7.3\times10^{-1}$ |
| A279V | $9.8\times10^{-1}$ | $9.1\times10^{-1}$ | $2.9\times10^{5}$ | $9.6\times10^{-2}$ | 1.4 | $3.0\times10^{-1}$ | $9.9\times10^{-1}$ | N/A | $8.8\times10^{-1}$ | $1.5\times10^{2}$ | $7.5\times10^{-1}$ |
| D280G | 1.3 | $4.1\times10^{-1}$ | $4.6\times10^{5}$ | $5.1\times10^{-1}$ | 1.1 | $6.8\times10^{-1}$ | N/A | 1.2 | $4.2\times10^{-2}$ | $1.0\times10^{2}$ | $4.7\times10^{-1}$ |
| D280V | $4.8\times10^{-1}$ | **$4.0\times10^{-3}$** | $8.6\times10^{5}$ | $1.4\times10^{-1}$ | $6.3\times10^{-1}$ | $1.3\times10^{-1}$ | $9.3\times10^{-1}$ | N/A | $1.1\times10^{-1}$ | $1.6\times10^{2}$ | $7.4\times10^{-1}$ |
| A281G | 1.2 | $5.2\times10^{-1}$ | $1.9\times10^{5}$ | $6.2\times10^{-2}$ | 1.6 | $1.9\times10^{-1}$ | N/A | 1.0 | $4.8\times10^{-1}$ | $1.7\times10^{2}$ | $6.5\times10^{-1}$ |
| A281V | $6.5\times10^{-1}$ | $1.3\times10^{-1}$ | $3.2\times10^{5}$ | $1.7\times10^{-1}$ | 1.9 | $5.9\times10^{-2}$ | $9.9\times10^{-1}$ | N/A | $8.2\times10^{-1}$ | $1.4\times10^{2}$ | $9.5\times10^{-1}$ |
| A282G | $6.0\times10^{-1}$ | $1.2\times10^{-1}$ | $2.7\times10^{5}$ | $2.1\times10^{-1}$ | 1.8 | $1.0\times10^{-1}$ | $8.8\times10^{-1}$ | N/A | $6.8\times10^{-2}$ | $8.3\times10^{1}$ | $4.4\times10^{-1}$ |
| A282V | $6.1\times10^{-1}$ | $2.4\times10^{-2}$ | $7.0\times10^{5}$ | $5.4\times10^{-1}$ | $7.5\times10^{-1}$ | $3.6\times10^{-1}$ | $8.7\times10^{-1}$ | N/A | **$1.7\times10^{-3}$** | $1.4\times10^{2}$ | $9.4\times10^{-1}$ |
| I283G | $3.5\times10^{-2}$ | **0** | $1.6\times10^{5}$ | $8.0\times10^{-2}$ | – | – | $6.4\times10^{-1}$ | N/A | **0** | $1.2\times10^{2}$ | $6.9\times10^{-1}$ |
| I283V | 1.0 | $9.4\times10^{-1}$ | $5.4\times10^{5}$ | $7.4\times10^{-1}$ | 1.0 | $9.5\times10^{-1}$ | $9.5\times10^{-1}$ | N/A | $2.3\times10^{-1}$ | $1.7\times10^{2}$ | $5.6\times10^{-1}$ |
| D284G | 1.2 | $3.4\times10^{-1}$ | $2.5\times10^{5}$ | **$2.9\times10^{-2}$** | 2.3 | **$4.2\times10^{-4}$** | $9.5\times10^{-1}$ | N/A | $1.6\times10^{-1}$ | $8.5\times10^{1}$ | $9.9\times10^{-2}$ |
| D284V | 1.4 | $1.0\times10^{-1}$ | $3.1\times10^{5}$ | $8.1\times10^{-2}$ | 1.4 | $2.4\times10^{-1}$ | N/A | 1.1 | $1.1\times10^{-1}$ | $1.7\times10^{2}$ | $4.4\times10^{-1}$ |
| G285A | 1.7 | $3.5\times10^{-2}$ | $3.4\times10^{5}$ | $1.5\times10^{-1}$ | 1.1 | $8.1\times10^{-1}$ | N/A | 1.0 | $6.4\times10^{-1}$ | $1.1\times10^{2}$ | $5.2\times10^{-1}$ |
| G285V | $4.5\times10^{-1}$ | **0** | $2.5\times10^{5}$ | $5.3\times10^{-2}$ | 1.7 | $9.2\times10^{-2}$ | 1.0 | N/A | $7.3\times10^{-1}$ | $1.6\times10^{2}$ | $7.2\times10^{-1}$ |
| N286G | 1.4 | $2.0\times10^{-1}$ | $2.1\times10^{5}$ | $6.6\times10^{-2}$ | 1.7 | $1.0\times10^{-1}$ | $9.7\times10^{-1}$ | N/A | $5.2\times10^{-1}$ | $1.2\times10^{2}$ | $7.5\times10^{-1}$ |
| N286V | 1.8 | $2.0\times10^{-2}$ | $2.4\times10^{5}$ | $9.5\times10^{-2}$ | 1.3 | $4.1\times10^{-1}$ | N/A | 1.1 | $1.5\times10^{-1}$ | $1.2\times10^{2}$ | $6.4\times10^{-1}$ |
| Q287G | 1.0 | $8.2\times10^{-1}$ | $4.7\times10^{5}$ | $4.5\times10^{-1}$ | 1.4 | $2.1\times10^{-1}$ | $9.4\times10^{-1}$ | N/A | $1.7\times10^{-1}$ | $1.0\times10^{2}$ | $4.1\times10^{-1}$ |
| Q287V | 1.2 | $3.6\times10^{-1}$ | $2.7\times10^{5}$ | $7.4\times10^{-2}$ | 1.6 | $9.6\times10^{-2}$ | N/A | 1.1 | $3.9\times10^{-2}$ | $1.9\times10^{2}$ | $2.4\times10^{-1}$ |

| Variant | $f_a$ | $f_a$ $p$–value | $(k_{cat}/K_M)^{MeP}_{chem.}$ ($M^{-1}s^{-1}$) | $(k_{cat}/K_M)^{MeP}_{chem.}$ $p$–value | FC1 | FC1 $p$–value | FC2 | FC3 | FC2/3 $p$–value | FC4 ($s^{-1}$) | FC4 $p$–value |
|---|---|---|---|---|---|---|---|---|---|---|---|
| M288G | $1.4\times10^{-1}$ | **0** | $1.6\times10^{5}$ | **$4.1\times10^{-2}$** | – | – | $9.9\times10^{-1}$ | N/A | $4.9\times10^{-1}$ | $1.0\times10^{2}$ | $4.6\times10^{-1}$ |
| M288V | – | – | – | – | – | – | $9.3\times10^{-1}$ | N/A | $3.4\times10^{-1}$ | – | – |
| G289A | $8.0\times10^{-1}$ | $5.4\times10^{-1}$ | $2.2\times10^{5}$ | $1.2\times10^{-1}$ | 2.5 | **$4.7\times10^{-3}$** | $9.7\times10^{-1}$ | N/A | $4.7\times10^{-1}$ | $8.9\times10^{1}$ | $3.4\times10^{-1}$ |
| V290A | 1.1 | $5.3\times10^{-1}$ | $4.9\times10^{5}$ | $3.0\times10^{-1}$ | 1.1 | $6.7\times10^{-1}$ | N/A | 1.1 | $7.5\times10^{-2}$ | $1.5\times10^{2}$ | $8.8\times10^{-1}$ |
| V290G | 1.1 | $7.4\times10^{-1}$ | $1.5\times10^{6}$ | **$1.4\times10^{-2}$** | $3.1\times10^{-1}$ | **$6.5\times10^{-4}$** | N/A | 1.0 | $6.7\times10^{-1}$ | $1.1\times10^{2}$ | $5.9\times10^{-1}$ |
| D291G | 1.3 | $3.3\times10^{-1}$ | $3.1\times10^{5}$ | $3.1\times10^{-1}$ | 1.9 | $6.3\times10^{-2}$ | $9.0\times10^{-1}$ | N/A | $2.6\times10^{-1}$ | $1.2\times10^{2}$ | $6.6\times10^{-1}$ |
| D291V | 1.1 | $7.3\times10^{-1}$ | $6.0\times10^{4}$ | **$3.4\times10^{-2}$** | 4.5 | **$4.7\times10^{-3}$** | $9.4\times10^{-1}$ | N/A | $3.4\times10^{-1}$ | $1.4\times10^{2}$ | 1.0 |
| D292G | 1.3 | $2.1\times10^{-1}$ | $6.1\times10^{5}$ | $9.8\times10^{-1}$ | $8.9\times10^{-1}$ | $7.2\times10^{-1}$ | N/A | 1.0 | $7.3\times10^{-1}$ | $1.2\times10^{2}$ | $5.8\times10^{-1}$ |
| D292V | $8.9\times10^{-1}$ | $6.8\times10^{-1}$ | $3.5\times10^{5}$ | $1.9\times10^{-1}$ | – | – | N/A | 1.1 | $2.1\times10^{-1}$ | $1.3\times10^{2}$ | $9.2\times10^{-1}$ |
| I293G | 1.2 | $5.2\times10^{-1}$ | $3.8\times10^{5}$ | $1.7\times10^{-1}$ | 1.6 | $1.1\times10^{-1}$ | $9.9\times10^{-1}$ | N/A | $7.9\times10^{-1}$ | $1.1\times10^{2}$ | $5.1\times10^{-1}$ |
| I293V | $6.3\times10^{-1}$ | $5.6\times10^{-2}$ | $9.5\times10^{5}$ | $7.3\times10^{-2}$ | $6.1\times10^{-1}$ | $7.7\times10^{-2}$ | N/A | 1.0 | $8.9\times10^{-1}$ | $1.6\times10^{2}$ | $7.1\times10^{-1}$ |
| T294G | $4.6\times10^{-1}$ | $4.7\times10^{-2}$ | $2.6\times10^{5}$ | $2.1\times10^{-1}$ | 2.0 | $6.7\times10^{-2}$ | $8.6\times10^{-1}$ | N/A | $2.4\times10^{-2}$ | $9.3\times10^{1}$ | $5.3\times10^{-1}$ |
| T294V | 1.3 | $2.4\times10^{-1}$ | $2.9\times10^{5}$ | $1.1\times10^{-1}$ | 1.9 | $5.5\times10^{-2}$ | N/A | 1.1 | $2.9\times10^{-2}$ | $1.6\times10^{2}$ | $5.8\times10^{-1}$ |
| D295G | $4.7\times10^{-1}$ | $2.2\times10^{-2}$ | $3.6\times10^{5}$ | $3.1\times10^{-1}$ | 1.4 | $3.4\times10^{-1}$ | $8.0\times10^{-1}$ | N/A | **$5.3\times10^{-3}$** | $8.3\times10^{1}$ | $3.4\times10^{-1}$ |
| D295V | $1.2\times10^{-1}$ | **0** | $1.5\times10^{5}$ | **$3.3\times10^{-2}$** | – | – | $8.1\times10^{-1}$ | N/A | **$1.\times10^{-4}$** | $1.1\times10^{2}$ | $5.0\times10^{-1}$ |
| F296G | $1.1\times10^{-1}$ | **0** | $1.6\times10^{5}$ | **$4.4\times10^{-2}$** | 3.1 | $2.9\times10^{-2}$ | $7.0\times10^{-1}$ | N/A | **0** | $1.0\times10^{2}$ | $4.2\times10^{-1}$ |
| F296V | $6.9\times10^{-1}$ | $2.1\times10^{-1}$ | $6.3\times10^{5}$ | $9.4\times10^{-1}$ | – | – | $7.8\times10^{-1}$ | N/A | **$9.0\times10^{-4}$** | $1.1\times10^{2}$ | $6.3\times10^{-1}$ |
| L297G | $\leq 5.4\times10^{-3}$ | **0** | $\leq 4.0\times10^{5}$ | $2.9\times10^{-1}$ | – | – | $7.4\times10^{-1}$ | N/A | $2.8\times10^{-2}$ | $\geq 8.5\times10^{1}$ | $3.8\times10^{-1}$ |
| L297V | 1.2 | $4.9\times10^{-1}$ | $3.0\times10^{5}$ | $6.7\times10^{-2}$ | 1.5 | $2.1\times10^{-1}$ | N/A | 1.0 | $7.2\times10^{-1}$ | $1.8\times10^{2}$ | $4.4\times10^{-1}$ |
| T298G | $8.3\times10^{-1}$ | $4.0\times10^{-1}$ | $3.9\times10^{5}$ | $3.1\times10^{-1}$ | 1.4 | $3.0\times10^{-1}$ | $8.0\times10^{-1}$ | N/A | **$6.0\times10^{-4}$** | $1.1\times10^{2}$ | $4.8\times10^{-1}$ |
| T298V | 1.8 | $1.3\times10^{-2}$ | $5.3\times10^{5}$ | $6.7\times10^{-1}$ | $7.3\times10^{-1}$ | $3.7\times10^{-1}$ | $9.4\times10^{-1}$ | N/A | $2.8\times10^{-1}$ | $1.2\times10^{2}$ | $6.5\times10^{-1}$ |
| V299A | $7.2\times10^{-1}$ | $2.2\times10^{-1}$ | $1.4\times10^{5}$ | $6.3\times10^{-2}$ | 4.0 | **$6.0\times10^{-3}$** | $8.2\times10^{-1}$ | N/A | $4.6\times10^{-2}$ | $1.4\times10^{2}$ | $9.3\times10^{-1}$ |
| V299G | $9.7\times10^{-3}$ | **0** | $1.3\times10^{6}$ | **$4.5\times10^{-2}$** | – | – | $6.3\times10^{-1}$ | N/A | **0** | $1.2\times10^{2}$ | $6.4\times10^{-1}$ |
| N300G | 2.5 | **$3.0\times10^{-3}$** | $2.6\times10^{5}$ | $1.5\times10^{-1}$ | $3.9\times10^{-1}$ | $1.1\times10^{-2}$ | $4.1\times10^{-1}$ | N/A | **$2.0\times10^{-4}$** | $6.4\times10^{1}$ | $1.3\times10^{-1}$ |
| N300V | $9.8\times10^{-2}$ | **0** | $2.8\times10^{4}$ | **$4.5\times10^{-2}$** | $1.9\times10^{1}$ | **$4.0\times10^{-4}$** | $8.3\times10^{-1}$ | N/A | **$3.6\times10^{-3}$** | $3.0\times10^{1}$ | $5.7\times10^{-2}$ |
| L301G | $1.1\times10^{-1}$ | **0** | $1.4\times10^{5}$ | **$2.8\times10^{-2}$** | – | – | $9.8\times10^{-1}$ | N/A | $7.2\times10^{-1}$ | $9.8\times10^{1}$ | $3.4\times10^{-1}$ |
| L301V | $4.3\times10^{-1}$ | $1.4\times10^{-2}$ | $7.6\times10^{5}$ | $5.1\times10^{-1}$ | $6.4\times10^{-1}$ | $3.2\times10^{-1}$ | N/A | 1.3 | **$3.2\times10^{-3}$** | $2.2\times10^{2}$ | $1.7\times10^{-1}$ |
| A302G | 1.2 | $5.4\times10^{-1}$ | $9.1\times10^{4}$ | $6.9\times10^{-2}$ | 3.9 | $1.1\times10^{-2}$ | N/A | 1.6 | **$4.0\times10^{-4}$** | $7.2\times10^{1}$ | $2.4\times10^{-1}$ |
| A302V | $2.2\times10^{-1}$ | **0** | $2.9\times10^{4}$ | **$2.9\times10^{-2}$** | $2.1\times10^{2}$ | **0** | N/A | 1.3 | **0** | $1.2\times10^{2}$ | $7.0\times10^{-1}$ |
| S303G | 1.2 | $5.8\times10^{-1}$ | $2.9\times10^{5}$ | $1.1\times10^{-1}$ | 1.7 | $1.1\times10^{-1}$ | N/A | 1.2 | $3.2\times10^{-2}$ | $1.6\times10^{2}$ | $8.2\times10^{-1}$ |
| S303V | $6.4\times10^{-1}$ | $4.6\times10^{-2}$ | $1.3\times10^{5}$ | **$3.4\times10^{-2}$** | 1.3 | $3.4\times10^{-1}$ | N/A | 1.1 | $8.2\times10^{-2}$ | $8.9\times10^{1}$ | $2.1\times10^{-1}$ |
| T304G | $\leq 2.0$ | $1.3\times10^{-2}$ | $\leq 1.6\times10^{5}$ | $5.2\times10^{-2}$ | $\leq 2.2$ | $1.1\times10^{-2}$ | $7.0\times10^{-1}$ | N/A | **$4.0\times10^{-4}$** | $\geq 8.9\times10^{1}$ | $2.9\times10^{-1}$ |
| T304V | $8.4\times10^{-2}$ | **0** | $1.2\times10^{6}$ | **$2.3\times10^{-2}$** | – | – | N/A | 1.2 | **$1.\times10^{-4}$** | $1.4\times10^{2}$ | 1.0 |
| D305G | $\leq 7.3\times10^{-2}$ | **0** | $\leq 4.0\times10^{4}$ | **$2.2\times10^{-2}$** | – | – | $\leq 3.3\times10^{-3}$ | N/A | **$4.0\times10^{-4}$** | – | – |
| D305V | – | – | – | – | – | – | – | – | – | – | – |
| Y306G | 4.1 | **0** | $1.9\times10^{4}$ | **$1.2\times10^{-2}$** | $4.5\times10^{1}$ | **0** | $2.3\times10^{-1}$ | N/A | **0** | $1.2\times10^{1}$ | $5.5\times10^{-2}$ |
| Y306V | $6.2\times10^{-1}$ | $5.0\times10^{-2}$ | $9.1\times10^{4}$ | **$8.0\times10^{-3}$** | $1.1\times10^{1}$ | **0** | $8.4\times10^{-1}$ | N/A | **$6.6\times10^{-3}$** | $1.1\times10^{2}$ | $4.7\times10^{-1}$ |
| V307A | 1.5 | $5.4\times10^{-2}$ | $1.8\times10^{5}$ | **$3.1\times10^{-2}$** | 2.1 | $1.9\times10^{-2}$ | N/A | 1.4 | **$2.0\times10^{-4}$** | $1.6\times10^{2}$ | $7.1\times10^{-1}$ |
| V307G | 1.4 | $2.0\times10^{-1}$ | $1.0\times10^{5}$ | $6.4\times10^{-2}$ | 2.4 | $2.5\times10^{-2}$ | N/A | 1.4 | **$8.0\times10^{-4}$** | $8.3\times10^{1}$ | $2.6\times10^{-1}$ |
| G308A | $4.5\times10^{-1}$ | **$8.0\times10^{-3}$** | $6.3\times10^{5}$ | $9.2\times10^{-1}$ | 1.7 | $8.6\times10^{-2}$ | $8.4\times10^{-1}$ | N/A | **$1.2\times10^{-3}$** | $1.2\times10^{2}$ | $7.3\times10^{-1}$ |
| G308V | $1.6\times10^{-1}$ | **0** | $4.5\times10^{5}$ | $3.6\times10^{-1}$ | – | – | $7.7\times10^{-1}$ | N/A | **0** | $1.1\times10^{2}$ | $5.0\times10^{-1}$ |
| H309G | $2.2\times10^{-1}$ | **0** | $4.6\times10^{5}$ | $4.9\times10^{-1}$ | – | – | N/A | 1.3 | **0** | $2.3\times10^{2}$ | $6.3\times10^{-2}$ |
| H309V | $\leq 3.7\times10^{-3}$ | **0** | $\leq 2.8\times10^{5}$ | $7.1\times10^{-2}$ | – | – | N/A | 1.6 | **0** | $\geq 2.0\times10^{3}$ | **0** |
| N310G | $9.1\times10^{-1}$ | $7.1\times10^{-1}$ | $2.7\times10^{5}$ | $1.1\times10^{-1}$ | 2.6 | **$3.3\times10^{-3}$** | $9.7\times10^{-1}$ | N/A | $5.0\times10^{-1}$ | $1.2\times10^{2}$ | $7.6\times10^{-1}$ |
| N310V | $9.9\times10^{-1}$ | $8.9\times10^{-1}$ | $4.4\times10^{5}$ | $3.0\times10^{-1}$ | $8.5\times10^{-1}$ | $6.7\times10^{-1}$ | $7.7\times10^{-1}$ | N/A | **$2.0\times10^{-4}$** | $9.2\times10^{1}$ | $2.5\times10^{-1}$ |
| F311G | $3.7\times10^{-1}$ | **$2.0\times10^{-3}$** | $8.2\times10^{5}$ | $2.9\times10^{-1}$ | $6.8\times10^{-1}$ | $3.3\times10^{-1}$ | $6.6\times10^{-1}$ | N/A | **$1.\times10^{-4}$** | $1.0\times10^{2}$ | $4.6\times10^{-1}$ |
| F311V | $5.6\times10^{-1}$ | $1.4\times10^{-2}$ | $5.5\times10^{5}$ | $7.7\times10^{-1}$ | $8.2\times10^{-1}$ | $6.1\times10^{-1}$ | $9.7\times10^{-1}$ | N/A | $3.8\times10^{-1}$ | $1.4\times10^{2}$ | $9.9\times10^{-1}$ |
| G312A | $7.3\times10^{-1}$ | $2.0\times10^{-1}$ | $4.1\times10^{5}$ | $3.7\times10^{-1}$ | 1.5 | $1.8\times10^{-1}$ | $9.1\times10^{-1}$ | N/A | $6.6\times10^{-2}$ | $8.6\times10^{1}$ | $3.1\times10^{-1}$ |
| G312V | – | – | – | – | – | – | – | – | – | – | – |
| P313G | $7.2\times10^{-1}$ | $2.8\times10^{-1}$ | $4.4\times10^{5}$ | $5.1\times10^{-1}$ | 1.8 | $3.3\times10^{-2}$ | $8.8\times10^{-1}$ | N/A | $2.3\times10^{-2}$ | $7.1\times10^{1}$ | $1.8\times10^{-1}$ |
| P313V | 1.2 | $4.2\times10^{-1}$ | $6.0\times10^{5}$ | $9.8\times10^{-1}$ | $6.6\times10^{-1}$ | $1.9\times10^{-1}$ | N/A | 1.1 | $4.2\times10^{-2}$ | $1.7\times10^{2}$ | $5.8\times10^{-1}$ |
| N314G | $7.4\times10^{-1}$ | $1.5\times10^{-1}$ | $3.8\times10^{5}$ | $1.7\times10^{-1}$ | 1.2 | $6.1\times10^{-1}$ | $6.6\times10^{-1}$ | N/A | **$1.\times10^{-4}$** | $6.6\times10^{1}$ | $1.3\times10^{-1}$ |
| N314V | 1.4 | $1.9\times10^{-1}$ | $9.3\times10^{4}$ | **$2.4\times10^{-2}$** | 7.1 | **0** | $7.2\times10^{-1}$ | N/A | **$2.0\times10^{-4}$** | $8.2\times10^{1}$ | $1.8\times10^{-1}$ |
| S315G | $5.4\times10^{-1}$ | $2.2\times10^{-2}$ | $3.2\times10^{5}$ | $1.9\times10^{-1}$ | 1.0 | $8.8\times10^{-1}$ | $7.5\times10^{-1}$ | N/A | **0** | $1.1\times10^{2}$ | $5.0\times10^{-1}$ |
| S315V | $3.0\times10^{-1}$ | **$1.0\times10^{-3}$** | $4.6\times10^{4}$ | $9.4\times10^{-2}$ | 7.9 | $1.1\times10^{-2}$ | $3.4\times10^{-1}$ | N/A | **0** | $6.0\times10^{1}$ | $1.5\times10^{-1}$ |
| I316G | $4.6\times10^{-1}$ | $1.2\times10^{-2}$ | $3.3\times10^{5}$ | $2.0\times10^{-1}$ | – | – | $6.1\times10^{-1}$ | N/A | **$1.\times10^{-4}$** | $9.1\times10^{1}$ | $3.4\times10^{-1}$ |
| I316V | 1.2 | $4.6\times10^{-1}$ | $4.1\times10^{5}$ | $2.7\times10^{-1}$ | $8.8\times10^{-1}$ | $7.4\times10^{-1}$ | N/A | 1.1 | $2.1\times10^{-1}$ | $1.5\times10^{2}$ | $9.0\times10^{-1}$ |
| E317G | $2.4\times10^{-1}$ | **0** | $8.6\times10^{4}$ | **$8.0\times10^{-3}$** | 3.9 | **$1.1\times10^{-3}$** | $3.8\times10^{-1}$ | N/A | **0** | $4.6\times10^{1}$ | $6.2\times10^{-2}$ |
| E317V | $4.0\times10^{-1}$ | **0** | $4.5\times10^{4}$ | **$1.3\times10^{-2}$** | 6.0 | **$8.7\times10^{-4}$** | $2.5\times10^{-1}$ | N/A | **0** | $3.0\times10^{1}$ | **$2.2\times10^{-2}$** |
| V318A | 1.5 | $1.1\times10^{-1}$ | $1.3\times10^{5}$ | **$2.5\times10^{-2}$** | 2.9 | **$4.3\times10^{-3}$** | N/A | 1.2 | $1.5\times10^{-2}$ | $1.7\times10^{2}$ | $5.5\times10^{-1}$ |
| V318G | $8.6\times10^{-1}$ | $5.5\times10^{-1}$ | $2.9\times10^{5}$ | $1.3\times10^{-1}$ | $8.5\times10^{-1}$ | $6.5\times10^{-1}$ | $8.9\times10^{-1}$ | N/A | $3.6\times10^{-2}$ | $1.1\times10^{2}$ | $5.5\times10^{-1}$ |
| E319G | $3.7\times10^{-1}$ | **0** | $3.6\times10^{5}$ | $2.8\times10^{-1}$ | 1.6 | $1.7\times10^{-1}$ | $7.8\times10^{-1}$ | N/A | **$2.0\times10^{-4}$** | $1.2\times10^{2}$ | $6.6\times10^{-1}$ |
| E319V | $4.7\times10^{-1}$ | **$7.0\times10^{-3}$** | $4.2\times10^{5}$ | $3.7\times10^{-1}$ | 1.6 | $2.2\times10^{-1}$ | $8.4\times10^{-1}$ | N/A | $2.0\times10^{-2}$ | $1.4\times10^{2}$ | $9.8\times10^{-1}$ |
| D320G | $1.5\times10^{-1}$ | **0** | $1.9\times10^{5}$ | **$3.6\times10^{-2}$** | 2.2 | $2.3\times10^{-2}$ | $5.9\times10^{-1}$ | N/A | **0** | $7.7\times10^{1}$ | $1.1\times10^{-1}$ |
| D320V | $1.1\times10^{-1}$ | **0** | $5.5\times10^{4}$ | **$1.9\times10^{-2}$** | 7.7 | **$2.7\times10^{-3}$** | $5.0\times10^{-1}$ | N/A | **0** | $6.0\times10^{1}$ | $1.3\times10^{-1}$ |
| T321G | 1.3 | $3.1\times10^{-1}$ | $2.0\times10^{5}$ | $1.8\times10^{-1}$ | 1.8 | $6.5\times10^{-2}$ | $6.5\times10^{-1}$ | N/A | $1.9\times10^{-2}$ | $9.0\times10^{1}$ | $4.0\times10^{-1}$ |
| T321V | 1.2 | $4.6\times10^{-1}$ | $4.5\times10^{5}$ | $3.3\times10^{-1}$ | 1.1 | $8.3\times10^{-1}$ | N/A | 1.1 | $7.8\times10^{-2}$ | $1.4\times10^{2}$ | $9.9\times10^{-1}$ |
| Y322G | $\leq 4.3\times10^{-2}$ | **0** | $\leq 2.4\times10^{5}$ | **$4.8\times10^{-2}$** | – | – | N/A | 1.4 | **0** | $\geq 1.3\times10^{2}$ | $8.9\times10^{-1}$ |
| Y322V | 1.7 | $2.2\times10^{-2}$ | $8.6\times10^{4}$ | **$2.5\times10^{-2}$** | 3.4 | **$3.9\times10^{-3}$** | N/A | 1.6 | **$6.0\times10^{-4}$** | $2.1\times10^{2}$ | $1.4\times10^{-1}$ |
| L323G | $4.2\times10^{-1}$ | $3.5\times10^{-2}$ | $2.8\times10^{5}$ | $2.3\times10^{-1}$ | – | – | $8.4\times10^{-1}$ | N/A | $1.2\times10^{-2}$ | $1.3\times10^{2}$ | $8.7\times10^{-1}$ |
| L323V | 1.3 | $3.2\times10^{-1}$ | $3.7\times10^{5}$ | $2.3\times10^{-1}$ | 1.0 | 1.0 | N/A | 1.1 | $1.0\times10^{-1}$ | $1.4\times10^{2}$ | 1.0 |
| R324G | $6.9\times10^{-1}$ | $1.6\times10^{-1}$ | $2.1\times10^{5}$ | $1.3\times10^{-1}$ | 2.2 | $2.8\times10^{-2}$ | $7.1\times10^{-1}$ | N/A | **$8.0\times10^{-4}$** | $1.0\times10^{2}$ | $4.8\times10^{-1}$ |
| R324V | 1.2 | $5.2\times10^{-1}$ | $9.5\times10^{4}$ | **$3.2\times10^{-2}$** | 4.3 | **$8.8\times10^{-4}$** | $9.6\times10^{-1}$ | N/A | $3.6\times10^{-1}$ | $1.3\times10^{2}$ | $7.8\times10^{-1}$ |
| L325G | $1.6\times10^{-2}$ | **0** | $2.6\times10^{5}$ | $6.2\times10^{-2}$ | – | – | N/A | 1.1 | $1.3\times10^{-1}$ | $1.3\times10^{2}$ | $7.9\times10^{-1}$ |
| L325V | $8.1\times10^{-1}$ | $3.9\times10^{-1}$ | $2.1\times10^{5}$ | **$4.6\times10^{-2}$** | 2.6 | **$9.4\times10^{-3}$** | N/A | 1.4 | **$1.\times10^{-4}$** | $1.9\times10^{2}$ | $2.4\times10^{-1}$ |

| Variant | $f_a$ | $f_a$ $p$–value | $(k_{cat}/K_M)^{MeP}_{chem.}$ ($M^{-1}s^{-1}$) | $(k_{cat}/K_M)^{MeP}_{chem.}$ $p$–value | FC1 | FC1 $p$–value | FC2 | FC3 | FC2/3 $p$–value | FC4 ($s^{-1}$) | FC4 $p$–value |
|---|---|---|---|---|---|---|---|---|---|---|---|
| D326G | $\le 1.2\times10^{-2}$ | **0** | $\le 1.7\times10^{5}$ | **$4.7\times10^{-2}$** | – | – | $9.9\times10^{-1}$ | N/A | $6.6\times10^{-1}$ | $\ge 1.0\times10^{2}$ | $4.1\times10^{-1}$ |
| D326V | – | – | – | – | – | – | – | – | – | – | – |
| R327G | 1.3 | $3.3\times10^{-1}$ | $1.7\times10^{5}$ | $7.6\times10^{-2}$ | 3.3 | **$6.0\times10^{-4}$** | $9.4\times10^{-1}$ | N/A | $2.5\times10^{-1}$ | $9.1\times10^{1}$ | $4.0\times10^{-1}$ |
| R327V | 1.5 | $5.7\times10^{-2}$ | $2.1\times10^{5}$ | **$4.6\times10^{-2}$** | 2.2 | $1.8\times10^{-2}$ | N/A | 1.1 | $1.8\times10^{-1}$ | $1.8\times10^{2}$ | $3.4\times10^{-1}$ |
| D328G | $7.2\times10^{-1}$ | $1.4\times10^{-1}$ | $2.2\times10^{5}$ | $6.3\times10^{-2}$ | 2.8 | **$2.9\times10^{-3}$** | $9.7\times10^{-1}$ | N/A | $5.2\times10^{-1}$ | $1.1\times10^{2}$ | $4.9\times10^{-1}$ |
| D328V | $4.5\times10^{-1}$ | **$6.0\times10^{-3}$** | $3.0\times10^{5}$ | $1.8\times10^{-1}$ | 2.8 | **$6.9\times10^{-3}$** | N/A | 1.0 | $7.6\times10^{-1}$ | $1.3\times10^{2}$ | $9.0\times10^{-1}$ |
| L329G | $\le 2.0\times10^{-3}$ | **0** | $\le 1.0\times10^{6}$ | $1.3\times10^{-1}$ | – | – | – | – | – | – | – |
| L329V | 1.1 | $5.2\times10^{-1}$ | $3.2\times10^{5}$ | $7.8\times10^{-2}$ | 1.6 | $1.3\times10^{-1}$ | N/A | 1.0 | $5.8\times10^{-1}$ | $1.3\times10^{2}$ | $8.8\times10^{-1}$ |
| A330G | 1.6 | $6.3\times10^{-2}$ | $2.6\times10^{5}$ | $8.8\times10^{-2}$ | 1.3 | $3.2\times10^{-1}$ | $8.9\times10^{-1}$ | N/A | $4.9\times10^{-2}$ | $1.3\times10^{2}$ | $9.0\times10^{-1}$ |
| A330V | 1.2 | $4.9\times10^{-1}$ | $2.5\times10^{5}$ | $7.5\times10^{-2}$ | 1.5 | $2.3\times10^{-1}$ | N/A | 1.1 | $1.4\times10^{-1}$ | $1.7\times10^{2}$ | $5.3\times10^{-1}$ |
| D331G | 1.1 | $7.6\times10^{-1}$ | $2.7\times10^{5}$ | $1.4\times10^{-1}$ | 2.3 | **$5.0\times10^{-3}$** | $9.5\times10^{-1}$ | N/A | $6.0\times10^{-1}$ | $1.2\times10^{2}$ | $7.0\times10^{-1}$ |
| D331V | $6.5\times10^{-1}$ | $6.7\times10^{-2}$ | $8.2\times10^{5}$ | $2.6\times10^{-1}$ | $7.3\times10^{-1}$ | $2.7\times10^{-1}$ | $9.8\times10^{-1}$ | N/A | $5.9\times10^{-1}$ | $1.7\times10^{2}$ | $5.6\times10^{-1}$ |
| F332G | $\le 5.3\times10^{-3}$ | **0** | $\le 1.0\times10^{6}$ | **$4.6\times10^{-2}$** | – | – | $9.3\times10^{-1}$ | N/A | $1.1\times10^{-1}$ | $\ge 1.0\times10^{2}$ | $4.3\times10^{-1}$ |
| F332V | 1.1 | $7.6\times10^{-1}$ | $5.6\times10^{4}$ | **$2.4\times10^{-2}$** | 5.7 | **$1.1\times10^{-4}$** | $9.0\times10^{-1}$ | N/A | $1.0\times10^{-1}$ | $1.0\times10^{2}$ | $5.0\times10^{-1}$ |
| F333G | $\le 4.7\times10^{-2}$ | **0** | $\le 1.3\times10^{5}$ | $5.7\times10^{-2}$ | – | – | $7.3\times10^{-1}$ | N/A | **0** | $\ge 1.2\times10^{2}$ | $7.1\times10^{-1}$ |
| F333V | 1.4 | $2.1\times10^{-1}$ | $7.1\times10^{4}$ | **$2.6\times10^{-2}$** | 3.9 | **$1.2\times10^{-3}$** | N/A | 1.0 | $5.7\times10^{-1}$ | $1.5\times10^{2}$ | $8.7\times10^{-1}$ |
| N334G | $8.6\times10^{-1}$ | $5.0\times10^{-1}$ | $5.5\times10^{5}$ | $7.2\times10^{-1}$ | 1.4 | $2.9\times10^{-1}$ | $8.5\times10^{-1}$ | N/A | **$7.3\times10^{-3}$** | $1.4\times10^{2}$ | $9.3\times10^{-1}$ |
| N334V | $9.6\times10^{-1}$ | $8.2\times10^{-1}$ | $8.7\times10^{5}$ | $1.4\times10^{-1}$ | $5.1\times10^{-1}$ | $2.2\times10^{-2}$ | N/A | 1.0 | $6.2\times10^{-1}$ | $1.7\times10^{2}$ | $5.0\times10^{-1}$ |
| N335G | 1.0 | $8.9\times10^{-1}$ | $5.9\times10^{5}$ | $8.9\times10^{-1}$ | 1.1 | $6.3\times10^{-1}$ | $9.9\times10^{-1}$ | N/A | $9.2\times10^{-1}$ | $9.2\times10^{1}$ | $3.6\times10^{-1}$ |
| N335V | 1.5 | $7.4\times10^{-2}$ | $4.1\times10^{5}$ | $2.3\times10^{-1}$ | $9.9\times10^{-1}$ | $9.7\times10^{-1}$ | N/A | 1.1 | $7.4\times10^{-2}$ | $1.6\times10^{2}$ | $6.9\times10^{-1}$ |
| L336G | $\le 3.2\times10^{-3}$ | **0** | $\le 7.1\times10^{5}$ | $6.4\times10^{-1}$ | – | – | N/A | 1.1 | $4.5\times10^{-1}$ | $\ge 8.0\times10^{1}$ | $2.7\times10^{-1}$ |
| L336V | $5.5\times10^{-1}$ | $5.4\times10^{-2}$ | $8.4\times10^{5}$ | $2.4\times10^{-1}$ | $8.8\times10^{-1}$ | $7.0\times10^{-1}$ | $9.1\times10^{-1}$ | N/A | $1.4\times10^{-1}$ | $2.2\times10^{2}$ | $1.6\times10^{-1}$ |
| D337G | $9.7\times10^{-2}$ | **0** | $2.4\times10^{5}$ | **$4.2\times10^{-2}$** | – | – | $9.5\times10^{-1}$ | N/A | $2.0\times10^{-1}$ | $1.2\times10^{2}$ | $7.2\times10^{-1}$ |
| D337V | $1.8\times10^{-1}$ | **0** | $5.4\times10^{4}$ | $5.0\times10^{-2}$ | – | – | $7.3\times10^{-1}$ | N/A | **$8.0\times10^{-4}$** | $1.4\times10^{2}$ | $9.7\times10^{-1}$ |
| K338G | $9.3\times10^{-1}$ | $7.2\times10^{-1}$ | $2.4\times10^{5}$ | $9.1\times10^{-2}$ | 2.6 | $1.6\times10^{-2}$ | $9.2\times10^{-1}$ | N/A | $9.0\times10^{-2}$ | $9.6\times10^{1}$ | $3.1\times10^{-1}$ |
| K338V | 1.3 | $3.3\times10^{-1}$ | $2.8\times10^{5}$ | $8.6\times10^{-2}$ | 1.9 | $4.3\times10^{-2}$ | N/A | 1.1 | $2.6\times10^{-1}$ | $2.0\times10^{2}$ | $2.4\times10^{-1}$ |
| K339G | $6.7\times10^{-1}$ | $2.6\times10^{-1}$ | $3.3\times10^{5}$ | $2.8\times10^{-1}$ | 1.4 | $2.5\times10^{-1}$ | $8.2\times10^{-1}$ | N/A | **$7.2\times10^{-3}$** | $8.9\times10^{1}$ | $4.3\times10^{-1}$ |
| K339V | $7.8\times10^{-1}$ | $2.9\times10^{-1}$ | $5.0\times10^{5}$ | $5.3\times10^{-1}$ | 1.0 | $9.9\times10^{-1}$ | $9.6\times10^{-1}$ | N/A | $3.9\times10^{-1}$ | $1.4\times10^{2}$ | $9.5\times10^{-1}$ |
| V340A | 1.3 | $3.4\times10^{-1}$ | $8.5\times10^{4}$ | **$4.3\times10^{-2}$** | 5.2 | **$1.9\times10^{-3}$** | $9.0\times10^{-1}$ | N/A | $1.3\times10^{-1}$ | $1.5\times10^{2}$ | $8.0\times10^{-1}$ |
| V340G | $8.6\times10^{-2}$ | **0** | $2.5\times10^{5}$ | $8.1\times10^{-2}$ | – | – | N/A | 1.2 | **0** | $1.4\times10^{2}$ | 1.0 |
| G341A | $4.4\times10^{-1}$ | **$3.0\times10^{-3}$** | $5.5\times10^{5}$ | $7.7\times10^{-1}$ | $8.3\times10^{-1}$ | $6.0\times10^{-1}$ | $9.4\times10^{-1}$ | N/A | $3.0\times10^{-1}$ | $9.5\times10^{1}$ | $4.0\times10^{-1}$ |
| G341V | $7.4\times10^{-1}$ | $2.0\times10^{-1}$ | $6.2\times10^{4}$ | **$8.0\times10^{-3}$** | 4.6 | **$7.4\times10^{-4}$** | $9.9\times10^{-1}$ | N/A | $8.7\times10^{-1}$ | $1.3\times10^{2}$ | $8.4\times10^{-1}$ |
| K342G | 1.0 | $9.9\times10^{-1}$ | $3.3\times10^{5}$ | $1.9\times10^{-1}$ | 1.5 | $1.1\times10^{-1}$ | $8.8\times10^{-1}$ | N/A | $1.1\times10^{-1}$ | $8.8\times10^{1}$ | $3.7\times10^{-1}$ |
| K342V | 1.3 | $1.2\times10^{-1}$ | $3.6\times10^{5}$ | $1.4\times10^{-1}$ | 1.3 | $3.4\times10^{-1}$ | N/A | 1.1 | $1.9\times10^{-1}$ | $1.5\times10^{2}$ | $8.8\times10^{-1}$ |
| G343A | 1.4 | $3.1\times10^{-1}$ | $6.2\times10^{5}$ | $9.6\times10^{-1}$ | $8.0\times10^{-1}$ | $5.9\times10^{-1}$ | 1.0 | N/A | $9.8\times10^{-1}$ | $1.0\times10^{2}$ | $5.1\times10^{-1}$ |
| G343V | 1.4 | $1.6\times10^{-1}$ | $1.5\times10^{5}$ | **$3.1\times10^{-2}$** | 2.5 | $1.3\times10^{-2}$ | N/A | 1.0 | $6.8\times10^{-1}$ | $1.6\times10^{2}$ | $7.1\times10^{-1}$ |
| N344G | $9.4\times10^{-1}$ | $7.6\times10^{-1}$ | $3.4\times10^{5}$ | $1.6\times10^{-1}$ | 1.4 | $2.0\times10^{-1}$ | $8.2\times10^{-1}$ | N/A | $1.2\times10^{-2}$ | $9.0\times10^{1}$ | $3.0\times10^{-1}$ |
| N344V | $2.4\times10^{-1}$ | **0** | $6.8\times10^{5}$ | $7.3\times10^{-1}$ | – | – | $8.6\times10^{-1}$ | N/A | $2.0\times10^{-2}$ | $1.3\times10^{2}$ | $9.2\times10^{-1}$ |
| Y345G | $6.2\times10^{-2}$ | **0** | $4.9\times10^{5}$ | $5.7\times10^{-1}$ | – | – | $8.3\times10^{-1}$ | N/A | **$8.0\times10^{-4}$** | $1.0\times10^{2}$ | $4.7\times10^{-1}$ |
| Y345V | 1.0 | $9.9\times10^{-1}$ | $2.4\times10^{5}$ | $6.4\times10^{-2}$ | 2.0 | $2.7\times10^{-2}$ | N/A | 1.1 | $1.8\times10^{-1}$ | $1.6\times10^{2}$ | $5.8\times10^{-1}$ |
| L346G | $3.3\times10^{-1}$ | **0** | $3.5\times10^{5}$ | $3.0\times10^{-1}$ | 1.6 | $1.5\times10^{-1}$ | $7.9\times10^{-1}$ | N/A | **$2.6\times10^{-3}$** | $1.2\times10^{2}$ | $6.5\times10^{-1}$ |
| L346V | 1.4 | $1.7\times10^{-1}$ | $3.8\times10^{5}$ | $2.6\times10^{-1}$ | 1.2 | $6.5\times10^{-1}$ | N/A | 1.0 | $4.0\times10^{-1}$ | $1.5\times10^{2}$ | $7.4\times10^{-1}$ |
| V347A | 1.6 | $6.7\times10^{-2}$ | $4.2\times10^{5}$ | $3.0\times10^{-1}$ | $7.1\times10^{-1}$ | $3.3\times10^{-1}$ | N/A | 1.0 | $6.0\times10^{-1}$ | $1.5\times10^{2}$ | $8.5\times10^{-1}$ |
| V347G | $\le 8.8\times10^{-1}$ | $6.4\times10^{-1}$ | $\le 9.7\times10^{4}$ | **$3.4\times10^{-2}$** | $\le 2.3$ | $1.3\times10^{-2}$ | $8.8\times10^{-1}$ | N/A | $2.8\times10^{-2}$ | $\ge 1.2\times10^{2}$ | $6.3\times10^{-1}$ |
| F348G | $1.0\times10^{-1}$ | **0** | $2.8\times10^{5}$ | $9.6\times10^{-2}$ | – | – | $9.5\times10^{-1}$ | N/A | $1.8\times10^{-1}$ | $9.0\times10^{1}$ | $2.9\times10^{-1}$ |
| F348V | $4.8\times10^{-1}$ | **$7.0\times10^{-3}$** | $4.7\times10^{5}$ | $4.1\times10^{-1}$ | $6.5\times10^{-1}$ | $1.7\times10^{-1}$ | N/A | 1.0 | $9.7\times10^{-1}$ | $1.4\times10^{2}$ | $9.5\times10^{-1}$ |
| L349G | $8.0\times10^{-2}$ | **0** | $2.4\times10^{5}$ | $6.3\times10^{-2}$ | – | – | N/A | 1.2 | **$1.\times10^{-4}$** | $1.3\times10^{2}$ | $8.2\times10^{-1}$ |
| L349V | $4.7\times10^{-1}$ | **$3.0\times10^{-3}$** | $5.6\times10^{5}$ | $7.9\times10^{-1}$ | – | – | $8.7\times10^{-1}$ | N/A | **$7.6\times10^{-3}$** | $1.7\times10^{2}$ | $5.6\times10^{-1}$ |
| S350G | $3.6\times10^{-1}$ | **$1.0\times10^{-3}$** | $4.5\times10^{5}$ | $4.4\times10^{-1}$ | 1.5 | $1.8\times10^{-1}$ | N/A | 1.2 | **$1.\times10^{-4}$** | $1.3\times10^{2}$ | $8.8\times10^{-1}$ |
| S350V | $6.3\times10^{-1}$ | $2.5\times10^{-2}$ | $2.5\times10^{5}$ | $6.2\times10^{-2}$ | $9.2\times10^{-1}$ | $7.9\times10^{-1}$ | N/A | 1.3 | **$9.0\times10^{-4}$** | $1.6\times10^{2}$ | $6.5\times10^{-1}$ |
| A351G | $7.6\times10^{-1}$ | $3.3\times10^{-1}$ | $2.1\times10^{5}$ | $1.1\times10^{-1}$ | – | – | $6.6\times10^{-1}$ | N/A | **$1.\times10^{-4}$** | $1.5\times10^{2}$ | $9.2\times10^{-1}$ |
| A351V | $7.2\times10^{-1}$ | $1.3\times10^{-1}$ | $6.4\times10^{4}$ | **$5.0\times10^{-3}$** | – | – | $7.4\times10^{-1}$ | N/A | **$1.\times10^{-4}$** | $8.5\times10^{1}$ | $1.8\times10^{-1}$ |
| D352G | $8.7\times10^{-2}$ | **0** | $5.9\times10^{6}$ | **0** | 1.6 | $1.2\times10^{-1}$ | $3.8\times10^{-2}$ | N/A | **0** | – | – |
| D352V | – | – | – | – | – | – | – | – | – | – | – |
| H353G | – | – | – | – | – | – | – | – | – | – | – |
| H353V | $6.6\times10^{-1}$ | $9.0\times10^{-2}$ | $1.2\times10^{5}$ | **$3.1\times10^{-2}$** | 1.0 | $9.0\times10^{-1}$ | $8.1\times10^{-1}$ | N/A | **$6.9\times10^{-3}$** | $8.5\times10^{1}$ | $2.3\times10^{-1}$ |
| G354A | $2.4\times10^{-1}$ | **0** | $4.4\times10^{5}$ | $5.0\times10^{-1}$ | 4.7 | **$1.4\times10^{-3}$** | $9.7\times10^{-1}$ | N/A | $4.6\times10^{-1}$ | $7.4\times10^{1}$ | $2.2\times10^{-1}$ |
| G354V | – | – | – | – | – | – | – | – | – | – | – |
| A355G | 1.4 | $2.4\times10^{-1}$ | $3.3\times10^{5}$ | $2.1\times10^{-1}$ | 2.1 | $1.3\times10^{-2}$ | $9.4\times10^{-1}$ | N/A | $3.7\times10^{-1}$ | $1.1\times10^{2}$ | $5.9\times10^{-1}$ |
| A355V | $3.0\times10^{-1}$ | **0** | $6.2\times10^{5}$ | $9.6\times10^{-1}$ | – | – | $8.6\times10^{-1}$ | N/A | $1.5\times10^{-2}$ | $1.6\times10^{2}$ | $6.8\times10^{-1}$ |
| A356G | 1.3 | $4.1\times10^{-1}$ | $4.1\times10^{5}$ | $2.6\times10^{-1}$ | 2.5 | **$2.3\times10^{-3}$** | $7.7\times10^{-1}$ | N/A | **0** | $1.1\times10^{2}$ | $5.0\times10^{-1}$ |
| A356V | $7.4\times10^{-1}$ | $9.0\times10^{-2}$ | $4.8\times10^{5}$ | $3.6\times10^{-1}$ | $9.5\times10^{-1}$ | $8.1\times10^{-1}$ | $7.5\times10^{-1}$ | N/A | **0** | $1.1\times10^{2}$ | $3.6\times10^{-1}$ |
| H357G | 3.4 | **$3.0\times10^{-3}$** | $6.3\times10^{5}$ | $9.2\times10^{-1}$ | $3.0\times10^{-1}$ | **$7.0\times10^{-3}$** | $5.1\times10^{-1}$ | N/A | **$9.0\times10^{-4}$** | $5.3\times10^{1}$ | $2.3\times10^{-1}$ |
| H357V | $5.7\times10^{-1}$ | $2.0\times10^{-2}$ | $3.6\times10^{5}$ | $1.9\times10^{-1}$ | 1.5 | $2.2\times10^{-1}$ | $7.7\times10^{-1}$ | N/A | **$1.9\times10^{-3}$** | $6.7\times10^{1}$ | $1.4\times10^{-1}$ |
| S358G | $6.7\times10^{-1}$ | $1.0\times10^{-1}$ | $5.1\times10^{5}$ | $6.7\times10^{-1}$ | 1.0 | $9.9\times10^{-1}$ | $7.5\times10^{-1}$ | N/A | **0** | $5.1\times10^{1}$ | $8.9\times10^{-2}$ |
| S358V | $4.8\times10^{-1}$ | $1.6\times10^{-2}$ | $2.4\times10^{5}$ | $1.2\times10^{-1}$ | 1.9 | $9.2\times10^{-2}$ | $8.1\times10^{-1}$ | N/A | **$5.7\times10^{-3}$** | $1.2\times10^{2}$ | $6.6\times10^{-1}$ |
| V359A | $6.6\times10^{-1}$ | $6.7\times10^{-2}$ | $2.4\times10^{5}$ | $8.2\times10^{-2}$ | 2.1 | $3.9\times10^{-2}$ | $9.2\times10^{-1}$ | N/A | $1.6\times10^{-1}$ | $1.6\times10^{2}$ | $6.9\times10^{-1}$ |
| V359G | $4.3\times10^{-1}$ | **$5.0\times10^{-3}$** | $8.7\times10^{5}$ | $1.8\times10^{-1}$ | $5.3\times10^{-1}$ | $3.9\times10^{-2}$ | $6.3\times10^{-1}$ | N/A | **$2.0\times10^{-4}$** | $1.1\times10^{2}$ | $5.5\times10^{-1}$ |
| G360A | $8.4\times10^{-1}$ | $5.2\times10^{-1}$ | $6.5\times10^{5}$ | $8.5\times10^{-1}$ | $9.3\times10^{-1}$ | $7.8\times10^{-1}$ | $8.4\times10^{-1}$ | N/A | **$4.2\times10^{-3}$** | $1.4\times10^{2}$ | $9.9\times10^{-1}$ |
| G360V | $4.9\times10^{-1}$ | **$2.0\times10^{-3}$** | $7.1\times10^{5}$ | $5.5\times10^{-1}$ | $6.3\times10^{-1}$ | $1.5\times10^{-1}$ | $9.5\times10^{-1}$ | N/A | $1.7\times10^{-1}$ | $1.3\times10^{2}$ | $7.9\times10^{-1}$ |
| F361G | $4.5\times10^{-2}$ | **0** | $1.9\times10^{5}$ | **$2.6\times10^{-2}$** | 2.8 | $3.3\times10^{-2}$ | $5.0\times10^{-1}$ | N/A | **0** | $5.3\times10^{1}$ | **$4.6\times10^{-2}$** |
| F361V | $3.4\times10^{-1}$ | **$1.0\times10^{-3}$** | $2.5\times10^{4}$ | **$2.1\times10^{-2}$** | $1.8\times10^{1}$ | **$5.0\times10^{-5}$** | $8.3\times10^{-1}$ | N/A | **$8.0\times10^{-3}$** | $1.1\times10^{2}$ | $5.5\times10^{-1}$ |
| M362G | $1.5\times10^{-1}$ | **0** | $1.3\times10^{6}$ | **$2.4\times10^{-2}$** | – | – | $6.6\times10^{-1}$ | N/A | **0** | $5.6\times10^{1}$ | $1.5\times10^{-1}$ |
| M362V | $1.6\times10^{-1}$ | **0** | $6.8\times10^{4}$ | **$1.7\times10^{-2}$** | – | – | N/A | 1.0 | $4.4\times10^{-1}$ | $1.5\times10^{2}$ | $7.6\times10^{-1}$ |
| Q363G | $6.3\times10^{-1}$ | $8.3\times10^{-2}$ | $5.0\times10^{5}$ | $6.0\times10^{-1}$ | $7.5\times10^{-1}$ | $3.7\times10^{-1}$ | $7.2\times10^{-1}$ | N/A | **$1.\times10^{-4}$** | $9.7\times10^{1}$ | $4.1\times10^{-1}$ |

| Variant | $f_a$ | $f_a$ $p$–value | $(k_{cat}/K_M)^{MeP}_{chem.}$ ($M^{-1}s^{-1}$) | $(k_{cat}/K_M)^{MeP}_{chem.}$ $p$–value | FC1 | FC1 $p$–value | FC2 | FC3 | FC2/3 $p$–value | FC4 ($s^{-1}$) | FC4 $p$–value |
|---|---|---|---|---|---|---|---|---|---|---|---|
| Q363V | 1.5 | $4.3\times10^{-2}$ | $2.1\times10^{5}$ | **$3.5\times10^{-2}$** | 2.1 | $2.7\times10^{-2}$ | N/A | 1.0 | $4.0\times10^{-1}$ | $1.2\times10^{2}$ | $7.0\times10^{-1}$ |
| A364V | 1.2 | $4.7\times10^{-1}$ | $1.8\times10^{5}$ | **$3.8\times10^{-2}$** | 2.7 | **$1.5\times10^{-3}$** | N/A | 1.1 | $4.9\times10^{-2}$ | $1.6\times10^{2}$ | $7.0\times10^{-1}$ |
| H365G | $3.8\times10^{-1}$ | **0** | $7.7\times10^{5}$ | $4.3\times10^{-1}$ | $7.2\times10^{-1}$ | $2.7\times10^{-1}$ | $9.5\times10^{-1}$ | N/A | $3.1\times10^{-1}$ | $1.1\times10^{2}$ | $5.9\times10^{-1}$ |
| H365V | $9.2\times10^{-1}$ | $7.6\times10^{-1}$ | $2.3\times10^{5}$ | $7.1\times10^{-2}$ | 2.8 | **$5.1\times10^{-3}$** | N/A | 1.0 | $2.5\times10^{-1}$ | $1.5\times10^{2}$ | $7.9\times10^{-1}$ |
| K366G | ≤ $3.8\times10^{-1}$ | **$4.0\times10^{-3}$** | ≤ $6.0\times10^{5}$ | $9.7\times10^{-1}$ | ≤ 1.4 | $3.3\times10^{-1}$ | $8.1\times10^{-1}$ | N/A | **$6.0\times10^{-4}$** | ≥ $9.7\times10^{1}$ | $4.2\times10^{-1}$ |
| K366V | $4.3\times10^{-1}$ | **$4.0\times10^{-3}$** | $5.3\times10^{4}$ | **$2.0\times10^{-2}$** | 8.4 | **$1.8\times10^{-4}$** | N/A | 1.0 | $3.2\times10^{-1}$ | $1.6\times10^{2}$ | $7.4\times10^{-1}$ |
| M367G | $1.0\times10^{-1}$ | **0** | $3.6\times10^{5}$ | $1.6\times10^{-1}$ | 2.0 | $6.1\times10^{-2}$ | $5.8\times10^{-1}$ | N/A | **0** | $7.5\times10^{1}$ | $1.6\times10^{-1}$ |
| M367V | $4.8\times10^{-1}$ | **$3.0\times10^{-3}$** | $2.0\times10^{5}$ | **$4.2\times10^{-2}$** | 2.5 | $1.2\times10^{-2}$ | N/A | 1.1 | $4.0\times10^{-2}$ | $1.5\times10^{2}$ | $7.8\times10^{-1}$ |
| P368G | $2.4\times10^{-1}$ | **0** | $2.2\times10^{5}$ | $1.1\times10^{-1}$ | 3.1 | **$4.2\times10^{-3}$** | $8.1\times10^{-1}$ | N/A | **$3.0\times10^{-4}$** | $9.7\times10^{1}$ | $3.9\times10^{-1}$ |
| P368V | $8.4\times10^{-1}$ | $4.9\times10^{-1}$ | $1.7\times10^{5}$ | **$3.7\times10^{-2}$** | 3.7 | **$2.8\times10^{-4}$** | $9.4\times10^{-1}$ | N/A | $1.6\times10^{-1}$ | $1.4\times10^{2}$ | $9.5\times10^{-1}$ |
| T369G | 1.7 | $3.6\times10^{-2}$ | $2.8\times10^{5}$ | $1.0\times10^{-1}$ | 1.1 | $7.0\times10^{-1}$ | $8.6\times10^{-1}$ | N/A | **$8.9\times10^{-3}$** | $1.2\times10^{2}$ | $7.1\times10^{-1}$ |
| T369V | 1.2 | $5.4\times10^{-1}$ | $3.2\times10^{5}$ | $1.3\times10^{-1}$ | 1.1 | $8.6\times10^{-1}$ | $9.0\times10^{-1}$ | N/A | $1.1\times10^{-1}$ | $1.4\times10^{2}$ | $9.9\times10^{-1}$ |
| G370A | $3.0\times10^{-1}$ | **0** | $8.6\times10^{5}$ | $2.2\times10^{-1}$ | $6.6\times10^{-1}$ | $2.0\times10^{-1}$ | $9.2\times10^{-1}$ | N/A | $8.1\times10^{-2}$ | $1.1\times10^{2}$ | $4.8\times10^{-1}$ |
| G370V | $8.2\times10^{-2}$ | **0** | $6.3\times10^{4}$ | **$3.8\times10^{-2}$** | – | – | – | – | – | – | – |
| F371G | $1.7\times10^{-1}$ | **0** | $1.5\times10^{6}$ | **$4.0\times10^{-3}$** | $3.9\times10^{-1}$ | **$8.2\times10^{-3}$** | $9.0\times10^{-1}$ | N/A | $5.3\times10^{-2}$ | $2.1\times10^{2}$ | $1.6\times10^{-1}$ |
| F371V | $9.3\times10^{-1}$ | $6.2\times10^{-1}$ | $6.1\times10^{5}$ | $9.9\times10^{-1}$ | $5.3\times10^{-1}$ | $6.1\times10^{-2}$ | N/A | 1.0 | $9.4\times10^{-1}$ | $3.2\times10^{2}$ | **$7.0\times10^{-3}$** |
| F372G | $6.5\times10^{-1}$ | $5.9\times10^{-2}$ | $8.0\times10^{5}$ | $2.9\times10^{-1}$ | 1.3 | $4.5\times10^{-1}$ | $9.6\times10^{-1}$ | N/A | $4.1\times10^{-1}$ | $1.6\times10^{2}$ | $7.0\times10^{-1}$ |
| F372V | $9.8\times10^{-1}$ | $9.6\times10^{-1}$ | $5.3\times10^{5}$ | $6.1\times10^{-1}$ | $7.9\times10^{-1}$ | $4.0\times10^{-1}$ | N/A | 1.1 | $4.4\times10^{-2}$ | $1.9\times10^{2}$ | $2.8\times10^{-1}$ |
| V373A | $8.5\times10^{-1}$ | $5.7\times10^{-1}$ | $6.3\times10^{5}$ | $9.1\times10^{-1}$ | $5.2\times10^{-1}$ | $2.6\times10^{-2}$ | N/A | 1.2 | **$6.6\times10^{-3}$** | $1.6\times10^{2}$ | $6.6\times10^{-1}$ |
| V373G | $7.1\times10^{-1}$ | $2.7\times10^{-1}$ | $9.2\times10^{5}$ | $2.0\times10^{-1}$ | $7.3\times10^{-1}$ | $4.1\times10^{-1}$ | $8.8\times10^{-1}$ | N/A | $3.4\times10^{-2}$ | $9.5\times10^{1}$ | $4.7\times10^{-1}$ |
| E374G | 1.5 | $9.2\times10^{-2}$ | $3.1\times10^{5}$ | $1.4\times10^{-1}$ | 1.9 | $2.9\times10^{-2}$ | $8.2\times10^{-1}$ | N/A | **$6.7\times10^{-3}$** | $1.1\times10^{2}$ | $6.0\times10^{-1}$ |
| E374V | 1.5 | $9.2\times10^{-2}$ | $3.8\times10^{5}$ | $2.3\times10^{-1}$ | 1.6 | $1.1\times10^{-1}$ | $9.1\times10^{-1}$ | N/A | $1.1\times10^{-1}$ | $1.7\times10^{2}$ | $5.2\times10^{-1}$ |
| D375G | $6.2\times10^{-1}$ | $5.3\times10^{-2}$ | $3.6\times10^{5}$ | $2.2\times10^{-1}$ | 2.1 | $2.0\times10^{-2}$ | $7.9\times10^{-1}$ | N/A | **$3.0\times10^{-4}$** | $1.1\times10^{2}$ | $5.6\times10^{-1}$ |
| D375V | $7.6\times10^{-1}$ | $2.4\times10^{-1}$ | $1.3\times10^{5}$ | $5.1\times10^{-2}$ | 3.8 | **$6.4\times10^{-3}$** | 1.0 | N/A | $8.7\times10^{-1}$ | $1.5\times10^{2}$ | $7.8\times10^{-1}$ |
| M376G | $3.2\times10^{-1}$ | **$4.0\times10^{-3}$** | $9.0\times10^{5}$ | $2.1\times10^{-1}$ | 1.1 | $6.7\times10^{-1}$ | $9.3\times10^{-1}$ | N/A | $1.7\times10^{-1}$ | $1.8\times10^{2}$ | $5.7\times10^{-1}$ |
| M376V | 1.1 | $6.6\times10^{-1}$ | $7.0\times10^{5}$ | $6.0\times10^{-1}$ | $6.0\times10^{-1}$ | $1.4\times10^{-1}$ | N/A | 1.0 | $6.8\times10^{-1}$ | $1.2\times10^{2}$ | $7.2\times10^{-1}$ |
| K377G | 1.3 | $4.5\times10^{-1}$ | $4.5\times10^{5}$ | $4.5\times10^{-1}$ | $9.3\times10^{-1}$ | $7.6\times10^{-1}$ | $9.9\times10^{-1}$ | N/A | $7.8\times10^{-1}$ | $1.3\times10^{2}$ | $9.2\times10^{-1}$ |
| K377V | $9.8\times10^{-1}$ | $8.4\times10^{-1}$ | $6.2\times10^{5}$ | $9.7\times10^{-1}$ | $6.5\times10^{-1}$ | $2.7\times10^{-1}$ | $9.3\times10^{-1}$ | N/A | $1.5\times10^{-1}$ | $1.9\times10^{2}$ | $2.1\times10^{-1}$ |
| K378G | $9.5\times10^{-1}$ | $7.9\times10^{-1}$ | $7.5\times10^{5}$ | $4.6\times10^{-1}$ | 1.3 | $3.8\times10^{-1}$ | $9.6\times10^{-1}$ | N/A | $3.5\times10^{-1}$ | $1.1\times10^{2}$ | $4.3\times10^{-1}$ |
| K378V | 1.2 | $5.2\times10^{-1}$ | $2.6\times10^{5}$ | $9.3\times10^{-2}$ | 2.1 | $3.2\times10^{-2}$ | N/A | 1.1 | $2.5\times10^{-2}$ | $2.3\times10^{2}$ | $1.0\times10^{-1}$ |
| E379G | 1.4 | $2.3\times10^{-1}$ | $3.0\times10^{5}$ | $1.2\times10^{-1}$ | 1.4 | $1.5\times10^{-1}$ | $9.1\times10^{-1}$ | N/A | $7.4\times10^{-2}$ | $1.0\times10^{2}$ | $4.6\times10^{-1}$ |
| E379V | 1.5 | $1.1\times10^{-1}$ | $4.5\times10^{5}$ | $4.5\times10^{-1}$ | 1.3 | $5.4\times10^{-1}$ | N/A | 1.1 | $3.7\times10^{-1}$ | $1.8\times10^{2}$ | $3.2\times10^{-1}$ |
| M380G | 1.0 | $9.8\times10^{-1}$ | $3.2\times10^{5}$ | $6.2\times10^{-2}$ | 1.4 | $1.6\times10^{-1}$ | $8.9\times10^{-1}$ | N/A | **$3.3\times10^{-3}$** | $1.3\times10^{2}$ | $8.3\times10^{-1}$ |
| M380V | $9.9\times10^{-1}$ | $9.4\times10^{-1}$ | $3.7\times10^{5}$ | $1.7\times10^{-1}$ | 1.9 | $6.0\times10^{-2}$ | N/A | 1.0 | $5.8\times10^{-1}$ | $1.7\times10^{2}$ | $5.4\times10^{-1}$ |
| N381G | $9.7\times10^{-1}$ | $8.4\times10^{-1}$ | $5.7\times10^{5}$ | $8.5\times10^{-1}$ | $9.7\times10^{-1}$ | $9.5\times10^{-1}$ | N/A | 1.0 | $4.5\times10^{-1}$ | $8.3\times10^{1}$ | $2.5\times10^{-1}$ |
| N381V | $4.1\times10^{-1}$ | **$3.0\times10^{-3}$** | $4.2\times10^{5}$ | $3.5\times10^{-1}$ | 1.2 | $5.9\times10^{-1}$ | $8.9\times10^{-1}$ | N/A | $3.1\times10^{-2}$ | $1.5\times10^{2}$ | $7.5\times10^{-1}$ |
| A382G | $9.5\times10^{-1}$ | $8.4\times10^{-1}$ | $1.2\times10^{6}$ | **$3.8\times10^{-2}$** | $6.6\times10^{-1}$ | $1.3\times10^{-1}$ | N/A | 1.0 | $7.0\times10^{-1}$ | $1.3\times10^{2}$ | $8.4\times10^{-1}$ |
| A382V | 1.3 | $1.9\times10^{-1}$ | $4.2\times10^{5}$ | $2.9\times10^{-1}$ | 1.0 | $9.1\times10^{-1}$ | N/A | 1.1 | $3.8\times10^{-2}$ | $1.8\times10^{2}$ | $3.5\times10^{-1}$ |
| K383G | $8.1\times10^{-1}$ | $4.4\times10^{-1}$ | $6.9\times10^{5}$ | $7.3\times10^{-1}$ | $6.4\times10^{-1}$ | $1.1\times10^{-1}$ | $8.5\times10^{-1}$ | N/A | $1.3\times10^{-2}$ | $8.9\times10^{1}$ | $3.9\times10^{-1}$ |
| K383V | 1.1 | $5.7\times10^{-1}$ | $2.3\times10^{5}$ | **$4.6\times10^{-2}$** | 1.8 | $2.9\times10^{-2}$ | N/A | 1.1 | **$7.0\times10^{-3}$** | $1.7\times10^{2}$ | $4.2\times10^{-1}$ |
| L384G | $8.2\times10^{-1}$ | $4.9\times10^{-1}$ | $2.6\times10^{5}$ | $1.7\times10^{-1}$ | 2.6 | **$2.9\times10^{-3}$** | $8.0\times10^{-1}$ | N/A | **$4.7\times10^{-3}$** | $7.7\times10^{1}$ | $2.2\times10^{-1}$ |
| L384V | $8.8\times10^{-1}$ | $6.4\times10^{-1}$ | $4.7\times10^{5}$ | $5.5\times10^{-1}$ | 1.2 | $6.5\times10^{-1}$ | N/A | 1.1 | $4.7\times10^{-1}$ | $1.5\times10^{2}$ | $8.9\times10^{-1}$ |
| K385G | 1.1 | $8.0\times10^{-1}$ | $4.2\times10^{5}$ | $2.9\times10^{-1}$ | 1.0 | $9.4\times10^{-1}$ | N/A | 1.2 | **$3.9\times10^{-3}$** | $1.8\times10^{2}$ | $4.4\times10^{-1}$ |
| K385V | 1.5 | $4.0\times10^{-2}$ | $2.9\times10^{5}$ | **$4.2\times10^{-2}$** | 1.8 | $7.4\times10^{-2}$ | N/A | 1.2 | **$1.7\times10^{-3}$** | $1.4\times10^{2}$ | $9.4\times10^{-1}$ |
| Q386G | $8.1\times10^{-2}$ | **0** | $1.1\times10^{6}$ | $8.8\times10^{-2}$ | – | – | $7.3\times10^{-1}$ | N/A | **$3.0\times10^{-4}$** | $1.1\times10^{2}$ | $6.4\times10^{-1}$ |
| Q386V | 1.5 | $1.7\times10^{-1}$ | $3.1\times10^{5}$ | $2.8\times10^{-1}$ | 1.8 | $5.3\times10^{-2}$ | N/A | 1.0 | $6.3\times10^{-1}$ | $1.4\times10^{2}$ | $9.7\times10^{-1}$ |
| K387G | $5.9\times10^{-1}$ | $3.2\times10^{-2}$ | $6.8\times10^{5}$ | $7.1\times10^{-1}$ | 1.2 | $4.7\times10^{-1}$ | $9.0\times10^{-1}$ | N/A | $6.6\times10^{-2}$ | $1.3\times10^{2}$ | $8.3\times10^{-1}$ |
| K387V | 1.3 | $2.4\times10^{-1}$ | $3.2\times10^{5}$ | $1.1\times10^{-1}$ | 1.4 | $2.3\times10^{-1}$ | N/A | 1.1 | $2.4\times10^{-2}$ | $2.2\times10^{2}$ | $7.5\times10^{-2}$ |
| F388G | $2.9\times10^{-1}$ | **0** | $5.2\times10^{5}$ | $7.0\times10^{-1}$ | – | – | $9.6\times10^{-1}$ | N/A | $4.1\times10^{-1}$ | $1.3\times10^{2}$ | $8.3\times10^{-1}$ |
| F388V | 1.2 | $4.4\times10^{-1}$ | $1.4\times10^{5}$ | **$4.0\times10^{-2}$** | 3.6 | **$1.0\times10^{-3}$** | N/A | 1.1 | $1.7\times10^{-1}$ | $1.5\times10^{2}$ | $8.9\times10^{-1}$ |
| G389A | 1.3 | $3.4\times10^{-1}$ | $6.4\times10^{5}$ | $9.0\times10^{-1}$ | $7.0\times10^{-1}$ | $1.9\times10^{-1}$ | N/A | 1.0 | $6.3\times10^{-1}$ | $1.1\times10^{2}$ | $5.3\times10^{-1}$ |
| G389V | $8.1\times10^{-1}$ | $3.5\times10^{-1}$ | $4.1\times10^{5}$ | $2.9\times10^{-1}$ | 1.5 | $2.7\times10^{-1}$ | N/A | 1.1 | $2.5\times10^{-1}$ | $1.4\times10^{2}$ | $9.6\times10^{-1}$ |
| A390G | 1.1 | $7.7\times10^{-1}$ | $5.1\times10^{5}$ | $6.8\times10^{-1}$ | 1.1 | $7.0\times10^{-1}$ | $9.5\times10^{-1}$ | N/A | $2.3\times10^{-1}$ | $1.0\times10^{2}$ | $4.5\times10^{-1}$ |
| A390V | 1.5 | $9.6\times10^{-2}$ | $2.7\times10^{5}$ | $9.2\times10^{-2}$ | 2.0 | $7.7\times10^{-2}$ | N/A | 1.2 | $1.6\times10^{-2}$ | $1.6\times10^{2}$ | $6.8\times10^{-1}$ |
| D391G | $8.6\times10^{-1}$ | $5.5\times10^{-1}$ | $4.9\times10^{5}$ | $5.6\times10^{-1}$ | 1.1 | $8.0\times10^{-1}$ | $8.1\times10^{-1}$ | N/A | $3.0\times10^{-2}$ | $1.5\times10^{2}$ | $9.3\times10^{-1}$ |
| D391V | $9.3\times10^{-1}$ | $7.3\times10^{-1}$ | $2.7\times10^{5}$ | $5.4\times10^{-2}$ | 1.5 | $1.6\times10^{-1}$ | $9.9\times10^{-1}$ | N/A | $7.3\times10^{-1}$ | $1.6\times10^{2}$ | $7.2\times10^{-1}$ |
| N392G | 1.0 | $9.0\times10^{-1}$ | $1.7\times10^{6}$ | **$4.0\times10^{-3}$** | $2.4\times10^{-1}$ | **$2.2\times10^{-4}$** | N/A | 1.1 | $1.6\times10^{-1}$ | $1.0\times10^{2}$ | $4.7\times10^{-1}$ |
| N392V | $3.7\times10^{-1}$ | **0** | $7.3\times10^{5}$ | $4.4\times10^{-1}$ | 1.1 | $8.5\times10^{-1}$ | N/A | 1.0 | $4.5\times10^{-1}$ | $1.4\times10^{2}$ | $9.4\times10^{-1}$ |
| I393G | 1.0 | $8.4\times10^{-1}$ | $3.1\times10^{5}$ | $1.9\times10^{-1}$ | 1.4 | $2.7\times10^{-1}$ | $8.9\times10^{-1}$ | N/A | $6.7\times10^{-2}$ | $7.6\times10^{1}$ | $2.7\times10^{-1}$ |
| I394G | ≤ $2.4\times10^{-1}$ | **0** | ≤ $3.4\times10^{5}$ | $2.3\times10^{-1}$ | ≤ 4.7 | **$2.1\times10^{-3}$** | $9.2\times10^{-1}$ | N/A | $1.6\times10^{-1}$ | ≥ $1.5\times10^{2}$ | $8.0\times10^{-1}$ |
| I394V | 1.4 | $8.1\times10^{-2}$ | $3.2\times10^{5}$ | $6.6\times10^{-2}$ | 1.6 | $1.0\times10^{-1}$ | N/A | 1.1 | $7.5\times10^{-2}$ | $1.5\times10^{2}$ | $8.1\times10^{-1}$ |
| A395G | 1.0 | $9.8\times10^{-1}$ | $9.5\times10^{5}$ | $1.2\times10^{-1}$ | $5.6\times10^{-1}$ | $1.4\times10^{-1}$ | $9.4\times10^{-1}$ | N/A | $2.3\times10^{-1}$ | $8.2\times10^{1}$ | $3.2\times10^{-1}$ |
| A395V | 1.9 | **$6.0\times10^{-3}$** | $2.0\times10^{5}$ | **$4.3\times10^{-2}$** | 3.1 | **$2.0\times10^{-3}$** | N/A | 1.1 | $2.5\times10^{-2}$ | $1.4\times10^{2}$ | $9.4\times10^{-1}$ |
| A396G | $8.7\times10^{-1}$ | $7.1\times10^{-1}$ | $5.6\times10^{5}$ | $8.4\times10^{-1}$ | $5.6\times10^{-1}$ | $6.1\times10^{-2}$ | $7.9\times10^{-1}$ | N/A | **$8.2\times10^{-3}$** | $1.1\times10^{2}$ | $5.3\times10^{-1}$ |
| A396V | $8.8\times10^{-1}$ | $5.8\times10^{-1}$ | $5.3\times10^{5}$ | $6.3\times10^{-1}$ | $8.1\times10^{-1}$ | $4.9\times10^{-1}$ | $9.5\times10^{-1}$ | N/A | $3.8\times10^{-1}$ | $1.4\times10^{2}$ | $9.5\times10^{-1}$ |
| A397G | $8.5\times10^{-1}$ | $7.1\times10^{-1}$ | $6.0\times10^{5}$ | $9.8\times10^{-1}$ | 1.1 | $6.8\times10^{-1}$ | $9.4\times10^{-1}$ | N/A | $3.5\times10^{-1}$ | $8.0\times10^{1}$ | $4.1\times10^{-1}$ |
| A397V | 1.6 | $2.6\times10^{-2}$ | $8.5\times10^{5}$ | $1.6\times10^{-1}$ | $7.4\times10^{-1}$ | $1.8\times10^{-1}$ | $9.1\times10^{-1}$ | N/A | $2.5\times10^{-2}$ | $1.6\times10^{2}$ | $6.2\times10^{-1}$ |
| M398G | $7.5\times10^{-1}$ | $3.0\times10^{-1}$ | $3.4\times10^{5}$ | $2.0\times10^{-1}$ | $7.2\times10^{-1}$ | $3.4\times10^{-1}$ | N/A | 1.1 | $6.0\times10^{-2}$ | $6.2\times10^{1}$ | $1.4\times10^{-1}$ |
| M398V | $8.3\times10^{-1}$ | $4.9\times10^{-1}$ | $4.2\times10^{5}$ | $3.1\times10^{-1}$ | 1.3 | $4.4\times10^{-1}$ | N/A | 1.0 | $8.2\times10^{-1}$ | $1.3\times10^{2}$ | $7.5\times10^{-1}$ |
| N399G | $3.4\times10^{-1}$ | **0** | $6.5\times10^{5}$ | $8.4\times10^{-1}$ | $8.0\times10^{-1}$ | $5.0\times10^{-1}$ | $9.4\times10^{-1}$ | N/A | $2.0\times10^{-1}$ | $4.5\times10^{1}$ | $8.4\times10^{-2}$ |
| N399V | $6.2\times10^{-1}$ | $7.8\times10^{-2}$ | $3.2\times10^{4}$ | **$1.2\times10^{-2}$** | – | – | $9.3\times10^{-1}$ | N/A | $1.4\times10^{-1}$ | $7.6\times10^{1}$ | $2.5\times10^{-1}$ |
| Y400G | ≤ 2.4 | $1.1\times10^{-2}$ | ≤ $1.8\times10^{5}$ | $9.9\times10^{-2}$ | ≤ 1.2 | $7.4\times10^{-1}$ | N/A | 1.2 | **$8.1\times10^{-3}$** | ≥ $1.1\times10^{2}$ | $6.5\times10^{-1}$ |
| Y400V | 1.6 | $3.4\times10^{-2}$ | $1.3\times10^{5}$ | **$2.3\times10^{-2}$** | 1.2 | $5.7\times10^{-1}$ | N/A | 1.3 | **$9.0\times10^{-4}$** | $1.4\times10^{2}$ | $9.2\times10^{-1}$ |
| Q401G | $7.5\times10^{-1}$ | $2.8\times10^{-1}$ | $5.7\times10^{5}$ | $8.8\times10^{-1}$ | $7.5\times10^{-1}$ | $3.3\times10^{-1}$ | $9.1\times10^{-1}$ | N/A | $8.2\times10^{-2}$ | $4.5\times10^{1}$ | $8.0\times10^{-2}$ |
| Q401V | $5.9\times10^{-1}$ | $2.9\times10^{-2}$ | $2.0\times10^{5}$ | $7.0\times10^{-2}$ | – | – | N/A | 1.1 | $1.4\times10^{-2}$ | $1.1\times10^{2}$ | $5.5\times10^{-1}$ |

| Variant | $f_a$ | $f_a$ $p$–value | $(k_{cat}/K_M)^{MeP}_{chem.}$ ($M^{-1}s^{-1}$) | $(k_{cat}/K_M)^{MeP}_{chem.}$ $p$–value | FC1 | FC1 $p$–value | FC2 | FC3 | FC2/3 $p$–value | FC4 ($s^{-1}$) | FC4 $p$–value |
|---|---|---|---|---|---|---|---|---|---|---|---|
| V402A | 1.6 | $4.0\times10^{-2}$ | $1.4\times10^{5}$ | **$2.5\times10^{-2}$** | 2.4 | **$7.9\times10^{-3}$** | N/A | 1.0 | $4.4\times10^{-1}$ | $1.4\times10^{2}$ | $9.6\times10^{-1}$ |
| V402G | $6.0\times10^{-1}$ | $3.5\times10^{-2}$ | $2.8\times10^{5}$ | $1.1\times10^{-1}$ | 2.0 | $2.5\times10^{-2}$ | $8.9\times10^{-1}$ | N/A | $1.3\times10^{-2}$ | $9.3\times10^{1}$ | $3.4\times10^{-1}$ |
| Y403G | $3.0\times10^{-1}$ | **$1.0\times10^{-3}$** | $7.9\times10^{5}$ | $4.3\times10^{-1}$ | – | – | $7.0\times10^{-1}$ | N/A | **$1.\times10^{-4}$** | $5.1\times10^{1}$ | $1.5\times10^{-1}$ |
| Y403V | $9.5\times10^{-1}$ | $8.2\times10^{-1}$ | $1.7\times10^{5}$ | **$3.0\times10^{-2}$** | 2.6 | **$4.5\times10^{-3}$** | N/A | 1.1 | $1.4\times10^{-1}$ | $1.2\times10^{2}$ | $7.2\times10^{-1}$ |
| F404G | $5.1\times10^{-1}$ | $3.2\times10^{-2}$ | $2.1\times10^{5}$ | $1.0\times10^{-1}$ | 1.4 | $3.2\times10^{-1}$ | $8.6\times10^{-1}$ | N/A | $8.2\times10^{-2}$ | $1.2\times10^{2}$ | $8.0\times10^{-1}$ |
| F404V | 1.6 | $3.6\times10^{-2}$ | $1.5\times10^{5}$ | **$3.4\times10^{-2}$** | 3.0 | **$2.2\times10^{-3}$** | N/A | 1.2 | $1.4\times10^{-2}$ | $1.3\times10^{2}$ | $8.6\times10^{-1}$ |
| D405G | $5.7\times10^{-1}$ | $8.7\times10^{-2}$ | $1.6\times10^{5}$ | $1.1\times10^{-1}$ | 3.8 | **$5.4\times10^{-3}$** | $7.8\times10^{-1}$ | N/A | **$2.8\times10^{-3}$** | $9.1\times10^{1}$ | $4.3\times10^{-1}$ |
| D405V | $8.7\times10^{-1}$ | $5.6\times10^{-1}$ | $1.7\times10^{4}$ | **$8.0\times10^{-3}$** | $1.2\times10^{1}$ | **$3.0\times10^{-4}$** | N/A | 1.0 | $6.0\times10^{-1}$ | $1.3\times10^{2}$ | $7.9\times10^{-1}$ |
| R406G | $9.7\times10^{-1}$ | $8.7\times10^{-1}$ | $2.8\times10^{5}$ | $2.3\times10^{-1}$ | 2.0 | $6.3\times10^{-2}$ | N/A | 1.2 | $2.2\times10^{-2}$ | $1.5\times10^{2}$ | $8.9\times10^{-1}$ |
| R406V | 1.5 | $9.4\times10^{-2}$ | $2.8\times10^{5}$ | $6.9\times10^{-2}$ | 1.6 | $9.3\times10^{-2}$ | N/A | 1.2 | $2.2\times10^{-2}$ | $2.1\times10^{2}$ | $1.2\times10^{-1}$ |
| K407G | 1.1 | $6.3\times10^{-1}$ | $7.5\times10^{5}$ | $5.2\times10^{-1}$ | $8.5\times10^{-1}$ | $6.6\times10^{-1}$ | $9.4\times10^{-1}$ | N/A | $1.9\times10^{-1}$ | $1.3\times10^{2}$ | $8.7\times10^{-1}$ |
| K407V | 1.3 | $2.0\times10^{-1}$ | $3.3\times10^{5}$ | $9.8\times10^{-2}$ | 1.2 | $4.9\times10^{-1}$ | N/A | 1.1 | $5.6\times10^{-2}$ | $1.6\times10^{2}$ | $5.8\times10^{-1}$ |
| V408A | 1.5 | $1.2\times10^{-1}$ | $5.9\times10^{5}$ | $9.2\times10^{-1}$ | $9.5\times10^{-1}$ | $8.6\times10^{-1}$ | N/A | 1.1 | $1.1\times10^{-1}$ | $1.7\times10^{2}$ | $5.6\times10^{-1}$ |
| V408G | $9.8\times10^{-1}$ | $8.6\times10^{-1}$ | $2.0\times10^{5}$ | $7.9\times10^{-2}$ | 2.6 | $1.9\times10^{-2}$ | N/A | 1.1 | $8.7\times10^{-2}$ | $1.1\times10^{2}$ | $4.4\times10^{-1}$ |
| L409G | $2.8\times10^{-1}$ | **0** | $3.3\times10^{5}$ | $1.7\times10^{-1}$ | 2.1 | $1.3\times10^{-2}$ | $9.6\times10^{-1}$ | N/A | $4.2\times10^{-1}$ | $9.9\times10^{1}$ | $3.6\times10^{-1}$ |
| L409V | $9.3\times10^{-1}$ | $7.4\times10^{-1}$ | $1.4\times10^{5}$ | $5.6\times10^{-2}$ | 2.7 | $2.0\times10^{-2}$ | N/A | 1.2 | $1.2\times10^{-2}$ | $1.6\times10^{2}$ | $7.3\times10^{-1}$ |
| A410G | 1.3 | $4.0\times10^{-1}$ | $6.5\times10^{5}$ | $8.6\times10^{-1}$ | $7.4\times10^{-1}$ | $2.3\times10^{-1}$ | $9.8\times10^{-1}$ | N/A | $6.4\times10^{-1}$ | $1.2\times10^{2}$ | $6.9\times10^{-1}$ |
| A410V | 1.2 | $4.4\times10^{-1}$ | $3.0\times10^{5}$ | $1.2\times10^{-1}$ | 1.4 | $2.4\times10^{-1}$ | N/A | 1.1 | $2.7\times10^{-1}$ | $1.3\times10^{2}$ | $7.9\times10^{-1}$ |
| D411G | 1.0 | $9.4\times10^{-1}$ | $4.2\times10^{5}$ | $3.5\times10^{-1}$ | 1.2 | $5.0\times10^{-1}$ | N/A | 1.1 | $2.5\times10^{-1}$ | $1.4\times10^{2}$ | $9.9\times10^{-1}$ |
| D411V | 1.2 | $1.9\times10^{-1}$ | $2.7\times10^{5}$ | **$4.0\times10^{-3}$** | 1.5 | $1.1\times10^{-1}$ | N/A | 1.0 | $2.4\times10^{-1}$ | $1.6\times10^{2}$ | $5.9\times10^{-1}$ |
| S412G | $9.4\times10^{-1}$ | $8.1\times10^{-1}$ | $3.2\times10^{5}$ | $1.4\times10^{-1}$ | 1.6 | $1.2\times10^{-1}$ | $9.5\times10^{-1}$ | N/A | $2.4\times10^{-1}$ | $1.1\times10^{2}$ | $5.3\times10^{-1}$ |
| S412V | 1.3 | $1.7\times10^{-1}$ | $4.1\times10^{5}$ | $2.2\times10^{-1}$ | 1.2 | $5.0\times10^{-1}$ | N/A | 1.1 | $6.9\times10^{-2}$ | $2.1\times10^{2}$ | $9.8\times10^{-2}$ |
| K413G | 1.1 | $5.7\times10^{-1}$ | $3.5\times10^{5}$ | $2.3\times10^{-1}$ | 2.3 | $1.8\times10^{-2}$ | $9.4\times10^{-1}$ | N/A | $1.4\times10^{-1}$ | $1.2\times10^{2}$ | $7.0\times10^{-1}$ |
| K413V | 1.5 | $7.2\times10^{-2}$ | $2.0\times10^{5}$ | **$4.0\times10^{-2}$** | 1.9 | $2.7\times10^{-2}$ | N/A | 1.1 | $2.1\times10^{-1}$ | $1.6\times10^{2}$ | $7.3\times10^{-1}$ |
| L414G | $3.8\times10^{-1}$ | **$3.0\times10^{-3}$** | $2.4\times10^{5}$ | $1.1\times10^{-1}$ | 2.7 | **$5.8\times10^{-3}$** | $9.8\times10^{-1}$ | N/A | $5.3\times10^{-1}$ | $9.8\times10^{1}$ | $4.2\times10^{-1}$ |
| L414V | $6.5\times10^{-1}$ | $8.8\times10^{-2}$ | $4.2\times10^{5}$ | $3.0\times10^{-1}$ | 1.6 | $9.3\times10^{-2}$ | N/A | 1.0 | $5.5\times10^{-1}$ | $1.7\times10^{2}$ | $5.5\times10^{-1}$ |
| E415G | $4.9\times10^{-1}$ | $1.3\times10^{-2}$ | $4.1\times10^{5}$ | $3.2\times10^{-1}$ | 3.2 | **$8.7\times10^{-4}$** | $9.6\times10^{-1}$ | N/A | $3.6\times10^{-1}$ | $2.1\times10^{2}$ | $2.2\times10^{-1}$ |
| E415V | $9.2\times10^{-1}$ | $7.5\times10^{-1}$ | $6.7\times10^{5}$ | $7.3\times10^{-1}$ | $6.0\times10^{-1}$ | $9.5\times10^{-2}$ | N/A | 1.0 | $5.8\times10^{-1}$ | $1.5\times10^{2}$ | $7.6\times10^{-1}$ |
| L416G | $2.7\times10^{-1}$ | **$1.0\times10^{-3}$** | $2.1\times10^{6}$ | **$9.0\times10^{-3}$** | – | – | N/A | 1.1 | $1.8\times10^{-1}$ | $1.3\times10^{2}$ | $9.1\times10^{-1}$ |
| L416V | $3.6\times10^{-1}$ | **$1.0\times10^{-3}$** | $2.8\times10^{5}$ | $7.7\times10^{-2}$ | 2.8 | **$3.5\times10^{-3}$** | N/A | 1.1 | $1.1\times10^{-1}$ | $1.6\times10^{2}$ | $7.0\times10^{-1}$ |
| D417G | $8.1\times10^{-1}$ | $3.8\times10^{-1}$ | $3.9\times10^{5}$ | $2.7\times10^{-1}$ | 1.5 | $1.6\times10^{-1}$ | N/A | 1.0 | $5.8\times10^{-1}$ | $9.9\times10^{1}$ | $4.3\times10^{-1}$ |
| D417V | $8.8\times10^{-1}$ | $6.0\times10^{-1}$ | $2.8\times10^{5}$ | $8.7\times10^{-2}$ | 1.7 | $1.2\times10^{-1}$ | N/A | 1.2 | $2.1\times10^{-2}$ | $1.6\times10^{2}$ | $6.3\times10^{-1}$ |
| D418G | $9.2\times10^{-1}$ | $7.0\times10^{-1}$ | $2.8\times10^{5}$ | $1.5\times10^{-1}$ | 2.5 | **$3.9\times10^{-3}$** | N/A | 1.1 | $1.6\times10^{-1}$ | $1.1\times10^{2}$ | $6.5\times10^{-1}$ |
| D418V | 1.6 | $8.9\times10^{-2}$ | $2.0\times10^{5}$ | $5.9\times10^{-2}$ | 2.7 | **$3.9\times10^{-3}$** | N/A | 1.1 | $1.2\times10^{-1}$ | $1.6\times10^{2}$ | $7.6\times10^{-1}$ |
| V419A | 1.1 | $7.5\times10^{-1}$ | $3.1\times10^{5}$ | $1.1\times10^{-1}$ | 1.6 | $1.3\times10^{-1}$ | N/A | 1.1 | $1.6\times10^{-1}$ | $1.5\times10^{2}$ | $8.2\times10^{-1}$ |
| V419G | $3.7\times10^{-1}$ | **$4.0\times10^{-3}$** | $2.0\times10^{5}$ | $1.0\times10^{-1}$ | 3.3 | **$2.7\times10^{-3}$** | $7.9\times10^{-1}$ | N/A | **$8.5\times10^{-3}$** | $8.3\times10^{1}$ | $3.3\times10^{-1}$ |
| R420G | $3.3\times10^{-1}$ | **0** | $3.0\times10^{5}$ | $1.9\times10^{-1}$ | 1.9 | $5.8\times10^{-2}$ | $9.7\times10^{-1}$ | N/A | $4.1\times10^{-1}$ | $9.1\times10^{1}$ | $3.2\times10^{-1}$ |
| R420V | $6.5\times10^{-1}$ | $4.4\times10^{-2}$ | $3.4\times10^{5}$ | $1.5\times10^{-1}$ | – | – | N/A | 1.2 | **$1.0\times10^{-3}$** | $1.6\times10^{2}$ | $6.1\times10^{-1}$ |
| D421G | $9.6\times10^{-1}$ | $8.5\times10^{-1}$ | $3.4\times10^{5}$ | $2.5\times10^{-1}$ | 1.8 | $5.5\times10^{-2}$ | N/A | 1.0 | $7.9\times10^{-1}$ | $1.5\times10^{2}$ | $8.7\times10^{-1}$ |
| D421V | $6.8\times10^{-1}$ | $1.1\times10^{-1}$ | $3.1\times10^{5}$ | $1.2\times10^{-1}$ | 1.6 | $1.4\times10^{-1}$ | $9.7\times10^{-1}$ | N/A | $5.1\times10^{-1}$ | $1.4\times10^{2}$ | $9.1\times10^{-1}$ |
| Y422G | $6.2\times10^{-1}$ | $6.9\times10^{-2}$ | $2.0\times10^{5}$ | $8.7\times10^{-2}$ | 3.1 | **$2.1\times10^{-3}$** | $9.7\times10^{-1}$ | N/A | $3.3\times10^{-1}$ | $7.0\times10^{1}$ | $2.0\times10^{-1}$ |
| Y422V | 1.1 | $7.1\times10^{-1}$ | $4.5\times10^{5}$ | $4.6\times10^{-1}$ | 1.4 | $2.7\times10^{-1}$ | N/A | 1.1 | $3.8\times10^{-1}$ | $1.7\times10^{2}$ | $5.8\times10^{-1}$ |
| V423A | 1.1 | $5.9\times10^{-1}$ | $3.4\times10^{5}$ | $1.2\times10^{-1}$ | 1.3 | $4.9\times10^{-1}$ | N/A | 1.0 | $4.6\times10^{-1}$ | $1.4\times10^{2}$ | $9.9\times10^{-1}$ |
| V423G | $4.5\times10^{-1}$ | **$2.0\times10^{-3}$** | $3.3\times10^{5}$ | $1.9\times10^{-1}$ | 2.3 | $2.4\times10^{-2}$ | $9.3\times10^{-1}$ | N/A | $1.9\times10^{-1}$ | $1.0\times10^{2}$ | $5.1\times10^{-1}$ |
| M424G | $3.1\times10^{-1}$ | **0** | $8.2\times10^{5}$ | $3.0\times10^{-1}$ | $8.5\times10^{-1}$ | $6.4\times10^{-1}$ | $7.3\times10^{-1}$ | N/A | **$2.0\times10^{-4}$** | $8.6\times10^{1}$ | $3.2\times10^{-1}$ |
| M424V | $8.1\times10^{-1}$ | $4.1\times10^{-1}$ | $9.7\times10^{5}$ | $5.9\times10^{-2}$ | $5.9\times10^{-1}$ | $9.7\times10^{-2}$ | $9.8\times10^{-1}$ | N/A | $5.5\times10^{-1}$ | $1.7\times10^{2}$ | $5.4\times10^{-1}$ |
| T425V | $9.2\times10^{-1}$ | $7.4\times10^{-1}$ | $5.9\times10^{5}$ | $9.0\times10^{-1}$ | $8.2\times10^{-1}$ | $5.9\times10^{-1}$ | N/A | 1.0 | $5.4\times10^{-1}$ | $2.0\times10^{2}$ | $2.3\times10^{-1}$ |
| E426G | $8.9\times10^{-1}$ | $6.3\times10^{-1}$ | $2.4\times10^{5}$ | $8.8\times10^{-2}$ | 1.5 | $3.1\times10^{-1}$ | $8.7\times10^{-1}$ | N/A | **$6.2\times10^{-3}$** | $1.3\times10^{2}$ | $7.6\times10^{-1}$ |
| E426V | 1.3 | $2.9\times10^{-1}$ | $2.3\times10^{5}$ | $5.3\times10^{-2}$ | 1.6 | $1.8\times10^{-1}$ | N/A | 1.1 | $3.2\times10^{-1}$ | $1.5\times10^{2}$ | $8.8\times10^{-1}$ |
| L427G | $\leq 1.7\times10^{-1}$ | **0** | $\leq 5.3\times10^{5}$ | $7.1\times10^{-1}$ | $\leq 1.6$ | $1.6\times10^{-1}$ | $8.2\times10^{-1}$ | N/A | **$2.4\times10^{-3}$** | $\geq 8.9\times10^{1}$ | $3.2\times10^{-1}$ |
| L427V | $9.8\times10^{-1}$ | $9.1\times10^{-1}$ | $2.2\times10^{5}$ | **$3.8\times10^{-2}$** | 1.8 | $9.1\times10^{-2}$ | N/A | 1.1 | $1.5\times10^{-1}$ | $1.4\times10^{2}$ | $9.8\times10^{-1}$ |
| K428G | $4.0\times10^{-1}$ | **$5.0\times10^{-3}$** | $6.4\times10^{5}$ | $8.8\times10^{-1}$ | 1.8 | $8.0\times10^{-2}$ | N/A | 1.0 | $6.6\times10^{-1}$ | $2.0\times10^{2}$ | $2.7\times10^{-1}$ |
| K428V | $6.2\times10^{-1}$ | $2.1\times10^{-2}$ | $2.4\times10^{5}$ | **$3.2\times10^{-2}$** | 2.6 | **$9.0\times10^{-4}$** | $9.2\times10^{-1}$ | N/A | $3.3\times10^{-2}$ | $1.4\times10^{2}$ | $9.6\times10^{-1}$ |
| K429G | 1.0 | 1.0 | $4.3\times10^{5}$ | $3.7\times10^{-1}$ | 1.3 | $3.4\times10^{-1}$ | $9.2\times10^{-1}$ | N/A | $1.2\times10^{-1}$ | $1.6\times10^{2}$ | $7.0\times10^{-1}$ |
| K429V | 1.4 | $9.4\times10^{-2}$ | $1.5\times10^{5}$ | **$1.5\times10^{-2}$** | 3.0 | **$2.6\times10^{-3}$** | N/A | 1.2 | $2.7\times10^{-2}$ | $1.5\times10^{2}$ | $8.3\times10^{-1}$ |
| E430G | $6.6\times10^{-1}$ | $3.7\times10^{-2}$ | $1.8\times10^{5}$ | **$4.5\times10^{-2}$** | 2.1 | $1.2\times10^{-2}$ | N/A | 1.3 | **0** | $1.3\times10^{2}$ | $6.9\times10^{-1}$ |
| E430V | $9.6\times10^{-1}$ | $8.3\times10^{-1}$ | $4.5\times10^{5}$ | $4.6\times10^{-1}$ | 1.1 | $7.3\times10^{-1}$ | N/A | 1.3 | **$5.0\times10^{-4}$** | $1.4\times10^{2}$ | $9.9\times10^{-1}$ |
| P431G | $7.8\times10^{-1}$ | $3.5\times10^{-1}$ | $2.4\times10^{5}$ | $7.4\times10^{-2}$ | 2.0 | $3.8\times10^{-2}$ | N/A | 1.0 | $4.9\times10^{-1}$ | $1.0\times10^{2}$ | $4.7\times10^{-1}$ |
| P431V | 1.5 | $7.7\times10^{-2}$ | $1.3\times10^{5}$ | **$4.3\times10^{-2}$** | 3.8 | **$2.9\times10^{-3}$** | N/A | 1.1 | $7.2\times10^{-2}$ | $1.5\times10^{2}$ | $8.2\times10^{-1}$ |
| S432G | $8.2\times10^{-1}$ | $4.1\times10^{-1}$ | $5.7\times10^{5}$ | $8.5\times10^{-1}$ | $6.5\times10^{-1}$ | $1.4\times10^{-1}$ | N/A | 1.2 | $1.0\times10^{-2}$ | $1.3\times10^{2}$ | $8.3\times10^{-1}$ |
| S432V | $2.6\times10^{-1}$ | **0** | $6.1\times10^{5}$ | $9.7\times10^{-1}$ | – | – | N/A | 1.1 | **$4.3\times10^{-3}$** | $1.6\times10^{2}$ | $7.3\times10^{-1}$ |
| V433A | $7.5\times10^{-1}$ | $2.7\times10^{-1}$ | $4.1\times10^{5}$ | $3.6\times10^{-1}$ | 1.1 | $7.1\times10^{-1}$ | N/A | 1.2 | **$4.6\times10^{-3}$** | $1.7\times10^{2}$ | $4.9\times10^{-1}$ |
| V433G | $1.1\times10^{-1}$ | **0** | $3.1\times10^{5}$ | $9.0\times10^{-2}$ | 1.3 | $4.4\times10^{-1}$ | N/A | 1.2 | **$4.0\times10^{-4}$** | $1.3\times10^{2}$ | $8.9\times10^{-1}$ |
| L434G | $2.5\times10^{-1}$ | **0** | $1.4\times10^{5}$ | **$2.5\times10^{-2}$** | 4.1 | **$3.1\times10^{-4}$** | $5.5\times10^{-1}$ | N/A | **0** | $6.2\times10^{1}$ | $8.3\times10^{-2}$ |
| L434V | $5.0\times10^{-1}$ | **$3.0\times10^{-3}$** | $3.2\times10^{4}$ | **$5.0\times10^{-3}$** | $1.1\times10^{1}$ | **$6.0\times10^{-5}$** | $9.7\times10^{-1}$ | N/A | $3.3\times10^{-1}$ | $1.1\times10^{2}$ | $5.4\times10^{-1}$ |
| Y435G | $3.8\times10^{-2}$ | **0** | $1.3\times10^{5}$ | $9.0\times10^{-2}$ | – | – | $5.6\times10^{-1}$ | N/A | **0** | $1.3\times10^{2}$ | $9.0\times10^{-1}$ |
| Y435V | $1.4\times10^{-1}$ | **0** | $1.1\times10^{5}$ | **$3.0\times10^{-2}$** | 3.9 | **$7.0\times10^{-3}$** | $8.0\times10^{-1}$ | N/A | **$3.0\times10^{-4}$** | $1.2\times10^{2}$ | $6.6\times10^{-1}$ |
| V436A | $9.3\times10^{-1}$ | $7.4\times10^{-1}$ | $4.7\times10^{5}$ | $4.1\times10^{-1}$ | 1.0 | $9.8\times10^{-1}$ | N/A | 1.1 | $1.5\times10^{-1}$ | $1.8\times10^{2}$ | $3.8\times10^{-1}$ |
| V436G | $3.2\times10^{-1}$ | **0** | $3.4\times10^{5}$ | $2.4\times10^{-1}$ | 2.1 | $5.0\times10^{-2}$ | $7.7\times10^{-1}$ | N/A | **$1.2\times10^{-3}$** | $8.0\times10^{1}$ | $2.6\times10^{-1}$ |
| L437G | $2.9\times10^{-1}$ | **0** | $3.6\times10^{5}$ | $2.0\times10^{-1}$ | 1.8 | $1.1\times10^{-1}$ | N/A | 1.0 | $6.6\times10^{-1}$ | $8.6\times10^{1}$ | $2.5\times10^{-1}$ |
| L437V | 1.5 | $6.8\times10^{-2}$ | $6.0\times10^{5}$ | $9.2\times10^{-1}$ | $7.0\times10^{-1}$ | $1.6\times10^{-1}$ | N/A | 1.1 | $4.7\times10^{-2}$ | $1.7\times10^{2}$ | $4.2\times10^{-1}$ |
| S438G | $5.3\times10^{-1}$ | $1.8\times10^{-2}$ | $4.4\times10^{5}$ | $3.9\times10^{-1}$ | 1.6 | $1.3\times10^{-1}$ | N/A | 1.0 | $2.3\times10^{-1}$ | $1.0\times10^{2}$ | $4.4\times10^{-1}$ |
| S438V | $7.7\times10^{-1}$ | $2.5\times10^{-1}$ | $2.3\times10^{5}$ | $6.4\times10^{-2}$ | 1.7 | $8.1\times10^{-2}$ | N/A | 1.0 | $5.3\times10^{-1}$ | $1.4\times10^{2}$ | $9.1\times10^{-1}$ |
| T439G | $2.6\times10^{-1}$ | **0** | $3.6\times10^{5}$ | $2.0\times10^{-1}$ | $8.9\times10^{-1}$ | $7.7\times10^{-1}$ | $8.3\times10^{-1}$ | N/A | **$1.4\times10^{-3}$** | $8.4\times10^{1}$ | $2.3\times10^{-1}$ |
| T439V | 1.4 | $1.8\times10^{-1}$ | $1.9\times10^{5}$ | $8.3\times10^{-2}$ | 1.7 | $1.1\times10^{-1}$ | N/A | 1.2 | **$2.9\times10^{-3}$** | $1.3\times10^{2}$ | $8.4\times10^{-1}$ |

| Variant | $f_a$ | $f_a$ $p$–value | $(k_{cat}/K_M)^{MeP}_{chem.}$ ($M^{-1}s^{-1}$) | $(k_{cat}/K_M)^{MeP}_{chem.}$ $p$–value | FC1 | FC1 $p$–value | FC2 | FC3 | FC2/3 $p$–value | FC4 ($s^{-1}$) | FC4 $p$–value |
|---|---|---|---|---|---|---|---|---|---|---|---|
| D440G | 1.1 | $5.4\times10^{-1}$ | $2.7\times10^{5}$ | $1.2\times10^{-1}$ | 1.2 | $4.4\times10^{-1}$ | $8.8\times10^{-1}$ | N/A | $1.7\times10^{-2}$ | $1.1\times10^{2}$ | $5.1\times10^{-1}$ |
| D440V | $2.1\times10^{-1}$ | **0** | $5.6\times10^{5}$ | $8.2\times10^{-1}$ | 1.6 | $3.3\times10^{-1}$ | N/A | 1.0 | $8.3\times10^{-1}$ | $1.2\times10^{2}$ | $6.9\times10^{-1}$ |
| E441G | $8.7\times10^{-1}$ | $5.3\times10^{-1}$ | $3.9\times10^{5}$ | $2.4\times10^{-1}$ | 1.7 | $4.8\times10^{-2}$ | $8.6\times10^{-1}$ | N/A | $2.5\times10^{-2}$ | $1.1\times10^{2}$ | $5.6\times10^{-1}$ |
| E441V | $9.6\times10^{-1}$ | $6.5\times10^{-1}$ | $4.2\times10^{5}$ | $1.8\times10^{-1}$ | $9.9\times10^{-1}$ | $9.5\times10^{-1}$ | N/A | 1.1 | $2.6\times10^{-2}$ | $1.4\times10^{2}$ | $8.8\times10^{-1}$ |
| I442G | $1.1\times10^{-1}$ | **0** | $4.1\times10^{5}$ | $3.1\times10^{-1}$ | – | – | $7.7\times10^{-1}$ | N/A | **0** | $6.8\times10^{1}$ | $1.3\times10^{-1}$ |
| I442V | $8.0\times10^{-1}$ | $3.4\times10^{-1}$ | $4.1\times10^{5}$ | $2.7\times10^{-1}$ | – | – | N/A | 1.1 | $4.6\times10^{-1}$ | $1.5\times10^{2}$ | $8.7\times10^{-1}$ |
| W443G | 1.0 | $9.1\times10^{-1}$ | $4.9\times10^{5}$ | $6.0\times10^{-1}$ | 1.5 | $1.3\times10^{-1}$ | N/A | 1.1 | $1.8\times10^{-1}$ | $1.4\times10^{2}$ | $9.7\times10^{-1}$ |
| W443V | 1.4 | $1.5\times10^{-1}$ | $2.4\times10^{5}$ | $6.4\times10^{-2}$ | 1.7 | $1.1\times10^{-1}$ | N/A | 1.1 | $1.3\times10^{-1}$ | $1.8\times10^{2}$ | $4.6\times10^{-1}$ |
| E444G | 1.0 | $9.2\times10^{-1}$ | $4.2\times10^{5}$ | $3.5\times10^{-1}$ | 1.7 | $1.3\times10^{-1}$ | N/A | 1.2 | **$7.0\times10^{-4}$** | $1.5\times10^{2}$ | $8.1\times10^{-1}$ |
| E444V | 1.0 | $7.8\times10^{-1}$ | $3.8\times10^{5}$ | $2.2\times10^{-1}$ | $7.3\times10^{-1}$ | $3.1\times10^{-1}$ | N/A | 1.2 | **$2.1\times10^{-3}$** | $1.7\times10^{2}$ | $5.8\times10^{-1}$ |
| S445G | $1.6\times10^{-1}$ | **0** | $6.1\times10^{5}$ | 1.0 | 1.4 | $2.9\times10^{-1}$ | N/A | 1.0 | $8.2\times10^{-1}$ | $1.3\times10^{2}$ | $8.3\times10^{-1}$ |
| S445V | 1.0 | $8.7\times10^{-1}$ | $3.4\times10^{5}$ | $4.0\times10^{-1}$ | 1.5 | $2.2\times10^{-1}$ | N/A | 1.0 | $4.5\times10^{-1}$ | $1.5\times10^{2}$ | $8.4\times10^{-1}$ |
| S446G | $3.6\times10^{-1}$ | **$2.0\times10^{-3}$** | $4.0\times10^{5}$ | $3.3\times10^{-1}$ | 1.2 | $6.5\times10^{-1}$ | $7.4\times10^{-1}$ | N/A | **0** | $1.1\times10^{2}$ | $5.1\times10^{-1}$ |
| S446V | 1.5 | $4.8\times10^{-2}$ | $2.9\times10^{5}$ | $8.5\times10^{-2}$ | 1.8 | $7.1\times10^{-2}$ | $9.7\times10^{-1}$ | N/A | $5.4\times10^{-1}$ | $1.5\times10^{2}$ | $7.5\times10^{-1}$ |
| I447G | $\leq 8.1\times10^{-3}$ | **0** | $\leq 1.9\times10^{5}$ | **$4.3\times10^{-2}$** | – | – | $5.6\times10^{-1}$ | N/A | **0** | $\geq 1.0\times10^{2}$ | $4.8\times10^{-1}$ |
| I447V | 1.1 | $7.8\times10^{-1}$ | $5.5\times10^{5}$ | $7.2\times10^{-1}$ | $7.4\times10^{-1}$ | $3.9\times10^{-1}$ | N/A | 1.1 | $1.4\times10^{-1}$ | $1.3\times10^{2}$ | $8.0\times10^{-1}$ |
| P448G | $5.4\times10^{-2}$ | **0** | $2.9\times10^{5}$ | $1.0\times10^{-1}$ | – | – | $7.9\times10^{-1}$ | N/A | **0** | $1.1\times10^{2}$ | $4.7\times10^{-1}$ |
| E449G | $7.5\times10^{-1}$ | $1.8\times10^{-1}$ | $4.4\times10^{5}$ | $3.1\times10^{-1}$ | 1.4 | $1.8\times10^{-1}$ | $6.3\times10^{-1}$ | N/A | **$1.\times10^{-4}$** | $1.1\times10^{2}$ | $5.0\times10^{-1}$ |
| E449V | $7.9\times10^{-1}$ | $3.5\times10^{-1}$ | $2.1\times10^{5}$ | $1.2\times10^{-1}$ | 3.2 | $1.5\times10^{-2}$ | $8.4\times10^{-1}$ | N/A | **$5.0\times10^{-3}$** | $1.2\times10^{2}$ | $6.4\times10^{-1}$ |
| P450G | 1.4 | $2.5\times10^{-1}$ | $2.7\times10^{5}$ | $1.3\times10^{-1}$ | 1.6 | $8.5\times10^{-2}$ | N/A | 1.0 | $4.7\times10^{-1}$ | $8.8\times10^{1}$ | $4.2\times10^{-1}$ |
| P450V | 1.3 | $3.4\times10^{-1}$ | $4.9\times10^{5}$ | $5.2\times10^{-1}$ | $8.7\times10^{-1}$ | $7.0\times10^{-1}$ | $9.1\times10^{-1}$ | N/A | $8.7\times10^{-2}$ | $1.4\times10^{2}$ | $9.9\times10^{-1}$ |
| I451G | $8.4\times10^{-2}$ | **0** | $1.3\times10^{5}$ | **$4.2\times10^{-2}$** | $1.0\times10^{1}$ | **$2.6\times10^{-3}$** | $4.6\times10^{-1}$ | N/A | **0** | $7.7\times10^{1}$ | $1.5\times10^{-1}$ |
| I451V | $7.1\times10^{-1}$ | $1.5\times10^{-1}$ | $6.6\times10^{5}$ | $7.7\times10^{-1}$ | $6.8\times10^{-1}$ | $2.2\times10^{-1}$ | $9.2\times10^{-1}$ | N/A | $5.6\times10^{-2}$ | $1.4\times10^{2}$ | $9.4\times10^{-1}$ |
| K452G | $1.4\times10^{-1}$ | **0** | $2.2\times10^{5}$ | $6.4\times10^{-2}$ | 2.3 | $2.6\times10^{-2}$ | $7.6\times10^{-1}$ | N/A | **0** | $1.1\times10^{2}$ | $6.0\times10^{-1}$ |
| K452V | $1.1\times10^{-1}$ | **0** | $2.3\times10^{5}$ | **$4.3\times10^{-2}$** | – | – | $8.2\times10^{-1}$ | N/A | **$6.6\times10^{-3}$** | $1.3\times10^{2}$ | $9.3\times10^{-1}$ |
| S453G | 1.0 | $9.5\times10^{-1}$ | $7.2\times10^{5}$ | $6.2\times10^{-1}$ | $6.8\times10^{-1}$ | $1.7\times10^{-1}$ | $7.2\times10^{-1}$ | N/A | **$1.\times10^{-4}$** | $1.0\times10^{2}$ | $3.5\times10^{-1}$ |
| S453V | 1.1 | $7.5\times10^{-1}$ | $1.3\times10^{5}$ | **$2.0\times10^{-2}$** | 4.1 | **$7.9\times10^{-4}$** | $9.2\times10^{-1}$ | N/A | $1.2\times10^{-1}$ | $1.3\times10^{2}$ | $7.3\times10^{-1}$ |
| R454G | $5.9\times10^{-2}$ | **0** | $1.1\times10^{5}$ | **$2.6\times10^{-2}$** | – | – | $8.5\times10^{-1}$ | N/A | **$7.0\times10^{-4}$** | $9.6\times10^{1}$ | $3.0\times10^{-1}$ |
| R454V | $5.7\times10^{-2}$ | **0** | $2.0\times10^{5}$ | **$3.7\times10^{-2}$** | – | – | $5.8\times10^{-1}$ | N/A | **0** | $1.3\times10^{2}$ | $7.0\times10^{-1}$ |
| V455A | 1.2 | $4.6\times10^{-1}$ | $1.3\times10^{5}$ | **$2.7\times10^{-2}$** | 2.0 | $3.7\times10^{-2}$ | N/A | 1.1 | $5.3\times10^{-2}$ | $1.2\times10^{2}$ | $7.1\times10^{-1}$ |
| V455G | $1.4\times10^{-1}$ | **0** | $1.5\times10^{5}$ | **$4.2\times10^{-2}$** | 2.0 | $2.9\times10^{-2}$ | $8.5\times10^{-1}$ | N/A | **$2.0\times10^{-4}$** | $1.0\times10^{2}$ | $4.3\times10^{-1}$ |
| I456G | $1.7\times10^{-1}$ | **0** | $2.0\times10^{5}$ | **$4.2\times10^{-2}$** | 1.8 | $4.7\times10^{-2}$ | $5.7\times10^{-1}$ | N/A | **0** | $8.6\times10^{1}$ | $2.1\times10^{-1}$ |
| I456V | $8.1\times10^{-1}$ | $2.6\times10^{-1}$ | $6.4\times10^{5}$ | $8.6\times10^{-1}$ | $6.4\times10^{-1}$ | $1.7\times10^{-1}$ | N/A | 1.0 | $2.6\times10^{-1}$ | $1.7\times10^{2}$ | $4.7\times10^{-1}$ |
| N457G | $3.8\times10^{-1}$ | **0** | $3.3\times10^{5}$ | $1.7\times10^{-1}$ | 1.5 | $1.8\times10^{-1}$ | $8.9\times10^{-1}$ | N/A | $1.8\times10^{-2}$ | $9.8\times10^{1}$ | $3.4\times10^{-1}$ |
| N457V | $1.3\times10^{-1}$ | **0** | $6.7\times10^{4}$ | **$2.6\times10^{-2}$** | $1.0\times10^{1}$ | **$1.2\times10^{-3}$** | $6.6\times10^{-1}$ | N/A | **0** | $1.2\times10^{2}$ | $6.0\times10^{-1}$ |
| G458A | $1.8\times10^{-1}$ | **0** | $7.8\times10^{5}$ | $4.7\times10^{-1}$ | 2.1 | $5.1\times10^{-2}$ | $4.8\times10^{-1}$ | N/A | **$1.\times10^{-4}$** | $9.2\times10^{1}$ | $3.7\times10^{-1}$ |
| G458V | $2.3\times10^{-1}$ | **0** | $9.1\times10^{4}$ | $6.2\times10^{-2}$ | 4.2 | **$5.5\times10^{-3}$** | $3.3\times10^{-1}$ | N/A | **0** | $4.8\times10^{1}$ | **$4.6\times10^{-2}$** |
| Y459G | $\leq 1.6\times10^{-2}$ | **0** | $\leq 1.2\times10^{5}$ | **$3.7\times10^{-2}$** | – | – | $8.8\times10^{-1}$ | N/A | $1.7\times10^{-2}$ | $\geq 1.0\times10^{2}$ | $4.5\times10^{-1}$ |
| Y459V | $4.0\times10^{-1}$ | **$5.0\times10^{-3}$** | $2.3\times10^{5}$ | $9.2\times10^{-2}$ | 1.6 | $2.0\times10^{-1}$ | $9.7\times10^{-1}$ | N/A | $6.0\times10^{-1}$ | $1.1\times10^{2}$ | $5.6\times10^{-1}$ |
| N460V | $2.2\times10^{-1}$ | **0** | $6.2\times10^{5}$ | $9.7\times10^{-1}$ | – | – | $9.4\times10^{-1}$ | N/A | $2.1\times10^{-1}$ | $1.7\times10^{2}$ | $6.8\times10^{-1}$ |
| W461G | $1.3\times10^{-1}$ | **0** | $1.0\times10^{6}$ | $7.9\times10^{-2}$ | 1.5 | $2.5\times10^{-1}$ | $6.8\times10^{-1}$ | N/A | **$6.0\times10^{-4}$** | $9.7\times10^{1}$ | $4.4\times10^{-1}$ |
| W461V | $1.2\times10^{-1}$ | **0** | $1.1\times10^{6}$ | $5.9\times10^{-2}$ | 1.1 | $8.3\times10^{-1}$ | $7.2\times10^{-1}$ | N/A | **0** | $1.0\times10^{2}$ | $5.4\times10^{-1}$ |
| K462G | 1.3 | $4.4\times10^{-1}$ | $2.4\times10^{5}$ | $1.5\times10^{-1}$ | 2.5 | **$2.6\times10^{-3}$** | N/A | 1.0 | $2.4\times10^{-1}$ | $9.7\times10^{1}$ | $3.3\times10^{-1}$ |
| K462V | $6.5\times10^{-1}$ | $1.2\times10^{-1}$ | $6.8\times10^{5}$ | $7.3\times10^{-1}$ | $7.0\times10^{-1}$ | $2.8\times10^{-1}$ | N/A | 1.0 | $8.6\times10^{-1}$ | $1.3\times10^{2}$ | $8.3\times10^{-1}$ |
| R463G | $1.8\times10^{-1}$ | **0** | $1.5\times10^{5}$ | **$1.7\times10^{-2}$** | 2.2 | $3.8\times10^{-2}$ | N/A | 1.1 | $9.9\times10^{-2}$ | $3.6\times10^{1}$ | **$2.8\times10^{-2}$** |
| R463V | $3.2\times10^{-1}$ | **0** | $3.9\times10^{5}$ | $2.4\times10^{-1}$ | $9.9\times10^{-1}$ | $9.9\times10^{-1}$ | $8.4\times10^{-1}$ | N/A | **$3.8\times10^{-3}$** | $1.3\times10^{2}$ | $8.5\times10^{-1}$ |
| S464G | 1.1 | $7.7\times10^{-1}$ | $5.2\times10^{5}$ | $6.4\times10^{-1}$ | $9.4\times10^{-1}$ | $8.5\times10^{-1}$ | $7.3\times10^{-1}$ | N/A | **$1.7\times10^{-3}$** | $8.0\times10^{1}$ | $4.1\times10^{-1}$ |
| S464V | – | – | – | – | – | – | – | – | – | – | – |
| G465A | $1.7\times10^{-1}$ | **0** | $5.1\times10^{5}$ | $6.2\times10^{-1}$ | 1.2 | $4.8\times10^{-1}$ | $6.8\times10^{-1}$ | N/A | **0** | $8.0\times10^{1}$ | $2.8\times10^{-1}$ |
| G465V | $\leq 4.6\times10^{-3}$ | **0** | $\leq 1.3\times10^{5}$ | $6.4\times10^{-2}$ | – | – | $1.0\times10^{-1}$ | N/A | **$4.0\times10^{-4}$** | $\geq 1.9\times10^{1}$ | $6.4\times10^{-2}$ |
| D466G | $3.7\times10^{-1}$ | **$1.0\times10^{-3}$** | $1.9\times10^{5}$ | $6.8\times10^{-2}$ | 2.9 | **$4.7\times10^{-3}$** | N/A | 1.1 | $5.7\times10^{-2}$ | $9.6\times10^{1}$ | $3.8\times10^{-1}$ |
| D466V | $7.4\times10^{-1}$ | $2.0\times10^{-1}$ | $5.0\times10^{4}$ | **$1.3\times10^{-2}$** | 5.2 | **$4.1\times10^{-4}$** | N/A | 1.2 | **$5.1\times10^{-3}$** | $1.3\times10^{2}$ | $8.3\times10^{-1}$ |
| I467G | $2.2\times10^{-1}$ | **0** | $2.2\times10^{5}$ | $6.9\times10^{-2}$ | 1.4 | $4.5\times10^{-1}$ | N/A | 1.0 | $7.9\times10^{-1}$ | $1.2\times10^{2}$ | $6.4\times10^{-1}$ |
| I467V | $6.6\times10^{-1}$ | $6.7\times10^{-2}$ | $7.4\times10^{5}$ | $4.4\times10^{-1}$ | $5.9\times10^{-1}$ | $1.1\times10^{-1}$ | N/A | 1.0 | $9.6\times10^{-1}$ | $1.3\times10^{2}$ | $7.7\times10^{-1}$ |
| Q468G | 1.6 | $7.2\times10^{-2}$ | $2.5\times10^{5}$ | $8.4\times10^{-2}$ | 1.6 | $1.1\times10^{-1}$ | N/A | 1.1 | $1.2\times10^{-1}$ | $1.5\times10^{2}$ | $9.1\times10^{-1}$ |
| Q468V | $4.2\times10^{-1}$ | **0** | $9.2\times10^{5}$ | $1.0\times10^{-1}$ | $4.9\times10^{-1}$ | $3.1\times10^{-2}$ | N/A | 1.3 | **$2.8\times10^{-3}$** | $1.3\times10^{2}$ | $8.7\times10^{-1}$ |
| I469G | $9.6\times10^{-1}$ | $8.3\times10^{-1}$ | $1.1\times10^{5}$ | $7.6\times10^{-2}$ | 2.6 | $1.6\times10^{-2}$ | N/A | 1.0 | $3.7\times10^{-1}$ | $1.2\times10^{2}$ | $7.7\times10^{-1}$ |
| I469V | $5.7\times10^{-1}$ | $2.5\times10^{-2}$ | $9.9\times10^{5}$ | $6.7\times10^{-2}$ | $7.4\times10^{-1}$ | $2.8\times10^{-1}$ | 1.0 | N/A | $9.5\times10^{-1}$ | $1.5\times10^{2}$ | $9.2\times10^{-1}$ |
| I470G | $1.3\times10^{-1}$ | **0** | $1.7\times10^{5}$ | $6.9\times10^{-2}$ | 3.3 | **$8.1\times10^{-3}$** | $7.0\times10^{-1}$ | N/A | **$4.0\times10^{-4}$** | $5.9\times10^{1}$ | $1.4\times10^{-1}$ |
| I470V | $3.3\times10^{-1}$ | **0** | $1.6\times10^{6}$ | **$4.0\times10^{-3}$** | $5.2\times10^{-1}$ | $5.8\times10^{-2}$ | $9.8\times10^{-1}$ | N/A | $4.4\times10^{-1}$ | $1.5\times10^{2}$ | $8.3\times10^{-1}$ |
| S471G | 1.6 | $5.6\times10^{-2}$ | $4.0\times10^{5}$ | $3.0\times10^{-1}$ | 1.1 | $8.4\times10^{-1}$ | $8.2\times10^{-1}$ | N/A | **$1.3\times10^{-3}$** | $1.2\times10^{2}$ | $6.7\times10^{-1}$ |
| S471V | 1.1 | $8.1\times10^{-1}$ | $1.1\times10^{6}$ | $1.1\times10^{-1}$ | $3.5\times10^{-1}$ | **$3.3\times10^{-3}$** | $9.9\times10^{-1}$ | N/A | $7.1\times10^{-1}$ | $1.4\times10^{2}$ | $9.9\times10^{-1}$ |
| K472G | $4.3\times10^{-1}$ | **$7.0\times10^{-3}$** | $6.7\times10^{5}$ | $7.6\times10^{-1}$ | $9.7\times10^{-1}$ | $9.3\times10^{-1}$ | $7.6\times10^{-1}$ | N/A | **$1.\times10^{-4}$** | $7.4\times10^{1}$ | $2.1\times10^{-1}$ |
| K472V | $2.6\times10^{-1}$ | **0** | $1.4\times10^{6}$ | **$1.2\times10^{-2}$** | – | – | $8.2\times10^{-1}$ | N/A | **$3.0\times10^{-4}$** | $1.2\times10^{2}$ | $6.2\times10^{-1}$ |
| D473G | 1.3 | $3.6\times10^{-1}$ | $4.4\times10^{5}$ | $3.8\times10^{-1}$ | 1.5 | $2.3\times10^{-1}$ | $9.2\times10^{-1}$ | N/A | $1.4\times10^{-1}$ | $1.2\times10^{2}$ | $6.1\times10^{-1}$ |
| D473V | $4.8\times10^{-1}$ | **$5.0\times10^{-3}$** | $1.8\times10^{4}$ | **$1.0\times10^{-2}$** | $6.1\times10^{1}$ | **0** | $5.1\times10^{-1}$ | N/A | **$3.0\times10^{-4}$** | $1.1\times10^{2}$ | $5.6\times10^{-1}$ |
| G474A | $5.6\times10^{-1}$ | $3.8\times10^{-2}$ | $8.8\times10^{5}$ | $1.7\times10^{-1}$ | $6.1\times10^{-1}$ | $9.6\times10^{-2}$ | $8.1\times10^{-1}$ | N/A | **$2.0\times10^{-4}$** | $6.9\times10^{1}$ | $1.9\times10^{-1}$ |
| G474V | $2.1\times10^{-1}$ | **0** | $2.3\times10^{5}$ | $1.4\times10^{-1}$ | – | – | $6.9\times10^{-1}$ | N/A | **0** | $1.0\times10^{2}$ | $5.4\times10^{-1}$ |
| Y475G | $1.5\times10^{-1}$ | **0** | $2.8\times10^{5}$ | $1.3\times10^{-1}$ | 1.5 | $2.7\times10^{-1}$ | $6.5\times10^{-1}$ | N/A | **0** | $9.5\times10^{1}$ | $3.9\times10^{-1}$ |
| Y475V | $5.0\times10^{-1}$ | $3.4\times10^{-2}$ | $1.0\times10^{6}$ | $8.7\times10^{-2}$ | $4.1\times10^{-1}$ | $1.3\times10^{-2}$ | N/A | 1.0 | $6.3\times10^{-1}$ | $1.6\times10^{2}$ | $7.1\times10^{-1}$ |
| L476G | $9.2\times10^{-1}$ | $6.7\times10^{-1}$ | $9.6\times10^{5}$ | **$4.9\times10^{-2}$** | $3.1\times10^{-1}$ | **$2.0\times10^{-5}$** | $5.3\times10^{-1}$ | N/A | **0** | $1.0\times10^{2}$ | $3.5\times10^{-1}$ |
| L476V | $9.2\times10^{-1}$ | $8.3\times10^{-1}$ | $6.6\times10^{5}$ | $8.2\times10^{-1}$ | $7.2\times10^{-1}$ | $2.1\times10^{-1}$ | N/A | 1.3 | **0** | $1.7\times10^{2}$ | $6.1\times10^{-1}$ |
| S477G | 1.1 | $7.5\times10^{-1}$ | $3.4\times10^{5}$ | $2.5\times10^{-1}$ | 3.0 | **$6.0\times10^{-5}$** | $8.2\times10^{-1}$ | N/A | $1.3\times10^{-2}$ | $1.3\times10^{2}$ | $7.8\times10^{-1}$ |
| S477V | $5.0\times10^{-1}$ | $3.6\times10^{-2}$ | $9.4\times10^{5}$ | $1.5\times10^{-1}$ | – | – | $9.0\times10^{-1}$ | N/A | $4.5\times10^{-2}$ | $1.8\times10^{2}$ | $4.6\times10^{-1}$ |
| A478G | $9.4\times10^{-1}$ | $8.1\times10^{-1}$ | $5.0\times10^{5}$ | $6.0\times10^{-1}$ | 1.4 | $2.3\times10^{-1}$ | 1.0 | N/A | $8.2\times10^{-1}$ | $1.1\times10^{2}$ | $4.7\times10^{-1}$ |

| Variant | $f_a$ | $f_a$ *p*-value | $(k_{cat}/K_M)^{MeP}_{chem.}$ ($M^{-1}s^{-1}$) | $(k_{cat}/K_M)^{MeP}_{chem.}$ *p*-value | FC1 | FC1 *p*-value | FC2 | FC3 | FC2/3 *p*-value | FC4 ($s^{-1}$) | FC4 *p*-value |
|---|---|---|---|---|---|---|---|---|---|---|---|
| A478V | $7.8\times10^{-1}$ | $3.4\times10^{-1}$ | $4.0\times10^{5}$ | $2.7\times10^{-1}$ | $7.5\times10^{-1}$ | $4.1\times10^{-1}$ | N/A | 1.1 | $1.2\times10^{-1}$ | $1.5\times10^{2}$ | $8.4\times10^{-1}$ |
| Y479G | 1.7 | $1.3\times10^{-1}$ | $6.4\times10^{5}$ | $9.2\times10^{-1}$ | $4.1\times10^{-1}$ | $1.1\times10^{-2}$ | $8.7\times10^{-1}$ | N/A | $1.4\times10^{-2}$ | $1.1\times10^{2}$ | $7.3\times10^{-1}$ |
| Y479V | $8.0\times10^{-1}$ | $2.6\times10^{-1}$ | $8.8\times10^{5}$ | $8.7\times10^{-2}$ | $2.8\times10^{-1}$ | **$3.0\times10^{-5}$** | N/A | 1.0 | $2.8\times10^{-1}$ | $2.0\times10^{2}$ | $1.5\times10^{-1}$ |
| S480G | 1.2 | $6.1\times10^{-1}$ | $4.1\times10^{5}$ | $3.6\times10^{-1}$ | 2.0 | $2.1\times10^{-2}$ | $6.6\times10^{-1}$ | N/A | **$3.0\times10^{-4}$** | $1.1\times10^{2}$ | $5.1\times10^{-1}$ |
| S480V | $6.7\times10^{-1}$ | $9.2\times10^{-2}$ | $1.1\times10^{6}$ | **$2.8\times10^{-2}$** | $2.8\times10^{-1}$ | $1.1\times10^{-2}$ | N/A | 1.2 | **$8.7\times10^{-3}$** | $2.6\times10^{2}$ | **$4.1\times10^{-2}$** |
| K481G | 1.3 | $3.2\times10^{-1}$ | $4.9\times10^{5}$ | $5.4\times10^{-1}$ | $9.3\times10^{-1}$ | $8.1\times10^{-1}$ | $9.8\times10^{-1}$ | N/A | $5.7\times10^{-1}$ | $9.6\times10^{1}$ | $3.4\times10^{-1}$ |
| K481V | $9.2\times10^{-1}$ | $8.1\times10^{-1}$ | $5.6\times10^{5}$ | $8.2\times10^{-1}$ | $7.8\times10^{-1}$ | $4.7\times10^{-1}$ | N/A | 1.1 | $7.4\times10^{-2}$ | $1.6\times10^{2}$ | $7.4\times10^{-1}$ |
| K482G | $8.0\times10^{-1}$ | $3.0\times10^{-1}$ | $4.3\times10^{5}$ | $3.2\times10^{-1}$ | $7.7\times10^{-1}$ | $3.1\times10^{-1}$ | $7.8\times10^{-1}$ | N/A | **0** | $1.1\times10^{2}$ | $5.6\times10^{-1}$ |
| K482V | $9.9\times10^{-1}$ | $9.3\times10^{-1}$ | $5.8\times10^{5}$ | $9.2\times10^{-1}$ | $7.6\times10^{-1}$ | $2.7\times10^{-1}$ | N/A | 1.0 | $4.7\times10^{-1}$ | $1.5\times10^{2}$ | $7.9\times10^{-1}$ |
| G483A | $3.3\times10^{-1}$ | **0** | $3.3\times10^{5}$ | $1.2\times10^{-1}$ | 2.6 | **$1.1\times10^{-3}$** | $5.0\times10^{-1}$ | N/A | **$1.\times10^{-4}$** | $1.5\times10^{2}$ | $9.3\times10^{-1}$ |
| G483V | $6.7\times10^{-1}$ | $1.7\times10^{-1}$ | $2.3\times10^{5}$ | $1.1\times10^{-1}$ | $8.7\times10^{-1}$ | $5.9\times10^{-1}$ | $3.2\times10^{-1}$ | N/A | **$2.0\times10^{-4}$** | $5.6\times10^{1}$ | $2.6\times10^{-1}$ |
| T484G | 1.1 | $6.8\times10^{-1}$ | $2.4\times10^{5}$ | $8.7\times10^{-2}$ | 1.1 | $8.7\times10^{-1}$ | $4.5\times10^{-1}$ | N/A | **0** | $7.3\times10^{1}$ | $1.8\times10^{-1}$ |
| T484V | $5.9\times10^{-1}$ | **$9.0\times10^{-3}$** | $5.9\times10^{5}$ | $9.2\times10^{-1}$ | $9.7\times10^{-1}$ | $9.6\times10^{-1}$ | $9.1\times10^{-1}$ | N/A | $6.7\times10^{-2}$ | $1.8\times10^{2}$ | $3.3\times10^{-1}$ |
| T485G | $8.8\times10^{-1}$ | $6.3\times10^{-1}$ | $5.0\times10^{5}$ | $5.4\times10^{-1}$ | 1.7 | $8.8\times10^{-2}$ | $6.5\times10^{-1}$ | N/A | **$1.\times10^{-4}$** | $1.3\times10^{2}$ | $8.9\times10^{-1}$ |
| T485V | $4.7\times10^{-1}$ | **$6.0\times10^{-3}$** | $1.1\times10^{6}$ | **$3.4\times10^{-2}$** | – | – | $6.5\times10^{-1}$ | N/A | **0** | $2.1\times10^{2}$ | $1.7\times10^{-1}$ |
| H486G | $\leq 3.2\times10^{-2}$ | **0** | $\leq 3.7\times10^{4}$ | **$7.0\times10^{-3}$** | – | – | – | – | – | – | – |
| H486V | $\leq 4.9\times10^{-3}$ | **$1.0\times10^{-3}$** | $\leq 2.0\times10^{5}$ | $1.7\times10^{-1}$ | – | – | – | – | – | – | – |
| S487G | $\leq 9.1\times10^{-1}$ | $8.2\times10^{-1}$ | $\leq 4.6\times10^{5}$ | $5.5\times10^{-1}$ | $\leq 4.5$ | **$2.5\times10^{-4}$** | $6.3\times10^{-1}$ | N/A | **$2.0\times10^{-4}$** | $\geq 1.2\times10^{2}$ | $7.4\times10^{-1}$ |
| S487V | 1.2 | $3.4\times10^{-1}$ | $1.5\times10^{5}$ | **$2.2\times10^{-2}$** | – | – | N/A | 1.7 | **$1.\times10^{-4}$** | $1.5\times10^{2}$ | $9.0\times10^{-1}$ |
| V488A | 2.3 | **$9.0\times10^{-3}$** | $3.7\times10^{5}$ | $2.0\times10^{-1}$ | 1.3 | $3.8\times10^{-1}$ | $9.2\times10^{-1}$ | N/A | $3.3\times10^{-1}$ | $1.3\times10^{2}$ | $8.0\times10^{-1}$ |
| V488G | 2.0 | **$8.0\times10^{-3}$** | $2.5\times10^{5}$ | $7.6\times10^{-2}$ | 4.8 | **$2.0\times10^{-5}$** | $5.9\times10^{-1}$ | N/A | **0** | $9.3\times10^{1}$ | $2.9\times10^{-1}$ |
| W489G | $2.5\times10^{-1}$ | **0** | $2.7\times10^{5}$ | $6.6\times10^{-2}$ | 4.3 | **$3.2\times10^{-4}$** | $2.0\times10^{-1}$ | N/A | **0** | $3.5\times10^{1}$ | **$4.6\times10^{-2}$** |
| W489V | $4.6\times10^{-1}$ | **$7.0\times10^{-3}$** | $3.0\times10^{4}$ | **$1.4\times10^{-2}$** | – | – | $4.9\times10^{-1}$ | N/A | **$2.0\times10^{-4}$** | $9.3\times10^{1}$ | $3.1\times10^{-1}$ |
| N490G | 1.3 | $3.2\times10^{-1}$ | $1.6\times10^{5}$ | $5.3\times10^{-2}$ | 3.7 | **$2.5\times10^{-4}$** | $7.2\times10^{-1}$ | N/A | **$1.\times10^{-4}$** | $9.1\times10^{1}$ | $3.0\times10^{-1}$ |
| N490V | $7.1\times10^{-1}$ | $8.5\times10^{-2}$ | $2.6\times10^{5}$ | **$4.6\times10^{-2}$** | 2.1 | **$8.8\times10^{-3}$** | $7.9\times10^{-1}$ | N/A | **0** | $1.0\times10^{2}$ | $3.7\times10^{-1}$ |
| S491G | $6.3\times10^{-1}$ | $1.1\times10^{-1}$ | $4.2\times10^{5}$ | $5.2\times10^{-1}$ | 1.1 | $7.4\times10^{-1}$ | $8.7\times10^{-1}$ | N/A | **$6.5\times10^{-3}$** | $1.0\times10^{2}$ | $4.5\times10^{-1}$ |
| S491V | $7.1\times10^{-1}$ | $1.2\times10^{-1}$ | $4.0\times10^{5}$ | $2.4\times10^{-1}$ | $7.1\times10^{-1}$ | $2.9\times10^{-1}$ | $8.2\times10^{-1}$ | N/A | **$1.\times10^{-4}$** | $8.8\times10^{1}$ | $2.3\times10^{-1}$ |
| Y492G | $9.1\times10^{-1}$ | $7.1\times10^{-1}$ | $2.4\times10^{5}$ | $9.0\times10^{-2}$ | 2.2 | $1.1\times10^{-2}$ | $4.1\times10^{-1}$ | N/A | **$1.\times10^{-4}$** | $5.3\times10^{1}$ | $1.1\times10^{-1}$ |
| Y492V | $5.3\times10^{-1}$ | $1.3\times10^{-2}$ | $5.1\times10^{5}$ | $5.9\times10^{-1}$ | – | – | $9.8\times10^{-1}$ | N/A | $5.7\times10^{-1}$ | $9.2\times10^{1}$ | $3.2\times10^{-1}$ |
| D493G | $5.4\times10^{-1}$ | $1.2\times10^{-2}$ | $6.9\times10^{4}$ | **$2.2\times10^{-2}$** | 2.1 | $1.1\times10^{-1}$ | $2.2\times10^{-1}$ | N/A | **$1.\times10^{-4}$** | $4.1\times10^{1}$ | $6.5\times10^{-2}$ |
| D493V | $8.8\times10^{-1}$ | $5.4\times10^{-1}$ | $9.4\times10^{4}$ | **$1.8\times10^{-2}$** | 1.7 | $1.4\times10^{-1}$ | $3.5\times10^{-1}$ | N/A | **0** | $8.4\times10^{1}$ | $1.8\times10^{-1}$ |
| S494G | $5.9\times10^{-1}$ | $3.9\times10^{-2}$ | $3.4\times10^{5}$ | $1.8\times10^{-1}$ | 2.1 | $2.7\times10^{-2}$ | $6.8\times10^{-1}$ | N/A | **$1.\times10^{-4}$** | $1.0\times10^{2}$ | $4.9\times10^{-1}$ |
| S494V | $2.0\times10^{-1}$ | **0** | $3.6\times10^{5}$ | $1.3\times10^{-1}$ | 1.0 | $9.5\times10^{-1}$ | $3.9\times10^{-1}$ | N/A | **0** | $1.1\times10^{2}$ | $5.1\times10^{-1}$ |
| H495G | – | – | – | – | – | – | – | – | – | – | – |
| H495V | $1.5\times10^{-1}$ | **0** | $1.0\times10^{6}$ | $1.6\times10^{-1}$ | – | – | $8.2\times10^{-1}$ | N/A | $1.9\times10^{-2}$ | $1.6\times10^{2}$ | $8.1\times10^{-1}$ |
| I496G | $\leq 5.3\times10^{-3}$ | **0** | $\leq 2.5\times10^{5}$ | $1.1\times10^{-1}$ | – | – | $6.4\times10^{-1}$ | N/A | **$1.1\times10^{-3}$** | $\geq 1.4\times10^{2}$ | $9.3\times10^{-1}$ |
| I496V | $2.9\times10^{-1}$ | **0** | $1.2\times10^{6}$ | **$3.9\times10^{-2}$** | $6.8\times10^{-1}$ | $2.4\times10^{-1}$ | $8.5\times10^{-1}$ | N/A | **$8.1\times10^{-3}$** | $1.5\times10^{2}$ | $8.3\times10^{-1}$ |
| P497G | $1.5\times10^{-1}$ | **0** | $6.4\times10^{5}$ | $8.7\times10^{-1}$ | – | – | $9.2\times10^{-1}$ | N/A | $2.8\times10^{-1}$ | $1.3\times10^{2}$ | $9.3\times10^{-1}$ |
| L498G | $\leq 6.0\times10^{-1}$ | $9.5\times10^{-2}$ | $\leq 2.3\times10^{5}$ | $1.1\times10^{-1}$ | $\leq 5.4$ | **$5.0\times10^{-5}$** | $7.5\times10^{-1}$ | N/A | **$4.0\times10^{-4}$** | $\geq 8.8\times10^{1}$ | $3.8\times10^{-1}$ |
| L498V | $4.0\times10^{-1}$ | **$3.0\times10^{-3}$** | $6.9\times10^{5}$ | $6.5\times10^{-1}$ | $6.8\times10^{-1}$ | $2.1\times10^{-1}$ | N/A | 1.1 | $1.1\times10^{-1}$ | $1.7\times10^{2}$ | $6.4\times10^{-1}$ |
| L499G | $8.0\times10^{-1}$ | $4.6\times10^{-1}$ | $1.7\times10^{5}$ | $7.8\times10^{-2}$ | 2.1 | $5.6\times10^{-2}$ | $9.7\times10^{-1}$ | N/A | $6.3\times10^{-1}$ | $1.3\times10^{2}$ | $8.7\times10^{-1}$ |
| L499V | $6.9\times10^{-1}$ | $1.2\times10^{-1}$ | $7.1\times10^{5}$ | $5.3\times10^{-1}$ | $7.3\times10^{-1}$ | $3.3\times10^{-1}$ | N/A | 1.1 | $6.9\times10^{-2}$ | $1.5\times10^{2}$ | $7.6\times10^{-1}$ |
| F500G | $\leq 5.1\times10^{-4}$ | **0** | $\leq 2.2\times10^{6}$ | **$1.0\times10^{-3}$** | – | – | – | – | – | – | – |
| F500V | $3.2\times10^{-1}$ | **$1.0\times10^{-3}$** | $3.3\times10^{5}$ | $2.3\times10^{-1}$ | – | – | $6.8\times10^{-1}$ | N/A | **0** | $1.2\times10^{2}$ | $7.2\times10^{-1}$ |
| M501G | $5.5\times10^{-1}$ | $5.6\times10^{-2}$ | $6.0\times10^{4}$ | **$4.9\times10^{-2}$** | 3.3 | **$5.3\times10^{-3}$** | $9.5\times10^{-1}$ | N/A | $3.5\times10^{-1}$ | $1.1\times10^{2}$ | $6.7\times10^{-1}$ |
| M501V | $7.8\times10^{-1}$ | $3.4\times10^{-1}$ | $2.4\times10^{5}$ | $2.5\times10^{-1}$ | – | – | $9.6\times10^{-1}$ | N/A | $3.8\times10^{-1}$ | $1.4\times10^{2}$ | 1.0 |
| G502A | $1.6\times10^{-1}$ | **$2.0\times10^{-3}$** | $4.9\times10^{5}$ | $6.4\times10^{-1}$ | – | – | $9.0\times10^{-1}$ | N/A | $2.1\times10^{-1}$ | $1.3\times10^{2}$ | $9.2\times10^{-1}$ |
| G502V | $\leq 1.1\times10^{-3}$ | **0** | $\leq 1.0\times10^{7}$ | **$1.0\times10^{-3}$** | – | – | – | – | – | – | – |
| W503G | $4.0\times10^{-1}$ | **$5.0\times10^{-3}$** | $3.5\times10^{5}$ | $2.3\times10^{-1}$ | 1.5 | $2.7\times10^{-1}$ | $9.8\times10^{-1}$ | N/A | $5.6\times10^{-1}$ | $8.4\times10^{1}$ | $2.5\times10^{-1}$ |
| W503V | $3.0\times10^{-1}$ | **0** | $3.3\times10^{5}$ | $2.3\times10^{-1}$ | 2.4 | $3.9\times10^{-2}$ | $9.7\times10^{-1}$ | N/A | $5.4\times10^{-1}$ | $1.8\times10^{2}$ | $3.8\times10^{-1}$ |
| G504A | $8.7\times10^{-1}$ | $5.4\times10^{-1}$ | $3.1\times10^{5}$ | $1.4\times10^{-1}$ | 1.3 | $3.7\times10^{-1}$ | $9.9\times10^{-1}$ | N/A | $6.7\times10^{-1}$ | $1.2\times10^{2}$ | $6.6\times10^{-1}$ |
| G504V | $2.6\times10^{-1}$ | **0** | $8.1\times10^{5}$ | $3.0\times10^{-1}$ | – | – | N/A | 1.0 | $3.7\times10^{-1}$ | $1.6\times10^{2}$ | $6.9\times10^{-1}$ |
| I505G | $1.3\times10^{-1}$ | **0** | $1.9\times10^{5}$ | $5.2\times10^{-2}$ | – | – | N/A | 1.0 | $3.6\times10^{-1}$ | $1.1\times10^{2}$ | $4.8\times10^{-1}$ |
| I505V | 1.3 | $2.6\times10^{-1}$ | $4.6\times10^{5}$ | $3.3\times10^{-1}$ | $8.2\times10^{-1}$ | $5.0\times10^{-1}$ | N/A | 1.1 | $6.2\times10^{-2}$ | $1.4\times10^{2}$ | $9.8\times10^{-1}$ |
| K506G | 1.6 | $7.4\times10^{-2}$ | $2.7\times10^{5}$ | $1.1\times10^{-1}$ | 1.6 | $8.6\times10^{-2}$ | $9.6\times10^{-1}$ | N/A | $3.9\times10^{-1}$ | $1.3\times10^{2}$ | $7.8\times10^{-1}$ |
| K506V | $8.4\times10^{-1}$ | $4.4\times10^{-1}$ | $4.3\times10^{5}$ | $3.7\times10^{-1}$ | 1.0 | $9.6\times10^{-1}$ | N/A | 1.1 | $1.7\times10^{-2}$ | $1.6\times10^{2}$ | $5.8\times10^{-1}$ |
| Q507G | 1.1 | $5.9\times10^{-1}$ | $4.5\times10^{5}$ | $3.6\times10^{-1}$ | 1.0 | $9.5\times10^{-1}$ | $8.6\times10^{-1}$ | N/A | **$9.2\times10^{-3}$** | $1.1\times10^{2}$ | $5.6\times10^{-1}$ |
| Q507V | 1.2 | $4.9\times10^{-1}$ | $5.8\times10^{5}$ | $8.6\times10^{-1}$ | $7.0\times10^{-1}$ | $2.7\times10^{-1}$ | N/A | 1.0 | $6.7\times10^{-1}$ | $1.8\times10^{2}$ | $5.1\times10^{-1}$ |
| G508A | $4.9\times10^{-1}$ | **$6.0\times10^{-3}$** | $5.1\times10^{5}$ | $6.3\times10^{-1}$ | 1.3 | $3.6\times10^{-1}$ | $8.4\times10^{-1}$ | N/A | **$6.3\times10^{-3}$** | $1.2\times10^{2}$ | $6.7\times10^{-1}$ |
| G508V | $1.7\times10^{-1}$ | **0** | $7.1\times10^{5}$ | $5.9\times10^{-1}$ | $7.0\times10^{-1}$ | $2.9\times10^{-1}$ | $8.1\times10^{-1}$ | N/A | **0** | $1.2\times10^{2}$ | $5.9\times10^{-1}$ |
| E509G | $1.5\times10^{-1}$ | **0** | $1.1\times10^{6}$ | $1.0\times10^{-1}$ | $6.2\times10^{-1}$ | $2.4\times10^{-1}$ | $7.0\times10^{-1}$ | N/A | **0** | $9.6\times10^{1}$ | $3.8\times10^{-1}$ |
| E509V | $4.0\times10^{-1}$ | **$5.0\times10^{-3}$** | $5.1\times10^{5}$ | $6.5\times10^{-1}$ | $8.6\times10^{-1}$ | $7.0\times10^{-1}$ | $8.9\times10^{-1}$ | N/A | $4.5\times10^{-2}$ | $1.2\times10^{2}$ | $7.3\times10^{-1}$ |
| S510G | $8.8\times10^{-1}$ | $6.0\times10^{-1}$ | $3.0\times10^{5}$ | $2.0\times10^{-1}$ | 1.5 | $2.1\times10^{-1}$ | $8.8\times10^{-1}$ | N/A | $3.4\times10^{-2}$ | $9.4\times10^{1}$ | $3.2\times10^{-1}$ |
| S510V | $1.3\times10^{-1}$ | **0** | $2.6\times10^{5}$ | $1.1\times10^{-1}$ | – | – | $7.4\times10^{-1}$ | N/A | **$3.2\times10^{-3}$** | $1.4\times10^{2}$ | $9.6\times10^{-1}$ |
| N511G | 1.2 | $4.7\times10^{-1}$ | $2.2\times10^{5}$ | $9.7\times10^{-2}$ | 2.2 | $2.6\times10^{-2}$ | $9.9\times10^{-1}$ | N/A | $7.7\times10^{-1}$ | $1.1\times10^{2}$ | $5.8\times10^{-1}$ |
| N511V | 1.6 | $7.8\times10^{-2}$ | $5.9\times10^{5}$ | $9.5\times10^{-1}$ | $6.7\times10^{-1}$ | $2.6\times10^{-1}$ | $9.9\times10^{-1}$ | N/A | $5.7\times10^{-1}$ | $2.0\times10^{2}$ | $2.5\times10^{-1}$ |
| Q512G | $7.0\times10^{-1}$ | $1.7\times10^{-1}$ | $3.0\times10^{5}$ | $2.1\times10^{-1}$ | 2.9 | **$2.0\times10^{-3}$** | $8.8\times10^{-1}$ | N/A | $3.3\times10^{-2}$ | $1.1\times10^{2}$ | $5.2\times10^{-1}$ |
| Q512V | $8.0\times10^{-1}$ | $3.3\times10^{-1}$ | $8.3\times10^{5}$ | $2.0\times10^{-1}$ | $5.8\times10^{-1}$ | $9.3\times10^{-2}$ | N/A | 1.0 | $7.1\times10^{-1}$ | $1.3\times10^{2}$ | $7.9\times10^{-1}$ |
| P513G | $9.4\times10^{-1}$ | $7.5\times10^{-1}$ | $2.3\times10^{5}$ | $7.7\times10^{-2}$ | 2.1 | $2.1\times10^{-2}$ | $9.6\times10^{-1}$ | N/A | $3.5\times10^{-1}$ | $1.1\times10^{2}$ | $5.4\times10^{-1}$ |
| P513V | $6.2\times10^{-1}$ | **$5.0\times10^{-3}$** | $5.7\times10^{5}$ | $7.8\times10^{-1}$ | $8.3\times10^{-1}$ | $4.2\times10^{-1}$ | N/A | 1.0 | $5.3\times10^{-1}$ | $1.7\times10^{2}$ | $5.3\times10^{-1}$ |
| Y514G | $1.2\times10^{-1}$ | **0** | $4.5\times10^{5}$ | $3.4\times10^{-1}$ | – | – | $8.9\times10^{-1}$ | N/A | $1.3\times10^{-2}$ | $8.3\times10^{1}$ | $2.3\times10^{-1}$ |
| Y514V | $5.2\times10^{-1}$ | $1.9\times10^{-2}$ | $5.3\times10^{5}$ | $7.2\times10^{-1}$ | $9.1\times10^{-1}$ | $7.8\times10^{-1}$ | $9.3\times10^{-1}$ | N/A | $2.8\times10^{-1}$ | $1.5\times10^{2}$ | $8.3\times10^{-1}$ |
| H515G | 1.3 | $3.9\times10^{-1}$ | $6.8\times10^{5}$ | $7.6\times10^{-1}$ | $4.6\times10^{-1}$ | $1.3\times10^{-2}$ | $8.5\times10^{-1}$ | N/A | $1.9\times10^{-2}$ | $6.7\times10^{1}$ | $2.0\times10^{-1}$ |
| H515V | 1.0 | $8.0\times10^{-1}$ | $1.9\times10^{5}$ | $6.4\times10^{-2}$ | 2.8 | $1.2\times10^{-2}$ | $9.1\times10^{-1}$ | N/A | $1.2\times10^{-1}$ | $1.5\times10^{2}$ | $7.9\times10^{-1}$ |
| M516G | 1.1 | $6.8\times10^{-1}$ | $3.8\times10^{5}$ | $2.4\times10^{-1}$ | 1.5 | $2.4\times10^{-1}$ | $3.1\times10^{-1}$ | N/A | **$2.0\times10^{-4}$** | $8.6\times10^{1}$ | $2.8\times10^{-1}$ |

| Variant | $f_a$ | $f_a$ *p*–value | $(k_{cat}/K_M)^{MeP}_{chem.}$ ($M^{-1}s^{-1}$) | $(k_{cat}/K_M)^{MeP}_{chem.}$ *p*–value | FC1 | FC1 *p*–value | FC2 | FC3 | FC2/3 *p*–value | FC4 ($s^{-1}$) | FC4 *p*–value |
|---|---|---|---|---|---|---|---|---|---|---|---|
| M516V | $4.5\times10^{-1}$ | **$6.0\times10^{-3}$** | $6.9\times10^{5}$ | $6.9\times10^{-1}$ | $7.8\times10^{-1}$ | $5.1\times10^{-1}$ | $8.4\times10^{-1}$ | N/A | **$4.4\times10^{-3}$** | $1.4\times10^{2}$ | $9.5\times10^{-1}$ |
| T517G | $3.3\times10^{-1}$ | **$1.0\times10^{-3}$** | $4.4\times10^{5}$ | $4.0\times10^{-1}$ | – | – | $8.4\times10^{-1}$ | N/A | **$5.3\times10^{-3}$** | $9.7\times10^{1}$ | $4.2\times10^{-1}$ |
| T517V | $1.3\times10^{-1}$ | **0** | $9.2\times10^{5}$ | $8.3\times10^{-2}$ | $7.6\times10^{-1}$ | $4.5\times10^{-1}$ | $6.1\times10^{-1}$ | N/A | **0** | $1.2\times10^{2}$ | $6.0\times10^{-1}$ |
| D518G | $1.9\times10^{-1}$ | **0** | $4.2\times10^{5}$ | $3.8\times10^{-1}$ | 1.6 | $2.6\times10^{-1}$ | $6.2\times10^{-1}$ | N/A | **0** | $1.2\times10^{2}$ | $7.3\times10^{-1}$ |
| D518V | $\leq 1.3\times10^{-2}$ | **0** | $\leq 1.3\times10^{6}$ | **$3.3\times10^{-2}$** | – | – | $7.0\times10^{-1}$ | N/A | **$6.0\times10^{-4}$** | $\geq 8.2\times10^{1}$ | $2.8\times10^{-1}$ |
| I519G | $1.1\times10^{-1}$ | **0** | $1.2\times10^{5}$ | $6.2\times10^{-2}$ | – | – | $5.7\times10^{-1}$ | N/A | **$7.0\times10^{-4}$** | $1.1\times10^{2}$ | $5.4\times10^{-1}$ |
| I519V | $6.4\times10^{-1}$ | $6.5\times10^{-2}$ | $5.3\times10^{5}$ | $7.0\times10^{-1}$ | $9.6\times10^{-1}$ | $8.7\times10^{-1}$ | $9.5\times10^{-1}$ | N/A | $2.3\times10^{-1}$ | $1.4\times10^{2}$ | $9.5\times10^{-1}$ |
| A520G | 1.3 | $3.8\times10^{-1}$ | $2.8\times10^{5}$ | $1.6\times10^{-1}$ | 1.3 | $3.7\times10^{-1}$ | $7.1\times10^{-1}$ | N/A | **$1.\times10^{-4}$** | $1.1\times10^{2}$ | $4.9\times10^{-1}$ |
| A520V | $4.8\times10^{-1}$ | **$6.0\times10^{-3}$** | $7.8\times10^{5}$ | $4.1\times10^{-1}$ | – | – | N/A | 1.0 | $5.6\times10^{-1}$ | $1.3\times10^{2}$ | $9.1\times10^{-1}$ |
| P521G | $5.8\times10^{-1}$ | $7.7\times10^{-2}$ | $3.4\times10^{5}$ | $2.3\times10^{-1}$ | 1.9 | $2.9\times10^{-2}$ | $7.5\times10^{-1}$ | N/A | **$7.0\times10^{-4}$** | $8.2\times10^{1}$ | $3.3\times10^{-1}$ |
| P521V | 1.5 | $1.1\times10^{-1}$ | $3.4\times10^{5}$ | $1.3\times10^{-1}$ | 1.0 | $9.6\times10^{-1}$ | $9.8\times10^{-1}$ | N/A | $7.0\times10^{-1}$ | $1.3\times10^{2}$ | $8.0\times10^{-1}$ |
| T522G | $2.0\times10^{-2}$ | **0** | $2.5\times10^{5}$ | $1.0\times10^{-1}$ | – | – | $9.4\times10^{-1}$ | N/A | $2.4\times10^{-1}$ | $1.2\times10^{2}$ | $6.3\times10^{-1}$ |
| T522V | $1.6\times10^{-1}$ | **0** | $1.7\times10^{5}$ | $9.8\times10^{-2}$ | 2.6 | $4.9\times10^{-2}$ | $5.6\times10^{-1}$ | N/A | **$1.3\times10^{-3}$** | $7.5\times10^{1}$ | $2.7\times10^{-1}$ |
| V523A | $8.0\times10^{-1}$ | $3.7\times10^{-1}$ | $1.0\times10^{5}$ | **$4.5\times10^{-2}$** | 4.0 | **$5.3\times10^{-3}$** | $8.9\times10^{-1}$ | N/A | $3.3\times10^{-2}$ | $1.5\times10^{2}$ | $8.2\times10^{-1}$ |
| V523G | $\leq 1.0\times10^{-3}$ | **0** | $\leq 1.3\times10^{6}$ | **$1.6\times10^{-2}$** | – | – | – | – | – | – | – |
| S524G | 1.0 | $8.5\times10^{-1}$ | $4.0\times10^{5}$ | $2.5\times10^{-1}$ | 1.5 | $2.7\times10^{-1}$ | $8.4\times10^{-1}$ | N/A | **$5.5\times10^{-3}$** | $8.7\times10^{1}$ | $2.4\times10^{-1}$ |
| S524V | $7.1\times10^{-1}$ | $1.3\times10^{-1}$ | $6.4\times10^{5}$ | $8.4\times10^{-1}$ | 1.1 | $8.5\times10^{-1}$ | $9.8\times10^{-1}$ | N/A | $5.4\times10^{-1}$ | $1.4\times10^{2}$ | $9.6\times10^{-1}$ |
| S525G | $\leq 6.0\times10^{-1}$ | $4.0\times10^{-2}$ | $\leq 3.8\times10^{5}$ | $2.5\times10^{-1}$ | $\leq 2.0$ | $3.0\times10^{-2}$ | $9.4\times10^{-1}$ | N/A | $2.7\times10^{-1}$ | $\geq 1.5\times10^{2}$ | $9.0\times10^{-1}$ |
| S525V | $6.7\times10^{-1}$ | $1.3\times10^{-1}$ | $9.3\times10^{5}$ | $1.1\times10^{-1}$ | $5.9\times10^{-1}$ | $1.4\times10^{-1}$ | N/A | 1.0 | $5.0\times10^{-1}$ | $1.6\times10^{2}$ | $7.1\times10^{-1}$ |
| L526G | $1.6\times10^{-1}$ | **0** | $3.4\times10^{5}$ | $1.4\times10^{-1}$ | 1.5 | $1.9\times10^{-1}$ | $8.4\times10^{-1}$ | N/A | **$5.0\times10^{-4}$** | $8.4\times10^{1}$ | $2.2\times10^{-1}$ |
| L526V | 1.1 | $6.2\times10^{-1}$ | $6.2\times10^{5}$ | $9.8\times10^{-1}$ | $8.4\times10^{-1}$ | $4.8\times10^{-1}$ | N/A | 1.0 | $7.8\times10^{-1}$ | $1.8\times10^{2}$ | $4.9\times10^{-1}$ |
| L527G | $1.0\times10^{-1}$ | **0** | $1.2\times10^{5}$ | **$2.8\times10^{-2}$** | – | – | $8.5\times10^{-1}$ | N/A | **$3.6\times10^{-3}$** | $9.9\times10^{1}$ | $3.9\times10^{-1}$ |
| L527V | 1.1 | $6.6\times10^{-1}$ | $2.8\times10^{5}$ | **$4.4\times10^{-2}$** | $9.3\times10^{-1}$ | $8.3\times10^{-1}$ | $9.7\times10^{-1}$ | N/A | $4.8\times10^{-1}$ | $1.2\times10^{2}$ | $5.7\times10^{-1}$ |
| K528G | $1.1\times10^{-1}$ | **0** | $1.8\times10^{5}$ | **$4.0\times10^{-2}$** | 3.1 | **$4.2\times10^{-3}$** | $7.0\times10^{-1}$ | N/A | **0** | $7.6\times10^{1}$ | $1.3\times10^{-1}$ |
| K528V | $1.8\times10^{-1}$ | **0** | $9.1\times10^{5}$ | $1.5\times10^{-1}$ | – | – | $6.8\times10^{-1}$ | N/A | **$6.0\times10^{-4}$** | $1.2\times10^{2}$ | $7.3\times10^{-1}$ |
| I529G | $1.5\times10^{-1}$ | **0** | $4.1\times10^{5}$ | $2.6\times10^{-1}$ | $9.1\times10^{-1}$ | $8.2\times10^{-1}$ | $6.2\times10^{-1}$ | N/A | **0** | $8.1\times10^{1}$ | $2.0\times10^{-1}$ |
| I529V | $9.4\times10^{-1}$ | $8.4\times10^{-1}$ | $6.1\times10^{5}$ | $9.9\times10^{-1}$ | $8.1\times10^{-1}$ | $5.6\times10^{-1}$ | $9.9\times10^{-1}$ | N/A | $6.9\times10^{-1}$ | $9.7\times10^{1}$ | $3.8\times10^{-1}$ |
| Q530G | 1.2 | $3.3\times10^{-1}$ | $5.0\times10^{4}$ | $5.6\times10^{-2}$ | 6.0 | $1.2\times10^{-2}$ | $7.6\times10^{-1}$ | N/A | **$1.\times10^{-4}$** | $8.8\times10^{1}$ | $2.6\times10^{-1}$ |
| Q530V | $5.6\times10^{-1}$ | $1.1\times10^{-2}$ | $6.8\times10^{5}$ | $6.5\times10^{-1}$ | $3.9\times10^{-1}$ | **$3.4\times10^{-3}$** | $7.0\times10^{-1}$ | N/A | **0** | $1.1\times10^{2}$ | $4.6\times10^{-1}$ |
| F531G | 1.5 | $1.3\times10^{-1}$ | $2.6\times10^{5}$ | $1.5\times10^{-1}$ | 1.4 | $2.2\times10^{-1}$ | $8.2\times10^{-1}$ | N/A | **$2.0\times10^{-3}$** | $8.9\times10^{1}$ | $2.9\times10^{-1}$ |
| F531V | $6.8\times10^{-1}$ | $1.2\times10^{-1}$ | $5.2\times10^{5}$ | $7.0\times10^{-1}$ | $7.8\times10^{-1}$ | $4.7\times10^{-1}$ | $9.4\times10^{-1}$ | N/A | $2.3\times10^{-1}$ | $1.3\times10^{2}$ | $8.2\times10^{-1}$ |
| P532G | 1.3 | $4.2\times10^{-1}$ | $1.6\times10^{5}$ | $6.3\times10^{-2}$ | 3.0 | **$1.2\times10^{-3}$** | $6.6\times10^{-1}$ | N/A | **$1.\times10^{-4}$** | $7.9\times10^{1}$ | $2.1\times10^{-1}$ |
| P532V | $2.6\times10^{-1}$ | **0** | $3.4\times10^{5}$ | $2.4\times10^{-1}$ | – | – | $5.7\times10^{-1}$ | N/A | **0** | $1.3\times10^{2}$ | $7.7\times10^{-1}$ |
| S533G | 1.1 | $8.1\times10^{-1}$ | $2.4\times10^{5}$ | $9.1\times10^{-2}$ | 2.3 | **$6.6\times10^{-3}$** | $5.9\times10^{-1}$ | N/A | **$3.0\times10^{-4}$** | $9.3\times10^{1}$ | $3.5\times10^{-1}$ |
| S533V | $\leq 4.2\times10^{-2}$ | **0** | $\leq 8.0\times10^{4}$ | **$1.7\times10^{-2}$** | – | – | – | – | – | – | – |
| G534A | 1.5 | $6.1\times10^{-2}$ | $2.5\times10^{5}$ | $7.9\times10^{-2}$ | 2.9 | **$4.6\times10^{-3}$** | $7.3\times10^{-1}$ | N/A | **0** | $9.8\times10^{1}$ | $3.4\times10^{-1}$ |
| G534V | $3.3\times10^{-1}$ | **0** | $6.1\times10^{5}$ | $9.9\times10^{-1}$ | 1.4 | $3.0\times10^{-1}$ | $3.9\times10^{-1}$ | N/A | **$1.\times10^{-4}$** | $9.7\times10^{1}$ | $3.9\times10^{-1}$ |
| A535G | 1.5 | $1.5\times10^{-1}$ | $2.7\times10^{5}$ | $1.3\times10^{-1}$ | 2.4 | **$3.0\times10^{-3}$** | $9.7\times10^{-1}$ | N/A | $6.6\times10^{-1}$ | $8.5\times10^{1}$ | $2.6\times10^{-1}$ |
| A535V | $7.9\times10^{-1}$ | $4.0\times10^{-1}$ | $8.6\times10^{5}$ | $2.2\times10^{-1}$ | $7.2\times10^{-1}$ | $2.2\times10^{-1}$ | N/A | 1.0 | $3.6\times10^{-1}$ | $2.1\times10^{2}$ | $1.7\times10^{-1}$ |
| V536A | $8.9\times10^{-1}$ | $6.5\times10^{-1}$ | $1.2\times10^{5}$ | $5.3\times10^{-2}$ | 2.6 | $2.4\times10^{-2}$ | $8.6\times10^{-1}$ | N/A | $3.5\times10^{-2}$ | $1.4\times10^{2}$ | $9.2\times10^{-1}$ |
| V536G | $\leq 2.2\times10^{-1}$ | **0** | $\leq 2.8\times10^{5}$ | $1.6\times10^{-1}$ | $\leq 1.3$ | $4.6\times10^{-1}$ | $6.5\times10^{-1}$ | N/A | **$1.\times10^{-4}$** | $\geq 7.9\times10^{1}$ | $2.5\times10^{-1}$ |
| G537A | $8.1\times10^{-2}$ | **0** | $6.4\times10^{5}$ | $8.5\times10^{-1}$ | – | – | $6.7\times10^{-1}$ | N/A | **0** | $1.4\times10^{2}$ | $9.4\times10^{-1}$ |
| G537V | $\leq 6.2\times10^{-2}$ | **0** | $\leq 1.2\times10^{5}$ | **$2.7\times10^{-2}$** | – | – | $4.3\times10^{-1}$ | N/A | **0** | $\geq 5.7\times10^{1}$ | $1.1\times10^{-1}$ |
| K538G | 1.6 | $1.0\times10^{-1}$ | $1.5\times10^{5}$ | $5.7\times10^{-2}$ | 3.5 | **$4.8\times10^{-4}$** | $7.7\times10^{-1}$ | N/A | **$5.4\times10^{-3}$** | $8.0\times10^{1}$ | $3.1\times10^{-1}$ |
| K538V | 1.3 | $3.7\times10^{-1}$ | $3.3\times10^{5}$ | $1.2\times10^{-1}$ | $9.4\times10^{-1}$ | $8.3\times10^{-1}$ | $9.6\times10^{-1}$ | N/A | $3.4\times10^{-1}$ | $1.2\times10^{2}$ | $6.9\times10^{-1}$ |
| P539G | 1.1 | $6.4\times10^{-1}$ | $4.9\times10^{5}$ | $5.4\times10^{-1}$ | $8.0\times10^{-1}$ | $4.1\times10^{-1}$ | N/A | 1.0 | $4.3\times10^{-1}$ | $1.4\times10^{2}$ | $9.8\times10^{-1}$ |
| P539V | 1.3 | $2.5\times10^{-1}$ | $7.1\times10^{5}$ | $6.0\times10^{-1}$ | $5.6\times10^{-1}$ | $3.4\times10^{-2}$ | N/A | 1.1 | $2.7\times10^{-1}$ | $1.4\times10^{2}$ | $9.4\times10^{-1}$ |
| I540G | $4.3\times10^{-2}$ | **0** | $1.6\times10^{5}$ | $9.6\times10^{-2}$ | – | – | $5.8\times10^{-1}$ | N/A | **$1.0\times10^{-3}$** | $1.0\times10^{2}$ | $4.6\times10^{-1}$ |
| I540V | $5.6\times10^{-1}$ | $2.2\times10^{-2}$ | $7.2\times10^{5}$ | $5.5\times10^{-1}$ | $6.4\times10^{-1}$ | $1.6\times10^{-1}$ | $9.7\times10^{-1}$ | N/A | $5.2\times10^{-1}$ | $1.6\times10^{2}$ | $7.5\times10^{-1}$ |
| T541G | $4.6\times10^{-1}$ | **$2.0\times10^{-3}$** | $5.1\times10^{5}$ | $6.5\times10^{-1}$ | 1.4 | $4.2\times10^{-1}$ | $8.4\times10^{-1}$ | N/A | **$1.9\times10^{-3}$** | $9.4\times10^{1}$ | $3.6\times10^{-1}$ |
| T541V | $9.6\times10^{-1}$ | $7.5\times10^{-1}$ | $5.7\times10^{5}$ | $7.9\times10^{-1}$ | $7.8\times10^{-1}$ | $4.8\times10^{-1}$ | N/A | 1.1 | $1.2\times10^{-1}$ | $1.5\times10^{2}$ | $9.1\times10^{-1}$ |
| E542G | $3.6\times10^{-1}$ | **0** | $5.9\times10^{5}$ | $9.5\times10^{-1}$ | 1.4 | $2.9\times10^{-1}$ | N/A | 1.1 | $1.4\times10^{-1}$ | $1.0\times10^{2}$ | $4.5\times10^{-1}$ |
| E542V | $5.7\times10^{-1}$ | $4.3\times10^{-2}$ | $4.4\times10^{5}$ | $4.0\times10^{-1}$ | 1.4 | $4.0\times10^{-1}$ | $8.6\times10^{-1}$ | N/A | $2.5\times10^{-2}$ | $1.3\times10^{2}$ | $7.8\times10^{-1}$ |
| V543A | 1.6 | $4.8\times10^{-2}$ | $3.6\times10^{5}$ | $1.7\times10^{-1}$ | 1.1 | $6.5\times10^{-1}$ | N/A | 1.1 | $1.6\times10^{-1}$ | $1.9\times10^{2}$ | $3.1\times10^{-1}$ |
| V543G | $6.2\times10^{-1}$ | $4.1\times10^{-2}$ | $2.1\times10^{5}$ | $5.9\times10^{-2}$ | 2.1 | $1.9\times10^{-2}$ | $9.5\times10^{-1}$ | N/A | $2.7\times10^{-1}$ | $1.2\times10^{2}$ | $5.9\times10^{-1}$ |
| I544G | $6.0\times10^{-1}$ | $8.6\times10^{-2}$ | $5.0\times10^{5}$ | $6.4\times10^{-1}$ | 1.0 | $9.8\times10^{-1}$ | $6.9\times10^{-1}$ | N/A | **$8.0\times10^{-4}$** | $7.7\times10^{1}$ | $2.9\times10^{-1}$ |
| I544V | $4.4\times10^{-1}$ | **$1.0\times10^{-3}$** | $8.4\times10^{5}$ | $2.4\times10^{-1}$ | $7.6\times10^{-1}$ | $3.1\times10^{-1}$ | $9.7\times10^{-1}$ | N/A | $5.8\times10^{-1}$ | $1.4\times10^{2}$ | 1.0 |
| G545A | 1.6 | $1.1\times10^{-1}$ | $3.3\times10^{5}$ | $1.4\times10^{-1}$ | 1.5 | $1.2\times10^{-1}$ | $9.2\times10^{-1}$ | N/A | $2.0\times10^{-1}$ | $8.7\times10^{1}$ | $2.9\times10^{-1}$ |
| G545V | 1.4 | $6.5\times10^{-2}$ | $4.3\times10^{5}$ | $2.4\times10^{-1}$ | $8.6\times10^{-1}$ | $7.2\times10^{-1}$ | N/A | 1.1 | $2.5\times10^{-1}$ | $1.6\times10^{2}$ | $5.6\times10^{-1}$ |
| R546G | 1.4 | $2.3\times10^{-1}$ | $3.1\times10^{5}$ | $1.4\times10^{-1}$ | 1.2 | $6.4\times10^{-1}$ | $9.7\times10^{-1}$ | N/A | $3.5\times10^{-1}$ | $1.5\times10^{2}$ | $8.9\times10^{-1}$ |
| R546V | $9.3\times10^{-1}$ | $6.9\times10^{-1}$ | $5.7\times10^{5}$ | $7.0\times10^{-1}$ | $8.6\times10^{-1}$ | $6.7\times10^{-1}$ | N/A | 1.0 | $4.3\times10^{-1}$ | $1.5\times10^{2}$ | $8.4\times10^{-1}$ |
